# Phylo-geo-network and haplogroup analysis of 611 novel Coronavirus (nCov-2019) genomes from India

**DOI:** 10.1101/2020.09.03.281774

**Authors:** Rezwanuzzaman Laskar, Safdar Ali

**Author notes:** **Corresponding author:** Dr Safdar Ali, Assistant Professor, Department of Biological Sciences, Aliah University, IIA/27, Newtown, Kolkata 700160, India.

## Abstract

The novel Coronavirus from Wuhan China discovered in December 2019 (nCOV-2019) has since developed into a global epidemic with major concerns about the possibility of the virus evolving into something even more sinister. In the present study we constructed the phylo-geo-network of nCOV-2019 genomes from across India to understand the viral evolution in the country. A total of 611 full length genomes were extracted from different states of India from the EpiCov repository of GISAID initiative and NCBI. Their alignment uncovered 270 parsimony informative sites. Further, 339 genomes were divided into 51 haplogroups. The network revealed the core haplogroup as that of reference sequence NC_045512.2 (Haplogroup A1) with 157 identical sequences present across 16 states. The rest were having not more than ten identical sequences across not more than three locations. Interestingly, some locations with fewer samples have more haplogroups and most haplogroups (41) are localized exclusively to any one state only, suggesting the local evolution of viruses. The two most common lineages are B6 and B1 (Pangolin) whereas clade A2a (Covidex) appears to be the most predominant in India. However, since the pandemic is still emerging, the final outcome will be clear later only.

## Introduction

Coronaviruses belonging to the family Coronaviridae have been named so owing to the electron microscopic structure resemblance of their virion structure to that of a crown. The spikes present on the virion surface provide for the resemblance. Their genome has a positive single strand RNA of 26 to 32kb in length and are known to infect a wide range of hosts (Cavanagh, 2007; Ismail et al., 2003; Lai and Cavanagh, 1997; Su et al., 2016; Weiss and Navas-Martin, 2005). A novel Coronavirus which has the potential to infect humans has been identified from Wuhan China in December 2019. It was subsequently referred to as nCOV-2019 (novel *Coronavirus* 2019) and since its emergence it has developed into a global epidemic. As of 28th August 2020, there were 33,10,234 cases and 60,472 deaths in India due to nCOV-2019 (https://www.mygov.in/covid-19). At the same time, as per WHO there have been 24,021,218 cases and 821,462 deaths globally (https://www.worldometers.info/coronavirus/). The nCOV-2019 is different from earlier Coronavirus outbreaks, severe acute respiratory syndrome (SARS) coronavirus in 2002 and Middle East respiratory syndrome (MERS) coronavirus in 2012 predominantly due to its extremely high transmission rates. The patients infected with nCOV-2019 have been observed to have varied symptoms ranging from normal flu like symptoms to high fever to invasive lesions (Chan et al., 2020; Huang et al., 2020; Peiris et al., 2004; Zaki et al., 2012; Zhu et al., 2020).

The nCOV-2019 belongs to genus Betacoronavirus and sub genus Sarbecovirus with suggested origin in bats. Various theories are in discussion about how it reached humans but nothing can be said with surety just yet (Lu et al., 2020; Zhou et al., 2020). However, the ever-increasing number of people being infected globally provides for the most conducive environments for the virus to evolve. The availability of full genome sequences for nCOV-2019 in GISAID has aided the study of these evolving sequences with both global and local perspectives (Shu and McCauley, 2017a).

At present, we build and analyze the phylo-geo-network of nCOV-2019 in India based on the publicly available full-length sequences of nCOV-2019 from India. We performed the haplogroup analysis and phylogenetic lineage study in addition to their defining mutations and geographical distributions. This assumes significance with India rapidly moving up the ladder in sharing the global burden of nCOV-2019 cases and is expected to continue so in near future owing to its demographic and health care structure.

## Materials and Methods

### Sequence Acquisition

Genome sequences of nCOV-2019 in FASTA format was assessed from the EpiCov™ repository (www.epicov.org) of GISAID initiative (Shu and McCauley, 2017b) and NCBI (www.ncbi.nlm.nih.gov).

On 6th June, 2020 we retrieved 611 FASTA sequence congregations along their rational meta data from GISAID EpiCoV server using the data filter ~ virus name: hCoV-19 - Host: Human - Location: Asia/India – Complete – High Coverage and use the genome ID by excluding the first part i.e. “EPI_ISL_” of GISAID accession ID. One sequence from the epicenter, Wuhan, China (NC_045512.2) was taken as reference. Details of the geographical distribution of the sequences and their accession numbers are provided in Figure 1 and Supplementary file 1 respectively. Location data of GISAID are used to identify the state of origin in India, and wherein state name is unavailable, state address of the originating lab has been used.

**Figure 1:**
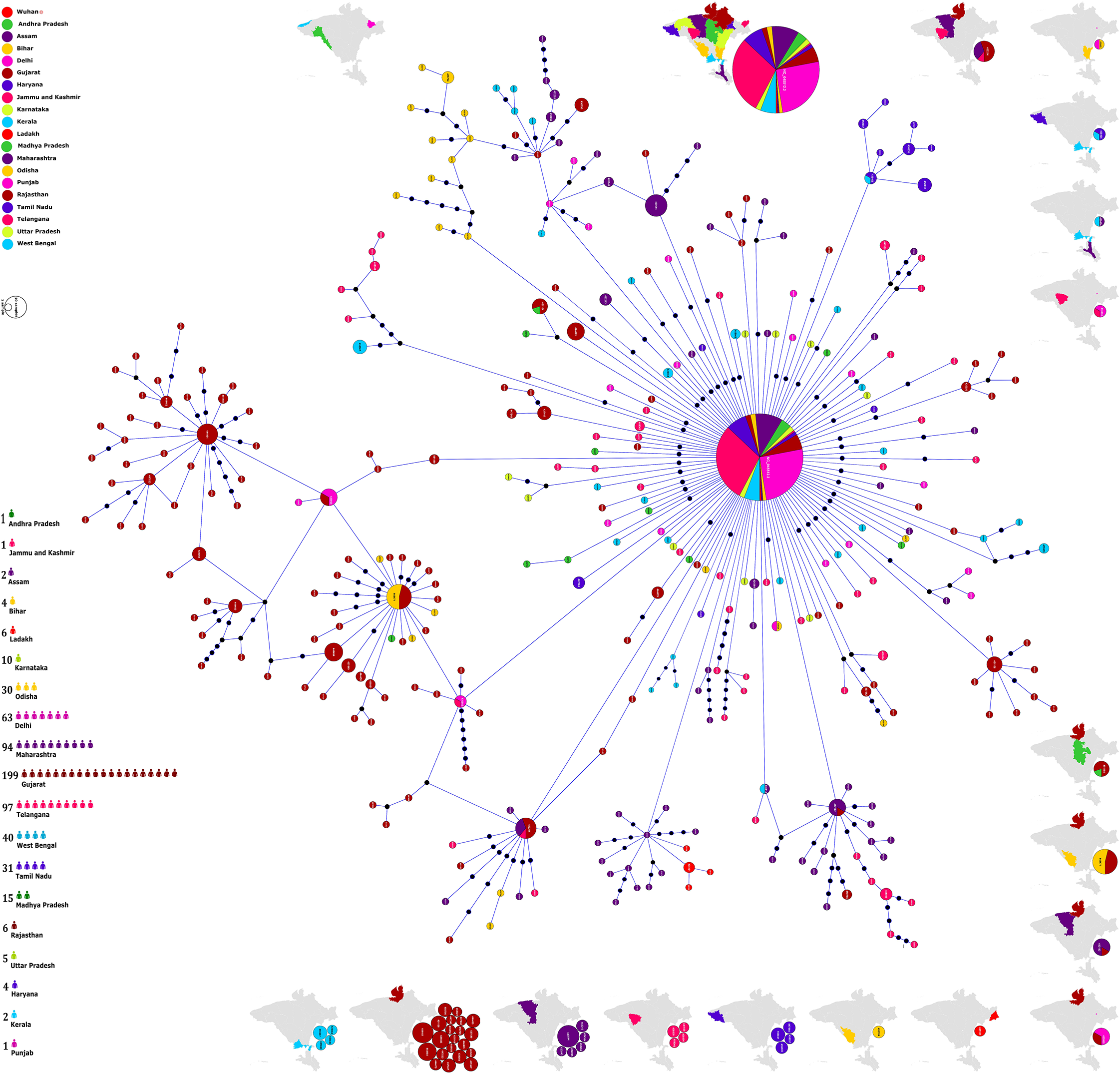
Phylogenomic geographic network of nCOV-2019 genomes from India. The nodes have been named after the Accession No of the defining sequences representing a particular cluster. The diameter of the circle represents the number of samples present therein. more the samples, longer the diameter. The different locations within India have been represented by color coding and the number of sequences from each state are shown in the bottom scale of the graph. Also shown are the distribution of haplogroups across different states in the maps on the periphery. On the right side are haplogroups present only in one state only whereas others include those present on multiple locations. Maps are generated and powered by Bing (©Geo Names, Microsoft, TomTom) through MS Excel 2019.

### Sequence Alignment

The congregations are aligned with the FFT-NS-fragment method using rapid calculation of full-length MSA of closely related viral genomes, a light-weight algorithm of MAFFT v7 web-server (https://mafft.cbrc.jp/alignment/software/closelyrelatedviralgenomes.html) (Katoh et al., 2018) and keeping alignment size exactly throughout the reference sequence. The nucleotide transformation sites of the alignment were further studied using MEGA X (Kumar et al., 2018)

### Phylogenetic Network Analysis

Aligned sequences were used to generated parsimony based TCS networks (Clement et al., 2002) implemented in Population Analysis with Reticulate Trees (PopART v1.7) software (Leigh and Bryant, 2015) where over 5 percent sites contain undefined states and will be masked. A map of haplotypes was also drawn using the same software with geotags and traits label coding.

### Genome Annotation

The tool IGLSF (Alam et al., 2019) arranges the location of variable sites according to genes. Using the software DNAPlotter (Carver et al., 2009) we used the Artemis (Carver et al., 2012) to annotate the genome and visualized it as a circular plot.

### Lineage and Subtyping Analysis

In the predefined cluster, using distinct nomenclature methods only a certain sequence belongs to the haplogroups have been classified into different lineage and subtype. Lineages that contribute most of the global spread have been assigned through Pangolin (Phylogenetic Assignment of Named Global Outbreak Lineages) Web (https://pangolin.cog-uk.io/), using nomenclature implemented by Rambaut, et al. (Rambaut et al., 2020). Viral subtypes of the studied Indian population were achieved using ‘SARS Cov 2 Nextstrain’ classification model of Covidex (https://cacciabue.shinyapps.io/shiny2/), a web-based subtyping tool (Cacciabue et al., 2020).

### Sequence Statistics

Multiple metrics were used to assess the population genetics to decipher the phylogenetic relationship. We calculated Tajima’s D (Tajima, 1989) statistic to test mutation-drift equilibrium and Pi value, segregating sites, parsimony-informative sites to measure DNA polymorphism among sequences using PopART statistics (Leigh and Bryant, 2015).

## Results and Discussion

### Phylogenetic network analysis

The alignment of genomes and their subsequent analysis revealed a total of 493 segregating sites of which 270 were parsimony informative (PI) sites. The incidence of sites and their distribution across gaps and ambiguous sequences and statistical evaluation has been summarized in Table 1. A negative value of Tajimas D statistic suggests the significance of these sites in evolution of these genomes. The reported phylo-geo-network herein has been built using the 152 PI sites excluding the gaps and ambiguous sequences.

**Table 1:** Some key statistical parameters observed in the study.

| **S No** | **Network Type** | **Number of segregating sites** | **Number of parsimony-informative sites** |  | **Nucleotide diversity** | **Tajima's D statistic** |
| --- | --- | --- | --- | --- | --- | --- |
|  |  |  | Excluding* | Including* |  |  |
| 1 | TCS | 493 | 152 | 270 | $\pi = 0.00120683$ | $D = -1.82662$<br>$p(D \geq -1.82662) = 0.982906$ |

\* Gaps and Ambiguous/Missing (Details in Supplementary file 3)

The phylo-geo-network analysis of the studied genomes has been represented in Figure 1. Several observations can be drawn from this analysis and the data forming the basis of this figure. First, the core of the network with maximum genomes (157) is the node of reference sequence of nCOV-2019 from Wuhan, China with accession no NC_045512.2. The fact that this accounts for over one fourth (25.7%) of the total studied sequences is a clear indication that in spite of many reported variations, the original nCOV-2019 genome continues to be the dominantly prevalent form. Interestingly, there was one sequence with genome id 458080 from Telangana which was hundred percent identical to the Wuhan reference sequence (Supplementary files 1 and 4). Though the absence of travel history for most of the studied patients and the sequences only being a partial representation of the patients present makes the conclusion subjective, but it does indicate about arrival of the virus directly from China to India. Though the variations are fast accumulating in the virus, it’s the original one that still prevails, at least in the Indian context. Viral evolution is a dynamic and fast process but unless due selection advantage is offered, a new form wouldn’t take over.

Secondly, the distribution of sequences from across India (Figure 1) don’t corroborate with the incidence scenarios but are a reflection of the ground level preparations and activity in getting the genomes sequenced. For instance, the under-representation of Maharashtra and Tamil Nadu in the present data set in-spite of being the two most affected states. However, assuming that the virus has an equal chance of evolving anywhere, we believe the number of sequences analyzed are apt for giving a glimpse of the ongoing viral evolution.

Thirdly, when we analyzed the distribution of PI sites across the genome and found it to be non-uniform in nature. We studied the distribution in the form of strike-rate of PI sites which we define as the number of bases after which there will be another PI site. This is to say that a region with a strike rate of 20 would mean a PI site every 20 bases and so on. Thus, a lower strike rate will infer a higher density of the PI sites in the region (Table 2). Based on our analysis, the Envelope and Spike protein have a PI strike rate of 45 and 115 respectively (Table 2). Before drawing any conclusions, we need to understand that a higher incidence of PI sites doesn’t necessarily corroborate to driving the evolutionary process as their impact on protein functionality needs to be ascertained first. However, it does indicate the potential genomic regions for the same which herein appear to be Envelope and Spike protein.

**Table 2:** Distribution of parsimony informative sites across the genome of nCOV-2019.

| **S No** | **Genome Region** | **Start position** | **End position** | **Size (bp)** | **No of parsimony sites** | **Strike-rate of parsimony sites\*** | **Position of PI Sites including the gaps and ambiguous sequences** |
| --- | --- | --- | --- | --- | --- | --- | --- |
| **1** | 5'UTR | 1 | 265 | 265 | 9 | 29.4 | 22, 55, 56, 94, 106, 218, 219, 222, 241 |
| **2** | ORF1a | 266 | 13483 | 13218 | 100 | 132.2 | 506, 635, 771, 875, 884, 1059, 1094, 1191, 1218, 1281, 1397, 1589, 1599, 1707, 1820, 1846, 2143, 2368, 2480, 2558, 2632, 2836, 3037, 3039, 3054, 3085, 3426, 3472, 3634, 3686, 3737, 3817, 4067, 4084, 4255, 4354, 4372, 4444, 4679, 4809, 4866, 4893, 4965, 5029, 5062, 5139, 5572, 5700, 5826, 6081, 6310, 6312, 6402, 6466, 6541, 6573, 6616, 6868, 6989, 7319, 7392, 7600, 7945, 8022, 8026, 8080, 8296, 8460, 8653, 8782, 8917, 8950, 9389, 9438, 9628, 9693, 10138, 10277, 10369, 10478, 10479, 10679, 10702, 10771, 10815, 11074, 11083, 11200, 11306, 11335, 11457, 11572, 11620, 12076, 12439, 12616, 12685, 12757, 13458 |
| **3** | ORF1ab | 13468 | 21555 | 8088 | 58 | 139.4 | 13585, 13617, 13730, 13859, 14130, 14181, 14274, 14408, 14425, 14673, 14805, 15324, 15435, 15451, 15708, 16017, 16078, 16355, 16393, 16626, 16738, 16852, 16887, 16993, 17135, 17440, 17722, 17747, 17858, 17959, 18052, 18129, 18380, 18395, 18457, 18486, 18511, 18877, 19086, 19185, 19344, 19417, 19524, 19679, 19684, 19816, 19872, 19983, 20006, 20063, 20087, 20151, 20355, 20773, 21004, 21137, 21550, 21551 |
| **4** | S protein | 21563 | 25384 | 3822 | 33 | 115.8 | 21575, 21627, 21628, 21646, 21724, 21792, 21795, 21890, 22289, 22343, 22374, 22444, 22468, 22530, 22663, 23120, 23236, 23277, 23111, 23403, 23593, 23638, 23678, 23815, 23821, 23929, 24811, 24933, 25098, 25290, 25314, 25381 |
| **5** | ORF3a | 25393 | 26220 | 828 | 10 | 82.8 | 25445, 25461, 25513, 25528, 25563, 25596, 25613, 25855, 25904, 26144 |
| **6** | NC | 26221 | 26244 | 24 | 1 | 24 | 26226 |
| **7** | E | 26245 | 26472 | 228 | 5 | 45.6 | 26330, 26338, 26375, 26376, 26467 |
| **8** | M | 26523 | 27191 | 669 | 6 | 111.5 | 26530, 26681, 26730, 26735, 27110, 27191 |
| **9** | ORF6 | 27202 | 27387 | 186 | 5 | 37.2 | 27213, 27379, 27382, 27383, 27384 |
| **10** | ORF7a | 27394 | 27759 | 366 | 1 | 366 | 27613 |
| **11** | ORF7b | 27756 | 27887 | 132 | 1 | 132 | 27874 |
| **12** | NC | 27888 | 27893 | 6 | 1 | 6 | 27889 |
| **13** | ORF8 | 27894 | 28259 | 366 | 7 | 52.3 | 28001, 28077, 28083, 28114, 28221, 28253, 28254 |
| **14** | N | 28274 | 29533 | 1260 | 20 | 63 | 28289, 28311, 28312, 28326, 28371, 28396, 28688, 28795, 28854, 28878, 28881, 28882, 28883, 28948, 29039, 29188, 29197, 29236, 29451, 29474 |
| **15** | NC | 29534 | 29557 | 24 | 3 | 8 | 29543, 29555, 29557 |
| **16** | ORF10 | 29558 | 29674 | 117 | 0 |  |  |
| **17** | 3'UTR | 29675 | 29903 | 229 | 10 | 22.9 | 29722, 29734, 29742, 29743, 29774, 29827, 29829, 29830, 29870, 29874 |
|  | **Total** |  |  |  | **270** |  |  |

\*Calculated by (Size/No of parsimony sites in the region)

### Haplogroup analysis and distribution

The network tree construction was accompanied by haplogroup determination of the studied genomes. The nodes representing haplogroups in phylo-geo-network in Figure 1 have been named as per accession number of the sequence defining the haplogroup. The nodal haplogroup represented by the Wuhan reference sequence NC_045512.2 has two maxima associated with it. The number of sequences therein as represented by the diameter of the circle (157 sequences) and total number of locations (16 states) in which the sequences are distributed. The details of distribution of all identical sequences have been summarized in Figure 2a and Supplementary file 2.

**Figure 2:**
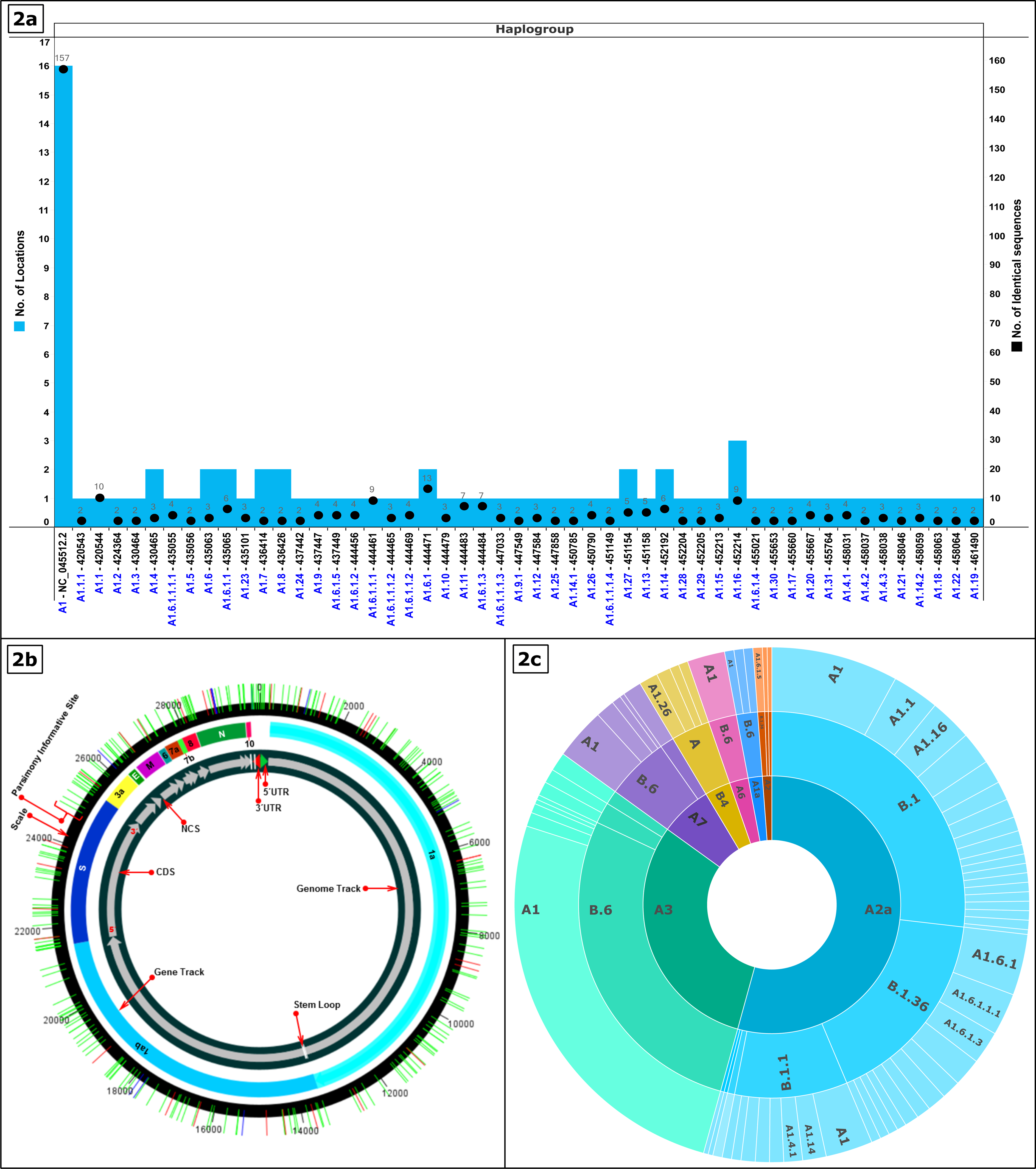
**a) Prevalence and geographical distribution of 51 haplogroups of nCOV-2019 genomes in India.** The number of identical sequences present in a haplogroup are shown on primary vertical axis whereas number of locations wherein its distribution is shown on secondary vertical axes. Note the maximum prevalence and widespread distribution of NC_045512.2 containing haplogroup (A1). For details of haplogroup IDs, identical sequences and locations please refer to Supplementary file 2. **b) Distribution of parsimony informative sites across the nCOV-2019 genome.** The nCOV-2019 genome has been represented circularly along with the locations of different genes/ORFs/Non coding regions have been represented. PI sites are shown as lines traversing the circle. **2c) Lineage and Subtype Analysis of nCOV-2019 genomes in India.** The outermost circle represents haplogroups reported in the study whereas the middle circle depicts lineage prediction by Pangolin web. The innermost circle is the clade analysis by Covidex web-tool.

Of the 611 studied genomes, the 51 haplogroups account for 339 genomes. At this juncture, we would like to note about the sequences left out of haplogroups. They belong to haplotypes which may converge to an existing haplogroup or emerge as a new one as the pandemic progresses. Due to the high mutation rate of viruses and with ever increasing incidence of the diseases the virus is replicating more and more and new polymorphisms are being generated every day. These variations are changing the haplotype and haplogroup profile on a regular basis.

We propose the nomenclature of the 51 observed haplogroups as per the path used to construct the network. We will explain the haplogroup nomenclature by taking a couple of examples. The haplogroup having NC_045512.2 was named A1 as the core of the network. From this cluster many haplogroups emerged and so on. The haplogroup A1.1 (420544) is defined by five positions; 241 (C→T), 3037 (C→T), 4809 (C→T), 14408 (C→T) and 23403 (A→G). However, as we move to haplogroup A1.1.1 (420543), in addition to the above mutations, another one at position 8782 (C→T) is present which becomes the defining polymorphism for this haplogroup. Similarly, haplogroup A1.6 (435063) is defined by positions 241(C→T), 1059 (C→T), 3037 (C→T), 14408 (C→T), 23403 (A→G) and 25563 (G→T). Subsequently haplogroup A1.6.1 (444471) is characterized by mutation at positions 18877 (C→T) and 26735 (C→T) and haplogroup A1.6.1.1 by additional mutations at 22444 (C→T) and 28854 (C→T). The haplogroup lineage thus defined clearly indicates that A1 is the most prevalent one while A1.6 is the most evolving one as it has the maximum number of steps going up to A1.6.1.1.1.4 reflective of five steps and stages of mutations/PI sites. The position of all the observed PI sites has been listed in Table 2/Figure 2b and their details are summarized in Supplementary file 3. The haplogroup nomenclature has been listed in correlation with their genome IDs and location in Table 3. If we observe the PI sites reported in the study, it includes most of the commonly reported sites from across the world besides some novel ones. However, we aren’t emphasizing on the novelty of sites due to the fast-changing scenario and rapidly emerging data.

**Table 3:** Details of Haplogroups: Geographical distribution and Phylogenetic lineage.

| **Haplogroup** | **Node Label/Genome ID** | **State** | **Most Common Countries** | **Lineage analysis (Rambaut et al., 2020)** | **Subtype analysis-SARS Cov 2 Nextstrain (Hadfield et al., 2018)** |
| --- | --- | --- | --- | --- | --- |
| Proposed | Assigned by (GISAID/NCBI) | Assigned by Database (GISAID/NCBI) | Assigned by Pangolin Web-server | Prediction by Pangolin Web-server | Prediction by Covidex Web-server |
| **A1** | NC_045512.2 | 1. Assam<br>2. Bihar<br>3. Delhi<br>4. Gujarat<br>5. Haryana<br>6. Jammu<br>7. Karnataka<br>8. Madhya Pradesh<br>9. Maharashtra<br>10. Odisha<br>11. Rajasthan<br>12. Tamil Nadu<br>13. Telangana<br>14. Uttar Pradesh<br>15. West Bengal<br>16. Wuhan, China | 1. Australia, Singapore, USA<br>2. India, Singapore, Australia<br>3. UK, Australia, USA<br>4. UK, China, USA<br>5. UK, Spain, Australia<br>6. UK, USA, Australia<br>7. UK, USA, China | 1. B<br>2. B.1<br>3. B.1.1<br>4. B.1.5<br>5. B.6 | 1. A1a<br>2. A2<br>3. A2a<br>4. A3<br>5. A6<br>6. A7 |
| **A1.1** | 420544 | Maharashtra | UK, USA, Australia | B.1 | A2a |
| **A1.1.1** | 420543 | Maharashtra | UK, USA, Australia | B.1 | A2a |
| **A1.10** | 444479 | Gujarat | UK, USA, Australia | B.1 | A2a |
| **A1.11** | 444483 | Gujarat | UK, USA, Australia | B.1 | A2a |
| **A1.12** | 447584 | Tamil Nadu | India, Singapore, Australia | B.6 | A3 |
| **A1.13** | 451158 | Gujarat | UK, USA, Australia | B.1 | A2a |
| **A1.14** | 452192 | 1. Gujarat<br>2. Maharashtra | UK, Australia, USA<br>UK, USA, Australia | 1. B.1<br>2. B.1.1 | A2a |
| **A1.14.1** | 450785 | Gujarat | UK, USA, Australia | B.1 | A2a |
| **A1.14.2** | 458059 | Telangana | UK, Australia, USA | B.1.1 | A2a |
| **A1.15** | 452213 | Maharashtra | Australia, UK, Turkey | B.4 | A3 |
| **A1.16** | 452214 | 1. Gujarat<br>2. Maharashtra<br>3. Telangana | UK, USA, Australia | B.1 | A2a |
| **A1.17** | 455660 | West Bengal | UK, USA, Australia | B.1 | A2a |
| **A1.18** | 458063 | Telangana | India, Singapore, Australia | B.6 | A7 |
| **A1.19** | 461490 | Gujarat | UK, USA, Australia | B.1 | A2a |
| **A1.2** | 424364 | Maharashtra | UK, USA, Australia | B.1 | A2a |
| **A1.20** | 455667 | West Bengal | UK, USA, Australia | B.1 | A2a |
| **A1.21** | 458046 | Telangana | UK, Australia, Gambia | B.1.1.8 | A2a |
| **A1.22** | 458064 | Telangana | UK, USA, Australia | B.1 | A2a |
| **A1.23** | 435101 | Ladakh | Australia, UK, Turkey | B.4 | A3 |
| **A1.24** | 437442 | Gujarat | Australia, Singapore, USA | B.6 | A1a |
| **A1.25** | 447858 | Telangana | India, Singapore, Australia | B.6 | A3 |
| **A1.26** | 450790 | Gujarat | China, South_Korea, USA | A | B4 |
| **A1.27** | 451154 | 1. Gujarat<br>2. Madhya Pradesh | Australia, Singapore, USA<br>India, Singapore, Australia | B.6 | 1. A3<br>2. A7 |
| **A1.28** | 452204 | Maharashtra | China, South_Korea, USA | A | B4 |
| **A1.29** | 452205 | Maharashtra | China, South_Korea, USA | A | B4 |
| **A1.3** | 430464 | West Bengal | UK, Australia, USA | B.1.1 | A2a |
| **A1.30** | 455653 | West Bengal | UK, USA, Australia | B.1 | A2a |
| **A1.31** | 455764 | Odisha | China, South_Korea, USA | A | B4 |
| **A1.4** | 430465 | 1. Tamil Nadu<br>2. West Bengal | UK, Australia, USA | B.1.1 | A2a |
| **A1.4.1** | 458031 | Tamil Nadu | UK, Australia, USA | B.1.1 | A2a |
| **A1.4.2** | 458037 | Tamil Nadu | UK, Australia, USA | B.1.1 | A2a |
| **A1.4.3** | 458038 | Tamil Nadu | UK, Australia, USA | B.1.1 | A2a |
| **A1.5** | 435056 | Gujarat | UK, USA, Australia | B.1 | A2a |
| **A1.6** | 435063 | 1. Delhi<br>2. Telangana | UK, USA, Australia | B.1 | A2a |
| **A1.6.1** | 444471 | 1. Gujarat<br>2. Odisha | Saudi_Arabia, UK, Turkey<br>Turkey, Finland, UK | B.1.36 | A2a |
| **A1.6.1.1** | 435065 | 1. Delhi<br>2. Gujarat | Saudi_Arabia, UK, Turkey<br>Turkey, Finland, UK | B.1.36 | A2a |
| **A1.6.1.1.1** | 444461 | Gujarat | Saudi_Arabia, UK, Turkey<br>Turkey, Finland, UK | B.1.36 | A2a |
| **A1.6.1.1.1.1** | 435055 | Gujarat | Turkey, Finland, UK | B.1.36 | A2a |
| **A1.6.1.1.1.2** | 444465 | Gujarat | Turkey, Finland, UK | B.1.36 | A2a |
| **A1.6.1.1.1.3** | 447033 | Gujarat | Saudi_Arabia, UK, Turkey<br>Turkey, Finland, UK | B.1.36 | A2a |
| **A1.6.1.1.1.4** | 451149 | Gujarat | Turkey, Finland, UK | B.1.36 | A2a |
| **A1.6.1.1.2** | 444469 | Gujarat | Turkey, Finland, UK | B.1.36 | A2a |
| **A1.6.1.2** | 444456 | Gujarat | Turkey, Finland, UK | B.1.36 | A2a |
| **A1.6.1.3** | 444484 | Gujarat | Turkey, Finland, UK | B.1.36 | A2a |
| **A1.6.1.4** | 455021 | Gujarat | Saudi_Arabia, UK, Turkey | B.1.36 | A2a |
| **A1.6.1.5** | 437449 | Gujarat | Turkey, Finland, UK | B.1.36 | 1. A2<br>2. A2a |
| **A1.7** | 436414 | 1. Assam<br>2. West Bengal | India, Singapore, Australia | B.6 | A1a |
| **A1.8** | 436426 | 1. Bihar<br>2. Delhi | India, Singapore, Australia | B.6 | 1. A3<br>2. A7 |
| **A1.9** | 437447 | Gujarat | UK, USA, Australia | B.1 | A2a |
| **A1.9.1** | 447549 | Gujarat | UK, USA, Australia | B.1 | A2a |

The geographical distribution of the haplogroups can be looked at from two different aspects. To begin with, which haplogroup is found in which location. Herein, A1 (NC_045512.2) haplogroup as already mentioned was most widely prevalent with 157 sequences distributed across 16 locations. All other haplogroups had ten or fewer genomes spread across one to three locations (Figure 2a). The scenario is more interesting if we inverse the analysis as in which location had how many haplogroups. Gujarat with a maximal representation of 199 genomes had 27 different haplogroups but this isn’t the norm as in more sequences would mean more haplogroups. Delhi (63 genomes, 3 haplogroups), Maharashtra (94 genomes, 9 haplogroups) and West Bengal (40 genomes,7 haplogroups) exhibit the non-linearity of the same. Also, 41 haplogroups have a single location only led by Gujarat (21), Maharashtra (6), West Bengal, Telangana, Tamil Nadu (4 each) and Ladakh, Orissa (1 each). Three states Punjab, Andhra Pradesh and Kerala don’t have any haplogroup so far. The distribution of haplogroups across states has been shown in Figure 1 and Supplementary file 2. The fact that some locations with fewer samples have more haplogroups and most haplogroups are localized exclusively to a single state is a clear indication about the local evolution of viruses. However, since the pandemic is still emerging, the final outcome will be clear only at a later stage.

### Lineage and Subtype Analysis

We also ascertained the lineage and subtype of the observed sequences through Pangolin and Covidex respectively. Also, the presence of lineages in India across the world was studied. The fact that phylogenetic lineage of nCOV-2019 genomes from India exhibits its relation with diverse countries like USA, Australia, UK, Singapore, China and Turkey is reflective of the global nature of the pandemic. Most of it can be attributed to international air travel and diverse regulations across countries. The three most common lineages in India as predicted by Pangolin are B6, B1 and B1.36 whereas clade A2a appears to be the most predominant one as predicted by Covidex (Figure 2c, Table 3, Supplementary file 4). These lineages can shift with increasing incidences and accumulating variations which requires regular monitoring. However, proper recording of both national and international travel history for all the patients will go a long way in unveiling the true path of viral evolution.

## Conclusions

The understanding of emergence and evolution of nCOV-2019 pandemic in India is an apt set up to understand viral divergence and evolution due to its huge population and diversity. As of now, the virus most prevalent in India is of the same haplogroup as the nCOV-2019 reference sequence from Wuhan indicating absence of any significant novel emerging strain. The two most common lineages are B6 and B1 whereas clade A2a appears to be the most predominant one in Indian context. However, with ever increasing incidence the situation needs to be monitored regularly.

## Authors’ contributions

RL performed the multiple sequence alignment and phylogenomic tree evaluation. SA supervised the whole study and prepared the manuscript.

## Acknowledgements

The authors thank the Department of Biological Sciences, Aliah University, Kolkata, India for all the financial and infrastructural support provided. Authors acknowledge all the authors associated with originating and submitting laboratories of the sequences from GISAID’s EpiFlu™ (www.gisaid.org) Database on which this research is based.

## Ethics approval

Not Applicable.

## Details of Supplementary Files

Supplementary File 1: Details of nCovid 2019 genomes used in the study

Supplementary file 2: Details of identical sequences in the study and their geographical distribution

Supplementary files 3: Details of parsimony informative sites including gaps and ambiguous sequences observed in the study

Supplementary file 4: Details of lineage analysis of studied genomes

