## Supplementary files 1-4 for "Phylo-geo-network and haplogroup analysis of 611 novel Coronavirus (nCov-2019) genomes from India"

**Supplementary File 1: Details of nCovid 2019 genomes used in the study**

| S No | Genome ID | Accession ID | Virus name | Location | State | Collection date | Originating lab |
|---|---|---|---|---|---|---|---|
| 1 | 413522 | EPI_ISL_413522 | hCoV-19/India/1-27/2020 | Asia / India / Kerala | Kerala | 2020-01-27 | Indian Council of Medical Research - National Institute of Virology |
| 2 | 413523 | EPI_ISL_413523 | hCoV-19/India/1-31/2020 | Asia / India / Kerala | Kerala | 2020-01-31 | Indian Council of Medical Research-National Institute of Virology |
| 3 | 420543 | EPI_ISL_420543 | hCoV-19/India/763/2020 | Asia / India | Maharashtra | 2020-03-03 | National Influenza Center, Indian Council of Medical Research - National Institute of Virology |
| 4 | 420544 | EPI_ISL_420544 | hCoV-19/India/2020763/2020 | Asia / India | Maharashtra | 2020 | Indian Council of Medical Research-National Institute of Virology, Microbial Containment Complex |
| 5 | 420545 | EPI_ISL_420545 | hCoV-19/India/770/2020 | Asia / India | Maharashtra | 2020-03-03 | National Influenza Center, Indian Council of Medical Research - National Institute of Virology |
| 6 | 420546 | EPI_ISL_420546 | hCoV-19/India/2020770/2020 | Asia / India | Maharashtra | 2020 | Indian Council of Medical Research-National Institute of Virology, Microbial Containment Complex |
| 7 | 420547 | EPI_ISL_420547 | hCoV-19/India/772/2020 | Asia / India | Maharashtra | 2020-03-03 | National Influenza Center, Indian Council of Medical Research - National Institute of Virology |
| 8 | 420548 | EPI_ISL_420548 | hCoV-19/India/2020772/2020 | Asia / India | Maharashtra | 2020 | Indian Council of Medical Research-National Institute of Virology, Microbial Containment Complex |
| 9 | 420549 | EPI_ISL_420549 | hCoV-19/India/773/2020 | Asia / India | Maharashtra | 2020-03-03 | National Influenza Center, Indian Council of Medical Research - National Institute of Virology |
| 10 | 420550 | EPI_ISL_420550 | hCoV-19/India/2020773/2020 | Asia / India | Maharashtra | 2020 | Indian Council of Medical Research-National Institute of Virology, Microbial Containment Complex |
| 11 | 420551 | EPI_ISL_420551 | hCoV-19/India/777/2020 | Asia / India | Maharashtra | 2020-03-03 | National Influenza Center, Indian Council of Medical Research - National Institute of Virology |
| 12 | 420552 | EPI_ISL_420552 | hCoV-19/India/2020777/2020 | Asia / India | Maharashtra | 2020 | Indian Council of Medical Research-National Institute of Virology, Microbial Containment Complex |
| 13 | 420553 | EPI_ISL_420553 | hCoV-19/India/781/2020 | Asia / India | Maharashtra | 2020-03-03 | National Influenza Center, Indian Council of Medical Research - National Institute of Virology |
| 14 | 420554 | EPI_ISL_420554 | hCoV-19/India/2020781/2020 | Asia / India | Maharashtra | 2020 | Indian Council of Medical Research-National Institute of Virology, Microbial Containment Complex |
| 15 | 420555 | EPI_ISL_420555 | hCoV-19/India/c32/2020 | Asia / India | Maharashtra | 2020-03-03 | National Influenza Center, Indian Council of Medical Research - National Institute of Virology |
| 16 | 420556 | EPI_ISL_420556 | hCoV-19/India/2020c32/2020 | Asia / India | Maharashtra | 2020 | Indian Council of Medical Research-National Institute of Virology, Microbial Containment Complex |
| 17 | 421662 | EPI_ISL_421662 | hCoV-19/India/1073/2020 | Asia / India | Maharashtra | 2020-03-10 | National Influenza Center, Indian Council of Medical Research - National Institute of Virology |
| 18 | 421663 | EPI_ISL_421663 | hCoV-19/India/1093/2020 | Asia / India | Maharashtra | 2020-03-10 | National Influenza Center, Indian Council of Medical Research - National Institute of Virology |
| 19 | 421664 | EPI_ISL_421664 | hCoV-19/India/1100/2020 | Asia / India | Maharashtra | 2020-03-10 | National Influenza Center, Indian Council of Medical Research - National Institute of Virology |
| 20 | 421665 | EPI_ISL_421665 | hCoV-19/India/1104/2020 | Asia / India | Maharashtra | 2020-03-10 | National Influenza Center, Indian Council of Medical Research - National Institute of Virology |
| 21 | 421666 | EPI_ISL_421666 | hCoV-19/India/1111/2020 | Asia / India | Maharashtra | 2020-03-10 | National Influenza Center, Indian Council of Medical Research - National Institute of Virology |
| 22 | 421667 | EPI_ISL_421667 | hCoV-19/India/1115/2020 | Asia / India | Maharashtra | 2020-03-10 | National Influenza Center, Indian Council of Medical Research - National Institute of Virology |
| 23 | 421668 | EPI_ISL_421668 | hCoV-19/India/1125/2020 | Asia / India | Maharashtra | 2020-03-10 | National Influenza Center, Indian Council of Medical Research - National Institute of Virology |
| 24 | 421669 | EPI_ISL_421669 | hCoV-19/India/1616/2020 | Asia / India | Maharashtra | 2020-03-12 | National Influenza Center, Indian Council of Medical Research - National Institute of Virology |
| 25 | 421670 | EPI_ISL_421670 | hCoV-19/India/1617/2020 | Asia / India | Maharashtra | 2020-03-12 | National Influenza Center, Indian Council of Medical Research - National Institute of Virology |
| 26 | 421671 | EPI_ISL_421671 | hCoV-19/India/1621/2020 | Asia / India | Maharashtra | 2020-03-12 | National Influenza Center, Indian Council of Medical Research - National Institute of Virology |
| 27 | 421672 | EPI_ISL_421672 | hCoV-19/India/1644/2020 | Asia / India | Maharashtra | 2020-03-12 | National Influenza Center, Indian Council of Medical Research - National Institute of Virology |
| 28 | 424361 | EPI_ISL_424361 | hCoV-19/India/1063/2020 | Asia / India | Maharashtra | 2020-03-10 | National Influenza Center, Indian Council of Medical Research - National Institute of Virology |
| 29 | 424362 | EPI_ISL_424362 | hCoV-19/India/1135/2020 | Asia / India | Maharashtra | 2020-03-10 | National Influenza Center, Indian Council of Medical Research - National Institute of Virology |
| 30 | 424363 | EPI_ISL_424363 | hCoV-19/India/1652/2020 | Asia / India | Maharashtra | 2020-03-12 | National Influenza Center, Indian Council of Medical Research - National Institute of Virology |
| 31 | 424364 | EPI_ISL_424364 | hCoV-19/India/3118/2020 | Asia / India | Maharashtra | 2020-03-17 | National Influenza Center, Indian Council of Medical Research - National Institute of Virology |
| 32 | 424365 | EPI_ISL_424365 | hCoV-19/India/3239/2020 | Asia / India | Maharashtra | 2020-03-17 | National Influenza Center, Indian Council of Medical Research - National Institute of Virology |
| 33 | 426179 | EPI_ISL_426179 | hCoV-19/India/c31/2020 | Asia / India | Maharashtra | 2020-03-02 | National Influenza Center, Indian Council of Medical Research - National Institute of Virology |
| 34 | 426414 | EPI_ISL_426414 | hCoV-19/India/GBRC1/2020 | Asia / India / Gujarat | Gujarat | 2020-04-05 | Sir M P Shah Government Medical College |
| 35 | 426415 | EPI_ISL_426415 | hCoV-19/India/GBRC1s/2020 | Asia / India / Gujarat | Gujarat | 2020-04-05 | Sir M P Shah Government Medical College, Jamnagar |
| 36 | 428479 | EPI_ISL_428479 | hCoV-19/India/nimh-0113/2020 | Asia / India / Karnataka | Karnataka | 2020-04-06 | District Surveillence Unit |
| 37 | 428480 | EPI_ISL_428480 | hCoV-19/India/nimh-0116/2020 | Asia / India / Karnataka | Karnataka | 2020-04-06 | District Surveillence Unit |
| 38 | 428481 | EPI_ISL_428481 | hCoV-19/India/nimh-0130/2020 | Asia / India / Karnataka | Karnataka | 2020-04-06 | District Surveillence Unit |
| 39 | 428482 | EPI_ISL_428482 | hCoV-19/India/nimh-0182/2020 | Asia / India / Karnataka | Karnataka | 2020-04-08 | District Surveillence Unit |
| 40 | 428483 | EPI_ISL_428483 | hCoV-19/India/nimh-0318/2020 | Asia / India / Karnataka | Karnataka | 2020-04-10 | District Surveillence Unit |
| 41 | 428484 | EPI_ISL_428484 | hCoV-19/India/nimh-0351/2020 | Asia / India / Karnataka | Karnataka | 2020-04-10 | District Surveillence Unit |
| 42 | 428485 | EPI_ISL_428485 | hCoV-19/India/nimh-0834/2020 | Asia / India / Karnataka | Karnataka | 2020-04-12 | District Surveillence Unit |
| 43 | 428486 | EPI_ISL_428486 | hCoV-19/India/nimh-0996/2020 | Asia / India / Karnataka | Karnataka | 2020-04-14 | District Surveillence Unit |
| 44 | 428487 | EPI_ISL_428487 | hCoV-19/India/nimh-1071/2020 | Asia / India / Karnataka | Karnataka | 2020-04-14 | District Surveillence Unit |
| 45 | 430464 | EPI_ISL_430464 | hCoV-19/India/S3/2020 | Asia / India / West Bengal / Kolkata | West Bengal | 2020-03-21 | ICMR-National Institute of Cholera and Enteric Diseases |
| 46 | 430465 | EPI_ISL_430465 | hCoV-19/India/S5/2020 | Asia / India / West Bengal / Darjeeling | West Bengal | 2020-03-28 | ICMR-National Institute of Cholera and Enteric Diseases |
| 47 | 430466 | EPI_ISL_430466 | hCoV-19/India/S6/2020 | Asia / India / West Bengal / Tehatta | West Bengal | 2020-03-26 | ICMR-National Institute of Cholera and Enteric Diseases |
| 48 | 430467 | EPI_ISL_430467 | hCoV-19/India/S11/2020 | Asia / India / West Bengal / East Medinipur | West Bengal | 2020-04-03 | ICMR-National Institute of Cholera and Enteric Diseases |
| 49 | 430468 | EPI_ISL_430468 | hCoV-19/India/S2/2020 | Asia / India / West_Bengal / Kolkata | West Bengal | 2020-03-21 | ICMR-National Institute of Cholera and Enteric Diseases |
| 50 | 431101 | EPI_ISL_431101 | hCoV-19/India/GMC-RK100/2020 | Asia / India / Telangana / Hyderabad | Telangana | 2020-03-01 | Department of Microbiology,Gandhi Medical College and Hospital |
| 51 | 431102 | EPI_ISL_431102 | hCoV-19/India/GMC-KN318/2020 | Asia / India / Telangana / Hyderabad | Telangana | 2020-03-11 | Department of MicroBiology,Gandhi Medical College and Hospital,Secendrabad,Hyderabad,India |
| 52 | 431103 | EPI_ISL_431103 | hCoV-19/India/GMC-KN443/2020 | Asia / India / Telangana / Hyderabad | Telangana | 2020-03-16 | Department of Microbiology, Gandhi Medical College and Hospital, Secendrabad, Hyderabad, India |
| 53 | 431117 | EPI_ISL_431117 | hCoV-19/India/GMC-TC469/2020 | Asia / India / Telangana / Hyderabad | Telangana | 2020-03-20 | Department of Microbiology, Gandhi Medical College and Hospital, Secendrabad, Hyderabad, India |
| 54 | 435049 | EPI_ISL_435049 | hCoV-19/India/GBRC2/2020 | Asia / India / Gujarat / Ahmedabad | Gujarat | 2020-04-13 | B.J. Medical College and Civil hospital |
| 55 | 435050 | EPI_ISL_435050 | hCoV-19/India/GBRC3/2020 | Asia / India / Gujarat / Ahmedabad | Gujarat | 2020-04-13 | B.J. Medical College and Civil hospital |
| 56 | 435051 | EPI_ISL_435051 | hCoV-19/India/GBRC4/2020 | Asia / India / Gujarat / Ahmedabad | Gujarat | 2020-04-13 | B.J. Medical College and Civil hospital |
| 57 | 435052 | EPI_ISL_435052 | hCoV-19/India/GBRC5/2020 | Asia / India / Gujarat / Ahmedabad | Gujarat | 2020-04-13 | B.J. Medical College and Civil hospital |
| 58 | 435053 | EPI_ISL_435053 | hCoV-19/India/GBRC6/2020 | Asia / India / Gujarat / Ahmedabad | Gujarat | 2020-04-07 | B.J. Medical College and Civil hospital |
| 59 | 435054 | EPI_ISL_435054 | hCoV-19/India/GBRC7/2020 | Asia / India / Gujarat / Ahmedabad | Gujarat | 2020-04-14 | B.J. Medical College and Civil hospital |
| 60 | 435055 | EPI_ISL_435055 | hCoV-19/India/GBRC8/2020 | Asia / India / Gujarat / Gandhinagar | Gujarat | 2020-04-22 | Gujarat Biotechnology Research Centre |
| 61 | 435056 | EPI_ISL_435056 | hCoV-19/India/GBRC9/2020 | Asia / India / Gujarat / Mansa | Gujarat | 2020-04-21 | Gujarat Biotechnology Research Centre |
| 62 | 435060 | EPI_ISL_435060 | hCoV-19/India/NCDC-01257/2020 | Asia / India / Uttar Pradesh / Noida | Uttar Pradesh | 2020-03-12 | National Centre for Disease control (NCDC), CSIR-Institute of Genomics and Integrative Biology (CSIR-IGIB) |
| 63 | 435061 | EPI_ISL_435061 | hCoV-19/India/NCDC-01537/2020 | Asia / India / Delhi | Delhi | 2020-03-16 | National Centre for Disease control (NCDC), CSIR-Institute of Genomics and Integrative Biology (CSIR-IGIB) |
| 64 | 435062 | EPI_ISL_435062 | hCoV-19/India/NCDC-01538/2020 | Asia / India / Punjab | Punjab | 2020-03-16 | National Centre for Disease control (NCDC), CSIR-Institute of Genomics and Integrative Biology (CSIR-IGIB) |
| 65 | 435063 | EPI_ISL_435063 | hCoV-19/India/NCDC-01326/2020 | Asia / India / Delhi | Delhi | 2020-03-13 | National Centre for Disease control (NCDC), CSIR-Institute of Genomics and Integrative Biology (CSIR-IGIB) |
| 66 | 435064 | EPI_ISL_435064 | hCoV-19/India/NCDC-01705/2020 | Asia / India / Delhi | Delhi | 2020-03-18 | National Centre for Disease control (NCDC), CSIR-Institute of Genomics and Integrative Biology (CSIR-IGIB) |
| 67 | 435065 | EPI_ISL_435065 | hCoV-19/India/NCDC-01475/2020 | Asia / India / Delhi | Delhi | 2020-03-15 | National Centre for Disease control (NCDC), CSIR-Institute of Genomics and Integrative Biology (CSIR-IGIB) |
| 68 | 435066 | EPI_ISL_435066 | hCoV-19/India/NCDC-01711/2020 | Asia / India / Delhi | Delhi | 2020-03-18 | National Centre for Disease control (NCDC), CSIR-Institute of Genomics and Integrative Biology (CSIR-IGIB) |
| 69 | 435067 | EPI_ISL_435067 | hCoV-19/India/NCDC-01712/2020 | Asia / India / Delhi | Delhi | 2020-03-18 | National Centre for Disease control (NCDC), CSIR-Institute of Genomics and Integrative Biology (CSIR-IGIB) |
| 70 | 435068 | EPI_ISL_435068 | hCoV-19/India/NCDC-01757/2020 | Asia / India / Delhi | Delhi | 2020-03-18 | National Centre for Disease control (NCDC), CSIR-Institute of Genomics and Integrative Biology (CSIR-IGIB) |
| 71 | 435069 | EPI_ISL_435069 | hCoV-19/India/NCDC-01744/2020 | Asia / India / Delhi | Delhi | 2020-03-18 | National Centre for Disease control (NCDC), CSIR-Institute of Genomics and Integrative Biology (CSIR-IGIB) |
| 72 | 435070 | EPI_ISL_435070 | hCoV-19/India/NCDC-01710/2020 | Asia / India / Delhi | Delhi | 2020-03-18 | National Centre for Disease control (NCDC), CSIR-Institute of Genomics and Integrative Biology (CSIR-IGIB) |
| 73 | 435071 | EPI_ISL_435071 | hCoV-19/India/NCDC-02105/2020 | Asia / India / Delhi | Delhi | 2020-03-25 | National Centre for Disease control (NCDC), CSIR-Institute of Genomics and Integrative Biology (CSIR-IGIB) |
| 74 | 435072 | EPI_ISL_435072 | hCoV-19/India/NCDC-02155/2020 | Asia / India / Delhi | Delhi | 2020-03-26 | National Centre for Disease control (NCDC), CSIR-Institute of Genomics and Integrative Biology (CSIR-IGIB) |
| 75 | 435074 | EPI_ISL_435074 | hCoV-19/India/NCDC-02250/2020 | Asia / India / West Bengal | West Bengal | 2020-03-28 | National Centre for Disease control (NCDC), CSIR-Institute of Genomics and Integrative Biology (CSIR-IGIB) |

| | | | | | | | |
|---|---|---|---|---|---|---|---|
| 76 | 435078 | EPI_ISL_435078 | hCoV-19/India/NCDC-02311/2020 | Asia / India / Tamil Nadu | Tamil Nadu | 2020-03-29 | National Centre for Disease control (NCDC), CSIR-Institute of Genomics and Integrative Biology (CSIR-IGIB) |
| 77 | 435080 | EPI_ISL_435080 | hCoV-19/India/NCDC-02334/2020 | Asia / India / Tamil Nadu | Tamil Nadu | 2020-03-29 | National Centre for Disease control (NCDC), CSIR-Institute of Genomics and Integrative Biology (CSIR-IGIB) |
| 78 | 435081 | EPI_ISL_435081 | hCoV-19/India/NCDC-02248/2020 | Asia / India / West Bengal | West Bengal | 2020-03-28 | National Centre for Disease control (NCDC), CSIR-Institute of Genomics and Integrative Biology (CSIR-IGIB) |
| 79 | 435082 | EPI_ISL_435082 | hCoV-19/India/NCDC-02322/2020 | Asia / India / Uttar Pradesh | Uttar Pradesh | 2020-03-29 | National Centre for Disease control (NCDC), CSIR-Institute of Genomics and Integrative Biology (CSIR-IGIB) |
| 80 | 435083 | EPI_ISL_435083 | hCoV-19/India/NCDC-02242/2020 | Asia / India / Tamil Nadu | Tamil Nadu | 2020-03-28 | National Centre for Disease control (NCDC), CSIR-Institute of Genomics and Integrative Biology (CSIR-IGIB) |
| 81 | 435084 | EPI_ISL_435084 | hCoV-19/India/NCDC-02309/2020 | Asia / India / Tamil Nadu | Tamil Nadu | 2020-03-29 | National Centre for Disease control (NCDC), CSIR-Institute of Genomics and Integrative Biology (CSIR-IGIB) |
| 82 | 435085 | EPI_ISL_435085 | hCoV-19/India/NCDC-02310/2020 | Asia / India / Mumbai | Maharashtra | 2020-03-29 | National Centre for Disease control (NCDC), CSIR-Institute of Genomics and Integrative Biology (CSIR-IGIB) |
| 83 | 435086 | EPI_ISL_435086 | hCoV-19/India/NCDC-02315/2020 | Asia / India / Mumbai | Maharashtra | 2020-03-29 | National Centre for Disease control (NCDC), CSIR-Institute of Genomics and Integrative Biology (CSIR-IGIB) |
| 84 | 435087 | EPI_ISL_435087 | hCoV-19/India/NCDC-02333/2020 | Asia / India / Tamil Nadu | Tamil Nadu | 2020-03-29 | National Centre for Disease control (NCDC), CSIR-Institute of Genomics and Integrative Biology (CSIR-IGIB) |
| 85 | 435088 | EPI_ISL_435088 | hCoV-19/India/NCDC-02336/2020 | Asia / India / Odisha | Odisha | 2020-03-29 | National Centre for Disease control (NCDC), CSIR-Institute of Genomics and Integrative Biology (CSIR-IGIB) |
| 86 | 435090 | EPI_ISL_435090 | hCoV-19/India/NCDC-02326/2020 | Asia / India / Jammu | Jammu | 2020-03-29 | National Centre for Disease control (NCDC), CSIR-Institute of Genomics and Integrative Biology (CSIR-IGIB) |
| 87 | 435091 | EPI_ISL_435091 | hCoV-19/India/NCDC-02245/2020 | Asia / India / Tamil Nadu | Tamil Nadu | 2020-03-28 | National Centre for Disease control (NCDC), CSIR-Institute of Genomics and Integrative Biology (CSIR-IGIB) |
| 88 | 435092 | EPI_ISL_435092 | hCoV-19/India/NCDC-02244/2020 | Asia / India / Tamil Nadu | Tamil Nadu | 2020-03-28 | National Centre for Disease control (NCDC), CSIR-Institute of Genomics and Integrative Biology (CSIR-IGIB) |
| 89 | 435093 | EPI_ISL_435093 | hCoV-19/India/NCDC-02312/2020 | Asia / India / Tamil Nadu | Tamil Nadu | 2020-03-29 | National Centre for Disease control (NCDC), CSIR-Institute of Genomics and Integrative Biology (CSIR-IGIB) |
| 90 | 435094 | EPI_ISL_435094 | hCoV-19/India/NCDC-02327/2020 | Asia / India / Tamil Nadu | Tamil Nadu | 2020-03-29 | National Centre for Disease control (NCDC), CSIR-Institute of Genomics and Integrative Biology (CSIR-IGIB) |
| 91 | 435095 | EPI_ISL_435095 | hCoV-19/India/NCDC-02328/2020 | Asia / India / Tamil Nadu | Tamil Nadu | 2020-03-29 | National Centre for Disease control (NCDC), CSIR-Institute of Genomics and Integrative Biology (CSIR-IGIB) |
| 92 | 435096 | EPI_ISL_435096 | hCoV-19/India/NCDC-02332/2020 | Asia / India / Tamil Nadu | Tamil Nadu | 2020-03-29 | National Centre for Disease control (NCDC), CSIR-Institute of Genomics and Integrative Biology (CSIR-IGIB) |
| 93 | 435097 | EPI_ISL_435097 | hCoV-19/India/NCDC-02252/2020 | Asia / India / West Bengal | West Bengal | 2020-03-28 | National Centre for Disease control (NCDC), CSIR-Institute of Genomics and Integrative Biology (CSIR-IGIB) |
| 94 | 435098 | EPI_ISL_435098 | hCoV-19/India/NCDC-02240/2020 | Asia / India | Delhi | 2020-03-28 | National Centre for Disease control (NCDC), CSIR-Institute of Genomics and Integrative Biology (CSIR-IGIB) |
| 95 | 435099 | EPI_ISL_435099 | hCoV-19/India/NCDC-02320/2020 | Asia / India / Uttar Pradesh | Uttar Pradesh | 2020-03-29 | National Centre for Disease control (NCDC), CSIR-Institute of Genomics and Integrative Biology (CSIR-IGIB) |
| 96 | 435100 | EPI_ISL_435100 | hCoV-19/India/NCDC-02321/2020 | Asia / India / Uttar Pradesh | Uttar Pradesh | 2020-03-29 | National Centre for Disease control (NCDC), CSIR-Institute of Genomics and Integrative Biology (CSIR-IGIB) |
| 97 | 435101 | EPI_ISL_435101 | hCoV-19/India/NCDC-01441/2020 | Asia / India / Ladakh | Ladakh | 2020-03-15 | National Centre for Disease control (NCDC), CSIR-Institute of Genomics and Integrative Biology (CSIR-IGIB) |
| 98 | 435102 | EPI_ISL_435102 | hCoV-19/India/NCDC-01444/2020 | Asia / India / Ladakh | Ladakh | 2020-03-15 | National Centre for Disease control (NCDC), CSIR-Institute of Genomics and Integrative Biology (CSIR-IGIB) |
| 99 | 435103 | EPI_ISL_435103 | hCoV-19/India/NCDC-01604/2020 | Asia / India / Ladakh | Ladakh | 2020-03-17 | National Centre for Disease control (NCDC), CSIR-Institute of Genomics and Integrative Biology (CSIR-IGIB) |
| 100 | 435104 | EPI_ISL_435104 | hCoV-19/India/NCDC-01614/2020 | Asia / India / Ladakh | Ladakh | 2020-03-17 | National Centre for Disease control (NCDC), CSIR-Institute of Genomics and Integrative Biology (CSIR-IGIB) |
| 101 | 435105 | EPI_ISL_435105 | hCoV-19/India/NCDC-01616/2020 | Asia / India / Ladakh | Ladakh | 2020-03-17 | National Centre for Disease control (NCDC), CSIR-Institute of Genomics and Integrative Biology (CSIR-IGIB) |
| 102 | 435106 | EPI_ISL_435106 | hCoV-19/India/NCDC-01760/2020 | Asia / India / Ladakh | Ladakh | 2020-03-18 | National Centre for Disease control (NCDC), CSIR-Institute of Genomics and Integrative Biology (CSIR-IGIB) |
| 103 | 435108 | EPI_ISL_435108 | hCoV-19/India/NCDC-01457/2020 | Asia / India / Delhi | Delhi | 2020-03-15 | National Centre for Disease control (NCDC), CSIR-Institute of Genomics and Integrative Biology (CSIR-IGIB) |
| 104 | 435109 | EPI_ISL_435109 | hCoV-19/India/NCDC-01638/2020 | Asia / India / Delhi | Delhi | 2020-03-17 | National Centre for Disease control (NCDC), CSIR-Institute of Genomics and Integrative Biology (CSIR-IGIB) |
| 105 | 435110 | EPI_ISL_435110 | hCoV-19/India/NCDC-01501/2020 | Asia / India / Delhi | Delhi | 2020-03-16 | National Centre for Disease control (NCDC), CSIR-Institute of Genomics and Integrative Biology (CSIR-IGIB) |
| 106 | 435111 | EPI_ISL_435111 | hCoV-19/India/NCDC-02415/2020 | Asia / India / Delhi | Delhi | 2020-03-30 | National Centre for Disease control (NCDC), CSIR-Institute of Genomics and Integrative Biology (CSIR-IGIB) |
| 107 | 435112 | EPI_ISL_435112 | hCoV-19/India/NCDC-02251/2020 | Asia / India / Bihar | Bihar | 2020-03-28 | National Centre for Disease control (NCDC), CSIR-Institute of Genomics and Integrative Biology (CSIR-IGIB) |
| 108 | 436413 | EPI_ISL_436413 | hCoV-19/India/NCDC-2443/2020 | Asia / India / Uttar Pradesh | Uttar Pradesh | 2020-03-30 | National Centre for Disease control (NCDC) |
| 109 | 436414 | EPI_ISL_436414 | hCoV-19/India/NCDC-2502/2020 | Asia / India / West Bengal | West Bengal | 2020-03-31 | National Centre for Disease control (NCDC) |
| 110 | 436415 | EPI_ISL_436415 | hCoV-19/India/NCDC-2507/2020 | Asia / India / Delhi | Delhi | 2020-03-31 | National Centre for Disease control (NCDC) |
| 111 | 436417 | EPI_ISL_436417 | hCoV-19/India/NCDC-2515/2020 | Asia / India / Bihar | Bihar | 2020-03-31 | National Centre for Disease control (NCDC) |
| 112 | 436418 | EPI_ISL_436418 | hCoV-19/India/NCDC-2517/2020 | Asia / India / Tamil Nadu | Tamil Nadu | 2020-03-31 | National Centre for Disease control (NCDC) |
| 113 | 436419 | EPI_ISL_436419 | hCoV-19/India/NCDC-2519/2020 | Asia / India / Bihar | Bihar | 2020-03-31 | National Centre for Disease control (NCDC) |
| 114 | 436420 | EPI_ISL_436420 | hCoV-19/India/NCDC-2522/2020 | Asia / India / Rajasthan | Rajasthan | 2020-03-31 | National Centre for Disease control (NCDC) |
| 115 | 436421 | EPI_ISL_436421 | hCoV-19/India/NCDC-2525/2020 | Asia / India / Assam | Assam | 2020-03-31 | National Centre for Disease control (NCDC) |
| 116 | 436422 | EPI_ISL_436422 | hCoV-19/India/NCDC-2535/2020 | Asia / India / Assam | Assam | 2020-03-31 | National Centre for Disease control (NCDC) |
| 117 | 436424 | EPI_ISL_436424 | hCoV-19/India/NCDC-3163/2020 | Asia / India / Delhi | Delhi | 2020-04-05 | National Centre for Disease control (NCDC) |
| 118 | 436425 | EPI_ISL_436425 | hCoV-19/India/NCDC-3170/2020 | Asia / India / Delhi | Delhi | 2020-04-05 | National Centre for Disease control (NCDC) |
| 119 | 436426 | EPI_ISL_436426 | hCoV-19/India/NCDC-3175/2020 | Asia / India / Delhi | Delhi | 2020-04-05 | National Centre for Disease control (NCDC) |
| 120 | 436428 | EPI_ISL_436428 | hCoV-19/India/NCDC-3181/2020 | Asia / India / Delhi | Delhi | 2020-04-05 | National Centre for Disease control (NCDC) |
| 121 | 436429 | EPI_ISL_436429 | hCoV-19/India/NCDC-3190/2020 | Asia / India / Delhi | Delhi | 2020-04-05 | National Centre for Disease control (NCDC) |
| 122 | 436430 | EPI_ISL_436430 | hCoV-19/India/NCDC-3195/2020 | Asia / India / Delhi | Delhi | 2020-04-05 | National Centre for Disease control (NCDC) |
| 123 | 436431 | EPI_ISL_436431 | hCoV-19/India/NCDC-3230/2020 | Asia / India / Delhi | Delhi | 2020-04-05 | National Centre for Disease control (NCDC) |
| 124 | 436432 | EPI_ISL_436432 | hCoV-19/India/NCDC-3264/2020 | Asia / India / Delhi | Delhi | 2020-04-05 | National Centre for Disease control (NCDC) |
| 125 | 436433 | EPI_ISL_436433 | hCoV-19/India/NCDC-3271/2020 | Asia / India / Delhi | Delhi | 2020-04-05 | National Centre for Disease control (NCDC) |
| 126 | 436434 | EPI_ISL_436434 | hCoV-19/India/NCDC-3279/2020 | Asia / India / Delhi | Delhi | 2020-04-05 | National Centre for Disease control (NCDC) |
| 127 | 436435 | EPI_ISL_436435 | hCoV-19/India/NCDC-3394/2020 | Asia / India / Delhi | Delhi | 2020-04-09 | National Centre for Disease control (NCDC) |
| 128 | 436436 | EPI_ISL_436436 | hCoV-19/India/NCDC-3400/2020 | Asia / India / Delhi | Delhi | 2020-04-06 | National Centre for Disease control (NCDC) |
| 129 | 436437 | EPI_ISL_436437 | hCoV-19/India/NCDC-3831/2020 | Asia / India / Delhi | Delhi | 2020-04-08 | National Centre for Disease control (NCDC) |
| 130 | 436440 | EPI_ISL_436440 | hCoV-19/India/NCDC-3936/2020 | Asia / India / Andhra Pradesh | Andhra Pradesh | 2020-04-09 | National Centre for Disease control (NCDC) |
| 131 | 436444 | EPI_ISL_436444 | hCoV-19/India/NCDC-3965/2020 | Asia / India / Maharashtra | Maharashtra | 2020-04-09 | National Centre for Disease control (NCDC) |
| 132 | 436445 | EPI_ISL_436445 | hCoV-19/India/NCDC-3982/2020 | Asia / India / Delhi | Delhi | 2020-04-09 | National Centre for Disease control (NCDC) |
| 133 | 436447 | EPI_ISL_436447 | hCoV-19/India/NCDC-4055/2020 | Asia / India / Karnataka | Karnataka | 2020-04-10 | National Centre for Disease control (NCDC) |
| 134 | 436448 | EPI_ISL_436448 | hCoV-19/India/NCDC-4138/2020 | Asia / India / Delhi | Delhi | 2020-04-12 | National Centre for Disease control (NCDC) |
| 135 | 436449 | EPI_ISL_436449 | hCoV-19/India/NCDC-4155/2020 | Asia / India / Bihar | Bihar | 2020-04-12 | National Centre for Disease control (NCDC) |
| 136 | 436450 | EPI_ISL_436450 | hCoV-19/India/NCDC-4189/2020 | Asia / India / Delhi | Delhi | 2020-04-13 | National Centre for Disease control (NCDC) |
| 137 | 436451 | EPI_ISL_436451 | hCoV-19/India/NCDC-4205/2020 | Asia / India / Delhi | Delhi | 2020-04-13 | National Centre for Disease control (NCDC) |
| 138 | 436452 | EPI_ISL_436452 | hCoV-19/India/NCDC-4208/2020 | Asia / India / Delhi | Delhi | 2020-04-14 | National Centre for Disease control (NCDC) |
| 139 | 436453 | EPI_ISL_436453 | hCoV-19/India/NCDC-4444/2020 | Asia / India / Madhya Pradesh | Madhya Pradesh | 2020-04-16 | National Centre for Disease control (NCDC) |
| 140 | 436454 | EPI_ISL_436454 | hCoV-19/India/NCDC-4475/2020 | Asia / India / Delhi | Delhi | 2020-04-13 | National Centre for Disease control (NCDC) |
| 141 | 436455 | EPI_ISL_436455 | hCoV-19/India/NCDC-4476/2020 | Asia / India / Delhi | Delhi | 2020-04-13 | National Centre for Disease control (NCDC) |
| 142 | 436456 | EPI_ISL_436456 | hCoV-19/India/NCDC-4846/2020 | Asia / India / Madhya Pradesh | Madhya Pradesh | 2020-04-20 | National Centre for Disease control (NCDC) |
| 143 | 436457 | EPI_ISL_436457 | hCoV-19/India/NCDC-4859/2020 | Asia / India / Madhya Pradesh | Madhya Pradesh | 2020-04-20 | National Centre for Disease control (NCDC) |
| 144 | 436458 | EPI_ISL_436458 | hCoV-19/India/NCDC-4867/2020 | Asia / India / Madhya Pradesh | Madhya Pradesh | 2020-04-16 | National Centre for Disease control (NCDC) |
| 145 | 436459 | EPI_ISL_436459 | hCoV-19/India/NCDC-4874/2020 | Asia / India / Madhya Pradesh | Madhya Pradesh | 2020-04-20 | National Centre for Disease control (NCDC) |
| 146 | 436460 | EPI_ISL_436460 | hCoV-19/India/NCDC-4876/2020 | Asia / India / Madhya Pradesh | Madhya Pradesh | 2020-04-20 | National Centre for Disease control (NCDC) |
| 147 | 436461 | EPI_ISL_436461 | hCoV-19/India/NCDC-4877/2020 | Asia / India / Madhya Pradesh | Madhya Pradesh | 2020-04-20 | National Centre for Disease control (NCDC) |
| 148 | 436462 | EPI_ISL_436462 | hCoV-19/India/NCDC-4878/2020 | Asia / India / Madhya Pradesh | Madhya Pradesh | 2020-04-16 | National Centre for Disease control (NCDC) |
| 149 | 436463 | EPI_ISL_436463 | hCoV-19/India/NCDC-4879/2020 | Asia / India / Madhya Pradesh | Madhya Pradesh | 2020-04-16 | National Centre for Disease control (NCDC) |
| 150 | 437438 | EPI_ISL_437438 | hCoV-19/India/GBRC10/2020 | Asia / India / Gujarat / Surat | Gujarat | 2020-04-24 | Department of MicroBiology, Government Medical College, Surat |
| 151 | 437440 | EPI_ISL_437440 | hCoV-19/India/GBRC12/2020 | Asia / India / Gujarat / Surat | Gujarat | 2020-04-26 | Department of MicroBiology, Government Medical College, Surat |
| 152 | 437441 | EPI_ISL_437441 | hCoV-19/India/GBRC13/2020 | Asia / India / Gujarat / Surat | Gujarat | 2020-04-26 | Department of MicroBiology, Government Medical College, Surat |

| | | | | | | | |
|---|---|---|---|---|---|---|---|
| 153 | 437442 | EPI_ISL_437442 | hCoV-19/India/GBRC14/2020 | Asia / India / Gujarat / Surat | Gujarat | 2020-04-27 | Department of MicroBiology, Government Medical College, Surat |
| 154 | 437444 | EPI_ISL_437444 | hCoV-19/India/GBRC16/2020 | Asia / India / Gujarat / Surat | Gujarat | 2020-04-26 | Department of MicroBiology, Government Medical College, Surat |
| 155 | 437445 | EPI_ISL_437445 | hCoV-19/India/GBRC17a/2020 | Asia / India / Gujarat / Ahmedabad | Gujarat | 2020-04-26 | B.J. Medical College and Civil hospital |
| 156 | 437446 | EPI_ISL_437446 | hCoV-19/India/GBRC17b/2020 | Asia / India / Gujarat / Ahmedabad | Gujarat | 2020-04-26 | B.J. Medical College and Civil hospital |
| 157 | 437447 | EPI_ISL_437447 | hCoV-19/India/GBRC18/2020 | Asia / India / Gujarat / Ahmedabad | Gujarat | 2020-04-26 | B.J. Medical College and Civil hospital |
| 158 | 437448 | EPI_ISL_437448 | hCoV-19/India/GBRC19/2020 | Asia / India / Gujarat / Ahmedabad | Gujarat | 2020-04-26 | B.J. Medical College and Civil hospital |
| 159 | 437449 | EPI_ISL_437449 | hCoV-19/India/GBRC20/2020 | Asia / India / Gujarat / Ahmedabad | Gujarat | 2020-04-26 | B.J. Medical College and Civil hospital |
| 160 | 437450 | EPI_ISL_437450 | hCoV-19/India/GBRC21/2020 | Asia / India / Gujarat / Ahmedabad | Gujarat | 2020-04-26 | B.J. Medical College and Civil hospital |
| 161 | 437451 | EPI_ISL_437451 | hCoV-19/India/GBRC22/2020 | Asia / India / Gujarat / Ahmedabad | Gujarat | 2020-04-26 | B.J. Medical College and Civil hospital |
| 162 | 437452 | EPI_ISL_437452 | hCoV-19/India/GBRC23/2020 | Asia / India / Gujarat / Ahmedabad | Gujarat | 2020-04-26 | B.J. Medical College and Civil hospital |
| 163 | 437453 | EPI_ISL_437453 | hCoV-19/India/GBRC24a/2020 | Asia / India / Gujarat / Ahmedabad | Gujarat | 2020-04-26 | B.J. Medical College and Civil hospital |
| 164 | 437454 | EPI_ISL_437454 | hCoV-19/India/GBRC24b/2020 | Asia / India / Gujarat / Ahmedabad | Gujarat | 2020-04-26 | B.J. Medical College and Civil hospital |
| 165 | 437539 | EPI_ISL_437539 | hCoV-19/India/S12/2020 | Asia / India / West Bengal / East Medinipur | West Bengal | 2020-04-06 | ICMR-National Institute of Cholera and Enteric Diseases |
| 166 | 437626 | EPI_ISL_437626 | hCoV-19/India/GMC-KP1125/2020 | Asia / India / Telangana / Hyderabad | Telangana | 2020-03-24 | Department of Microbiology,Gandhi Medical College and Hospital |
| 167 | 438138 | EPI_ISL_438138 | hCoV-19/India/GMC-RR1191/2020 | Asia / India / Telangana / Hyderabad | Telangana | 2020-03-25 | Department of Microbiology,Gandhi Medical College and Hospital |
| 168 | 438139 | EPI_ISL_438139 | hCoV-19/India/GMC-RK1090/2020 | Asia / India / Telangana / Hyderabad | Telangana | 2020-03-25 | Department of Microbiology,Gandhi Medical College and Hospital,Hyderabad |
| 169 | 444456 | EPI_ISL_444456 | hCoV-19/India/GBRC25/2020 | Asia / India / Gujarat / Ahmedabad | Gujarat | 2020-04-26 | B.J. Medical College and Civil hospital |
| 170 | 444457 | EPI_ISL_444457 | hCoV-19/India/GBRC26/2020 | Asia / India / Gujarat / Ahmedabad | Gujarat | 2020-04-26 | B.J. Medical College and Civil hospital |
| 171 | 444458 | EPI_ISL_444458 | hCoV-19/India/GBRC27/2020 | Asia / India / Gujarat / Ahmedabad | Gujarat | 2020-04-26 | B.J. Medical College and Civil hospital |
| 172 | 444459 | EPI_ISL_444459 | hCoV-19/India/GBRC28/2020 | Asia / India / Gujarat / Ahmedabad | Gujarat | 2020-04-30 | B.J. Medical College and Civil hospital |
| 173 | 444460 | EPI_ISL_444460 | hCoV-19/India/GBRC29/2020 | Asia / India / Gujarat / Ahmedabad | Gujarat | 2020-04-30 | B.J. Medical College and Civil hospital |
| 174 | 444461 | EPI_ISL_444461 | hCoV-19/India/GBRC30/2020 | Asia / India / Gujarat / Ahmedabad | Gujarat | 2020-04-29 | B.J. Medical College and Civil hospital |
| 175 | 444462 | EPI_ISL_444462 | hCoV-19/India/GBRC31/2020 | Asia / India / Gujarat / Ahmedabad | Gujarat | 2020-04-29 | B.J. Medical College and Civil hospital |
| 176 | 444463 | EPI_ISL_444463 | hCoV-19/India/GBRC32/2020 | Asia / India / Gujarat / Ahmedabad | Gujarat | 2020-04-29 | B.J. Medical College and Civil hospital |
| 177 | 444464 | EPI_ISL_444464 | hCoV-19/India/GBRC33/2020 | Asia / India / Gujarat / Ahmedabad | Gujarat | 2020-04-29 | B.J. Medical College and Civil hospital |
| 178 | 444465 | EPI_ISL_444465 | hCoV-19/India/GBRC34/2020 | Asia / India / Gujarat / Ahmedabad | Gujarat | 2020-04-29 | B.J. Medical College and Civil hospital |
| 179 | 444466 | EPI_ISL_444466 | hCoV-19/India/GBRC35/2020 | Asia / India / Gujarat / Ahmedabad | Gujarat | 2020-04-29 | B.J. Medical College and Civil hospital |
| 180 | 444467 | EPI_ISL_444467 | hCoV-19/India/GBRC36/2020 | Asia / India / Gujarat / Ahmedabad | Gujarat | 2020-04-29 | B.J. Medical College and Civil hospital |
| 181 | 444468 | EPI_ISL_444468 | hCoV-19/India/GBRC37/2020 | Asia / India / Gujarat / Ahmedabad | Gujarat | 2020-04-29 | B.J. Medical College and Civil hospital |
| 182 | 444469 | EPI_ISL_444469 | hCoV-19/India/GBRC38/2020 | Asia / India / Gujarat / Ahmedabad | Gujarat | 2020-04-29 | B.J. Medical College and Civil hospital |
| 183 | 444470 | EPI_ISL_444470 | hCoV-19/India/GBRC39/2020 | Asia / India / Gujarat / Ahmedabad | Gujarat | 2020-04-29 | B.J. Medical College and Civil hospital |
| 184 | 444471 | EPI_ISL_444471 | hCoV-19/India/GBRC40/2020 | Asia / India / Gujarat / Ahmedabad | Gujarat | 2020-04-29 | B.J. Medical College and Civil hospital |
| 185 | 444472 | EPI_ISL_444472 | hCoV-19/India/GBRC41/2020 | Asia / India / Gujarat / Ahmedabad | Gujarat | 2020-04-29 | B.J. Medical College and Civil hospital |
| 186 | 444473 | EPI_ISL_444473 | hCoV-19/India/GBRC42/2020 | Asia / India / Gujarat / Ahmedabad | Gujarat | 2020-04-29 | B.J. Medical College and Civil hospital |
| 187 | 444474 | EPI_ISL_444474 | hCoV-19/India/GBRC43/2020 | Asia / India / Gujarat / Ahmedabad | Gujarat | 2020-04-29 | B.J. Medical College and Civil hospital |
| 188 | 444475 | EPI_ISL_444475 | hCoV-19/India/GBRC44/2020 | Asia / India / Gujarat / Ahmedabad | Gujarat | 2020-04-29 | B.J. Medical College and Civil hospital |
| 189 | 444476 | EPI_ISL_444476 | hCoV-19/India/GBRC45/2020 | Asia / India / Gujarat / Ahmedabad | Gujarat | 2020-04-29 | B.J. Medical College and Civil hospital |
| 190 | 444477 | EPI_ISL_444477 | hCoV-19/India/GBRC46/2020 | Asia / India / Gujarat / Ahmedabad | Gujarat | 2020-04-29 | B.J. Medical College and Civil hospital |
| 191 | 444478 | EPI_ISL_444478 | hCoV-19/India/GBRC47/2020 | Asia / India / Gujarat / Ahmedabad | Gujarat | 2020-04-30 | B.J. Medical College and Civil hospital |
| 192 | 444479 | EPI_ISL_444479 | hCoV-19/India/GBRC48/2020 | Asia / India / Gujarat / Ahmedabad | Gujarat | 2020-04-29 | B.J. Medical College and Civil hospital |
| 193 | 444480 | EPI_ISL_444480 | hCoV-19/India/GBRC49/2020 | Asia / India / Gujarat / Ahmedabad | Gujarat | 2020-04-30 | B.J. Medical College and Civil hospital |
| 194 | 444481 | EPI_ISL_444481 | hCoV-19/India/GBRC50/2020 | Asia / India / Gujarat / Ahmedabad | Gujarat | 2020-04-30 | B.J. Medical College and Civil hospital |
| 195 | 444482 | EPI_ISL_444482 | hCoV-19/India/GBRC51/2020 | Asia / India / Gujarat / Prantij | Gujarat | 2020-05-02 | Gujarat Biotechnology Research Centre |
| 196 | 444483 | EPI_ISL_444483 | hCoV-19/India/GBRC52/2020 | Asia / India / Gujarat / Prantij | Gujarat | 2020-05-02 | Gujarat Biotechnology Research Centre |
| 197 | 444484 | EPI_ISL_444484 | hCoV-19/India/GBRC53/2020 | Asia / India / Gujarat / Modasa | Gujarat | 2020-05-04 | Gujarat Biotechnology Research Centre |
| 198 | 444485 | EPI_ISL_444485 | hCoV-19/India/GBRC54/2020 | Asia / India / Gujarat / Modasa | Gujarat | 2020-05-04 | Gujarat Biotechnology Research Centre |
| 199 | 444486 | EPI_ISL_444486 | hCoV-19/India/GBRC55/2020 | Asia / India / Gujarat / Dhansura | Gujarat | 2020-05-04 | Gujarat Biotechnology Research Centre |
| 200 | 447030 | EPI_ISL_447030 | hCoV-19/India/GBRC56a/2020 | Asia / India / Gujarat / Ahmedabad | Gujarat | 2020-05-03 | B.J. Medical College and Civil hospital |
| 201 | 447031 | EPI_ISL_447031 | hCoV-19/India/GBRC56b/2020 | Asia / India / Gujarat / Ahmedabad | Gujarat | 2020-05-03 | B.J. Medical College and Civil hospital |
| 202 | 447032 | EPI_ISL_447032 | hCoV-19/India/GBRC57/2020 | Asia / India / Gujarat / Ahmedabad | Gujarat | 2020-05-03 | B.J. Medical College and Civil hospital |
| 203 | 447033 | EPI_ISL_447033 | hCoV-19/India/GBRC58/2020 | Asia / India / Gujarat / Ahmedabad | Gujarat | 2020-05-03 | B.J. Medical College and Civil hospital |
| 204 | 447034 | EPI_ISL_447034 | hCoV-19/India/GBRC59/2020 | Asia / India / Gujarat / Ahmedabad | Gujarat | 2020-05-03 | B.J. Medical College and Civil hospital |
| 205 | 447035 | EPI_ISL_447035 | hCoV-19/India/GBRC60/2020 | Asia / India / Gujarat / Ahmedabad | Gujarat | 2020-05-03 | B.J. Medical College and Civil hospital |
| 206 | 447036 | EPI_ISL_447036 | hCoV-19/India/GBRC61/2020 | Asia / India / Gujarat / Ahmedabad | Gujarat | 2020-05-03 | B.J. Medical College and Civil hospital |
| 207 | 447037 | EPI_ISL_447037 | hCoV-19/India/GBRC62/2020 | Asia / India / Gujarat / Ahmedabad | Gujarat | 2020-05-03 | B.J. Medical College and Civil hospital |
| 208 | 447038 | EPI_ISL_447038 | hCoV-19/India/GBRC63/2020 | Asia / India / Gujarat / Ahmedabad | Gujarat | 2020-05-03 | B.J. Medical College and Civil hospital |
| 209 | 447039 | EPI_ISL_447039 | hCoV-19/India/GBRC64/2020 | Asia / India / Gujarat / Ahmedabad | Gujarat | 2020-05-03 | B.J. Medical College and Civil hospital |
| 210 | 447040 | EPI_ISL_447040 | hCoV-19/India/GBRC65a/2020 | Asia / India / Gujarat / Ahmedabad | Gujarat | 2020-05-03 | B.J. Medical College and Civil hospital |
| 211 | 447041 | EPI_ISL_447041 | hCoV-19/India/GBRC65b/2020 | Asia / India / Gujarat / Ahmedabad | Gujarat | 2020-05-03 | B.J. Medical College and Civil hospital |
| 212 | 447042 | EPI_ISL_447042 | hCoV-19/India/GBRC66/2020 | Asia / India / Gujarat / Ahmedabad | Gujarat | 2020-05-03 | B.J. Medical College and Civil hospital |
| 213 | 447043 | EPI_ISL_447043 | hCoV-19/India/GBRC67a/2020 | Asia / India / Gujarat / Ahmedabad | Gujarat | 2020-05-03 | B.J. Medical College and Civil hospital |
| 214 | 447044 | EPI_ISL_447044 | hCoV-19/India/GBRC67b/2020 | Asia / India / Gujarat / Ahmedabad | Gujarat | 2020-05-03 | B.J. Medical College and Civil hospital |
| 215 | 447045 | EPI_ISL_447045 | hCoV-19/India/GBRC68/2020 | Asia / India / Gujarat / Ahmedabad | Gujarat | 2020-05-03 | B.J. Medical College and Civil hospital |
| 216 | 447046 | EPI_ISL_447046 | hCoV-19/India/GBRC69/2020 | Asia / India / Gujarat / Ahmedabad | Gujarat | 2020-05-04 | B.J. Medical College and Civil hospital |
| 217 | 447047 | EPI_ISL_447047 | hCoV-19/India/GBRC70/2020 | Asia / India / Gujarat / Gandhinagar | Gujarat | 2020-04-29 | GMERS Medical College and Hospital, Gandhinagar |
| 218 | 447048 | EPI_ISL_447048 | hCoV-19/India/GBRC71/2020 | Asia / India / Gujarat / Gandhinagar | Gujarat | 2020-04-27 | GMERS Medical College and Hospital, Gandhinagar |
| 219 | 447049 | EPI_ISL_447049 | hCoV-19/India/GBRC72a/2020 | Asia / India / Gujarat / Gandhinagar | Gujarat | 2020-04-29 | GMERS Medical College and Hospital, Gandhinagar |
| 220 | 447050 | EPI_ISL_447050 | hCoV-19/India/GBRC72b/2020 | Asia / India / Gujarat / Gandhinagar | Gujarat | 2020-04-29 | GMERS Medical College and Hospital, Gandhinagar |
| 221 | 447051 | EPI_ISL_447051 | hCoV-19/India/GBRC73/2020 | Asia / India / Gujarat / Gandhinagar | Gujarat | 2020-04-29 | GMERS Medical College and Hospital, Gandhinagar |
| 222 | 447052 | EPI_ISL_447052 | hCoV-19/India/GBRC74/2020 | Asia / India / Gujarat / Gandhinagar | Gujarat | 2020-05-02 | GMERS Medical College and Hospital, Gandhinagar |
| 223 | 447053 | EPI_ISL_447053 | hCoV-19/India/GBRC75/2020 | Asia / India / Gujarat / Gandhinagar | Gujarat | 2020-04-29 | GMERS Medical College and Hospital, Gandhinagar |
| 224 | 447534 | EPI_ISL_447534 | hCoV-19/India/GBRC76/2020 | Asia / India / Gujarat / Modasa | Gujarat | 2020-05-05 | Gujarat Biotechnology Research Centre |
| 225 | 447535 | EPI_ISL_447535 | hCoV-19/India/GBRC77/2020 | Asia / India / Gujarat / Modasa | Gujarat | 2020-05-05 | Gujarat Biotechnology Research Centre |
| 226 | 447536 | EPI_ISL_447536 | hCoV-19/India/GBRC78a/2020 | Asia / India / Gujarat / Modasa | Gujarat | 2020-05-05 | Gujarat Biotechnology Research Centre |
| 227 | 447537 | EPI_ISL_447537 | hCoV-19/India/GBRC78b/2020 | Asia / India / Gujarat / Modasa | Gujarat | 2020-05-05 | Gujarat Biotechnology Research Centre |
| 228 | 447538 | EPI_ISL_447538 | hCoV-19/India/GBRC79a/2020 | Asia / India / Gujarat / Modasa | Gujarat | 2020-05-05 | Gujarat Biotechnology Research Centre |
| 229 | 447539 | EPI_ISL_447539 | hCoV-19/India/GBRC79b/2020 | Asia / India / Gujarat / Modasa | Gujarat | 2020-05-05 | Gujarat Biotechnology Research Centre |

| | | | | | | | |
|---|---|---|---|---|---|---|---|
| 230 | 447540 | EPI_ISL_447540 | hCoV-19/India/GBRC81a/2020 | Asia / India / Gujarat / Modasa | Gujarat | 2020-05-05 | Gujarat Biotechnology Research Centre |
| 231 | 447541 | EPI_ISL_447541 | hCoV-19/India/GBRC81b/2020 | Asia / India / Gujarat / Modasa | Gujarat | 2020-05-05 | Gujarat Biotechnology Research Centre |
| 232 | 447542 | EPI_ISL_447542 | hCoV-19/India/GBRC82a/2020 | Asia / India / Gujarat / Modasa | Gujarat | 2020-05-05 | Gujarat Biotechnology Research Centre |
| 233 | 447543 | EPI_ISL_447543 | hCoV-19/India/GBRC82b/2020 | Asia / India / Gujarat / Modasa | Gujarat | 2020-05-05 | Gujarat Biotechnology Research Centre |
| 234 | 447544 | EPI_ISL_447544 | hCoV-19/India/GBRC83/2020 | Asia / India / Gujarat / Modasa | Gujarat | 2020-05-05 | Gujarat Biotechnology Research Centre |
| 235 | 447545 | EPI_ISL_447545 | hCoV-19/India/GBRC84/2020 | Asia / India / Gujarat / Prantij | Gujarat | 2020-05-05 | Gujarat Biotechnology Research Centre |
| 236 | 447546 | EPI_ISL_447546 | hCoV-19/India/GBRC85/2020 | Asia / India / Gujarat / Himatnagar | Gujarat | 2020-05-05 | Gujarat Biotechnology Research Centre |
| 237 | 447547 | EPI_ISL_447547 | hCoV-19/India/GBRC86/2020 | Asia / India / Gujarat / Gandhinagar | Gujarat | 2020-04-28 | GMERS Medical College and Hospital, Gandhinagar |
| 238 | 447548 | EPI_ISL_447548 | hCoV-19/India/GBRC87/2020 | Asia / India / Gujarat / Dahegam | Gujarat | 2020-04-28 | GMERS Medical College and Hospital, Gandhinagar |
| 239 | 447549 | EPI_ISL_447549 | hCoV-19/India/GBRC88a/2020 | Asia / India / Gujarat / Dahegam | Gujarat | 2020-05-03 | GMERS Medical College and Hospital, Gandhinagar |
| 240 | 447550 | EPI_ISL_447550 | hCoV-19/India/GBRC88b/2020 | Asia / India / Gujarat / Dahegam | Gujarat | 2020-05-03 | GMERS Medical College and Hospital, Gandhinagar |
| 241 | 447551 | EPI_ISL_447551 | hCoV-19/India/GBRC89/2020 | Asia / India / Gujarat / Gandhinagar | Gujarat | 2020-04-25 | GMERS Medical College and Hospital, Gandhinagar |
| 242 | 447552 | EPI_ISL_447552 | hCoV-19/India/GBRC90/2020 | Asia / India / Gujarat / Gandhinagar | Gujarat | 2020-05-02 | GMERS Medical College and Hospital, Gandhinagar |
| 243 | 447553 | EPI_ISL_447553 | hCoV-19/India/GBRC91/2020 | Asia / India / Gujarat / Gandhinagar | Gujarat | 2020-04-27 | GMERS Medical College and Hospital, Gandhinagar |
| 244 | 447554 | EPI_ISL_447554 | hCoV-19/India/GBRC92/2020 | Asia / India / Gujarat / Gandhinagar | Gujarat | 2020-04-26 | GMERS Medical College and Hospital, Gandhinagar |
| 245 | 447555 | EPI_ISL_447555 | hCoV-19/India/GBRC93/2020 | Asia / India / Gujarat / Mansa | Gujarat | 2020-04-28 | GMERS Medical College and Hospital, Gandhinagar |
| 246 | 447556 | EPI_ISL_447556 | hCoV-19/India/CCMB_J021/2020 | Asia / India / Telangana | Telangana | 2020-03-30 | CSIR-Centre for Cellular and Molecular Biology |
| 247 | 447557 | EPI_ISL_447557 | hCoV-19/India/CCMB_J009/2020 | Asia / India / Telangana | Telangana | 2020-03-30 | CSIR-Centre for Cellular and Molecular Biology |
| 248 | 447558 | EPI_ISL_447558 | hCoV-19/India/CCMB_J014/2020 | Asia / India / Telangana | Telangana | 2020-03-30 | CSIR-Centre for Cellular and Molecular Biology |
| 249 | 447559 | EPI_ISL_447559 | hCoV-19/India/CCMB_J048/2020 | Asia / India / Telangana | Telangana | 2020-03-31 | CSIR-Centre for Cellular and Molecular Biology |
| 250 | 447560 | EPI_ISL_447560 | hCoV-19/India/CCMB_J043/2020 | Asia / India / Telangana | Telangana | 2020-03-31 | CSIR-Centre for Cellular and Molecular Biology |
| 251 | 447561 | EPI_ISL_447561 | hCoV-19/India/CCMB_J067/2020 | Asia / India / Telangana | Telangana | 2020-03-31 | CSIR-Centre for Cellular and Molecular Biology |
| 252 | 447562 | EPI_ISL_447562 | hCoV-19/India/CCMB_J068/2020 | Asia / India / Telangana | Telangana | 2020-03-31 | CSIR-Centre for Cellular and Molecular Biology |
| 253 | 447563 | EPI_ISL_447563 | hCoV-19/India/CCMB_J057/2020 | Asia / India / Telangana | Telangana | 2020-03-31 | CSIR-Centre for Cellular and Molecular Biology |
| 254 | 447564 | EPI_ISL_447564 | hCoV-19/India/CCMB_J212/2020 | Asia / India / Telangana | Telangana | 2020-04-01 | CSIR-Centre for Cellular and Molecular Biology |
| 255 | 447565 | EPI_ISL_447565 | hCoV-19/India/CCMB_J278/2020 | Asia / India / Telangana | Telangana | 2020-04-01 | CSIR-Centre for Cellular and Molecular Biology |
| 256 | 447566 | EPI_ISL_447566 | hCoV-19/India/CCMB_J131/2020 | Asia / India / Telangana | Telangana | 2020-04-01 | CSIR-Centre for Cellular and Molecular Biology |
| 257 | 447567 | EPI_ISL_447567 | hCoV-19/India/CCMB_J304/2020 | Asia / India / Telangana | Telangana | 2020-04-01 | CSIR-Centre for Cellular and Molecular Biology |
| 258 | 447568 | EPI_ISL_447568 | hCoV-19/India/CCMB_J166/2020 | Asia / India / Telangana | Telangana | 2020-04-01 | CSIR-Centre for Cellular and Molecular Biology |
| 259 | 447569 | EPI_ISL_447569 | hCoV-19/India/CCMB_J206/2020 | Asia / India / Telangana | Telangana | 2020-04-01 | CSIR-Centre for Cellular and Molecular Biology |
| 260 | 447570 | EPI_ISL_447570 | hCoV-19/India/CCMB_J207/2020 | Asia / India / Telangana | Telangana | 2020-04-01 | CSIR-Centre for Cellular and Molecular Biology |
| 261 | 447571 | EPI_ISL_447571 | hCoV-19/India/CCMB_J224/2020 | Asia / India / Telangana | Telangana | 2020-04-01 | CSIR-Centre for Cellular and Molecular Biology |
| 262 | 447572 | EPI_ISL_447572 | hCoV-19/India/CCMB_J230/2020 | Asia / India / Telangana | Telangana | 2020-04-01 | CSIR-Centre for Cellular and Molecular Biology |
| 263 | 447573 | EPI_ISL_447573 | hCoV-19/India/CCMB_J118/2020 | Asia / India / Telangana | Telangana | 2020-04-01 | CSIR-Centre for Cellular and Molecular Biology |
| 264 | 447574 | EPI_ISL_447574 | hCoV-19/India/CCMB_J208/2020 | Asia / India / Telangana | Telangana | 2020-04-01 | CSIR-Centre for Cellular and Molecular Biology |
| 265 | 447575 | EPI_ISL_447575 | hCoV-19/India/CCMB_J458/2020 | Asia / India / Telangana | Telangana | 2020-04-02 | CSIR-Centre for Cellular and Molecular Biology |
| 266 | 447576 | EPI_ISL_447576 | hCoV-19/India/CCMB_J232/2020 | Asia / India / Telangana | Telangana | 2020-04-02 | CSIR-Centre for Cellular and Molecular Biology |
| 267 | 447577 | EPI_ISL_447577 | hCoV-19/India/CCMB_J321/2020 | Asia / India / Telangana | Telangana | 2020-04-02 | CSIR-Centre for Cellular and Molecular Biology |
| 268 | 447578 | EPI_ISL_447578 | hCoV-19/India/CCMB_J327/2020 | Asia / India / Telangana | Telangana | 2020-04-02 | CSIR-Centre for Cellular and Molecular Biology |
| 269 | 447579 | EPI_ISL_447579 | hCoV-19/India/CCMB_K128/2020 | Asia / India / Telangana | Telangana | 2020-04-10 | CSIR-Centre for Cellular and Molecular Biology |
| 270 | 447580 | EPI_ISL_447580 | hCoV-19/India/CCMB_K496/2020 | Asia / India / Telangana | Telangana | 2020-04-13 | CSIR-Centre for Cellular and Molecular Biology |
| 271 | 447581 | EPI_ISL_447581 | hCoV-19/India/CCMB_K600B/2020 | Asia / India / Telangana | Telangana | 2020-04-14 | CSIR-Centre for Cellular and Molecular Biology |
| 272 | 447582 | EPI_ISL_447582 | hCoV-19/India/CCMB_K601/2020 | Asia / India / Telangana | Telangana | 2020-04-14 | CSIR-Centre for Cellular and Molecular Biology |
| 273 | 447583 | EPI_ISL_447583 | hCoV-19/India/CCMB_L007/2020 | Asia / India / Telangana | Telangana | 2020-04-16 | CSIR-Centre for Cellular and Molecular Biology |
| 274 | 447584 | EPI_ISL_447584 | hCoV-19/India/CCMB_C1-12/2020 | Asia / India / Tamil Nadu | Tamil Nadu | 2020-04-16 | Tamil Nadu Veterinary and Animal Sciences University |
| 275 | 447585 | EPI_ISL_447585 | hCoV-19/India/CCMB_C2-13/2020 | Asia / India / Tamil Nadu | Tamil Nadu | 2020-04-16 | Tamil Nadu Veterinary and Animal Sciences University |
| 276 | 447586 | EPI_ISL_447586 | hCoV-19/India/CCMB_C3-14/2020 | Asia / India / Tamil Nadu | Tamil Nadu | 2020-04-16 | Tamil Nadu Veterinary and Animal Sciences University |
| 277 | 447587 | EPI_ISL_447587 | hCoV-19/India/CCMB_C4-15/2020 | Asia / India / Tamil Nadu | Tamil Nadu | 2020-04-20 | Tamil Nadu Veterinary and Animal Sciences University |
| 278 | 447847 | EPI_ISL_447847 | hCoV-19/India/CCMB_J045/2020 | Asia / India / Telangana | Telangana | 2020-03-31 | CSIR-Centre for Cellular and Molecular Biology |
| 279 | 447848 | EPI_ISL_447848 | hCoV-19/India/CCMB_J165/2020 | Asia / India / Telangana | Telangana | 2020-04-01 | CSIR-Centre for Cellular and Molecular Biology |
| 280 | 447849 | EPI_ISL_447849 | hCoV-19/India/CCMB_J107/2020 | Asia / India / Telangana | Telangana | 2020-04-01 | CSIR-Centre for Cellular and Molecular Biology |
| 281 | 447850 | EPI_ISL_447850 | hCoV-19/India/CCMB_J132/2020 | Asia / India / Telangana | Telangana | 2020-04-01 | CSIR-Centre for Cellular and Molecular Biology |
| 282 | 447851 | EPI_ISL_447851 | hCoV-19/India/CCMB_J223/2020 | Asia / India / Telangana | Telangana | 2020-04-01 | CSIR-Centre for Cellular and Molecular Biology |
| 283 | 447852 | EPI_ISL_447852 | hCoV-19/India/CCMB_J235/2020 | Asia / India / Telangana | Telangana | 2020-04-01 | CSIR-Centre for Cellular and Molecular Biology |
| 284 | 447853 | EPI_ISL_447853 | hCoV-19/India/CCMB_J247/2020 | Asia / India / Telangana | Telangana | 2020-04-01 | CSIR-Centre for Cellular and Molecular Biology |
| 285 | 447854 | EPI_ISL_447854 | hCoV-19/India/CCMB_J350/2020 | Asia / India / Telangana | Telangana | 2020-04-02 | CSIR-Centre for Cellular and Molecular Biology |
| 286 | 447855 | EPI_ISL_447855 | hCoV-19/India/CCMB_J318/2020 | Asia / India / Telangana | Telangana | 2020-04-02 | CSIR-Centre for Cellular and Molecular Biology |
| 287 | 447856 | EPI_ISL_447856 | hCoV-19/India/CCMB_J374/2020 | Asia / India / Telangana | Telangana | 2020-04-02 | CSIR-Centre for Cellular and Molecular Biology |
| 288 | 447857 | EPI_ISL_447857 | hCoV-19/India/CCMB_J390/2020 | Asia / India / Telangana | Telangana | 2020-04-02 | CSIR-Centre for Cellular and Molecular Biology |
| 289 | 447858 | EPI_ISL_447858 | hCoV-19/India/CCMB_J812/2020 | Asia / India / Telangana | Telangana | 2020-04-06 | CSIR-Centre for Cellular and Molecular Biology |
| 290 | 447859 | EPI_ISL_447859 | hCoV-19/India/CCMB_K771/2020 | Asia / India / Telangana | Telangana | 2020-04-15 | CSIR-Centre for Cellular and Molecular Biology |
| 291 | 447860 | EPI_ISL_447860 | hCoV-19/India/CCMB_K849/2020 | Asia / India / Telangana | Telangana | 2020-04-16 | CSIR-Centre for Cellular and Molecular Biology |
| 292 | 447861 | EPI_ISL_447861 | hCoV-19/India/CCMB_L149/2020 | Asia / India / Telangana | Telangana | 2020-04-16 | CSIR-Centre for Cellular and Molecular Biology |
| 293 | 447862 | EPI_ISL_447862 | hCoV-19/India/CCMB_J047/2020 | Asia / India / Telangana | Telangana | 2020-03-31 | CSIR-Centre for Cellular and Molecular Biology |
| 294 | 447863 | EPI_ISL_447863 | hCoV-19/India/CCMB_K599/2020 | Asia / India / Telangana | Telangana | 2020-04-14 | CSIR-Centre for Cellular and Molecular Biology |
| 295 | 447864 | EPI_ISL_447864 | hCoV-19/India/CCMB_L044/2020 | Asia / India / Telangana | Telangana | 2020-04-16 | CSIR-Centre for Cellular and Molecular Biology |
| 296 | 447865 | EPI_ISL_447865 | hCoV-19/India/CCMB_K499/2020 | Asia / India / Telangana | Telangana | 2020-04-14 | CSIR-Centre for Cellular and Molecular Biology |
| 297 | 447866 | EPI_ISL_447866 | hCoV-19/India/CCMB_K500/2020 | Asia / India / Telangana | Telangana | 2020-04-14 | CSIR-Centre for Cellular and Molecular Biology |
| 298 | 450321 | EPI_ISL_450321 | hCoV-19/India/CCMB_NIV1/2020 | Asia / India / Maharashtra | Maharashtra | 2020-04-04 | NIV Pune |
| 299 | 450322 | EPI_ISL_450322 | hCoV-19/India/CCMB_NIV2/2020 | Asia / India / Maharashtra | Maharashtra | 2020-04-04 | NIV Pune |
| 300 | 450323 | EPI_ISL_450323 | hCoV-19/India/CCMB_NIV3/2020 | Asia / India / Maharashtra | Maharashtra | 2020-03-27 | NIV Pune |
| 301 | 450324 | EPI_ISL_450324 | hCoV-19/India/CCMB_NIV4/2020 | Asia / India / Maharashtra | Maharashtra | 2020-03-26 | NIV Pune |
| 302 | 450325 | EPI_ISL_450325 | hCoV-19/India/CCMB_NIV5/2020 | Asia / India / Maharashtra | Maharashtra | 2020-03-17 | NIV Pune |
| 303 | 450326 | EPI_ISL_450326 | hCoV-19/India/CCMB_J125/2020 | Asia / India / Telangana | Telangana | 2020-04-01 | CSIR-Centre for Cellular and Molecular Biology |
| 304 | 450327 | EPI_ISL_450327 | hCoV-19/India/CCMB_J199/2020 | Asia / India / Telangana | Telangana | 2020-04-01 | CSIR-Centre for Cellular and Molecular Biology |
| 305 | 450328 | EPI_ISL_450328 | hCoV-19/India/CCMB_J283/2020 | Asia / India / Telangana | Telangana | 2020-04-01 | CSIR-Centre for Cellular and Molecular Biology |
| 306 | 450329 | EPI_ISL_450329 | hCoV-19/India/CCMB_J288/2020 | Asia / India / Telangana | Telangana | 2020-04-01 | CSIR-Centre for Cellular and Molecular Biology |

| | | | | | | | |
|---|---|---|---|---|---|---|---|
| 307 | 450330 | EPI_ISL_450330 | hCoV-19/India/CCMB_J375/2020 | Asia / India / Telangana | Telangana | 2020-04-02 | CSIR-Centre for Cellular and Molecular Biology |
| 308 | 450331 | EPI_ISL_450331 | hCoV-19/India/CCMB_J220/2020 | Asia / India / Telangana | Telangana | 2020-04-01 | CSIR-Centre for Cellular and Molecular Biology |
| 309 | 450332 | EPI_ISL_450332 | hCoV-19/India/CCMB_J222/2020 | Asia / India / Telangana | Telangana | 2020-04-01 | CSIR-Centre for Cellular and Molecular Biology |
| 310 | 450781 | EPI_ISL_450781 | hCoV-19/India/GBRC94/2020 | Asia / India / Gujarat / Junagadh | Gujarat | 2020-05-09 | Government Medical College-Bhavnagar |
| 311 | 450782 | EPI_ISL_450782 | hCoV-19/India/GBRC95a/2020 | Asia / India / Gujarat / Ahmedabad | Gujarat | 2020-04-11 | Government Medical College-Bhavnagar |
| 312 | 450783 | EPI_ISL_450783 | hCoV-19/India/GBRC95b/2020 | Asia / India / Gujarat / Ahmedabad | Gujarat | 2020-04-11 | Government Medical College-Bhavnagar |
| 313 | 450784 | EPI_ISL_450784 | hCoV-19/India/GBRC96/2020 | Asia / India / Gujarat / Botad | Gujarat | 2020-05-09 | Government Medical College-Bhavnagar |
| 314 | 450785 | EPI_ISL_450785 | hCoV-19/India/GBRC97a/2020 | Asia / India / Gujarat / Una | Gujarat | 2020-05-10 | Pandit Deendayal Upadhyay Government Medical College, Rajkot |
| 315 | 450786 | EPI_ISL_450786 | hCoV-19/India/GBRC97b/2020 | Asia / India / Gujarat / Una | Gujarat | 2020-05-10 | Pandit Deendayal Upadhyay Government Medical College, Rajkot |
| 316 | 450787 | EPI_ISL_450787 | hCoV-19/India/GBRC98/2020 | Asia / India / Gujarat / Una | Gujarat | 2020-05-11 | Pandit Deendayal Upadhyay Government Medical College, Rajkot |
| 317 | 450788 | EPI_ISL_450788 | hCoV-19/India/GBRC99/2020 | Asia / India / Gujarat / Rajkot | Gujarat | 2020-05-06 | Pandit Deendayal Upadhyay Government Medical College, Rajkot |
| 318 | 450789 | EPI_ISL_450789 | hCoV-19/India/GBRC100/2020 | Asia / India / Gujarat / Kodinar | Gujarat | 2020-05-10 | Pandit Deendayal Upadhyay Government Medical College, Rajkot |
| 319 | 450790 | EPI_ISL_450790 | hCoV-19/India/GBRC101/2020 | Asia / India / Gujarat / Rajkot | Gujarat | 2020-04-28 | Pandit Deendayal Upadhyay Government Medical College, Rajkot |
| 320 | 450791 | EPI_ISL_450791 | hCoV-19/India/GBRC102/2020 | Asia / India / Gujarat / Rajkot | Gujarat | 2020-04-27 | Pandit Deendayal Upadhyay Government Medical College, Rajkot |
| 321 | 451149 | EPI_ISL_451149 | hCoV-19/India/GBRC104/2020 | Asia / India / Gujarat / Jamnagar | Gujarat | 2020-05-07 | M.P Shah Government Medocal college Jamnagar |
| 322 | 451150 | EPI_ISL_451150 | hCoV-19/India/GBRC105/2020 | Asia / India / Gujarat / Jamnagar | Gujarat | 2020-05-06 | M.P Shah Government Medocal college Jamnagar |
| 323 | 451151 | EPI_ISL_451151 | hCoV-19/India/GBRC106/2020 | Asia / India / Gujarat / Jamnagar | Gujarat | 2020-05-05 | M.P Shah Government Medocal college Jamnagar |
| 324 | 451152 | EPI_ISL_451152 | hCoV-19/India/GBRC107/2020 | Asia / India / Gujarat / Jamnagar | Gujarat | 2020-05-09 | M.P Shah Government Medocal college Jamnagar |
| 325 | 451153 | EPI_ISL_451153 | hCoV-19/India/GBRC108/2020 | Asia / India / Gujarat / Jamnagar | Gujarat | 2020-05-10 | M.P Shah Government Medocal college Jamnagar |
| 326 | 451154 | EPI_ISL_451154 | hCoV-19/India/GBRC109/2020 | Asia / India / Gujarat / Dahod | Gujarat | 2020-05-03 | Government Medical College, Vadodara |
| 327 | 451155 | EPI_ISL_451155 | hCoV-19/India/GBRC110/2020 | Asia / India / Gujarat / Vadodara | Gujarat | 2020-05-03 | Government Medical College, Vadodara |
| 328 | 451156 | EPI_ISL_451156 | hCoV-19/India/GBRC111/2020 | Asia / India / Gujarat / Dahod | Gujarat | 2020-05-01 | Government Medical College, Vadodara |
| 329 | 451157 | EPI_ISL_451157 | hCoV-19/India/GBRC112/2020 | Asia / India / Gujarat / Vadodara | Gujarat | 2020-05-02 | Government Medical College, Vadodara |
| 330 | 451158 | EPI_ISL_451158 | hCoV-19/India/GBRC113/2020 | Asia / India / Gujarat / Vadodara | Gujarat | 2020-05-03 | Government Medical College, Vadodara |
| 331 | 451159 | EPI_ISL_451159 | hCoV-19/India/GBRC114/2020 | Asia / India / Gujarat / Dahod | Gujarat | 2020-05-03 | Government Medical College, Vadodara |
| 332 | 451160 | EPI_ISL_451160 | hCoV-19/India/GBRC115/2020 | Asia / India / Gujarat / Vadodara | Gujarat | 2020-05-03 | Government Medical College, Vadodara |
| 333 | 451161 | EPI_ISL_451161 | hCoV-19/India/GBRC116/2020 | Asia / India / Gujarat / Dahod | Gujarat | 2020-05-03 | Government Medical College, Vadodara |
| 334 | 451162 | EPI_ISL_451162 | hCoV-19/India/GBRC117/2020 | Asia / India / Gujarat / Vadodara | Gujarat | 2020-05-03 | Government Medical College, Vadodara |
| 335 | 451163 | EPI_ISL_451163 | hCoV-19/India/GBRC118/2020 | Asia / India / Gujarat / Vadodara | Gujarat | 2020-05-02 | Government Medical College, Vadodara |
| 336 | 451666 | EPI_ISL_451666 | hCoV-19/India/GBRC103/2020 | Asia / India / Gujarat / Jamnagar | Gujarat | 2020-05-04 | M.P Shah Government Medocal college Jamnagar |
| 337 | 452192 | EPI_ISL_452192 | hCoV-19/India/NIV-10380/2020 | Asia / India / Maharashtra | Maharashtra | 2020-04-16 | NIV Influenza |
| 338 | 452193 | EPI_ISL_452193 | hCoV-19/India/NIV-10419/2020 | Asia / India / Maharashtra | Maharashtra | 2020-04-16 | NIV Influenza |
| 339 | 452195 | EPI_ISL_452195 | hCoV-19/India/NIV-11681/2020 | Asia / India / Maharashtra | Maharashtra | 2020-04-19 | NIV Influenza |
| 340 | 452196 | EPI_ISL_452196 | hCoV-19/India/NIV-11683/2020 | Asia / India / Maharashtra | Maharashtra | 2020-04-19 | NIV Influenza |
| 341 | 452197 | EPI_ISL_452197 | hCoV-19/India/NIV-11687/2020 | Asia / India / Maharashtra | Maharashtra | 2020-04-19 | NIV Influenza |
| 342 | 452198 | EPI_ISL_452198 | hCoV-19/India/NIV-11855/2020 | Asia / India / Maharashtra | Maharashtra | 2020-04-20 | NIV Influenza |
| 343 | 452199 | EPI_ISL_452199 | hCoV-19/India/NIV-12019/2020 | Asia / India / Maharashtra | Maharashtra | 2020-04-20 | NIV Influenza |
| 344 | 452200 | EPI_ISL_452200 | hCoV-19/India/NIV-12024/2020 | Asia / India / Maharashtra | Maharashtra | 2020-04-20 | NIV Influenza |
| 345 | 452201 | EPI_ISL_452201 | hCoV-19/India/NIV-12067/2020 | Asia / India / Maharashtra | Maharashtra | 2020-04-20 | NIV Influenza |
| 346 | 452202 | EPI_ISL_452202 | hCoV-19/India/NIV-2810/2020 | Asia / India / Maharashtra | Maharashtra | 2020-03-16 | NIV Influenza |
| 347 | 452203 | EPI_ISL_452203 | hCoV-19/India/NIV-4367/2020 | Asia / India / Maharashtra | Maharashtra | 2020-03-22 | NIV Influenza |
| 348 | 452204 | EPI_ISL_452204 | hCoV-19/India/NIV-5109/2020 | Asia / India / Maharashtra | Maharashtra | 2020-03-26 | NIV Influenza |
| 349 | 452205 | EPI_ISL_452205 | hCoV-19/India/NIV-5119/2020 | Asia / India / Maharashtra | Maharashtra | 2020-03-26 | NIV Influenza |
| 350 | 452207 | EPI_ISL_452207 | hCoV-19/India/NIV-7439/2020 | Asia / India / Maharashtra | Maharashtra | 2020-04-05 | NIV Influenza |
| 351 | 452208 | EPI_ISL_452208 | hCoV-19/India/NIV-7511/2020 | Asia / India / Maharashtra | Maharashtra | 2020-04-05 | NIV Influenza |
| 352 | 452209 | EPI_ISL_452209 | hCoV-19/India/NIV-7786/2020 | Asia / India / Maharashtra | Maharashtra | 2020-04-06 | NIV Influenza |
| 353 | 452210 | EPI_ISL_452210 | hCoV-19/India/NIV-7830/2020 | Asia / India / Maharashtra | Maharashtra | 2020-04-06 | NIV Influenza |
| 354 | 452211 | EPI_ISL_452211 | hCoV-19/India/NIV-9157/2020 | Asia / India / Maharashtra | Maharashtra | 2020-04-13 | NIV Influenza |
| 355 | 452212 | EPI_ISL_452212 | hCoV-19/India/NIV-9768/2020 | Asia / India / Maharashtra | Maharashtra | 2020-04-15 | NIV Influenza |
| 356 | 452213 | EPI_ISL_452213 | hCoV-19/India/NIV-981/2020 | Asia / India / Maharashtra | Maharashtra | 2020-03-10 | NIV Influenza |
| 357 | 452214 | EPI_ISL_452214 | hCoV-19/India/NIV-QA-709/2020 | Asia / India / Maharashtra | Maharashtra | 2020-04-07 | NIV Influenza |
| 358 | 452215 | EPI_ISL_452215 | hCoV-19/India/NIV-QC-797/2020 | Asia / India / Maharashtra | Maharashtra | 2020-04-22 | NIV Influenza |
| 359 | 452216 | EPI_ISL_452216 | hCoV-19/India/NIV-QC-802/2020 | Asia / India / Maharashtra | Maharashtra | 2020-04-27 | NIV Influenza |
| 360 | 452217 | EPI_ISL_452217 | hCoV-19/India/NIV-QC-800/2020 | Asia / India / Maharashtra | Maharashtra | 2020-04-26 | NIV Influenza |
| 361 | 452790 | EPI_ISL_452790 | hCoV-19/India/NIHSAD-2604-22/2020 | Asia / India / Madhya Pradesh | Madhya Pradesh | 2020-04-24 | ICAR-National Institute of High Security Animal Diseases |
| 362 | 452791 | EPI_ISL_452791 | hCoV-19/India/NIHSAD-2904-107/2020 | Asia / India / Madhya Pradesh | Madhya Pradesh | 2020-04-27 | ICAR-National Institute of High Security Animal Diseases |
| 363 | 452792 | EPI_ISL_452792 | hCoV-19/India/NIHSAD-0105-77/2020 | Asia / India / Madhya Pradesh | Madhya Pradesh | 2020-04-30 | ICAR-National Institute of High Security Animal Diseases |
| 364 | 452793 | EPI_ISL_452793 | hCoV-19/India/NIHSAD-0405-24/2020 | Asia / India / Madhya Pradesh | Madhya Pradesh | 2020-05-03 | ICAR-National Institute of High Security Animal Diseases |
| 365 | 452794 | EPI_ISL_452794 | hCoV-19/India/NIHSAD-0905-216/2020 | Asia / India / Madhya Pradesh | Madhya Pradesh | 2020-05-08 | ICAR-National Institute of High Security Animal Diseases |
| 366 | 452795 | EPI_ISL_452795 | hCoV-19/India/NIHSAD-1005-52/2020 | Asia / India / Madhya Pradesh | Madhya Pradesh | 2020-05-09 | ICAR-National Institute of High Security Animal Diseases |
| 367 | 454524 | EPI_ISL_454524 | hCoV-19/India/NIV-12058/2020 | Asia / India / Maharashtra | Maharashtra | 2020-04-20 | NIV Influenza |
| 368 | 454525 | EPI_ISL_454525 | hCoV-19/India/NIV-1738/2020 | Asia / India / Maharashtra | Maharashtra | 2020-03-12 | NIV Influenza |
| 369 | 454526 | EPI_ISL_454526 | hCoV-19/India/NIV-1804/2020 | Asia / India / Maharashtra | Maharashtra | 2020-03-13 | NIV Influenza |
| 370 | 454527 | EPI_ISL_454527 | hCoV-19/India/NIV-2387/2020 | Asia / India / Maharashtra | Maharashtra | 2020-03-13 | NIV Influenza |
| 371 | 454528 | EPI_ISL_454528 | hCoV-19/India/NIV-3472/2020 | Asia / India / Maharashtra | Maharashtra | 2020-03-17 | NIV Influenza |
| 372 | 454529 | EPI_ISL_454529 | hCoV-19/India/NIV-4130/2020 | Asia / India / Maharashtra | Maharashtra | 2020-03-21 | NIV Influenza |
| 373 | 454531 | EPI_ISL_454531 | hCoV-19/India/NIV-4369/2020 | Asia / India / Maharashtra | Maharashtra | 2020-03-22 | NIV Influenza |
| 374 | 454532 | EPI_ISL_454532 | hCoV-19/India/NIV-4379/2020 | Asia / India / Maharashtra | Maharashtra | 2020-03-23 | NIV Influenza |
| 375 | 454533 | EPI_ISL_454533 | hCoV-19/India/NIV-4438/2020 | Asia / India / Maharashtra | Maharashtra | 2020-03-23 | NIV Influenza |
| 376 | 454534 | EPI_ISL_454534 | hCoV-19/India/NIV-4622/2020 | Asia / India / Maharashtra | Maharashtra | 2020-03-24 | NIV Influenza |
| 377 | 454536 | EPI_ISL_454536 | hCoV-19/India/NIV-5102/2020 | Asia / India / Maharashtra | Maharashtra | 2020-03-26 | NIV Influenza |
| 378 | 454537 | EPI_ISL_454537 | hCoV-19/India/NIV-5104/2020 | Asia / India / Maharashtra | Maharashtra | 2020-03-26 | NIV Influenza |
| 379 | 454540 | EPI_ISL_454540 | hCoV-19/India/NIV-6216/2020 | Asia / India / Maharashtra | Maharashtra | 2020-04-01 | NIV Influenza |
| 380 | 454542 | EPI_ISL_454542 | hCoV-19/India/NIV-6315/2020 | Asia / India / Maharashtra | Maharashtra | 2020-03-30 | NIV Influenza |
| 381 | 454543 | EPI_ISL_454543 | hCoV-19/India/NIV-6423/2020 | Asia / India / Maharashtra | Maharashtra | 2020-03-31 | NIV Influenza |
| 382 | 454544 | EPI_ISL_454544 | hCoV-19/India/NIV-6614/2020 | Asia / India / Maharashtra | Maharashtra | 2020-04-02 | NIV Influenza |
| 383 | 454546 | EPI_ISL_454546 | hCoV-19/India/NIV-6964/2020 | Asia / India / Maharashtra | Maharashtra | 2020-04-03 | NIV Influenza |

| | | | | | | | |
|---|---|---|---|---|---|---|---|
| 384 | 454547 | EPI_ISL_454547 | hCoV-19/India/NIV-7044/2020 | Asia / India / Maharashtra | Maharashtra | 2020-04-04 | NIV Influenza |
| 385 | 454549 | EPI_ISL_454549 | hCoV-19/India/NIV-7558/2020 | Asia / India / Maharashtra | Maharashtra | 2020-04-05 | NIV Influenza |
| 386 | 454551 | EPI_ISL_454551 | hCoV-19/India/NIV-7893/2020 | Asia / India / Maharashtra | Maharashtra | 2020-04-07 | NIV Influenza |
| 387 | 454552 | EPI_ISL_454552 | hCoV-19/India/NIV-7966/2020 | Asia / India / Maharashtra | Maharashtra | 2020-04-06 | NIV Influenza |
| 388 | 454557 | EPI_ISL_454557 | hCoV-19/India/NIV-9154/2020 | Asia / India / Maharashtra | Maharashtra | 2020-04-13 | NIV Influenza |
| 389 | 454560 | EPI_ISL_454560 | hCoV-19/India/NIV-701_3/2020 | Asia / India / Maharashtra | Maharashtra | 2020-04-06 | NIV Influenza |
| 390 | 454563 | EPI_ISL_454563 | hCoV-19/India/NIV-QA-707/2020 | Asia / India / Maharashtra | Maharashtra | 2020-04-10 | NIV Influenza |
| 391 | 454564 | EPI_ISL_454564 | hCoV-19/India/NIV-QA-708/2020 | Asia / India / Maharashtra | Maharashtra | 2020-04-04 | NIV Influenza |
| 392 | 454565 | EPI_ISL_454565 | hCoV-19/India/NIV-QA-710/2020 | Asia / India / Maharashtra | Maharashtra | 2020-03-31 | NIV Influenza |
| 393 | 454566 | EPI_ISL_454566 | hCoV-19/India/NIV-QC-795/2020 | Asia / India / Maharashtra | Maharashtra | 2020-04-21 | NIV Influenza |
| 394 | 454567 | EPI_ISL_454567 | hCoV-19/India/NIV-QC-796/2020 | Asia / India / Maharashtra | Maharashtra | 2020-04-22 | NIV Influenza |
| 395 | 454568 | EPI_ISL_454568 | hCoV-19/India/NIV-QC-798/2020 | Asia / India / Maharashtra | Maharashtra | 2020-04-25 | NIV Influenza |
| 396 | 454569 | EPI_ISL_454569 | hCoV-19/India/NIV-QC-799/2020 | Asia / India / Maharashtra | Maharashtra | 2020-04-26 | NIV Influenza |
| 397 | 454570 | EPI_ISL_454570 | hCoV-19/India/NIV-QC-801/2020 | Asia / India / Maharashtra | Maharashtra | 2020-04-26 | NIV Influenza |
| 398 | 454830 | EPI_ISL_454830 | hCoV-19/India/SMSCOV109/2020 | Asia / India / Rajasthan / Jaipur | Rajasthan | 2020-04-23 | SMS Medical College, Jaipur |
| 399 | 454831 | EPI_ISL_454831 | hCoV-19/India/SMSCOV141/2020 | Asia / India / Rajasthan / Jaipur | Rajasthan | 2020-04-29 | SMS Medical College, Jaipur |
| 400 | 454832 | EPI_ISL_454832 | hCoV-19/India/SMSCOV161/2020 | Asia / India / Rajasthan / Jaipur | Rajasthan | 2020-04-21 | SMS Medical College, Jaipur |
| 401 | 454833 | EPI_ISL_454833 | hCoV-19/India/SMSCOV175/2020 | Asia / India / Rajasthan / Jaipur | Rajasthan | 2020-04-27 | SMS Medical College, Jaipur |
| 402 | 454858 | EPI_ISL_454858 | hCoV-19/India/THSTI-BAL_39/2020 | Asia / India / Haryana | Haryana | 2020-04-07 | Translational Health Science and Technology Institute -ESIC medical college and hospital, Faridabad |
| 403 | 454862 | EPI_ISL_454862 | hCoV-19/India/THSTI-BAL_231/2020 | Asia / India / Haryana | Haryana | 2020-04-11 | Translational Health Science and Technology Institute -ESIC medical college and hospital, Faridabad |
| 404 | 454866 | EPI_ISL_454866 | hCoV-19/India/THSTI-BAL_914/2020 | Asia / India / Haryana | Haryana | 2020-04-16 | Translational Health Science and Technology Institute -ESIC medical college and hospital, Faridabad |
| 405 | 454867 | EPI_ISL_454867 | hCoV-19/India/THSTI-BAL_918/2020 | Asia / India / Haryana | Haryana | 2020-04-16 | Translational Health Science and Technology Institute -ESIC medical college and hospital, Faridabad |
| 406 | 455015 | EPI_ISL_455015 | hCoV-19/India/GBRC80/2020 | Asia / India / Gujarat / Rajkot | Gujarat | 2020-04-28 | Pandit Deendayal Upadhyay Government Medical College, Rajkot |
| 407 | 455016 | EPI_ISL_455016 | hCoV-19/India/GBRC119/2020 | Asia / India / Gujarat / Rajkot | Gujarat | 2020-04-27 | Pandit Deendayal Upadhyay Government Medical College, Rajkot |
| 408 | 455017 | EPI_ISL_455017 | hCoV-19/India/GBRC120/2020 | Asia / India / Gujarat / Vadodara | Gujarat | 2020-05-02 | Government Medical College, Vadodara |
| 409 | 455018 | EPI_ISL_455018 | hCoV-19/India/GBRC121/2020 | Asia / India / Gujarat / Vadodara | Gujarat | 2020-05-02 | Government Medical College, Vadodara |
| 410 | 455019 | EPI_ISL_455019 | hCoV-19/India/GBRC122/2020 | Asia / India / Gujarat / Vadodara | Gujarat | 2020-05-02 | Government Medical College, Vadodara |
| 411 | 455020 | EPI_ISL_455020 | hCoV-19/India/GBRC123/2020 | Asia / India / Gujarat / Vadodara | Gujarat | 2020-05-02 | Government Medical College, Vadodara |
| 412 | 455021 | EPI_ISL_455021 | hCoV-19/India/GBRC124/2020 | Asia / India / Gujarat / Vadodara | Gujarat | 2020-05-02 | Government Medical College, Vadodara |
| 413 | 455022 | EPI_ISL_455022 | hCoV-19/India/GBRC125/2020 | Asia / India / Gujarat / Vadodara | Gujarat | 2020-05-02 | Government Medical College, Vadodara |
| 414 | 455023 | EPI_ISL_455023 | hCoV-19/India/GBRC126/2020 | Asia / India / Gujarat / Vadodara | Gujarat | 2020-05-02 | Government Medical College, Vadodara |
| 415 | 455024 | EPI_ISL_455024 | hCoV-19/India/GBRC127/2020 | Asia / India / Gujarat / Vadodara | Gujarat | 2020-05-02 | Government Medical College, Vadodara |
| 416 | 455025 | EPI_ISL_455025 | hCoV-19/India/GBRC128/2020 | Asia / India / Gujarat / Vadodara | Gujarat | 2020-05-02 | Government Medical College, Vadodara |
| 417 | 455026 | EPI_ISL_455026 | hCoV-19/India/GBRC129/2020 | Asia / India / Gujarat / Vadodara | Gujarat | 2020-05-02 | Government Medical College, Vadodara |
| 418 | 455027 | EPI_ISL_455027 | hCoV-19/India/GBRC130/2020 | Asia / India / Gujarat / Vadodara | Gujarat | 2020-05-02 | Government Medical College, Vadodara |
| 419 | 455478 | EPI_ISL_455478 | hCoV-19/India/RMRC49/2020 | Asia / India / Odisha | Odisha | 2020-04-09 | REGIONAL VRDL,ICMR-RMRC BBSR |
| 420 | 455640 | EPI_ISL_455640 | hCoV-19/India/S4/2020 | Asia / India / West_Bengal / East Medinipur | West Bengal | 2020-03-27 | ICMR-National Institute of Cholera and Enteric Diseases |
| 421 | 455643 | EPI_ISL_455643 | hCoV-19/India/S13/2020 | Asia / India / West_Bengal / North 24 Parganas | West Bengal | 2020-03-30 | ICMR-National Institute of Cholera and Enteric Diseases |
| 422 | 455644 | EPI_ISL_455644 | hCoV-19/India/S14/2020 | Asia / India / West_Bengal / East Medinipur | West Bengal | 2020-04-02 | ICMR-National Institute of Cholera and Enteric Diseases |
| 423 | 455645 | EPI_ISL_455645 | hCoV-19/India/S15/2020 | Asia / India / West_Bengal / Kolkata | West Bengal | 2020-04-03 | ICMR-National Institute of Cholera and Enteric Diseases |
| 424 | 455646 | EPI_ISL_455646 | hCoV-19/India/S16/2020 | Asia / India / West_Bengal / Howrah | West Bengal | 2020-04-03 | ICMR-National Institute of Cholera and Enteric Diseases |
| 425 | 455647 | EPI_ISL_455647 | hCoV-19/India/S17/2020 | Asia / India / West_Bengal / Kolkata | West Bengal | 2020-04-19 | ICMR-National Institute of Cholera and Enteric Diseases |
| 426 | 455648 | EPI_ISL_455648 | hCoV-19/India/S18/2020 | Asia / India / West_Bengal / Kolkata | West Bengal | 2020-04-20 | ICMR-National Institute of Cholera and Enteric Diseases |
| 427 | 455649 | EPI_ISL_455649 | hCoV-19/India/S19/2020 | Asia / India / West_Bengal / Kolkata | West Bengal | 2020-04-20 | ICMR-National Institute of Cholera and Enteric Diseases |
| 428 | 455650 | EPI_ISL_455650 | hCoV-19/India/S20/2020 | Asia / India / West_Bengal / Kolkata | West Bengal | 2020-04-20 | ICMR-National Institute of Cholera and Enteric Diseases |
| 429 | 455651 | EPI_ISL_455651 | hCoV-19/India/S21/2020 | Asia / India / West_Bengal / Kolkata | West Bengal | 2020-04-21 | ICMR-National Institute of Cholera and Enteric Diseases |
| 430 | 455652 | EPI_ISL_455652 | hCoV-19/India/S22/2020 | Asia / India / West_Bengal / South 24 Parganas | West Bengal | 2020-04-21 | ICMR-National Institute of Cholera and Enteric Diseases |
| 431 | 455653 | EPI_ISL_455653 | hCoV-19/India/S23/2020 | Asia / India / West_Bengal / South 24 Parganas | West Bengal | 2020-04-21 | ICMR-National Institute of Cholera and Enteric Diseases |
| 432 | 455655 | EPI_ISL_455655 | hCoV-19/India/S25/2020 | Asia / India / Rajasthan / Bikaner | Rajasthan | 2020-04-30 | ICMR-National Institute of Cholera and Enteric Diseases |
| 433 | 455656 | EPI_ISL_455656 | hCoV-19/India/S27/2020 | Asia / India / West_Bengal / South 24 Parganas | West Bengal | 2020-04-30 | ICMR-National Institute of Cholera and Enteric Diseases |
| 434 | 455658 | EPI_ISL_455658 | hCoV-19/India/S29/2020 | Asia / India / West_Bengal / South 24 Parganas | West Bengal | 2020-04-30 | ICMR-National Institute of Cholera and Enteric Diseases |
| 435 | 455659 | EPI_ISL_455659 | hCoV-19/India/S30/2020 | Asia / India / West_Bengal / South 24 Parganas | West Bengal | 2020-05-01 | ICMR-National Institute of Cholera and Enteric Diseases |
| 436 | 455660 | EPI_ISL_455660 | hCoV-19/India/S31/2020 | Asia / India / West_Bengal / South 24 Parganas | West Bengal | 2020-05-01 | ICMR-National Institute of Cholera and Enteric Diseases |
| 437 | 455662 | EPI_ISL_455662 | hCoV-19/India/S33/2020 | Asia / India / West_Bengal / Howrah | West Bengal | 2020-05-02 | ICMR-National Institute of Cholera and Enteric Diseases |
| 438 | 455663 | EPI_ISL_455663 | hCoV-19/India/S34/2020 | Asia / India / West_Bengal / South 24 Parganas | West Bengal | 2020-05-02 | ICMR-National Institute of Cholera and Enteric Diseases |
| 439 | 455667 | EPI_ISL_455667 | hCoV-19/India/S46/2020 | Asia / India / West_Bengal / South 24 Parganas | West Bengal | 2020-05-03 | ICMR-National Institute of Cholera and Enteric Diseases |
| 440 | 455668 | EPI_ISL_455668 | hCoV-19/India/S48/2020 | Asia / India / West_Bengal / North 24 Parganas | West Bengal | 2020-05-05 | ICMR-National Institute of Cholera and Enteric Diseases |
| 441 | 455669 | EPI_ISL_455669 | hCoV-19/India/S49/2020 | Asia / India / West_Bengal / Kolkata | West Bengal | 2020-05-03 | ICMR-National Institute of Cholera and Enteric Diseases |
| 442 | 455670 | EPI_ISL_455670 | hCoV-19/India/S50/2020 | Asia / India / West_Bengal / Kolkata | West Bengal | 2020-05-03 | ICMR-National Institute of Cholera and Enteric Diseases |
| 443 | 455671 | EPI_ISL_455671 | hCoV-19/India/S51/2020 | Asia / India / West_Bengal / Kolkata | West Bengal | 2020-05-03 | ICMR-National Institute of Cholera and Enteric Diseases |
| 444 | 455672 | EPI_ISL_455672 | hCoV-19/India/S52/2020 | Asia / India / West_Bengal / North 24 Parganas | West Bengal | 2020-05-03 | ICMR-National Institute of Cholera and Enteric Diseases |
| 445 | 455673 | EPI_ISL_455673 | hCoV-19/India/S53/2020 | Asia / India / West_Bengal / South 24 Parganas | West Bengal | 2020-05-02 | ICMR-National Institute of Cholera and Enteric Diseases |
| 446 | 455674 | EPI_ISL_455674 | hCoV-19/India/S55/2020 | Asia / India / West_Bengal / Kolkata | West Bengal | 2020-05-03 | ICMR-National Institute of Cholera and Enteric Diseases |
| 447 | 455675 | EPI_ISL_455675 | hCoV-19/India/S56/2020 | Asia / India / West_Bengal / North 24 Parganas | West Bengal | 2020-05-03 | ICMR-National Institute of Cholera and Enteric Diseases |
| 448 | 455676 | EPI_ISL_455676 | hCoV-19/India/S58/2020 | Asia / India / West_Bengal / Kolkata | West Bengal | 2020-05-03 | ICMR-National Institute of Cholera and Enteric Diseases |
| 449 | 455678 | EPI_ISL_455678 | hCoV-19/India/S61/2020 | Asia / India / West_Bengal / North 24 Parganas | West Bengal | 2020-04-30 | ICMR-National Institute of Cholera and Enteric Diseases |
| 450 | 455679 | EPI_ISL_455679 | hCoV-19/India/S62/2020 | Asia / India / West_Bengal / East Medinipur | West Bengal | 2020-05-04 | ICMR-National Institute of Cholera and Enteric Diseases |
| 451 | 455749 | EPI_ISL_455749 | hCoV-19/India/RMRC103/2020 | Asia / India / Odisha | Odisha | 2020-05-06 | REGIONAL VRDL,ICMR-RMRC BBSR |
| 452 | 455751 | EPI_ISL_455751 | hCoV-19/India/RMRC106/2020 | Asia / India / Odisha | Odisha | 2020-05-06 | REGIONAL VRDL,ICMR-RMRC BBSR |
| 453 | 455752 | EPI_ISL_455752 | hCoV-19/India/RMRC108/2020 | Asia / India / Odisha | Odisha | 2020-05-06 | REGIONAL VRDL,ICMR-RMRC BBSR |
| 454 | 455754 | EPI_ISL_455754 | hCoV-19/India/RMRC112/2020 | Asia / India / Odisha | Odisha | 2020-05-06 | REGIONAL VRDL,ICMR-RMRC BBSR |
| 455 | 455755 | EPI_ISL_455755 | hCoV-19/India/RMRC154/2020 | Asia / India / Odisha | Odisha | 2020-05-07 | REGIONAL VRDL,ICMR-RMRC BBSR |
| 456 | 455757 | EPI_ISL_455757 | hCoV-19/India/RMRC156/2020 | Asia / India / Odisha | Odisha | 2020-05-07 | REGIONAL VRDL,ICMR-RMRC BBSR |
| 457 | 455758 | EPI_ISL_455758 | hCoV-19/India/RMRC157/2020 | Asia / India / Odisha | Odisha | 2020-05-07 | REGIONAL VRDL,ICMR-RMRC BBSR |
| 458 | 455760 | EPI_ISL_455760 | hCoV-19/India/RMRC159/2020 | Asia / India / Odisha | Odisha | 2020-05-07 | REGIONAL VRDL,ICMR-RMRC BBSR |
| 459 | 455761 | EPI_ISL_455761 | hCoV-19/India/RMRC160/2020 | Asia / India / Odisha | Odisha | 2020-05-07 | REGIONAL VRDL,ICMR-RMRC BBSR |
| 460 | 455763 | EPI_ISL_455763 | hCoV-19/India/RMRC163/2020 | Asia / India / Odisha | Odisha | 2020-05-07 | REGIONAL VRDL,ICMR-RMRC BBSR |

| | | | | | | | |
|---|---|---|---|---|---|---|---|
| 461 | 455764 | EPI_ISL_455764 | hCoV-19/India/RMRC164/2020 | Asia / India / Odisha | Odisha | 2020-05-07 | REGIONAL VRDL,ICMR-RMRC BBSR |
| 462 | 455765 | EPI_ISL_455765 | hCoV-19/India/RMRC165/2020 | Asia / India / Odisha | Odisha | 2020-05-07 | REGIONAL VRDL,ICMR-RMRC BBSR |
| 463 | 455766 | EPI_ISL_455766 | hCoV-19/India/RMRC166/2020 | Asia / India / Odisha | Odisha | 2020-05-07 | REGIONAL VRDL,ICMR-RMRC BBSR |
| 464 | 455767 | EPI_ISL_455767 | hCoV-19/India/RMRC167/2020 | Asia / India / Odisha | Odisha | 2020-05-07 | REGIONAL VRDL,ICMR-RMRC BBSR |
| 465 | 455768 | EPI_ISL_455768 | hCoV-19/India/RMRC169/2020 | Asia / India / Odisha | Odisha | 2020-05-07 | REGIONAL VRDL,ICMR-RMRC BBSR |
| 466 | 455770 | EPI_ISL_455770 | hCoV-19/India/RMRC171/2020 | Asia / India / Odisha | Odisha | 2020-05-07 | REGIONAL VRDL,ICMR-RMRC BBSR |
| 467 | 455771 | EPI_ISL_455771 | hCoV-19/India/RMRC2/2020 | Asia / India / Odisha | Odisha | 2020-03-19 | REGIONAL VRDL,ICMR-RMRC BBSR |
| 468 | 455775 | EPI_ISL_455775 | hCoV-19/India/RMRC24/2020 | Asia / India / Odisha | Odisha | 2020-04-04 | REGIONAL VRDL,ICMR-RMRC BBSR |
| 469 | 455776 | EPI_ISL_455776 | hCoV-19/India/RMRC27/2020 | Asia / India / Odisha | Odisha | 2020-04-04 | REGIONAL VRDL,ICMR-RMRC BBSR |
| 470 | 455777 | EPI_ISL_455777 | hCoV-19/India/RMRC28/2020 | Asia / India / Odisha | Odisha | 2020-04-04 | REGIONAL VRDL,ICMR-RMRC BBSR |
| 471 | 455778 | EPI_ISL_455778 | hCoV-19/India/RMRC30/2020 | Asia / India / Odisha | Odisha | 2020-04-04 | REGIONAL VRDL,ICMR-RMRC BBSR |
| 472 | 455779 | EPI_ISL_455779 | hCoV-19/India/RMRC38/2020 | Asia / India / Odisha | Odisha | 2020-04-04 | REGIONAL VRDL,ICMR-RMRC BBSR |
| 473 | 455780 | EPI_ISL_455780 | hCoV-19/India/RMRC46/2020 | Asia / India / Odisha | Odisha | 2020-04-08 | REGIONAL VRDL,ICMR-RMRC BBSR |
| 474 | 455782 | EPI_ISL_455782 | hCoV-19/India/RMRC5/2020 | Asia / India / Odisha | Odisha | 2020-04-02 | REGIONAL VRDL,ICMR-RMRC BBSR |
| 475 | 455783 | EPI_ISL_455783 | hCoV-19/India/RMRC6/2020 | Asia / India / Odisha | Odisha | 2020-04-02 | REGIONAL VRDL,ICMR-RMRC BBSR |
| 476 | 455784 | EPI_ISL_455784 | hCoV-19/India/RMRC7/2020 | Asia / India / Odisha | Odisha | 2020-04-02 | REGIONAL VRDL,ICMR-RMRC BBSR |
| 477 | 455786 | EPI_ISL_455786 | hCoV-19/India/RMRC90/2020 | Asia / India / Odisha | Odisha | 2020-05-04 | REGIONAL VRDL,ICMR-RMRC BBSR |
| 478 | 455787 | EPI_ISL_455787 | hCoV-19/India/RMRC92/2020 | Asia / India / Odisha | Odisha | 2020-05-04 | REGIONAL VRDL,ICMR-RMRC BBSR |
| 479 | 458030 | EPI_ISL_458030 | hCoV-19/India/CCMB_C12/2020 | Asia / India / Tamil Nadu | Tamil Nadu | 2020-04-26 | King Institute of Preventive Medicine & Research |
| 480 | 458031 | EPI_ISL_458031 | hCoV-19/India/CCMB_C6/2020 | Asia / India / Tamil Nadu | Tamil Nadu | 2020-04-26 | King Institute of Preventive Medicine & Research |
| 481 | 458032 | EPI_ISL_458032 | hCoV-19/India/CCMB_C8/2020 | Asia / India / Tamil Nadu | Tamil Nadu | 2020-04-27 | King Institute of Preventive Medicine & Research |
| 482 | 458033 | EPI_ISL_458033 | hCoV-19/India/CCMB_C5/2020 | Asia / India / Tamil Nadu | Tamil Nadu | 2020-04-29 | King Institute of Preventive Medicine & Research |
| 483 | 458034 | EPI_ISL_458034 | hCoV-19/India/CCMB_C11/2020 | Asia / India / Tamil Nadu | Tamil Nadu | 2020-04-29 | King Institute of Preventive Medicine & Research |
| 484 | 458035 | EPI_ISL_458035 | hCoV-19/India/CCMB_C13/2020 | Asia / India / Tamil Nadu | Tamil Nadu | 2020-04-29 | King Institute of Preventive Medicine & Research |
| 485 | 458036 | EPI_ISL_458036 | hCoV-19/India/CCMB_C10/2020 | Asia / India / Tamil Nadu | Tamil Nadu | 2020-04-29 | King Institute of Preventive Medicine & Research |
| 486 | 458037 | EPI_ISL_458037 | hCoV-19/India/CCMB_C7/2020 | Asia / India / Tamil Nadu | Tamil Nadu | 2020-04-29 | King Institute of Preventive Medicine & Research |
| 487 | 458038 | EPI_ISL_458038 | hCoV-19/India/CCMB_C9/2020 | Asia / India / Tamil Nadu | Tamil Nadu | 2020-04-29 | King Institute of Preventive Medicine & Research |
| 488 | 458039 | EPI_ISL_458039 | hCoV-19/India/CCMB_C16/2020 | Asia / India / Tamil Nadu | Tamil Nadu | 2020-05-06 | King Institute of Preventive Medicine & Research |
| 489 | 458040 | EPI_ISL_458040 | hCoV-19/India/CCMB_C19/2020 | Asia / India / Tamil Nadu | Tamil Nadu | 2020-05-06 | King Institute of Preventive Medicine & Research |
| 490 | 458041 | EPI_ISL_458041 | hCoV-19/India/CCMB_C15/2020 | Asia / India / Tamil Nadu | Tamil Nadu | 2020-05-06 | King Institute of Preventive Medicine & Research |
| 491 | 458042 | EPI_ISL_458042 | hCoV-19/India/CCMB_C18/2020 | Asia / India / Tamil Nadu | Tamil Nadu | 2020-05-06 | King Institute of Preventive Medicine & Research |
| 492 | 458043 | EPI_ISL_458043 | hCoV-19/India/CCMB_C14/2020 | Asia / India / Tamil Nadu | Tamil Nadu | 2020-05-06 | King Institute of Preventive Medicine & Research |
| 493 | 458044 | EPI_ISL_458044 | hCoV-19/India/CCMB_C17/2020 | Asia / India / Tamil Nadu | Tamil Nadu | 2020-05-06 | King Institute of Preventive Medicine & Research |
| 494 | 458045 | EPI_ISL_458045 | hCoV-19/India/CCMB_L1017/2020 | Asia / India / Telangana | Telangana | 2020-05-11 | CSIR-Centre for Cellular and Molecular Biology |
| 495 | 458046 | EPI_ISL_458046 | hCoV-19/India/CCMB_L1021/2020 | Asia / India / Telangana | Telangana | 2020-05-11 | CSIR-Centre for Cellular and Molecular Biology |
| 496 | 458047 | EPI_ISL_458047 | hCoV-19/India/CCMB_L1022/2020 | Asia / India / Telangana | Telangana | 2020-05-11 | CSIR-Centre for Cellular and Molecular Biology |
| 497 | 458048 | EPI_ISL_458048 | hCoV-19/India/CCMB_L1024/2020 | Asia / India / Telangana | Telangana | 2020-05-11 | CSIR-Centre for Cellular and Molecular Biology |
| 498 | 458049 | EPI_ISL_458049 | hCoV-19/India/CCMB_L1025/2020 | Asia / India / Telangana | Telangana | 2020-05-11 | CSIR-Centre for Cellular and Molecular Biology |
| 499 | 458050 | EPI_ISL_458050 | hCoV-19/India/CCMB_L1026/2020 | Asia / India / Telangana | Telangana | 2020-05-11 | CSIR-Centre for Cellular and Molecular Biology |
| 500 | 458051 | EPI_ISL_458051 | hCoV-19/India/CCMB_L1062/2020 | Asia / India / Telangana | Telangana | 2020-05-12 | CSIR-Centre for Cellular and Molecular Biology |
| 501 | 458052 | EPI_ISL_458052 | hCoV-19/India/CCMB_L1063/2020 | Asia / India / Telangana | Telangana | 2020-05-12 | CSIR-Centre for Cellular and Molecular Biology |
| 502 | 458053 | EPI_ISL_458053 | hCoV-19/India/CCMB_L1064/2020 | Asia / India / Telangana | Telangana | 2020-05-12 | CSIR-Centre for Cellular and Molecular Biology |
| 503 | 458054 | EPI_ISL_458054 | hCoV-19/India/CCMB_L1073/2020 | Asia / India / Telangana | Telangana | 2020-05-13 | CSIR-Centre for Cellular and Molecular Biology |
| 504 | 458055 | EPI_ISL_458055 | hCoV-19/India/CCMB_L1082/2020 | Asia / India / Telangana | Telangana | 2020-05-13 | CSIR-Centre for Cellular and Molecular Biology |
| 505 | 458056 | EPI_ISL_458056 | hCoV-19/India/CCMB_L1085/2020 | Asia / India / Telangana | Telangana | 2020-05-13 | CSIR-Centre for Cellular and Molecular Biology |
| 506 | 458057 | EPI_ISL_458057 | hCoV-19/India/CCMB_L1089/2020 | Asia / India / Telangana | Telangana | 2020-05-13 | CSIR-Centre for Cellular and Molecular Biology |
| 507 | 458058 | EPI_ISL_458058 | hCoV-19/India/CCMB_L1090/2020 | Asia / India / Telangana | Telangana | 2020-05-13 | CSIR-Centre for Cellular and Molecular Biology |
| 508 | 458059 | EPI_ISL_458059 | hCoV-19/India/CCMB_L1100/2020 | Asia / India / Telangana | Telangana | 2020-05-13 | CSIR-Centre for Cellular and Molecular Biology |
| 509 | 458060 | EPI_ISL_458060 | hCoV-19/India/CCMB_L1103/2020 | Asia / India / Telangana | Telangana | 2020-05-13 | CSIR-Centre for Cellular and Molecular Biology |
| 510 | 458061 | EPI_ISL_458061 | hCoV-19/India/CCMB_L1108/2020 | Asia / India / Telangana | Telangana | 2020-05-13 | CSIR-Centre for Cellular and Molecular Biology |
| 511 | 458062 | EPI_ISL_458062 | hCoV-19/India/CCMB_L299/2020 | Asia / India / Telangana | Telangana | 2020-04-20 | CSIR-Centre for Cellular and Molecular Biology |
| 512 | 458063 | EPI_ISL_458063 | hCoV-19/India/CCMB_L301/2020 | Asia / India / Telangana | Telangana | 2020-04-20 | CSIR-Centre for Cellular and Molecular Biology |
| 513 | 458064 | EPI_ISL_458064 | hCoV-19/India/CCMB_L988/2020 | Asia / India / Telangana | Telangana | 2020-05-11 | CSIR-Centre for Cellular and Molecular Biology |
| 514 | 458065 | EPI_ISL_458065 | hCoV-19/India/CCMB_L989/2020 | Asia / India / Telangana | Telangana | 2020-05-11 | CSIR-Centre for Cellular and Molecular Biology |
| 515 | 458066 | EPI_ISL_458066 | hCoV-19/India/CCMB_O1/2020 | Asia / India / Telangana | Telangana | 2020-04-13 | Osmania Medical College |
| 516 | 458067 | EPI_ISL_458067 | hCoV-19/India/CCMB_O2/2020 | Asia / India / Telangana | Telangana | 2020-04-15 | Osmania Medical College |
| 517 | 458068 | EPI_ISL_458068 | hCoV-19/India/CCMB_O3/2020 | Asia / India / Telangana | Telangana | 2020-04-14 | Osmania Medical College |
| 518 | 458069 | EPI_ISL_458069 | hCoV-19/India/CCMB_O4/2020 | Asia / India / Telangana | Telangana | 2020-04-03 | Osmania Medical College |
| 519 | 458070 | EPI_ISL_458070 | hCoV-19/India/CCMB_K272/2020 | Asia / India / Telangana | Telangana | 2020-04-12 | CSIR-Centre for Cellular and Molecular Biology |
| 520 | 458071 | EPI_ISL_458071 | hCoV-19/India/CCMB_J130/2020 | Asia / India / Telangana | Telangana | 2020-04-01 | CSIR-Centre for Cellular and Molecular Biology |
| 521 | 458072 | EPI_ISL_458072 | hCoV-19/India/CCMB_J812_P1/2020 | Asia / India / Telangana | Telangana | 2020-04-08 | CSIR-Centre for Cellular and Molecular Biology |
| 522 | 458073 | EPI_ISL_458073 | hCoV-19/India/CCMB_L276_P1/2020 | Asia / India / Telangana | Telangana | 2020-04-20 | CSIR-Centre for Cellular and Molecular Biology |
| 523 | 458074 | EPI_ISL_458074 | hCoV-19/India/CCMB_L301_P1/2020 | Asia / India / Telangana | Telangana | 2020-04-20 | CSIR-Centre for Cellular and Molecular Biology |
| 524 | 458075 | EPI_ISL_458075 | hCoV-19/India/CCMB_O2_P1/2020 | Asia / India / Telangana | Telangana | 2020-04-15 | CSIR-Centre for Cellular and Molecular Biology |
| 525 | 458076 | EPI_ISL_458076 | hCoV-19/India/CCMB_L302_P1/2020 | Asia / India / Telangana | Telangana | 2020-04-20 | CSIR-Centre for Cellular and Molecular Biology |
| 526 | 458077 | EPI_ISL_458077 | hCoV-19/India/CCMB_L040_P1/2020 | Asia / India / Telangana | Telangana | 2020-04-18 | CSIR-Centre for Cellular and Molecular Biology |
| 527 | 458080 | EPI_ISL_458080 | hCoV-19/India/CCMB_J300/2020 | Asia / India / Telangana | Telangana | 2020-04-03 | CSIR-Centre for Cellular and Molecular Biology |
| 528 | 458086 | EPI_ISL_458086 | hCoV-19/India/GBRC131/2020 | Asia / India / Gujarat / Ahmedabad | Gujarat | 2020-05-24 | B.J. Medical College and Civil hospital |
| 529 | 458087 | EPI_ISL_458087 | hCoV-19/India/GBRC132/2020 | Asia / India / Gujarat / Ahmedabad | Gujarat | 2020-05-24 | B.J. Medical College and Civil hospital |
| 530 | 458088 | EPI_ISL_458088 | hCoV-19/India/GBRC133/2020 | Asia / India / Gujarat / Ahmedabad | Gujarat | 2020-05-24 | B.J. Medical College and Civil hospital |
| 531 | 458089 | EPI_ISL_458089 | hCoV-19/India/GBRC134/2020 | Asia / India / Gujarat / Ahmedabad | Gujarat | 2020-05-24 | B.J. Medical College and Civil hospital |
| 532 | 458090 | EPI_ISL_458090 | hCoV-19/India/GBRC135a/2020 | Asia / India / Gujarat / Ahmedabad | Gujarat | 2020-05-24 | B.J. Medical College and Civil hospital |
| 533 | 458091 | EPI_ISL_458091 | hCoV-19/India/GBRC135b/2020 | Asia / India / Gujarat / Ahmedabad | Gujarat | 2020-05-24 | B.J. Medical College and Civil hospital |
| 534 | 458092 | EPI_ISL_458092 | hCoV-19/India/GBRC136a/2020 | Asia / India / Gujarat / Ahmedabad | Gujarat | 2020-05-24 | B.J. Medical College and Civil hospital |
| 535 | 458093 | EPI_ISL_458093 | hCoV-19/India/GBRC136b/2020 | Asia / India / Gujarat / Ahmedabad | Gujarat | 2020-05-24 | B.J. Medical College and Civil hospital |
| 536 | 458094 | EPI_ISL_458094 | hCoV-19/India/GBRC137a/2020 | Asia / India / Gujarat / Ahmedabad | Gujarat | 2020-05-24 | B.J. Medical College and Civil hospital |
| 537 | 458095 | EPI_ISL_458095 | hCoV-19/India/GBRC137b/2020 | Asia / India / Gujarat / Ahmedabad | Gujarat | 2020-05-24 | B.J. Medical College and Civil hospital |

| | | | | | | | |
|---|---|---|---|---|---|---|---|
| 538 | 458096 | EPI_ISL_458096 | hCoV-19/India/GBRC138/2020 | Asia / India / Gujarat / Ahmedabad | Gujarat | 2020-05-24 | B.J. Medical College and Civil hospital |
| 539 | 458097 | EPI_ISL_458097 | hCoV-19/India/GBRC139a/2020 | Asia / India / Gujarat / Ahmedabad | Gujarat | 2020-05-24 | B.J. Medical College and Civil hospital |
| 540 | 458098 | EPI_ISL_458098 | hCoV-19/India/GBRC139b/2020 | Asia / India / Gujarat / Ahmedabad | Gujarat | 2020-05-24 | B.J. Medical College and Civil hospital |
| 541 | 458099 | EPI_ISL_458099 | hCoV-19/India/GBRC140/2020 | Asia / India / Gujarat / Ahmedabad | Gujarat | 2020-05-24 | B.J. Medical College and Civil hospital |
| 542 | 458100 | EPI_ISL_458100 | hCoV-19/India/GBRC141a/2020 | Asia / India / Gujarat / Ahmedabad | Gujarat | 2020-05-24 | B.J. Medical College and Civil hospital |
| 543 | 458101 | EPI_ISL_458101 | hCoV-19/India/GBRC141b/2020 | Asia / India / Gujarat / Ahmedabad | Gujarat | 2020-05-24 | B.J. Medical College and Civil hospital |
| 544 | 458102 | EPI_ISL_458102 | hCoV-19/India/GBRC142/2020 | Asia / India / Gujarat / Ahmedabad | Gujarat | 2020-05-24 | B.J. Medical College and Civil hospital |
| 545 | 458103 | EPI_ISL_458103 | hCoV-19/India/GBRC143/2020 | Asia / India / Gujarat / Bayad | Gujarat | 2020-05-05 | Gujarat Biotechnology Research Centre |
| 546 | 458104 | EPI_ISL_458104 | hCoV-19/India/GBRC144/2020 | Asia / India / Gujarat / Prantij | Gujarat | 2020-05-06 | Gujarat Biotechnology Research Centre |
| 547 | 458105 | EPI_ISL_458105 | hCoV-19/India/GBRC145/2020 | Asia / India / Gujarat / Prantij | Gujarat | 2020-05-06 | Gujarat Biotechnology Research Centre |
| 548 | 458106 | EPI_ISL_458106 | hCoV-19/India/GBRC146a/2020 | Asia / India / Gujarat / Himatnagar | Gujarat | 2020-05-16 | Gujarat Biotechnology Research Centre |
| 549 | 458107 | EPI_ISL_458107 | hCoV-19/India/GBRC146b/2020 | Asia / India / Gujarat / Himatnagar | Gujarat | 2020-05-16 | Gujarat Biotechnology Research Centre |
| 550 | 458108 | EPI_ISL_458108 | hCoV-19/India/GBRC147/2020 | Asia / India / Gujarat / Khedbrahma | Gujarat | 2020-05-16 | Gujarat Biotechnology Research Centre |
| 551 | 458109 | EPI_ISL_458109 | hCoV-19/India/GBRC148a/2020 | Asia / India / Gujarat / Khedbrahma | Gujarat | 2020-05-17 | Gujarat Biotechnology Research Centre |
| 552 | 458110 | EPI_ISL_458110 | hCoV-19/India/GBRC148b/2020 | Asia / India / Gujarat / Khedbrahma | Gujarat | 2020-05-17 | Gujarat Biotechnology Research Centre |
| 553 | 458111 | EPI_ISL_458111 | hCoV-19/India/GBRC149/2020 | Asia / India / Gujarat / Himatnagar | Gujarat | 2020-05-17 | Gujarat Biotechnology Research Centre |
| 554 | 458112 | EPI_ISL_458112 | hCoV-19/India/GBRC150a/2020 | Asia / India / Gujarat / Himatnagar | Gujarat | 2020-05-18 | Gujarat Biotechnology Research Centre |
| 555 | 458113 | EPI_ISL_458113 | hCoV-19/India/GBRC150b/2020 | Asia / India / Gujarat / Himatnagar | Gujarat | 2020-05-18 | Gujarat Biotechnology Research Centre |
| 556 | 458298 | EPI_ISL_458298 | hCoV-19/India/CCMB_J714/2020 | Asia / India / Telangana | Telangana | 2020-04-06 | CSIR-Centre for Cellular and Molecular Biology |
| 557 | 459911 | EPI_ISL_459911 | hCoV-19/India/MaxCov0003_CSIR-IGIB/2020 | Asia / India / Delhi | Delhi | 2020-05-09 | Devki Devi Foundation, a unit of Max Healthcare |
| 558 | 459913 | EPI_ISL_459913 | hCoV-19/India/MaxCov0010_CSIR-IGIB/2020 | Asia / India / Delhi | Delhi | 2020-05-03 | Devki Devi Foundation, a unit of Max Healthcare |
| 559 | 459914 | EPI_ISL_459914 | hCoV-19/India/MaxCov0012_CSIR-IGIB/2020 | Asia / India / Delhi | Delhi | 2020-05-01 | Devki Devi Foundation, a unit of Max Healthcare |
| 560 | 459915 | EPI_ISL_459915 | hCoV-19/India/MaxCov0013_CSIR-IGIB/2020 | Asia / India / Delhi | Delhi | 2020-05-12 | Devki Devi Foundation, a unit of Max Healthcare |
| 561 | 459916 | EPI_ISL_459916 | hCoV-19/India/MaxCov0014_CSIR-IGIB/2020 | Asia / India / Delhi | Delhi | 2020-05-12 | Devki Devi Foundation, a unit of Max Healthcare |
| 562 | 459917 | EPI_ISL_459917 | hCoV-19/India/MaxCov0015_CSIR-IGIB/2020 | Asia / India / Delhi | Delhi | 2020-05-08 | Devki Devi Foundation, a unit of Max Healthcare |
| 563 | 459918 | EPI_ISL_459918 | hCoV-19/India/MaxCov0016_CSIR-IGIB/2020 | Asia / India / Delhi | Delhi | 2020-05-09 | Devki Devi Foundation, a unit of Max Healthcare |
| 564 | 459919 | EPI_ISL_459919 | hCoV-19/India/MaxCov0017_CSIR-IGIB/2020 | Asia / India / Delhi | Delhi | 2020-05-21 | Devki Devi Foundation, a unit of Max Healthcare |
| 565 | 459920 | EPI_ISL_459920 | hCoV-19/India/MaxCov0018_CSIR-IGIB/2020 | Asia / India / Delhi | Delhi | 2020-05-21 | Devki Devi Foundation, a unit of Max Healthcare |
| 566 | 459921 | EPI_ISL_459921 | hCoV-19/India/MaxCov0019_CSIR-IGIB/2020 | Asia / India / Delhi | Delhi | 2020-05-08 | Devki Devi Foundation, a unit of Max Healthcare |
| 567 | 459922 | EPI_ISL_459922 | hCoV-19/India/MaxCov0020_CSIR-IGIB/2020 | Asia / India / Delhi | Delhi | 2020-05-10 | Devki Devi Foundation, a unit of Max Healthcare |
| 568 | 459923 | EPI_ISL_459923 | hCoV-19/India/MaxCov0021_CSIR-IGIB/2020 | Asia / India / Delhi | Delhi | 2020-05-08 | Devki Devi Foundation, a unit of Max Healthcare |
| 569 | 459924 | EPI_ISL_459924 | hCoV-19/India/MaxCov0022_CSIR-IGIB/2020 | Asia / India / Delhi | Delhi | 2020-05-08 | Devki Devi Foundation, a unit of Max Healthcare |
| 570 | 459926 | EPI_ISL_459926 | hCoV-19/India/MaxCov0024_CSIR-IGIB/2020 | Asia / India / Delhi | Delhi | 2020 | Devki Devi Foundation, a unit of Max Healthcare |
| 571 | 459927 | EPI_ISL_459927 | hCoV-19/India/MaxCov0025_CSIR-IGIB/2020 | Asia / India / Delhi | Delhi | 2020 | Devki Devi Foundation, a unit of Max Healthcare |
| 572 | 459932 | EPI_ISL_459932 | hCoV-19/India/MaxCov0030_CSIR-IGIB/2020 | Asia / India / Delhi | Delhi | 2020-05-13 | Devki Devi Foundation, a unit of Max Healthcare |
| 573 | 459933 | EPI_ISL_459933 | hCoV-19/India/MaxCov0031_CSIR-IGIB/2020 | Asia / India / Delhi | Delhi | 2020-05-05 | Devki Devi Foundation, a unit of Max Healthcare |
| 574 | 459934 | EPI_ISL_459934 | hCoV-19/India/MaxCov0032_CSIR-IGIB/2020 | Asia / India / Delhi | Delhi | 2020-05-05 | Devki Devi Foundation, a unit of Max Healthcare |
| 575 | 459935 | EPI_ISL_459935 | hCoV-19/India/MaxCov0033_CSIR-IGIB/2020 | Asia / India / Delhi | Delhi | 2020-05-06 | Devki Devi Foundation, a unit of Max Healthcare |
| 576 | 459937 | EPI_ISL_459937 | hCoV-19/India/MaxCov0035_CSIR-IGIB/2020 | Asia / India / Delhi | Delhi | 2020-05-10 | Devki Devi Foundation, a unit of Max Healthcare |
| 577 | 459938 | EPI_ISL_459938 | hCoV-19/India/MaxCov0036_CSIR-IGIB/2020 | Asia / India / Delhi | Delhi | 2020-05-11 | Devki Devi Foundation, a unit of Max Healthcare |
| 578 | 459939 | EPI_ISL_459939 | hCoV-19/India/MaxCov0037_CSIR-IGIB/2020 | Asia / India / Delhi | Delhi | 2020-05-11 | Devki Devi Foundation, a unit of Max Healthcare |
| 579 | 459940 | EPI_ISL_459940 | hCoV-19/India/MaxCov0038_CSIR-IGIB/2020 | Asia / India / Delhi | Delhi | 2020-05-10 | Devki Devi Foundation, a unit of Max Healthcare |
| 580 | 459941 | EPI_ISL_459941 | hCoV-19/India/MaxCov0039_CSIR-IGIB/2020 | Asia / India / Delhi | Delhi | 2020-05-11 | Devki Devi Foundation, a unit of Max Healthcare |
| 581 | 459942 | EPI_ISL_459942 | hCoV-19/India/MaxCov0040_CSIR-IGIB/2020 | Asia / India / Delhi | Delhi | 2020-04-27 | Devki Devi Foundation, a unit of Max Healthcare |
| 582 | 459943 | EPI_ISL_459943 | hCoV-19/India/MaxCov0041_CSIR-IGIB/2020 | Asia / India / Delhi | Delhi | 2020-05-01 | Devki Devi Foundation, a unit of Max Healthcare |
| 583 | 461478 | EPI_ISL_461478 | hCoV-19/India/GBRC151a/2020 | Asia / India / Gujarat / Vadodara | Gujarat | 2020-05-02 | Government Medical College, Vadodara |
| 584 | 461479 | EPI_ISL_461479 | hCoV-19/India/GBRC151b/2020 | Asia / India / Gujarat / Vadodara | Gujarat | 2020-05-02 | Government Medical College, Vadodara |
| 585 | 461480 | EPI_ISL_461480 | hCoV-19/India/GBRC152/2020 | Asia / India / Gujarat / Vadodara | Gujarat | 2020-05-02 | Government Medical College, Vadodara |
| 586 | 461481 | EPI_ISL_461481 | hCoV-19/India/GBRC153a/2020 | Asia / India / Gujarat / Rajkot | Gujarat | 2020-04-27 | Pandit Deendayal Upadhyay Government Medical College, Rajkot |
| 587 | 461482 | EPI_ISL_461482 | hCoV-19/India/GBRC153b/2020 | Asia / India / Gujarat / Rajkot | Gujarat | 2020-04-27 | Pandit Deendayal Upadhyay Government Medical College, Rajkot |
| 588 | 461483 | EPI_ISL_461483 | hCoV-19/India/GBRC154/2020 | Asia / India / Gujarat / Ahmedabad | Gujarat | 2020-05-27 | B.J. Medical College and Civil hospital |
| 589 | 461484 | EPI_ISL_461484 | hCoV-19/India/GBRC155/2020 | Asia / India / Gujarat / Ahmedabad | Gujarat | 2020-05-27 | B.J. Medical College and Civil hospital |
| 590 | 461485 | EPI_ISL_461485 | hCoV-19/India/GBRC156/2020 | Asia / India / Gujarat / Ahmedabad | Gujarat | 2020-05-27 | B.J. Medical College and Civil hospital |
| 591 | 461486 | EPI_ISL_461486 | hCoV-19/India/GBRC157a/2020 | Asia / India / Gujarat / Ahmedabad | Gujarat | 2020-05-27 | B.J. Medical College and Civil hospital |
| 592 | 461487 | EPI_ISL_461487 | hCoV-19/India/GBRC157b/2020 | Asia / India / Gujarat / Ahmedabad | Gujarat | 2020-05-27 | B.J. Medical College and Civil hospital |
| 593 | 461488 | EPI_ISL_461488 | hCoV-19/India/GBRC158/2020 | Asia / India / Gujarat / Ahmedabad | Gujarat | 2020-05-27 | B.J. Medical College and Civil hospital |
| 594 | 461489 | EPI_ISL_461489 | hCoV-19/India/GBRC159a/2020 | Asia / India / Gujarat / Ahmedabad | Gujarat | 2020-05-27 | B.J. Medical College and Civil hospital |
| 595 | 461490 | EPI_ISL_461490 | hCoV-19/India/GBRC159b/2020 | Asia / India / Gujarat / Ahmedabad | Gujarat | 2020-05-27 | B.J. Medical College and Civil hospital |
| 596 | 461491 | EPI_ISL_461491 | hCoV-19/India/GBRC160a/2020 | Asia / India / Gujarat / Ahmedabad | Gujarat | 2020-05-27 | B.J. Medical College and Civil hospital |
| 597 | 461492 | EPI_ISL_461492 | hCoV-19/India/GBRC160b/2020 | Asia / India / Gujarat / Ahmedabad | Gujarat | 2020-05-27 | B.J. Medical College and Civil hospital |
| 598 | 461493 | EPI_ISL_461493 | hCoV-19/India/GBRC161a/2020 | Asia / India / Gujarat / Ahmedabad | Gujarat | 2020-05-27 | B.J. Medical College and Civil hospital |
| 599 | 461494 | EPI_ISL_461494 | hCoV-19/India/GBRC161b/2020 | Asia / India / Gujarat / Ahmedabad | Gujarat | 2020-05-27 | B.J. Medical College and Civil hospital |
| 600 | 461495 | EPI_ISL_461495 | hCoV-19/India/GBRC162a/2020 | Asia / India / Gujarat / Ahmedabad | Gujarat | 2020-05-27 | B.J. Medical College and Civil hospital |
| 601 | 461496 | EPI_ISL_461496 | hCoV-19/India/GBRC162b/2020 | Asia / India / Gujarat / Ahmedabad | Gujarat | 2020-05-27 | B.J. Medical College and Civil hospital |
| 602 | 461497 | EPI_ISL_461497 | hCoV-19/India/GBRC163a/2020 | Asia / India / Gujarat / Ahmedabad | Gujarat | 2020-05-27 | B.J. Medical College and Civil hospital |
| 603 | 461498 | EPI_ISL_461498 | hCoV-19/India/GBRC163b/2020 | Asia / India / Gujarat / Ahmedabad | Gujarat | 2020-05-27 | B.J. Medical College and Civil hospital |
| 604 | 461499 | EPI_ISL_461499 | hCoV-19/India/GBRC164/2020 | Asia / India / Gujarat / Ahmedabad | Gujarat | 2020-05-27 | B.J. Medical College and Civil hospital |
| 605 | 461500 | EPI_ISL_461500 | hCoV-19/India/GBRC165/2020 | Asia / India / Gujarat / Ahmedabad | Gujarat | 2020-05-27 | B.J. Medical College and Civil hospital |
| 606 | 461501 | EPI_ISL_461501 | hCoV-19/India/GBRC166a/2020 | Asia / India / Gujarat / Ahmedabad | Gujarat | 2020-05-27 | B.J. Medical College and Civil hospital |
| 607 | 461502 | EPI_ISL_461502 | hCoV-19/India/GBRC166b/2020 | Asia / India / Gujarat / Ahmedabad | Gujarat | 2020-05-27 | B.J. Medical College and Civil hospital |
| 608 | 461503 | EPI_ISL_461503 | hCoV-19/India/GBRC167a/2020 | Asia / India / Gujarat / Ahmedabad | Gujarat | 2020-05-27 | B.J. Medical College and Civil hospital |
| 609 | 461504 | EPI_ISL_461504 | hCoV-19/India/GBRC167b/2020 | Asia / India / Gujarat / Ahmedabad | Gujarat | 2020-05-27 | B.J. Medical College and Civil hospital |
| 610 | 461505 | EPI_ISL_461505 | hCoV-19/India/GBRC168a/2020 | Asia / India / Gujarat / Ahmedabad | Gujarat | 2020-05-27 | B.J. Medical College and Civil hospital |
| 611 | 461506 | EPI_ISL_461506 | hCoV-19/India/GBRC168b/2020 | Asia / India / Gujarat / Ahmedabad | Gujarat | 2020-05-27 | B.J. Medical College and Civil hospital |

Supplementary file 2: Details of identical sequences in the study and their geographical distribution

| Haplogroup ID | Node Label | No of Identical Sequence | Genome IDs of Identical Sequences | Date of sequence submission | No of Locations | Details of locations |
|---|---|---|---|---|---|---|
| **A1** | NC_045512.2 | 157 | NC_045512.2 | 2019-12-00 | 16 | Wuhan |
| | | | 420555 | 2020-03-03 | | Maharashtra |
| | | | 420556 | 2020-03-03 | | Maharashtra |
| | | | 426179 | 2020-03-02 | | Maharashtra |
| | | | 428479 | 2020-04-06 | | Karnataka |
| | | | 428484 | 2020-04-10 | | Karnataka |
| | | | 430467 | 2020-04-03 | | West Bengal |
| | | | 431102 | 2020-03-11 | | Telangana |
| | | | 431103 | 2020-03-16 | | Telangana |
| | | | 431117 | 2020-03-20 | | Telangana |
| | | | 435049 | 2020-04-13 | | Gujarat |
| | | | 435054 | 2020-04-14 | | Gujarat |
| | | | 435070 | 2020-03-18 | | Delhi |
| | | | 435071 | 2020-03-25 | | Delhi |
| | | | 435074 | 2020-03-28 | | West Bengal |
| | | | 435078 | 2020-03-29 | | Tamil Nadu |
| | | | 435080 | 2020-03-29 | | Tamil Nadu |
| | | | 435081 | 2020-03-28 | | West Bengal |
| | | | 435083 | 2020-03-28 | | Tamil Nadu |
| | | | 435084 | 2020-03-29 | | Tamil Nadu |
| | | | 435085 | 2020-03-29 | | Maharashtra |
| | | | 435086 | 2020-03-29 | | Maharashtra |
| | | | 435087 | 2020-03-29 | | Tamil Nadu |
| | | | 435088 | 2020-03-29 | | Odisha |
| | | | 435090 | 2020-03-29 | | Jammu |
| | | | 435091 | 2020-03-28 | | Tamil Nadu |
| | | | 435092 | 2020-03-28 | | Tamil Nadu |
| | | | 435093 | 2020-03-29 | | Tamil Nadu |
| | | | 435094 | 2020-03-29 | | Tamil Nadu |
| | | | 435095 | 2020-03-29 | | Tamil Nadu |
| | | | 435096 | 2020-03-29 | | Tamil Nadu |
| | | | 435097 | 2020-03-28 | | West Bengal |
| | | | 435098 | 2020-03-28 | | Delhi |
| | | | 435099 | 2020-03-29 | | Uttar Pradesh |
| | | | 435100 | 2020-03-29 | | Uttar Pradesh |
| | | | 435112 | 2020-03-28 | | Bihar |
| | | | 436413 | 2020-03-30 | | Uttar Pradesh |
| | | | 436417 | 2020-03-31 | | Bihar |
| | | | 436418 | 2020-03-31 | | Tamil Nadu |
| | | | 436420 | 2020-03-31 | | Rajasthan |
| | | | 436422 | 2020-03-31 | | Assam |
| | | | 436424 | 2020-04-05 | | Delhi |
| | | | 436428 | 2020-04-05 | | Delhi |
| | | | 436429 | 2020-04-05 | | Delhi |
| | | | 436430 | 2020-04-05 | | Delhi |
| | | | 436431 | 2020-04-05 | | Delhi |
| | | | 436433 | 2020-04-05 | | Delhi |
| | | | 436435 | 2020-04-09 | | Delhi |
| | | | 436436 | 2020-04-06 | | Delhi |
| | | | 436437 | 2020-04-08 | | Delhi |
| | | | 436445 | 2020-04-09 | | Delhi |
| | | | 436447 | 2020-04-10 | | Karnataka |
| | | | 436448 | 2020-04-12 | | Delhi |
| | | | 436451 | 2020-04-13 | | Delhi |
| | | | 436453 | 2020-04-16 | | Madhya Pradesh |
| | | | 436455 | 2020-04-13 | | Delhi |
| | | | 436460 | 2020-04-20 | | Madhya Pradesh |
| | | | 436461 | 2020-04-20 | | Madhya Pradesh |
| | | | 436462 | 2020-04-16 | | Madhya Pradesh |
| | | | 436463 | 2020-04-16 | | Madhya Pradesh |
| | | | 437539 | 2020-04-06 | | West Bengal |
| | | | 438138 | 2020-03-25 | | Telangana |
| | | | 438139 | 2020-03-25 | | Telangana |
| | | | 444481 | 2020-04-30 | | Gujarat |
| | | | 447031 | 2020-05-03 | | Gujarat |
| | | | 447032 | 2020-05-03 | | Gujarat |
| | | | 447038 | 2020-05-03 | | Gujarat |
| | | | 447047 | 2020-04-29 | | Gujarat |
| | | | 447554 | 2020-04-26 | | Gujarat |
| | | | 447556 | 2020-03-30 | | Telangana |
| | | | 447557 | 2020-03-30 | | Telangana |
| | | | 447558 | 2020-03-30 | | Telangana |

| | | |
|---|---|---|
| 447559 | 2020-03-31 | Telangana |
| 447560 | 2020-03-31 | Telangana |
| 447561 | 2020-03-31 | Telangana |
| 447562 | 2020-03-31 | Telangana |
| 447564 | 2020-04-01 | Telangana |
| 447565 | 2020-04-01 | Telangana |
| 447566 | 2020-04-01 | Telangana |
| 447567 | 2020-04-01 | Telangana |
| 447574 | 2020-04-01 | Telangana |
| 447575 | 2020-04-02 | Telangana |
| 447577 | 2020-04-02 | Telangana |
| 447582 | 2020-04-14 | Telangana |
| 447847 | 2020-03-31 | Telangana |
| 447850 | 2020-04-01 | Telangana |
| 447854 | 2020-04-02 | Telangana |
| 447855 | 2020-04-02 | Telangana |
| 447856 | 2020-04-02 | Telangana |
| 447857 | 2020-04-02 | Telangana |
| 447860 | 2020-04-16 | Telangana |
| 447861 | 2020-04-16 | Telangana |
| 447862 | 2020-03-31 | Telangana |
| 447863 | 2020-04-14 | Telangana |
| 447865 | 2020-04-14 | Telangana |
| 450321 | 2020-04-04 | Maharashtra |
| 450322 | 2020-04-04 | Maharashtra |
| 450323 | 2020-03-27 | Maharashtra |
| 450326 | 2020-04-01 | Telangana |
| 450327 | 2020-04-01 | Telangana |
| 450328 | 2020-04-01 | Telangana |
| 450329 | 2020-04-01 | Telangana |
| 450330 | 2020-04-02 | Telangana |
| 450331 | 2020-04-01 | Telangana |
| 450332 | 2020-04-01 | Telangana |
| 452208 | 2020-04-05 | Maharashtra |
| 452209 | 2020-04-06 | Maharashtra |
| 452790 | 2020-04-24 | Madhya Pradesh |
| 454532 | 2020-03-23 | Maharashtra |
| 454533 | 2020-03-23 | Maharashtra |
| 454543 | 2020-03-31 | Maharashtra |
| 454546 | 2020-04-03 | Maharashtra |
| 454547 | 2020-04-04 | Maharashtra |
| 454565 | 2020-03-31 | Maharashtra |
| 454832 | 2020-04-21 | Rajasthan |
| 454833 | 2020-04-27 | Rajasthan |
| 454862 | 2020-04-11 | Haryana |
| 454867 | 2020-04-16 | Haryana |
| 455643 | 2020-03-30 | West Bengal |
| 455644 | 2020-04-02 | West Bengal |
| 455656 | 2020-04-30 | West Bengal |
| 455679 | 2020-05-04 | West Bengal |
| 455786 | 2020-05-04 | Odisha |
| 455787 | 2020-05-04 | Odisha |
| 458062 | 2020-04-20 | Telangana |
| 458066 | 2020-04-13 | Telangana |
| 458069 | 2020-04-03 | Telangana |
| 458071 | 2020-04-01 | Telangana |
| 458073 | 2020-04-20 | Telangana |
| 458076 | 2020-04-20 | Telangana |
| 458080 | 2020-04-03 | Telangana |
| 458298 | 2020-04-06 | Telangana |
| 459913 | 2020-05-03 | Delhi |
| 459915 | 2020-05-12 | Delhi |
| 459916 | 2020-05-12 | Delhi |
| 459917 | 2020-05-08 | Delhi |
| 459918 | 2020-05-09 | Delhi |
| 459919 | 2020-05-21 | Delhi |
| 459920 | 2020-05-21 | Delhi |
| 459921 | 2020-05-08 | Delhi |
| 459922 | 2020-05-10 | Delhi |
| 459923 | 2020-05-08 | Delhi |
| 459924 | 2020-05-08 | Delhi |
| 459926 | 2020-05-08 | Delhi |
| 459927 | 2020-05-08 | Delhi |
| 459932 | 2020-05-13 | Delhi |

| | | | | | | |
|---|---|---|---|---|---|---|
| | | | 459933 | 2020-05-05 | | Delhi |
| | | | 459934 | 2020-05-05 | | Delhi |
| | | | 459935 | 2020-05-06 | | Delhi |
| | | | 459937 | 2020-05-10 | | Delhi |
| | | | 459938 | 2020-05-11 | | Delhi |
| | | | 459939 | 2020-05-11 | | Delhi |
| | | | 459940 | 2020-05-10 | | Delhi |
| | | | 459941 | 2020-05-11 | | Delhi |
| | | | 459942 | 2020-04-27 | | Delhi |
| | | | 459943 | 2020-05-01 | | Delhi |
| | | | 461505 | 2020-05-27 | | Gujarat |
| **A1.1** | 420544 | 10 | 420544 | 2020-03-03 | 1 | Maharashtra |
| | | | 420546 | 2020-03-03 | | Maharashtra |
| | | | 420547 | 2020-03-03 | | Maharashtra |
| | | | 420548 | 2020-03-03 | | Maharashtra |
| | | | 420549 | 2020-03-03 | | Maharashtra |
| | | | 420550 | 2020-03-03 | | Maharashtra |
| | | | 420551 | 2020-03-03 | | Maharashtra |
| | | | 420552 | 2020-03-03 | | Maharashtra |
| | | | 420553 | 2020-03-03 | | Maharashtra |
| | | | 420554 | 2020-03-03 | | Maharashtra |
| **A1.1.1** | 420543 | 2 | 420543 | 2020-03-03 | 1 | Maharashtra |
| | | | 420545 | 2020-03-03 | | Maharashtra |
| **A1.10** | 444479 | 3 | 444479 | 2020-04-29 | 1 | Gujarat |
| | | | 447052 | 2020-05-02 | | Gujarat |
| | | | 447547 | 2020-04-28 | | Gujarat |
| **A1.11** | 444483 | 7 | 444483 | 2020-05-02 | 1 | Gujarat |
| | | | 447544 | 2020-05-05 | | Gujarat |
| | | | 447545 | 2020-05-05 | | Gujarat |
| | | | 458104 | 2020-05-06 | | Gujarat |
| | | | 458105 | 2020-05-06 | | Gujarat |
| | | | 458106 | 2020-05-16 | | Gujarat |
| | | | 458107 | 2020-05-16 | | Gujarat |
| **A1.12** | 447584 | 3 | 447584 | 2020-04-16 | 1 | Tamil Nadu |
| | | | 447585 | 2020-04-16 | | Tamil Nadu |
| | | | 447586 | 2020-04-16 | | Tamil Nadu |
| **A1.13** | 451158 | 5 | 451158 | 2020-05-03 | 1 | Gujarat |
| | | | 455019 | 2020-05-02 | | Gujarat |
| | | | 455022 | 2020-05-02 | | Gujarat |
| | | | 455024 | 2020-05-02 | | Gujarat |
| | | | 455025 | 2020-05-02 | | Gujarat |
| **A1.14** | 452192 | 6 | 452192 | 2020-04-16 | 2 | Maharashtra |
| | | | 452199 | 2020-04-20 | | Maharashtra |
| | | | 452207 | 2020-04-05 | | Maharashtra |
| | | | 452211 | 2020-04-13 | | Maharashtra |
| | | | 454557 | 2020-04-13 | | Maharashtra |
| | | | 458109 | 2020-05-17 | | Gujarat |
| **A1.14.1** | 450785 | 2 | 450785 | 2020-05-10 | 1 | Gujarat |
| | | | 450786 | 2020-05-10 | | Gujarat |
| **A1.14.2** | 458059 | 3 | 458059 | 2020-05-13 | 1 | Telangana |
| | | | 458060 | 2020-05-13 | | Telangana |
| | | | 458061 | 2020-05-13 | | Telangana |
| **A1.15** | 452213 | 3 | 452213 | 2020-03-10 | 1 | Maharashtra |
| | | | 454525 | 2020-03-12 | | Maharashtra |
| | | | 454526 | 2020-03-13 | | Maharashtra |
| **A1.16** | 452214 | 9 | 452214 | 2020-04-07 | 3 | Maharashtra |
| | | | 454529 | 2020-03-21 | | Maharashtra |
| | | | 454570 | 2020-04-26 | | Maharashtra |
| | | | 458053 | 2020-05-12 | | Telangana |
| | | | 458108 | 2020-05-16 | | Gujarat |
| | | | 461481 | 2020-04-27 | | Gujarat |
| | | | 461483 | 2020-05-27 | | Gujarat |
| | | | 461493 | 2020-05-27 | | Gujarat |
| | | | 461496 | 2020-05-27 | | Gujarat |
| **A1.17** | 455660 | 2 | 455660 | 2020-05-01 | 1 | West Bengal |
| | | | 455678 | 2020-04-30 | | West Bengal |
| **A1.18** | 458063 | 2 | 458063 | 2020-04-20 | 1 | Telangana |
| | | | 458074 | 2020-04-20 | | Telangana |
| **A1.19** | 461490 | 2 | 461490 | 2020-05-27 | 1 | Gujarat |
| | | | 461503 | 2020-05-27 | | Gujarat |
| **A1.2** | 424364 | 2 | 424364 | 2020-03-17 | 1 | Maharashtra |
| | | | 424365 | 2020-03-17 | | Maharashtra |
| **A1.20** | 455667 | 4 | 455667 | 2020-05-03 | 1 | West Bengal |
| | | | 455673 | 2020-05-02 | | West Bengal |

| | | | | | | |
|---|---|---|---|---|---|---|
| | | | 455674 | 2020-05-03 | | West Bengal |
| | | | 455676 | 2020-05-03 | | West Bengal |
| **A1.21** | 458046 | 2 | 458046 | 2020-05-11 | 1 | Telangana |
| | | | 458048 | 2020-05-11 | | Telangana |
| **A1.22** | 458064 | 2 | 458064 | 2020-05-11 | 1 | Telangana |
| | | | 458065 | 2020-05-11 | | Telangana |
| **A1.23** | 435101 | 3 | 435101 | 2020-03-15 | 1 | Ladakh |
| | | | 435102 | 2020-03-15 | | Ladakh |
| | | | 435103 | 2020-03-17 | | Ladakh |
| **A1.24** | 437442 | 2 | 437442 | 2020-04-27 | 1 | Gujarat |
| | | | 437444 | 2020-04-26 | | Gujarat |
| **A1.25** | 447858 | 2 | 447858 | 2020-04-06 | 1 | Telangana |
| | | | 458072 | 2020-04-08 | | Telangana |
| **A1.26** | 450790 | 4 | 450790 | 2020-04-28 | 1 | Gujarat |
| | | | 450791 | 2020-04-27 | | Gujarat |
| | | | 455015 | 2020-04-28 | | Gujarat |
| | | | 455016 | 2020-04-27 | | Gujarat |
| **A1.27** | 451154 | 5 | 451154 | 2020-05-03 | 2 | Gujarat |
| | | | 451156 | 2020-05-01 | | Gujarat |
| | | | 451159 | 2020-05-03 | | Gujarat |
| | | | 451161 | 2020-05-03 | | Gujarat |
| | | | 452793 | 2020-05-03 | | Madhya Pradesh |
| **A1.28** | 452204 | 2 | 452204 | 2020-03-26 | 1 | Maharashtra |
| | | | 454531 | 2020-03-22 | | Maharashtra |
| **A1.29** | 452205 | 2 | 452205 | 2020-03-26 | 1 | Maharashtra |
| | | | 454534 | 2020-03-24 | | Maharashtra |
| **A1.3** | 430464 | 2 | 430464 | 2020-03-21 | 1 | West Bengal |
| | | | 430468 | 2020-03-21 | | West Bengal |
| **A1.30** | 455653 | 2 | 455653 | 2020-04-21 | 1 | West Bengal |
| | | | 455675 | 2020-05-03 | | West Bengal |
| **A1.31** | 455764 | 3 | 455764 | 2020-05-07 | 1 | Odisha |
| | | | 455766 | 2020-05-07 | | Odisha |
| | | | 455767 | 2020-05-07 | | Odisha |
| **A1.4** | 430465 | 3 | 430465 | 2020-03-28 | 2 | West Bengal |
| | | | 458030 | 2020-04-26 | | Tamil Nadu |
| | | | 458032 | 2020-04-27 | | Tamil Nadu |
| **A1.4.1** | 458031 | 4 | 458031 | 2020-04-26 | 1 | Tamil Nadu |
| | | | 458033 | 2020-04-29 | | Tamil Nadu |
| | | | 458034 | 2020-04-29 | | Tamil Nadu |
| | | | 458040 | 2020-05-06 | | Tamil Nadu |
| **A1.4.2** | 458037 | 2 | 458037 | 2020-04-29 | 1 | Tamil Nadu |
| | | | 458044 | 2020-05-06 | | Tamil Nadu |
| **A1.4.3** | 458038 | 3 | 458038 | 2020-04-29 | 1 | Tamil Nadu |
| | | | 458039 | 2020-05-06 | | Tamil Nadu |
| | | | 458041 | 2020-05-06 | | Tamil Nadu |
| **A1.5** | 435056 | 2 | 435056 | 2020-04-21 | 1 | Gujarat |
| | | | 450788 | 2020-05-06 | | Gujarat |
| **A1.6** | 435063 | 3 | 435063 | 2020-03-13 | 2 | Delhi |
| | | | 435064 | 2020-03-18 | | Delhi |
| | | | 437626 | 2020-03-24 | | Telangana |
| **A1.6.1** | 444471 | 13 | 444471 | 2020-04-29 | 2 | Gujarat |
| | | | 444482 | 2020-05-02 | | Gujarat |
| | | | 447555 | 2020-04-28 | | Gujarat |
| | | | 451160 | 2020-05-03 | | Gujarat |
| | | | 455775 | 2020-04-04 | | Odisha |
| | | | 455777 | 2020-04-04 | | Odisha |
| | | | 455778 | 2020-04-04 | | Odisha |
| | | | 455779 | 2020-04-04 | | Odisha |
| | | | 455780 | 2020-04-08 | | Odisha |
| | | | 455782 | 2020-04-02 | | Odisha |
| | | | 455784 | 2020-04-02 | | Odisha |
| | | | 458087 | 2020-05-24 | | Gujarat |
| | | | 458089 | 2020-05-24 | | Gujarat |
| **A1.6.1.1** | 435065 | 6 | 435065 | 2020-03-15 | 2 | Delhi |
| | | | 435066 | 2020-03-18 | | Delhi |
| | | | 435068 | 2020-03-18 | | Delhi |
| | | | 435069 | 2020-03-18 | | Delhi |
| | | | 447041 | 2020-05-03 | | Gujarat |
| | | | 458095 | 2020-05-24 | | Gujarat |
| **A1.6.1.1.1** | 444461 | 9 | 444461 | 2020-04-29 | 1 | Gujarat |
| | | | 444472 | 2020-04-29 | | Gujarat |
| | | | 447039 | 2020-05-03 | | Gujarat |
| | | | 447040 | 2020-05-03 | | Gujarat |
| | | | 458088 | 2020-05-24 | | Gujarat |

| | | | | | | |
|---|---|---|---|---|---|---|
| | | | 458094 | 2020-05-24 | | Gujarat |
| | | | 458097 | 2020-05-24 | | Gujarat |
| | | | 458098 | 2020-05-24 | | Gujarat |
| | | | 461500 | 2020-05-27 | | Gujarat |
| **A1.6.1.1.1.1** | 435055 | 4 | 435055 | 2020-04-22 | 1 | Gujarat |
| | | | 447050 | 2020-04-29 | | Gujarat |
| | | | 447051 | 2020-04-29 | | Gujarat |
| | | | 447548 | 2020-04-28 | | Gujarat |
| **A1.6.1.1.1.2** | 444465 | 3 | 444465 | 2020-04-29 | 1 | Gujarat |
| | | | 444466 | 2020-04-29 | | Gujarat |
| | | | 447035 | 2020-05-03 | | Gujarat |
| **A1.6.1.1.1.3** | 447033 | 3 | 447033 | 2020-05-03 | 1 | Gujarat |
| | | | 451153 | 2020-05-10 | | Gujarat |
| | | | 458112 | 2020-05-18 | | Gujarat |
| **A1.6.1.1.1.4** | 451149 | 2 | 451149 | 2020-05-07 | 1 | Gujarat |
| | | | 451151 | 2020-05-05 | | Gujarat |
| **A1.6.1.1.2** | 444469 | 4 | 444469 | 2020-04-29 | 1 | Gujarat |
| | | | 447034 | 2020-05-03 | | Gujarat |
| | | | 447044 | 2020-05-03 | | Gujarat |
| | | | 447546 | 2020-05-05 | | Gujarat |
| **A1.6.1.2** | 444456 | 4 | 444456 | 2020-04-26 | 1 | Gujarat |
| | | | 444457 | 2020-04-26 | | Gujarat |
| **A1.6.1.3** | 444484 | 7 | 444484 | 2020-05-04 | 1 | Gujarat |
| | | | 444485 | 2020-05-04 | | Gujarat |
| | | | 447535 | 2020-05-05 | | Gujarat |
| | | | 447540 | 2020-05-05 | | Gujarat |
| | | | 447541 | 2020-05-05 | | Gujarat |
| | | | 447542 | 2020-05-05 | | Gujarat |
| | | | 447543 | 2020-05-05 | | Gujarat |
| **A1.6.1.4** | 455021 | 2 | 455021 | 2020-05-02 | 1 | Gujarat |
| | | | 455027 | 2020-05-02 | | Gujarat |
| **A1.6.1.5** | 437449 | 4 | 437449 | 2020-04-26 | 1 | Gujarat |
| | | | 444470 | 2020-04-29 | | Gujarat |
| | | | 447536 | 2020-05-05 | | Gujarat |
| | | | 447538 | 2020-05-05 | | Gujarat |
| **A1.7** | 436414 | 2 | 436414 | 2020-03-31 | 2 | West Bengal |
| | | | 436421 | 2020-03-31 | | Assam |
| **A1.8** | 436426 | 2 | 436426 | 2020-04-05 | 2 | Delhi |
| | | | 436449 | 2020-04-12 | | Bihar |
| **A1.9** | 437447 | 4 | 437447 | 2020-04-26 | 1 | Gujarat |
| | | | 437448 | 2020-04-26 | | Gujarat |
| | | | 444460 | 2020-04-30 | | Gujarat |
| | | | 447550 | 2020-05-03 | | Gujarat |
| **A1.9.1** | 447549 | 2 | 447549 | 2020-05-03 | 1 | Gujarat |
| | | | 447551 | 2020-04-25 | | Gujarat |

**Supplementary files 3: Details of parsimony informative sites including gaps and ambiguous sequences observed in the study**

| | | |
|---|---|---|
| | Pi Site Excluding Ai | 152 |
| | Pi site Combine (G | 270 |

| | S No | 1 | 2 | 3 | 4 | 5 | 6 | 7 | 8 | 9 | 10 | 11 | 12 | 13 | 14 | 15 | 16 | 17 | 18 | 19 | 20 | 21 | 22 | 23 | 24 |
|---|---|---|---|---|---|---|---|---|---|---|---|---|---|---|---|---|---|---|---|---|---|---|---|---|---|
| Haplogroup ID | Gene Location | 5'UTR | 5'UTR | 5'UTR | 5'UTR | 5'UTR | 5'UTR | 5'UTR | 5'UTR | 5'UTR | ORF1a | ORF1a | ORF1a | ORF1a | ORF1a | ORF1a | ORF1a | ORF1a | ORF1a | ORF1a | ORF1a | ORF1a | ORF1a | ORF1a | ORF1a |
| | Genome ID | 22 | 55 | 56 | 94 | 106 | 218 | 219 | 222 | 241 | 313 | 506 | 635 | 771 | 875 | 884 | 1059 | 1094 | 1191 | 1218 | 1281 | 1397 | 1589 | 1599 | 1707 |
| **A1** | NC 045512.2 | A | A | G | G | C | C | G | C | C | C | C | C | T | C | C | C | T | C | C | C | G | G | G | C |
| | 413522 | . | . | . | . | . | . | . | . | . | . | . | . | . | . | . | . | . | . | . | . | . | . | . | . |
| | 413523 | . | . | . | . | . | . | . | . | . | . | . | . | . | . | . | . | . | . | . | . | . | . | . | . |
| **A1.1.1** | 420543 | . | . | . | . | . | . | . | . | T | . | . | . | . | . | . | . | . | . | . | . | . | . | . | . |
| **A1.1** | 420544 | . | . | . | . | . | . | . | . | T | . | . | . | . | . | . | . | . | . | . | . | . | . | . | . |
| | 420545 | . | . | . | . | . | . | . | . | T | . | . | . | . | . | . | . | . | . | . | . | . | . | . | . |
| | 420546 | . | . | . | . | . | . | . | . | T | . | . | . | . | . | . | . | . | . | . | . | . | . | . | . |
| | 420547 | . | . | . | . | . | . | . | . | T | . | . | . | . | . | . | . | . | . | . | . | . | . | . | . |
| | 420548 | . | . | . | . | . | . | . | . | T | . | . | . | . | . | . | . | . | . | . | . | . | . | . | . |
| | 420549 | . | . | . | . | . | . | . | . | T | . | . | . | . | . | . | . | . | . | . | . | . | . | . | . |
| | 420550 | . | . | . | . | . | . | . | . | T | . | . | . | . | . | . | . | . | . | . | . | . | . | . | . |
| | 420551 | . | . | . | . | . | . | . | . | T | . | . | . | . | . | . | . | . | . | . | . | . | . | . | . |
| | 420552 | . | . | . | . | . | . | . | . | T | . | . | . | . | . | . | . | . | . | . | . | . | . | . | . |
| | 420553 | . | . | . | . | . | . | . | . | T | . | . | . | . | . | . | . | . | . | . | . | . | . | . | . |
| | 420554 | . | . | . | . | . | . | . | . | T | . | . | . | . | . | . | . | . | . | . | . | . | . | . | . |
| | 420555 | . | . | . | . | . | . | . | . | T | T | . | . | . | . | . | . | . | . | . | . | . | . | . | . |
| | 420556 | . | . | . | . | . | . | . | . | T | T | . | . | . | . | . | . | . | . | . | . | . | . | . | . |
| | 421662 | . | . | . | . | . | . | . | . | . | . | . | . | . | . | T | . | . | . | . | . | A | . | . | . |
| | 421663 | - | . | . | . | . | . | . | . | . | . | . | . | . | . | T | . | . | . | . | . | A | . | . | . |
| | 421664 | . | . | . | . | . | . | . | . | . | . | . | . | . | . | T | . | . | . | . | . | A | . | . | . |
| | 421665 | . | . | . | . | . | . | . | . | . | . | . | . | . | . | T | . | . | . | . | . | A | . | . | . |
| | 421666 | . | . | . | . | . | . | . | . | . | . | . | . | . | . | . | . | . | . | . | . | A | . | . | . |
| | 421667 | . | . | . | . | . | . | . | . | . | . | . | . | . | . | T | . | . | . | . | . | A | . | . | . |
| | 421668 | - | . | . | . | . | . | . | . | . | . | . | . | . | . | . | . | . | . | . | . | A | . | . | . |
| | 421669 | . | . | . | . | . | . | . | . | . | . | . | . | . | . | T | . | . | . | . | . | A | . | . | . |
| | 421670 | . | . | . | . | . | . | . | . | . | . | . | . | . | . | T | . | . | . | . | . | A | . | . | . |
| | 421671 | . | . | . | . | . | . | . | . | . | . | . | . | . | . | T | . | . | . | . | . | A | . | . | . |
| | 421672 | . | . | . | . | . | . | . | . | . | . | . | . | . | . | T | . | . | . | . | . | A | . | . | . |
| | 424361 | . | . | . | . | . | . | . | . | . | . | . | . | . | . | T | . | . | . | . | . | A | . | . | . |
| | 424362 | . | . | . | . | . | . | . | . | T | . | . | . | . | . | . | . | . | . | . | . | . | . | . | . |
| | 424363 | - | . | . | . | . | . | . | . | . | . | . | . | . | . | T | . | . | . | . | . | A | . | . | . |
| **A1.2** | 424364 | . | . | . | . | . | . | . | . | T | . | . | . | . | . | . | . | . | . | . | . | . | . | . | . |
| | 424365 | . | . | . | . | . | . | . | . | T | . | . | . | . | . | . | . | . | . | . | . | . | . | . | . |
| | 426179 | . | . | . | . | . | . | . | . | T | T | . | . | . | . | . | . | . | . | . | . | . | . | . | . |
| | 426414 | . | . | . | . | . | . | . | . | T | . | . | . | . | . | . | . | . | . | . | . | . | . | . | . |
| | 426415 | . | . | . | . | . | . | . | . | T | . | . | . | . | . | . | . | . | . | . | . | . | . | . | . |
| | 428479 | - | . | . | . | . | . | . | . | T | T | . | . | . | . | . | . | . | . | . | . | . | . | . | . |
| | 428480 | - | . | . | . | . | . | . | . | T | T | . | . | . | . | . | . | . | . | . | . | ? | ? | ? | ? |
| | 428481 | - | . | . | . | . | . | . | . | T | T | . | . | . | . | . | . | . | . | . | . | . | . | . | . |
| | 428482 | - | . | . | . | . | . | . | . | T | T | . | . | . | . | . | . | . | . | . | . | . | . | . | . |
| | 428483 | - | . | . | . | . | . | . | . | . | . | . | . | . | . | . | . | . | . | . | . | . | . | . | . |
| | 428484 | - | . | . | . | . | . | . | . | . | . | . | . | . | . | . | . | . | . | . | . | . | . | . | . |
| | 428485 | - | . | . | . | . | . | . | . | Y | T | . | . | . | . | . | . | . | . | . | . | ? | ? | ? | ? |
| | 428486 | - | . | . | . | . | . | . | . | . | . | . | . | . | . | . | . | . | . | . | . | . | . | . | . |
| | 428487 | - | . | . | . | . | . | . | . | . | . | . | . | . | . | . | . | . | . | . | . | . | . | . | . |
| **A1.3** | 430464 | . | . | . | . | . | . | . | . | T | . | . | . | . | . | . | . | . | . | . | . | . | . | . | . |
| **A1.4** | 430465 | . | . | . | . | . | . | . | . | T | . | . | . | . | . | . | . | . | . | . | . | . | . | . | . |
| | 430466 | . | . | . | . | . | . | . | . | T | . | . | . | . | . | . | . | . | . | . | . | . | . | . | . |
| | 430467 | . | . | . | . | . | . | . | . | T | . | . | . | . | . | . | . | . | . | . | . | . | . | . | . |
| | 430468 | . | . | . | . | . | . | . | . | T | . | . | . | . | . | . | . | . | . | . | . | . | . | . | . |
| | 431101 | . | . | . | . | . | . | . | . | . | . | . | . | . | . | . | . | . | . | . | . | . | . | . | . |
| | 431102 | . | . | . | . | . | . | . | . | T | . | . | . | . | . | . | . | . | . | . | . | . | . | . | . |
| | 431103 | . | . | . | . | . | . | . | . | . | . | . | . | . | . | . | . | . | . | . | . | . | . | . | Y |
| | 431117 | . | . | . | . | . | . | . | . | T | . | . | . | . | . | . | . | . | . | . | . | . | . | . | . |
| | 435049 | ? | . | . | . | . | . | . | . | T | . | . | . | . | . | . | . | . | . | . | . | . | . | . | . |
| | 435050 | ? | . | . | . | . | . | . | . | T | . | . | . | . | . | . | . | . | . | . | . | . | . | . | . |

| | | | | | | | | | | | | | | | | | | | | | | | | | |
|---|---|---|---|---|---|---|---|---|---|---|---|---|---|---|---|---|---|---|---|---|---|---|---|---|---|
| | 435051 | ? | . | . | . | . | . | . | . | T | . | . | . | . | . | . | T | . | . | . | . | . | . | . | . |
| | 435052 | ? | . | . | . | . | . | . | . | T | . | . | . | . | . | . | T | . | . | . | . | . | . | . | ? |
| | 435053 | ? | . | . | . | . | . | . | . | T | . | . | . | . | . | . | . | . | . | . | . | . | . | . | . |
| | 435054 | ? | . | . | . | . | . | . | . | T | . | . | . | . | . | . | . | . | . | . | . | . | . | . | ? |
| **A1.6.1.1.1.1** | 435055 | ? | . | . | . | . | . | . | . | T | . | . | . | . | . | . | . | . | . | . | . | . | . | . | . |
| **A1.5** | 435056 | ? | . | . | . | . | . | . | . | T | . | . | . | . | . | . | . | . | . | . | . | . | . | . | . |
| | 435060 | ? | . | . | . | . | . | . | . | T | . | . | . | . | . | . | . | . | . | . | . | g | . | . | . |
| | 435061 | . | . | . | . | . | . | . | . | T | . | . | . | . | . | . | . | . | . | . | . | . | . | . | . |
| | 435062 | . | . | . | . | . | . | . | . | T | . | . | . | . | . | . | . | . | . | . | . | . | . | . | . |
| **A1.6** | 435063 | . | . | . | . | . | . | . | . | T | . | . | . | . | . | . | T | . | . | . | . | . | . | . | . |
| | 435064 | . | . | . | . | . | . | . | . | T | . | . | . | . | . | . | T | . | . | . | . | . | . | . | . |
| **A1.6.1.1** | 435065 | a | . | . | . | . | . | . | . | T | . | . | . | . | . | . | . | . | . | . | . | . | . | . | . |
| | 435066 | . | . | . | . | . | . | . | . | T | . | . | . | . | . | . | . | . | . | . | . | . | . | . | . |
| | 435067 | . | . | . | . | . | . | . | . | T | . | . | . | . | . | . | . | . | . | . | . | . | . | . | . |
| | 435068 | . | . | . | . | . | . | . | . | T | . | . | . | . | . | . | . | . | . | . | . | . | . | . | . |
| | 435069 | . | . | . | . | . | . | . | . | T | . | . | . | . | . | . | . | . | . | . | . | . | . | . | . |
| | 435070 | . | . | . | . | . | . | . | . | T | . | . | . | . | . | . | . | . | . | . | . | . | . | . | . |
| | 435071 | . | . | . | . | . | . | . | . | T | . | . | . | . | . | . | . | . | . | . | . | . | . | . | . |
| | 435072 | ? | a | g | g | c | c | . | . | T | c | . | . | . | . | . | . | . | . | . | . | . | . | . | . |
| | 435074 | ? | . | . | . | . | . | . | . | . | . | . | . | . | . | . | . | . | . | . | . | . | . | . | . |
| | 435078 | ? | . | . | . | . | . | . | . | c | . | . | . | . | . | Y | . | . | . | . | . | . | . | . | . |
| | 435080 | a | . | . | . | . | . | . | . | . | . | . | . | . | . | . | . | . | . | . | . | . | . | . | . |
| | 435081 | . | . | . | . | . | . | . | . | ? | . | . | . | . | . | . | . | . | . | . | . | . | . | . | . |
| | 435082 | . | . | . | g | . | . | . | . | . | . | . | . | . | . | . | . | . | . | . | . | . | . | . | . |
| | 435083 | . | . | . | . | . | . | . | . | . | . | . | . | . | . | . | . | . | . | . | . | g | . | . | . |
| | 435084 | . | . | . | . | . | . | . | . | . | . | . | . | . | . | . | . | . | . | . | . | . | . | . | . |
| | 435085 | . | . | . | . | . | . | . | . | . | . | . | . | . | . | . | . | . | . | . | Y | . | . | . | . |
| | 435086 | . | . | . | . | . | . | . | . | . | . | . | . | . | . | . | . | . | . | . | . | g | . | . | . |
| | 435087 | ? | . | . | . | . | . | . | . | . | . | . | . | . | . | . | . | . | . | . | . | . | . | . | . |
| | 435088 | . | . | . | . | . | . | . | . | . | . | . | . | . | . | . | . | . | . | . | T | . | . | . | . |
| | 435090 | . | . | . | . | . | . | . | . | . | . | . | . | . | . | . | . | . | . | . | . | . | . | . | T |
| | 435091 | . | . | . | . | . | . | . | . | . | . | . | . | . | . | . | . | . | . | . | . | . | . | . | T |
| | 435092 | . | . | . | . | . | . | . | . | . | . | . | . | . | . | . | . | . | . | . | . | . | . | . | . |
| | 435093 | . | . | . | . | . | . | . | . | . | . | . | . | t | . | . | . | . | . | . | . | . | . | . | . |
| | 435094 | . | . | . | . | . | . | . | . | . | . | . | . | . | . | . | . | . | . | . | . | . | . | . | . |
| | 435095 | . | . | . | . | . | . | . | . | . | . | . | . | . | . | . | . | . | . | . | . | . | . | . | . |
| | 435096 | . | . | . | . | . | . | . | . | . | . | . | . | . | . | . | . | . | . | . | . | . | . | . | . |
| | 435097 | . | . | . | . | . | . | . | . | . | . | . | . | . | . | . | . | . | . | . | . | . | . | . | . |
| | 435098 | a | . | . | . | . | . | . | . | . | . | . | . | . | . | . | . | . | . | . | . | . | . | . | . |
| | 435099 | . | . | . | . | . | . | . | . | . | . | . | . | . | . | . | . | . | . | . | . | . | . | . | . |
| | 435100 | . | . | . | . | . | . | . | . | . | . | . | . | . | . | . | . | . | . | . | . | . | . | . | . |
| **A1.23** | 435101 | a | . | . | . | . | . | . | . | . | . | . | . | . | . | T | . | . | . | . | . | A | . | . | . |
| | 435102 | . | . | . | . | . | . | . | . | . | . | . | . | . | . | T | . | . | . | . | . | A | . | . | . |
| | 435103 | . | . | . | . | . | . | . | . | . | . | . | . | . | . | T | . | . | . | . | . | A | . | . | . |
| | 435104 | . | . | . | . | . | . | . | . | . | . | . | . | . | . | T | . | . | . | . | . | A | . | . | . |
| | 435105 | . | . | . | . | . | . | . | . | . | . | . | . | . | . | T | . | . | . | . | . | A | . | . | . |
| | 435106 | ? | . | . | . | . | . | . | . | . | . | . | . | . | . | T | . | . | . | . | . | A | . | . | . |
| | 435108 | . | . | . | . | . | . | . | . | . | . | . | . | . | . | . | . | . | . | . | . | . | . | . | . |
| | 435109 | ? | . | . | . | . | . | . | . | . | . | . | . | . | . | . | . | . | . | . | . | . | . | . | . |
| | 435110 | ? | . | . | . | . | . | . | . | . | . | . | . | . | . | . | . | . | . | . | . | . | . | . | . |
| | 435111 | . | . | . | . | . | . | . | . | . | . | . | . | . | . | . | . | . | . | . | . | . | . | . | . |
| | 435112 | . | . | . | . | . | . | . | . | . | c | . | . | . | . | . | . | . | . | . | . | . | . | . | ? |
| | 436413 | ? | . | . | . | . | . | . | . | T | . | . | . | . | . | . | . | . | . | . | . | . | . | . | . |
| **A1.7** | 436414 | . | . | . | . | . | . | . | . | . | . | . | . | . | . | . | . | . | . | . | . | . | . | . | . |
| | 436415 | . | . | . | . | . | . | . | . | . | . | . | . | . | . | . | . | . | . | . | . | . | . | . | . |
| | 436417 | a | . | . | . | . | . | . | . | . | . | . | . | . | . | . | . | . | . | . | . | . | . | . | . |
| | 436418 | ? | . | . | . | . | . | . | . | . | . | . | . | . | . | . | . | . | . | . | . | . | . | . | . |
| | 436419 | ? | . | . | . | . | . | . | . | . | . | . | . | . | . | . | . | . | . | . | . | . | . | . | . |
| | 436420 | . | . | . | . | . | . | . | . | . | . | . | . | . | . | . | . | . | . | . | . | . | . | . | . |
| | 436421 | ? | . | . | . | . | . | . | . | . | . | . | . | . | . | . | . | . | . | . | . | . | . | . | . |
| | 436422 | . | . | . | . | . | . | . | . | . | . | . | . | . | . | . | . | . | . | . | . | . | . | . | . |
| | 436424 | . | . | . | . | . | . | . | . | . | . | . | . | . | . | . | . | . | . | . | . | . | . | . | . |

| | | | | | | | | | | | | | | | | | | | | | | | | | |
|---|---|---|---|---|---|---|---|---|---|---|---|---|---|---|---|---|---|---|---|---|---|---|---|---|---|
| | 436425 | a | . | . | . | . | . | . | . | . | . | . | . | . | . | . | . | . | . | . | . | . | . | . | T |
| **A1.8** | 436426 | . | . | . | . | . | . | . | . | . | . | . | . | . | . | . | . | . | . | . | . | . | . | . | . |
| | 436428 | ? | . | . | . | . | . | . | . | . | . | . | . | . | . | . | . | . | . | . | . | . | . | . | . |
| | 436429 | ? | . | . | . | . | . | . | . | . | . | . | . | . | . | . | . | . | . | . | . | . | . | . | . |
| | 436430 | . | . | . | . | . | . | . | . | . | . | . | . | . | . | . | . | . | . | . | . | . | . | . | T |
| | 436431 | . | . | . | . | . | . | . | . | . | . | . | . | . | . | . | Y | . | . | . | . | . | . | . | Y |
| | 436432 | . | . | . | . | . | . | . | . | . | . | . | . | . | . | . | . | . | . | . | . | . | . | . | . |
| | 436433 | a | . | . | . | . | . | . | . | . | . | . | . | . | . | . | . | . | . | . | . | . | . | . | . |
| | 436434 | . | . | . | . | . | . | . | . | . | . | . | . | . | . | . | . | . | . | . | . | . | . | . | . |
| | 436435 | . | . | . | . | . | . | . | . | . | . | . | . | . | . | . | . | . | . | . | . | . | . | . | . |
| | 436436 | ? | . | . | . | . | . | . | . | . | . | . | . | . | . | . | . | . | . | . | . | . | . | . | Y |
| | 436437 | ? | . | . | . | . | . | . | . | . | . | . | . | . | . | . | . | . | . | . | Y | . | . | . | . |
| | 436440 | a | . | . | . | . | . | . | . | . | . | . | . | . | . | . | . | . | . | . | . | . | . | . | . |
| | 436444 | . | . | . | . | . | . | . | Y | . | . | . | . | . | . | . | . | . | . | . | . | . | . | . | . |
| | 436445 | . | . | . | . | . | . | . | . | . | . | . | . | . | . | . | . | . | . | . | . | . | . | . | Y |
| | 436447 | . | . | . | . | . | . | . | . | . | . | . | . | . | . | . | . | . | . | . | . | . | . | . | . |
| | 436448 | . | . | . | . | . | . | . | . | . | . | . | . | . | . | . | . | . | . | . | . | . | . | . | . |
| | 436449 | . | . | . | . | . | . | . | . | . | . | . | . | . | . | . | . | . | . | . | . | . | . | . | . |
| | 436450 | . | . | . | . | . | . | . | . | . | . | . | . | . | . | . | . | . | . | . | . | . | . | . | . |
| | 436451 | . | . | . | . | . | . | . | . | . | . | . | . | . | . | . | . | . | . | . | . | . | . | . | . |
| | 436452 | . | . | . | . | . | . | . | . | . | . | . | . | . | . | . | . | . | . | . | . | . | . | . | . |
| | 436453 | . | . | . | . | . | . | . | . | T | . | . | . | . | . | . | . | . | . | . | . | . | . | . | . |
| | 436454 | . | . | . | . | . | . | . | . | T | . | . | . | . | . | . | . | . | . | . | . | . | . | . | . |
| | 436455 | . | . | . | . | . | . | . | . | T | . | c | . | . | . | . | . | . | . | . | . | . | . | . | . |
| | 436456 | . | . | . | . | . | . | . | . | T | . | . | . | . | . | . | . | . | . | . | . | . | . | . | . |
| | 436457 | a | . | . | . | . | . | . | . | T | . | . | . | . | . | . | . | . | . | . | . | . | . | . | . |
| | 436458 | ? | . | . | . | . | . | . | . | ? | . | . | . | . | . | . | . | . | . | . | . | . | . | . | . |
| | 436459 | . | . | . | . | . | . | . | . | T | . | . | . | . | . | . | . | . | . | . | . | . | . | . | . |
| | 436460 | . | . | . | . | . | . | . | . | T | . | . | . | . | . | . | . | . | . | . | . | . | . | . | . |
| | 436461 | a | . | . | . | . | . | . | . | T | . | . | . | . | . | . | . | . | . | . | . | . | . | . | . |
| | 436462 | a | . | . | . | . | . | . | . | T | . | . | . | . | . | . | . | . | . | . | . | . | . | . | . |
| | 436463 | ? | . | . | . | . | . | . | . | T | . | . | . | . | . | . | . | . | M | ? | . | . | . | . | . |
| | 437438 | ? | . | . | . | . | . | . | . | . | . | . | . | . | . | . | . | . | . | . | . | . | . | . | . |
| | 437440 | ? | . | . | . | . | . | . | . | T | . | . | . | . | . | . | . | . | . | . | . | . | . | . | . |
| | 437441 | ? | . | . | . | . | . | . | . | . | . | . | . | . | . | . | . | . | . | . | . | . | . | . | ? |
| **A1.24** | 437442 | ? | . | . | . | . | . | . | . | . | . | . | . | . | . | . | . | . | . | . | . | . | . | . | ? |
| | 437444 | ? | . | . | . | . | . | . | . | . | . | . | . | . | . | . | . | . | . | . | . | . | . | . | . |
| | 437445 | ? | . | . | . | . | . | . | . | T | . | . | . | . | . | . | . | . | . | . | . | . | . | . | . |
| | 437446 | ? | . | . | . | . | . | . | . | T | . | . | . | . | . | . | . | . | . | . | . | . | . | . | . |
| **A1.9** | 437447 | ? | . | . | . | . | . | . | . | T | . | . | . | . | . | . | . | . | . | . | . | . | . | . | . |
| | 437448 | ? | . | . | . | . | . | . | . | T | . | . | . | . | . | . | . | . | . | . | . | . | . | . | . |
| **A1.6.1.5** | 437449 | ? | . | . | . | . | . | . | . | T | . | . | . | . | . | . | . | . | . | . | . | . | . | . | . |
| | 437450 | ? | . | . | . | . | . | . | . | T | . | . | . | . | . | . | . | . | . | . | . | . | . | . | ? |
| | 437451 | ? | . | . | . | . | . | . | . | T | . | . | . | . | . | . | . | . | . | . | . | . | . | . | ? |
| | 437452 | ? | . | . | . | . | . | . | . | T | . | . | . | . | . | . | . | . | . | . | . | . | . | . | . |
| | 437453 | ? | . | . | . | . | . | . | . | . | . | . | . | . | . | . | . | . | . | . | . | . | . | . | . |
| | 437454 | ? | . | . | . | . | . | . | . | T | . | . | . | . | . | . | . | . | . | . | . | . | . | . | . |
| | 437539 | ? | . | . | . | . | . | . | . | T | . | . | . | . | . | . | . | . | . | . | . | . | . | . | . |
| | 437626 | . | . | . | . | . | . | . | . | T | . | . | . | . | . | . | . | . | . | . | . | . | . | . | . |
| | 438138 | . | . | . | . | . | . | . | . | . | . | . | . | . | . | . | . | . | . | . | . | . | . | . | . |
| | 438139 | . | . | . | . | . | . | . | . | . | . | . | . | . | . | . | . | . | . | . | . | . | . | . | . |
| **A1.6.1.2** | 444456 | ? | . | . | . | . | . | . | . | T | . | . | . | . | . | . | . | . | . | . | . | . | . | . | . |
| | 444457 | ? | . | . | . | . | . | . | . | T | . | . | . | . | . | . | . | . | . | . | . | . | . | . | . |
| | 444458 | ? | . | . | . | . | . | . | . | T | . | . | . | . | . | . | . | . | . | . | . | . | . | . | . |
| | 444459 | . | . | . | . | . | . | . | . | T | . | . | . | . | . | . | . | . | . | . | . | . | . | . | . |
| | 444460 | . | . | . | . | . | . | . | . | T | . | . | . | . | . | . | . | . | . | . | . | . | . | . | . |
| **A1.6.1.1.1** | 444461 | ? | . | . | . | . | . | . | . | T | . | . | . | . | . | . | . | . | . | . | . | . | . | . | . |
| | 444462 | ? | . | . | . | . | . | . | . | T | . | . | . | . | . | . | . | . | . | . | . | . | . | . | . |
| | 444463 | ? | . | . | . | . | . | . | . | T | . | . | . | . | . | . | . | . | . | . | . | . | . | . | . |
| | 444464 | . | . | . | . | . | . | . | . | T | . | . | . | . | . | . | . | . | . | . | . | . | . | . | . |
| **A1.6.1.1.1.2** | 444465 | . | . | . | . | . | . | . | . | T | . | . | . | . | . | . | . | . | . | . | . | . | . | . | . |
| | 444466 | ? | . | . | . | . | . | . | . | T | . | . | . | . | . | . | . | . | . | . | . | . | . | . | . |

| | | | | | | | | | | | | | | | | | | | | | | | | | |
|---|---|---|---|---|---|---|---|---|---|---|---|---|---|---|---|---|---|---|---|---|---|---|---|---|---|
| | 444467 | . | . | . | . | . | . | . | . | T | . | . | . | . | . | . | . | . | . | . | . | . | . | . | . |
| | 444468 | ? | . | . | . | . | . | . | . | T | . | . | . | . | . | . | . | . | . | . | . | . | . | . | . |
| **A1.6.1.1.2** | 444469 | ? | . | . | . | . | . | . | . | T | . | . | . | . | . | . | . | . | . | . | . | . | . | . | . |
| | 444470 | . | . | . | . | . | . | . | . | T | . | . | . | . | . | . | . | . | . | . | . | . | . | . | . |
| **A1.6.1** | 444471 | ? | . | . | . | . | . | . | . | T | . | . | . | . | . | . | . | . | . | . | . | . | . | . | . |
| | 444472 | ? | . | . | . | . | . | . | . | T | . | . | . | . | . | . | . | . | . | . | . | . | . | . | . |
| | 444473 | . | . | . | . | . | . | . | . | T | . | . | . | . | . | . | . | . | . | . | . | . | . | . | . |
| | 444474 | ? | . | . | . | . | . | . | . | T | . | . | . | . | . | . | . | . | . | . | . | . | . | . | . |
| | 444475 | ? | . | . | . | . | . | . | . | T | . | . | . | . | . | . | . | . | . | . | . | . | . | . | . |
| | 444476 | ? | . | . | . | . | . | . | . | T | . | . | . | . | . | . | . | . | . | . | . | . | . | . | . |
| | 444477 | ? | . | . | . | . | . | . | . | T | . | . | . | . | . | . | . | . | . | . | . | . | . | . | . |
| | 444478 | ? | . | . | . | . | . | . | . | T | . | . | . | . | . | . | . | . | . | . | . | . | . | . | . |
| **A1.10** | 444479 | ? | . | . | . | . | . | . | . | T | . | . | . | . | . | . | . | . | . | . | . | . | . | . | . |
| | 444480 | ? | . | . | . | . | . | . | . | T | . | . | . | . | . | . | . | . | . | . | . | . | . | . | . |
| | 444481 | ? | . | . | . | . | . | . | . | T | . | . | . | . | . | . | . | . | . | . | . | . | . | . | . |
| | 444482 | ? | . | . | . | . | . | . | . | T | . | . | . | . | . | . | . | . | . | . | . | . | . | . | . |
| **A1.11** | 444483 | . | . | . | . | . | . | . | . | T | . | . | . | . | . | . | . | . | . | . | . | . | . | . | . |
| | 444484 | . | . | . | . | . | . | . | . | T | . | . | . | . | . | . | . | . | . | . | . | . | . | . | . |
| | 444485 | ? | . | . | . | . | . | . | . | T | . | . | . | . | . | . | . | . | . | . | . | . | . | . | . |
| | 444486 | ? | . | . | . | . | . | . | . | T | . | . | . | . | . | . | . | . | . | . | . | . | . | . | . |
| | 447030 | ? | . | . | . | . | . | . | . | T | . | . | . | . | . | . | . | . | . | . | . | . | . | . | . |
| | 447031 | ? | . | . | . | . | . | . | . | T | . | . | . | . | . | . | . | . | . | . | . | . | . | . | . |
| | 447032 | ? | . | . | . | . | . | . | . | T | . | . | . | . | . | . | . | . | . | . | . | . | . | . | . |
| **A1.6.1.1.1.3** | 447033 | ? | . | . | . | . | . | . | . | T | . | . | . | . | . | . | . | . | . | . | . | . | . | . | . |
| | 447034 | ? | . | . | . | . | . | . | . | T | . | . | . | . | . | . | . | . | . | . | . | . | . | . | . |
| | 447035 | ? | . | . | . | . | . | . | . | T | . | . | . | . | . | . | . | . | . | . | . | . | . | . | . |
| | 447036 | ? | . | . | . | . | . | . | . | T | . | . | . | . | . | . | . | . | . | . | . | . | . | . | . |
| | 447037 | ? | . | . | . | . | . | . | . | T | . | . | . | . | . | . | . | . | . | . | . | . | . | . | . |
| | 447038 | ? | . | . | . | . | . | . | . | T | . | . | . | . | . | . | . | . | T | . | . | . | . | . | . |
| | 447039 | ? | . | . | . | . | . | . | . | T | . | . | . | . | . | . | . | . | . | . | . | . | . | . | . |
| | 447040 | ? | . | . | . | . | . | . | . | T | . | . | . | . | . | . | . | . | . | . | . | . | . | . | . |
| | 447041 | ? | . | . | . | . | . | . | . | T | . | . | . | . | . | . | . | . | . | . | . | . | . | . | . |
| | 447042 | ? | . | . | . | . | . | . | . | T | . | . | . | . | . | . | . | . | . | . | . | . | . | . | . |
| | 447043 | ? | . | . | . | . | . | . | . | T | . | . | . | . | . | . | . | . | . | . | . | . | . | . | . |
| | 447044 | ? | . | . | . | . | . | . | . | T | . | . | . | . | . | . | . | . | . | . | . | . | . | . | . |
| | 447045 | ? | . | . | . | . | . | . | . | T | . | . | . | . | . | . | . | . | . | . | . | . | . | . | . |
| | 447046 | ? | . | . | . | . | . | . | . | T | . | . | . | . | . | . | . | . | . | . | . | . | . | . | . |
| | 447047 | ? | . | . | . | . | . | . | . | T | . | . | . | . | . | . | . | . | . | . | . | . | . | . | . |
| | 447048 | ? | . | . | . | . | . | . | . | T | . | . | . | . | . | . | . | . | . | . | . | . | . | . | . |
| | 447049 | ? | . | . | . | . | . | . | . | T | . | . | . | . | . | . | . | . | . | . | . | . | . | . | . |
| | 447050 | ? | . | . | . | . | . | . | . | T | . | . | . | . | . | . | . | . | . | . | . | . | . | . | . |
| | 447051 | ? | . | . | . | . | . | . | . | T | . | . | . | . | . | . | . | . | . | . | . | . | . | . | . |
| | 447052 | ? | . | . | . | . | . | . | . | T | . | . | . | . | . | . | . | . | . | . | . | . | . | . | . |
| | 447053 | ? | . | . | . | . | . | . | . | T | . | . | . | . | . | . | . | . | . | . | . | . | . | . | . |
| | 447534 | ? | . | . | . | . | . | . | . | T | . | . | . | . | . | . | . | . | . | . | . | . | . | . | . |
| | 447535 | ? | . | . | . | . | . | . | . | T | . | . | . | . | . | . | . | . | . | . | . | . | . | . | . |
| | 447536 | ? | . | . | . | . | . | . | . | T | . | . | . | . | . | . | . | . | . | . | . | . | . | . | . |
| | 447537 | ? | . | . | . | . | . | . | . | T | . | . | . | . | . | . | . | . | . | . | . | . | . | . | . |
| | 447538 | ? | . | . | . | . | . | . | . | T | . | . | . | . | . | . | . | . | . | . | . | . | . | . | . |
| | 447539 | ? | . | . | . | . | . | . | . | T | . | . | . | . | . | . | . | . | . | . | . | . | . | . | . |
| | 447540 | ? | . | . | . | . | . | . | . | T | . | . | . | . | . | . | . | . | . | . | . | . | . | . | . |
| | 447541 | ? | . | . | . | . | . | . | . | T | . | . | . | . | . | . | . | . | . | . | . | . | . | . | . |
| | 447542 | ? | . | . | . | . | . | . | . | T | . | . | . | . | . | . | . | . | . | . | . | . | . | . | . |
| | 447543 | ? | . | . | . | . | . | . | . | T | . | . | . | . | . | . | . | . | . | . | . | . | . | . | . |
| | 447544 | ? | . | . | . | . | . | . | . | T | . | . | . | . | . | . | . | . | . | . | . | . | . | . | . |
| | 447545 | ? | . | . | . | . | . | . | . | T | . | . | . | . | . | . | . | . | . | . | . | . | . | . | . |
| | 447546 | ? | . | . | . | . | . | . | . | T | . | . | . | . | . | . | . | . | . | . | . | . | . | . | . |
| | 447547 | ? | . | . | . | . | . | . | . | T | . | . | . | . | . | . | . | . | . | . | . | . | . | . | . |
| | 447548 | ? | . | . | . | . | . | . | . | T | . | . | . | . | . | . | . | . | . | . | . | . | . | . | . |
| **A1.9.1** | 447549 | ? | . | . | . | . | T | . | . | T | . | . | . | . | . | . | . | . | . | . | . | . | . | . | . |
| | 447550 | ? | . | . | . | . | . | . | . | T | . | . | . | . | . | . | . | . | . | . | . | . | . | . | . |
| | 447551 | ? | . | . | . | . | T | . | . | T | . | . | . | . | . | . | . | . | . | . | . | . | . | . | . |

| | | | | | | | | | | | | | | | | | | | | | | | | | |
|---|---|---|---|---|---|---|---|---|---|---|---|---|---|---|---|---|---|---|---|---|---|---|---|---|---|
| | 447552 | ? | . | . | . | . | . | . | . | T | . | . | . | . | . | . | . | . | . | . | . | . | . | . | . |
| | 447553 | ? | . | . | . | . | . | . | . | T | . | . | . | . | . | . | . | . | . | . | . | . | . | . | . |
| | 447554 | ? | . | . | . | . | . | . | . | T | . | . | . | . | . | . | . | . | . | . | . | . | . | . | . |
| | 447555 | ? | . | . | . | . | . | . | . | T | . | . | . | . | . | . | . | . | . | . | . | . | . | . | . |
| | 447556 | . | . | . | . | . | . | . | . | . | . | . | . | . | . | . | . | . | . | . | . | . | . | . | . |
| | 447557 | - | . | . | . | . | . | . | . | . | . | . | . | . | . | . | . | . | . | . | . | . | . | . | . |
| | 447558 | - | . | . | . | . | . | . | . | . | . | . | . | . | . | . | . | . | . | . | . | . | . | . | . |
| | 447559 | . | . | . | . | . | . | . | . | . | . | . | . | . | . | . | . | . | . | . | T | . | . | . | . |
| | 447560 | - | . | . | . | . | . | . | . | . | . | . | . | . | . | . | . | . | . | . | . | . | . | . | . |
| | 447561 | - | . | . | . | . | . | . | . | . | . | . | . | . | . | . | . | . | . | . | . | . | . | . | T |
| | 447562 | - | . | . | . | . | . | . | . | . | . | . | . | . | . | . | . | . | . | . | . | . | . | . | . |
| | 447563 | - | . | . | . | . | . | . | . | . | . | . | . | . | . | . | . | . | . | . | . | . | . | . | . |
| | 447564 | . | . | . | . | . | . | . | . | . | . | . | . | . | . | . | . | . | . | . | . | . | . | . | . |
| | 447565 | - | . | . | . | . | . | . | . | . | . | . | . | . | . | . | . | . | . | . | . | . | . | . | . |
| | 447566 | - | . | . | . | . | . | . | . | . | . | . | . | . | . | . | . | . | . | . | . | . | . | . | . |
| | 447567 | - | . | . | . | . | . | . | . | . | . | . | . | . | . | . | . | . | . | . | . | . | . | . | . |
| | 447568 | - | . | . | . | . | . | . | . | T | . | . | . | . | . | . | . | . | . | . | . | . | . | . | . |
| | 447569 | - | . | . | . | . | . | . | . | . | . | . | . | . | . | . | . | . | . | . | . | . | . | . | . |
| | 447570 | - | . | . | . | . | . | . | . | . | . | . | . | . | . | . | . | . | . | . | . | . | . | . | . |
| | 447571 | - | . | . | . | . | . | . | . | . | . | . | . | . | . | . | . | . | . | . | . | . | . | . | . |
| | 447572 | - | . | . | . | . | . | . | . | . | . | T | . | . | . | . | . | . | . | . | . | . | . | . | . |
| | 447573 | - | C | T | . | . | . | . | . | . | . | . | . | . | . | . | . | . | . | . | . | . | . | . | . |
| | 447574 | - | . | . | . | . | . | . | . | . | . | . | . | . | . | . | . | . | . | . | . | . | . | . | . |
| | 447575 | . | . | . | . | . | . | . | . | . | . | . | . | . | . | . | . | . | . | . | . | . | . | . | . |
| | 447576 | - | . | . | . | . | . | . | . | . | . | . | . | . | . | . | . | . | . | . | . | . | . | . | . |
| | 447577 | - | . | . | . | . | . | . | . | . | . | . | . | . | . | . | . | . | . | . | . | . | . | . | . |
| | 447578 | - | . | . | . | . | . | . | . | . | . | . | . | . | . | . | . | . | . | . | . | . | . | . | . |
| | 447579 | - | C | T | . | . | . | . | . | . | . | . | . | . | . | . | . | . | . | . | . | . | . | . | . |
| | 447580 | - | . | . | . | . | . | . | . | . | . | . | . | . | . | . | . | . | . | . | . | . | . | . | . |
| | 447581 | - | . | . | . | . | . | . | . | . | . | . | . | . | . | . | . | . | . | . | . | . | . | . | T |
| | 447582 | - | . | . | . | . | . | . | . | . | . | . | . | . | . | . | . | . | . | . | . | . | . | . | . |
| | 447583 | - | . | . | . | . | . | . | . | . | . | . | . | . | . | . | . | . | . | . | . | . | . | . | . |
| **A1.12** | 447584 | - | . | . | . | . | . | . | . | . | . | . | . | . | . | . | . | . | . | . | . | . | . | . | . |
| | 447585 | - | . | . | . | . | . | . | . | . | . | . | . | . | . | . | . | . | . | . | . | . | . | . | . |
| | 447586 | - | . | . | . | . | . | . | . | . | . | . | . | . | . | . | . | . | . | . | . | . | . | . | . |
| | 447587 | - | . | . | . | . | . | . | . | T | . | . | T | . | . | . | . | . | . | . | . | . | . | . | . |
| | 447847 | - | . | . | . | . | . | . | . | . | . | . | . | . | . | . | . | . | . | . | . | . | . | . | . |
| | 447848 | - | . | . | . | . | . | . | . | . | . | . | . | . | . | . | . | . | . | . | . | . | . | . | . |
| | 447849 | - | . | . | . | . | . | . | . | . | . | . | T | . | . | . | . | . | . | . | . | . | . | . | . |
| | 447850 | - | . | . | . | . | . | . | . | . | . | . | . | . | . | . | . | . | . | . | . | . | . | . | . |
| | 447851 | - | . | . | . | . | . | . | . | . | . | . | . | . | . | . | . | . | . | . | . | . | . | . | . |
| | 447852 | - | . | . | . | . | . | . | . | . | . | . | . | . | . | . | . | . | . | . | T | . | . | . | . |
| | 447853 | - | . | . | . | . | . | . | . | . | . | . | . | . | . | . | . | . | . | . | . | . | . | . | . |
| | 447854 | - | . | . | . | . | . | . | . | . | . | . | . | . | . | . | . | . | . | . | . | . | . | . | . |
| | 447855 | - | . | . | . | . | . | . | . | . | . | . | . | . | . | . | . | . | . | . | . | . | . | . | T |
| | 447856 | - | . | . | . | . | . | . | . | . | . | . | . | . | . | . | . | . | . | . | . | . | . | . | . |
| | 447857 | - | . | . | . | . | . | . | . | . | . | . | . | . | . | . | . | . | . | . | . | . | . | . | . |
| **A1.25** | 447858 | - | . | . | . | . | . | . | . | . | . | . | . | . | . | . | . | . | . | . | . | . | . | . | . |
| | 447859 | . | . | . | . | . | . | . | . | . | . | . | . | . | . | . | . | . | . | . | . | . | . | . | . |
| | 447860 | - | . | . | . | . | . | . | . | . | . | . | . | . | . | . | . | . | . | . | . | . | . | . | . |
| | 447861 | - | . | . | . | . | . | . | . | . | . | . | . | . | . | . | . | . | . | . | . | . | . | . | . |
| | 447862 | . | . | . | . | . | . | . | . | . | . | . | . | . | . | . | . | . | . | . | . | . | . | . | . |
| | 447863 | - | . | . | . | . | . | . | . | . | . | . | . | . | . | . | . | . | . | . | . | . | . | . | . |
| | 447864 | - | . | . | . | . | . | . | . | . | . | . | . | . | . | . | . | . | . | . | . | . | . | . | . |
| | 447865 | - | . | . | . | . | . | . | . | . | . | . | . | . | . | . | . | . | . | . | . | . | . | . | . |
| | 447866 | - | . | . | . | . | . | . | . | . | . | T | . | . | . | . | . | . | . | . | . | . | . | . | . |
| | 450321 | - | . | . | . | . | . | . | . | . | . | . | . | . | . | . | . | . | . | . | . | . | . | . | . |
| | 450322 | - | . | . | . | . | . | . | . | . | . | . | . | . | . | . | . | . | . | . | . | . | . | . | . |
| | 450323 | - | . | . | . | . | . | . | . | T | . | . | . | . | . | . | . | . | . | . | . | . | . | . | . |
| | 450324 | - | . | . | . | . | . | . | . | . | . | . | . | . | . | . | . | . | . | . | . | . | . | . | . |
| | 450325 | - | . | . | . | . | . | . | . | T | . | . | . | . | . | . | . | . | . | . | . | . | . | . | . |
| | 450326 | - | . | . | . | . | . | . | . | . | . | . | . | . | . | . | . | . | . | . | . | . | . | . | . |

| | | | | | | | | | | | | | | | | | | | | | | | | | |
|---|---|---|---|---|---|---|---|---|---|---|---|---|---|---|---|---|---|---|---|---|---|---|---|---|---|
| | 450327 | . | . | . | . | . | . | . | . | . | . | . | . | . | . | . | . | . | . | . | . | . | . | . | . |
| | 450328 | - | . | . | . | . | . | . | . | . | . | . | . | . | . | . | . | . | . | . | . | . | . | . | . |
| | 450329 | - | . | . | . | . | . | . | . | . | . | . | . | . | . | . | . | . | . | . | . | . | . | . | . |
| | 450330 | - | . | . | . | . | . | . | . | . | . | . | . | . | . | . | . | . | . | . | . | . | . | . | . |
| | 450331 | - | . | . | . | . | . | . | . | . | . | . | . | . | . | . | . | . | . | . | . | . | . | . | . |
| | 450332 | . | . | . | . | . | . | . | . | . | . | . | . | . | . | . | . | . | . | . | T | . | . | . | . |
| | 450781 | ? | . | . | . | . | . | . | . | T | . | . | . | . | . | . | . | . | . | . | . | . | . | . | . |
| | 450782 | ? | . | . | . | . | . | . | . | T | . | . | . | . | . | . | . | . | . | . | . | . | . | . | . |
| | 450783 | ? | . | . | . | . | . | . | . | T | . | . | . | . | . | . | . | . | . | . | . | . | . | . | . |
| | 450784 | ? | . | . | . | . | . | . | . | . | . | . | . | . | . | . | . | . | . | . | . | . | . | . | . |
| **A1.14.1** | 450785 | ? | . | . | . | . | . | . | . | T | T | . | . | . | T | . | . | . | . | . | . | . | . | . | . |
| | 450786 | ? | . | . | . | . | . | . | . | T | T | . | . | . | T | . | . | . | . | . | . | . | . | . | . |
| | 450787 | . | . | . | . | . | . | . | . | . | . | . | . | . | . | . | . | . | . | . | . | . | . | . | . |
| | 450788 | ? | . | . | . | . | . | . | . | T | . | . | . | . | . | . | . | . | . | . | . | . | . | . | . |
| | 450789 | ? | . | . | . | . | . | . | . | T | . | . | . | . | . | . | . | . | . | . | . | . | . | . | . |
| **A1.26** | 450790 | ? | . | . | . | . | . | . | . | . | . | . | . | C | . | . | . | . | . | . | . | . | . | . | . |
| | 450791 | ? | . | . | . | . | . | . | . | . | . | . | . | C | . | . | . | . | . | . | . | . | . | . | . |
| **A1.6.1.1.1.4** | 451149 | ? | . | . | . | . | . | . | . | T | . | . | . | . | . | . | . | . | . | . | . | . | . | . | . |
| | 451150 | . | . | . | . | . | . | . | . | T | . | . | . | . | . | . | . | . | . | . | . | . | . | . | . |
| | 451151 | ? | . | . | . | . | . | . | . | T | . | . | . | . | . | . | . | . | . | . | . | . | . | . | . |
| | 451152 | ? | . | . | . | . | . | . | . | T | . | . | . | . | . | . | . | . | . | . | . | . | . | . | . |
| | 451153 | ? | . | . | . | . | . | . | . | T | . | . | . | . | . | . | . | . | . | . | . | . | . | . | . |
| **A1.27** | 451154 | ? | . | . | . | . | . | . | . | . | . | . | . | . | . | . | . | . | . | . | . | . | . | . | . |
| | 451155 | ? | . | . | . | . | . | . | . | T | . | . | . | . | . | . | . | . | . | . | . | . | . | . | . |
| | 451156 | ? | . | . | . | . | . | . | . | . | . | . | . | . | . | . | . | . | . | . | . | . | . | . | . |
| | 451157 | ? | . | . | . | . | . | . | . | T | . | . | . | . | . | . | . | . | . | . | . | . | . | . | . |
| **A1.13** | 451158 | ? | . | . | . | . | . | . | . | T | . | . | . | . | . | . | . | . | . | . | . | . | . | . | . |
| | 451159 | . | . | . | . | . | . | . | . | . | . | . | . | . | . | . | . | . | . | . | . | . | . | . | . |
| | 451160 | ? | . | . | . | . | . | . | . | T | . | . | . | . | . | . | . | . | . | . | . | . | . | . | . |
| | 451161 | ? | . | . | . | . | . | . | . | . | . | . | . | . | . | . | . | . | . | . | . | . | . | . | . |
| | 451162 | ? | . | . | . | . | . | . | . | T | . | . | . | . | . | . | . | . | . | . | . | . | . | . | . |
| | 451163 | ? | . | . | . | . | . | . | . | T | . | . | . | . | . | . | . | . | . | . | . | . | . | . | . |
| | 451666 | ? | . | . | . | . | . | . | . | . | . | . | . | . | . | . | . | . | . | . | T | . | . | . | . |
| **A1.14** | 452192 | - | a | g | g | c | c | g | c | t | t | c | c | t | c | c | c | t | c | c | c | g | g | g | c |
| | 452193 | - | a | g | g | c | c | g | c | t | t | c | c | t | c | c | c | t | c | c | c | g | g | g | c |
| | 452195 | - | a | g | g | c | c | g | c | t | c | c | c | t | c | c | c | t | c | c | c | g | g | g | c |
| | 452196 | - | a | g | g | c | c | g | c | t | c | c | c | t | c | c | c | t | c | c | c | g | g | g | c |
| | 452197 | - | a | g | g | c | c | g | c | t | c | c | c | t | c | c | c | t | c | c | c | g | g | g | c |
| | 452198 | - | a | g | g | c | c | g | c | t | t | c | c | t | c | c | c | t | c | c | c | g | g | g | c |
| | 452199 | - | a | g | g | c | c | g | c | t | t | c | c | t | c | c | c | t | c | c | c | g | g | g | c |
| | 452200 | - | a | g | g | c | c | g | c | t | t | c | c | t | c | c | c | t | c | c | c | g | g | g | c |
| | 452201 | - | a | g | g | c | c | g | c | t | t | c | c | t | c | c | c | t | c | c | c | g | g | g | c |
| | 452202 | - | a | g | g | c | c | g | c | c | c | c | c | t | c | c | c | t | c | c | c | g | g | g | c |
| | 452203 | - | a | g | g | c | c | g | c | c | c | c | c | t | c | c | c | t | c | c | c | g | g | g | c |
| **A1.28** | 452204 | - | a | g | g | c | c | g | c | c | c | c | c | t | c | c | c | t | c | c | c | g | g | g | c |
| **A1.29** | 452205 | - | a | g | g | c | c | g | c | c | c | c | c | t | c | c | c | t | c | c | c | g | g | g | c |
| | 452207 | - | a | g | g | c | c | g | c | t | t | c | c | t | c | c | c | t | c | c | c | g | g | g | c |
| | 452208 | - | a | g | g | c | c | g | c | c | c | c | c | t | c | c | c | t | c | c | c | g | g | g | c |
| | 452209 | - | a | g | g | c | c | g | c | c | c | c | c | t | c | c | c | t | c | c | c | g | g | g | c |
| | 452210 | - | a | g | g | c | c | g | c | t | t | c | c | t | c | c | c | t | c | c | c | g | g | g | c |
| | 452211 | - | a | g | g | c | c | g | c | t | t | c | c | t | c | c | c | t | c | c | c | g | g | g | c |
| | 452212 | - | a | g | g | c | c | g | c | t | t | c | c | t | c | c | c | t | c | c | c | g | g | g | c |
| **A1.15** | 452213 | - | a | g | g | c | c | g | c | c | c | c | c | t | c | c | c | t | c | c | c | a | g | g | c |
| **A1.16** | 452214 | - | a | g | g | c | c | g | c | t | c | c | c | t | c | c | c | t | c | c | c | g | g | g | c |
| | 452215 | - | a | g | g | c | c | g | c | c | c | c | c | t | c | c | c | t | c | c | c | g | g | g | c |
| | 452216 | - | a | g | g | c | c | g | c | t | t | c | c | t | c | c | c | t | c | c | c | g | g | g | c |
| | 452217 | - | a | g | g | c | c | g | c | t | c | c | c | t | c | c | c | t | c | c | c | g | g | g | c |
| | 452790 | - | . | . | . | . | . | . | . | T | . | . | . | . | . | . | . | . | . | . | . | . | . | . | . |
| | 452791 | - | . | . | . | . | . | . | . | T | . | . | . | . | . | . | . | . | . | . | . | . | . | . | . |
| | 452792 | - | . | . | . | . | . | . | . | T | . | . | . | . | . | . | . | . | . | . | . | . | . | . | . |
| | 452793 | - | . | . | . | . | . | . | . | . | . | . | . | . | . | . | . | . | . | . | . | . | . | . | . |
| | 452794 | - | . | . | . | . | . | . | . | T | . | . | . | . | . | . | . | . | . | . | . | . | . | . | . |

| | | | | | | | | | | | | | | | | | | | | | | | | | |
|---|---|---|---|---|---|---|---|---|---|---|---|---|---|---|---|---|---|---|---|---|---|---|---|---|---|
| | 452795 | . | . | . | . | . | . | . | . | . | . | . | . | . | . | . | . | . | . | . | . | . | . | . | . |
| | 454524 | ? | . | . | . | . | . | . | . | T | T | . | . | . | . | . | . | . | . | . | . | . | . | . | . |
| | 454525 | ? | . | . | . | . | . | . | . | . | . | . | . | . | . | . | . | . | . | . | . | A | . | . | . |
| | 454526 | ? | . | . | . | . | . | . | . | . | . | . | . | . | . | . | . | . | . | . | . | A | . | . | . |
| | 454527 | ? | . | . | . | . | . | . | . | T | . | . | . | . | . | . | . | . | . | . | . | . | . | . | . |
| | 454528 | ? | . | . | . | . | . | . | . | . | . | . | . | . | . | T | . | . | . | . | . | A | . | . | . |
| | 454529 | ? | . | . | . | . | . | . | . | T | . | . | . | . | . | . | . | . | . | . | . | . | . | . | . |
| | 454531 | ? | . | . | . | . | . | . | . | . | . | . | . | . | . | . | . | . | . | . | . | . | . | . | . |
| | 454532 | ? | . | . | . | . | . | . | . | T | . | . | . | . | . | . | . | . | . | . | . | . | . | . | . |
| | 454533 | ? | . | . | . | . | . | . | . | T | . | . | . | . | . | . | . | . | . | . | . | . | . | . | . |
| | 454534 | ? | . | . | . | . | . | . | . | . | . | . | . | . | . | . | . | . | . | . | . | . | . | . | . |
| | 454536 | ? | . | . | . | . | . | . | . | . | . | . | . | . | . | . | . | . | . | . | . | . | . | . | . |
| | 454537 | ? | . | . | . | . | . | . | . | . | . | . | . | . | . | . | . | . | . | . | . | . | . | . | . |
| | 454540 | ? | . | . | . | . | . | . | . | . | . | . | . | . | . | . | . | . | . | . | . | . | . | . | . |
| | 454542 | ? | . | . | . | . | . | . | . | . | . | . | . | . | . | . | . | . | . | . | . | . | . | . | . |
| | 454543 | ? | . | . | . | . | . | . | . | . | . | . | . | . | . | . | . | . | . | . | . | . | . | . | T |
| | 454544 | ? | . | . | . | . | . | . | . | . | . | . | . | . | . | . | . | . | . | . | . | . | . | . | . |
| | 454546 | ? | . | . | . | . | . | . | . | . | . | . | . | . | . | . | . | . | . | . | . | . | . | . | . |
| | 454547 | ? | . | . | . | . | . | . | . | . | . | . | . | . | . | . | . | . | . | . | . | . | . | . | . |
| | 454549 | ? | . | . | . | . | . | . | . | . | . | . | . | . | . | . | . | . | . | . | . | . | . | . | . |
| | 454551 | ? | . | . | . | . | . | . | . | . | . | . | . | . | . | . | . | . | . | . | . | . | . | . | . |
| | 454552 | ? | . | . | . | . | . | . | . | . | . | . | . | . | . | . | . | . | . | . | . | . | . | . | . |
| | 454557 | ? | . | . | . | . | . | . | . | T | T | . | . | . | . | . | . | . | . | . | . | . | . | . | . |
| | 454560 | ? | . | . | . | . | . | . | . | T | . | . | . | . | . | . | . | . | . | . | . | . | . | . | . |
| | 454563 | ? | . | . | . | . | . | . | . | T | T | . | . | . | T | . | . | . | . | . | . | . | . | . | . |
| | 454564 | ? | . | . | . | . | . | . | . | T | T | . | . | . | . | . | . | . | . | . | . | . | . | . | . |
| | 454565 | ? | . | . | . | . | . | . | . | T | . | . | . | . | . | . | . | . | . | . | . | . | . | . | . |
| | 454566 | ? | . | . | . | . | . | . | . | T | . | . | . | . | . | . | . | . | . | . | . | . | . | . | . |
| | 454567 | ? | . | . | . | . | . | . | . | T | T | . | . | . | . | . | . | . | . | . | . | . | . | . | . |
| | 454568 | ? | . | . | . | . | . | . | . | . | . | . | . | . | . | . | . | . | . | . | T | . | . | . | . |
| | 454569 | ? | . | . | . | . | . | . | . | T | . | . | . | . | . | . | T | . | . | . | . | . | . | . | . |
| | 454570 | ? | . | . | . | . | . | . | . | T | . | . | . | . | . | . | . | . | . | . | . | . | . | . | . |
| | 454830 | . | . | . | . | . | . | . | . | . | . | . | . | . | . | . | . | . | . | . | . | . | . | . | . |
| | 454831 | . | . | . | . | . | . | . | . | T | . | . | . | . | . | . | . | . | . | . | . | . | . | . | . |
| | 454832 | . | . | . | . | . | . | . | . | . | . | . | . | . | . | . | . | . | . | . | . | . | . | . | . |
| | 454833 | . | . | . | . | . | . | . | . | . | . | . | . | . | . | . | . | . | . | . | . | . | . | . | . |
| | 454858 | ? | . | . | . | . | . | . | . | . | . | . | . | . | . | . | . | . | . | . | . | . | . | . | . |
| | 454862 | ? | . | . | . | . | . | . | . | . | . | . | . | . | . | . | . | . | . | . | . | . | . | . | . |
| | 454866 | ? | . | . | . | . | . | . | . | . | . | . | . | . | . | . | . | . | . | . | . | . | . | . | . |
| | 454867 | ? | . | . | . | . | . | . | . | . | . | . | . | . | . | . | . | . | . | . | . | . | . | . | . |
| | 455015 | ? | . | . | . | . | . | . | . | . | . | . | . | C | . | . | . | . | . | . | . | . | . | . | . |
| | 455016 | ? | . | . | . | . | . | . | . | . | . | . | . | C | . | . | . | . | . | . | . | . | . | . | . |
| | 455017 | ? | . | . | . | . | . | . | . | T | . | . | . | . | . | . | . | . | . | . | . | . | . | . | . |
| | 455018 | ? | . | . | . | . | . | . | . | T | . | . | . | . | . | . | . | . | . | . | . | . | . | . | . |
| | 455019 | ? | . | . | . | . | . | . | . | T | . | . | . | . | . | . | . | . | . | . | . | . | . | . | . |
| | 455020 | ? | . | . | . | . | . | . | . | T | . | . | . | . | . | . | . | . | . | . | . | . | . | . | . |
| **A1.6.1.4** | 455021 | ? | . | . | . | . | . | . | . | T | . | . | . | . | . | . | . | . | . | . | . | . | . | . | . |
| | 455022 | ? | . | . | . | . | . | . | . | T | . | . | . | . | . | . | . | . | . | . | . | . | . | . | . |
| | 455023 | ? | . | . | . | . | . | . | . | T | . | . | . | . | . | . | . | . | . | . | . | . | . | . | . |
| | 455024 | ? | . | . | . | . | . | . | . | T | . | . | . | . | . | . | . | . | . | . | . | . | . | . | . |
| | 455025 | ? | . | . | . | . | . | . | . | T | . | . | . | . | . | . | . | . | . | . | . | . | . | . | . |
| | 455026 | ? | . | . | . | . | . | . | . | T | . | . | . | . | . | . | . | . | . | . | . | . | . | . | . |
| | 455027 | . | . | . | . | . | . | . | . | T | . | . | . | . | . | . | . | . | . | . | . | . | . | . | . |
| | 455478 | ? | . | . | . | . | . | . | . | T | . | . | . | . | . | . | . | . | . | . | . | . | . | . | . |
| | 455640 | ? | . | . | . | . | . | . | . | T | . | . | . | . | . | . | . | . | . | . | . | . | . | . | . |
| | 455643 | . | . | . | . | . | . | . | . | T | . | . | . | . | . | . | . | . | . | . | . | . | . | . | . |
| | 455644 | . | . | . | . | . | . | . | . | T | . | . | . | . | . | . | . | . | . | . | . | . | . | . | . |
| | 455645 | . | . | . | . | . | . | . | . | . | . | . | . | . | . | . | . | . | . | . | . | . | . | . | . |
| | 455646 | . | . | . | . | . | . | . | . | T | . | . | . | . | . | . | . | . | . | . | . | . | . | . | . |
| | 455647 | . | . | . | . | . | . | . | . | T | . | . | . | . | . | . | . | . | . | . | . | . | . | . | . |
| | 455648 | . | . | . | . | . | . | . | . | T | . | . | . | . | . | . | . | . | . | . | . | . | . | . | . |
| | 455649 | . | . | . | . | . | . | . | . | . | . | . | . | . | . | . | . | . | . | . | . | . | . | . | . |

| | | | | | | | | | | | | | | | | | | | | | | | | | |
|---|---|---|---|---|---|---|---|---|---|---|---|---|---|---|---|---|---|---|---|---|---|---|---|---|---|
| | 455650 | . | . | . | . | . | . | . | . | T | . | . | . | . | . | . | . | . | . | . | . | . | . | . | . |
| | 455651 | . | . | . | . | . | . | . | . | T | . | . | . | . | . | . | . | . | . | . | . | . | . | . | . |
| | 455652 | . | . | . | . | . | . | . | . | T | . | . | . | . | . | . | . | . | . | . | . | . | . | . | . |
| **A1.30** | 455653 | . | . | . | T | . | . | . | . | T | . | . | . | . | . | . | . | . | . | . | . | . | . | . | . |
| | 455655 | . | . | . | . | . | . | . | . | T | . | . | . | . | . | . | . | . | . | . | . | . | . | . | . |
| | 455656 | . | . | . | . | . | . | . | . | . | . | . | . | . | . | . | . | . | . | . | . | . | . | . | . |
| | 455658 | . | . | . | . | . | . | . | . | . | . | . | . | . | . | . | . | . | T | . | . | . | . | . | . |
| | 455659 | . | . | . | . | . | . | . | . | T | . | . | . | . | . | . | . | . | . | . | . | . | . | . | . |
| **A1.17** | 455660 | . | . | . | . | . | . | . | . | T | . | . | . | . | . | . | . | . | . | . | . | . | A | . | . |
| | 455662 | . | . | . | . | . | . | . | . | T | . | . | . | . | . | . | . | . | . | . | . | . | . | . | . |
| | 455663 | . | . | . | . | . | . | . | . | T | . | . | . | . | . | . | . | . | . | . | . | . | . | . | . |
| **A1.20** | 455667 | . | . | . | . | T | . | . | . | T | . | . | . | . | . | . | . | . | . | . | . | . | . | . | . |
| | 455668 | . | . | . | . | . | . | . | . | T | . | . | . | . | . | . | . | . | . | . | . | . | . | . | . |
| | 455669 | . | . | . | . | . | . | . | . | T | . | . | . | . | . | . | . | . | . | . | . | . | . | . | . |
| | 455670 | . | . | . | . | . | . | . | . | T | . | . | . | . | . | . | . | . | T | . | . | . | . | . | . |
| | 455671 | . | . | . | . | . | . | . | . | . | . | . | . | . | . | . | . | . | . | . | . | . | . | . | . |
| | 455672 | . | . | . | . | . | . | . | . | T | . | . | . | . | . | . | . | . | . | . | . | . | . | . | . |
| | 455673 | . | . | . | . | T | . | . | . | T | . | . | . | . | . | . | . | . | . | . | . | . | . | . | . |
| | 455674 | . | . | . | . | T | . | . | . | T | . | . | . | . | . | . | . | . | . | . | . | . | . | . | . |
| | 455675 | . | . | . | T | . | . | . | . | T | . | . | . | . | . | . | . | . | . | . | . | . | . | . | . |
| | 455676 | . | . | . | . | T | . | . | . | T | . | . | . | . | . | . | . | . | . | . | . | . | . | . | . |
| | 455678 | . | . | . | . | . | . | . | . | T | . | . | . | . | . | . | . | . | . | . | . | . | A | . | . |
| | 455679 | . | . | . | . | . | . | . | . | T | . | . | . | . | . | . | . | . | . | . | . | . | . | . | . |
| | 455749 | ? | . | . | . | . | . | . | . | . | . | . | . | . | . | . | . | . | . | . | . | . | . | . | . |
| | 455751 | ? | . | . | . | . | . | . | . | . | . | . | . | . | . | . | . | . | . | . | . | . | . | . | . |
| | 455752 | ? | . | . | . | . | . | . | . | . | . | . | . | . | . | . | . | . | . | . | . | . | . | . | . |
| | 455754 | ? | . | . | . | . | . | . | . | T | . | . | . | . | . | . | . | . | . | . | . | . | . | . | . |
| | 455755 | ? | . | . | . | . | . | . | . | T | . | . | . | . | . | . | . | . | . | . | . | . | . | . | . |
| | 455757 | ? | . | . | . | . | . | . | . | T | . | . | . | . | . | . | . | . | . | . | . | . | . | . | . |
| | 455758 | ? | . | . | . | . | . | . | . | . | . | . | . | . | . | . | . | . | . | . | . | . | . | . | . |
| | 455760 | ? | . | . | . | . | . | . | . | . | . | . | . | . | . | . | . | . | . | . | . | . | . | . | . |
| | 455761 | . | . | . | . | . | . | . | . | . | . | . | . | . | . | . | . | . | . | . | . | . | . | . | . |
| | 455763 | . | . | . | . | . | . | . | . | . | . | . | . | . | . | . | . | . | . | . | . | . | . | . | . |
| **A1.31** | 455764 | . | . | . | . | . | . | . | . | . | . | . | . | . | . | . | . | . | . | . | . | . | . | . | . |
| | 455765 | ? | . | . | . | . | . | . | . | T | . | . | . | . | . | . | . | . | . | . | . | . | . | . | . |
| | 455766 | . | . | . | . | . | . | . | . | . | . | . | . | . | . | . | . | . | . | . | . | . | . | . | . |
| | 455767 | ? | . | . | . | . | . | . | . | . | . | . | . | . | . | . | . | . | . | . | . | . | . | . | . |
| | 455768 | ? | . | . | . | . | . | . | . | . | . | . | . | . | . | . | . | . | . | . | . | . | . | . | . |
| | 455770 | ? | . | . | . | . | . | . | . | . | . | . | . | . | . | . | . | . | . | . | . | . | . | . | . |
| | 455771 | ? | . | . | . | . | . | . | . | T | T | . | . | . | . | . | . | . | . | . | . | . | . | . | . |
| | 455775 | ? | . | . | . | . | . | . | . | T | . | . | . | . | . | . | . | . | . | . | . | . | . | . | . |
| | 455776 | . | . | . | . | . | . | . | . | T | . | . | . | . | . | . | . | . | . | . | . | . | . | . | . |
| | 455777 | ? | . | . | . | . | . | . | . | T | . | . | . | . | . | . | . | . | . | . | . | . | . | . | . |
| | 455778 | ? | . | . | . | . | . | . | . | T | . | . | . | . | . | . | . | . | . | . | . | . | . | . | . |
| | 455779 | . | . | . | . | . | . | . | . | T | . | . | . | . | . | . | . | . | . | . | . | . | . | . | . |
| | 455780 | ? | . | . | . | . | . | . | . | T | . | . | . | . | . | . | . | . | . | . | . | . | . | . | . |
| | 455782 | ? | . | . | . | . | . | . | . | T | . | . | . | . | . | . | . | . | . | . | . | . | . | . | . |
| | 455783 | ? | . | . | . | . | . | . | . | T | . | . | . | . | . | . | . | . | . | . | . | . | . | . | . |
| | 455784 | . | . | . | . | . | . | . | . | T | . | . | . | . | . | . | . | . | . | . | . | . | . | . | . |
| | 455786 | ? | . | . | . | . | . | . | . | T | . | . | . | . | . | . | . | . | . | . | . | . | . | . | . |
| | 455787 | ? | . | . | . | . | . | . | . | T | . | . | . | . | . | . | . | . | . | . | . | . | . | . | . |
| | 458030 | - | . | . | . | . | . | . | . | T | . | . | . | . | . | . | . | . | . | . | . | . | . | . | . |
| **A1.4.1** | 458031 | - | . | . | . | . | . | . | . | T | . | . | . | . | . | . | . | . | . | . | . | . | . | . | . |
| | 458032 | - | . | . | . | . | . | . | . | T | . | . | . | . | . | . | . | . | . | . | . | . | . | . | . |
| | 458033 | - | . | . | . | . | . | . | . | T | . | . | . | . | . | . | . | . | . | . | . | . | . | . | . |
| | 458034 | - | . | . | . | . | . | . | . | T | . | . | . | . | . | . | . | . | . | . | . | . | . | . | . |
| | 458035 | - | . | . | . | . | . | . | . | . | . | . | . | . | . | . | . | . | . | . | . | . | . | . | . |
| | 458036 | - | . | . | . | . | . | . | . | T | . | . | . | . | . | . | . | . | . | . | . | . | . | . | . |
| **A1.4.2** | 458037 | - | . | . | . | . | . | . | . | T | . | . | . | . | . | . | . | C | . | . | . | . | . | . | . |
| **A1.4.3** | 458038 | - | . | . | . | . | . | . | . | T | . | . | . | . | . | . | . | . | . | T | . | . | . | . | . |
| | 458039 | - | . | . | . | . | . | . | . | T | . | . | . | . | . | . | . | . | . | T | . | . | . | . | . |
| | 458040 | - | . | . | . | . | . | . | . | T | . | . | . | . | . | . | . | . | . | . | . | . | . | . | . |

| | | | | | | | | | | | | | | | | | | | | | | | | | |
|---|---|---|---|---|---|---|---|---|---|---|---|---|---|---|---|---|---|---|---|---|---|---|---|---|---|
| | 458041 | - | . | . | . | . | . | . | . | T | . | . | . | . | . | . | . | . | . | T | . | . | . | . | . |
| | 458042 | - | . | . | . | . | . | . | . | T | . | . | . | . | . | . | . | . | . | T | . | . | . | . | . |
| | 458043 | - | . | . | . | . | . | . | . | T | . | . | . | . | . | . | . | C | . | . | . | . | . | . | . |
| | 458044 | - | . | . | . | . | . | . | . | T | . | . | . | . | . | . | . | C | . | . | . | . | . | . | . |
| | 458045 | - | . | . | . | . | . | . | . | T | . | . | . | . | . | . | . | . | . | . | . | . | . | . | . |
| **A1.21** | 458046 | - | . | . | . | . | . | . | . | T | . | . | . | . | . | . | . | . | . | . | . | . | . | . | . |
| | 458047 | - | . | . | . | . | . | . | . | T | . | . | . | . | . | . | . | . | . | . | . | . | . | . | . |
| | 458048 | - | . | . | . | . | . | . | . | T | . | . | . | . | . | . | . | . | . | . | . | . | . | . | . |
| | 458049 | - | . | . | . | . | . | . | . | T | . | . | . | . | . | . | . | . | . | . | . | . | . | . | . |
| | 458050 | - | . | . | . | . | . | . | . | T | . | . | . | . | . | . | . | . | . | . | . | . | . | . | . |
| | 458051 | - | . | . | . | . | . | . | . | T | . | . | . | . | . | . | . | . | . | . | . | . | . | . | . |
| | 458052 | - | . | . | . | . | . | . | . | T | . | . | . | . | . | . | . | . | . | . | . | . | . | . | . |
| | 458053 | - | . | . | . | . | . | . | . | T | . | . | . | . | . | . | . | . | . | . | . | . | . | . | . |
| | 458054 | - | . | . | . | . | . | . | . | T | T | . | . | . | . | . | . | . | . | . | . | . | . | . | . |
| | 458055 | - | . | . | . | . | . | . | . | T | T | . | . | . | . | . | . | . | . | . | . | . | . | . | . |
| | 458056 | - | . | . | . | . | . | . | . | T | T | . | . | . | . | . | . | . | . | . | . | . | . | . | . |
| | 458057 | - | . | . | . | . | . | . | . | T | T | . | . | . | . | . | . | . | . | . | . | . | . | . | . |
| | 458058 | - | . | . | . | . | . | . | . | T | T | . | . | . | . | . | . | . | . | . | . | . | . | . | . |
| **A1.14.2** | 458059 | - | . | . | . | . | . | . | . | T | T | . | . | . | . | . | . | . | . | . | . | . | . | . | . |
| | 458060 | - | . | . | . | . | . | . | . | T | T | . | . | . | . | . | . | . | . | . | . | . | . | . | . |
| | 458061 | - | . | . | . | . | . | . | . | T | T | . | . | . | . | . | . | . | . | . | . | . | . | . | . |
| | 458062 | - | . | . | . | . | . | . | . | . | . | . | . | . | . | . | . | . | . | . | . | . | . | . | . |
| **A1.18** | 458063 | - | . | . | . | . | . | . | . | . | . | . | . | . | . | . | . | . | . | . | . | . | . | . | . |
| **A1.22** | 458064 | - | . | . | . | . | . | . | . | T | . | . | . | . | . | . | . | . | . | . | . | . | . | T | . |
| | 458065 | - | . | . | . | . | . | . | . | T | . | . | . | . | . | . | . | . | . | . | . | . | . | T | . |
| | 458066 | - | . | . | . | . | . | . | . | . | . | . | . | . | . | . | . | . | . | . | . | . | . | . | . |
| | 458067 | - | . | . | . | . | . | . | . | . | . | . | . | . | . | . | . | . | . | . | . | . | . | . | . |
| | 458068 | - | . | . | . | . | . | . | . | . | . | . | . | . | . | . | . | . | . | . | . | . | . | . | . |
| | 458069 | - | . | . | . | . | . | . | . | . | . | . | . | . | . | . | . | . | . | . | . | . | . | . | . |
| | 458070 | - | . | . | . | . | . | . | . | . | . | . | . | . | . | . | . | . | . | . | . | . | . | . | . |
| | 458071 | - | . | . | . | . | . | . | . | . | . | . | . | . | . | . | . | . | . | . | . | . | . | . | . |
| | 458072 | - | . | . | . | . | . | . | . | . | . | . | . | . | . | . | . | . | . | . | . | . | . | . | . |
| | 458073 | - | . | . | . | . | . | . | . | . | . | . | . | . | . | . | . | . | . | . | . | . | . | . | . |
| | 458074 | - | . | . | . | . | . | . | . | . | . | . | . | . | . | . | . | . | . | . | . | . | . | . | . |
| | 458075 | - | . | . | . | . | . | . | . | . | . | . | . | . | . | . | . | . | . | . | . | . | . | . | . |
| | 458076 | - | . | . | . | . | . | . | . | . | . | . | . | . | . | . | . | . | . | . | . | . | . | . | . |
| | 458077 | - | . | . | . | . | . | . | . | . | . | . | . | . | . | . | . | . | . | . | . | . | . | . | . |
| | 458080 | - | . | . | . | . | . | . | . | . | . | . | . | . | . | . | . | . | . | . | . | . | . | . | . |
| | 458086 | ? | . | . | . | . | . | . | . | T | . | . | . | . | . | . | . | . | . | . | . | . | . | . | . |
| | 458087 | ? | . | . | . | . | . | . | . | T | . | . | . | . | . | . | . | . | . | . | . | . | . | . | . |
| | 458088 | ? | . | . | . | . | . | . | . | T | . | . | . | . | . | . | . | . | . | . | . | . | . | . | . |
| | 458089 | ? | . | . | . | . | . | . | . | T | . | . | . | . | . | . | . | . | . | . | . | . | . | . | . |
| | 458090 | ? | . | . | . | . | . | . | . | T | . | . | . | . | . | . | . | . | . | . | . | . | . | . | . |
| | 458091 | ? | . | . | . | . | . | . | . | T | . | . | . | . | . | . | . | . | . | . | . | . | . | . | . |
| | 458092 | ? | . | . | . | . | . | . | . | T | . | . | . | . | . | . | . | . | . | . | . | . | . | . | . |
| | 458093 | ? | . | . | . | . | . | . | . | T | . | . | . | . | . | . | . | . | . | . | . | . | . | . | . |
| | 458094 | ? | . | . | . | . | . | . | . | T | . | . | . | . | . | . | . | . | . | . | . | . | . | . | . |
| | 458095 | ? | . | . | . | . | . | . | . | T | . | . | . | . | . | . | . | . | . | . | . | . | . | . | . |
| | 458096 | . | . | . | . | . | . | . | . | T | . | . | . | . | . | . | . | . | . | . | . | . | . | . | . |
| | 458097 | ? | . | . | . | . | . | . | . | T | . | . | . | . | . | . | . | . | . | . | . | . | . | . | . |
| | 458098 | ? | . | . | . | . | . | . | . | T | . | . | . | . | . | . | . | . | . | . | . | . | . | . | . |
| | 458099 | ? | . | . | . | . | . | . | . | T | . | . | . | . | . | . | . | . | . | . | . | . | . | . | . |
| | 458100 | ? | . | . | . | . | . | . | . | T | . | . | . | . | . | . | . | . | . | . | . | . | . | . | . |
| | 458101 | ? | . | . | . | . | . | . | . | T | . | . | . | . | . | . | . | . | . | . | . | . | . | . | . |
| | 458102 | ? | . | . | . | . | . | . | . | T | . | . | . | . | . | . | . | . | . | . | . | . | . | . | . |
| | 458103 | ? | . | . | . | . | . | T | . | T | . | . | . | . | . | . | . | . | . | . | . | . | . | . | . |
| | 458104 | . | . | . | . | . | . | . | . | T | . | . | . | . | . | . | . | . | . | . | . | . | . | . | . |
| | 458105 | ? | . | . | . | . | . | . | . | T | . | . | . | . | . | . | . | . | . | . | . | . | . | . | . |
| | 458106 | ? | . | . | . | . | . | . | . | T | . | . | . | . | . | . | . | . | . | . | . | . | . | . | . |
| | 458107 | ? | . | . | . | . | . | . | . | T | . | . | . | . | . | . | . | . | . | . | . | . | . | . | . |
| | 458108 | . | . | . | . | . | . | . | . | T | . | . | . | . | . | . | . | . | . | . | . | . | . | . | . |
| | 458109 | ? | . | . | . | . | . | . | . | T | . | . | . | . | . | . | . | . | . | . | . | . | . | . | . |

| | | | | | | | | | | | | | | | | | | | | | | | | | |
|---|---|---|---|---|---|---|---|---|---|---|---|---|---|---|---|---|---|---|---|---|---|---|---|---|---|
| | 458110 | ? | . | . | . | . | . | . | . | T | . | . | . | . | . | . | . | . | . | . | . | . | . | . | . |
| | 458111 | . | . | . | . | . | . | . | . | T | . | . | . | . | . | . | . | . | . | . | . | . | . | . | . |
| | 458112 | . | . | . | . | . | . | . | . | T | . | . | . | . | . | . | . | . | . | . | . | . | . | . | . |
| | 458113 | . | . | . | . | . | . | . | . | T | . | . | . | . | . | . | . | . | . | . | . | . | . | . | . |
| | 458298 | - | . | . | . | . | . | . | . | . | . | . | . | . | . | . | . | . | . | . | . | . | . | . | . |
| | 459911 | ? | . | . | . | . | . | . | . | . | . | . | . | . | . | . | . | . | . | . | . | . | . | . | . |
| | 459913 | ? | . | . | . | . | . | . | . | T | Y | . | . | . | . | . | . | . | . | . | . | . | . | . | . |
| | 459914 | ? | ? | ? | . | . | . | . | . | T | T | . | . | . | . | . | . | . | . | . | . | . | . | . | Y |
| | 459915 | . | . | . | . | . | . | . | . | . | . | . | . | . | . | . | . | . | . | . | . | . | . | . | Y |
| | 459916 | a | . | . | . | . | . | . | . | Y | Y | . | . | . | . | . | . | . | . | . | . | . | . | . | Y |
| | 459917 | . | . | . | . | . | . | . | . | . | . | . | . | . | . | . | . | . | . | . | . | . | . | . | Y |
| | 459918 | . | . | . | . | . | . | . | . | . | . | . | . | . | . | . | . | . | . | . | . | . | . | . | Y |
| | 459919 | . | . | . | . | . | . | . | . | . | . | . | . | . | . | . | . | . | . | . | . | . | . | . | Y |
| | 459920 | . | . | . | . | . | . | . | . | . | . | . | . | . | . | . | . | . | . | . | . | . | . | . | Y |
| | 459921 | . | . | . | . | . | . | . | . | . | . | . | . | . | . | . | . | . | . | . | . | . | . | . | Y |
| | 459922 | . | . | . | . | . | . | . | . | . | . | . | . | . | . | . | . | . | . | . | . | . | . | . | Y |
| | 459923 | ? | . | . | . | . | . | . | . | . | . | . | . | . | . | . | . | . | . | . | . | . | . | . | Y |
| | 459924 | . | . | . | . | . | . | . | . | Y | Y | . | . | . | . | . | . | . | . | . | . | . | . | . | . |
| | 459926 | . | . | . | . | . | . | . | . | . | . | . | . | . | . | . | . | . | . | . | . | . | . | . | Y |
| | 459927 | . | . | . | . | . | . | . | . | . | . | . | . | . | . | . | . | . | . | . | . | . | . | . | Y |
| | 459932 | ? | . | . | . | . | . | . | . | . | . | . | . | . | . | . | . | . | . | . | . | . | . | . | Y |
| | 459933 | . | . | . | . | . | . | . | . | . | . | . | . | . | . | . | . | . | . | . | . | . | . | . | Y |
| | 459934 | . | . | . | . | . | . | . | . | . | . | . | . | . | . | . | ? | . | . | . | . | . | . | . | Y |
| | 459935 | . | . | . | . | . | . | . | . | . | . | . | . | . | . | . | . | . | . | . | . | . | . | . | Y |
| | 459937 | . | . | . | . | . | . | . | . | . | . | . | . | . | . | . | . | . | . | . | . | . | . | . | Y |
| | 459938 | . | . | . | . | . | . | . | . | . | . | . | . | . | . | . | . | . | . | . | . | . | . | . | Y |
| | 459939 | . | . | . | . | . | . | . | . | . | . | . | . | . | . | . | . | . | . | . | . | . | . | . | Y |
| | 459940 | . | . | . | . | . | . | . | . | . | . | . | . | . | . | . | . | . | . | . | . | . | . | . | Y |
| | 459941 | . | . | . | . | . | . | . | . | . | . | . | . | . | . | . | . | . | . | . | . | . | . | . | Y |
| | 459942 | . | . | . | . | . | . | . | . | . | . | . | . | . | . | . | . | . | . | . | . | . | . | . | Y |
| | 459943 | . | . | . | . | . | . | . | . | . | . | . | . | . | . | . | . | . | . | . | . | . | . | . | Y |
| | 461478 | ? | . | . | . | . | . | . | . | T | . | . | . | . | . | . | . | . | . | . | . | . | . | . | . |
| | 461479 | ? | . | . | . | . | . | . | . | T | . | . | . | . | . | . | . | . | . | . | . | . | . | . | . |
| | 461480 | ? | . | . | . | . | . | . | . | T | . | . | . | . | . | . | . | . | . | . | . | . | . | . | . |
| | 461481 | . | . | . | . | . | . | . | . | T | . | . | . | . | . | . | . | . | . | . | . | . | . | . | . |
| | 461482 | . | . | . | . | . | . | . | . | T | . | . | . | . | . | . | . | . | . | . | . | . | . | . | . |
| | 461483 | ? | . | . | . | . | . | . | . | T | . | . | . | . | . | . | . | . | . | . | . | . | . | . | . |
| | 461484 | . | . | . | . | . | . | . | . | T | . | . | . | . | . | . | . | . | . | . | . | . | . | . | . |
| | 461485 | ? | . | . | . | . | . | . | . | T | . | . | . | . | . | . | . | . | . | . | . | . | . | . | . |
| | 461486 | ? | . | . | . | . | . | . | . | T | . | . | . | . | . | . | . | . | . | . | . | . | . | . | . |
| | 461487 | ? | . | . | . | . | . | . | . | T | . | . | . | . | . | . | . | . | . | . | . | . | . | . | . |
| | 461488 | ? | . | . | . | . | . | . | . | T | . | . | . | . | . | . | . | . | . | . | . | . | . | . | . |
| | 461489 | ? | . | . | . | . | . | . | . | T | . | . | . | . | . | . | . | . | . | . | . | . | . | . | . |
| **A1.19** | 461490 | ? | . | . | . | . | . | . | . | T | . | . | . | . | . | . | . | . | . | . | . | . | . | . | . |
| | 461491 | ? | . | . | . | . | . | . | . | T | . | . | . | . | . | . | . | . | . | . | . | . | . | . | . |
| | 461492 | ? | . | . | . | . | . | T | . | T | . | . | . | . | . | . | . | . | . | . | . | . | . | . | . |
| | 461493 | T | . | . | . | . | . | . | . | T | . | . | . | . | . | . | . | . | . | . | . | . | . | . | . |
| | 461494 | T | . | . | . | . | . | T | . | T | . | . | . | . | . | . | . | . | . | . | . | . | . | . | . |
| | 461495 | ? | . | . | . | . | . | . | . | T | . | . | . | . | . | . | . | . | . | . | . | . | . | . | . |
| | 461496 | ? | . | . | . | . | . | . | . | T | . | . | . | . | . | . | . | . | . | . | . | . | . | . | . |
| | 461497 | . | . | . | . | . | . | . | . | T | . | . | . | . | . | . | . | . | . | . | . | . | . | . | . |
| | 461498 | . | . | . | . | . | . | . | . | T | . | . | . | . | . | . | . | . | . | . | . | . | . | . | . |
| | 461499 | ? | . | . | . | . | . | . | . | T | . | . | . | . | . | . | . | . | . | . | . | . | . | . | . |
| | 461500 | ? | . | . | . | . | . | . | . | T | . | . | . | . | . | . | . | . | . | . | . | . | . | . | . |
| | 461501 | ? | . | . | . | . | . | . | T | T | . | . | . | . | . | . | . | . | . | . | . | . | . | . | . |
| | 461502 | ? | . | . | . | . | . | . | T | T | . | . | . | . | . | . | . | . | . | . | . | . | . | . | . |
| | 461503 | ? | . | . | . | . | . | . | . | T | . | . | . | . | . | . | . | . | . | . | . | . | . | . | . |
| | 461504 | ? | . | . | . | . | . | T | . | T | . | . | . | . | . | . | . | . | . | . | . | . | . | . | . |
| | 461505 | ? | . | . | . | . | . | . | . | T | . | . | . | . | . | . | . | . | . | . | . | . | . | . | . |
| | 461506 | ? | . | . | . | . | . | . | . | T | . | . | . | . | . | . | . | . | . | . | . | . | . | . | . |

| 25 | 26 | 27 | 28 | 29 | 30 | 31 | 32 | 33 | 34 | 35 | 36 | 37 | 38 | 39 | 40 | 41 | 42 | 43 | 44 | 45 | 46 | 47 | 48 | 49 | 50 | 51 | 52 |
|---|---|---|---|---|---|---|---|---|---|---|---|---|---|---|---|---|---|---|---|---|---|---|---|---|---|---|---|
| ORF1a | ORF1a | ORF1a | ORF1a | ORF1a | ORF1a | ORF1a | ORF1a | ORF1a | ORF1a | ORF1a | ORF1a | ORF1a | ORF1a | ORF1a | ORF1a | ORF1a | ORF1a | ORF1a | ORF1a | ORF1a | ORF1a | ORF1a | ORF1a | ORF1a | ORF1a | ORF1a | ORF1a |
| 1820 | 1846 | 2143 | 2368 | 2480 | 2558 | 2632 | 2836 | 3037 | 3039 | 3054 | 3085 | 3426 | 3472 | 3634 | 3686 | 3737 | 3817 | 4067 | 4084 | 4255 | 4354 | 4372 | 4444 | 4679 | 4809 | 4866 | 4893 |
| G | A | C | A | A | C | G | C | C | A | A | G | C | A | C | C | C | C | G | C | G | G | A | G | C | C | G | C |
| . | . | . | . | . | . | . | . | . | . | . | . | . | . | . | . | . | . | . | . | . | . | . | . | . | . | . | . |
| . | . | . | . | . | . | . | . | . | . | . | . | . | . | . | . | . | . | . | . | . | . | . | . | . | . | . | . |
| . | . | . | . | . | . | . | . | T | . | . | . | . | . | . | . | . | . | . | . | . | . | . | . | . | T | . | . |
| . | . | . | . | . | . | . | . | T | . | . | . | . | . | . | . | . | . | . | . | . | . | . | . | . | T | . | . |
| . | . | . | . | . | . | . | . | T | . | . | . | . | . | . | . | . | . | . | . | . | . | . | . | . | T | . | . |
| . | . | . | . | . | . | . | . | T | . | . | . | . | . | . | . | . | . | . | . | . | . | . | . | . | T | . | . |
| . | . | . | . | . | . | . | . | T | . | . | . | . | . | . | . | . | . | . | . | . | . | . | . | . | T | . | . |
| . | . | . | . | . | . | . | . | T | . | . | . | . | . | . | . | . | . | . | . | . | . | . | . | . | T | . | . |
| . | . | . | . | . | . | . | . | T | . | . | . | . | . | . | . | . | . | . | . | . | . | . | . | . | T | . | . |
| . | . | . | . | . | . | . | . | T | . | . | . | . | . | . | . | . | . | . | . | . | . | . | . | . | T | . | . |
| . | . | . | . | . | . | . | . | T | . | . | . | . | . | . | . | . | . | . | . | . | . | . | . | . | T | . | . |
| . | . | . | . | . | . | . | . | T | . | . | . | . | . | . | . | . | . | . | . | . | . | . | . | . | T | . | . |
| . | . | . | . | . | . | . | . | T | . | . | . | . | . | . | . | . | . | . | . | . | . | . | . | . | T | . | . |
| . | . | . | . | . | . | . | . | T | . | . | . | . | . | . | . | . | . | . | . | . | . | . | . | . | T | . | . |
| . | . | . | . | . | . | . | . | T | . | . | . | . | . | . | . | . | . | . | . | . | . | . | . | . | . | . | . |
| . | . | . | . | . | . | . | . | T | . | . | . | . | . | . | . | . | . | . | . | . | . | . | . | . | . | . | . |
| . | . | . | . | . | . | . | . | . | . | . | . | . | G | . | . | . | . | . | . | . | . | . | . | . | . | . | . |
| . | . | . | . | . | . | . | . | . | . | . | . | . | G | . | . | . | . | . | . | . | . | . | . | . | . | . | . |
| . | . | . | . | . | . | . | . | . | . | . | . | . | G | . | . | . | . | . | . | . | . | . | . | . | . | . | . |
| . | . | . | . | . | . | . | . | . | . | . | . | . | . | . | . | . | . | . | . | . | . | . | . | . | . | . | . |
| . | . | . | . | . | . | . | . | . | . | . | . | . | . | . | . | . | . | . | . | . | . | . | . | . | . | . | . |
| . | . | . | . | . | . | . | . | . | . | . | . | . | G | . | . | . | . | . | . | . | . | . | . | . | . | . | . |
| . | . | . | . | . | . | . | . | . | . | . | . | . | . | . | . | . | . | . | . | . | . | . | . | . | . | . | . |
| . | . | . | . | . | . | . | . | . | . | . | . | . | . | . | . | . | . | . | . | . | . | . | . | . | . | . | . |
| . | . | . | . | . | . | . | . | . | . | . | . | . | . | . | . | . | . | . | . | . | . | . | . | . | . | . | . |
| . | . | . | . | . | . | . | . | . | . | . | . | . | . | . | . | . | . | . | . | . | . | . | . | . | . | . | . |
| . | . | . | . | . | . | . | . | . | . | . | . | . | . | . | . | . | . | . | . | . | . | . | . | . | . | . | . |
| . | . | . | . | . | . | . | . | . | . | . | . | . | . | . | . | . | . | . | . | . | . | . | . | . | . | . | . |
| . | . | . | . | . | . | . | . | T | . | . | . | . | . | . | . | . | . | . | . | . | . | . | . | . | T | . | . |
| . | . | . | . | . | . | . | . | . | . | . | . | . | . | . | . | . | . | . | . | . | . | . | . | . | . | . | . |
| . | . | . | . | . | . | . | . | T | . | . | . | . | . | . | . | . | . | . | . | T | . | . | . | . | . | . | . |
| . | . | . | . | . | . | . | . | T | . | . | . | . | . | . | . | . | . | . | . | T | . | . | . | . | . | . | . |
| . | . | . | . | . | . | . | . | T | . | . | . | . | . | . | . | . | . | . | . | . | . | . | . | . | . | . | . |
| . | . | . | . | . | . | . | . | T | . | . | . | . | . | . | . | . | . | . | . | . | . | . | . | . | . | . | . |
| . | . | . | . | . | . | . | . | T | . | . | . | . | . | . | . | . | . | . | . | . | . | . | . | . | . | . | . |
| . | . | . | . | . | . | . | . | T | . | . | . | . | . | . | . | . | . | . | . | . | . | . | . | . | . | . | . |
| ? | ? | . | . | . | . | ? | . | T | . | . | . | . | . | . | . | . | . | . | . | . | . | . | . | . | . | . | . |
| . | . | . | . | . | . | . | . | T | . | . | . | . | . | . | . | . | . | . | . | . | . | . | . | . | . | . | . |
| . | . | . | . | . | . | . | . | T | . | . | . | . | . | . | . | . | . | . | . | . | . | . | . | . | . | . | . |
| . | . | . | . | . | . | . | . | . | . | . | . | . | . | . | . | . | . | . | . | . | . | . | . | . | . | . | . |
| . | . | . | . | . | . | . | . | . | . | . | . | . | . | . | . | . | . | . | . | . | . | . | . | . | . | . | . |
| ? | ? | . | . | . | . | ? | . | ? | . | . | . | . | . | . | . | . | . | . | . | . | . | . | . | . | . | . | . |
| . | . | . | . | . | . | . | . | . | . | . | . | . | . | . | . | . | . | . | . | . | . | . | . | . | . | . | . |
| . | . | . | . | . | . | . | . | . | . | . | . | . | . | . | . | . | . | . | . | . | . | . | . | . | . | . | . |
| . | . | . | . | . | . | . | . | T | . | . | . | . | . | . | . | . | . | . | . | . | . | . | . | . | . | . | . |
| . | . | . | . | . | . | . | . | T | . | . | . | . | . | . | . | . | . | . | . | . | . | . | . | . | . | . | . |
| . | . | . | . | . | . | . | . | T | . | . | . | . | . | . | . | . | . | . | . | . | . | . | . | . | . | . | . |
| . | . | . | . | . | . | . | . | T | . | . | . | . | . | . | . | . | . | . | . | . | . | . | . | . | . | . | . |
| . | . | . | . | . | . | . | . | T | . | . | . | . | . | . | . | . | . | . | . | . | . | . | . | . | . | . | . |
| . | . | . | . | . | . | . | . | . | . | . | . | . | . | . | . | . | . | . | . | . | . | . | . | . | . | . | . |
| . | . | . | . | . | . | . | . | T | . | . | . | . | . | . | . | . | . | . | . | . | . | . | . | . | . | . | . |
| . | . | . | . | . | . | . | . | . | . | . | . | . | . | . | . | . | . | . | . | . | . | . | . | . | . | . | . |
| . | . | . | . | . | . | . | . | T | . | . | . | . | . | . | . | . | . | . | . | . | . | . | . | . | . | . | . |
| . | . | . | . | . | . | . | . | T | . | . | . | . | . | . | . | . | . | . | . | . | . | . | . | . | . | . | . |
| . | . | . | . | . | . | . | . | T | . | . | . | . | . | . | . | . | . | . | . | . | . | . | . | . | . | . | . |

| | | | | | | | | | | | | | | | | | | | | | | | | | | | |
|---|---|---|---|---|---|---|---|---|---|---|---|---|---|---|---|---|---|---|---|---|---|---|---|---|---|---|---|
| . | . | . | . | . | . | . | . | T | . | . | . | . | . | . | . | . | . | . | . | . | . | . | . | . | . | . | . |
| . | . | . | . | . | . | . | . | T | . | . | . | . | . | . | . | . | . | . | . | . | . | . | . | . | . | . | . |
| . | . | . | . | . | . | . | . | T | . | . | . | . | . | . | . | . | . | . | . | . | . | . | . | . | . | . | . |
| . | . | . | . | . | . | . | . | T | . | . | . | . | . | . | . | . | . | . | . | . | . | . | . | . | . | . | . |
| . | . | . | . | . | . | . | T | T | . | . | . | . | . | . | . | . | . | . | . | . | . | . | . | . | . | . | . |
| . | . | . | . | . | . | . | . | T | . | . | . | . | . | . | . | . | . | . | . | . | . | . | . | . | . | T | . |
| . | . | . | . | . | . | g | . | T | . | . | . | . | . | . | . | . | . | . | . | . | . | . | . | . | . | . | . |
| . | . | . | . | . | . | . | . | T | . | . | . | . | . | . | . | . | . | . | . | . | . | . | . | . | . | . | . |
| . | . | . | . | . | . | . | . | T | . | . | . | . | . | . | . | . | . | . | . | . | . | . | . | . | . | . | . |
| . | . | . | . | . | . | . | . | T | . | . | . | . | . | . | . | . | . | . | . | . | . | . | . | . | . | . | . |
| . | . | . | . | . | . | . | . | T | . | . | . | . | . | . | . | . | . | . | . | . | . | . | . | . | . | . | . |
| . | . | . | . | . | . | . | . | T | . | . | . | . | . | . | . | . | . | . | . | . | . | . | . | . | . | . | . |
| . | . | . | . | . | . | . | . | T | . | . | . | . | . | . | . | . | . | . | . | . | . | . | . | . | . | . | . |
| . | . | . | . | . | . | . | . | T | . | . | . | . | . | . | . | . | . | . | . | . | . | . | . | . | . | . | . |
| . | . | . | . | . | . | . | . | T | . | . | . | . | . | . | . | . | . | . | . | . | . | . | . | . | . | . | . |
| . | . | . | . | . | . | . | . | T | . | . | . | . | . | . | . | . | . | . | . | . | . | . | . | . | . | . | . |
| . | . | . | . | . | . | . | . | T | . | . | . | . | . | . | . | . | . | . | . | . | . | . | . | . | . | . | . |
| . | . | . | . | . | . | . | . | T | . | . | . | . | . | . | . | . | . | . | . | . | . | . | . | . | . | . | . |
| . | . | . | . | . | . | g | c | T | . | . | . | . | . | . | . | . | . | . | . | . | . | . | . | . | . | . | . |
| . | . | . | . | . | . | . | . | . | . | . | . | . | . | . | . | . | . | . | . | . | . | . | . | . | . | . | . |
| . | . | . | . | . | . | g | . | . | . | . | . | . | . | . | . | . | . | . | . | . | . | . | . | . | . | . | . |
| . | . | . | . | . | . | g | . | . | . | . | . | . | . | . | . | . | ? | . | . | . | . | . | . | . | . | . | . |
| . | . | . | . | . | . | . | . | . | . | . | . | . | . | . | . | . | . | . | . | . | . | . | . | . | . | . | . |
| . | . | . | . | . | . | . | . | . | . | . | . | . | . | . | . | . | . | . | . | . | . | . | . | . | . | . | . |
| . | . | . | . | . | . | . | . | . | . | . | . | . | . | . | . | . | . | . | . | . | . | . | . | . | . | . | . |
| . | . | . | . | . | . | . | . | . | . | . | . | . | . | . | . | . | . | . | . | . | . | . | . | . | . | . | . |
| . | . | . | . | . | . | . | . | . | . | . | . | . | . | . | . | . | . | . | . | . | . | . | . | . | . | . | . |
| . | . | . | . | . | . | . | . | . | . | . | . | . | . | . | . | . | . | . | . | . | . | . | . | . | . | . | . |
| . | . | . | . | . | . | ? | . | . | . | . | . | . | . | . | . | . | . | . | . | . | . | . | . | . | . | . | . |
| . | . | . | . | . | . | g | . | . | . | . | . | . | . | c | . | . | . | . | . | . | . | . | . | . | . | . | . |
| . | . | . | . | . | . | . | . | . | . | . | . | . | . | c | . | . | . | . | . | . | . | . | . | . | . | . | . |
| . | . | . | . | . | . | . | . | . | . | . | . | . | . | . | . | . | . | . | . | . | . | . | . | . | . | . | . |
| . | . | . | . | . | . | . | . | . | . | . | . | . | . | . | . | . | . | . | . | . | . | . | . | . | . | . | . |
| . | . | . | . | . | . | . | . | . | . | . | . | . | . | . | . | . | . | . | . | . | . | . | . | . | . | . | . |
| . | . | . | . | . | . | . | . | . | . | . | . | . | . | . | . | . | . | . | . | . | . | . | . | . | . | . | . |
| . | . | . | . | . | . | . | . | . | . | . | . | . | . | . | . | . | . | . | . | . | . | . | . | . | . | . | . |
| . | . | . | . | . | . | . | . | . | . | . | . | . | . | . | . | . | . | . | . | . | . | . | . | . | . | . | . |
| . | . | . | . | . | . | . | . | . | . | . | . | . | . | . | . | . | . | . | . | . | . | . | . | . | . | . | . |
| . | . | . | . | . | . | . | . | . | . | . | . | . | . | . | . | . | . | . | . | . | . | . | . | . | . | . | . |
| . | . | . | . | . | . | . | . | . | . | . | . | . | . | c | c | c | . | . | . | . | . | . | . | . | . | . | . |
| . | . | . | . | . | . | . | . | . | . | . | . | . | . | . | . | . | . | . | . | . | . | . | . | . | . | . | . |
| . | . | . | . | . | . | . | . | . | . | . | . | . | . | . | . | . | . | . | . | . | . | . | . | . | . | . | . |
| . | . | . | . | . | . | . | . | . | . | . | . | . | . | . | . | . | . | . | . | . | . | . | . | . | . | . | . |
| . | . | . | . | . | . | . | . | . | . | . | . | . | . | . | . | . | . | . | . | . | . | . | . | . | . | . | . |
| . | . | . | . | . | . | . | . | . | . | . | . | . | . | . | . | . | . | . | . | . | . | . | . | . | . | . | . |
| . | . | . | . | . | . | . | . | . | . | . | . | . | . | . | . | . | . | . | . | . | . | . | . | . | . | . | . |
| . | . | . | . | . | . | . | . | . | . | . | . | . | . | . | . | . | . | . | . | . | . | . | . | . | . | . | . |
| . | . | . | . | G | T | . | . | . | . | . | . | . | . | . | . | . | . | . | . | . | . | . | . | . | . | . | . |
| . | . | . | . | G | T | . | . | . | . | . | . | . | . | . | . | . | . | . | . | . | . | . | . | . | . | . | . |
| . | . | . | . | . | . | . | . | . | . | . | . | . | . | . | . | . | . | . | . | . | . | . | . | . | . | . | . |
| . | . | . | . | . | . | . | . | . | . | . | . | . | . | . | . | . | . | . | . | . | . | . | . | . | . | . | . |
| . | . | . | . | . | . | g | . | . | . | . | . | . | . | . | . | . | . | . | . | . | . | . | . | . | . | . | . |
| . | . | . | . | . | . | . | . | T | . | . | . | . | . | . | . | . | . | . | . | . | . | . | . | . | . | . | . |
| . | . | . | . | . | . | . | . | . | . | . | . | . | . | c | c | c | . | . | . | . | . | . | . | . | . | . | . |
| . | . | . | . | . | . | . | . | . | . | . | . | . | . | . | . | . | . | . | . | . | . | . | . | . | . | . | . |
| . | . | . | . | . | . | . | . | . | . | . | . | . | . | . | . | . | . | . | . | . | . | . | . | . | . | . | . |
| . | . | . | a | . | . | . | . | . | . | . | . | . | . | . | . | . | . | . | . | . | . | . | . | . | . | . | . |
| . | . | . | . | . | . | . | . | . | . | . | . | . | . | . | . | . | . | . | . | . | . | . | . | . | . | . | . |
| . | . | . | . | . | . | . | . | . | . | . | . | . | . | . | . | . | . | . | . | . | . | . | . | . | . | . | . |
| . | . | . | . | . | . | . | . | . | . | . | . | . | . | . | . | . | . | . | . | . | . | . | . | . | . | . | . |
| . | . | . | . | . | . | . | . | . | . | . | . | . | . | . | . | . | . | . | . | . | . | . | . | . | . | . | . |
| . | . | . | . | . | . | . | . | . | . | . | . | . | . | . | . | . | . | . | . | . | . | . | . | . | . | . | . |

| | | | | | | | | | | | | | | | | | | | | | | | | | | | |
|---|---|---|---|---|---|---|---|---|---|---|---|---|---|---|---|---|---|---|---|---|---|---|---|---|---|---|---|
| . | . | . | . | . | . | . | . | . | . | . | . | . | . | . | . | . | . | . | . | . | . | . | . | . | . | . | . |
| . | . | . | . | . | . | . | . | . | . | . | . | . | . | . | . | . | . | . | . | . | . | . | . | . | . | . | . |
| . | . | . | . | . | . | . | . | . | . | . | . | . | . | . | . | . | . | . | . | . | . | . | . | . | . | . | . |
| . | . | . | . | . | . | g | . | . | . | . | . | . | . | . | . | . | . | . | . | . | . | . | . | . | . | . | . |
| . | . | . | . | . | . | . | . | . | . | . | . | . | . | . | . | . | . | . | . | . | . | . | . | . | . | . | . |
| . | . | . | . | . | . | . | . | . | . | . | . | c | . | . | . | . | . | . | . | . | . | . | . | . | . | . | . |
| g | . | . | . | . | . | . | . | . | . | . | . | . | . | . | . | . | . | . | . | . | . | . | . | . | . | . | . |
| A | . | . | . | . | . | . | . | . | . | . | . | . | . | . | . | . | . | . | . | . | . | . | . | . | . | . | . |
| . | . | . | . | . | . | g | . | . | . | . | . | . | . | . | . | . | . | . | . | . | . | . | . | . | . | . | . |
| . | . | . | . | . | . | . | . | . | . | . | . | . | . | . | . | . | . | . | . | g | . | . | . | . | . | . | . |
| . | . | . | . | . | . | . | . | . | . | . | . | . | . | . | . | . | . | . | . | . | . | . | . | . | . | . | . |
| . | . | . | . | . | . | g | . | . | . | . | . | . | . | . | . | . | . | . | . | . | . | . | . | . | . | . | . |
| . | . | . | . | . | . | . | . | . | . | . | . | . | . | . | . | . | . | . | . | . | . | . | . | . | . | . | . |
| . | . | . | . | . | . | . | . | . | . | . | . | . | . | . | . | . | . | . | . | . | . | . | . | . | . | . | . |
| . | . | . | . | . | . | . | . | . | . | . | . | . | . | . | . | . | . | . | . | . | . | . | . | . | . | . | . |
| . | . | . | . | . | . | . | . | . | . | . | . | . | . | . | . | . | . | . | . | . | . | . | . | . | . | . | . |
| . | . | . | . | . | . | . | . | . | . | . | . | . | . | . | . | . | . | . | . | . | . | . | . | . | . | . | . |
| . | . | . | . | . | . | . | . | . | . | . | . | . | . | . | . | . | . | . | . | . | . | . | . | . | . | . | . |
| . | . | . | . | . | . | . | . | . | . | . | . | . | . | . | . | . | . | . | . | . | . | . | . | . | . | . | . |
| A | . | . | . | . | . | . | . | . | . | . | . | . | . | . | . | . | . | . | . | . | . | . | . | . | . | . | . |
| . | . | . | . | . | . | . | . | . | . | . | . | . | . | . | c | c | . | . | . | . | . | . | . | . | . | . | . |
| . | . | . | . | . | . | . | . | T | . | . | . | . | . | . | . | . | . | . | . | . | . | . | . | . | . | . | . |
| . | . | . | . | . | . | . | . | T | . | . | . | . | . | . | . | . | . | . | . | . | . | . | . | . | . | . | . |
| . | . | . | . | . | . | . | . | T | . | . | . | . | . | . | . | . | . | . | . | . | . | . | . | . | . | . | . |
| . | . | . | . | . | . | . | . | T | . | . | . | . | . | . | . | . | . | . | . | . | . | . | . | . | . | . | . |
| . | . | . | . | . | . | . | . | T | . | . | . | . | . | . | . | . | . | . | . | . | . | . | . | . | . | . | . |
| . | . | . | . | . | . | . | . | T | . | . | . | . | . | . | . | . | . | . | . | . | . | . | . | . | . | . | . |
| . | . | . | . | . | . | . | . | T | . | . | . | . | . | . | . | . | . | . | . | . | . | . | . | . | . | . | . |
| . | . | . | . | . | . | . | . | T | . | . | . | . | . | . | . | . | . | . | . | . | . | . | . | . | . | . | . |
| . | . | . | . | . | . | . | . | T | . | . | . | . | . | . | . | . | . | . | . | . | . | . | . | . | . | . | . |
| . | . | . | . | . | . | . | . | T | . | . | . | . | . | . | . | . | . | . | . | . | . | . | . | . | . | . | . |
| . | . | . | . | . | . | . | . | T | . | . | . | . | . | . | . | . | . | . | . | . | . | . | . | . | . | . | . |
| A | . | . | . | . | . | . | . | . | . | . | . | . | . | . | . | . | . | . | . | . | . | . | . | . | . | . | . |
| . | . | . | . | . | . | . | . | T | . | . | . | . | . | . | . | . | . | . | . | . | . | . | . | . | . | . | . |
| . | . | . | . | . | . | . | . | . | . | . | . | . | . | . | . | . | . | . | . | . | . | . | . | . | . | . | . |
| . | . | . | . | . | . | . | . | . | . | . | . | . | . | . | . | . | . | . | . | ? | . | . | . | . | . | . | . |
| . | . | . | . | . | . | . | . | . | . | . | . | . | . | . | . | . | . | . | . | . | . | . | . | . | . | . | . |
| . | . | . | . | . | . | . | T | T | . | . | . | . | . | . | . | . | . | . | . | . | . | . | . | . | . | . | . |
| . | . | . | . | . | . | . | T | T | . | . | . | . | . | . | . | . | . | . | . | . | . | . | . | . | . | . | . |
| . | . | . | . | . | . | . | . | T | . | . | . | . | . | . | . | . | . | . | . | . | . | . | . | . | . | . | . |
| . | . | . | . | . | . | . | . | T | . | . | . | . | . | . | . | . | . | . | . | . | . | . | . | . | . | . | . |
| . | . | . | . | . | . | . | . | T | . | . | . | . | . | . | . | . | . | . | . | . | . | . | . | . | . | . | . |
| . | . | . | . | . | . | . | . | T | . | . | . | . | . | . | . | . | . | . | . | . | . | . | . | . | . | . | . |
| . | . | . | . | . | . | . | . | T | . | . | . | . | . | . | . | . | . | . | . | . | . | . | . | . | . | . | . |
| . | . | . | . | . | . | . | T | T | . | . | . | . | . | . | . | . | . | . | . | . | . | . | . | . | . | . | . |
| . | . | . | . | . | . | . | . | . | . | . | . | . | . | . | . | . | . | . | . | . | . | . | . | . | . | . | . |
| . | . | . | . | . | . | . | . | T | . | . | . | . | . | . | . | . | . | . | . | . | . | . | . | . | . | . | . |
| . | . | . | . | . | . | . | . | T | . | . | . | . | . | . | . | . | . | . | . | . | . | . | . | . | . | . | . |
| . | . | . | . | . | . | . | . | T | . | . | . | . | . | . | . | . | . | . | . | . | . | . | . | . | . | . | . |
| . | . | . | . | . | . | . | . | . | . | . | . | . | . | . | . | . | . | . | . | . | . | . | . | . | . | . | . |
| . | . | . | . | . | . | . | . | . | . | . | . | . | . | . | . | . | . | . | . | . | . | . | . | . | . | . | . |
| . | . | . | . | . | . | . | . | T | . | . | . | . | . | . | . | . | . | . | . | . | . | . | . | . | . | . | . |
| . | . | . | . | . | . | . | . | T | . | . | . | . | . | . | . | . | . | . | . | . | . | . | . | . | . | . | . |
| . | . | . | . | . | . | . | . | T | . | . | . | . | . | . | . | . | . | . | . | . | . | . | . | . | . | T | . |
| . | . | . | . | . | . | . | . | T | . | . | . | . | . | . | . | . | . | . | . | . | . | . | . | . | . | . | . |
| . | . | . | . | . | . | . | . | T | . | . | . | . | . | . | . | . | . | . | . | . | . | . | . | . | . | . | . |
| . | . | . | . | . | . | . | T | T | . | . | . | . | . | . | . | . | . | . | . | . | . | . | . | . | . | . | . |
| . | . | . | . | . | . | . | . | T | . | . | . | . | . | . | . | T | . | . | . | . | . | . | . | . | . | T | . |
| . | . | . | . | . | . | . | . | T | . | . | . | . | . | . | . | . | . | . | . | . | . | . | . | . | . | . | . |
| . | . | . | . | . | . | . | T | T | . | . | . | . | . | . | . | . | . | . | . | . | . | . | . | . | . | . | . |
| . | . | . | . | . | . | . | T | T | . | . | . | . | . | . | . | . | . | . | . | . | . | . | . | . | . | . | . |
| . | . | . | . | . | . | . | T | T | . | . | . | . | . | . | . | . | . | . | . | . | . | . | . | . | . | . | . |

| | | | | | | | | | | | | | | | | | | | | | | | | | | | |
|---|---|---|---|---|---|---|---|---|---|---|---|---|---|---|---|---|---|---|---|---|---|---|---|---|---|---|---|
| . | . | . | . | . | . | . | . | T | . | . | . | . | . | . | . | . | . | . | . | . | . | . | . | . | . | . | . |
| . | . | . | . | . | . | . | . | T | . | . | . | . | . | . | . | . | . | . | . | . | . | . | . | . | . | . | . |
| . | . | . | . | . | . | . | . | T | . | . | . | . | . | . | . | . | . | . | . | . | . | . | . | . | . | . | . |
| . | . | . | . | . | . | . | . | T | . | . | . | . | . | . | . | . | . | . | . | . | . | . | . | . | . | . | . |
| . | . | . | . | . | . | . | . | T | . | . | . | . | . | . | . | . | . | . | . | . | . | . | . | . | . | . | . |
| . | . | . | . | . | . | . | T | T | . | . | . | . | . | . | . | . | . | . | . | . | . | . | . | . | . | . | . |
| . | . | . | . | . | . | . | . | T | . | . | . | . | . | . | . | . | . | . | . | . | . | . | . | . | . | . | . |
| . | . | . | . | . | . | . | . | T | . | . | . | . | . | . | . | . | . | . | . | . | . | . | . | . | . | . | . |
| . | . | . | . | . | . | . | . | T | . | . | . | . | . | . | . | T | . | . | . | . | . | . | . | . | . | T | . |
| . | . | . | . | . | . | . | T | T | . | . | . | . | . | . | . | . | . | . | . | . | . | . | . | . | . | . | . |
| . | . | . | . | . | . | . | . | T | . | . | . | . | . | . | . | . | . | . | . | . | . | . | . | . | . | . | . |
| . | . | . | . | . | . | . | . | T | . | . | . | . | . | . | . | . | . | . | . | . | . | . | . | . | . | T | . |
| . | . | . | . | . | . | . | . | T | . | . | . | . | . | . | . | . | . | . | . | . | . | . | . | . | . | . | . |
| . | . | . | . | . | . | . | . | T | . | . | . | . | . | . | . | . | . | . | . | . | . | . | . | . | . | . | . |
| . | . | . | . | . | . | . | . | T | . | . | . | . | . | . | . | . | . | . | . | . | . | . | . | . | . | . | . |
| . | . | . | . | . | . | . | . | T | . | . | . | . | . | . | . | . | . | . | . | . | . | . | . | . | . | . | . |
| . | . | . | . | . | . | . | . | T | . | . | . | . | . | . | . | . | . | . | . | . | . | . | . | . | . | . | . |
| . | . | . | . | . | . | . | . | T | . | . | . | . | . | . | . | . | . | . | . | . | . | . | . | . | . | . | . |
| . | . | . | . | . | . | . | . | T | . | . | . | . | . | . | . | . | . | . | . | . | . | . | . | . | . | . | . |
| . | . | . | . | . | . | . | T | T | . | . | . | . | . | . | . | . | . | . | . | . | . | . | . | . | . | . | . |
| . | . | . | . | . | . | . | . | T | . | . | . | . | . | . | . | . | . | . | . | . | . | . | . | . | . | . | . |
| . | . | . | . | . | . | . | . | T | . | . | . | . | . | . | . | . | . | . | . | . | . | . | . | . | . | . | . |
| . | . | . | . | . | . | . | . | T | . | . | . | . | . | . | . | . | . | . | . | . | . | . | . | . | . | . | . |
| . | . | . | . | . | . | . | T | T | . | . | . | . | . | . | . | . | . | . | . | . | . | . | . | . | . | . | . |
| . | . | . | . | . | . | . | . | T | . | . | . | . | . | . | . | . | . | . | . | . | . | . | . | . | . | . | . |
| . | . | . | . | . | . | . | T | T | . | . | . | . | . | . | . | . | . | . | . | . | . | . | . | . | . | . | . |
| . | . | . | . | . | . | . | . | T | . | . | . | . | . | . | . | . | . | . | . | . | . | . | . | . | . | . | . |
| . | . | . | . | . | . | . | . | T | . | . | . | . | . | . | . | . | . | . | . | . | . | . | . | . | . | . | . |
| . | . | . | . | . | . | . | . | T | . | . | . | . | . | . | . | . | . | . | . | . | . | . | . | . | . | . | . |
| . | . | . | . | . | . | . | T | T | . | . | . | . | . | . | . | . | . | . | . | . | . | . | . | . | . | . | . |
| . | . | . | . | . | . | . | T | T | . | . | . | . | . | . | . | . | . | . | . | . | . | . | . | . | . | . | . |
| . | . | . | . | . | . | . | . | T | . | . | . | . | . | . | . | . | . | . | . | . | . | . | . | . | . | . | . |
| . | . | . | . | . | . | . | . | T | . | . | . | . | . | . | . | . | . | . | . | . | . | . | . | . | . | . | . |
| . | . | . | . | . | . | . | . | T | . | . | . | . | . | . | . | . | . | . | . | . | . | . | . | . | . | . | . |
| . | . | . | . | . | . | . | . | T | . | . | . | . | . | . | . | . | . | . | . | . | . | . | . | . | . | . | . |
| . | . | . | . | . | . | . | T | T | . | . | . | . | . | . | . | . | . | . | . | . | . | . | . | . | . | . | . |
| . | . | . | . | . | . | . | T | T | . | . | . | . | . | . | . | . | . | . | . | . | . | . | . | . | . | . | . |
| . | . | . | . | . | . | . | . | T | . | . | . | . | . | . | . | . | . | . | . | . | . | . | . | . | . | . | . |
| . | . | . | . | . | . | . | T | T | . | . | . | . | . | . | . | . | . | . | . | . | . | . | . | . | . | . | . |
| . | . | . | . | . | . | . | T | T | . | . | . | . | . | . | . | . | . | . | . | . | . | . | . | . | . | . | . |
| . | . | . | . | . | . | . | T | T | . | . | . | . | . | . | . | . | . | . | . | . | . | . | . | . | . | . | . |
| . | . | . | . | . | . | . | T | T | . | . | . | . | . | . | . | . | . | . | . | . | . | . | . | . | . | . | . |
| . | . | . | . | . | . | . | . | T | . | . | . | . | . | . | . | . | . | . | . | . | . | . | . | . | . | . | . |
| . | . | . | . | . | . | . | . | T | . | . | . | . | . | . | . | . | . | . | . | . | . | . | . | . | . | . | . |
| . | . | . | . | . | . | . | . | T | . | . | . | . | . | . | . | . | . | . | . | . | . | . | . | . | . | . | . |
| . | . | . | . | . | . | . | . | T | . | . | . | . | . | . | . | . | . | . | . | . | . | . | . | . | . | . | . |
| . | . | . | . | . | . | . | . | T | . | . | . | . | . | . | . | . | . | . | . | . | . | . | . | . | . | . | . |
| . | . | . | . | . | . | . | . | T | . | . | . | . | . | . | . | . | . | . | . | . | . | . | . | . | . | . | . |
| . | . | . | . | . | . | . | . | T | . | . | . | . | . | . | . | . | . | . | . | . | . | . | . | . | . | . | . |
| . | . | . | . | . | . | . | . | T | . | . | . | . | . | . | . | . | . | . | . | . | . | . | . | . | . | . | . |
| . | . | . | . | . | . | . | . | T | . | . | . | . | . | . | . | . | . | . | . | . | . | . | . | . | . | . | . |
| . | . | . | . | . | . | . | . | T | . | . | . | . | . | . | . | . | . | . | . | . | . | . | . | . | . | . | . |
| . | . | . | . | . | . | . | . | T | . | . | . | . | . | . | . | . | . | . | . | . | . | . | . | . | . | . | . |
| . | . | . | . | . | . | . | . | T | . | . | . | . | . | . | . | . | . | . | . | . | . | . | . | . | . | . | . |
| . | . | . | . | . | . | . | . | T | . | . | . | . | . | . | . | . | . | . | . | . | . | . | . | . | . | . | . |
| . | . | . | . | . | . | . | . | T | . | . | . | . | . | . | . | . | . | . | . | . | . | . | . | . | . | . | . |
| . | . | . | . | . | . | . | . | T | . | . | . | . | . | . | . | . | . | . | . | . | . | . | . | . | . | . | . |
| . | . | . | . | . | . | . | . | T | . | . | . | . | . | . | . | . | . | . | . | . | . | . | . | . | . | . | . |
| . | . | . | . | . | . | . | T | T | . | . | . | . | . | . | . | . | . | . | . | . | . | . | . | . | . | . | . |
| . | . | . | . | . | . | . | . | T | . | . | . | . | . | . | . | . | . | . | . | . | . | . | . | . | . | . | . |
| . | . | . | . | . | . | . | . | T | . | . | . | . | . | . | . | . | . | . | . | . | . | . | . | . | . | . | . |
| . | . | . | . | . | . | . | . | T | . | . | . | . | . | . | . | . | . | . | . | . | . | . | . | . | . | . | . |

| | | | | | | | | | | | | | | | | | | | | | | | | | | | |
|---|---|---|---|---|---|---|---|---|---|---|---|---|---|---|---|---|---|---|---|---|---|---|---|---|---|---|---|
| . | . | . | . | . | . | . | . | T | . | . | . | . | . | . | . | . | . | . | . | . | . | . | . | . | . | . | . |
| . | . | . | . | . | . | . | T | T | . | . | . | . | . | . | . | . | . | . | . | . | . | . | . | . | . | . | . |
| . | . | . | . | . | . | . | . | T | . | . | . | . | . | . | . | . | . | . | . | . | . | . | . | . | . | . | . |
| . | . | . | . | . | . | . | . | T | . | . | . | . | . | . | . | . | . | . | . | . | . | . | . | . | . | . | . |
| . | . | . | . | . | . | . | . | . | . | . | . | . | . | . | . | . | . | . | . | . | . | . | . | . | . | . | . |
| . | . | . | . | . | . | . | . | . | . | . | . | . | . | . | . | . | . | . | . | . | . | . | . | . | . | . | . |
| . | . | . | . | . | . | . | . | . | . | . | . | . | . | . | . | . | . | . | . | . | . | . | . | . | . | . | . |
| . | . | . | . | . | . | . | . | . | . | . | . | . | . | . | . | . | . | . | . | . | . | . | . | . | . | . | . |
| . | . | . | . | . | . | . | . | . | . | . | . | . | . | . | . | . | . | . | . | . | . | . | . | . | . | . | . |
| . | . | . | . | . | . | . | . | . | . | . | . | . | . | . | . | . | . | . | . | . | . | . | . | . | . | . | . |
| . | . | . | . | . | . | . | . | . | . | . | . | . | . | . | . | . | . | . | . | . | . | . | . | . | . | . | . |
| . | . | . | . | . | . | . | . | . | . | . | . | . | . | . | . | . | . | . | . | . | . | . | . | . | . | . | . |
| . | . | . | . | . | . | . | . | . | . | . | . | . | . | . | . | . | . | . | . | . | . | . | . | . | . | . | . |
| . | . | . | . | . | . | . | . | . | . | . | . | . | . | . | . | . | . | . | . | . | . | . | . | . | . | . | . |
| . | . | . | . | . | . | . | . | . | . | . | . | . | . | . | . | . | . | . | . | . | . | . | . | . | . | . | . |
| . | . | . | . | . | . | . | . | . | . | . | . | . | . | . | . | . | . | . | . | . | . | . | . | . | . | . | . |
| . | . | . | . | . | . | . | . | . | . | . | . | . | . | . | . | . | . | . | . | . | . | . | . | . | . | . | . |
| . | . | . | . | . | . | . | . | . | . | . | . | . | . | . | . | . | . | . | . | . | . | . | . | . | . | . | . |
| A | . | . | . | . | . | . | . | . | . | . | . | . | . | . | . | . | . | . | . | . | . | . | . | . | . | . | . |
| . | . | . | . | . | . | . | . | . | . | . | . | . | . | . | . | . | . | . | . | . | . | . | . | . | . | . | . |
| . | . | . | . | . | . | . | . | . | . | . | . | . | . | . | . | . | . | . | . | . | . | . | . | . | . | . | . |
| . | . | . | . | . | . | . | . | . | . | . | . | . | . | . | . | . | . | . | . | . | . | . | . | . | . | . | . |
| . | . | . | . | . | . | . | . | . | . | . | . | . | . | . | . | . | . | . | . | . | . | . | . | . | . | . | . |
| . | . | . | . | . | . | . | . | . | . | . | . | . | . | . | . | . | . | . | . | . | . | . | . | . | . | . | . |
| . | . | . | . | . | . | . | . | . | . | . | . | . | . | . | . | . | . | . | . | . | . | . | . | . | . | . | . |
| . | . | . | . | . | . | . | . | . | . | . | . | . | . | . | . | . | . | . | . | . | . | . | . | . | . | . | . |
| . | . | . | . | . | . | . | . | . | . | . | . | . | . | . | . | . | . | . | . | . | . | . | . | . | . | . | . |
| . | . | . | . | . | . | . | . | . | . | . | . | . | . | . | . | . | . | . | . | . | . | . | . | . | . | . | . |
| . | . | . | . | . | . | . | . | . | . | . | . | . | . | . | . | . | . | . | . | . | . | . | . | . | . | . | . |
| . | . | . | . | . | . | . | . | . | . | . | . | . | . | . | . | . | . | . | . | . | . | . | . | . | . | . | . |
| . | . | . | . | . | . | . | . | . | . | . | . | . | . | . | . | . | . | . | . | . | . | . | . | . | . | . | . |
| . | . | . | . | . | . | T | . | . | . | . | . | . | . | . | . | . | . | . | . | . | . | . | . | . | . | . | . |
| . | . | . | . | . | . | T | . | . | . | . | . | . | . | . | . | . | . | . | . | . | . | . | . | . | . | . | . |
| . | . | . | . | . | . | T | . | . | . | . | . | . | . | . | . | . | . | . | . | . | . | . | . | . | . | . | . |
| . | . | . | . | . | . | . | . | T | . | . | . | . | . | . | . | . | . | . | . | . | . | . | . | . | . | . | . |
| . | . | . | . | . | . | . | . | . | . | . | . | . | . | . | . | . | . | . | . | . | . | . | . | . | . | . | . |
| . | . | . | . | . | . | . | . | . | . | . | . | . | . | . | . | . | . | . | . | . | . | . | . | . | . | . | . |
| . | . | . | . | . | . | . | . | . | . | . | . | . | . | . | . | . | . | . | . | . | . | . | . | . | . | . | . |
| . | . | . | . | . | . | . | . | . | . | . | . | . | . | . | . | . | . | . | . | . | . | . | . | . | . | . | . |
| . | . | . | . | . | . | . | . | . | . | . | . | . | . | . | . | . | . | . | . | . | . | . | . | . | . | . | . |
| . | . | . | . | . | . | . | . | . | . | . | . | . | . | . | . | . | . | . | . | . | . | . | . | . | . | . | . |
| . | . | . | . | . | . | . | . | . | . | . | . | . | . | . | . | . | . | . | . | . | . | . | . | . | . | . | . |
| . | . | . | . | . | . | . | . | . | . | . | . | . | . | . | . | . | . | . | . | . | . | . | . | . | . | . | . |
| . | . | . | . | . | . | . | . | . | . | . | . | . | . | . | . | . | . | . | . | . | . | . | . | . | . | . | . |
| . | . | . | . | . | . | . | . | . | . | . | . | . | . | . | . | . | . | . | . | . | . | . | . | . | . | . | . |
| . | . | . | . | . | . | . | . | . | . | . | . | . | . | . | T | . | . | . | . | . | . | . | . | . | . | . | . |
| A | T | . | . | . | . | . | . | . | . | . | . | . | . | . | . | . | . | . | . | . | . | . | . | . | . | . | . |
| . | . | . | . | . | . | . | . | . | . | . | . | . | . | . | . | . | . | . | . | . | . | . | . | . | . | . | . |
| . | . | . | . | . | . | . | . | . | . | . | . | . | . | . | . | . | . | . | . | . | . | . | . | . | . | . | . |
| A | . | . | . | . | . | . | . | . | . | . | . | . | . | . | . | . | . | . | . | . | . | . | . | . | . | . | . |
| . | . | . | . | . | . | . | . | . | . | . | . | . | . | . | . | . | . | . | . | . | . | . | . | . | . | . | . |
| . | . | . | . | . | . | . | . | . | . | . | . | . | . | . | . | . | . | . | . | T | . | . | . | . | . | . | . |
| A | T | . | . | . | . | . | . | . | . | . | . | . | . | . | . | . | . | . | . | . | . | . | . | . | . | . | . |
| . | . | . | . | . | . | . | . | . | . | . | . | . | . | . | . | . | . | . | . | . | . | . | . | . | . | . | . |
| . | . | . | . | . | . | . | . | . | . | . | . | . | . | . | . | . | . | . | . | . | . | . | . | . | . | . | . |
| . | . | . | . | . | . | . | . | . | . | . | . | . | . | . | . | . | . | . | . | . | . | . | . | . | . | . | . |
| . | . | . | . | . | . | . | . | T | . | . | . | . | . | . | . | . | . | . | . | . | . | . | . | . | . | . | . |
| . | . | . | . | . | . | . | . | . | . | . | . | . | . | . | . | . | . | . | . | . | . | . | . | . | . | . | . |
| . | . | . | . | . | . | . | . | T | . | . | . | . | . | . | . | . | . | . | . | . | . | . | . | . | . | . | . |
| A | . | . | . | . | . | . | . | . | . | . | . | . | . | . | . | . | . | . | . | . | . | . | . | . | . | . | . |

| | | | | | | | | | | | | | | | | | | | | | | | | | | | |
|---|---|---|---|---|---|---|---|---|---|---|---|---|---|---|---|---|---|---|---|---|---|---|---|---|---|---|---|
| . | . | . | . | . | . | . | . | . | . | . | . | . | . | . | . | . | . | . | . | . | . | . | . | . | . | . | . |
| . | . | . | . | . | . | . | . | . | . | . | . | . | . | . | . | . | . | . | . | . | . | . | . | . | . | . | . |
| . | . | . | . | . | . | . | . | . | . | . | . | . | . | . | . | . | . | . | . | . | . | . | . | . | . | . | . |
| . | . | . | . | . | . | . | . | . | . | . | . | . | . | . | . | . | . | . | . | . | . | . | . | . | . | . | . |
| . | . | . | . | . | . | . | . | . | . | . | . | . | . | . | . | . | . | . | . | . | . | . | . | . | . | . | . |
| . | . | . | . | . | . | . | . | . | . | . | . | . | . | . | . | . | . | . | . | . | . | . | . | . | . | . | . |
| . | . | . | . | . | . | . | . | T | . | . | . | . | . | T | . | . | . | . | . | . | . | . | . | . | . | . | . |
| . | . | . | . | . | . | . | . | T | . | . | . | . | . | . | . | . | . | . | . | . | . | . | . | . | . | . | . |
| . | . | . | . | . | . | . | . | T | . | . | . | . | . | . | . | . | . | . | . | . | . | . | . | . | . | . | . |
| . | . | . | . | . | . | . | . | . | . | . | . | . | . | . | . | . | . | . | . | . | . | . | . | . | . | . | . |
| . | . | . | . | . | . | . | . | T | . | . | . | . | . | . | . | . | . | . | . | . | . | . | . | . | . | . | . |
| . | . | . | . | . | . | . | . | T | . | . | . | . | . | . | . | . | . | . | . | . | . | . | . | . | . | . | . |
| A | . | . | . | . | . | . | . | . | . | . | . | . | . | . | . | . | . | . | . | . | . | . | . | . | . | . | . |
| . | . | . | . | . | . | . | . | T | . | . | . | . | . | . | . | . | . | . | . | . | . | . | . | . | . | T | . |
| . | . | . | . | . | . | . | . | T | . | . | . | . | . | . | . | . | . | . | . | . | . | . | . | . | . | . | . |
| . | . | . | . | . | . | . | . | . | . | . | . | . | . | . | . | . | . | . | . | . | . | . | . | . | . | . | . |
| . | . | . | . | . | . | . | . | . | . | . | . | . | . | . | . | . | . | . | . | . | . | . | . | . | . | . | . |
| . | . | . | . | . | . | . | T | T | . | . | . | . | . | . | . | . | . | . | . | . | . | . | . | . | . | . | . |
| . | . | . | . | . | . | . | . | T | . | . | . | . | . | T | . | . | . | . | . | . | . | . | . | . | . | . | . |
| . | . | . | . | . | . | . | T | T | . | . | . | . | . | . | . | . | . | . | . | . | . | . | . | . | . | . | . |
| . | . | . | . | . | . | . | T | T | . | . | . | . | . | . | . | . | . | . | . | . | . | . | . | . | . | . | . |
| . | . | . | . | . | . | . | T | T | . | . | . | . | . | . | . | . | . | . | . | . | . | . | . | . | . | . | . |
| . | . | . | . | . | . | . | . | . | . | . | . | . | . | . | . | . | . | . | . | . | . | . | T | . | . | . | . |
| . | . | . | . | . | . | . | . | T | . | . | . | . | . | . | . | . | . | . | T | . | . | . | . | . | . | . | . |
| . | . | . | . | . | . | . | . | . | . | . | . | . | . | . | . | . | . | . | . | . | . | . | T | . | . | . | . |
| . | . | . | . | . | . | . | . | T | . | . | . | . | . | . | . | . | . | . | T | . | . | . | . | . | . | . | . |
| . | . | . | . | . | . | . | . | T | . | . | . | . | . | . | . | . | . | . | T | . | . | . | . | . | . | . | . |
| . | . | . | . | . | . | . | . | . | . | . | . | . | . | . | . | . | . | . | . | . | . | . | T | . | . | . | . |
| . | . | . | . | . | . | . | . | T | . | . | . | . | . | . | . | . | . | . | . | . | . | . | . | . | . | . | . |
| . | . | . | . | . | . | . | . | . | . | . | . | . | . | . | . | . | . | . | . | . | . | . | T | . | . | . | . |
| . | . | . | . | . | . | . | T | T | . | . | . | . | . | . | . | . | . | . | . | . | . | . | . | . | . | . | . |
| . | . | . | . | . | . | . | . | T | . | . | . | . | . | . | . | . | . | . | T | . | . | . | . | . | . | . | . |
| . | . | . | . | . | . | . | . | . | . | . | . | . | . | . | . | . | . | . | . | . | . | . | . | . | . | . | . |
| g | a | c | a | a | c | g | c | t | a | a | g | c | a | c | c | c | c | g | c | g | g | a | g | c | c | g | c |
| g | a | c | a | a | c | g | c | t | a | a | g | c | a | c | c | c | c | g | c | g | g | a | g | c | c | g | c |
| g | a | c | a | a | c | g | c | t | a | a | g | c | a | c | c | c | c | g | c | g | g | a | g | c | c | g | c |
| g | a | c | a | a | c | g | c | t | a | a | g | c | a | c | c | c | c | g | c | g | g | a | g | c | c | g | c |
| g | a | c | a | a | c | g | c | t | a | a | g | c | a | c | c | c | c | g | c | g | g | a | g | t | c | g | c |
| g | a | c | a | a | c | g | c | t | a | a | g | c | a | c | c | c | c | g | c | g | g | a | g | t | c | g | c |
| g | a | c | a | a | c | g | c | t | a | a | g | c | a | c | c | c | c | g | c | g | g | a | g | c | c | g | c |
| g | a | c | a | a | c | g | c | t | a | a | g | c | a | c | c | c | c | g | c | g | g | a | g | t | c | g | c |
| g | a | c | a | a | c | g | c | t | a | a | g | c | a | c | c | c | c | g | c | g | g | a | g | c | c | g | c |
| g | a | c | a | a | c | g | c | c | a | a | g | c | a | c | c | c | c | g | c | g | g | a | g | c | c | g | c |
| g | a | c | a | a | c | g | c | c | a | a | g | c | a | c | c | c | c | g | c | g | g | a | g | c | c | g | c |
| g | a | c | a | a | c | g | c | c | a | a | g | c | a | c | c | c | c | g | c | g | g | a | g | c | c | g | c |
| g | a | c | a | a | c | g | c | c | a | a | g | c | a | c | c | c | c | g | c | g | g | a | g | c | c | g | c |
| g | a | c | a | a | c | g | c | t | a | a | g | c | a | c | c | c | c | g | c | g | g | a | g | c | c | g | c |
| g | a | c | a | a | c | g | c | c | a | a | g | c | a | c | c | c | c | g | c | g | g | a | g | c | c | g | c |
| g | a | c | a | a | c | g | c | c | a | a | g | c | a | c | c | c | c | g | c | g | g | a | g | c | c | g | c |
| g | a | c | a | a | c | g | c | t | a | a | g | c | a | c | c | c | c | g | c | g | g | a | g | c | c | g | c |
| g | a | c | a | a | c | g | c | t | a | a | g | c | a | c | c | c | c | g | c | g | g | a | g | c | c | g | c |
| g | a | c | a | a | c | g | c | t | a | a | g | c | a | c | c | c | c | g | c | g | g | a | g | c | c | g | c |
| g | a | c | a | a | c | g | c | c | g | a | g | c | a | c | c | c | c | g | c | g | g | a | g | c | c | g | c |
| g | a | c | a | a | c | g | c | t | a | a | g | c | a | t | c | c | c | g | c | g | g | a | g | c | c | g | c |
| a | a | c | a | a | c | g | c | c | a | a | g | c | a | c | c | c | c | g | c | g | g | a | g | c | c | g | c |
| g | a | c | a | a | c | g | c | t | a | a | g | c | a | c | c | c | c | g | c | g | g | a | g | c | c | g | c |
| g | a | c | a | a | c | g | c | t | a | a | g | c | a | t | c | c | c | g | c | g | g | a | g | c | c | g | c |
| . | . | . | . | . | . | . | . | T | . | . | . | . | . | . | . | . | . | . | . | . | . | . | . | . | . | . | . |
| . | . | . | . | . | . | . | . | T | . | . | . | . | . | . | . | . | . | . | . | . | . | . | . | . | . | . | . |
| . | . | . | . | . | . | . | . | T | . | . | . | . | . | . | . | . | . | . | . | . | . | . | . | . | . | . | . |
| . | . | . | . | . | . | . | . | . | . | . | . | . | . | . | . | . | . | . | . | . | . | . | T | . | . | . | . |
| . | . | . | . | . | . | . | . | T | . | . | . | . | . | . | . | . | . | . | . | . | . | . | . | . | . | . | . |

```
. . . . . . . . . . . . . . . . . . . . . . . . . . . .
. . . . . . . . T . . . . . . . . . . . . . . . . . . .
. . . . . . . . . G . . . . . . . . . . . . . . . . . .
. . . . . . . . . G . . . . . . . . . . . . . . . . . .
. . . . . . . . T . . . . . . . . . . . . . . . . . . .
. . . ? . . . . . . . . . . . . . . . . . . . . . . . .
. . . . . . . . T . . . . . T . . . . . . . . . . . . .
. . . . . . . . . . . . . . . . . . . . . . . . . . . .
. . . . . . . . T . . . . . . . . . . . . . . . . . . .
. . . . . . . . T . . . . . . . . . . . . . . . . . . .
. . . . . . . . . . . . . . . . . . . . . . . . . . . .
. . . . . . . . . . . . . . . . . . . . . . . . . . . .
. . . . . . . . . . . . . . . . . . . . . . . . . . . .
. . . . . . . . . . . . . . . . . . . . . . . . . . . .
. . . . . . . . . . . . . . . . . . . . . . . . . . . .
. . . . . . . . . . . . . . . . . . . . . . . . . . . .
. . . . . . . . . . . . . . . . . . . . . . . . . . . .
. . . . . . . . . . . . . . . . . . . . . . . . . . . .
. . . . . . . . . . . . . . . . . . . . . . . . . . . .
. . . . . T . . . . . . . . . . . . . . . . . . . . . .
. . . . . . . . . . . . . . . . . . . . . . . . . . . .
. . . . . . . . . . . . . . . . . . . . ? . . . . . . .
. . . . . . . . T . . . . . . . . . . . . . . . . . . .
. . . . . . . . T . . . . . T . . . . . . . . . . . . .
. . . . . . . . T . . . . . . . . . . . . . . . . . . A
. . . . . . . . T . . . . . . . . . . . . . . . . . . .
. . . . . . . . T . . . . . . . . . . . . . . . . . . .
. . . . . . . . T . . . . . T . . . . . . . . . . . . .
. . . . . . . . T . . . . . . . . . . . . . . . . . . .
. . . . . . . . . . . . . . . . . . . . . . . . . . . .
. . . . . . . . T . . . . . T . . . . . . . . . . . . .
. . . . . . . . T . . . . . T . . . . . . . . . . . . .
. . . . . . . . . . . . . . . . . . . . . . . . . . . .
. . . . . . . . Y . . . . . . . . . . . . . . . . T . .
. . . . . . . . . . . . . . . . . . . . . . . . . . . .
. . . . . . . . . . . . . . . . . . . . . . . . . . . .
. . . . . . . . . . . . . . . . . . . . . . . . . . . .
. . . . . . . . . . . . . . . . . . . . . . . . . . . .
. . . . . . . . . . . . . . . . . . . . . . . . . . . .
. . . . . . . . . . . . . . . . . . . . . . . . . . . .
. . . . . . . . . . . . . . . . . . . . . . . . . . . .
. . . . . . . . . . . . . . . . . . . . . . . . . . . .
. . . . . . . . T . . . . . . . . . . . . . . . . . . .
. . . . . . . . T . . . . . . . . . . . T . . . . . . .
. . . . . . . . T . . . . . . . . . . . T . . . . . . .
. . . . . . . . T . . . . . . . . . . . T . . . . . . .
. . . . . . . . T . . . . . . . . . . . . . . . . . . .
. . . . . . . . T . . . . . . . . . . . T . . . . . . .
. . . . . . . . T . . . . . . . . . . . . . . . . . . .
. . . . . . . . T . . . . . . . . . . . T . . . . . . .
. . . . . . . . T . . . . . . . . . . . T . . . . . . .
. . . . . . . . T . . . . . . . . . . . T . . . . . . .
. . . . . . . . T . . . . . . . . . . . . . . . . . . .
. . . . . . . . T . . . . . . . . . . . . . . . . . . .
. . . . . . . . T . . . . . . . . . . . . . . . . . . .
. . . . . . . . T . . . . . . . . . . . . . . . . . . .
. . . . . . . . T . . . . . . . . . . . . . . . . . . .
. . . . . . . . . . . . . . . . . . . . . . . . . . . .
. . . . . . . . T . . . . . . . . . . . . . . . . . . .
. . . . . . . . T . . . . . . . . . . . . . . . . . . .
. . . . . . . . T . . . . . . . . . . . . . . . . . . .
. . . . . . . . . . . . . . . . . . . . . . . . . . . .
```

| | | | | | | | | | | | | | | | | | | | | | | | | | | | |
|---|---|---|---|---|---|---|---|---|---|---|---|---|---|---|---|---|---|---|---|---|---|---|---|---|---|---|---|
| . | . | . | . | . | . | . | . | T | . | . | . | . | . | . | . | . | . | . | . | . | . | . | . | . | . | . | . |
| . | . | . | . | . | . | . | . | T | . | . | . | . | . | . | . | . | . | . | . | . | . | . | . | . | . | . | . |
| . | . | . | . | . | . | . | . | T | . | . | . | . | . | . | . | . | . | . | . | . | . | . | . | . | . | . | . |
| . | . | . | . | . | . | . | . | T | . | . | . | . | . | . | . | . | . | . | . | . | . | . | . | . | . | . | . |
| . | . | . | . | . | . | . | . | T | . | . | . | . | . | . | . | . | . | . | . | . | . | . | . | . | . | . | . |
| A | . | . | . | . | . | . | . | . | . | . | . | . | . | . | . | . | . | . | . | . | . | . | . | . | . | . | . |
| . | . | . | . | . | . | . | . | . | . | . | . | . | . | . | . | . | . | . | . | . | . | . | . | . | . | . | . |
| . | . | . | . | . | . | . | . | T | . | . | . | . | . | . | . | . | . | . | . | . | . | . | . | . | . | . | . |
| . | . | . | T | . | . | . | . | T | . | . | . | . | . | . | . | . | . | . | . | . | . | . | . | . | . | . | T |
| . | . | . | . | . | . | . | . | T | . | . | . | . | . | . | . | . | . | . | . | . | . | . | . | . | . | . | . |
| . | . | . | . | . | . | . | . | T | . | . | . | . | . | . | . | . | . | . | . | . | . | . | . | . | . | . | . |
| . | . | . | . | . | . | . | . | T | . | . | . | . | . | . | . | . | . | . | . | . | . | . | . | . | . | . | . |
| . | . | . | . | . | . | . | . | T | . | . | . | . | . | . | . | . | . | . | . | . | . | . | . | . | . | . | . |
| . | . | . | . | . | . | . | . | T | . | . | . | . | . | . | . | . | . | . | . | . | . | . | . | . | . | . | . |
| . | . | . | . | . | . | . | . | T | . | . | . | . | . | . | . | . | . | . | . | . | . | . | . | . | . | . | . |
| . | . | . | . | . | . | . | . | . | . | . | . | . | . | . | . | . | . | . | . | . | . | . | . | . | . | . | . |
| . | . | . | . | . | . | . | . | T | . | . | . | . | . | . | . | . | . | . | . | . | . | . | . | . | . | . | . |
| . | . | . | . | . | . | . | . | T | . | . | . | . | . | . | . | . | . | . | . | . | . | . | . | . | . | . | . |
| . | . | . | . | . | . | . | . | T | . | . | . | . | . | . | . | . | . | . | . | . | . | . | . | . | . | . | . |
| . | . | . | . | . | . | . | . | T | . | . | . | . | . | . | . | . | . | . | . | . | . | . | . | . | . | . | . |
| . | . | . | . | . | . | . | . | T | . | . | . | . | . | . | . | . | . | . | . | . | . | . | . | . | . | . | . |
| . | . | . | T | . | . | . | . | T | . | . | . | . | . | . | . | . | . | . | . | . | . | . | . | . | . | . | T |
| . | . | . | . | . | . | . | . | T | . | . | . | . | . | . | . | . | . | . | . | . | . | . | . | . | . | . | . |
| . | . | T | . | . | . | . | . | . | . | . | . | . | . | . | . | . | . | . | . | . | . | . | . | . | . | . | . |
| . | . | T | . | . | . | . | . | . | . | . | . | . | . | . | . | . | . | . | . | . | . | . | . | . | . | . | . |
| . | . | T | . | . | . | . | . | . | . | . | . | . | . | . | . | . | . | . | . | . | . | . | . | . | . | . | . |
| . | . | . | . | . | . | . | . | T | . | . | T | . | . | T | . | . | . | . | . | . | . | . | . | . | . | . | . |
| . | . | . | . | . | . | . | . | T | . | . | . | . | . | . | . | . | . | . | . | . | . | . | . | . | . | . | . |
| . | . | . | . | . | . | . | . | T | . | . | . | . | . | . | . | . | . | . | . | . | . | . | . | . | . | . | . |
| . | . | T | . | . | . | . | . | . | . | . | . | . | . | . | . | . | . | . | . | . | . | . | . | . | . | . | . |
| . | . | T | . | . | . | . | . | . | . | . | . | . | . | . | . | . | . | . | . | . | . | . | . | . | . | . | . |
| . | . | T | . | . | . | . | . | . | . | . | . | . | . | . | . | . | . | . | . | . | . | . | . | . | . | . | . |
| . | . | T | . | . | . | . | . | . | . | . | . | . | . | . | . | . | . | . | . | . | . | . | . | . | . | . | . |
| . | . | T | . | . | . | . | . | . | . | . | . | . | . | . | . | . | . | . | . | . | . | . | . | . | . | . | . |
| . | . | . | . | . | . | . | . | T | . | . | T | . | . | T | . | . | . | . | . | . | . | . | . | . | . | . | . |
| . | . | T | . | . | . | . | . | . | . | . | . | . | . | . | . | . | . | . | . | . | . | . | . | . | . | . | . |
| . | . | T | . | . | . | . | . | . | . | . | . | . | . | . | . | . | . | . | . | . | . | . | . | . | . | . | . |
| . | . | . | . | . | . | . | . | . | . | . | . | . | . | . | . | . | . | . | . | . | . | . | . | . | . | . | . |
| . | . | T | . | . | . | . | . | . | . | . | . | . | . | . | . | . | . | . | . | . | . | . | . | . | . | . | . |
| . | . | . | . | . | . | . | . | T | . | . | . | . | . | . | . | . | . | . | . | . | . | . | . | . | . | . | . |
| . | . | . | . | . | . | . | . | T | . | . | . | . | . | . | . | . | . | . | . | . | . | . | . | . | . | . | . |
| . | . | . | . | . | . | . | . | T | . | . | . | . | . | . | . | . | . | . | . | . | . | . | . | . | . | . | . |
| . | . | . | . | . | . | . | . | T | . | . | . | . | . | . | . | . | . | . | . | . | . | . | . | . | . | . | . |
| . | . | . | . | . | . | . | . | T | . | . | . | . | . | . | . | . | . | . | . | . | . | . | . | . | . | . | . |
| . | . | . | . | . | . | . | . | T | . | . | . | . | . | . | . | . | . | . | . | . | . | . | . | . | . | . | . |
| . | . | . | . | . | . | . | . | T | . | . | . | . | . | . | . | . | . | . | . | . | . | . | . | . | . | . | . |
| . | . | . | . | . | . | . | . | T | . | . | . | . | . | . | . | . | . | . | . | . | . | . | . | . | . | . | . |
| . | . | . | . | . | . | . | . | T | . | . | . | . | . | . | . | . | . | . | . | . | . | . | . | . | . | . | . |
| . | . | . | . | . | . | . | . | T | . | . | . | . | . | . | . | . | . | . | . | . | . | . | . | . | . | . | . |
| . | . | . | . | . | . | . | . | T | . | . | . | . | . | . | . | . | . | . | . | . | . | . | . | . | . | . | . |
| . | . | . | . | . | . | . | . | T | . | . | . | . | . | . | . | . | . | . | . | . | . | . | . | . | . | . | . |
| . | . | . | . | . | . | . | . | T | . | . | . | . | . | . | . | . | . | . | . | . | . | . | . | . | . | . | . |
| . | . | . | . | . | . | . | . | T | . | . | . | . | . | . | . | . | . | . | . | . | . | . | . | . | . | . | . |
| . | . | . | . | . | . | . | . | T | . | . | . | . | . | . | . | . | . | . | . | . | . | . | . | . | . | . | . |
| . | . | . | . | . | . | . | . | T | . | . | . | . | . | . | . | . | . | . | . | . | . | . | . | . | . | . | . |
| . | . | . | . | . | . | . | . | T | . | . | . | . | . | . | . | . | . | . | . | . | . | . | . | . | . | . | . |
| . | . | . | . | . | . | . | . | . | . | . | . | . | . | . | . | . | . | . | . | . | . | . | . | . | . | . | . |
| . | . | . | . | . | . | . | . | T | . | . | . | . | . | . | . | . | . | . | . | . | . | . | . | . | . | . | . |
| . | . | . | . | . | . | . | . | T | . | . | . | . | . | . | . | . | . | . | . | . | . | . | . | . | . | . | . |
| . | . | . | . | . | . | . | . | T | . | . | . | . | . | . | . | . | . | . | . | . | . | . | . | . | . | . | . |
| . | . | . | . | . | . | . | . | T | . | . | . | . | . | . | . | . | . | . | . | . | . | . | . | . | . | . | . |
| . | . | . | . | . | . | . | . | T | . | . | . | . | . | . | . | . | . | . | . | . | . | . | . | . | . | . | . |

| | | | | | | | | | | | | | | | | | | | | | | | | | | | |
|---|---|---|---|---|---|---|---|---|---|---|---|---|---|---|---|---|---|---|---|---|---|---|---|---|---|---|---|
| . | . | . | . | . | . | . | . | T | . | . | . | . | . | . | . | . | . | . | . | . | . | . | . | . | . | . | . |
| . | . | . | . | . | . | . | . | T | . | . | . | . | . | . | . | . | . | . | . | . | . | . | . | . | . | . | . |
| . | . | . | . | . | . | . | . | T | . | . | . | . | . | . | . | . | . | . | . | . | . | . | . | . | . | . | . |
| . | . | . | . | . | . | . | . | T | . | . | . | . | . | . | . | . | . | . | . | . | . | . | . | . | . | . | . |
| . | . | . | . | . | . | . | . | T | . | . | . | . | . | . | . | . | . | . | . | . | A | . | . | . | . | . | . |
| . | . | . | . | . | . | . | . | T | . | . | . | . | . | . | . | . | . | . | . | . | A | . | . | . | . | . | . |
| . | . | . | . | . | . | . | . | T | . | . | . | . | . | . | . | . | . | . | . | . | A | . | . | . | . | . | . |
| . | . | . | . | . | . | . | . | T | . | . | . | . | . | . | . | . | . | . | . | . | A | . | . | . | . | . | . |
| . | . | . | . | . | . | . | . | T | . | . | . | . | . | . | . | . | . | . | . | . | . | . | . | . | . | . | . |
| . | . | . | . | . | . | . | . | T | . | . | . | . | . | . | . | . | . | . | . | . | A | . | . | . | . | . | . |
| . | . | . | . | . | . | . | . | T | . | . | . | . | . | T | . | . | . | . | . | . | . | . | . | . | . | . | . |
| . | . | . | . | . | . | . | . | T | . | . | . | . | . | T | . | . | . | . | . | . | . | . | . | . | . | . | . |
| . | . | . | . | . | . | . | . | T | . | . | . | . | . | T | . | . | . | . | . | . | . | . | . | . | . | . | . |
| . | . | . | . | . | . | . | . | T | . | G | . | . | . | . | . | . | . | . | . | . | . | . | . | . | . | . | . |
| . | . | . | . | . | . | . | . | T | . | G | . | . | . | . | . | . | . | . | . | . | . | G | . | . | . | . | . |
| . | . | . | . | . | . | . | . | T | . | . | . | . | . | . | . | . | . | . | . | . | . | . | . | . | . | . | . |
| . | . | . | . | . | . | . | . | T | . | . | . | . | . | . | . | . | . | . | . | . | . | . | . | . | . | . | . |
| . | . | . | . | . | . | . | . | T | . | . | . | . | . | . | . | . | . | . | . | . | . | G | . | . | . | . | . |
| . | . | . | . | . | . | . | . | T | . | . | . | . | . | . | . | . | . | . | . | . | . | G | . | . | . | . | . |
| . | . | . | . | . | . | . | . | T | . | . | . | . | . | . | . | . | . | . | . | . | . | G | . | . | . | . | . |
| . | . | . | . | . | . | . | . | T | . | . | . | . | . | . | . | . | . | . | . | . | . | G | . | . | . | . | . |
| A | . | . | . | . | . | . | . | . | . | . | . | . | . | . | . | . | . | . | . | . | . | . | . | . | . | . | . |
| A | . | . | . | . | . | . | . | . | . | . | . | . | . | . | . | . | . | . | . | . | . | . | . | . | . | . | . |
| . | . | . | . | . | . | . | . | T | . | . | . | . | . | . | . | . | . | . | . | . | . | . | . | . | . | . | . |
| . | . | . | . | . | . | . | . | T | . | . | . | . | . | . | . | . | . | . | . | . | . | . | . | . | . | . | . |
| . | . | . | . | . | . | . | . | . | . | . | . | . | . | . | . | . | . | . | . | . | . | . | . | . | . | . | . |
| . | . | . | . | . | . | . | . | . | . | . | . | . | . | . | . | . | . | . | . | . | . | . | . | . | . | . | . |
| . | . | . | . | . | . | . | . | . | . | . | . | . | . | . | . | . | . | . | . | . | . | . | . | . | . | . | . |
| A | . | . | . | . | . | . | . | . | . | . | . | . | . | . | . | . | . | . | . | . | . | . | . | . | . | . | . |
| . | . | . | . | . | . | . | . | . | . | . | . | . | . | . | . | . | . | . | . | . | . | . | . | . | . | . | . |
| . | . | . | . | . | . | . | . | . | . | . | . | . | . | . | . | . | . | . | . | . | . | . | . | . | . | . | . |
| . | . | . | . | . | . | . | . | . | . | . | . | . | . | . | T | . | . | . | . | . | . | . | . | . | . | . | . |
| A | . | . | . | . | . | . | . | . | . | . | . | . | . | . | . | . | . | . | . | . | . | . | . | . | . | . | . |
| A | . | . | . | . | . | . | . | . | . | . | . | . | . | . | . | . | . | . | . | . | . | . | . | . | . | . | . |
| . | . | . | . | . | . | . | . | . | . | . | . | . | . | . | . | . | . | . | . | . | . | . | . | . | . | . | . |
| A | . | . | . | . | . | . | . | . | . | . | . | . | . | . | . | . | . | . | . | . | . | . | . | . | . | . | . |
| . | . | . | . | . | . | . | . | . | . | . | . | . | . | . | . | . | . | . | . | . | . | . | . | . | . | . | . |
| . | . | . | . | . | . | . | . | . | . | . | . | . | . | . | . | . | . | . | . | . | . | . | . | . | . | . | . |
| . | . | . | . | . | . | . | T | T | . | . | . | . | . | . | . | . | . | . | . | . | . | . | . | . | . | . | . |
| . | . | . | . | . | . | . | . | T | . | . | . | . | . | . | . | . | . | . | . | . | . | . | . | . | . | . | . |
| . | . | . | . | . | . | . | T | T | . | . | . | . | . | . | . | . | . | . | . | . | . | . | . | . | . | . | . |
| . | . | . | . | . | . | . | . | T | . | . | . | . | . | . | . | . | . | . | . | . | . | . | . | . | . | . | . |
| . | . | . | . | . | . | . | T | T | . | . | . | T | . | . | . | . | . | . | . | . | . | . | . | . | . | . | . |
| . | . | . | . | . | . | . | T | T | . | . | . | T | . | . | . | . | . | . | . | . | . | . | . | . | . | . | . |
| . | . | . | . | . | . | . | T | T | . | . | . | . | . | . | . | . | . | . | . | . | . | . | . | . | . | . | . |
| . | . | . | . | . | . | . | . | T | . | . | . | . | . | . | . | . | . | . | . | . | . | . | . | . | . | . | . |
| . | . | . | . | . | . | . | T | T | . | . | . | . | . | . | . | . | . | . | . | . | . | . | . | . | . | . | . |
| . | . | . | . | . | . | . | . | T | . | . | . | . | . | . | . | . | . | . | . | . | . | . | . | . | . | . | . |
| . | . | . | . | . | . | . | . | T | . | . | . | . | . | . | . | . | . | . | . | . | . | . | . | . | . | . | . |
| . | . | . | . | . | . | . | T | T | . | . | . | . | . | . | . | . | . | . | . | . | . | . | . | . | . | . | . |
| . | . | . | . | . | . | . | T | T | . | . | . | . | . | . | . | . | . | . | . | . | . | . | . | . | . | . | . |
| . | . | . | . | . | . | . | T | T | . | . | . | . | . | . | . | . | . | . | . | . | . | . | . | . | . | . | . |
| . | . | . | . | . | . | . | T | T | . | . | . | . | . | . | . | . | . | . | . | . | . | . | . | . | . | . | . |
| . | . | . | . | . | . | . | . | T | . | . | . | . | . | . | . | . | . | . | . | . | . | . | . | . | . | . | . |
| . | . | . | . | . | . | . | . | T | . | . | . | . | . | . | . | . | . | . | . | . | . | . | . | . | . | . | . |
| . | . | . | . | . | . | . | T | T | . | . | . | . | . | . | . | . | . | . | . | . | . | . | . | . | . | . | . |
| . | . | . | . | . | . | . | . | T | . | . | . | . | . | . | . | . | . | . | . | . | . | . | . | . | . | . | . |
| . | . | . | . | . | . | . | . | T | . | . | . | . | . | . | . | . | . | . | . | . | . | . | . | . | . | . | . |
| . | . | . | . | . | . | . | . | T | . | . | . | . | . | . | . | . | . | . | . | . | . | . | . | . | . | . | . |
| . | . | . | . | . | . | . | . | T | . | . | . | . | . | . | . | . | . | . | . | A | . | . | . | . | . | . | . |
| . | . | . | . | . | . | . | . | T | . | . | . | . | . | T | . | . | . | . | . | . | . | . | . | . | . | . | . |
| . | . | . | . | . | . | . | . | T | . | . | . | . | . | . | . | . | . | . | . | . | . | . | . | . | . | . | . |

| | | | | | | | | | | | | | | | | | | | | | | | | | | | |
|---|---|---|---|---|---|---|---|---|---|---|---|---|---|---|---|---|---|---|---|---|---|---|---|---|---|---|---|
| . | . | . | . | . | . | . | . | . | . | . | . | . | . | . | . | . | . | . | . | . | . | . | . | . | . | . | . |
| . | . | . | . | . | . | . | . | T | . | . | . | . | . | . | . | . | . | . | . | . | . | . | . | . | . | . | . |
| . | . | . | . | . | . | . | T | T | . | . | . | . | . | . | . | . | T | . | . | . | . | . | . | . | . | . | . |
| . | . | . | . | . | . | . | T | T | . | . | . | . | . | . | . | . | T | . | . | . | . | . | . | . | . | . | . |
| A | T | . | . | . | . | . | . | . | . | . | . | . | . | . | . | . | . | . | . | . | . | . | . | . | . | . | . |
| . | . | . | . | . | . | . | . | . | . | . | . | . | . | . | . | . | . | . | . | . | . | . | . | . | . | . | . |
| . | . | . | . | . | . | . | . | T | . | . | . | . | . | . | . | . | . | . | . | . | . | . | . | . | . | . | . |
| . | . | . | . | . | . | . | . | T | . | . | . | . | . | . | . | . | . | . | . | . | . | . | . | . | . | . | . |
| . | . | . | . | . | . | . | . | . | . | . | . | . | . | . | . | . | . | . | . | . | . | . | . | . | . | . | . |
| . | . | . | . | . | . | . | . | Y | . | . | . | . | . | . | . | . | . | . | . | . | . | . | . | . | . | . | . |
| . | . | . | . | . | . | . | . | . | . | . | . | . | . | . | . | . | . | . | . | . | . | . | . | . | . | . | . |
| . | . | . | . | . | . | . | . | . | . | . | . | . | . | . | . | . | . | . | . | . | . | . | . | . | . | . | . |
| . | . | . | . | . | . | . | . | . | . | . | . | . | . | . | . | . | . | . | . | . | . | . | . | . | . | . | . |
| . | . | . | . | . | . | . | . | . | . | . | . | . | . | . | . | . | . | . | . | . | . | . | . | . | . | . | . |
| . | . | . | . | . | . | . | . | . | . | . | . | . | . | . | . | . | . | . | . | . | . | . | . | . | . | . | . |
| . | . | . | . | . | . | . | . | . | . | . | . | . | . | . | . | . | . | . | . | . | . | . | . | . | . | . | . |
| . | . | . | . | . | . | . | . | . | . | . | . | . | . | . | . | . | . | . | . | . | . | . | . | . | . | . | . |
| . | . | . | . | . | . | . | . | Y | . | . | . | . | . | . | . | . | . | . | . | . | . | . | . | . | . | . | . |
| . | . | . | . | . | . | . | . | . | . | . | . | . | . | . | . | . | . | . | . | . | . | . | . | . | . | . | . |
| . | . | . | . | . | . | . | . | . | . | . | . | . | . | . | . | . | . | . | . | . | . | . | . | . | . | . | . |
| . | . | . | . | . | . | . | . | . | . | . | . | . | . | . | . | . | . | . | . | . | . | . | . | . | . | . | . |
| . | . | . | . | . | . | . | . | . | . | . | . | . | . | . | . | . | . | . | . | . | . | . | . | . | . | . | . |
| . | . | . | . | . | . | . | . | . | . | . | . | . | . | . | . | . | . | . | . | . | . | . | . | . | . | . | . |
| . | . | . | . | . | . | . | . | . | . | . | . | . | . | . | . | . | . | . | . | . | . | . | . | . | . | . | . |
| . | . | . | . | . | . | . | . | . | . | . | . | . | . | . | . | . | . | . | . | . | . | . | . | . | . | . | . |
| . | . | . | . | . | . | . | . | . | . | . | . | . | . | . | . | . | . | . | . | . | . | . | . | . | . | . | . |
| . | . | . | . | . | . | . | . | . | . | . | . | . | . | . | . | . | . | . | . | . | . | . | . | . | . | . | . |
| . | . | . | . | . | . | . | . | . | . | . | . | . | . | . | . | . | . | . | . | . | . | . | . | . | . | . | . |
| . | . | . | . | . | . | . | . | . | . | . | . | . | . | . | . | . | . | . | . | . | . | . | . | . | . | . | . |
| . | . | . | . | . | . | . | . | . | . | . | . | . | . | . | . | . | . | . | . | . | . | . | . | . | . | . | . |
| . | . | . | . | . | . | . | . | . | . | . | . | . | . | . | . | . | . | . | . | . | . | . | . | . | . | . | . |
| . | . | . | . | . | . | . | . | T | . | . | . | . | . | . | . | . | . | . | T | . | . | . | . | . | . | . | . |
| . | . | . | . | . | . | . | . | T | . | . | . | . | . | . | . | . | . | . | T | . | . | . | . | . | . | . | . |
| . | . | . | . | . | . | . | . | T | . | . | . | . | . | . | . | . | . | . | . | . | . | . | . | . | . | . | . |
| . | . | . | . | . | . | . | . | T | . | . | . | . | . | T | . | . | . | . | . | . | . | . | . | . | . | . | . |
| . | . | . | . | . | . | . | . | T | . | . | . | . | . | . | . | . | . | . | . | . | . | . | . | . | . | . | . |
| . | . | . | . | . | . | . | . | T | . | . | . | . | . | T | . | . | . | . | . | . | . | . | . | . | . | . | . |
| . | . | . | . | . | . | . | T | T | . | . | . | . | . | . | . | . | . | . | . | . | . | . | . | . | . | . | . |
| . | . | . | . | . | . | . | . | T | . | . | . | . | . | . | . | . | . | . | . | . | . | . | . | . | . | . | . |
| . | . | . | . | . | . | . | . | T | . | . | . | . | . | . | . | . | . | . | . | . | . | . | . | . | . | . | . |
| . | . | . | . | . | . | . | . | T | . | . | . | . | . | T | . | . | . | . | . | . | . | . | . | . | . | . | . |
| . | . | . | . | . | . | . | T | T | . | . | . | . | . | . | . | . | . | . | . | . | . | . | . | . | . | . | . |
| . | . | . | . | . | . | . | T | T | . | . | . | . | . | . | . | . | . | . | . | . | . | . | . | . | . | . | . |
| . | . | . | . | . | . | . | . | T | . | . | . | . | . | . | . | . | . | . | . | . | . | . | . | . | . | . | . |
| . | . | . | . | . | . | . | . | T | . | . | . | . | . | . | . | . | . | . | . | . | . | . | . | . | . | . | . |
| . | . | . | . | . | . | . | T | T | . | . | . | . | . | T | . | . | . | . | . | . | . | . | . | . | . | . | . |
| . | . | . | . | . | . | . | . | T | . | . | . | . | . | T | . | . | . | . | . | . | . | . | . | . | . | . | . |
| . | . | . | . | . | . | . | . | T | . | . | . | . | . | T | . | . | . | . | . | . | . | . | . | . | . | . | . |
| . | . | . | . | . | . | . | . | T | . | . | . | . | . | . | . | . | . | . | . | . | . | . | . | . | . | . | . |
| . | . | . | . | . | . | . | . | T | . | . | . | . | . | T | . | . | . | . | . | . | . | . | . | . | . | . | . |
| . | . | . | . | . | . | . | . | T | . | . | . | . | . | . | . | . | . | A | . | . | . | . | . | . | . | . | . |
| . | . | . | . | . | . | . | . | T | . | . | . | . | . | . | . | . | . | A | . | . | . | . | . | . | . | . | . |
| . | . | . | . | . | . | . | T | T | . | . | . | . | . | . | . | . | . | . | . | . | . | . | . | . | . | . | . |
| . | . | . | . | . | . | . | T | T | . | . | . | . | . | . | . | . | . | . | . | . | . | . | . | . | . | . | . |
| . | . | . | . | . | . | . | T | T | . | . | . | . | . | . | . | . | . | . | . | . | . | . | . | . | . | . | . |
| . | . | . | . | . | . | . | T | T | . | . | . | . | . | . | . | . | . | . | . | . | . | . | . | . | . | . | . |
| . | . | . | . | . | . | . | . | T | . | . | . | . | . | . | . | . | . | . | . | . | . | . | . | . | . | . | . |
| . | . | . | . | . | . | . | T | T | . | . | . | . | . | T | . | . | . | . | . | . | . | . | . | . | . | . | . |
| . | . | . | . | . | . | . | . | T | . | . | . | . | . | . | . | . | . | . | . | . | . | . | . | . | . | . | . |
| . | . | . | . | . | . | . | T | T | . | . | . | . | . | . | . | . | . | . | . | . | . | . | . | . | . | . | . |

| 53 | 54 | 55 | 56 | 57 | 58 | 59 | 60 | 61 | 62 | 63 | 64 | 65 | 66 | 67 | 68 | 69 | 70 | 71 | 72 | 73 | 74 | 75 | 76 | 77 | 78 | 79 | 80 |
|---|---|---|---|---|---|---|---|---|---|---|---|---|---|---|---|---|---|---|---|---|---|---|---|---|---|---|---|
| ORF1a | ORF1a | ORF1a | ORF1a | ORF1a | ORF1a | ORF1a | ORF1a | ORF1a | ORF1a | ORF1a | ORF1a | ORF1a | ORF1a | ORF1a | ORF1a | ORF1a | ORF1a | ORF1a | ORF1a | ORF1a | ORF1a | ORF1a | ORF1a | ORF1a | ORF1a | ORF1a | ORF1a |
| 4965 | 5029 | 5062 | 5139 | 5572 | 5700 | 5826 | 6081 | 6310 | 6312 | 6402 | 6466 | 6541 | 6573 | 6616 | 6868 | 6989 | 7319 | 7392 | 7600 | 7945 | 8022 | 8026 | 8080 | 8296 | 8460 | 8653 | 8782 |
| C | G | G | A | G | C | C | A | C | C | C | A | C | C | A | T | T | A | C | C | C | T | A | A | T | C | G | C |
| . | . | . | . | . | . | . | . | . | . | . | . | . | . | . | . | . | . | . | . | . | . | . | . | . | . | . | . |
| . | . | . | . | . | . | . | . | . | . | . | . | . | . | . | . | . | . | . | . | . | . | . | . | . | . | . | T |
| . | . | . | . | . | . | . | . | . | . | . | . | . | . | . | . | . | . | . | . | . | . | . | . | . | . | . | T |
| . | . | . | . | . | . | . | . | . | . | . | . | . | . | . | . | . | . | . | . | . | . | . | . | . | . | . | . |
| . | . | . | . | . | . | . | . | . | . | . | . | . | . | . | . | . | . | . | . | . | . | . | . | . | . | . | T |
| . | . | . | . | . | . | . | . | . | . | . | . | . | . | . | . | . | . | . | . | . | . | . | . | . | . | . | . |
| . | . | . | . | . | . | . | . | . | . | . | . | . | . | . | . | . | . | . | . | . | . | . | . | . | . | . | . |
| . | . | . | . | . | . | . | . | . | . | . | . | . | . | . | . | . | . | . | . | . | . | . | . | . | . | . | . |
| . | . | . | . | . | . | . | . | . | . | . | . | . | . | . | . | . | . | . | . | . | . | . | . | . | . | . | . |
| . | . | . | . | . | . | . | . | . | . | . | . | . | . | . | . | . | . | . | . | . | . | . | . | . | . | . | . |
| . | . | . | . | . | . | . | . | . | . | . | . | . | . | . | . | . | . | . | . | . | . | . | . | . | . | . | . |
| . | . | . | . | . | . | . | . | . | . | . | . | . | . | . | . | . | . | . | . | . | . | . | . | . | . | . | . |
| . | . | . | . | . | . | . | . | . | . | . | . | . | . | . | . | . | . | . | . | . | . | . | . | . | . | . | . |
| . | . | . | . | . | . | . | . | . | . | . | . | . | . | . | . | . | . | . | . | . | . | . | . | . | . | . | . |
| . | . | . | . | . | . | . | . | . | . | . | . | . | . | . | . | . | . | . | . | . | . | . | . | . | . | . | . |
| . | . | . | . | . | . | . | . | . | . | . | . | . | . | . | . | . | . | . | . | . | . | . | . | . | . | . | . |
| . | . | . | . | . | . | . | . | . | . | . | . | . | . | . | . | . | . | . | . | . | . | . | . | . | . | . | . |
| . | . | . | . | . | . | . | . | . | . | . | . | . | . | . | . | . | . | . | . | . | . | . | . | . | . | T | . |
| . | . | . | . | . | . | . | . | . | . | . | . | . | . | . | . | . | . | . | . | . | . | . | . | . | . | T | . |
| . | . | . | . | . | . | . | . | . | . | . | . | . | . | . | . | . | . | . | . | . | . | . | . | . | . | T | . |
| . | . | . | . | . | . | . | . | . | . | . | . | . | . | . | . | . | . | . | . | . | . | . | . | . | . | T | . |
| . | . | . | . | . | . | . | . | . | . | . | . | . | . | . | . | . | . | . | . | . | . | . | . | . | . | . | . |
| . | . | . | . | . | . | . | . | . | . | . | . | . | . | . | . | . | . | . | . | . | . | . | . | . | . | T | . |
| . | . | . | . | . | . | . | . | . | . | . | . | . | . | . | . | . | . | . | . | . | . | . | . | . | . | . | . |
| . | . | . | . | . | . | . | . | . | . | . | . | . | . | . | . | . | . | . | . | . | . | . | . | . | . | T | . |
| . | . | . | . | . | . | . | . | . | . | . | . | . | . | . | . | . | . | . | . | . | . | . | . | . | . | T | . |
| . | . | . | . | . | . | . | . | . | . | . | . | . | . | . | . | . | . | . | . | . | . | . | . | . | . | T | . |
| . | . | . | . | . | . | . | . | . | . | . | . | . | . | . | . | . | . | . | . | . | . | . | . | . | . | T | . |
| . | . | . | . | . | . | . | . | . | . | . | . | . | . | . | . | . | . | . | . | . | . | . | . | . | . | T | . |
| . | . | . | . | . | . | . | . | . | . | . | . | . | . | . | . | . | . | . | . | . | . | . | . | . | . | . | . |
| . | . | . | . | . | . | . | . | . | . | . | . | . | . | . | . | . | . | . | . | . | . | . | . | . | . | T | . |
| . | . | . | . | . | . | . | . | . | . | . | . | . | . | . | . | . | . | . | . | . | . | . | . | . | . | . | . |
| . | . | . | . | . | . | . | . | . | . | . | . | . | . | . | . | . | . | . | . | . | . | . | . | . | . | . | . |
| . | . | . | . | . | . | . | . | . | . | . | . | . | . | . | . | . | . | . | . | . | . | . | . | . | . | . | . |
| . | . | . | . | . | . | . | . | . | . | . | G | . | . | . | . | . | . | . | . | . | . | . | . | . | . | . | . |
| . | . | . | . | . | . | . | . | . | . | . | G | . | . | . | . | . | . | . | . | . | . | . | . | . | . | . | . |
| . | . | . | . | . | . | . | . | . | . | . | . | . | . | . | . | . | . | T | . | . | . | . | . | . | . | . | . |
| . | . | . | T | . | . | . | . | . | . | . | . | . | . | . | . | . | . | T | . | . | G | . | . | . | . | . | . |
| . | . | . | . | . | . | . | . | . | . | . | . | . | . | . | . | . | . | T | . | . | . | . | . | . | . | . | . |
| . | . | . | . | . | . | . | . | . | . | . | . | . | . | . | . | . | . | T | . | . | . | . | . | . | . | . | . |
| . | . | . | . | . | . | . | . | . | A | . | . | . | . | . | . | . | . | T | . | . | . | . | . | . | . | . | . |
| . | . | . | . | . | . | . | . | . | . | . | . | . | . | . | . | . | . | T | . | . | . | . | . | . | . | . | . |
| . | . | . | T | . | . | . | . | . | . | . | . | . | . | . | . | . | . | T | . | . | G | . | . | . | . | . | . |
| . | . | . | . | . | . | . | . | . | A | . | . | . | . | . | . | . | . | T | . | . | . | . | . | . | . | . | . |
| . | . | . | . | . | . | . | . | . | A | . | . | . | . | . | . | . | . | T | . | . | . | . | . | . | . | . | . |
| . | . | . | . | . | . | . | . | . | . | . | . | . | . | . | . | . | . | . | . | . | . | . | . | . | . | . | . |
| . | . | . | . | . | . | . | . | . | . | . | . | . | . | . | . | . | . | . | . | . | . | . | . | . | . | . | . |
| . | . | . | . | . | . | . | . | . | . | . | . | . | . | . | . | . | . | . | . | . | . | . | . | . | . | . | . |
| . | . | . | . | . | . | . | . | . | . | . | . | . | . | . | . | . | . | . | . | . | . | . | . | . | . | . | . |
| . | . | . | . | . | . | . | . | . | . | . | . | . | . | . | . | . | . | . | . | . | . | . | . | . | . | . | . |
| . | . | . | . | . | . | . | . | . | . | . | . | . | . | . | . | . | . | . | . | . | . | . | . | . | . | . | . |
| . | . | . | . | . | . | . | . | . | . | . | . | . | . | . | . | . | . | . | . | . | . | . | . | . | . | . | . |
| . | . | . | . | . | . | . | . | ? | A | . | . | . | . | . | . | . | . | . | . | . | . | . | . | . | . | . | . |
| . | . | . | . | . | . | . | . | . | . | . | . | . | . | . | . | . | . | . | . | . | . | . | . | . | . | . | . |
| . | . | . | . | . | . | . | . | . | . | . | . | . | . | . | . | . | . | . | . | . | . | . | . | . | . | . | . |
| . | . | . | . | . | . | . | . | . | . | . | . | . | . | . | . | . | . | . | . | . | . | . | . | . | . | . | . |

| | | | | | | | | | | | | | | | | | | | | | | | | | | | |
|---|---|---|---|---|---|---|---|---|---|---|---|---|---|---|---|---|---|---|---|---|---|---|---|---|---|---|---|
| . | . | . | . | . | . | . | . | . | . | . | . | . | . | . | . | . | . | . | . | . | . | . | . | . | . | . | . |
| . | . | . | . | . | . | . | . | . | . | . | . | . | . | . | . | . | . | . | . | . | . | . | . | . | . | . | . |
| . | . | . | . | . | . | . | . | . | . | . | . | . | . | . | . | . | . | . | . | . | . | . | . | . | . | . | . |
| . | . | . | . | . | . | . | . | . | . | . | . | . | . | . | . | . | . | . | . | . | . | . | . | . | . | . | . |
| . | . | . | . | . | . | . | . | . | . | . | . | . | . | . | . | . | . | . | . | . | . | . | . | . | . | . | . |
| . | . | . | . | . | . | . | . | . | . | . | . | . | . | . | . | . | . | . | . | . | K | . | . | . | . | . | . |
| . | . | . | . | . | . | . | . | . | . | . | . | . | . | . | . | . | . | . | . | . | K | . | . | . | . | . | . |
| . | . | . | . | . | . | . | . | . | . | . | . | . | . | . | . | . | . | . | . | . | K | . | . | . | . | . | . |
| . | . | . | . | . | . | . | . | . | . | . | . | . | . | . | . | . | . | . | . | . | K | W | . | . | . | . | . |
| . | . | . | . | . | . | . | . | . | . | . | . | . | . | . | . | . | . | . | . | . | K | . | . | . | . | . | . |
| . | . | . | . | . | . | . | . | . | . | . | . | . | . | . | . | . | . | . | . | . | K | . | . | . | . | . | . |
| . | . | . | . | . | . | . | . | . | . | . | . | . | . | . | . | . | . | . | . | . | K | W | . | . | . | . | . |
| . | . | . | . | . | c | . | . | . | . | . | . | . | . | . | . | . | . | . | . | . | K | . | . | . | . | . | . |
| . | . | . | . | . | . | . | a | . | . | . | . | . | . | . | . | . | . | . | . | . | K | . | . | . | . | . | . |
| . | . | . | . | . | . | . | . | . | . | . | . | . | . | . | . | . | . | . | . | . | K | W | . | . | . | . | . |
| . | . | . | . | . | . | . | . | . | . | . | . | . | . | . | . | . | . | . | . | . | K | . | . | . | . | . | . |
| . | . | . | . | . | . | . | . | . | . | . | . | . | . | . | . | . | . | . | . | . | K | . | . | . | . | . | . |
| . | . | . | . | . | . | . | . | . | . | . | . | . | . | . | . | . | . | . | . | . | K | . | . | . | . | . | . |
| . | . | . | . | . | . | . | R | c | a | . | . | . | . | . | . | . | . | . | . | . | . | . | . | . | . | . | . |
| . | . | . | . | . | . | . | . | . | A | . | . | . | . | . | . | . | . | . | . | . | K | . | . | . | c | . | . |
| . | . | . | . | . | . | . | . | . | A | . | . | . | . | . | . | . | . | . | . | . | K | . | . | . | . | . | . |
| . | . | . | . | . | . | . | . | . | A | . | . | . | . | . | . | . | . | . | . | . | K | . | . | . | . | . | . |
| . | . | . | . | . | . | . | . | . | A | . | . | . | . | . | . | . | . | . | . | . | K | W | . | . | . | . | . |
| . | . | . | . | . | . | . | . | . | A | . | . | . | . | . | . | . | . | . | . | . | ? | . | . | . | c | . | . |
| . | . | . | . | . | . | . | . | . | A | . | . | . | . | . | . | . | . | . | . | . | K | . | . | . | . | . | . |
| . | . | . | . | . | . | . | . | . | A | . | . | . | . | . | . | . | . | . | . | . | K | . | . | . | . | . | . |
| . | . | . | . | . | . | . | . | . | A | . | . | . | . | . | . | . | . | . | . | . | K | W | . | . | . | . | . |
| . | . | . | . | . | . | . | . | . | A | . | . | . | . | . | . | . | . | . | . | . | G | W | . | . | . | . | . |
| . | . | . | . | . | . | . | . | . | A | . | . | . | . | . | . | . | . | . | . | . | K | . | . | . | . | . | . |
| . | . | . | . | . | . | . | . | A | A | . | . | . | . | . | . | . | . | . | . | . | K | W | . | . | . | . | . |
| . | . | . | . | . | . | . | . | A | A | . | . | . | . | . | . | . | . | . | . | . | K | . | . | . | . | . | . |
| . | . | . | . | . | . | . | . | . | A | . | . | . | . | . | . | . | . | . | . | . | . | . | . | . | . | . | . |
| . | . | . | . | . | . | . | R | M | A | . | . | . | . | . | . | . | . | . | . | . | . | . | . | . | . | . | . |
| . | . | . | . | . | . | . | . | M | A | . | . | . | . | . | . | . | . | . | . | . | . | . | . | . | . | . | . |
| . | . | . | . | . | . | . | . | . | A | . | . | . | . | . | . | . | . | . | . | . | K | . | . | . | . | . | . |
| . | . | . | . | . | . | . | . | . | A | . | . | . | . | . | . | . | . | . | . | . | . | . | . | . | . | . | . |
| . | . | . | . | . | . | . | . | . | A | . | . | . | . | . | . | . | . | . | . | . | . | . | . | . | . | . | . |
| . | . | . | . | . | . | . | R | . | A | . | . | . | . | . | . | . | . | . | . | . | K | . | . | . | . | . | . |
| . | . | . | . | . | . | . | . | . | A | . | . | . | . | . | . | . | . | . | . | . | ? | . | . | . | . | . | . |
| . | . | . | . | . | . | . | . | . | A | . | . | . | . | . | . | . | . | . | . | . | K | . | . | . | . | . | . |
| . | . | . | . | . | . | . | . | . | . | . | . | . | . | . | . | . | . | . | . | . | . | . | . | . | . | T | . |
| . | . | . | . | . | . | . | . | . | . | . | . | . | . | . | . | . | . | . | . | . | K | . | . | . | . | T | . |
| . | . | . | . | . | . | . | . | . | . | . | . | . | . | . | . | . | . | . | . | . | K | . | . | . | . | T | . |
| . | . | . | . | . | . | . | . | . | . | . | . | . | . | . | . | . | . | . | . | . | K | W | . | . | c | T | . |
| . | . | . | . | . | . | . | . | . | . | . | . | . | . | . | . | . | . | . | . | . | K | . | . | . | . | T | . |
| . | . | . | . | . | . | . | . | . | . | . | . | . | . | . | . | . | . | . | . | . | G | . | . | . | c | T | . |
| . | . | . | . | . | . | . | . | . | . | . | . | . | . | . | . | . | . | . | . | . | ? | . | . | . | . | . | . |
| . | . | . | . | . | . | . | . | . | . | . | . | . | . | . | . | . | . | . | . | . | G | T | . | . | . | . | . |
| . | . | . | . | . | . | . | . | . | . | . | . | . | . | . | . | . | . | . | . | . | G | T | . | . | . | . | T |
| . | . | . | . | . | . | . | . | . | . | . | . | . | . | . | . | . | . | . | . | . | . | . | . | . | . | . | T |
| . | . | . | . | . | . | . | . | M | A | . | . | . | . | . | . | . | . | . | . | . | K | . | . | . | . | . | . |
| . | . | . | . | . | . | . | . | . | . | . | . | . | . | . | . | . | . | . | . | . | K | . | . | . | . | . | . |
| . | . | . | . | . | . | . | . | . | A | . | . | . | . | . | . | . | . | . | . | . | ? | . | . | . | . | . | . |
| . | . | . | . | . | . | . | . | . | A | . | . | . | . | . | . | . | . | . | ? | . | K | . | . | . | . | . | . |
| . | . | . | . | . | . | . | a | . | A | . | . | . | . | . | . | . | . | . | . | . | . | . | . | . | . | . | . |
| . | . | . | . | . | . | . | R | . | A | . | . | . | . | . | . | . | . | . | . | . | . | . | . | . | . | . | . |
| . | . | . | . | . | . | . | . | . | A | . | . | . | . | . | . | . | . | . | . | . | K | . | . | . | . | . | . |
| . | . | . | . | . | . | . | . | . | A | . | . | . | . | . | . | . | . | . | . | . | K | . | . | . | . | . | . |
| . | . | . | . | . | . | . | . | . | A | . | . | . | . | . | . | . | . | . | . | . | K | . | . | . | . | . | . |
| . | . | . | . | . | . | . | . | A | A | . | . | . | . | . | . | . | . | . | . | . | . | . | . | . | . | . | . |
| . | . | . | . | . | . | . | . | A | A | . | . | . | . | . | . | . | . | . | . | . | ? | . | . | . | . | . | . |

| | | | | | | | | | | | | | | | | | | | | | | | | | | | |
|---|---|---|---|---|---|---|---|---|---|---|---|---|---|---|---|---|---|---|---|---|---|---|---|---|---|---|---|
| . | . | . | . | . | . | . | a | A | A | . | . | . | . | . | . | . | . | . | . | . | ? | . | . | . | . | . | . |
| . | . | . | . | . | . | . | . | A | A | . | . | . | . | . | . | . | . | . | T | . | K | W | . | . | . | . | . |
| . | . | . | . | . | . | . | . | . | A | . | . | . | . | . | . | . | . | . | . | . | K | . | . | . | . | . | . |
| . | . | . | . | . | . | . | . | . | ? | . | . | . | . | . | . | t | . | . | Y | . | K | W | . | . | . | . | . |
| . | . | . | . | . | . | . | . | A | A | . | . | . | . | . | . | . | . | . | . | . | K | . | . | . | . | . | . |
| . | . | . | . | . | . | . | . | M | A | . | . | . | . | . | . | . | . | . | . | . | . | . | . | . | . | . | . |
| . | . | . | . | . | . | . | . | . | A | . | . | . | . | . | . | . | . | . | . | . | . | . | . | . | . | . | . |
| . | . | . | . | . | . | . | . | A | A | . | . | . | . | . | . | . | . | . | . | . | K | W | . | . | . | . | . |
| . | . | . | . | . | . | . | . | ? | M | . | . | . | . | . | . | . | . | . | . | . | K | . | . | . | . | . | T |
| . | . | . | . | . | . | . | . | . | A | . | . | . | . | . | . | . | . | . | . | . | K | . | . | . | . | . | . |
| . | . | . | . | . | . | . | a | M | M | . | . | . | . | . | . | . | . | . | . | . | K | . | . | . | . | . | . |
| . | . | . | . | . | . | . | . | . | A | . | . | . | . | . | . | . | . | . | . | . | K | . | . | . | . | . | . |
| . | . | . | . | . | . | . | . | . | A | . | . | . | . | . | . | . | . | . | . | . | . | . | . | . | . | . | . |
| . | . | . | . | . | . | . | . | ? | A | . | . | . | . | . | . | . | . | . | . | . | . | . | . | . | . | . | . |
| . | . | . | . | . | . | . | . | ? | A | . | . | . | . | . | . | . | . | . | . | . | . | . | . | . | . | . | . |
| . | . | . | . | . | . | . | . | . | A | . | . | . | . | . | . | . | . | . | . | . | K | . | . | . | c | . | . |
| . | . | . | . | . | . | . | . | A | A | . | . | . | . | . | . | . | . | . | . | . | . | . | . | . | . | . | c |
| . | . | . | . | . | . | . | . | A | A | . | . | . | . | . | . | . | . | . | T | . | . | . | . | . | . | . | . |
| . | . | . | . | . | . | . | . | . | A | . | . | . | . | . | . | . | . | . | . | . | K | . | . | . | . | . | . |
| . | . | . | . | . | . | . | . | A | A | . | . | . | . | . | . | . | . | . | . | . | K | . | . | . | c | . | . |
| . | . | . | . | . | . | . | . | . | A | . | . | . | . | . | . | . | . | . | . | . | . | . | . | . | . | . | . |
| . | . | . | . | . | . | . | . | . | . | . | . | . | . | . | . | t | . | . | . | . | K | . | . | . | . | . | . |
| . | . | . | . | . | . | . | . | . | . | . | . | . | . | . | . | . | . | . | . | . | K | . | . | . | . | . | . |
| . | . | . | . | . | . | . | . | . | . | . | . | . | . | . | . | . | . | . | . | . | K | . | . | . | . | . | . |
| . | . | . | . | . | . | . | . | . | . | . | . | . | . | . | . | . | . | . | . | . | . | . | . | . | . | . | . |
| . | . | . | . | . | . | . | . | . | . | . | . | . | . | . | . | . | . | . | . | . | ? | . | . | . | . | . | . |
| . | . | . | . | . | . | . | . | . | . | . | . | . | . | . | . | . | . | . | . | . | . | . | . | . | . | . | . |
| . | . | . | . | . | . | . | . | . | . | . | . | . | . | . | . | . | . | . | . | . | K | . | . | . | c | . | . |
| . | . | . | . | . | . | . | . | . | . | . | . | . | . | . | . | . | . | . | . | . | K | . | . | . | c | . | . |
| . | . | . | . | . | . | . | . | . | . | . | . | . | . | . | . | . | . | . | . | . | K | . | . | . | . | . | . |
| . | . | . | . | . | . | . | . | . | . | . | . | . | . | . | . | . | . | . | . | . | K | . | . | . | c | . | . |
| . | . | . | . | . | . | . | . | . | ? | . | . | . | . | . | . | . | . | . | . | . | . | . | . | . | . | . | . |
| . | . | . | . | . | . | . | . | A | A | . | . | . | . | . | . | . | . | . | . | . | . | . | . | . | . | . | . |
| . | . | . | . | . | . | . | . | . | . | . | . | . | . | . | . | . | . | . | . | . | . | . | . | . | . | . | . |
| . | . | . | . | . | . | . | . | . | . | . | . | . | . | . | ? | . | . | ? | . | . | . | . | . | . | . | . | T |
| . | . | . | . | . | . | . | . | . | A | . | . | . | . | . | ? | . | . | . | . | . | . | . | . | . | . | . | . |
| . | . | . | . | . | . | . | . | . | A | . | . | . | . | . | . | . | . | . | . | . | . | . | . | . | . | . | . |
| . | . | . | . | . | . | . | . | . | . | . | . | . | . | . | . | . | . | . | . | . | . | . | . | . | . | . | . |
| . | . | . | . | . | . | . | . | . | . | . | . | . | . | . | . | . | . | . | . | . | . | . | . | . | . | . | . |
| . | . | . | . | . | . | . | . | . | . | . | . | . | . | . | . | . | . | . | . | . | . | . | . | . | . | . | . |
| . | . | . | . | . | . | . | . | . | . | . | . | . | . | . | . | . | . | . | . | . | . | . | . | . | . | . | . |
| . | . | . | . | . | . | . | . | . | . | . | . | . | . | . | . | . | . | . | . | . | . | . | . | . | . | . | . |
| . | . | . | . | . | . | . | . | . | . | . | . | . | . | . | ? | . | . | . | . | . | . | . | . | . | . | . | . |
| . | . | . | . | . | . | . | . | . | . | . | . | . | . | . | . | . | . | . | . | . | . | . | . | . | . | . | . |
| . | . | . | . | . | . | . | . | . | . | . | . | . | . | . | . | . | . | . | . | . | . | . | . | . | . | . | . |
| . | . | . | . | . | . | . | G | . | A | . | . | . | . | . | . | C | . | . | . | . | . | . | . | . | . | . | . |
| . | . | . | . | . | . | . | G | . | A | . | . | . | . | . | . | C | . | . | . | . | . | . | . | . | . | . | . |
| . | . | . | . | . | . | . | . | . | . | . | . | . | . | . | . | . | . | . | . | . | . | . | . | . | . | . | . |
| . | . | . | . | . | . | . | . | . | . | . | . | . | . | . | . | . | . | . | . | . | . | . | . | . | . | . | . |
| . | . | . | . | . | . | . | . | . | A | . | . | . | . | . | . | . | . | . | . | . | . | . | . | . | . | . | . |
| . | . | . | . | . | . | . | . | . | A | . | . | . | . | . | . | . | . | . | . | . | . | . | . | . | . | . | . |
| . | . | . | . | . | . | . | . | . | . | . | . | . | . | . | . | . | . | . | . | . | . | . | . | . | . | . | . |
| . | . | . | . | . | . | . | . | . | . | . | . | . | . | . | . | . | ? | . | . | . | . | . | . | . | . | . | . |
| . | . | . | . | . | . | . | . | . | . | . | . | . | . | . | . | . | . | . | . | . | . | . | . | . | . | . | . |
| . | . | . | . | . | . | . | . | . | . | . | . | . | . | . | . | . | . | . | . | . | . | . | . | . | . | . | . |
| . | . | . | . | . | . | . | . | . | . | . | . | . | . | . | . | . | . | . | . | . | . | . | . | . | . | . | . |
| . | . | . | . | . | . | . | . | . | . | . | . | . | . | . | . | . | . | . | . | . | . | . | . | . | . | . | . |
| . | . | . | . | . | . | . | . | . | . | . | . | . | . | . | . | . | . | . | . | . | . | . | . | . | . | . | . |
| . | . | . | . | . | . | . | . | . | . | . | . | . | . | . | . | . | . | . | . | . | . | . | . | . | . | . | . |
| . | . | . | . | . | . | . | . | . | . | . | . | . | . | . | . | . | . | . | . | . | . | . | . | . | . | . | . |
| . | . | . | . | . | . | . | . | . | . | . | . | . | . | . | . | . | . | . | . | . | . | . | . | . | . | . | . |
| . | . | . | . | . | . | . | . | . | . | . | . | . | . | . | . | . | . | . | . | . | . | . | . | . | . | . | . |

?

?

?

T

?

T

| | | | | | | | | | | | | | | | | | | | | | | | | | | | |
|---|---|---|---|---|---|---|---|---|---|---|---|---|---|---|---|---|---|---|---|---|---|---|---|---|---|---|---|
| . | . | . | . | . | . | . | . | . | . | . | . | . | . | . | . | . | . | . | . | . | . | . | . | . | . | . | . |
| . | . | . | . | . | . | . | . | . | . | . | . | . | . | . | . | . | . | . | . | . | . | . | . | . | . | . | . |
| . | . | . | . | . | . | . | . | . | . | . | . | . | . | . | . | . | . | . | . | . | . | . | . | . | . | . | . |
| . | . | . | . | . | . | . | . | . | . | . | . | . | . | . | . | . | . | . | . | . | . | . | . | . | . | . | . |
| . | . | . | . | . | . | . | . | . | A | . | . | . | . | . | . | . | . | . | . | . | . | . | . | . | . | . | . |
| . | . | . | . | . | . | . | . | . | A | . | . | . | . | . | . | . | . | . | . | . | . | . | . | . | . | . | . |
| . | . | . | . | . | . | . | . | . | A | . | . | . | . | . | . | . | . | . | . | . | . | . | . | . | . | . | . |
| . | . | . | . | . | . | . | . | . | A | . | . | . | . | . | . | . | . | . | . | . | . | . | . | . | . | . | . |
| . | . | . | . | . | . | . | . | . | A | . | . | . | . | . | . | . | . | . | . | . | . | . | . | . | . | . | . |
| . | . | . | . | . | . | . | . | . | A | . | . | . | . | . | . | . | . | . | . | . | . | . | . | . | . | . | . |
| . | . | . | . | . | . | . | . | . | A | . | . | . | . | . | . | . | . | . | . | . | . | . | . | . | . | . | . |
| . | . | . | . | . | . | . | . | . | A | . | . | . | . | . | . | . | . | . | . | . | . | . | . | . | . | . | . |
| . | . | . | . | . | . | . | . | . | A | . | . | . | . | . | . | . | . | . | . | . | . | . | . | . | . | . | . |
| . | . | . | . | . | . | . | . | . | . | . | . | . | . | . | . | . | . | . | . | . | . | . | . | . | . | . | . |
| . | . | . | . | . | . | . | . | . | A | . | . | . | . | . | . | . | . | . | . | . | . | . | . | . | . | . | . |
| . | . | . | . | . | . | . | G | . | A | . | . | . | . | . | . | . | . | . | . | . | . | . | . | . | . | . | . |
| . | . | . | . | . | . | . | . | . | A | . | . | . | . | . | . | . | . | . | . | . | . | . | . | . | . | . | . |
| . | . | . | . | . | . | . | . | A | A | . | . | . | . | . | . | . | . | . | . | . | . | . | . | . | . | . | . |
| . | . | . | . | . | . | . | . | A | A | . | . | . | . | . | . | . | . | . | . | . | . | . | . | . | . | . | . |
| . | . | . | . | . | . | . | . | . | A | . | . | . | . | . | . | . | . | . | . | . | . | . | . | . | . | . | . |
| . | . | . | . | . | . | . | . | . | A | . | . | . | . | . | . | . | . | . | . | . | . | . | . | . | . | . | . |
| . | . | . | . | . | . | . | . | . | A | . | . | . | . | . | . | . | . | . | . | . | . | . | . | . | . | . | . |
| . | . | . | . | . | . | . | . | . | A | . | . | . | . | . | . | . | . | . | . | . | . | . | . | . | . | . | . |
| . | . | . | . | . | . | . | G | . | A | . | . | . | . | . | . | . | . | . | . | . | . | . | . | . | . | . | . |
| . | . | . | . | . | . | . | . | . | A | . | . | . | . | . | . | . | . | . | . | . | . | . | . | . | . | . | . |
| . | . | . | . | . | . | . | . | A | A | . | . | . | . | . | . | . | . | . | . | . | . | . | . | . | . | . | . |
| . | . | . | . | . | . | . | . | . | A | . | . | . | . | . | . | . | . | . | . | . | . | . | . | . | . | . | . |
| . | . | . | . | . | . | . | . | . | A | . | . | . | . | . | . | . | . | . | . | . | . | . | . | . | . | . | . |
| . | . | . | . | . | . | . | . | . | A | . | . | . | . | . | . | . | . | . | . | . | . | . | . | . | . | . | . |
| . | . | . | . | . | . | . | . | A | A | . | . | . | . | . | . | . | . | . | . | . | . | . | . | . | . | . | . |
| . | . | . | . | . | . | . | . | . | A | . | . | . | . | . | . | . | . | . | . | . | . | . | . | . | . | . | . |
| . | . | . | . | . | . | . | . | . | A | . | . | . | . | . | . | . | . | . | . | . | . | . | . | . | . | . | . |
| . | . | . | . | . | . | . | . | . | A | . | . | . | . | . | . | . | . | . | . | . | . | . | . | . | . | . | . |
| . | . | . | . | . | . | . | . | . | A | . | . | . | . | . | . | . | . | . | . | . | . | . | . | . | . | . | . |
| . | . | . | . | . | . | . | . | . | A | . | . | . | . | . | . | . | . | . | . | . | . | . | . | . | . | . | . |
| . | . | . | . | . | . | . | . | . | . | . | . | . | . | . | . | . | . | . | . | . | . | . | . | . | . | . | . |
| . | . | . | . | . | . | . | G | . | A | . | . | . | . | . | . | . | . | . | . | . | . | . | . | . | . | . | . |
| . | . | . | . | . | . | . | . | . | A | . | . | . | . | . | . | . | . | . | . | . | . | . | . | . | . | . | . |
| . | . | . | . | . | . | . | . | . | A | . | . | . | . | . | . | . | . | . | . | . | . | . | . | . | . | . | . |
| . | . | . | . | . | . | . | . | . | A | . | . | . | . | . | . | . | . | . | . | . | . | . | . | . | . | . | . |
| . | . | . | . | . | . | . | G | . | . | . | . | . | . | . | . | . | . | . | . | . | . | . | . | . | . | . | . |
| . | . | . | . | . | . | . | . | . | A | . | . | . | . | . | . | . | . | . | . | . | . | . | . | . | . | . | . |
| . | . | . | . | . | . | . | . | . | . | . | . | . | . | . | . | . | . | . | . | . | . | . | . | . | . | . | . |
| . | . | . | . | . | . | . | G | . | A | . | . | . | . | . | . | . | . | . | . | . | . | . | . | . | . | . | . |
| . | . | . | . | . | . | . | . | A | A | . | . | . | . | . | . | . | . | . | . | . | . | . | . | . | . | . | . |
| . | . | . | . | . | . | . | . | . | A | . | . | . | . | . | . | . | . | . | . | . | . | . | . | . | . | . | . |
| . | . | . | . | . | . | . | . | A | A | . | . | . | . | . | . | . | . | . | . | . | . | . | . | . | . | . | . |
| . | . | . | . | . | . | . | . | . | A | . | . | . | . | . | . | . | . | . | . | . | . | . | . | . | . | . | . |
| . | . | . | . | . | . | . | . | A | A | . | . | . | . | . | . | . | . | . | . | . | . | . | . | . | . | . | . |
| . | . | . | . | . | . | . | . | . | A | . | . | . | . | . | . | . | . | . | . | . | . | . | . | . | . | . | . |
| . | . | . | . | . | . | . | . | . | A | . | . | . | . | . | . | . | . | . | . | . | . | . | . | . | . | . | . |
| . | . | . | . | . | . | . | . | A | A | . | . | . | . | . | . | . | . | . | . | . | . | . | . | . | . | . | . |
| . | . | . | . | . | . | . | G | . | A | . | . | . | . | . | . | . | . | . | . | . | . | . | . | . | . | . | . |
| . | . | . | . | . | . | . | . | . | A | . | . | . | . | . | . | . | . | . | . | . | . | . | . | . | . | . | . |
| . | . | . | . | . | . | . | . | . | A | . | . | . | . | . | . | . | . | . | . | . | . | . | . | . | . | . | . |
| . | . | . | . | . | . | . | . | . | A | . | . | . | . | . | . | . | . | . | . | . | . | . | . | . | . | . | . |
| . | . | . | . | . | . | . | G | . | A | . | . | . | . | . | . | . | . | . | . | . | . | . | . | . | . | . | . |
| . | . | . | . | . | . | . | . | . | A | . | . | . | . | . | . | . | . | . | . | . | . | . | . | . | . | . | . |
| . | . | . | . | . | . | . | . | . | . | . | . | . | . | . | . | . | . | . | . | . | . | . | . | . | . | . | . |
| . | . | . | . | . | . | . | . | . | . | . | . | . | . | . | . | . | . | . | . | . | . | . | . | . | . | . | T |
| . | . | . | . | . | . | . | . | . | . | . | . | . | . | . | . | . | . | . | . | . | . | . | . | . | . | . | . |
| . | . | . | . | . | . | . | . | A | A | . | . | . | . | . | . | . | . | . | . | . | . | . | . | . | . | . | . |

| | | | | | | | | | | | | | | | | | | | | | | | | | | | |
|---|---|---|---|---|---|---|---|---|---|---|---|---|---|---|---|---|---|---|---|---|---|---|---|---|---|---|---|
| . | . | . | . | . | . | . | . | . | A | . | . | . | . | . | . | . | . | . | . | . | . | . | . | . | . | . | . |
| . | . | . | . | . | . | . | . | . | A | . | . | . | . | . | . | . | . | . | . | . | . | . | . | . | . | . | . |
| . | . | . | . | . | . | . | . | . | A | . | . | . | . | . | . | . | . | . | . | . | . | . | . | . | . | . | . |
| . | . | . | . | . | . | . | . | . | A | . | . | . | . | . | . | . | . | . | . | . | . | . | . | . | . | . | . |
| . | . | . | . | . | . | . | . | . | A | . | . | . | . | . | . | . | . | . | . | . | . | . | . | . | . | . | . |
| . | . | . | . | . | . | . | . | . | A | . | . | . | . | . | . | . | . | . | . | . | . | . | . | . | . | . | . |
| . | . | . | . | . | . | . | . | . | . | . | . | . | . | . | . | . | . | . | . | . | . | . | . | . | . | . | . |
| . | . | . | . | . | . | . | . | . | . | . | . | . | . | . | . | . | . | . | . | . | . | . | . | . | . | . | . |
| . | . | . | . | . | . | . | . | . | . | . | . | . | . | . | . | . | . | . | . | . | . | . | . | . | . | . | . |
| . | . | . | . | ? | . | . | . | . | A | . | . | . | . | . | ? | . | . | . | . | . | . | . | . | . | . | . | . |
| . | . | . | . | . | A | . | . | . | . | . | . | . | . | . | . | . | . | . | . | . | . | . | . | . | . | . | . |
| . | . | . | . | . | A | . | . | . | . | . | . | . | . | . | . | . | . | . | . | . | . | . | . | . | . | . | . |
| . | . | . | . | . | . | . | . | A | A | . | . | . | . | . | . | . | . | . | . | . | . | . | . | . | . | . | . |
| . | . | . | . | . | . | . | . | . | . | . | . | . | . | . | . | . | . | . | . | . | . | . | . | . | . | . | . |
| . | . | . | . | . | . | . | . | . | . | . | . | . | . | . | ? | . | . | . | . | . | . | . | . | . | . | . | . |
| . | . | . | . | . | . | . | . | . | . | . | . | . | . | . | ? | . | . | . | . | . | . | . | . | . | . | . | T |
| . | . | . | . | . | . | . | . | . | . | . | . | . | . | . | . | . | ? | . | . | . | . | . | . | . | . | . | T |
| . | . | A | . | . | . | . | . | . | . | . | . | . | . | . | . | . | . | . | . | . | . | . | . | . | . | . | . |
| . | . | . | . | . | . | . | . | . | . | . | . | . | . | . | . | . | . | . | . | . | . | . | . | . | . | . | . |
| . | . | A | . | . | . | . | . | . | . | . | . | . | . | . | . | . | . | . | . | . | . | . | . | . | . | . | . |
| . | . | A | . | . | . | . | . | . | . | . | . | . | . | . | . | . | . | . | . | . | . | . | . | . | . | . | . |
| . | . | . | . | . | . | . | . | . | . | . | . | . | . | . | . | . | . | . | . | . | . | . | . | . | . | . | . |
| . | . | . | . | . | . | . | . | A | A | . | . | . | . | . | . | . | . | . | . | . | . | . | . | . | . | . | . |
| . | . | . | . | . | . | . | . | . | . | . | . | . | . | . | . | . | . | . | . | . | . | . | . | . | . | . | . |
| . | . | . | . | . | . | . | . | A | A | . | . | . | . | . | . | . | . | . | . | . | . | . | . | . | . | . | . |
| . | . | . | . | . | . | . | . | . | . | . | . | . | . | . | . | . | . | . | . | . | . | . | . | . | . | . | . |
| . | . | . | . | . | . | . | . | . | . | . | . | . | . | . | . | . | . | . | . | . | . | . | . | . | . | . | . |
| . | . | . | . | . | . | . | . | A | A | . | . | . | . | . | . | . | . | . | . | . | . | . | . | . | . | . | . |
| . | . | . | . | . | . | . | . | . | . | . | . | . | . | . | . | . | . | . | . | . | . | . | . | . | . | . | . |
| . | . | . | . | . | . | . | . | A | A | . | . | . | . | . | . | . | . | . | . | . | . | . | . | . | . | . | . |
| . | . | . | . | . | . | . | . | . | . | . | . | . | . | . | . | . | . | . | . | . | . | . | . | . | . | . | . |
| . | . | . | . | . | . | . | . | . | . | . | . | . | . | . | . | . | . | . | . | . | . | . | . | . | . | . | . |
| . | . | . | . | . | . | . | . | . | A | . | . | . | . | . | . | . | . | . | . | . | . | . | . | . | . | . | . |
| c | g | g | a | g | a | c | a | c | c | c | a | c | c | a | t | t | a | c | c | c | t | a | a | t | c | g | c |
| c | g | g | a | g | a | c | a | c | c | c | a | c | c | a | t | t | a | c | c | c | t | a | a | t | c | g | c |
| c | g | g | a | g | c | c | a | c | c | c | a | c | c | a | t | t | a | c | c | c | t | a | a | t | c | g | c |
| c | g | g | a | g | c | c | a | c | c | c | a | c | c | a | t | t | a | c | c | c | t | a | a | t | c | g | c |
| c | g | g | a | g | a | c | a | c | c | c | a | c | c | a | t | t | a | c | c | c | t | a | a | t | c | g | c |
| c | g | g | a | g | a | c | a | c | c | c | a | c | c | a | t | t | a | c | c | c | t | a | a | t | c | g | c |
| c | g | g | a | g | a | c | a | c | c | c | a | c | c | a | t | t | a | c | c | c | t | a | a | t | c | g | c |
| c | g | g | a | g | a | c | a | c | c | c | a | c | c | a | t | t | a | c | c | c | t | a | a | t | c | g | c |
| c | g | g | a | g | a | c | a | c | c | c | a | c | c | a | t | t | a | c | c | c | t | a | a | t | c | g | c |
| c | g | g | a | g | c | c | a | c | c | c | a | c | c | a | t | t | a | c | c | c | t | a | a | t | c | g | t |
| c | g | g | a | g | c | c | a | c | c | c | a | c | c | a | t | t | a | c | c | c | t | a | a | t | c | g | t |
| c | g | g | a | g | c | c | a | c | c | c | a | c | c | a | t | t | a | c | c | c | t | a | a | t | c | g | t |
| c | g | g | a | g | c | c | a | c | c | c | a | c | c | a | t | t | a | c | c | c | t | a | a | t | c | g | t |
| c | g | g | a | g | a | c | a | c | c | c | a | c | c | a | t | t | a | c | c | c | t | a | a | t | c | g | c |
| c | g | g | a | g | c | c | a | c | a | c | a | c | c | a | t | t | a | c | c | c | t | a | a | t | c | g | c |
| c | g | g | a | g | c | c | a | a | a | c | a | c | c | a | t | t | a | c | c | c | t | a | a | t | c | g | c |
| c | g | g | a | g | a | c | a | c | c | c | a | c | c | a | t | t | a | c | c | c | t | a | a | t | c | g | c |
| c | g | g | a | g | a | c | a | c | c | c | a | c | c | a | t | t | a | c | c | c | t | a | a | t | c | g | c |
| c | g | g | a | g | a | c | a | c | c | c | a | c | c | a | t | t | a | c | c | c | t | a | a | t | c | g | c |
| c | g | g | a | g | c | c | a | c | c | c | a | c | c | a | t | t | a | c | c | c | t | a | a | t | c | g | c |
| c | g | g | a | g | c | c | a | c | c | c | a | c | c | a | t | t | a | c | c | c | t | a | a | t | c | g | c |
| c | g | g | a | g | c | c | a | a | a | c | a | c | c | a | t | t | a | c | c | c | t | a | a | t | c | g | c |
| c | g | g | a | g | a | t | a | c | c | c | a | c | c | a | t | t | a | c | c | c | t | a | a | t | c | g | c |
| c | g | g | a | g | c | c | a | c | c | c | a | c | c | a | t | t | a | c | c | c | t | a | a | t | c | g | c |
| . | . | . | . | . | . | . | . | . | . | . | . | . | . | . | . | . | . | T | T | . | . | . | . | . | . | . | . |
| . | . | . | . | . | . | . | . | . | . | . | . | . | . | . | . | . | . | T | . | . | . | . | . | . | . | . | . |
| . | . | . | . | . | . | . | . | . | . | . | . | . | . | . | . | . | . | T | . | . | . | . | . | . | . | . | . |
| . | . | . | . | . | . | . | . | A | A | . | . | . | . | . | . | . | . | T | . | . | . | . | . | . | . | . | . |
| . | . | . | . | . | . | . | . | . | . | . | . | . | . | . | . | . | . | T | . | . | . | . | . | . | . | . | . |

| | | | | | | | | | | | | | | | | | | | | | | | | | | | |
|---|---|---|---|---|---|---|---|---|---|---|---|---|---|---|---|---|---|---|---|---|---|---|---|---|---|---|---|
| . | . | . | . | . | . | . | . | A | A | . | . | . | . | . | . | . | . | T | . | . | . | . | . | . | . | . | . |
| . | . | . | . | . | A | . | . | . | . | . | . | . | . | . | . | . | . | . | . | . | . | . | . | . | . | . | . |
| . | . | . | . | . | . | . | . | . | . | . | . | . | . | . | . | . | . | . | . | . | . | . | . | . | . | . | . |
| . | . | . | . | . | . | . | . | . | . | . | . | . | . | . | . | . | . | . | . | . | . | . | . | . | . | . | . |
| . | . | . | . | . | . | . | . | . | . | . | . | . | . | . | . | . | . | . | . | . | . | . | . | . | . | . | . |
| . | . | . | . | . | . | . | . | . | . | . | . | . | . | . | . | . | . | . | . | . | . | . | . | . | . | T | . |
| . | . | . | . | . | . | . | . | . | . | . | . | . | . | . | . | . | . | . | . | . | . | . | . | . | . | . | . |
| . | . | . | . | . | . | . | . | . | . | . | . | . | . | . | . | . | . | . | . | . | . | . | . | . | . | . | T |
| . | . | . | . | . | . | . | . | . | . | . | . | . | . | . | . | . | . | . | . | . | . | . | . | . | . | . | . |
| . | . | . | . | . | . | . | . | . | . | . | . | . | . | . | . | . | . | . | . | . | . | . | . | . | . | . | . |
| . | . | . | . | . | . | . | . | . | . | . | . | . | . | . | . | . | . | . | . | . | . | . | . | . | . | . | T |
| . | . | . | . | . | . | . | . | . | . | . | . | . | . | . | . | . | . | . | . | . | . | . | . | . | . | . | T |
| . | . | . | . | . | . | . | . | . | . | . | . | . | . | . | . | . | . | . | . | . | . | . | . | . | . | . | T |
| . | . | . | . | . | . | . | . | . | A | . | . | . | . | . | . | . | . | . | . | . | . | . | . | . | . | . | . |
| . | . | . | . | . | . | . | . | . | A | . | . | . | . | . | . | . | . | . | . | . | . | . | . | . | . | . | . |
| . | . | . | . | . | . | . | . | A | A | . | . | . | . | . | . | . | . | . | . | . | . | . | . | . | . | . | . |
| . | . | . | . | . | . | . | . | . | A | . | . | . | . | . | . | . | . | . | . | . | . | . | . | . | . | . | . |
| . | . | . | . | . | . | . | . | . | A | . | . | . | . | . | . | . | . | . | . | . | . | . | . | . | . | . | . |
| . | . | . | . | . | . | . | . | A | A | . | . | . | . | . | . | . | . | . | . | . | . | . | . | . | . | . | . |
| . | . | . | . | . | . | . | . | . | . | . | . | . | . | . | . | . | . | . | . | . | . | . | . | . | . | . | . |
| . | . | . | . | . | . | . | . | . | A | . | . | . | . | . | . | . | . | . | . | . | . | . | . | . | . | . | . |
| . | . | . | . | . | . | . | . | . | A | . | . | . | . | . | ? | . | . | . | . | . | . | . | . | . | . | . | . |
| . | . | . | . | . | A | . | . | . | . | . | . | . | . | . | . | . | . | . | . | . | . | . | . | . | . | . | . |
| . | . | . | . | . | . | . | . | . | . | . | . | . | . | . | . | . | . | . | . | . | . | . | . | . | . | . | . |
| . | . | . | . | . | A | . | . | . | . | . | . | . | . | . | . | . | . | . | . | . | . | . | . | . | . | . | . |
| . | . | . | . | . | A | . | . | . | . | . | . | . | . | . | . | . | . | . | . | . | . | . | . | . | . | . | . |
| . | . | . | . | . | . | . | . | . | . | . | . | . | . | . | . | . | . | . | . | . | . | . | . | . | . | . | . |
| . | . | . | . | . | . | . | . | . | . | . | . | . | . | . | . | . | . | . | . | . | . | . | . | . | . | . | . |
| . | . | . | . | . | A | . | . | . | . | . | . | . | . | . | . | . | . | . | . | . | . | . | . | . | . | . | . |
| . | . | . | . | . | . | . | . | . | A | . | . | . | . | . | . | . | . | . | . | . | . | . | . | . | . | . | . |
| . | . | . | . | . | . | . | . | . | . | . | . | . | . | . | . | . | . | . | . | . | . | . | . | . | . | . | . |
| . | . | . | . | . | . | . | . | . | . | . | . | . | . | . | . | . | . | . | . | . | . | . | . | . | . | . | . |
| . | . | . | . | . | . | . | . | . | A | . | . | . | . | . | . | . | . | . | . | . | . | . | . | . | . | . | . |
| . | . | . | . | . | . | . | . | M | M | . | . | . | . | . | . | . | . | . | . | . | . | . | . | . | . | . | . |
| . | . | . | . | . | . | . | . | . | A | . | . | . | . | . | . | . | . | . | . | . | . | . | . | . | . | . | . |
| . | . | . | . | . | . | . | . | . | A | . | . | . | . | . | . | . | . | . | . | . | . | . | . | . | . | . | . |
| . | . | . | . | . | . | . | . | . | A | . | . | . | . | . | . | . | . | . | . | . | . | . | . | . | . | . | . |
| . | . | . | . | . | . | . | . | . | A | . | . | . | . | . | . | . | . | . | . | . | . | . | . | . | . | . | . |
| . | . | . | . | . | . | . | . | . | A | . | . | . | . | . | . | . | . | . | . | . | . | . | . | . | . | . | . |
| . | . | . | . | . | . | . | . | . | A | . | . | . | . | . | . | . | . | . | . | . | . | . | . | . | . | . | . |
| . | . | . | . | . | . | . | . | . | . | . | . | . | . | . | ? | . | . | . | . | . | . | . | . | . | . | . | T |
| . | . | . | . | . | . | . | . | . | . | . | . | . | . | . | . | . | . | . | . | . | . | . | . | . | . | . | T |
| . | . | . | . | . | . | . | . | . | . | . | . | . | . | . | . | . | . | . | . | . | . | . | . | . | . | . | . |
| . | . | . | . | . | . | . | . | . | . | . | . | . | . | . | . | . | . | . | . | . | . | . | . | . | . | . | . |
| . | . | . | . | . | . | . | . | . | . | . | . | . | . | . | . | . | . | . | . | . | . | . | . | . | . | . | . |
| . | . | . | . | . | . | . | . | . | . | . | . | . | . | . | . | . | . | . | . | . | . | . | . | . | . | . | . |
| . | . | . | . | . | . | . | . | . | . | . | . | . | . | . | . | . | . | . | . | . | . | . | . | . | . | . | . |
| . | . | . | . | . | . | . | . | . | . | . | . | . | . | . | . | . | . | . | . | . | . | . | . | . | . | . | . |
| . | . | . | . | . | . | . | . | . | . | . | . | . | . | . | . | . | . | . | . | . | . | . | . | . | . | . | . |
| . | . | . | . | . | . | . | . | . | . | . | . | . | . | . | . | . | . | . | . | . | . | . | . | . | . | . | . |
| . | . | . | . | T | . | . | . | . | . | . | . | . | . | . | . | . | . | . | . | . | . | . | . | . | . | . | . |
| . | . | . | . | . | . | . | . | . | . | . | . | . | . | . | . | . | . | . | . | . | . | . | . | . | . | . | . |
| . | . | . | . | . | . | . | . | . | . | . | . | . | . | . | . | . | . | . | . | . | . | . | . | . | . | . | . |
| . | . | . | . | . | . | . | . | . | . | . | . | . | . | . | . | . | . | . | . | . | . | . | . | . | . | . | . |
| . | . | . | . | . | . | . | . | . | . | . | . | . | . | . | . | . | . | . | . | . | . | . | . | . | . | . | . |
| . | . | . | . | . | . | . | . | . | . | . | . | . | . | . | . | . | . | . | . | . | . | . | . | . | . | . | . |
| . | . | . | . | . | . | . | . | . | . | . | . | . | . | . | . | . | . | . | . | T | . | . | . | . | . | . | T |
| . | . | . | . | . | . | . | . | . | . | . | . | . | . | . | . | . | . | . | . | . | . | . | . | . | . | . | . |
| . | . | . | . | . | . | . | . | . | . | . | . | . | . | . | . | . | . | . | . | . | . | . | . | . | . | . | . |
| . | . | . | . | . | . | . | . | . | . | . | . | . | . | . | . | . | . | . | . | . | . | . | . | . | . | . | . |
| . | . | . | . | . | . | . | . | . | . | . | . | . | . | . | . | . | . | . | . | . | . | . | . | . | . | . | . |
| . | . | . | . | . | . | . | . | . | . | . | . | . | . | . | . | . | . | . | . | T | . | . | . | . | . | . | T |

| | | | | | | | | | | | | | | | | | | | | | | | | | | | |
|---|---|---|---|---|---|---|---|---|---|---|---|---|---|---|---|---|---|---|---|---|---|---|---|---|---|---|---|
| . | . | . | . | . | . | . | . | . | . | . | . | . | . | . | . | . | . | . | . | . | . | . | . | . | . | . | . |
| . | . | . | . | . | . | . | . | . | . | . | . | . | . | . | . | . | . | . | . | . | . | . | . | . | . | . | . |
| . | . | . | . | . | . | . | . | . | . | . | . | . | . | . | . | . | . | . | . | . | . | . | . | . | . | . | . |
| . | . | . | . | . | . | . | . | . | . | . | . | . | . | . | . | . | . | . | . | . | . | . | . | . | . | . | . |
| . | . | . | . | . | . | . | . | . | . | . | . | . | . | . | . | . | . | . | . | . | . | . | . | . | . | . | . |
| . | . | . | . | . | . | . | . | A | A | . | . | . | . | . | . | . | . | . | . | . | . | . | . | . | . | . | . |
| . | . | . | . | . | . | . | . | . | . | . | . | . | . | . | . | . | . | . | . | . | . | . | . | . | . | . | T |
| . | . | . | . | . | . | . | . | . | . | . | . | . | . | . | . | . | . | . | . | . | . | . | . | . | . | . | . |
| . | . | . | . | . | . | . | . | . | . | . | . | T | . | . | . | . | . | . | . | . | . | . | . | . | . | . | . |
| . | . | . | . | . | . | . | . | . | . | . | . | . | . | . | . | . | . | . | . | . | . | . | . | . | . | . | . |
| . | . | . | . | . | . | . | . | . | . | . | . | . | . | . | . | . | . | . | . | . | . | . | . | . | . | . | . |
| . | . | . | . | . | . | . | . | . | . | . | . | . | . | . | . | . | . | . | . | . | . | . | . | . | . | . | . |
| . | . | . | . | . | . | . | . | . | . | . | . | . | . | . | . | . | . | . | . | . | . | . | . | . | . | . | . |
| . | . | . | . | . | . | . | . | . | . | . | . | . | . | . | . | . | . | . | . | . | . | . | . | . | . | . | . |
| . | . | . | . | . | . | . | . | . | . | . | . | . | . | . | . | . | . | . | . | . | . | . | . | . | . | . | . |
| . | . | . | . | . | . | . | . | . | . | . | . | . | . | . | . | . | . | . | . | T | . | . | . | . | . | . | T |
| . | . | . | . | . | . | . | . | . | . | . | . | . | . | . | . | . | . | . | . | . | . | . | . | . | . | . | . |
| . | . | . | . | . | . | . | . | . | . | . | . | . | . | . | . | . | . | . | . | . | . | . | . | . | . | . | . |
| . | . | . | . | . | . | . | . | . | . | . | . | . | . | . | . | . | . | . | . | . | . | . | . | . | . | . | . |
| . | . | . | . | . | . | . | . | . | . | . | . | . | . | . | . | . | . | . | . | . | . | . | . | . | . | . | . |
| . | . | . | . | . | . | . | . | . | . | . | . | . | . | . | . | . | . | . | . | . | . | . | . | . | . | . | . |
| . | . | . | . | . | . | . | . | . | . | . | . | T | . | . | . | . | . | . | . | . | . | . | . | . | . | . | . |
| . | . | . | . | . | . | . | . | . | . | . | . | . | . | . | . | . | . | . | . | . | . | . | . | . | . | . | . |
| . | . | . | . | . | . | . | . | . | . | . | . | . | . | . | . | . | . | . | . | . | . | . | . | . | . | . | . |
| T | . | . | . | . | . | . | . | . | . | . | . | . | . | . | . | . | . | . | . | . | . | . | . | . | . | . | T |
| . | . | . | . | . | . | . | . | . | . | . | . | . | . | . | . | . | . | . | . | . | . | . | . | . | . | . | . |
| . | . | . | . | . | . | . | . | . | . | . | . | . | . | . | . | . | . | . | . | . | . | . | . | . | . | . | . |
| . | . | . | . | . | . | . | . | . | . | . | . | . | . | . | . | . | . | . | . | . | . | . | . | . | . | . | . |
| . | . | . | . | . | . | . | . | . | . | . | . | . | . | . | . | . | . | . | . | . | . | . | . | . | . | . | . |
| . | . | . | . | . | . | . | . | . | . | . | . | . | . | . | . | . | . | . | . | . | . | . | . | . | . | . | . |
| . | . | . | . | . | . | . | . | . | . | . | . | . | . | . | . | . | . | . | . | . | . | . | . | . | . | . | T |
| . | . | . | . | . | . | . | . | . | . | . | . | . | . | . | . | . | . | . | . | . | . | . | . | . | . | . | T |
| . | . | . | . | . | . | . | . | . | . | . | . | . | . | . | . | . | . | . | . | . | . | . | . | . | . | . | T |
| T | . | . | . | . | . | . | . | . | . | . | . | . | . | . | . | . | . | . | . | . | . | . | . | . | . | . | T |
| . | . | . | . | . | . | . | . | . | . | . | . | . | . | . | . | . | . | . | . | . | . | . | . | . | . | . | . |
| T | . | . | . | . | . | . | . | . | . | . | . | . | . | . | . | . | . | . | . | . | . | . | . | . | . | . | T |
| T | . | . | . | . | . | . | . | . | . | . | . | . | . | . | . | . | . | . | . | . | . | . | . | . | . | . | T |
| . | . | . | . | . | . | . | . | . | A | . | . | . | . | . | . | . | . | . | . | . | . | . | . | . | . | . | . |
| T | . | . | . | . | . | . | . | . | . | . | . | . | . | . | . | . | . | . | . | . | . | . | . | . | . | . | T |
| . | . | . | . | . | . | . | . | . | . | . | . | . | . | . | . | . | . | . | . | . | . | . | . | . | . | . | . |
| . | . | . | . | . | . | . | . | . | . | . | . | . | . | . | . | . | . | . | . | . | . | . | . | . | . | . | . |
| . | . | . | . | . | . | . | . | . | . | . | . | . | . | . | C | . | . | . | . | . | . | . | . | . | . | . | . |
| . | . | . | . | . | . | . | . | . | . | . | . | . | . | . | . | . | . | . | . | . | . | . | . | . | . | . | . |
| . | . | . | . | . | . | . | . | . | . | . | . | . | . | . | . | . | . | . | . | . | . | . | . | . | . | . | . |
| . | . | . | . | . | . | . | . | . | . | . | . | . | . | . | . | . | . | . | . | . | . | . | . | . | . | . | . |
| . | . | . | . | . | . | . | . | . | . | . | . | . | . | . | . | . | . | . | . | . | . | . | . | . | . | . | . |
| . | . | . | . | . | . | . | . | . | . | . | . | . | . | . | . | . | . | . | . | . | . | . | . | . | . | . | . |
| . | . | . | . | . | . | . | . | . | . | . | . | . | . | . | . | . | . | . | . | . | . | . | . | . | . | . | . |
| . | . | . | . | . | . | . | . | . | . | . | . | . | . | . | C | . | . | . | . | . | . | . | . | . | . | . | . |
| . | . | . | . | . | . | . | . | . | . | . | . | . | . | . | . | . | . | . | . | . | . | . | . | . | . | . | . |
| . | . | . | . | . | . | . | . | . | . | . | . | . | . | . | . | . | . | . | . | . | . | . | . | . | . | . | . |
| . | . | . | . | . | . | . | . | . | . | . | . | . | . | . | . | . | . | . | . | . | . | . | . | . | . | . | . |
| . | . | . | . | . | . | . | . | . | . | . | . | . | . | . | . | . | . | . | . | . | . | . | . | . | . | . | . |
| . | . | . | . | . | . | . | . | . | . | . | . | . | . | . | . | . | . | . | . | . | . | . | . | . | . | . | . |
| . | . | . | . | . | . | . | . | . | . | . | . | . | . | . | . | . | . | . | . | . | . | . | . | . | . | . | . |
| . | . | . | . | . | . | . | . | . | . | . | . | . | . | . | . | . | . | . | . | . | . | . | . | . | . | . | . |
| . | . | . | . | . | . | . | . | . | A | . | . | . | . | . | . | . | . | . | . | . | . | . | . | . | . | . | . |
| . | . | . | . | . | . | . | . | . | . | . | . | . | . | . | . | . | . | . | . | . | . | . | . | . | . | . | . |
| . | . | . | . | . | . | . | . | . | . | . | . | . | . | . | . | . | . | . | . | . | . | . | . | . | . | . | . |
| . | . | . | . | . | . | . | . | . | . | . | . | . | . | . | . | . | . | . | . | . | . | . | . | . | . | . | . |
| . | . | . | . | . | . | . | . | . | . | . | . | . | . | . | . | . | . | . | . | . | . | . | . | . | . | . | . |
| . | . | . | . | . | . | . | . | . | . | . | . | . | . | . | . | . | . | . | . | . | . | . | . | . | . | . | . |

| | | | | | | | | | | | | | | | | | | | | | | | | | | | |
|---|---|---|---|---|---|---|---|---|---|---|---|---|---|---|---|---|---|---|---|---|---|---|---|---|---|---|---|
| . | . | . | . | . | . | . | . | . | . | . | . | . | . | . | . | . | . | . | . | . | . | . | . | . | . | . | . |
| . | . | . | . | . | . | . | . | . | . | . | . | . | . | . | . | . | . | . | . | . | . | . | . | . | . | . | . |
| . | . | . | . | . | . | . | . | . | . | . | . | . | . | . | . | . | . | . | . | . | . | . | . | . | . | . | . |
| . | . | . | . | . | . | . | . | . | . | . | . | . | . | . | . | . | . | . | . | . | . | . | . | . | . | . | . |
| . | . | . | . | . | . | . | . | . | . | . | . | . | T | . | . | . | . | . | . | . | . | . | . | . | . | . | . |
| . | . | . | . | . | . | . | . | . | . | . | . | . | T | . | . | . | . | . | . | . | . | . | . | . | . | . | . |
| . | . | . | . | . | . | . | . | . | . | . | . | . | T | . | . | . | . | . | . | . | . | . | . | . | . | . | . |
| . | . | . | . | . | . | . | . | . | . | . | . | . | T | . | . | . | . | . | . | . | . | . | . | . | . | . | . |
| . | . | . | . | . | . | . | . | . | . | . | . | . | T | . | . | . | . | . | . | . | . | . | . | . | . | . | . |
| . | . | . | . | . | . | . | . | . | . | . | . | . | T | . | . | . | . | . | . | . | . | . | . | . | . | . | . |
| . | . | . | . | . | . | . | . | . | . | . | . | . | . | . | . | . | . | . | . | . | . | . | . | . | . | . | . |
| . | . | . | . | . | . | . | . | . | . | . | . | . | . | . | . | . | . | . | . | . | . | . | . | . | . | . | . |
| . | . | . | . | . | . | . | . | . | . | . | . | . | . | . | . | . | . | . | . | . | . | . | . | . | . | . | . |
| . | . | . | . | . | A | . | . | . | . | . | . | . | . | . | . | . | . | . | . | . | . | . | . | . | . | . | . |
| . | . | . | . | . | A | . | . | . | . | . | . | . | . | . | . | . | . | . | . | . | . | . | . | . | . | . | . |
| . | . | . | . | . | A | . | . | . | . | . | . | . | . | . | . | . | . | . | . | . | . | . | . | . | . | . | . |
| . | . | . | . | . | A | . | . | . | . | . | . | . | . | . | . | . | . | . | . | . | . | . | . | . | . | . | . |
| . | . | . | . | . | A | . | . | . | . | . | . | . | . | . | . | . | . | . | . | . | . | . | . | . | . | . | . |
| . | . | . | . | . | A | . | . | . | . | . | . | . | . | . | . | . | . | . | . | . | . | . | . | . | . | . | . |
| . | . | . | . | . | A | . | . | . | . | . | . | . | . | . | . | . | . | . | . | . | . | . | . | . | . | . | . |
| . | . | . | . | . | A | . | . | . | . | . | . | . | . | . | . | . | . | . | . | . | . | . | . | . | . | . | . |
| . | . | . | . | . | . | . | . | A | A | . | . | . | . | . | . | . | . | . | . | . | . | . | . | . | . | . | . |
| . | . | . | . | . | . | . | . | A | A | . | . | . | . | . | . | . | . | . | . | . | . | . | T | . | . | . | . |
| . | . | . | . | . | . | . | . | . | . | . | G | . | . | G | . | . | G | . | . | . | . | . | . | . | . | . | . |
| . | . | . | . | . | . | . | . | . | . | . | G | . | . | G | . | . | G | . | . | . | . | . | . | . | . | . | . |
| . | . | . | . | . | . | . | . | . | A | . | . | . | . | . | . | . | . | . | . | . | . | . | . | . | . | . | . |
| . | . | . | . | . | . | . | . | . | A | . | . | . | . | . | . | . | . | . | . | . | . | . | . | . | . | . | . |
| . | . | . | . | . | . | . | . | . | A | . | . | . | . | . | . | . | . | . | . | . | . | . | . | . | . | . | . |
| . | . | . | . | . | . | . | . | A | A | . | . | . | . | . | . | . | . | . | . | . | . | . | . | . | . | . | . |
| . | . | . | . | . | . | . | . | . | A | . | . | . | . | . | . | . | . | . | . | . | . | . | . | . | . | . | . |
| . | . | . | . | . | . | . | . | . | A | . | . | . | . | . | . | . | . | . | . | . | . | . | . | . | . | . | . |
| . | . | . | . | . | . | . | . | . | A | . | . | . | . | . | . | . | . | . | . | . | . | . | . | . | . | . | . |
| . | . | . | . | . | . | . | . | A | A | . | . | . | . | . | . | . | . | . | . | . | . | . | . | . | . | . | . |
| . | . | . | . | . | . | . | . | A | A | . | . | . | . | . | . | . | . | . | . | . | . | . | T | . | . | . | . |
| . | . | . | . | . | . | . | . | . | A | . | . | . | . | . | . | . | . | . | . | . | . | . | . | . | . | . | . |
| . | . | . | . | . | . | . | . | A | A | . | . | . | . | . | . | . | . | . | . | . | . | . | . | . | . | . | . |
| . | . | . | . | . | . | . | . | . | A | . | . | . | . | . | . | . | . | . | . | . | . | . | . | . | . | . | . |
| . | . | . | . | . | . | . | . | . | . | . | . | . | . | . | . | . | . | . | . | . | . | . | . | . | . | . | . |
| . | . | . | . | . | . | . | . | . | . | . | . | . | . | . | . | . | . | . | . | . | . | . | . | . | . | . | . |
| . | . | . | . | . | . | . | . | . | . | . | . | . | . | . | . | . | . | . | . | . | . | . | . | . | . | . | . |
| . | . | . | . | . | . | . | . | . | . | . | . | . | . | . | . | . | . | . | . | . | . | . | . | . | . | . | . |
| . | . | . | . | . | . | . | . | . | . | . | . | . | . | . | . | . | . | . | . | . | . | . | . | . | . | . | . |
| . | . | . | . | . | . | . | . | . | . | . | . | . | . | . | . | . | . | . | . | . | . | . | . | . | . | . | . |
| . | . | . | . | . | . | . | . | . | . | . | . | . | . | . | . | . | . | . | . | . | . | . | . | C | . | . | . |
| . | . | . | . | . | . | T | . | . | . | . | . | . | . | . | . | . | . | . | . | . | . | . | . | . | . | . | . |
| . | . | . | . | . | . | . | . | . | . | . | . | . | . | . | . | . | . | . | . | . | . | . | . | . | . | . | . |
| . | . | . | . | . | . | . | . | . | . | . | . | . | . | . | . | . | . | . | . | . | . | . | . | . | . | . | . |
| . | . | . | . | . | . | . | . | . | . | . | . | . | . | . | . | . | . | . | . | . | . | . | . | . | . | . | . |
| . | . | . | . | . | . | . | . | . | . | . | . | . | . | . | . | . | . | . | . | . | . | . | . | . | . | . | . |
| . | . | . | . | . | . | . | . | . | . | . | . | . | . | . | . | . | . | . | . | . | . | . | . | . | . | . | . |
| . | . | . | . | . | . | . | . | . | . | . | . | . | . | . | . | . | . | . | . | . | . | . | . | . | . | . | . |
| . | . | . | . | . | . | . | . | . | . | . | . | . | . | . | . | . | . | . | . | . | . | . | . | . | . | . | . |
| . | . | . | . | . | . | . | . | . | . | . | . | . | . | . | . | . | . | . | . | . | . | . | . | . | . | . | . |
| . | . | . | . | . | . | . | . | . | . | . | . | . | . | . | . | . | . | . | . | . | . | . | . | . | . | . | . |
| . | . | . | . | . | . | . | . | . | . | . | . | . | . | . | . | . | . | . | . | . | . | . | . | . | . | . | . |
| . | . | . | . | . | . | . | . | . | . | . | . | . | . | . | . | . | . | . | . | . | . | . | . | . | . | . | . |
| . | . | . | . | . | . | . | . | . | . | . | . | . | . | . | . | . | . | . | . | . | . | . | . | . | . | . | . |
| . | . | . | . | . | . | . | . | . | . | . | . | . | . | . | . | . | . | . | . | . | . | . | . | . | . | . | . |
| . | . | . | . | . | . | . | . | . | . | . | . | . | . | . | . | . | . | . | . | . | . | . | . | . | . | . | . |
| . | . | . | . | . | . | . | . | . | . | . | . | . | . | . | . | . | . | . | . | . | . | . | . | . | . | . | . |
| . | . | . | . | . | A | . | . | . | . | . | . | . | . | . | . | . | . | . | . | . | . | . | . | . | . | . | . |

| | | | | | | | | | | | | | | | | | | | | | | | | | | | |
|---|---|---|---|---|---|---|---|---|---|---|---|---|---|---|---|---|---|---|---|---|---|---|---|---|---|---|---|
| . | . | . | . | . | A | . | . | . | . | . | . | . | . | . | . | . | . | . | . | . | . | . | . | . | . | . | . |
| . | . | . | . | . | . | . | . | . | . | . | . | . | . | . | . | . | . | . | . | . | . | . | . | C | . | . | . |
| . | . | . | . | . | . | . | . | . | . | . | . | . | . | . | . | . | . | . | . | . | . | . | . | . | . | . | . |
| . | . | . | . | . | . | . | . | . | . | . | . | . | . | . | . | . | . | . | . | . | . | . | . | . | . | . | . |
| . | . | . | . | . | . | . | . | . | A | . | . | . | . | . | . | . | . | . | . | . | . | . | . | . | . | . | . |
| . | . | . | . | . | . | . | . | . | A | . | . | . | . | . | . | . | . | . | . | . | . | . | . | . | . | . | . |
| . | . | . | . | . | . | . | . | . | M | . | . | . | . | . | . | . | . | . | . | . | . | . | . | . | . | . | . |
| . | . | . | . | . | . | . | . | . | . | . | . | . | . | . | . | . | . | . | . | . | . | . | . | . | . | . | . |
| . | . | . | . | . | . | . | . | M | A | . | . | . | . | . | . | . | . | . | . | . | . | . | . | . | . | . | . |
| . | . | . | . | . | . | . | . | M | M | . | . | . | . | . | . | . | . | . | . | . | . | . | . | . | . | . | . |
| . | . | . | . | . | . | . | . | M | A | . | . | . | . | . | . | . | . | . | . | . | . | . | . | . | . | . | . |
| . | . | . | . | . | . | . | . | M | A | . | . | . | . | . | . | . | . | . | . | . | . | . | . | . | . | . | . |
| . | . | . | . | . | . | . | . | M | A | . | . | . | . | . | . | . | . | . | . | . | . | . | . | . | . | . | . |
| . | . | . | . | . | . | . | . | M | A | . | . | . | . | . | . | . | . | . | . | . | . | . | . | . | . | . | . |
| . | . | . | . | . | . | . | . | M | A | . | . | . | . | . | . | . | . | . | . | . | . | . | . | . | . | . | . |
| . | . | . | . | . | . | . | . | M | A | . | . | . | . | . | . | . | . | . | . | . | . | . | . | . | . | . | . |
| . | . | . | . | . | . | . | . | M | A | . | . | . | . | . | . | . | . | . | . | . | . | . | . | . | . | . | . |
| . | . | . | . | . | . | . | . | M | M | . | . | . | . | . | . | . | . | . | . | . | . | . | . | . | . | . | . |
| . | . | . | . | . | . | . | . | M | A | . | . | . | . | . | . | . | . | . | . | . | . | . | . | . | . | . | . |
| . | . | . | . | . | . | . | . | M | A | . | . | . | . | . | . | . | . | . | . | . | . | . | . | . | . | . | . |
| . | . | . | . | . | . | . | . | M | A | . | . | . | . | . | . | t | . | . | . | . | . | . | . | . | . | . | . |
| . | . | . | . | . | . | . | . | M | A | . | . | . | . | . | . | . | . | . | . | . | . | . | . | . | . | . | . |
| . | . | . | . | . | . | . | . | M | A | . | . | . | . | . | . | . | . | . | . | . | . | . | . | . | . | . | . |
| . | . | . | . | . | . | . | . | M | A | . | . | . | . | . | . | . | . | . | . | . | . | . | . | . | . | . | . |
| . | . | . | . | . | . | . | . | M | A | . | . | . | . | . | . | . | . | . | . | . | . | . | . | . | . | . | . |
| . | . | . | . | . | . | . | . | M | A | . | . | . | . | . | . | . | . | . | . | . | . | . | . | . | . | . | . |
| . | . | . | . | . | . | . | . | M | A | . | . | . | . | . | . | . | . | . | . | . | . | . | . | . | . | . | . |
| . | . | . | . | . | . | . | . | M | A | . | . | . | . | . | . | . | . | . | . | . | . | . | . | . | . | . | . |
| . | . | . | . | . | . | . | . | M | A | . | . | . | . | . | . | . | . | . | . | . | . | . | . | . | . | . | . |
| . | . | . | . | . | . | . | . | M | A | . | . | . | . | . | . | . | . | . | . | . | . | . | . | . | . | . | . |
| . | . | . | . | . | . | . | . | M | A | . | . | . | . | . | . | . | . | . | . | . | . | . | . | . | . | . | . |
| . | T | . | . | . | . | . | . | . | . | . | . | . | . | . | . | . | . | . | . | . | . | . | . | . | . | . | . |
| . | T | . | . | . | . | . | . | . | . | . | . | . | . | . | . | . | . | . | . | . | . | . | . | . | . | . | . |
| . | . | . | . | . | . | . | . | . | . | . | . | . | . | . | . | . | . | . | . | . | . | . | . | . | . | . | . |
| . | . | . | . | . | . | . | . | . | . | . | . | . | . | . | . | . | . | . | . | . | . | . | . | . | . | . | . |
| . | . | . | . | . | . | . | . | . | . | . | . | . | . | . | . | . | . | . | . | . | . | . | . | . | . | . | T |
| . | . | . | . | . | . | . | . | . | . | . | . | . | . | . | . | . | . | . | . | . | . | . | . | . | . | . | . |
| . | . | . | . | . | . | . | . | . | . | . | . | . | . | . | . | . | . | . | . | . | . | . | . | . | . | . | . |
| . | . | . | . | . | . | . | . | . | . | . | . | . | . | . | . | . | . | . | . | . | . | . | . | . | . | . | . |
| . | . | . | . | . | . | . | . | . | . | . | . | . | . | . | . | . | . | . | . | . | . | . | . | . | . | . | . |
| . | . | . | . | . | . | . | . | . | . | . | . | . | . | . | . | . | . | . | . | . | . | . | . | . | . | . | . |
| . | . | . | . | . | . | . | . | . | . | . | . | . | . | . | . | . | . | . | . | . | . | . | . | . | . | . | . |
| . | . | . | . | . | . | . | . | . | . | . | . | . | . | . | . | . | . | . | . | . | . | . | . | . | . | . | . |
| . | . | . | . | . | . | . | . | . | . | . | . | . | . | . | . | . | . | . | . | . | . | . | . | . | . | . | . |
| . | . | . | . | . | . | . | . | . | . | . | . | . | . | . | . | . | . | . | . | . | . | . | . | . | . | . | . |
| . | . | . | . | . | . | . | . | . | . | . | . | . | . | . | . | . | . | . | . | . | . | . | . | . | . | . | . |
| . | . | . | . | . | . | . | . | . | . | . | . | . | . | . | . | . | . | . | . | . | . | . | . | . | . | . | . |
| . | . | . | . | . | . | . | . | . | . | . | . | . | . | . | . | . | . | . | . | . | . | . | . | . | . | . | . |
| . | . | . | . | . | . | . | . | . | . | . | . | . | . | . | . | . | . | . | . | . | . | . | . | . | . | . | . |
| . | . | . | . | T | . | . | . | . | . | . | . | . | . | . | . | . | . | . | . | . | . | . | . | . | A | . | . |
| . | . | . | . | T | . | . | . | . | . | . | . | . | . | . | . | . | . | . | . | . | . | . | . | . | A | . | . |
| . | . | . | . | . | . | . | . | . | . | . | . | . | . | . | . | . | . | . | . | . | . | . | . | . | . | . | . |
| . | . | . | . | . | . | . | . | . | . | . | . | . | . | . | . | . | . | . | . | . | . | . | . | . | . | . | . |
| . | . | . | . | . | . | . | . | . | . | . | . | . | . | . | . | . | . | . | . | . | . | . | . | . | . | . | . |
| . | . | . | . | . | . | . | . | . | . | . | . | . | . | . | . | . | . | . | . | . | . | . | . | . | . | . | . |
| . | . | . | . | . | . | . | . | . | . | . | . | . | . | . | . | . | . | . | . | . | . | . | . | . | . | . | . |
| . | . | . | . | . | . | . | . | . | . | . | . | . | . | . | . | . | . | . | . | . | . | . | . | . | . | . | . |
| . | . | . | . | . | . | . | . | . | . | . | . | . | . | . | . | . | . | . | . | . | . | . | . | . | . | . | . |
| . | . | . | . | . | . | . | . | . | . | . | . | . | . | . | . | . | . | . | . | . | . | . | . | . | . | . | . |

| 81 | 82 | 83 | 84 | 85 | 86 | 87 | 88 | 89 | 90 | 91 | 92 | 93 | 94 | 95 | 96 | 97 | 98 | 99 | 100 | 101 | 102 | 103 | 104 | 105 | 106 | 107 |
|---|---|---|---|---|---|---|---|---|---|---|---|---|---|---|---|---|---|---|---|---|---|---|---|---|---|---|
| ORF1a | ORF1a | ORF1a | ORF1a | ORF1a | ORF1a | ORF1a | ORF1a | ORF1a | ORF1a | ORF1a | ORF1a | ORF1a | ORF1a | ORF1a | ORF1a | ORF1a | ORF1a | ORF1a | ORF1a | ORF1a | ORF1a | ORF1a | ORF1a | ORF1a | ORF1a | ORF1a |
| 8917 | 8950 | 9389 | 9438 | 9628 | 9693 | 10138 | 10277 | 10369 | 10478 | 10479 | 10679 | 10702 | 10771 | 10815 | 11074 | 11083 | 11200 | 11306 | 11335 | 11457 | 11572 | 11620 | 12076 | 12439 | 12616 | 12685 |
| C | C | G | C | T | C | C | C | C | A | A | T | C | T | C | C | G | C | G | G | T | C | C | C | C | C | G |
| . | . | . | . | . | . | . | . | . | . | . | . | . | . | . | . | . | . | . | . | . | . | . | . | . | . | . |
| . | . | . | . | . | . | . | . | . | . | . | . | . | . | . | . | . | . | . | . | . | . | . | . | . | . | . |
| . | . | . | . | . | . | . | . | . | . | . | . | . | . | . | . | . | . | . | . | . | . | . | . | . | . | . |
| . | . | . | . | . | . | . | . | . | . | . | . | . | . | . | . | . | . | . | . | . | . | . | . | . | . | . |
| . | . | . | . | . | . | . | . | . | . | . | . | . | . | . | . | . | . | . | . | . | . | . | . | . | . | . |
| . | . | . | . | . | . | . | . | . | . | . | . | . | . | . | . | . | . | . | . | . | . | . | . | . | . | . |
| . | . | . | . | . | . | . | . | . | . | . | . | . | . | . | . | . | . | . | . | . | . | . | . | . | . | . |
| . | . | . | . | . | . | . | . | . | . | . | . | . | . | . | . | . | . | . | . | . | . | . | . | . | . | . |
| . | . | . | . | . | . | . | . | . | . | . | . | . | . | . | . | . | . | . | . | . | . | . | . | . | . | . |
| . | . | . | . | . | . | . | . | . | . | . | . | . | . | . | . | . | . | . | . | . | . | . | . | . | . | . |
| . | . | . | . | . | . | . | . | . | . | . | . | . | . | . | . | . | . | . | . | . | . | . | . | . | . | . |
| . | . | . | . | . | . | . | . | . | . | . | . | . | . | . | . | . | . | . | . | . | . | . | . | . | . | . |
| . | . | . | . | . | . | . | . | . | . | . | . | . | . | . | . | . | . | . | . | . | . | . | . | . | . | . |
| . | . | . | . | . | . | . | . | . | . | . | . | . | . | . | . | . | . | . | . | . | . | . | . | . | . | . |
| . | . | . | . | . | . | . | . | . | . | . | . | . | . | . | . | . | . | . | . | . | . | . | . | . | . | . |
| . | . | . | . | . | . | . | . | . | . | . | . | . | . | . | . | . | . | . | . | . | . | . | . | . | . | . |
| . | . | . | . | . | . | . | . | . | . | . | . | . | . | . | . | T | . | . | . | . | . | . | . | . | . | . |
| . | . | . | . | . | . | . | . | . | . | . | . | . | . | . | . | T | . | . | . | . | . | . | . | . | . | . |
| . | . | . | . | . | . | . | . | . | . | . | . | . | . | . | . | T | . | . | . | . | . | . | . | . | . | . |
| . | . | . | . | . | . | . | . | . | . | . | . | . | . | . | . | T | . | . | . | . | . | . | . | . | . | . |
| . | . | . | . | . | . | . | . | . | . | . | . | . | . | . | . | T | . | . | . | . | . | . | . | . | . | . |
| . | . | . | . | . | . | . | . | . | . | . | . | . | . | . | . | T | . | . | . | . | . | . | . | . | . | . |
| . | . | . | . | . | . | . | . | . | . | . | . | . | . | . | T | T | . | . | . | . | . | . | . | . | . | . |
| . | . | . | . | . | . | . | . | . | . | . | . | . | . | . | . | T | . | . | . | . | . | . | . | . | . | . |
| . | . | . | . | . | . | . | . | . | . | . | . | . | . | . | . | T | . | . | . | . | . | . | . | . | . | . |
| . | . | . | . | . | . | . | . | . | . | . | . | . | . | . | . | T | . | . | . | . | . | . | . | . | . | . |
| . | . | . | . | . | . | . | . | . | . | . | . | . | . | . | . | T | . | . | . | . | . | . | . | . | . | . |
| . | . | . | . | . | . | . | . | . | . | . | . | . | . | . | . | T | . | . | . | . | . | . | . | . | . | . |
| . | . | . | . | . | . | . | . | . | . | . | . | . | . | . | . | . | . | . | . | . | . | . | . | . | . | . |
| . | . | . | . | . | . | . | . | . | . | . | . | . | . | . | . | T | . | . | . | . | . | . | . | . | . | . |
| . | . | . | . | . | . | . | . | . | . | . | . | . | . | . | . | . | . | . | . | . | . | . | . | . | . | . |
| . | . | . | . | . | . | . | . | . | . | . | . | . | . | . | . | . | . | . | . | . | . | . | . | . | . | . |
| . | . | . | . | . | . | . | . | . | . | . | . | . | . | . | . | . | . | . | . | . | . | . | . | . | . | . |
| . | . | . | . | . | . | . | . | . | . | . | . | . | . | . | . | . | . | . | . | . | . | . | . | . | . | . |
| . | . | . | . | . | . | . | . | . | . | . | . | . | . | . | . | . | . | . | . | . | . | . | . | . | . | . |
| . | . | . | . | . | . | . | . | . | . | . | . | . | . | . | . | . | . | . | . | . | . | . | . | . | . | . |
| . | . | . | . | . | . | . | . | . | . | . | . | . | . | . | . | . | . | . | . | . | . | . | . | . | . | . |
| . | . | . | . | . | T | . | . | . | . | . | . | . | . | . | . | . | . | . | . | . | . | . | . | . | . | . |
| . | . | . | . | . | . | . | . | . | . | . | . | . | . | . | . | . | . | . | . | . | . | . | . | . | . | . |
| . | . | . | . | . | . | . | . | . | . | . | . | . | . | . | . | T | . | . | . | . | . | . | . | . | . | . |
| . | . | . | . | . | . | . | . | . | . | . | . | . | . | . | . | ? | . | . | . | . | . | . | . | . | . | . |
| . | . | . | . | . | . | . | . | . | . | . | . | . | . | . | . | . | . | . | . | . | . | . | . | . | . | . |
| . | . | . | . | . | . | . | . | . | . | . | . | . | . | . | . | . | . | . | . | . | . | . | . | . | . | . |
| . | . | . | . | . | . | . | . | . | . | . | . | . | . | . | . | . | . | . | . | . | . | . | . | . | . | . |
| . | . | . | . | . | . | . | . | . | . | . | . | . | . | . | . | . | . | . | . | . | . | . | . | . | . | . |
| T | . | . | . | . | . | . | . | . | . | . | . | . | . | . | . | . | . | . | . | . | . | . | . | . | . | . |
| . | . | . | . | . | . | . | . | . | . | . | . | . | . | . | . | . | . | . | . | . | . | . | . | . | . | . |
| . | . | . | . | . | . | . | . | . | . | . | . | . | . | . | . | . | . | . | . | . | . | . | . | . | . | . |
| . | . | . | . | . | . | . | . | . | . | . | . | . | . | . | . | . | . | . | . | . | . | . | . | . | . | . |
| . | . | . | . | . | . | . | . | . | . | . | . | . | . | . | . | . | . | . | . | . | . | . | . | . | . | . |
| . | . | . | . | . | . | . | . | . | . | . | . | . | . | . | . | . | . | . | . | . | . | . | . | . | . | . |
| . | . | . | . | . | . | . | . | . | . | . | . | . | . | . | . | T | . | . | . | . | . | . | . | . | . | . |
| . | . | . | . | . | . | . | . | . | . | . | . | . | . | . | . | . | . | . | . | . | . | . | . | . | . | . |
| . | . | . | . | . | . | . | . | . | . | . | . | . | . | . | . | . | . | . | . | . | . | . | . | . | . | . |
| . | . | . | . | . | . | . | . | . | . | . | . | . | . | . | . | . | . | . | . | . | . | . | . | . | . | . |

| | | | | | | | | | | | | | | | | | | | | | | | | | | |
|---|---|---|---|---|---|---|---|---|---|---|---|---|---|---|---|---|---|---|---|---|---|---|---|---|---|---|
| . | . | . | . | . | . | . | . | . | . | . | . | . | . | . | . | . | . | . | . | . | . | . | . | . | . | . |
| . | . | . | . | . | . | . | . | . | . | . | . | . | . | . | . | . | . | . | . | . | . | . | . | . | . | . |
| . | . | . | . | . | . | . | . | . | . | . | . | . | . | . | . | . | . | . | . | . | . | . | . | . | . | . |
| . | . | . | . | . | . | . | . | . | . | . | . | . | . | . | . | . | . | . | . | . | . | . | . | . | . | . |
| . | . | . | . | . | . | . | . | . | . | . | . | . | . | . | . | . | . | . | . | . | . | . | . | . | . | . |
| . | . | . | . | . | . | . | . | . | . | . | . | . | . | . | . | . | . | . | . | . | . | . | . | . | . | . |
| . | . | . | . | . | . | . | . | . | . | . | . | . | . | . | . | . | . | . | . | . | . | . | . | . | . | . |
| . | . | . | . | . | . | . | . | . | . | . | . | . | . | . | . | . | . | . | . | . | . | . | . | . | . | . |
| . | . | . | . | . | . | . | . | . | . | . | . | . | . | . | . | . | . | . | . | . | . | . | . | . | . | . |
| . | . | . | . | . | . | . | . | . | . | . | . | . | . | . | . | . | . | . | . | . | . | . | . | . | . | . |
| . | . | . | . | . | . | . | . | . | . | . | . | . | . | . | . | . | . | . | . | . | . | . | . | . | . | . |
| . | . | . | . | . | . | . | . | . | . | . | . | . | . | . | . | . | . | . | . | . | . | . | . | . | . | . |
| . | . | . | . | . | . | . | . | . | . | . | . | . | . | . | . | . | . | . | . | . | . | . | . | . | . | . |
| . | . | . | . | . | . | . | . | . | . | . | . | . | . | . | . | . | . | . | . | . | . | . | . | . | . | . |
| . | . | . | . | . | . | . | . | . | . | . | . | . | . | . | . | . | . | . | . | . | . | . | . | . | . | . |
| . | . | . | . | . | . | . | . | . | . | . | . | . | . | . | . | . | . | . | . | . | . | . | . | . | . | . |
| . | . | . | . | . | . | . | . | . | . | . | . | . | . | . | . | . | . | . | . | . | . | . | . | . | . | . |
| . | . | . | . | . | . | . | . | . | . | . | . | . | . | . | . | . | . | . | . | . | . | . | . | . | . | . |
| . | . | . | . | . | . | . | . | . | . | . | . | . | . | . | . | . | . | . | . | . | . | . | . | . | . | . |
| . | . | . | . | . | . | . | . | . | . | . | . | . | . | . | . | T | . | . | . | . | . | . | . | . | . | . |
| . | . | . | . | . | . | . | . | . | . | . | . | . | . | . | . | K | . | . | . | . | . | . | . | . | . | . |
| . | . | . | c | . | . | . | . | . | . | . | . | . | . | . | . | T | . | . | . | . | . | . | . | . | . | . |
| . | . | . | . | . | . | . | . | . | . | . | . | . | . | . | . | T | . | . | . | . | . | . | . | . | c | g |
| . | . | . | . | . | . | . | . | . | . | . | . | . | . | . | . | T | . | . | . | . | . | . | . | . | . | . |
| . | . | . | . | . | . | . | . | . | . | . | . | . | . | . | . | T | . | . | . | . | . | . | . | . | . | . |
| . | . | . | . | . | . | . | . | . | . | . | . | . | . | . | . | T | . | . | . | . | . | . | . | . | . | . |
| . | . | . | . | . | . | . | . | . | . | . | . | . | . | . | . | T | . | . | . | . | . | . | . | . | . | . |
| . | . | . | . | . | . | . | . | . | . | . | . | . | . | . | . | T | . | . | . | . | . | . | . | . | . | . |
| . | . | . | . | . | . | . | . | c | . | . | . | . | . | . | . | T | . | . | . | . | . | . | . | . | . | . |
| . | . | . | . | . | . | . | . | . | . | . | . | . | . | . | . | T | . | . | . | . | . | . | . | . | . | . |
| . | . | . | . | . | . | . | . | . | . | . | . | . | . | . | . | T | . | . | . | . | . | . | . | . | . | . |
| . | . | . | . | . | . | . | . | . | . | . | . | . | . | . | . | T | . | . | . | . | . | . | . | . | . | . |
| . | . | . | . | . | . | . | . | . | . | . | . | . | . | . | . | T | . | . | . | . | . | . | . | . | . | . |
| . | . | . | . | . | . | . | . | . | . | . | . | . | . | c | . | T | . | . | . | . | . | . | . | . | . | K |
| . | . | . | . | . | . | . | . | . | . | . | . | . | . | . | . | ? | . | . | . | . | . | . | . | . | . | . |
| . | . | . | . | . | . | . | . | . | . | . | . | . | . | . | . | T | . | . | . | . | . | . | . | . | . | . |
| . | . | . | . | . | . | . | . | . | . | . | . | . | . | . | . | T | . | . | . | . | . | . | . | . | . | . |
| . | . | . | . | . | . | . | . | . | . | . | . | . | . | . | . | T | . | . | . | . | . | . | . | . | . | . |
| . | . | . | . | . | . | . | . | . | . | . | . | . | . | . | . | T | . | . | . | . | . | . | . | . | . | . |
| . | . | . | . | . | . | . | . | . | . | . | . | . | . | . | . | T | . | . | . | . | . | . | . | . | . | . |
| . | . | . | . | . | . | . | . | . | . | . | . | . | . | . | . | T | . | . | . | . | . | . | . | . | . | . |
| . | . | . | . | . | . | . | . | . | . | . | . | . | . | . | . | T | . | . | . | . | . | . | . | . | . | . |
| . | . | . | . | . | . | . | . | . | . | . | . | . | . | . | . | T | . | . | . | . | . | . | . | . | . | . |
| . | . | . | . | . | . | . | . | . | . | . | . | . | . | . | . | T | . | . | . | . | . | . | . | . | . | . |
| . | . | . | . | . | . | . | . | . | . | . | . | . | . | . | . | T | . | . | . | . | . | . | . | . | . | . |
| . | . | . | . | . | . | . | . | . | . | . | . | . | . | . | . | T | . | . | . | . | . | . | . | . | . | . |
| . | . | . | . | . | . | . | . | . | . | . | . | . | . | . | . | T | . | . | . | . | . | . | . | . | . | . |
| . | . | . | . | . | . | . | . | . | . | . | . | . | . | . | . | T | . | . | . | . | . | . | . | . | . | . |
| . | . | . | . | . | . | . | . | . | . | . | . | . | . | . | . | T | . | . | . | . | . | . | . | . | . | . |
| . | . | . | . | . | . | . | . | . | . | . | . | . | . | . | . | . | . | . | . | . | . | . | . | . | . | . |
| . | . | . | . | . | . | . | . | . | . | . | . | . | . | . | . | . | . | . | . | . | . | . | . | . | . | . |
| . | . | . | . | . | . | . | . | . | . | . | . | . | . | . | . | T | . | . | . | . | . | . | . | . | . | . |
| . | . | . | . | . | . | . | . | . | . | . | . | . | . | . | . | . | . | . | . | . | . | . | . | . | . | . |
| . | . | . | . | . | . | . | . | . | . | . | . | . | . | . | . | T | . | . | . | . | . | . | . | . | . | . |
| . | . | . | . | . | . | . | . | . | . | . | . | . | . | . | . | K | . | . | . | . | . | . | . | . | . | . |
| . | . | . | . | . | . | . | . | . | . | . | . | . | . | . | . | T | . | . | . | . | . | . | . | . | . | . |
| . | . | . | . | . | . | . | . | . | . | . | . | . | . | . | . | T | . | . | . | . | . | . | . | . | . | . |
| . | . | . | . | . | . | . | . | . | . | . | . | . | . | . | . | T | . | . | . | . | . | . | . | . | . | . |
| . | . | . | . | . | . | . | . | . | . | . | . | . | . | . | . | T | . | . | . | . | . | . | . | . | . | . |
| . | . | . | . | . | . | . | . | . | . | . | . | . | . | . | . | T | . | . | . | . | . | . | . | . | . | . |
| . | . | . | . | . | . | . | . | . | . | . | . | . | . | . | . | T | . | . | . | . | . | . | . | . | . | T |
| . | . | . | . | . | . | . | . | . | . | . | . | . | . | . | . | ? | . | . | . | . | . | . | . | . | . | T |

| | | | | | | | | | | | | | | | | | | | | | | | | | | |
|---|---|---|---|---|---|---|---|---|---|---|---|---|---|---|---|---|---|---|---|---|---|---|---|---|---|---|
| . | . | . | . | . | . | . | . | . | . | . | . | . | . | . | . | T | . | . | . | . | . | . | . | . | . | . |
| . | . | . | . | . | . | . | c | . | . | . | . | . | . | . | . | T | . | . | . | . | . | . | . | . | . | T |
| . | . | . | . | . | . | . | . | . | . | . | . | . | . | . | . | T | . | . | . | . | . | . | . | . | . | . |
| . | . | . | . | . | . | . | . | . | . | . | . | . | . | . | . | ? | . | . | . | . | . | . | . | . | . | . |
| . | . | . | . | . | . | . | . | . | . | . | . | . | . | . | . | T | . | . | . | . | . | . | . | . | . | . |
| . | . | . | . | . | . | . | . | . | . | . | . | . | . | . | . | T | . | . | . | . | . | . | . | . | . | . |
| . | . | . | . | . | . | . | . | . | . | . | . | . | . | . | . | T | . | . | . | . | . | . | . | . | . | . |
| . | . | . | . | . | . | . | . | . | . | . | . | . | . | . | . | T | . | . | . | . | . | . | . | . | . | . |
| . | . | . | . | . | . | . | . | . | . | . | . | . | . | . | . | . | . | . | . | . | . | . | . | . | . | . |
| . | . | . | . | . | . | . | . | . | . | . | . | . | . | . | . | T | . | . | . | . | . | . | . | . | . | . |
| . | . | . | . | . | . | . | . | . | . | . | . | . | . | . | . | K | . | . | . | . | . | . | . | . | . | . |
| . | . | . | . | . | . | . | . | . | . | . | . | . | . | . | . | T | . | . | . | . | . | . | . | . | . | . |
| . | . | . | . | . | . | . | . | . | . | . | . | . | . | . | . | T | . | . | . | . | . | . | . | . | . | . |
| . | . | . | . | . | . | . | . | . | . | . | . | . | . | . | . | K | . | . | . | . | . | . | . | . | . | . |
| . | . | . | . | . | . | . | . | . | . | . | . | . | . | . | . | K | . | . | . | . | . | . | . | . | . | . |
| . | . | . | . | . | . | . | . | . | . | . | . | . | . | . | . | T | . | . | . | . | . | . | . | . | . | . |
| . | . | . | . | . | . | . | . | . | . | . | . | . | . | . | . | T | . | . | . | . | . | . | . | . | . | T |
| . | . | . | . | . | . | . | . | . | . | . | . | . | . | . | . | T | . | . | . | . | . | . | . | . | . | T |
| . | . | . | . | . | . | . | . | . | . | . | . | . | . | . | . | T | . | . | . | . | . | . | . | . | . | . |
| . | . | . | . | . | . | . | . | . | . | . | . | . | . | . | . | T | . | . | . | . | . | . | c | . | . | . |
| . | . | . | . | . | . | . | . | . | . | . | . | . | . | . | . | T | . | . | . | . | . | . | . | . | . | . |
| . | . | . | . | . | . | . | . | . | . | . | . | . | . | . | . | . | . | . | . | . | . | . | . | . | . | . |
| . | . | . | . | . | . | . | . | . | . | . | . | . | . | . | . | . | . | . | . | . | . | . | . | . | . | . |
| . | . | . | . | . | . | . | . | . | . | . | . | . | . | . | . | . | . | . | . | . | . | . | . | . | . | . |
| . | . | . | . | . | . | . | . | . | . | . | . | . | . | . | . | . | . | . | . | . | . | . | . | . | . | . |
| . | . | . | . | . | . | . | . | . | . | . | . | . | . | . | . | . | . | . | . | . | . | . | . | . | . | . |
| . | . | . | . | . | . | . | . | . | . | . | . | . | . | . | . | . | . | . | . | . | . | . | . | . | . | . |
| . | . | . | . | . | . | . | . | . | . | . | . | . | . | . | . | . | . | . | . | . | . | . | . | . | . | . |
| . | . | . | . | . | . | . | . | . | . | . | . | . | . | . | . | . | . | . | . | . | . | . | . | . | . | . |
| . | . | . | . | . | . | . | . | . | . | . | . | . | . | . | . | . | . | . | . | . | . | . | . | . | . | . |
| . | . | . | . | . | . | . | . | . | . | . | . | . | . | . | . | . | . | . | . | . | . | . | . | . | . | . |
| . | . | . | . | . | . | . | . | . | . | . | . | . | . | . | . | . | . | . | . | . | . | . | . | . | . | . |
| . | . | . | . | . | . | . | . | . | . | . | . | . | . | . | . | T | . | . | . | . | . | . | . | . | . | . |
| . | . | . | . | . | . | . | . | . | . | . | . | . | . | . | . | T | . | . | . | . | . | . | . | . | . | . |
| . | . | . | . | . | . | . | . | . | . | . | . | . | . | . | . | . | . | . | . | . | . | . | . | . | . | . |
| . | . | . | . | . | . | . | . | . | . | . | . | . | . | . | . | T | T | . | . | . | . | . | . | . | . | . |
| . | . | . | . | . | . | . | . | . | . | . | . | . | . | . | . | T | T | . | . | . | . | . | . | . | . | . |
| . | . | . | . | . | . | . | . | . | . | . | . | . | . | . | . | . | . | . | . | . | . | . | . | . | . | . |
| . | . | . | . | . | . | . | . | . | . | . | . | . | . | . | . | . | . | . | . | . | . | . | . | . | . | . |
| . | . | . | . | . | . | . | . | . | . | . | . | . | . | . | . | . | . | . | . | . | . | . | . | . | . | . |
| . | . | . | . | . | . | . | . | . | . | . | . | . | . | . | . | . | . | . | . | . | . | . | . | . | . | . |
| . | . | . | . | . | . | . | . | . | C | T | . | . | . | . | . | . | . | . | . | . | . | . | . | . | . | . |
| . | . | . | . | . | . | . | . | . | . | . | . | . | . | . | . | . | . | . | . | . | . | . | . | . | . | . |
| . | . | . | . | . | . | . | . | . | . | . | . | . | . | . | . | . | . | . | . | . | . | . | . | . | . | . |
| . | . | . | . | . | . | . | . | . | . | . | . | . | . | . | . | . | . | . | . | . | . | . | . | . | . | . |
| . | . | . | . | . | . | . | . | . | . | . | . | . | . | . | . | . | . | . | . | . | . | . | . | . | . | . |
| . | . | . | . | . | . | . | . | . | . | . | . | . | . | . | . | ? | . | . | . | . | . | . | . | . | . | . |
| . | . | . | . | . | . | . | . | . | . | . | . | . | . | . | . | . | . | . | . | . | . | . | . | . | . | . |
| . | . | . | . | . | . | . | . | . | . | . | . | . | . | . | . | . | . | . | . | . | . | . | . | . | . | . |
| . | . | . | . | . | . | . | . | . | . | . | . | . | . | . | . | T | . | . | . | . | . | . | . | . | . | . |
| . | . | . | . | . | . | . | . | . | . | . | . | . | . | . | . | T | . | . | . | . | . | . | . | . | . | . |
| . | . | . | . | . | . | . | T | . | . | . | . | . | . | . | . | . | . | . | . | . | . | . | . | . | . | . |
| . | . | . | . | . | . | . | T | . | . | . | . | . | . | . | . | . | . | . | . | . | . | . | . | . | . | . |
| . | . | . | . | . | . | . | . | . | . | . | . | . | . | . | . | . | . | . | . | . | . | . | . | . | . | . |
| . | . | . | . | . | . | . | . | . | . | . | . | . | . | . | . | . | . | . | . | . | . | . | . | . | . | . |
| . | . | . | . | . | . | . | . | . | . | . | . | . | . | . | . | . | . | . | . | . | . | . | . | . | . | . |
| . | . | . | . | . | . | . | . | . | . | . | . | . | . | . | . | . | . | . | . | . | . | . | . | . | . | . |
| . | . | . | . | . | . | . | . | . | . | . | . | . | . | . | . | . | . | . | . | . | . | . | . | . | . | . |
| . | . | . | . | . | . | . | . | . | . | . | . | . | . | . | . | . | . | . | . | . | . | . | . | . | . | . |
| . | . | . | . | . | . | . | . | . | . | . | . | . | . | . | . | . | . | . | . | . | . | . | . | . | . | . |
| . | . | . | . | . | . | . | . | . | . | . | . | . | . | . | . | . | . | . | . | . | . | . | . | . | . | . |
| . | . | . | . | . | . | . | . | . | . | . | . | . | . | . | . | . | . | . | . | . | . | . | . | . | . | . |

| | | | | | | | | | | | | | | | | | | | | | | | | | | |
|---|---|---|---|---|---|---|---|---|---|---|---|---|---|---|---|---|---|---|---|---|---|---|---|---|---|---|
| . | . | . | . | . | . | . | . | . | . | . | . | . | . | . | . | . | . | . | . | . | . | . | . | . | . | . |
| . | . | . | . | . | . | . | . | . | . | . | . | . | . | . | . | . | . | . | . | . | . | . | . | . | . | . |
| . | . | . | . | . | . | . | . | . | . | . | . | . | . | . | . | . | . | . | . | . | . | . | . | . | . | . |
| . | . | . | . | . | . | . | . | . | C | T | . | . | . | . | . | . | . | . | . | . | . | . | . | . | . | . |
| . | . | . | . | . | . | . | . | . | . | . | . | . | . | . | . | . | . | . | . | . | . | . | . | . | . | . |
| . | . | . | . | . | . | . | . | . | . | . | . | . | . | . | . | . | . | . | . | . | . | . | . | . | . | . |
| . | . | . | . | . | . | . | . | . | . | . | . | . | . | . | . | . | . | . | . | . | . | . | . | . | . | . |
| . | . | . | . | . | . | . | . | . | . | . | . | . | . | . | . | . | . | . | . | . | . | . | . | . | . | . |
| . | . | . | . | . | . | . | . | . | . | . | . | . | . | . | . | . | . | . | . | . | . | . | . | . | . | . |
| . | . | . | . | . | . | . | . | . | . | . | . | . | . | . | . | . | . | . | . | . | . | . | . | . | . | . |
| . | . | . | . | . | . | . | . | . | . | . | . | . | . | . | . | . | . | . | . | . | . | . | . | . | . | . |
| . | . | . | . | . | . | . | . | . | . | . | . | . | . | . | . | . | . | . | . | . | . | . | . | . | . | . |
| . | . | . | . | . | . | . | . | . | . | . | . | . | . | . | . | . | . | . | . | . | . | . | . | . | . | . |
| . | . | . | . | . | . | . | . | . | . | . | . | . | . | . | . | . | . | . | . | . | . | . | . | . | . | . |
| . | . | . | . | . | . | . | . | . | . | . | . | . | . | . | . | . | . | . | . | . | . | . | . | . | . | . |
| . | . | . | . | . | . | . | . | . | . | . | . | . | . | . | . | . | . | . | . | . | . | . | . | . | . | . |
| . | . | . | T | . | . | . | . | . | . | . | . | . | . | . | . | . | . | . | . | . | . | . | . | . | . | . |
| . | . | . | . | . | . | . | . | . | . | . | . | . | . | . | . | . | . | . | . | . | . | . | . | . | T | . |
| . | . | . | . | . | . | . | . | . | . | . | . | . | . | . | . | . | . | . | . | . | . | . | . | . | T | . |
| . | . | . | . | . | . | . | . | . | . | . | . | . | . | . | . | . | . | . | . | . | . | . | . | . | . | . |
| . | . | . | . | . | . | . | . | . | . | . | . | . | . | . | . | . | . | . | . | . | . | . | . | . | . | . |
| . | . | . | . | . | . | . | . | . | . | . | . | . | . | . | . | . | . | . | . | . | . | . | . | . | . | . |
| . | . | . | . | . | . | . | . | . | . | . | . | . | . | . | . | . | . | . | . | . | . | . | . | . | . | . |
| . | . | . | . | . | . | . | . | . | . | . | . | . | . | . | . | . | . | . | . | . | . | . | . | . | . | . |
| . | . | . | . | . | . | . | . | . | . | . | . | . | . | . | . | . | . | . | . | . | . | . | . | . | . | . |
| . | . | . | . | . | . | . | . | . | . | . | . | . | . | . | . | . | . | . | . | . | . | . | . | . | . | . |
| . | . | . | . | . | . | . | . | . | . | . | . | . | . | . | . | . | . | . | . | . | . | . | . | . | . | . |
| . | . | . | . | . | . | . | . | . | . | . | . | . | . | . | . | . | . | . | . | . | . | . | . | . | . | . |
| . | . | . | . | . | . | . | . | . | . | . | . | . | . | . | . | . | . | . | . | . | . | . | . | . | . | . |
| . | . | . | . | . | . | . | . | . | . | . | . | . | . | . | . | . | . | . | . | . | . | . | . | . | . | . |
| . | . | . | . | . | . | . | . | . | . | . | . | . | . | . | . | . | . | . | . | . | . | . | . | . | . | . |
| . | . | . | . | . | . | . | . | . | . | . | . | . | . | . | . | . | . | . | . | . | . | . | . | . | . | . |
| . | . | . | . | . | . | . | . | . | . | . | . | . | . | . | . | . | . | . | . | . | . | . | . | . | . | . |
| . | . | . | . | . | . | . | . | . | . | . | . | . | . | . | . | . | . | . | . | . | . | . | . | . | . | . |
| . | . | . | . | . | . | . | . | . | . | . | . | . | . | . | . | . | . | . | . | . | . | . | . | . | . | . |
| . | . | . | . | . | . | . | . | . | . | . | . | . | . | . | . | . | . | . | . | . | . | . | . | . | . | . |
| . | . | . | . | . | . | . | . | . | . | . | . | . | . | . | . | . | . | . | . | . | . | . | . | . | . | . |
| . | . | . | . | . | . | . | . | . | . | . | . | . | . | . | . | . | . | . | . | . | . | . | . | . | . | . |
| . | . | . | . | . | . | . | . | . | . | . | . | . | . | . | . | . | . | . | . | . | . | . | . | . | . | . |
| . | . | . | . | . | . | . | . | . | . | . | . | . | . | . | . | . | . | . | . | . | . | . | . | . | . | . |
| . | . | . | . | . | . | . | . | . | . | . | . | . | . | . | . | . | . | . | . | . | . | . | . | . | . | . |
| . | . | . | . | . | . | . | . | . | . | . | . | . | . | . | . | . | . | . | . | . | . | . | . | . | . | . |
| . | . | . | . | . | . | . | . | . | . | . | . | . | . | . | . | . | . | . | . | . | . | . | . | . | . | . |
| . | . | . | . | . | . | . | . | . | . | . | . | . | . | . | . | . | . | . | . | . | . | . | . | . | T | . |
| . | . | . | . | . | . | . | . | . | . | . | . | . | . | . | . | . | . | . | . | . | . | . | . | . | T | . |
| . | . | . | . | . | . | . | . | . | C | T | . | . | . | . | . | . | . | . | . | . | . | . | . | . | . | . |
| . | . | . | . | . | . | . | . | . | . | . | . | . | . | . | . | . | . | . | . | . | . | . | . | . | . | . |
| . | . | . | . | . | . | . | . | . | C | T | . | . | . | . | . | . | . | . | . | . | . | . | . | . | . | . |
| . | . | . | . | . | . | . | . | . | C | T | . | . | . | . | . | . | . | . | . | . | . | . | . | . | . | . |
| . | . | . | . | . | . | . | . | . | . | . | . | . | . | . | . | . | . | . | . | . | . | . | . | . | T | . |
| . | . | . | . | . | . | . | . | . | . | . | . | . | . | . | . | T | . | . | . | . | . | . | . | . | T | . |
| . | . | . | . | . | . | . | . | . | . | . | . | . | . | . | . | T | . | . | . | . | . | . | . | . | T | . |
| . | . | . | . | . | . | . | . | . | . | . | . | . | . | . | . | . | . | . | . | . | . | . | . | . | T | . |
| . | . | . | T | . | . | . | . | . | . | . | . | . | . | . | . | . | . | . | . | . | . | . | . | . | . | . |
| . | . | . | T | . | . | . | . | . | . | . | . | . | . | . | . | . | . | . | . | . | . | . | . | . | . | . |
| . | . | . | . | . | . | . | . | . | . | . | . | . | . | . | . | . | . | . | . | . | . | . | . | . | . | . |
| . | . | . | . | . | . | . | . | . | . | . | . | . | . | . | . | . | . | . | . | . | . | . | . | . | . | . |
| . | . | . | . | . | . | . | . | . | . | . | . | . | . | . | . | . | . | . | . | . | . | . | . | . | . | . |
| . | . | . | . | . | . | . | . | . | . | . | . | . | . | . | . | . | . | . | . | . | . | . | . | . | . | . |
| . | . | . | . | . | . | . | . | . | . | . | . | . | . | . | . | . | . | . | . | . | . | . | . | . | . | . |
| . | . | . | . | . | . | . | . | . | . | . | . | . | . | . | . | . | . | . | . | . | . | . | . | . | . | . |

| | | | | | | | | | | | | | | | | | | | | | | | | | | |
|---|---|---|---|---|---|---|---|---|---|---|---|---|---|---|---|---|---|---|---|---|---|---|---|---|---|---|
| . | . | . | . | . | . | . | . | . | . | . | . | . | . | . | . | . | . | . | . | . | . | . | . | . | . | . |
| . | . | . | . | . | . | . | . | . | . | . | . | . | . | . | . | . | . | . | . | . | . | . | . | . | . | . |
| . | . | . | . | . | . | . | . | . | . | . | . | . | . | . | . | . | . | . | . | . | . | . | . | . | . | . |
| . | . | . | . | . | . | . | . | . | . | . | . | . | . | . | . | . | . | . | . | . | . | . | . | . | . | . |
| . | . | . | . | . | . | . | . | . | . | . | . | . | . | . | . | T | . | . | . | . | . | . | . | . | . | . |
| . | . | . | . | . | . | . | . | . | . | . | . | . | . | . | . | T | . | . | . | . | . | . | . | . | . | . |
| . | . | . | . | . | . | . | . | . | . | . | . | . | . | . | . | . | . | . | . | . | . | . | . | . | . | . |
| . | . | . | . | . | . | . | . | . | . | . | . | . | . | . | . | T | . | . | . | . | . | . | . | . | . | . |
| . | . | . | . | . | . | . | . | . | . | . | . | . | . | . | . | T | . | . | . | . | . | . | . | . | . | . |
| . | . | . | . | . | . | . | . | . | . | . | . | . | . | . | . | T | . | . | . | . | . | . | . | . | . | . |
| . | . | . | . | . | . | . | . | . | . | . | . | . | . | . | . | T | . | . | . | . | . | . | . | . | . | . |
| . | . | . | . | . | . | . | . | . | . | . | . | . | . | . | . | T | . | . | . | . | . | . | . | . | . | . |
| . | . | . | . | . | . | . | . | . | . | . | . | . | . | . | . | T | . | . | . | . | . | . | . | . | . | . |
| . | . | . | . | . | . | . | . | . | . | . | . | . | . | . | . | . | . | . | . | . | . | . | . | . | . | . |
| . | . | . | . | . | . | . | . | . | . | . | . | . | . | . | . | T | . | . | . | . | . | . | . | . | . | . |
| . | . | . | . | . | . | . | . | . | . | . | . | . | . | . | . | T | . | . | . | . | . | . | . | . | . | . |
| . | . | . | . | . | . | . | . | . | . | . | . | . | . | . | . | T | . | . | . | . | . | . | . | . | . | . |
| . | . | . | . | . | . | . | . | . | . | . | . | . | . | . | . | T | . | . | . | . | . | . | . | . | . | . |
| . | . | . | . | . | . | . | . | . | . | . | . | . | . | . | . | T | . | . | . | . | . | . | . | . | . | . |
| . | . | . | . | . | . | . | . | . | . | . | . | . | . | . | . | T | . | . | . | . | . | . | . | . | . | . |
| . | . | . | . | . | . | . | . | . | . | . | . | . | . | . | . | T | . | . | . | . | . | . | . | . | . | . |
| . | . | . | . | . | . | . | . | . | . | . | . | . | . | . | . | T | . | . | . | . | . | . | . | . | . | . |
| . | . | . | . | . | . | . | . | . | . | . | . | . | . | . | . | T | . | . | . | . | . | . | . | . | . | . |
| . | . | . | . | . | . | . | . | . | . | . | . | . | . | . | . | T | . | . | . | . | . | . | . | . | . | . |
| . | . | . | . | . | . | . | . | . | . | . | . | . | . | . | . | T | . | . | . | . | . | . | . | . | . | . |
| . | . | . | . | . | . | . | . | . | . | . | . | . | . | . | . | T | . | . | . | . | . | . | . | . | . | . |
| . | . | . | . | . | . | . | . | . | . | . | . | . | . | . | . | T | . | . | . | . | . | . | . | . | . | . |
| . | . | . | . | . | . | . | . | . | . | . | . | . | . | . | . | T | . | . | . | . | . | . | . | . | . | . |
| . | . | . | . | . | . | . | . | . | . | . | . | . | . | . | . | T | . | . | . | . | . | . | . | . | . | . |
| . | . | . | . | . | . | . | . | . | . | . | . | . | . | . | . | T | . | . | . | . | . | . | . | . | . | . |
| . | . | . | . | . | . | . | . | . | . | . | . | . | . | . | . | T | . | . | . | . | . | . | . | . | . | . |
| . | . | . | . | . | . | . | . | . | . | . | . | . | . | . | . | A | . | . | . | . | . | . | . | . | . | . |
| . | . | . | . | . | . | . | . | . | . | . | . | . | . | . | . | T | . | . | . | . | . | . | . | . | . | . |
| . | . | . | . | . | . | . | . | . | . | . | . | . | . | . | . | T | . | . | . | . | . | . | . | . | . | . |
| . | . | . | . | . | . | . | . | . | . | . | . | . | . | . | . | T | . | . | . | . | . | . | . | . | . | . |
| T | . | . | . | . | . | . | . | . | . | . | . | . | . | . | . | . | . | . | . | . | . | . | . | . | . | . |
| . | . | . | . | . | . | . | . | . | . | . | . | . | . | . | . | T | . | . | . | . | . | . | . | . | . | . |
| . | . | . | . | . | . | . | . | . | . | . | . | . | . | . | . | T | T | . | . | . | . | . | . | . | . | . |
| . | . | . | . | . | . | . | . | . | . | . | . | . | . | . | . | T | . | . | . | . | . | . | . | . | . | . |
| . | . | . | . | . | . | . | . | . | . | . | . | . | . | . | . | T | . | . | . | . | . | . | . | . | . | . |
| . | . | . | . | . | . | . | . | . | . | . | . | . | . | . | . | T | . | . | . | . | . | . | . | . | . | . |
| . | . | . | . | . | . | . | . | . | . | . | . | . | . | . | . | T | . | . | . | . | . | . | . | . | . | . |
| . | . | . | . | . | . | . | . | . | . | . | . | . | . | . | . | T | . | . | . | . | . | . | . | . | . | . |
| . | . | . | . | . | . | . | . | . | . | . | . | . | . | . | . | T | . | . | . | . | . | . | . | . | . | . |
| . | . | . | . | . | . | . | . | . | . | . | . | . | . | . | . | T | . | . | . | . | . | . | . | . | . | . |
| . | . | . | . | . | . | . | . | . | . | . | . | . | . | . | . | T | . | . | . | . | . | . | . | . | . | . |
| . | . | . | . | . | . | . | . | . | . | . | . | . | . | . | . | T | . | . | . | . | . | . | . | . | . | T |
| . | . | . | . | . | . | . | . | . | . | . | . | . | . | . | . | T | . | . | . | . | . | . | . | . | . | . |
| . | . | . | . | . | . | . | . | . | . | . | . | . | . | . | . | T | . | . | . | . | . | . | . | . | . | . |
| . | . | . | . | . | . | . | . | . | . | . | . | . | . | . | . | T | . | . | . | . | . | . | . | . | . | . |
| . | . | . | . | . | . | . | . | . | . | . | . | . | . | . | . | T | . | . | . | . | . | . | . | . | . | . |
| . | . | . | . | . | . | . | . | . | . | . | . | . | . | . | . | T | . | . | . | . | . | . | . | . | . | . |
| . | . | . | . | . | . | . | . | . | . | . | . | . | . | . | . | T | . | . | . | . | . | . | . | . | . | . |
| . | . | . | . | . | . | . | . | . | . | . | . | . | . | . | . | T | . | . | . | . | . | . | . | . | . | . |
| . | . | . | . | . | . | . | . | . | . | . | . | . | . | . | . | T | . | . | . | . | . | . | . | . | . | . |
| . | . | . | . | . | . | . | . | . | . | . | . | . | . | . | . | T | . | . | . | . | . | . | . | . | . | . |
| . | . | . | . | . | . | . | . | . | . | . | . | . | . | . | . | T | . | . | . | . | . | . | . | . | . | . |
| . | . | . | . | . | . | . | . | . | . | . | . | . | . | . | . | T | . | . | . | . | . | . | . | . | . | . |
| . | . | . | . | . | . | . | . | . | . | . | . | . | . | . | . | . | . | . | . | . | . | . | . | . | . | . |
| . | . | . | . | . | . | . | . | . | . | . | . | . | . | . | . | . | . | . | . | . | T | . | . | . | . | . |
| . | . | . | . | . | . | . | . | . | . | . | . | . | . | . | . | T | . | . | . | . | . | . | . | . | . | . |
| . | . | . | . | . | . | . | . | . | . | . | . | . | . | . | . | T | . | . | . | . | . | . | . | . | . | . |

| | | | | | | | | | | | | | | | | | | | | | | | | | | |
|---|---|---|---|---|---|---|---|---|---|---|---|---|---|---|---|---|---|---|---|---|---|---|---|---|---|---|
| . | . | . | . | . | . | . | . | . | . | . | . | . | . | . | . | T | . | . | . | . | . | . | . | . | . | . |
| . | . | . | . | . | . | . | . | . | . | . | . | . | . | . | . | T | . | . | . | . | . | . | . | . | . | . |
| . | . | . | . | . | . | . | . | . | . | . | . | . | . | . | . | T | . | . | . | . | . | . | . | . | . | . |
| . | . | . | . | . | . | . | . | . | . | . | . | . | . | . | . | T | . | . | . | . | . | . | . | . | . | . |
| . | . | . | . | . | . | . | . | . | . | . | . | . | . | . | . | T | . | . | . | . | . | . | . | . | . | . |
| . | . | . | . | . | . | . | . | . | . | . | . | . | . | . | . | T | . | . | . | . | . | . | . | . | . | . |
| . | . | . | . | . | . | . | . | . | . | . | . | . | . | . | . | . | . | . | . | . | . | . | . | . | . | . |
| . | . | . | . | . | . | . | . | . | . | . | . | . | . | . | . | . | . | . | . | . | . | . | . | . | . | . |
| . | . | . | . | . | . | . | . | . | . | . | . | . | . | . | . | . | . | . | . | . | . | . | . | . | T | . |
| . | . | . | . | . | . | . | . | . | . | . | . | . | . | . | . | T | . | . | . | . | . | . | . | . | . | . |
| . | . | . | . | . | . | . | . | . | . | . | . | . | . | . | . | T | . | . | . | . | . | . | . | . | . | . |
| . | . | . | . | . | . | . | . | . | . | . | . | . | . | . | . | T | . | . | . | . | . | . | . | . | . | . |
| . | . | . | . | . | . | . | . | . | . | . | . | . | . | . | . | T | . | . | . | . | . | . | . | . | . | . |
| . | . | . | . | . | . | . | . | . | . | . | . | . | . | . | . | . | . | . | . | . | . | . | . | . | . | . |
| . | . | . | . | . | . | . | . | . | . | . | . | . | . | . | . | . | . | . | . | . | . | . | . | . | . | . |
| . | . | . | . | . | . | . | . | . | . | . | . | . | . | . | . | . | . | A | . | . | . | . | . | . | . | . |
| . | . | . | . | . | . | . | . | . | . | . | . | . | . | . | . | . | . | A | . | . | . | . | . | . | . | . |
| . | . | . | . | . | . | . | . | . | . | . | . | . | . | . | . | T | . | . | . | . | . | . | . | . | . | . |
| . | . | . | . | . | . | . | . | . | . | . | . | . | . | . | . | . | . | . | . | . | . | . | . | . | . | . |
| . | . | . | . | . | . | . | . | . | . | . | . | . | . | . | . | . | . | . | . | . | . | . | . | . | . | . |
| . | . | . | . | . | . | . | . | . | . | . | . | . | . | . | . | . | . | . | . | . | . | . | . | . | . | . |
| . | . | . | . | . | . | . | . | . | . | . | . | . | . | . | . | . | . | . | . | . | . | . | . | . | . | . |
| . | . | . | . | . | . | . | . | . | . | . | . | . | . | . | . | T | . | . | . | . | . | . | . | . | . | . |
| . | . | . | . | . | . | . | . | . | . | . | . | . | . | . | . | . | . | . | . | . | . | . | . | . | . | . |
| . | . | . | . | . | . | . | . | . | . | . | . | . | . | . | . | T | . | . | . | . | . | . | . | . | . | . |
| . | . | . | . | . | . | . | . | . | . | . | . | . | . | . | . | . | . | . | . | . | . | . | . | . | . | . |
| . | . | . | . | . | . | . | . | . | . | . | . | . | . | . | . | . | . | . | . | . | . | . | . | . | . | . |
| . | . | . | . | . | . | . | . | . | . | . | . | . | . | . | . | T | . | . | . | . | . | . | . | . | . | . |
| . | . | . | . | . | . | . | . | . | . | . | . | . | . | . | . | . | . | . | . | . | . | . | . | . | . | . |
| . | . | . | . | . | . | . | . | . | . | . | . | . | . | . | . | T | . | . | . | . | . | . | . | . | . | . |
| . | . | . | . | . | . | . | . | . | . | . | . | . | . | . | . | . | . | . | . | . | . | . | T | . | . | . |
| . | . | . | . | . | . | . | . | . | . | . | . | . | . | . | . | . | . | . | . | . | . | . | . | . | . | . |
| . | . | . | . | . | . | . | . | . | . | . | . | . | . | . | . | T | . | . | . | . | . | . | . | . | . | . |
| c | c | g | c | t | c | c | c | c | a | a | t | c | t | c | c | g | c | g | g | t | c | c | c | c | c | g |
| c | c | g | c | t | c | c | c | c | a | a | t | c | t | c | c | g | c | g | g | t | c | c | c | c | c | g |
| c | c | g | c | c | c | c | c | c | a | a | t | c | t | c | c | g | c | g | g | t | c | c | c | c | c | g |
| c | c | g | c | c | c | c | c | c | a | a | t | c | t | c | c | g | c | g | g | t | c | c | c | c | c | g |
| c | c | g | c | t | c | c | c | c | a | a | t | c | t | c | c | g | c | g | g | t | c | c | c | c | c | g |
| c | c | g | c | t | c | c | c | c | a | a | t | c | t | c | c | g | c | g | g | t | c | c | c | c | c | g |
| c | c | g | c | t | c | c | c | c | a | a | t | c | t | c | c | g | c | g | g | t | c | c | c | c | c | g |
| c | c | g | c | t | c | c | c | c | a | a | t | c | t | c | c | g | c | g | g | t | c | c | c | c | c | g |
| c | c | g | c | t | c | c | c | c | a | a | t | c | t | c | c | g | c | g | g | t | c | c | c | c | c | g |
| c | c | g | c | t | c | c | c | c | a | a | t | c | c | c | c | g | c | g | g | t | c | c | c | c | c | g |
| c | c | g | c | t | c | c | c | c | a | a | t | c | t | c | c | g | c | g | g | t | c | c | c | c | c | g |
| c | c | g | c | t | c | c | c | c | a | a | t | c | t | c | c | g | c | g | g | t | c | c | c | c | c | g |
| c | c | g | c | t | c | c | c | c | a | a | t | c | t | c | c | g | c | g | g | t | c | c | c | c | c | g |
| c | c | g | c | t | c | c | c | c | a | a | t | c | t | c | c | g | c | g | g | t | c | c | c | c | c | g |
| c | c | g | c | t | c | c | c | c | a | a | t | c | t | c | c | t | c | g | g | t | c | c | c | c | c | g |
| c | c | g | c | t | c | c | c | c | a | a | t | c | t | c | c | t | c | g | g | t | c | c | c | c | c | g |
| c | c | g | c | t | c | c | c | c | a | a | t | c | t | c | c | g | c | g | g | t | c | c | c | c | c | g |
| c | c | g | c | t | c | c | c | c | a | a | t | c | t | c | c | g | c | g | g | t | c | c | c | c | c | g |
| c | c | g | c | t | c | c | c | c | a | a | t | c | t | c | c | g | c | g | g | t | c | c | c | c | c | g |
| c | c | g | c | t | c | c | c | c | a | a | t | c | t | c | c | t | c | g | g | t | c | c | c | c | c | g |
| c | c | g | c | t | c | c | c | c | a | a | t | c | t | c | c | g | c | g | g | t | c | c | c | c | c | g |
| c | c | g | c | t | c | c | c | c | a | a | t | c | t | c | c | t | c | g | g | t | c | c | c | c | c | g |
| c | c | g | c | t | c | c | c | c | a | a | t | c | t | c | c | g | c | g | g | t | c | c | c | c | c | g |
| c | c | g | c | t | c | c | c | c | a | a | t | c | t | c | c | g | c | g | g | t | c | c | c | c | c | g |
| . | . | . | . | . | . | . | . | . | . | . | . | . | . | . | . | . | . | . | . | . | . | . | . | . | . | . |
| . | . | . | . | . | . | . | . | . | . | . | . | . | . | . | . | . | . | . | . | . | . | . | . | . | . | . |
| . | . | . | . | . | . | . | . | . | . | . | . | . | . | . | . | . | . | . | . | . | . | . | . | . | . | . |
| . | . | . | . | . | . | . | . | . | . | . | . | . | . | . | Y | T | . | . | . | . | . | . | . | . | . | . |
| . | . | . | . | . | . | . | . | . | . | . | . | . | . | . | . | . | . | . | . | . | . | . | . | . | . | . |

| | | | | | | | | | | | | | | | | | | | | | | | | | | |
|---|---|---|---|---|---|---|---|---|---|---|---|---|---|---|---|---|---|---|---|---|---|---|---|---|---|---|
| . | . | . | . | . | . | . | . | . | . | . | . | . | . | . | . | T | . | . | . | . | . | . | . | . | . | . |
| . | . | . | . | . | . | . | . | . | . | . | . | . | . | . | . | . | . | . | . | . | . | . | . | . | . | . |
| . | . | . | . | . | . | . | . | . | . | . | . | . | . | . | . | T | . | . | . | . | . | . | . | . | . | . |
| . | . | . | . | . | . | . | . | . | . | . | . | . | . | . | . | T | . | . | . | . | . | . | . | . | . | . |
| . | . | . | . | . | . | . | . | . | . | . | . | . | . | . | . | . | . | . | . | . | . | . | . | . | . | . |
| . | . | . | . | . | . | . | . | . | . | . | . | . | . | . | . | T | . | . | . | . | . | . | . | . | . | . |
| . | . | . | . | . | . | . | . | . | . | . | . | . | . | . | . | . | . | . | . | . | . | . | . | . | . | . |
| . | . | . | . | . | . | . | . | . | . | . | . | . | . | . | . | . | . | . | . | . | . | . | . | . | . | . |
| . | . | . | . | . | . | . | . | . | . | . | . | . | . | . | . | . | . | . | . | . | . | . | . | . | . | . |
| . | . | . | . | . | . | . | . | . | . | . | . | . | . | . | . | . | . | . | . | . | . | . | . | . | . | . |
| . | . | . | . | . | . | . | . | . | . | . | . | . | . | . | . | . | . | . | . | . | . | . | . | . | . | . |
| . | . | . | . | . | . | . | . | . | . | . | . | . | . | . | . | . | . | . | . | . | . | . | . | . | . | . |
| . | . | . | . | . | . | . | . | . | . | . | . | . | . | . | . | . | . | . | . | . | . | . | . | . | . | . |
| . | . | . | . | . | . | . | . | . | . | . | . | . | . | . | T | T | . | . | . | . | . | . | . | . | . | . |
| . | . | . | . | . | . | . | . | . | . | . | . | . | . | . | . | T | . | . | . | . | . | . | . | . | . | . |
| . | . | . | . | . | . | . | . | . | . | . | . | . | . | . | . | T | . | . | . | . | . | . | . | . | . | . |
| . | . | . | . | . | . | . | . | . | . | . | . | . | . | . | . | T | . | . | . | . | . | . | . | . | . | . |
| . | . | . | . | . | . | . | . | . | . | . | . | . | . | . | . | T | . | . | . | . | . | . | . | . | . | . |
| . | . | . | . | . | . | . | . | . | . | . | . | . | . | . | . | T | . | . | . | . | . | . | . | . | . | . |
| . | . | . | . | . | . | . | . | . | . | . | . | . | . | . | . | T | . | . | . | . | . | . | . | . | . | . |
| . | . | . | . | . | . | . | . | . | . | . | . | . | . | . | . | T | . | . | . | . | . | . | . | . | . | . |
| . | . | . | . | . | . | . | . | . | . | . | . | . | . | . | . | T | . | . | . | . | . | . | . | . | . | . |
| . | . | . | . | . | . | . | . | . | . | . | . | . | . | . | . | . | . | . | . | . | . | . | . | . | . | . |
| . | . | . | . | . | . | . | . | . | . | . | . | . | . | . | . | . | . | . | . | . | . | . | . | . | . | . |
| . | . | . | . | . | . | . | . | . | . | . | . | . | . | . | . | . | . | . | . | . | . | . | . | . | . | . |
| . | . | . | . | . | . | . | . | . | . | . | . | . | . | . | . | . | . | . | . | . | . | . | . | . | . | . |
| . | . | . | . | . | . | . | . | . | . | . | . | . | . | . | . | . | . | . | . | . | . | . | . | . | . | . |
| . | . | . | . | . | . | . | . | . | . | . | . | . | . | . | . | . | . | . | . | . | . | . | . | . | . | . |
| . | . | . | . | . | . | . | . | . | . | . | . | . | C | . | . | . | . | . | . | . | . | . | . | . | . | . |
| . | . | . | . | . | . | . | . | . | . | . | . | . | . | . | . | T | . | . | . | . | . | . | . | . | . | . |
| . | . | . | . | . | . | . | . | . | . | . | . | . | . | . | . | . | . | . | . | . | . | . | . | . | . | . |
| . | . | . | . | . | . | . | . | . | . | . | . | . | . | . | . | . | . | . | . | . | . | . | . | . | . | . |
| . | . | . | . | . | . | . | . | . | . | . | . | . | . | . | . | T | . | . | . | . | . | . | . | . | . | . |
| . | . | . | . | . | . | . | . | . | . | . | . | . | . | . | . | K | . | . | . | . | . | . | . | . | . | . |
| . | . | . | . | . | . | . | . | . | . | . | . | . | . | . | . | T | . | . | . | . | . | . | . | . | . | . |
| . | . | . | . | . | . | . | . | . | . | . | . | . | . | . | . | T | . | . | . | . | . | . | . | . | . | . |
| . | . | . | . | . | . | . | . | . | . | . | . | . | . | . | . | T | . | . | . | . | . | . | . | . | . | . |
| . | . | . | . | . | . | . | . | . | . | . | . | . | . | . | . | T | . | . | . | . | . | . | . | . | . | . |
| . | . | . | . | . | . | . | . | . | . | . | . | . | . | . | . | T | . | . | . | . | . | . | . | . | . | . |
| . | . | . | . | . | . | . | . | . | . | . | . | . | . | . | . | T | . | . | . | . | . | . | . | . | . | . |
| . | . | . | . | . | . | . | . | . | . | . | . | . | . | . | . | . | . | A | . | . | . | . | . | . | . | . |
| . | . | . | . | . | . | . | . | . | . | . | . | . | . | . | . | . | . | A | . | . | . | . | . | . | . | . |
| . | . | . | . | . | . | . | . | . | . | . | . | . | . | . | . | . | . | . | T | . | . | T | . | . | . | . |
| . | . | . | . | . | . | . | . | . | . | . | . | . | . | . | . | . | . | . | . | . | . | . | . | . | . | . |
| . | . | . | . | . | . | . | . | . | . | . | . | . | . | . | . | . | . | . | . | . | . | . | . | . | . | . |
| . | . | . | . | . | . | . | . | . | . | . | . | . | . | . | . | . | . | . | . | . | . | . | . | . | . | . |
| . | . | . | . | . | . | . | . | . | . | . | . | . | . | . | . | . | . | . | . | . | . | T | . | . | . | . |
| . | . | . | . | . | . | . | . | . | . | . | . | . | . | . | . | . | . | . | . | . | . | . | . | . | . | . |
| . | . | . | . | . | . | . | . | . | . | . | . | . | . | . | . | . | . | . | T | . | . | T | . | . | . | . |
| . | . | . | . | . | . | . | . | . | . | . | . | . | . | . | . | . | . | . | . | . | . | . | . | . | . | . |
| . | . | . | . | . | . | . | . | . | . | . | . | . | . | . | . | . | . | . | . | . | . | . | . | . | . | . |
| . | . | . | . | . | . | . | . | . | . | . | . | . | . | . | . | . | . | . | . | . | . | . | . | . | . | . |
| . | . | . | . | . | . | . | . | . | . | . | . | . | . | . | . | . | . | . | . | . | . | T | . | . | . | . |
| . | . | . | . | . | . | . | . | . | . | . | . | . | . | . | . | . | . | . | . | . | . | . | . | . | . | . |
| . | . | . | . | . | . | . | . | . | . | . | . | . | . | . | . | . | . | . | . | . | . | . | . | . | . | . |
| . | . | . | . | . | . | . | . | . | . | . | . | . | . | . | . | . | . | . | . | . | . | . | . | . | . | . |
| . | . | . | . | . | . | . | . | . | . | . | . | . | . | . | . | . | . | . | . | . | . | . | . | . | . | . |
| . | . | . | . | . | . | . | . | . | . | . | . | . | . | . | . | . | . | . | . | . | . | . | . | . | . | . |
| . | . | . | . | . | . | . | . | . | . | . | . | . | . | . | . | . | . | . | . | . | . | . | . | . | . | . |
| . | . | . | . | . | . | . | . | . | . | . | . | . | . | . | . | . | . | . | . | . | . | . | . | . | . | . |
| . | . | . | . | . | . | . | . | . | . | . | . | . | . | . | . | . | . | . | . | . | . | . | . | . | . | . |
| . | . | . | . | . | . | . | . | . | . | . | . | . | . | . | . | . | . | . | . | . | . | . | . | . | . | . |

| | | | | | | | | | | | | | | | | | | | | | | | | | | |
|---|---|---|---|---|---|---|---|---|---|---|---|---|---|---|---|---|---|---|---|---|---|---|---|---|---|---|
| . | . | . | . | . | . | . | . | . | . | . | . | . | . | . | . | . | . | . | . | . | . | . | . | . | . | . |
| . | . | . | . | . | . | . | . | . | . | . | . | . | . | . | . | . | . | . | . | . | . | . | . | . | . | . |
| . | . | . | . | . | . | . | . | . | . | . | . | . | . | . | . | . | . | . | . | . | . | . | . | . | . | . |
| . | T | . | . | . | . | . | . | . | . | . | . | . | . | . | . | . | . | . | . | . | . | . | . | . | . | . |
| . | . | . | . | . | . | . | . | . | . | . | . | . | . | . | . | . | . | . | . | . | . | . | . | . | . | . |
| . | . | . | . | . | . | . | . | . | . | . | . | . | . | . | . | T | . | . | . | . | . | . | . | . | . | . |
| . | . | . | . | . | . | . | . | . | . | . | . | . | . | . | . | . | . | . | . | . | . | . | . | . | . | . |
| . | . | . | . | . | . | . | . | . | . | . | . | . | . | . | . | . | . | . | . | . | . | . | . | . | . | . |
| . | . | . | . | . | . | . | . | . | . | . | . | . | . | . | . | . | . | . | . | . | . | . | . | . | . | . |
| . | . | . | . | . | . | . | . | . | . | . | . | . | . | . | . | . | . | . | . | . | . | . | . | . | . | . |
| . | . | . | . | . | . | . | . | . | . | . | . | . | . | . | . | . | . | . | . | . | . | . | . | . | . | . |
| . | . | . | . | . | . | . | . | . | . | . | . | . | . | . | . | . | . | . | . | . | . | . | . | T | . | . |
| . | . | . | . | . | . | . | . | . | . | . | . | . | . | . | . | . | . | . | . | . | . | . | . | . | . | . |
| . | . | . | . | . | . | . | . | . | . | . | . | . | . | . | . | . | . | . | . | . | T | . | . | . | . | . |
| . | . | . | . | . | . | . | . | . | . | . | . | . | . | . | . | . | . | . | . | . | . | . | . | . | . | . |
| . | . | . | . | . | . | . | . | . | . | . | . | . | . | . | . | . | . | . | . | . | . | . | . | . | . | . |
| . | . | . | . | . | . | . | . | . | . | . | . | . | . | . | . | . | . | . | . | . | . | . | . | . | . | . |
| . | . | . | . | . | . | . | . | . | . | . | . | . | . | . | . | . | . | . | . | . | . | . | . | T | . | . |
| . | . | . | . | . | . | . | . | . | . | . | . | . | . | . | . | . | . | . | . | . | . | . | . | T | . | . |
| . | T | . | . | . | . | . | . | . | . | . | . | . | . | . | . | . | . | . | . | . | . | . | . | . | . | . |
| . | . | . | . | . | . | . | . | . | . | . | . | . | . | . | . | . | . | . | . | . | . | . | . | T | . | . |
| . | . | . | . | . | . | . | . | . | . | . | . | . | . | . | . | . | . | . | . | . | . | . | . | . | . | . |
| . | . | . | . | . | . | . | . | . | . | . | . | . | . | . | . | . | . | . | . | . | . | . | . | . | . | . |
| . | . | . | . | . | . | . | . | . | . | . | C | . | . | . | . | . | . | . | . | . | . | . | . | . | . | . |
| . | . | . | . | . | . | T | . | . | . | . | . | . | . | . | . | . | . | . | . | . | . | . | . | . | . | . |
| . | . | . | . | . | . | . | . | . | . | . | . | . | . | . | . | . | . | . | . | . | . | . | . | . | . | . |
| . | . | . | . | . | . | . | . | . | . | . | . | . | . | . | . | . | . | . | . | . | . | . | . | . | . | . |
| . | . | . | . | . | . | . | . | . | . | . | . | . | . | . | . | . | . | . | . | . | . | . | . | . | . | . |
| . | . | . | . | . | . | . | . | . | . | . | . | . | . | . | . | . | . | . | . | . | . | . | . | . | . | . |
| . | . | . | . | . | . | . | . | . | . | . | C | . | . | . | . | . | . | . | . | . | . | . | . | . | . | . |
| . | . | . | . | . | . | . | . | . | . | . | . | . | . | . | . | . | . | . | . | . | . | . | . | . | . | . |
| . | . | . | . | . | . | . | . | . | . | . | . | . | . | . | . | . | . | . | . | . | . | . | . | . | . | . |
| . | . | . | . | . | . | . | . | . | . | . | . | . | . | . | . | . | . | . | . | . | . | . | . | . | . | . |
| . | . | . | . | . | . | T | . | . | . | . | . | T | . | . | . | . | . | . | . | . | . | . | . | . | . | . |
| . | . | . | . | . | . | . | . | . | . | . | . | . | . | . | . | . | . | . | . | . | . | . | . | . | . | . |
| . | . | . | . | . | . | T | . | . | . | . | . | T | . | . | . | . | . | . | . | . | . | . | . | . | . | . |
| . | . | . | . | . | . | T | . | . | . | . | . | T | . | . | . | . | . | . | . | . | . | . | . | . | . | . |
| . | . | . | . | . | . | . | . | . | . | . | . | . | . | . | . | T | T | . | . | . | . | . | . | . | . | . |
| . | . | . | . | . | . | T | . | . | . | . | . | T | . | . | . | . | . | . | . | . | . | . | . | . | . | . |
| . | . | . | . | . | . | . | . | . | . | . | . | . | . | . | . | . | . | . | . | . | . | . | . | . | . | . |
| . | . | . | . | . | . | . | . | . | . | . | . | . | . | . | . | . | . | . | . | . | . | . | . | . | . | . |
| . | . | . | . | . | . | . | . | . | . | . | . | . | . | . | . | . | . | . | . | . | . | . | . | . | . | . |
| . | . | . | . | . | . | . | . | . | . | . | . | . | . | . | . | . | . | . | . | . | . | . | . | . | . | . |
| . | . | . | . | . | . | . | . | . | . | . | . | . | . | . | . | . | . | . | . | . | . | . | . | . | . | . |
| . | . | . | . | . | . | . | . | . | . | . | . | . | . | . | . | . | . | . | . | . | . | . | . | . | . | . |
| . | . | . | . | . | . | . | . | . | . | . | . | . | . | . | . | . | . | . | . | . | . | . | . | . | . | . |
| . | . | . | . | . | . | . | . | . | . | . | . | . | . | . | . | . | . | . | . | . | . | . | . | . | . | . |
| . | . | . | . | . | . | . | . | . | . | . | . | . | . | . | . | . | . | . | . | . | . | . | . | . | . | . |
| . | . | . | . | . | . | . | . | . | . | . | . | . | . | . | . | . | . | . | . | . | . | . | . | . | . | . |
| . | . | . | . | . | . | . | . | . | . | . | . | . | . | . | . | . | . | . | . | . | . | . | . | . | . | . |
| . | . | . | . | . | . | . | . | . | . | . | . | . | . | . | . | . | . | . | . | . | . | . | . | . | . | . |
| T | . | . | . | . | . | . | . | . | . | . | . | . | . | . | . | . | . | . | . | . | . | . | . | . | . | . |
| T | . | A | . | . | . | . | . | . | . | . | . | . | . | . | . | . | . | . | . | . | . | . | . | . | . | . |
| T | . | . | . | . | . | . | . | . | . | . | . | . | . | . | . | . | . | . | . | . | . | . | . | . | . | . |
| T | . | A | . | . | . | . | . | . | . | . | . | . | . | . | . | . | . | . | . | . | . | . | . | . | . | . |
| T | . | A | . | . | . | . | . | . | . | . | . | . | . | . | . | . | . | . | . | . | . | . | . | . | . | . |
| . | . | . | . | . | . | . | . | . | . | . | . | . | . | . | . | T | . | . | . | . | . | . | . | . | . | . |
| T | . | . | . | . | . | . | . | . | . | . | . | . | . | . | . | . | . | . | . | . | . | . | . | . | . | . |
| T | . | . | . | . | . | . | . | . | . | . | . | . | . | . | . | . | . | . | . | . | . | . | . | . | . | . |
| T | . | . | . | . | . | . | . | . | . | . | . | . | . | . | . | . | . | . | . | . | . | . | . | . | . | . |
| T | . | . | . | . | . | . | . | . | . | . | . | . | . | . | . | . | . | . | . | . | . | . | . | . | . | . |
| T | . | A | . | . | . | . | . | . | . | . | . | . | . | . | . | . | . | . | . | . | . | . | . | . | . | . |

```
T . . . . . . . . . . . . . . . . . . . . . . . . . .
T . . . . . . . . . . . . . . . . . . . . . . . . . .
T . . . . . . . . . . . . . . . . . . . . . . . . . .
T . . . . . . . . . . . . . . . . . . . . . . . . . .
. . . . . . . . . . . . . . . . . . . . . . . . T . .
. . . . . . . . . . . . . . . . . . . . . . . . T . .
. . . . . . . . . . . . . . . . . . . . . . . . . . .
. . . . . . . . . . . . . . . . . . . . . . . . T . .
. . . . . . . . . . . . . . . . . . . . . . . . . . .
. . . . . . . . . . . . . . . . . . . . . . . . T . .
. . . . . . . . . . . . . . . . . . . . . . . . . . .
. . . . . . . . . . . . . . . . . . . . . . . . . . .
. . . . . . . . . . . . . . . . . . . . . . . . . . .
. . . . . . . . . . . . . . . . . . . . . . . . . . .
. . . . . T . . . . . . . . . . . . . . . . . . . . .
. . . . . T . . . . . . . . T . . . . . . . . . . . .
. . . . . T . . . . . . . . T . . . . . . . . . . . .
. . . . . T . . . . . . . . . . . . . . . . . . . . .
. . . . . T . . . . . . . . . . . . . . . . . . . . .
. . . . . T . . . . . . . . . . . . . . . . . . . . .
. . . . . T . . . . . . . . . . . . . . . . . . . . .
. . . . . . . . . . . . . . . . T . . . . . . . . . .
. . . . . . . . . . . . . . . . T . . . . . . . . . .
. . . . . . . . . . . . . . . . . . . . . . . . . . .
. . . . . . . . . . . . . . . . . . . . . . . . . . .
. . . . . . . . . . . . . . . . T . . . . . . . . . .
. . . . . . . . . . . . . . . . T . . . . . . . . . .
. . . . . . . . . . . . . . . . T . . . . . . . . . .
. . . . . . . . . . . . . . . . T . . . . . . . . . .
. . . . . . . . . . . . . . . . T . . . . . . . . . .
. . . . . . . . . . . . . . . . T . . . . . . . . . .
. . . . . . . . . . . . . . . . T . . . . . . . . . .
. . . . . . . . . . . . . . . . T . . . . . . . . . .
. . . . . . . . . . . . . . . . T . . . . . . . . . .
. . . . . . . . . . . . . . . . T . . . . . . . . . .
. . . . . . . . . . . . . . . . T . . . . . . . . . .
. . . . . . . . . . . . . . . . A . . . . . . . . . .
. . . . . . . . . . . . . . . . . . . . . . . . . . .
. . . . . . . T . . . . . . . . . . . . . . . T . . .
. . . . . . . . . . . . . . . . . . . . . . . . . . .
. . . . . . . . . . . . . . . . . . . . . . . . . . .
. . . . . . . . . . . . . . . . . . . . . . . . . . .
. . . . . . . . T . . . . . . . . . . . . . . . . . .
. . . . . . . . T . . . . . . . . . . . . . . . . . .
. . . . . . . . . . . . . . . . . . . . . . . . . . .
. . . . . . . . . . . . . . . . . . . . . . . . . . .
. . . . . . . . . . . . . . . . . . . . . . . . . . .
. . . . . . . . . . . . . . . . . . . . . . . . . . .
. . . . . . . . . . . . . . . . . . . . . . . . . . .
. . . . . . . . . . . . . . . . . . . . . . . . . . .
. . . . . . . . . . . . . . . . . . . . . . . . . . .
. . . . . . . . . . . . . . . . . . . . . . . T . . .
. . . . . . . . . . . . . . . . . . . . . . . . . . .
. . . . . . . . . . . . . . . . . . . . . . . . . . .
. . . . . . . . . . . . . . . . . . . . . . . . . . .
. . . . . . . . . . . . . . . . . . . . . . . . . . .
. . . T . . . . . . . . . . . . . . . . . . . . . . .
. . . T . . . . . . . . . . . . . . . . . . . . . . .
. . . T . . . . . . . . . . . . . . . . . . . . . . .
. . . T . . . . . . . . . . . . . . . . . . . . . . .
. . . . . . . . . . . . . . . . . . . . . . . . . . .
. . . . . . . . . . . . . . . . . . . . . . . . . . .
```

| | | | | | | | | | | | | | | | | | | | | | | | | | | |
|---|---|---|---|---|---|---|---|---|---|---|---|---|---|---|---|---|---|---|---|---|---|---|---|---|---|---|
| . | . | . | . | . | . | . | . | . | . | . | . | . | . | . | . | . | . | . | . | . | . | . | . | . | . | . |
| . | . | . | . | . | . | . | . | . | . | . | . | . | . | . | . | . | . | . | . | . | . | . | T | . | . | . |
| . | . | . | . | . | . | . | . | . | . | . | . | . | . | . | . | . | . | . | . | . | . | . | . | . | . | . |
| . | . | . | . | . | . | . | . | . | . | . | . | . | . | . | . | . | . | . | . | . | . | . | . | . | . | . |
| . | . | . | . | . | . | . | . | . | . | . | . | . | . | . | . | T | . | . | . | . | . | . | . | . | . | . |
| . | . | . | . | . | . | . | . | . | . | . | . | . | . | . | . | T | . | . | . | . | . | . | . | . | . | . |
| . | . | . | . | . | . | . | . | . | . | . | . | . | . | . | . | K | . | . | . | . | . | . | . | . | . | . |
| . | . | . | . | . | . | . | . | . | . | . | . | . | . | . | . | . | . | . | . | . | . | . | . | . | . | . |
| . | . | . | . | . | . | . | . | . | . | . | . | . | . | . | . | T | . | . | . | . | . | . | . | . | . | K |
| . | . | . | . | . | . | . | . | . | . | . | . | . | . | . | . | K | . | . | . | . | . | . | . | . | . | K |
| . | . | . | . | . | . | . | . | . | . | . | . | . | . | . | . | T | . | . | . | . | . | . | . | . | . | K |
| . | . | . | . | . | . | . | . | . | . | . | . | . | . | . | . | T | . | . | . | . | . | . | . | . | . | K |
| . | . | . | . | . | . | . | . | . | . | . | . | . | . | . | . | T | . | . | . | . | . | . | . | . | . | K |
| . | . | . | . | . | . | . | . | . | . | . | . | . | . | . | . | T | . | . | . | . | . | . | . | . | . | K |
| . | . | . | . | . | . | . | . | . | . | . | . | . | . | . | . | T | . | . | . | . | . | . | . | . | . | K |
| . | . | . | . | . | . | . | . | . | . | . | . | . | . | . | . | T | . | . | . | . | . | . | . | . | . | K |
| . | . | . | . | . | . | . | . | . | . | . | . | . | . | . | . | T | . | . | . | . | . | . | . | . | . | K |
| . | . | . | . | . | . | . | . | . | . | . | . | . | . | . | . | K | . | . | . | . | . | . | . | . | . | ? |
| . | . | . | . | . | . | . | . | . | . | . | . | . | . | . | . | T | . | . | . | . | . | . | . | . | . | K |
| . | . | . | . | . | . | . | . | . | . | . | . | . | . | . | . | T | . | . | . | . | . | . | . | . | . | K |
| . | . | . | . | . | . | . | . | . | . | . | . | . | . | . | . | T | . | . | . | . | . | . | . | . | . | K |
| . | . | . | . | . | . | . | . | . | . | . | . | . | . | . | . | T | . | . | . | . | . | . | . | . | . | K |
| . | . | . | . | . | . | . | . | . | . | . | . | . | . | . | . | T | . | . | . | . | . | . | . | . | . | K |
| . | . | . | . | . | . | . | . | . | . | . | . | . | . | . | . | T | . | . | . | . | . | . | . | . | . | K |
| . | . | . | . | . | . | . | . | . | . | . | . | . | . | . | . | T | . | . | . | . | . | . | . | . | . | K |
| . | . | . | . | . | . | . | . | . | . | . | . | . | . | . | . | T | . | . | . | . | . | . | . | . | . | K |
| . | . | . | . | . | . | . | . | . | . | . | . | . | . | . | . | T | . | . | . | . | . | . | . | . | . | K |
| . | . | . | . | . | . | . | . | . | . | . | . | . | . | . | . | T | . | . | . | . | . | . | . | . | . | K |
| . | . | . | . | . | . | . | . | . | . | . | . | . | . | . | . | T | . | . | . | . | . | . | . | . | . | K |
| . | . | . | . | . | . | . | . | . | . | . | . | . | . | . | . | T | . | . | . | . | . | . | . | . | . | K |
| . | . | . | . | . | . | . | . | . | . | . | . | . | . | . | . | T | . | . | . | . | . | . | . | . | . | K |
| . | . | . | . | . | . | . | . | . | . | . | . | . | . | . | . | . | . | . | . | . | . | . | . | . | . | . |
| . | . | . | . | . | . | . | . | . | . | . | . | . | . | . | . | . | . | . | . | . | . | . | . | . | . | . |
| T | . | . | . | . | . | . | . | . | . | . | . | . | . | . | . | . | . | . | . | . | . | T | . | . | . | . |
| . | . | . | . | . | . | . | . | . | . | . | . | . | . | . | . | . | . | . | . | . | . | . | . | . | . | . |
| . | . | . | . | . | . | . | . | . | . | . | . | . | . | . | . | . | . | . | . | . | . | . | . | . | . | . |
| . | . | . | . | . | . | . | . | . | . | . | . | . | . | . | . | . | . | . | . | . | . | . | . | . | . | . |
| . | . | . | . | . | . | . | . | . | . | . | . | . | . | . | . | . | . | . | . | . | . | . | . | . | . | . |
| . | . | . | . | . | . | . | . | . | . | . | . | . | . | . | . | . | . | . | . | . | . | . | . | . | . | . |
| . | . | . | . | . | . | . | . | . | . | . | . | . | . | . | . | . | . | . | . | . | . | . | . | . | . | . |
| . | . | . | . | . | . | . | . | . | . | . | . | . | . | . | . | . | . | . | . | . | . | . | . | . | . | . |
| . | . | . | . | . | . | . | . | . | . | . | . | . | . | . | . | . | . | . | . | . | . | . | . | . | . | . |
| . | . | . | . | . | . | . | . | . | . | . | . | . | . | . | . | . | . | . | . | . | . | . | . | . | . | . |
| . | . | . | . | . | . | . | . | . | . | . | . | . | . | . | . | . | . | . | . | . | . | . | . | . | . | . |
| . | . | . | . | . | . | . | . | . | . | . | . | . | . | . | . | . | . | . | . | . | . | . | . | . | . | . |
| . | . | . | . | . | . | . | . | . | . | . | . | . | . | . | . | . | . | . | . | . | . | . | . | . | . | . |
| . | . | . | . | . | . | . | . | . | . | . | . | . | . | . | . | . | . | . | . | . | . | . | . | . | . | . |
| . | . | . | . | . | . | . | . | . | . | . | . | . | . | . | . | . | . | . | . | . | . | . | . | . | . | . |
| . | . | . | . | . | . | . | . | . | . | . | . | . | . | . | . | . | . | . | . | . | . | . | . | . | . | . |
| . | . | . | . | . | . | . | . | . | . | . | . | . | . | . | . | . | . | . | . | . | . | . | . | . | . | . |
| . | . | . | . | . | . | . | . | . | . | . | . | . | . | . | . | . | . | . | . | . | . | . | . | . | . | . |
| . | . | . | . | . | . | . | . | . | . | . | . | . | . | . | . | . | . | . | . | . | . | . | . | . | . | . |
| . | . | . | . | . | . | . | . | . | . | . | . | . | . | . | . | . | . | . | . | . | . | . | . | . | . | . |
| . | . | . | . | . | . | . | . | . | . | . | . | . | . | . | . | . | . | . | . | . | . | . | . | . | . | . |
| . | . | . | . | . | . | . | . | . | . | . | . | . | . | . | . | . | . | . | . | C | . | . | . | . | . | . |
| . | . | . | . | . | . | . | . | . | . | . | . | . | . | . | . | . | . | . | . | C | . | . | . | . | . | . |
| . | . | . | . | . | . | . | . | . | . | . | . | . | . | . | . | . | . | . | . | . | . | . | . | . | . | . |
| . | . | . | . | . | . | . | . | . | . | . | . | . | . | . | . | . | . | . | . | . | . | . | . | . | . | . |
| . | . | . | . | . | . | . | . | . | . | . | . | . | . | . | . | . | . | . | . | . | . | . | . | . | . | . |
| . | . | . | . | . | . | . | . | . | . | . | . | . | . | . | . | . | . | . | . | . | . | . | . | . | . | . |

| 108 | 109 | 110 | 111 | 112 | 113 | 114 | 115 | 116 | 117 | 118 | 119 | 120 | 121 | 122 | 123 | 124 | 125 | 126 | 127 | 128 | 129 | 130 | 131 | 132 | 133 |
|---|---|---|---|---|---|---|---|---|---|---|---|---|---|---|---|---|---|---|---|---|---|---|---|---|---|
| ORF1a | ORF1a | ORF1ab | ORF1ab | ORF1ab | ORF1ab | ORF1ab | ORF1ab | ORF1ab | ORF1ab | ORF1ab | ORF1ab | ORF1ab | ORF1ab | ORF1ab | ORF1ab | ORF1ab | ORF1ab | ORF1ab | ORF1ab | ORF1ab | ORF1ab | ORF1ab | ORF1ab | ORF1ab | ORF1ab |
| 12757 | 13458 | 13585 | 13617 | 13730 | 13859 | 14130 | 14181 | 14274 | 14408 | 14425 | 14673 | 14805 | 15324 | 15435 | 15451 | 15708 | 16017 | 16078 | 16355 | 16393 | 16626 | 16738 | 16852 | 16887 | 16993 |
| T | C | C | G | C | A | A | G | G | C | C | A | C | C | A | G | G | C | G | A | C | C | G | G | C | T |
| . | . | . | . | . | . | . | . | . | . | . | . | . | . | . | . | . | . | . | . | . | . | . | . | . | . |
| . | . | . | . | . | . | . | . | . | . | . | . | . | . | . | . | . | . | . | . | . | . | . | . | . | . |
| . | . | . | . | . | . | . | . | . | T | . | . | . | . | . | . | . | . | . | . | . | . | . | . | . | . |
| . | . | . | . | . | . | . | . | . | T | . | . | . | . | . | . | . | . | . | . | . | . | . | . | . | . |
| . | . | . | . | . | . | . | . | . | T | . | . | . | . | . | . | . | . | . | . | . | . | . | . | . | . |
| . | . | . | . | . | . | . | . | . | T | . | . | . | . | . | . | . | . | . | . | . | . | . | . | . | . |
| . | . | . | . | . | . | . | . | . | T | . | . | . | . | . | . | . | . | . | . | . | . | . | . | . | . |
| . | . | . | . | . | . | . | . | . | T | . | . | . | . | . | . | . | . | . | . | . | . | . | . | . | . |
| . | . | . | . | . | . | . | . | . | T | . | . | . | . | . | . | . | . | . | . | . | . | . | . | . | . |
| . | . | . | . | . | . | . | . | . | T | . | . | . | . | . | . | . | . | . | . | . | . | . | . | . | . |
| . | . | . | . | . | . | . | . | . | T | . | . | . | . | . | . | . | . | . | . | . | . | . | . | . | . |
| . | . | . | . | . | . | . | . | . | T | . | . | . | . | . | . | . | . | . | . | . | . | . | . | . | . |
| . | . | . | . | . | . | . | . | . | T | . | . | . | . | . | . | . | . | . | . | . | . | . | . | . | . |
| . | . | . | . | . | . | . | . | . | T | . | . | . | . | . | . | . | . | . | . | . | . | . | . | . | . |
| . | . | . | . | . | . | . | . | . | T | . | . | . | . | . | . | . | . | . | . | . | . | . | . | . | . |
| . | . | . | . | . | . | . | . | . | T | . | . | . | . | . | . | . | . | . | . | . | . | . | . | . | . |
| . | . | . | . | . | . | . | . | . | . | . | . | . | . | . | . | . | . | . | . | . | . | . | . | . | . |
| . | . | . | . | . | . | . | . | . | . | . | . | . | . | . | . | . | . | . | . | . | . | . | . | . | . |
| . | . | . | . | . | . | . | . | . | . | . | . | . | . | . | . | . | . | . | . | . | . | . | . | . | . |
| . | . | . | . | . | . | . | . | . | . | . | . | . | . | . | . | . | . | . | . | . | . | . | . | . | . |
| . | . | . | . | . | . | . | . | . | . | . | . | . | . | . | . | . | . | . | . | . | . | . | . | . | . |
| . | . | . | . | . | . | . | . | . | . | . | . | . | . | . | . | . | . | . | . | . | . | . | . | . | . |
| . | . | . | . | . | . | . | . | . | . | . | . | . | . | . | . | . | . | . | . | . | . | . | . | . | . |
| . | . | . | . | . | . | . | . | . | . | . | . | . | . | . | . | . | . | . | . | . | . | . | . | . | . |
| . | . | . | . | . | . | . | . | . | . | . | . | . | . | . | . | . | . | . | . | . | . | . | . | . | . |
| . | . | . | . | . | . | . | . | . | . | . | . | . | . | . | . | . | . | . | . | . | . | . | . | . | . |
| . | . | . | . | . | . | . | . | . | . | . | . | . | . | . | . | . | . | . | . | . | . | . | . | . | . |
| . | . | . | . | . | . | . | . | . | . | . | . | . | . | . | . | . | . | . | . | . | . | . | . | . | . |
| . | . | . | . | . | . | . | . | . | T | . | . | . | . | . | . | . | . | . | . | . | . | . | . | . | . |
| . | . | . | . | . | . | . | . | . | . | . | . | . | . | . | . | . | . | . | . | . | . | . | . | . | . |
| . | . | . | . | . | . | . | . | . | T | . | . | . | . | . | . | . | . | . | . | . | . | . | . | . | . |
| . | . | . | . | . | . | . | . | . | T | . | . | . | . | . | . | . | . | . | . | . | . | . | . | . | . |
| . | . | . | . | . | . | . | . | . | T | . | . | . | . | . | . | . | . | . | . | . | . | . | . | . | . |
| . | . | . | . | . | . | . | . | . | T | . | . | . | . | . | . | . | . | . | . | . | . | . | . | . | . |
| . | . | . | . | . | . | . | . | . | T | . | . | . | . | . | . | . | . | . | . | . | . | . | . | . | . |
| . | . | . | . | . | . | . | . | . | T | . | . | . | ? | . | . | . | . | . | . | . | . | . | ? | ? | ? |
| . | . | . | . | . | . | . | . | . | T | . | . | . | ? | ? | ? | . | . | . | . | . | . | . | ? | ? | ? |
| . | . | . | . | . | . | . | . | . | T | . | . | . | ? | G | . | . | . | . | . | . | . | . | . | . | . |
| . | . | . | . | . | . | . | . | . | T | . | . | . | ? | G | . | . | . | . | . | . | . | . | . | . | . |
| . | . | . | . | T | . | . | . | . | . | . | . | . | ? | R | . | . | . | . | . | . | . | . | ? | ? | ? |
| . | . | . | . | T | . | . | . | . | . | . | . | . | ? | ? | ? | . | . | . | . | . | . | . | ? | ? | ? |
| . | . | . | . | . | . | . | . | . | T | . | . | . | ? | ? | ? | . | . | . | . | . | . | . | ? | ? | ? |
| . | . | . | . | T | . | . | . | . | . | . | . | . | ? | G | . | . | . | . | . | . | . | . | ? | ? | . |
| . | . | . | . | T | . | . | . | . | . | . | . | . | ? | ? | ? | . | . | . | . | . | . | . | ? | ? | ? |
| . | . | . | . | . | . | . | . | . | T | . | . | . | . | . | . | . | . | . | . | . | . | . | . | . | . |
| . | . | . | . | . | . | . | . | . | T | . | . | . | . | . | . | . | . | . | . | . | . | . | . | . | . |
| . | . | . | . | . | . | . | . | . | T | . | . | . | . | . | . | . | . | . | . | . | . | . | . | . | . |
| . | . | . | . | T | . | . | . | . | T | . | . | . | . | . | . | . | . | . | . | . | . | . | . | . | . |
| . | . | . | . | . | . | . | . | . | T | . | . | . | . | . | . | . | . | . | . | . | . | . | . | . | . |
| . | . | . | . | . | . | . | . | . | . | . | . | . | T | . | . | . | . | . | . | . | . | . | . | . | . |
| . | . | . | . | . | . | . | . | . | T | . | . | . | . | . | . | . | . | . | . | . | . | . | . | . | . |
| . | . | . | . | T | . | . | . | . | . | . | . | . | . | . | . | . | . | . | . | . | . | . | . | . | . |
| . | . | . | . | . | . | . | . | . | T | . | . | . | . | . | . | . | . | . | . | . | . | . | . | . | . |
| . | . | . | . | . | . | . | . | . | T | . | . | . | . | . | . | . | . | . | . | . | . | . | . | . | . |
| . | . | . | . | . | . | . | . | . | T | . | . | . | . | . | . | . | . | . | . | . | . | . | . | . | . |

| | | | | | | | | | | | | | | | | | | | | | | | | | |
|---|---|---|---|---|---|---|---|---|---|---|---|---|---|---|---|---|---|---|---|---|---|---|---|---|---|
| . | . | . | . | . | . | . | . | . | T | . | . | . | . | . | . | . | . | . | . | . | . | . | . | . | . |
| . | . | . | . | . | . | . | . | . | T | . | . | . | . | . | . | . | . | . | . | . | . | . | . | . | . |
| . | . | . | . | . | . | . | . | . | T | . | . | . | . | . | . | . | . | . | . | . | . | . | . | . | . |
| . | . | . | . | . | . | . | . | . | T | . | . | . | . | . | . | . | . | . | . | . | . | . | . | . | . |
| . | . | . | . | . | . | . | . | . | T | . | . | . | . | . | . | . | . | A | . | . | . | . | . | . | . |
| . | . | . | . | . | . | . | . | . | T | A | . | . | . | . | . | . | . | . | . | . | . | . | . | . | . |
| . | . | . | . | . | . | . | . | . | T | . | . | . | . | . | . | . | . | . | . | . | . | . | . | . | . |
| . | . | . | . | . | . | . | . | . | T | . | . | . | . | . | . | . | . | . | . | . | . | . | . | . | . |
| . | . | . | . | . | . | . | . | . | T | . | . | . | . | . | . | . | . | . | . | . | . | . | . | . | . |
| . | . | . | . | . | . | . | . | . | T | . | . | . | . | . | . | . | . | . | . | . | . | . | . | . | . |
| . | . | . | . | . | . | . | . | . | T | . | . | . | . | . | . | . | . | . | a | . | . | . | . | . | . |
| . | . | . | . | . | . | . | . | . | T | . | . | . | . | . | . | . | . | . | . | . | . | . | . | . | . |
| . | . | . | . | . | . | . | . | . | T | . | . | . | . | . | . | . | . | . | . | . | . | . | . | . | . |
| . | . | . | . | . | . | . | . | . | T | . | . | . | . | . | . | . | . | . | . | . | . | . | . | . | . |
| . | . | . | . | . | . | . | . | . | T | . | . | . | . | . | . | . | . | . | . | . | . | . | . | . | . |
| . | . | . | . | . | . | . | . | . | T | . | . | . | . | . | . | . | . | . | . | . | . | . | . | . | . |
| . | . | . | . | . | . | . | . | . | T | . | . | . | . | ? | . | . | . | . | . | . | . | . | . | . | . |
| . | . | . | . | . | . | . | . | . | T | . | . | . | . | . | . | . | . | . | . | . | . | . | . | . | . |
| . | . | . | . | . | . | . | . | . | T | . | . | . | . | . | . | . | . | . | . | . | . | . | . | . | . |
| . | . | . | . | T | . | . | . | . | . | . | . | . | . | . | . | . | . | . | . | . | . | . | . | . | . |
| . | . | . | . | T | . | . | . | . | . | . | . | . | . | . | . | . | . | . | . | c | . | . | . | . | . |
| . | . | . | . | T | . | . | . | . | . | . | . | . | . | ? | . | . | . | . | . | . | . | . | . | . | . |
| . | . | . | . | T | . | . | . | . | . | . | . | . | . | . | . | . | . | . | . | . | . | . | . | . | . |
| . | . | . | . | T | . | . | . | . | . | . | . | . | . | . | . | . | . | . | . | . | . | . | . | . | . |
| . | . | . | . | T | . | . | . | . | . | . | . | . | . | . | . | . | . | . | . | . | . | . | . | . | . |
| . | . | . | . | T | . | . | . | . | . | . | . | . | . | . | . | . | . | . | . | . | . | . | . | . | . |
| . | . | . | . | T | . | . | . | . | . | . | . | . | . | . | . | . | . | . | . | . | . | . | . | . | . |
| . | . | . | . | T | . | . | . | . | . | . | . | . | . | . | . | . | . | . | . | . | . | . | . | . | . |
| . | . | . | . | T | . | . | . | . | . | . | . | . | . | ? | . | . | . | . | . | . | . | . | . | . | . |
| . | . | . | . | T | . | . | . | . | . | . | . | . | . | . | . | . | . | . | . | . | . | . | . | . | . |
| . | . | . | . | T | . | . | . | . | . | . | . | . | . | . | . | . | . | . | . | . | . | . | . | c | t |
| . | . | . | . | T | . | . | . | . | . | . | . | . | . | . | . | . | . | . | . | . | . | . | . | . | . |
| . | . | . | . | T | . | . | . | . | . | . | . | . | . | . | . | . | . | . | . | . | c | . | . | . | . |
| . | . | . | . | T | . | . | . | . | . | . | . | . | . | . | . | . | . | . | . | . | . | . | . | . | . |
| . | . | . | . | T | . | . | . | . | . | . | . | . | . | . | . | . | . | . | . | . | . | . | . | . | . |
| . | . | . | . | T | . | . | . | . | . | . | . | . | . | . | . | . | . | . | . | . | . | . | . | . | . |
| . | . | . | . | T | . | . | . | . | . | . | . | . | . | . | . | . | . | . | . | . | . | . | . | . | . |
| . | . | . | . | T | . | . | . | . | . | . | . | . | . | . | . | . | . | . | . | . | . | . | . | . | . |
| . | . | . | . | T | . | . | . | . | . | . | . | . | . | . | . | . | . | . | . | . | . | . | . | . | . |
| . | . | . | . | T | . | . | . | . | . | . | . | . | . | . | . | . | . | . | . | . | . | . | . | . | . |
| . | . | . | . | T | . | . | . | . | . | . | . | . | . | . | . | . | . | . | . | . | . | . | . | . | . |
| . | . | . | . | . | . | . | . | . | . | . | . | . | . | . | . | . | . | . | . | . | . | . | . | . | C |
| . | . | . | . | . | . | . | . | . | . | . | . | . | . | . | . | . | . | . | . | . | . | . | . | . | C |
| . | . | . | . | . | . | . | . | . | . | . | . | . | . | . | . | . | . | . | . | . | . | . | . | . | C |
| . | . | . | . | . | . | . | . | . | . | . | . | . | . | . | . | . | . | . | . | . | . | . | . | . | C |
| . | . | . | . | . | . | . | . | . | . | . | . | . | . | . | . | . | . | . | . | . | . | . | . | . | C |
| . | . | . | . | . | . | . | . | . | . | . | . | . | . | . | . | . | . | . | . | . | . | . | . | . | . |
| . | . | . | . | . | . | . | . | . | . | . | . | T | . | . | . | . | . | . | . | . | . | . | . | . | . |
| . | . | . | . | . | . | . | . | . | . | . | . | T | . | . | . | . | . | . | . | . | . | . | . | . | . |
| . | . | . | . | . | . | . | . | . | . | . | . | . | . | . | . | . | . | . | . | . | . | . | . | . | . |
| . | . | . | . | . | . | a | g | . | . | . | . | T | . | . | . | . | . | . | . | . | . | . | . | . | . |
| . | . | . | . | T | . | . | . | . | . | . | . | . | . | . | . | . | . | . | . | . | . | . | . | . | . |
| . | . | . | . | . | . | . | . | . | T | . | . | . | . | . | . | . | . | . | . | . | . | . | . | . | . |
| . | . | . | . | T | . | . | . | . | . | . | . | . | . | . | . | . | . | . | . | . | . | . | . | . | . |
| . | . | . | . | T | . | . | . | . | . | . | . | . | . | . | . | . | . | . | . | . | . | . | . | . | . |
| . | . | . | . | T | . | . | . | . | . | . | . | . | . | . | . | . | . | . | . | . | . | . | . | . | . |
| . | . | . | . | T | . | . | . | . | . | . | . | . | . | . | . | . | . | . | . | . | . | . | . | . | . |
| . | . | . | . | T | . | . | . | . | . | . | . | . | . | R | . | . | . | . | . | . | . | . | . | . | . |
| . | . | . | . | T | . | . | . | . | . | . | . | . | . | ? | . | . | . | . | . | . | . | . | . | . | . |
| . | . | . | . | T | . | . | . | . | . | . | . | . | . | ? | . | . | . | . | . | . | . | . | . | . | . |
| . | . | . | . | T | a | . | . | . | . | . | . | . | . | ? | . | . | . | . | . | . | . | . | . | . | . |
| . | . | . | . | T | . | . | . | . | . | . | . | . | . | . | . | . | . | . | . | . | . | . | . | . | . |

| | | | | | | | | | | | | | | | | | | | | | | | | | |
|---|---|---|---|---|---|---|---|---|---|---|---|---|---|---|---|---|---|---|---|---|---|---|---|---|---|
| . | . | . | . | T | . | . | . | . | . | . | . | . | . | . | . | . | . | . | . | . | . | . | . | . | . |
| . | . | . | . | T | . | . | . | . | . | . | . | . | . | . | . | . | . | . | G | . | . | . | . | . | . |
| . | . | . | . | T | . | . | . | . | . | . | . | . | . | ? | . | . | . | . | . | . | . | . | . | . | . |
| . | . | . | . | T | . | . | . | . | . | . | . | . | . | R | . | . | . | . | . | . | . | . | . | . | . |
| . | . | . | . | T | . | . | . | . | . | . | . | . | . | . | . | . | . | . | . | . | . | . | . | . | . |
| . | c | . | . | T | . | . | . | . | . | . | . | . | . | . | . | . | . | . | . | . | c | . | . | . | . |
| . | . | . | . | T | . | . | . | . | . | . | . | . | . | R | . | . | . | . | . | . | . | . | . | . | . |
| . | . | . | . | T | . | . | . | . | . | . | . | . | . | . | . | . | . | . | . | . | . | . | . | . | . |
| . | . | . | . | Y | . | . | . | . | ? | c | . | . | . | . | . | . | . | . | . | . | . | . | . | . | . |
| . | . | . | . | T | . | . | . | . | . | . | . | . | . | . | . | . | . | . | . | . | . | . | . | . | . |
| . | . | . | . | T | . | . | . | . | . | . | . | . | . | R | . | . | . | . | . | . | . | . | . | . | . |
| . | . | . | . | T | . | . | . | . | . | . | . | . | . | R | . | . | . | . | . | . | . | . | . | . | . |
| . | . | . | . | T | . | . | . | . | . | . | . | . | . | . | . | . | . | . | . | . | . | . | . | . | . |
| . | . | . | . | T | . | . | . | . | . | . | . | . | . | R | . | . | . | . | . | . | . | . | . | . | . |
| . | . | . | . | T | . | . | . | . | . | . | . | . | . | . | . | . | . | . | . | . | . | . | . | . | . |
| . | . | . | . | T | . | . | . | . | . | . | . | . | . | ? | . | . | . | . | . | . | . | . | . | . | . |
| . | . | . | . | T | . | . | . | . | . | . | . | . | . | . | . | . | . | . | . | . | . | . | . | . | . |
| . | . | . | . | T | . | . | . | . | . | . | . | . | . | . | . | . | . | . | G | . | . | . | . | . | . |
| . | . | . | . | T | . | . | . | . | . | . | . | . | . | ? | . | . | . | . | . | . | . | . | . | . | . |
| . | . | . | . | T | . | . | . | . | . | . | . | . | . | . | . | . | . | . | . | . | . | . | . | . | . |
| . | . | . | . | T | . | . | . | . | . | . | . | . | . | . | . | . | . | . | . | . | . | . | . | . | . |
| . | . | . | . | . | . | . | . | . | T | . | . | . | . | R | . | . | . | . | . | . | . | . | . | . | . |
| . | . | . | . | . | . | . | . | . | T | . | . | . | . | R | . | . | . | . | . | . | . | . | . | . | . |
| . | . | . | . | . | . | . | . | . | T | . | . | . | . | . | . | . | . | . | . | . | . | . | . | . | . |
| . | . | . | . | . | . | . | . | . | T | . | . | . | . | . | . | . | . | . | . | . | . | . | . | . | . |
| . | . | . | . | . | . | . | . | . | T | . | . | . | . | . | . | . | . | . | . | . | c | . | . | . | . |
| . | . | . | . | . | . | . | . | . | Y | . | . | . | . | R | . | . | . | . | . | . | . | . | . | . | . |
| . | . | . | . | . | . | . | . | . | T | . | . | . | . | R | . | . | . | . | . | . | . | . | . | . | . |
| . | . | . | . | . | . | . | . | . | T | . | . | . | . | ? | . | . | . | . | . | . | . | . | . | . | . |
| . | . | . | . | . | . | . | . | . | T | . | . | . | . | . | . | . | . | . | . | . | . | . | . | . | . |
| . | . | . | . | . | . | . | . | . | T | . | . | . | . | R | . | . | . | . | . | . | . | . | . | . | . |
| . | . | . | . | Y | . | . | . | . | Y | . | . | . | . | ? | . | . | . | . | . | . | . | . | . | . | . |
| . | . | . | . | T | . | . | . | . | . | . | . | . | . | . | . | . | . | . | . | . | . | . | . | . | . |
| . | . | . | . | . | . | . | . | . | T | . | . | . | . | . | . | . | . | . | . | . | . | . | . | . | . |
| . | . | . | . | . | . | . | . | . | . | . | . | . | . | . | . | . | . | . | . | . | . | . | . | . | . |
| . | . | . | . | T | . | . | . | . | . | . | . | . | . | . | . | . | T | . | . | . | . | . | . | . | . |
| . | . | . | . | T | . | . | . | . | . | . | . | . | . | . | . | . | T | . | . | . | . | . | . | . | . |
| . | . | . | . | . | . | . | . | C | T | . | . | . | . | . | . | . | . | . | . | . | . | . | . | . | . |
| . | . | . | . | . | . | . | . | C | T | . | . | . | . | . | . | . | . | . | . | . | . | . | . | . | . |
| . | . | . | . | . | . | . | . | . | T | . | G | . | . | . | . | . | . | . | . | . | . | . | . | . | . |
| . | . | . | . | . | . | . | . | . | T | . | G | . | . | . | . | . | . | . | . | . | . | . | . | . | . |
| . | . | . | . | . | . | . | . | . | T | . | . | . | . | . | . | . | . | . | . | . | . | . | . | . | . |
| . | . | . | . | . | . | . | . | . | T | . | . | . | . | . | . | . | . | . | . | . | . | . | . | . | . |
| . | . | . | . | . | . | . | . | . | T | . | . | . | . | . | . | . | . | . | . | . | . | . | . | . | . |
| . | . | . | . | . | . | . | . | . | T | . | . | . | . | . | . | . | . | . | . | . | . | . | . | . | . |
| . | . | . | . | T | . | . | . | . | . | . | . | . | . | . | . | . | . | . | . | . | . | . | . | . | . |
| . | . | . | . | T | G | . | . | . | T | . | . | . | . | . | . | . | . | A | . | . | . | . | . | . | . |
| . | . | . | . | T | . | . | . | . | T | . | . | . | . | . | . | . | . | . | . | . | . | . | . | . | . |
| . | . | . | . | . | . | . | . | . | T | . | . | . | . | . | . | . | . | . | . | . | . | . | . | . | . |
| . | . | . | . | T | . | . | . | . | . | . | . | . | . | . | . | . | . | . | . | . | . | . | . | . | . |
| . | . | . | . | T | . | . | . | . | . | . | . | . | . | . | . | . | . | . | . | . | . | . | . | . | . |
| . | . | . | . | . | . | . | . | . | T | . | . | . | . | . | . | . | . | . | . | . | . | . | . | . | . |
| . | . | . | . | . | . | . | . | . | T | . | . | . | . | . | . | . | . | . | . | . | . | . | . | . | . |
| . | . | . | . | . | . | . | . | . | T | A | . | . | . | . | . | . | . | . | . | . | . | . | . | . | . |
| . | . | . | . | . | . | . | . | . | T | . | . | . | . | . | . | . | . | A | . | . | . | . | . | . | . |
| . | . | . | . | . | . | . | . | . | T | . | G | . | . | . | . | . | . | . | . | . | . | . | . | . | . |
| . | . | . | . | . | . | . | . | . | T | . | . | . | . | . | . | . | . | . | . | . | . | . | . | . | . |
| . | . | . | . | . | . | . | . | . | T | A | . | . | . | . | . | . | . | . | . | . | . | . | . | . | . |
| . | . | . | . | . | . | . | . | . | T | . | . | . | . | . | . | . | . | . | . | . | . | . | . | . | . |
| . | . | . | . | . | . | . | . | . | T | . | . | . | . | . | . | . | . | . | . | . | . | . | . | . | . |
| . | . | . | . | . | . | . | . | . | T | . | . | . | . | . | . | . | . | . | . | . | . | . | . | . | . |
| . | . | . | . | . | . | . | . | . | T | . | . | . | . | . | . | . | . | . | . | . | . | . | . | . | . |

| | | | | | | | | | | | | | | | | | | | | | | | | | |
|---|---|---|---|---|---|---|---|---|---|---|---|---|---|---|---|---|---|---|---|---|---|---|---|---|---|
| . | . | . | . | . | . | . | . | . | T | . | . | . | . | . | . | . | . | . | . | . | . | . | . | . | . |
| . | . | . | . | . | . | . | . | . | T | . | . | . | . | . | . | . | . | . | . | . | . | . | . | . | . |
| . | . | . | . | . | . | . | . | . | T | . | . | . | . | . | . | . | . | A | . | . | . | . | . | . | . |
| . | . | . | . | . | . | . | . | . | T | . | . | . | . | . | . | . | . | . | . | . | . | . | . | . | . |
| . | . | . | . | . | . | . | . | . | T | . | . | . | . | . | . | . | . | . | . | . | . | . | . | . | . |
| . | . | . | . | . | . | . | . | . | T | . | . | . | . | . | . | . | . | . | . | . | . | . | . | . | . |
| . | . | . | . | . | . | . | . | . | T | . | . | . | . | . | . | . | . | . | . | . | . | . | . | . | . |
| . | . | . | . | . | . | . | . | . | T | . | . | . | . | . | . | . | . | . | . | . | . | . | . | . | . |
| . | . | . | . | . | . | . | . | . | T | A | . | . | . | . | . | . | . | . | . | . | . | . | . | . | . |
| . | . | . | . | . | . | . | . | . | T | . | . | . | . | . | . | . | . | . | . | . | . | . | . | . | . |
| . | . | . | . | . | . | . | . | . | T | . | . | . | . | . | . | . | . | . | . | . | . | . | . | . | . |
| . | . | . | . | . | . | . | . | . | T | A | . | . | . | . | . | . | . | . | . | . | . | . | . | . | . |
| . | . | . | . | . | . | . | . | . | T | . | . | . | . | . | . | . | . | . | . | . | . | . | . | . | . |
| . | . | . | . | . | . | . | . | . | T | . | . | . | . | . | . | . | . | . | . | . | . | . | . | . | . |
| . | . | . | . | . | . | . | . | . | T | . | . | . | . | . | . | . | . | . | . | . | . | . | . | . | . |
| . | . | . | . | . | . | . | . | . | T | . | . | . | . | . | . | . | . | . | . | . | . | . | . | . | . |
| . | . | . | . | . | . | . | . | . | T | . | . | . | . | . | . | . | . | . | . | . | . | . | . | . | . |
| . | . | . | . | . | . | . | . | . | T | . | . | . | . | . | . | . | . | . | . | . | . | . | . | . | . |
| . | . | . | . | . | . | . | . | . | T | . | . | . | . | . | . | . | . | . | . | . | . | . | . | . | . |
| . | . | . | . | . | . | . | . | . | T | . | . | . | . | . | . | . | . | . | . | . | . | . | . | . | . |
| . | . | . | . | . | . | . | . | . | T | . | . | . | . | . | . | . | . | . | . | . | . | . | . | . | . |
| . | . | . | . | . | . | . | . | . | T | . | . | . | . | . | . | . | . | . | . | . | . | . | . | . | . |
| . | . | . | . | . | . | . | . | . | T | . | . | . | . | . | . | . | . | . | . | . | . | . | . | . | . |
| . | . | . | . | . | . | . | . | . | T | . | . | . | . | . | . | . | . | . | . | . | . | . | . | . | . |
| . | . | . | . | . | . | . | . | . | T | . | . | . | . | . | . | . | . | A | . | . | . | . | . | . | . |
| . | . | . | . | . | . | . | . | . | T | . | . | . | . | . | . | . | . | . | . | . | . | . | . | . | . |
| . | . | . | . | . | . | . | . | . | T | . | . | . | . | . | . | . | . | . | . | . | . | . | . | . | . |
| . | . | . | . | . | . | . | . | . | T | . | . | . | . | . | . | . | . | . | . | . | . | . | . | . | . |
| . | . | . | . | . | . | . | . | . | T | . | . | . | . | . | . | . | . | . | . | . | . | . | . | . | . |
| . | . | . | . | . | . | . | . | . | T | . | . | . | . | . | . | . | . | . | . | . | . | . | . | . | . |
| . | . | . | . | . | . | . | . | . | T | . | . | . | . | . | . | . | . | . | . | . | . | . | . | T | . |
| . | . | . | . | . | . | . | . | . | T | . | . | . | . | . | . | . | . | . | . | . | . | . | . | T | . |
| . | . | . | . | . | . | . | . | . | T | . | . | . | . | . | . | . | . | . | . | . | . | . | . | . | . |
| . | . | . | . | . | . | . | . | . | T | . | . | . | . | . | . | . | . | A | . | . | . | . | . | . | . |
| . | . | . | . | . | . | . | . | . | T | . | . | . | . | . | . | . | . | A | . | . | . | . | . | . | . |
| . | . | . | . | . | . | . | . | . | T | . | . | . | . | . | . | . | . | . | . | . | . | . | . | . | . |
| . | . | . | . | . | . | . | . | . | T | . | . | . | . | . | . | . | . | . | . | . | . | . | . | . | . |
| . | . | . | . | . | . | . | . | . | T | . | . | . | . | . | . | . | . | . | . | . | . | . | . | . | . |
| . | . | . | . | . | . | . | . | . | T | . | . | . | . | . | . | . | . | . | . | . | . | . | . | . | . |
| . | . | . | . | . | . | . | . | . | T | . | . | . | . | . | . | . | . | A | . | . | . | . | . | . | . |
| . | . | . | . | . | . | . | . | . | T | . | . | . | . | . | . | . | . | A | . | . | . | . | . | . | . |
| . | . | . | . | . | . | . | . | . | T | . | . | . | . | . | . | . | . | A | . | . | . | . | . | . | . |
| . | . | . | . | . | . | . | . | . | T | . | . | . | . | . | . | . | . | . | . | . | . | . | . | . | . |
| . | . | . | . | . | . | . | . | . | T | . | . | . | . | . | . | . | . | . | . | . | . | . | T | . | . |
| . | . | . | . | . | . | . | . | . | T | . | . | . | . | . | . | . | . | . | . | . | . | . | . | . | . |
| . | . | . | . | . | . | . | . | . | T | . | . | . | . | . | . | . | . | . | . | . | . | . | . | . | . |
| . | . | . | . | . | . | . | . | . | . | . | . | . | . | . | . | . | . | . | . | . | . | . | . | . | . |
| . | . | . | . | . | . | . | . | . | T | . | . | . | . | . | . | . | . | . | . | . | . | . | . | . | . |
| . | . | . | . | . | . | . | . | . | . | . | . | . | . | . | . | . | . | . | . | . | . | . | . | . | . |
| . | . | . | . | . | . | . | . | . | T | . | . | . | . | . | . | . | . | . | . | . | . | . | . | . | . |
| . | . | . | . | . | . | . | . | . | T | . | . | . | . | . | . | . | . | . | . | . | . | . | . | . | . |
| . | . | . | . | . | . | . | . | . | T | . | . | . | . | . | . | . | . | . | . | . | . | . | . | . | . |
| . | . | . | . | . | . | . | . | . | T | . | . | . | . | . | . | . | . | . | . | . | . | . | . | . | . |
| . | . | . | . | . | . | . | . | . | T | . | . | . | . | . | . | . | . | . | . | . | . | . | . | . | . |
| . | . | . | . | . | . | . | . | . | T | . | . | . | . | . | . | . | . | . | . | . | . | . | . | . | . |
| . | . | . | . | . | . | . | . | . | T | . | . | . | . | . | . | . | . | . | . | . | . | . | . | . | . |
| . | . | . | . | . | . | . | . | . | T | . | . | . | . | . | . | . | . | A | . | . | . | . | . | . | . |
| . | . | . | . | . | . | . | . | . | T | . | . | . | . | . | . | . | . | . | . | . | . | . | . | . | . |
| . | . | . | . | . | . | . | . | . | T | . | . | . | . | . | . | . | . | A | . | . | . | . | . | . | . |
| . | . | . | . | . | . | . | . | . | T | . | G | . | . | . | . | . | . | . | . | . | . | . | . | . | . |
| . | . | . | . | . | . | . | . | . | T | . | G | . | . | . | . | . | . | . | . | . | . | . | . | . | . |
| . | . | . | . | . | . | . | . | . | T | . | G | . | . | . | . | . | . | . | . | . | . | . | . | . | . |

| | | | | | | | | | | | | | | | | | | | | | | | | | |
|---|---|---|---|---|---|---|---|---|---|---|---|---|---|---|---|---|---|---|---|---|---|---|---|---|---|
| . | . | . | . | . | . | . | . | . | T | . | . | . | . | . | . | . | . | . | . | . | . | . | T | . | . |
| . | . | . | . | . | . | . | . | . | T | . | . | . | . | . | . | . | . | . | . | . | . | . | . | . | . |
| . | . | . | . | . | . | . | . | . | T | . | . | . | . | . | . | . | . | . | . | . | . | . | . | . | . |
| . | . | . | . | . | . | . | . | . | T | . | . | . | . | . | . | . | . | . | . | . | . | . | . | . | . |
| . | . | . | . | T | . | . | . | . | . | . | . | . | . | . | . | . | . | . | . | . | . | . | . | . | . |
| . | . | . | . | T | . | . | . | . | . | . | . | . | . | . | . | . | . | . | . | . | . | . | . | . | . |
| . | . | . | . | T | . | . | . | . | . | . | . | . | . | . | . | . | . | . | . | . | . | . | . | . | . |
| . | . | . | . | T | . | . | . | . | . | . | . | . | . | . | . | . | . | . | . | . | . | . | . | . | . |
| . | . | . | . | T | . | . | . | . | . | . | . | . | . | . | . | . | . | . | . | . | . | . | . | . | . |
| . | . | . | . | T | . | . | . | . | . | . | . | . | . | . | . | . | . | . | . | . | . | . | . | . | . |
| . | . | . | . | T | . | . | . | . | . | . | . | . | . | . | . | . | . | . | . | . | . | . | . | . | . |
| . | . | . | . | T | . | . | . | . | . | . | . | . | . | . | . | . | . | . | . | . | . | . | . | . | . |
| . | . | . | . | T | . | . | . | . | . | . | . | . | . | . | . | . | . | . | . | . | . | . | . | . | . |
| . | . | . | . | T | . | . | . | . | . | . | . | . | . | . | . | . | . | . | . | . | . | . | . | . | . |
| . | . | . | . | T | . | . | . | . | . | . | . | . | . | . | . | . | . | . | . | . | . | . | . | . | . |
| . | . | . | . | T | . | . | . | . | . | . | . | . | . | . | . | . | . | . | . | . | . | . | . | . | . |
| . | . | . | . | T | . | . | . | . | . | . | . | . | . | . | . | . | . | . | . | . | . | . | . | . | . |
| . | . | . | . | T | . | . | . | . | . | . | . | . | . | . | . | . | . | . | . | . | . | . | . | . | . |
| . | . | . | . | T | . | . | . | . | . | . | . | . | . | . | . | . | . | . | . | . | . | . | . | . | . |
| . | . | . | . | T | . | . | . | . | . | . | . | . | . | . | . | . | . | . | . | . | . | . | . | . | . |
| . | . | . | . | T | . | . | . | . | . | . | . | . | . | . | . | . | . | . | . | . | . | . | . | . | . |
| . | . | . | . | T | . | . | . | . | . | . | . | . | . | . | . | . | . | . | . | . | . | . | . | . | . |
| . | . | . | . | T | . | . | . | . | . | . | . | . | . | . | . | . | . | . | . | . | . | . | . | . | . |
| . | . | . | . | T | . | . | . | . | . | . | . | . | . | . | . | . | . | . | . | . | . | . | . | . | . |
| . | . | . | . | T | . | . | . | . | . | . | . | . | . | . | . | . | . | . | . | . | . | . | . | . | . |
| . | . | . | . | T | . | . | . | . | . | . | . | . | . | . | . | . | . | . | . | . | . | . | . | . | . |
| . | . | . | . | T | . | . | . | . | . | . | . | . | . | . | . | . | . | . | . | . | . | . | . | . | . |
| . | . | . | . | T | . | . | . | . | . | . | . | . | . | . | . | . | . | . | . | . | . | . | . | . | . |
| . | . | . | . | T | . | . | . | . | . | . | . | . | . | . | . | . | . | . | . | . | . | . | . | . | . |
| . | . | . | . | T | . | . | . | . | . | . | . | . | . | . | . | . | . | . | . | . | . | . | . | . | . |
| . | T | . | . | T | . | . | . | . | . | . | . | . | . | . | . | . | . | . | . | . | . | . | . | . | . |
| . | . | . | . | T | . | . | . | . | . | . | . | . | . | . | . | . | . | . | . | . | . | . | . | . | . |
| . | . | . | . | T | . | . | . | . | . | . | . | . | . | . | . | . | . | . | . | . | . | . | . | . | . |
| . | . | . | . | T | . | . | . | . | . | . | . | . | . | . | . | . | . | . | . | . | . | . | . | . | . |
| . | . | . | . | . | . | . | . | . | T | . | . | . | . | . | . | . | . | . | . | . | . | . | . | . | . |
| . | . | . | . | T | . | . | . | . | . | . | . | . | . | . | . | . | . | . | . | . | . | . | . | . | . |
| . | . | . | . | T | . | . | . | . | . | . | . | . | . | . | . | . | . | . | . | . | . | . | . | . | . |
| . | . | . | . | T | . | . | . | . | . | . | . | . | . | . | . | . | . | . | . | . | . | . | . | . | . |
| . | . | . | . | T | . | . | . | . | . | . | . | . | . | . | . | . | . | . | . | . | . | . | . | . | . |
| . | . | . | . | T | . | . | . | . | . | . | . | . | . | . | . | . | . | . | . | . | . | . | . | . | . |
| . | . | . | . | T | . | . | . | . | . | . | . | . | . | . | . | . | . | . | . | . | . | . | . | . | . |
| . | . | . | . | T | . | . | . | . | . | . | . | . | . | . | . | . | . | . | . | . | . | . | . | . | . |
| . | . | . | . | T | . | . | . | . | . | . | . | . | . | . | . | . | . | . | . | . | . | . | . | . | . |
| . | . | . | . | T | . | . | . | . | . | . | . | . | . | . | . | . | . | . | . | . | . | . | . | . | . |
| . | . | . | . | T | . | . | . | . | . | . | . | . | . | . | . | . | . | . | . | . | . | . | . | . | . |
| . | . | . | . | T | . | . | . | . | . | . | . | . | . | . | . | . | . | . | . | . | . | . | . | . | . |
| . | . | . | . | T | . | . | . | . | . | . | . | . | . | . | . | . | . | . | . | . | . | . | . | . | . |
| . | . | . | . | T | . | . | . | . | . | . | . | . | . | . | . | . | . | . | . | . | . | . | . | . | . |
| . | . | . | . | T | . | . | . | . | . | . | . | . | . | . | . | . | . | . | . | . | . | . | . | . | . |
| . | . | . | . | T | . | . | . | . | . | . | . | . | . | . | . | . | . | . | . | . | . | . | . | . | . |
| . | . | . | . | T | . | . | . | . | . | . | . | . | . | . | . | . | . | . | . | . | . | . | . | . | . |
| . | . | . | . | T | . | . | . | . | . | . | . | . | . | . | . | . | . | . | . | . | . | . | . | . | . |
| . | T | . | . | T | . | . | . | . | . | . | . | . | . | . | . | . | . | . | . | . | . | . | . | . | . |
| . | . | . | . | T | . | . | . | . | . | . | . | . | . | . | . | . | . | . | . | . | . | . | . | . | . |
| . | . | . | . | T | . | . | . | . | . | . | . | . | . | . | . | . | . | . | . | . | . | . | . | . | . |
| . | . | . | . | T | . | . | . | . | . | . | . | . | . | . | . | . | . | . | . | . | . | . | . | . | . |
| . | . | . | . | T | . | . | . | . | . | . | . | . | . | . | . | . | . | . | . | . | . | . | . | . | . |
| . | . | . | . | . | . | . | . | . | T | . | . | . | . | . | . | . | . | . | . | . | . | . | . | . | . |
| . | . | . | . | . | . | . | . | . | . | . | . | . | . | . | . | . | . | . | . | . | . | . | . | . | . |
| . | . | . | . | . | . | . | . | . | T | . | . | . | . | . | . | . | . | . | . | . | . | . | . | . | . |
| . | . | . | . | T | . | . | . | . | . | . | . | . | . | . | . | . | . | . | . | . | . | . | . | . | . |

| | | | | | | | | | | | | | | | | | | | | | | | | | |
|---|---|---|---|---|---|---|---|---|---|---|---|---|---|---|---|---|---|---|---|---|---|---|---|---|---|
| . | . | . | . | T | . | . | . | . | . | . | . | . | . | . | . | . | . | . | . | . | . | . | . | . | . |
| . | . | . | . | T | . | . | . | . | . | . | . | . | . | . | . | . | . | . | . | . | . | . | . | . | . |
| . | . | . | . | T | . | . | . | . | . | . | . | . | . | . | . | . | . | . | . | . | . | . | . | . | . |
| . | . | . | . | T | . | . | . | . | . | . | . | . | . | . | . | . | . | . | . | . | . | . | . | . | . |
| . | . | . | . | T | . | . | . | . | . | . | . | . | . | . | . | . | . | . | . | . | . | . | . | . | . |
| . | . | . | . | T | . | . | . | . | . | . | . | . | . | . | . | . | . | . | . | . | . | . | . | . | . |
| . | . | . | . | . | . | . | . | . | T | . | . | . | T | . | . | . | . | . | . | . | . | . | . | . | . |
| . | . | . | . | . | . | . | . | . | T | . | . | . | . | . | . | . | . | . | . | . | . | . | . | . | . |
| . | . | . | . | . | . | . | . | . | T | . | . | . | . | . | . | . | . | A | . | . | . | . | . | . | . |
| . | . | . | . | ? | . | . | . | . | . | . | . | . | . | . | . | . | . | . | . | . | . | . | . | . | . |
| . | . | . | T | . | . | . | . | . | T | . | . | . | . | . | . | . | . | . | . | . | . | . | . | . | . |
| . | . | . | T | . | . | . | . | . | T | . | . | . | . | . | . | . | . | . | . | . | . | . | . | . | . |
| . | . | . | . | T | . | . | . | . | . | . | . | . | . | . | . | . | . | . | . | . | . | . | . | . | . |
| . | . | . | . | . | . | . | . | . | T | A | . | . | . | . | . | . | . | . | . | . | . | . | . | . | . |
| . | . | . | . | . | . | . | . | . | T | . | . | . | . | . | . | . | . | . | . | . | . | . | . | . | . |
| . | . | . | . | . | . | . | . | . | . | . | . | . | . | . | . | . | . | . | . | . | . | . | . | . | . |
| . | . | . | . | . | . | . | . | . | . | . | . | . | . | . | . | . | . | . | . | . | . | . | . | . | . |
| . | . | . | . | . | . | . | . | . | T | . | . | . | . | . | . | . | . | . | . | . | . | . | . | . | . |
| . | . | . | . | . | . | . | . | . | T | . | . | . | T | . | . | . | . | . | . | . | . | . | . | . | . |
| . | . | . | . | . | . | . | . | . | T | . | . | . | . | . | . | . | . | . | . | . | . | . | . | . | . |
| . | . | . | . | . | . | . | . | . | T | . | . | . | . | . | . | . | . | . | . | . | . | . | . | . | . |
| . | . | . | . | . | . | . | . | . | T | . | . | . | . | . | . | . | . | . | . | . | . | . | . | . | . |
| C | . | . | . | T | . | . | . | . | . | . | . | . | . | . | . | . | . | . | . | . | . | . | . | . | . |
| . | . | . | . | . | . | . | . | . | T | . | . | . | . | . | . | . | . | . | . | . | . | . | . | . | . |
| C | . | . | . | T | . | . | . | . | . | . | . | . | . | . | . | . | . | . | . | . | . | . | . | . | . |
| . | . | . | . | . | . | . | . | . | T | . | . | . | . | . | . | . | . | . | . | . | . | . | . | . | . |
| . | . | . | . | . | . | . | . | . | T | . | . | . | . | . | . | . | . | . | . | . | . | . | . | . | . |
| C | . | . | . | T | . | . | . | . | . | . | . | . | . | . | . | . | . | . | . | . | . | . | . | . | . |
| . | . | . | . | . | . | . | . | . | T | . | . | . | . | . | . | . | . | . | . | . | . | . | . | . | . |
| C | . | . | . | T | . | . | . | . | . | . | . | . | . | . | . | . | . | . | . | . | . | . | . | . | . |
| . | . | . | . | . | . | . | . | . | T | . | . | . | . | . | . | . | . | . | . | . | . | . | . | . | . |
| . | . | . | . | . | . | . | . | . | T | . | . | . | . | . | . | . | . | . | . | . | . | . | . | . | . |
| . | . | . | . | T | . | . | . | . | . | . | . | . | . | . | . | . | . | . | . | . | . | . | . | . | . |
| t | c | c | g | c | a | a | g | g | t | c | a | c | c | a | g | g | c | g | a | c | c | g | g | c | t |
| t | c | c | g | c | a | a | g | g | t | c | a | c | c | a | g | g | c | g | a | c | c | g | g | c | t |
| t | c | c | g | c | a | a | g | g | t | c | a | c | c | a | g | g | c | g | a | c | c | g | g | c | t |
| t | c | c | g | c | a | a | g | g | t | c | a | c | c | a | g | g | c | g | a | c | c | g | g | c | t |
| t | c | c | g | c | a | a | g | g | t | c | a | c | c | a | g | g | c | g | a | c | c | g | g | c | t |
| t | c | c | g | c | a | a | g | g | t | c | a | c | c | a | g | g | c | g | a | c | c | g | g | c | t |
| t | c | c | g | c | a | a | g | g | t | c | a | c | c | a | g | g | c | g | a | c | c | g | g | c | t |
| t | c | c | g | c | a | a | g | g | t | c | a | c | c | a | g | g | c | g | a | c | c | g | g | c | t |
| t | c | c | g | c | a | a | g | g | t | c | a | c | c | a | g | t | c | g | a | c | c | g | g | c | t |
| t | c | c | g | c | a | a | g | g | c | c | a | c | c | a | g | g | c | g | a | c | c | g | g | c | t |
| t | c | c | g | c | a | a | g | g | c | c | a | c | c | a | g | g | c | g | a | c | c | g | g | c | t |
| t | c | c | g | c | a | a | g | g | c | c | a | c | c | a | g | g | c | g | a | c | c | g | g | c | t |
| t | c | c | g | c | a | a | g | g | c | c | a | c | c | a | g | g | c | g | a | c | c | g | g | c | t |
| t | c | c | g | c | a | a | g | g | t | c | a | c | c | a | g | g | c | g | a | c | c | g | g | c | t |
| t | c | c | g | t | a | a | g | g | c | c | a | c | c | a | g | g | c | g | a | c | c | g | g | c | t |
| t | c | c | g | t | a | a | g | g | c | c | a | c | c | a | g | g | c | g | a | c | c | g | g | c | t |
| t | c | c | g | c | a | a | g | g | t | c | a | c | c | a | g | g | c | g | a | c | c | g | g | c | t |
| t | c | c | g | c | a | a | g | g | t | c | a | c | c | a | g | g | c | g | a | c | c | g | g | c | t |
| t | c | c | g | c | a | a | g | g | t | c | a | c | c | a | g | g | c | g | a | c | c | g | g | c | t |
| t | c | c | g | c | a | a | g | g | c | c | a | c | c | a | g | g | c | g | a | c | c | g | g | c | t |
| t | c | c | g | c | a | a | g | g | t | c | a | c | t | a | g | g | c | g | a | c | c | g | g | c | t |
| t | c | c | g | t | a | a | g | g | c | c | a | c | c | a | g | g | c | g | a | c | c | g | g | c | t |
| t | c | c | g | c | a | a | g | g | t | c | a | c | c | a | g | g | c | g | a | c | c | g | g | c | t |
| t | c | c | g | c | a | a | g | g | t | c | a | c | t | a | g | g | c | g | a | c | c | g | g | c | t |
| . | . | . | . | . | . | . | . | . | T | . | . | . | . | . | . | . | . | . | . | . | . | . | . | . | . |
| . | . | . | . | . | . | . | . | . | T | . | . | . | . | . | . | . | . | . | . | . | . | . | . | . | . |
| . | . | . | . | . | . | . | . | . | T | . | . | . | . | . | . | . | . | . | . | . | . | . | . | . | . |
| C | . | . | . | T | . | . | . | . | . | . | . | . | . | . | . | . | . | . | . | . | . | . | . | . | . |
| . | . | . | . | . | . | . | . | . | T | . | . | . | . | . | . | . | . | . | . | . | . | . | . | . | . |

| | | | | | | | | | | | | | | | | | | | | | | | | | |
|---|---|---|---|---|---|---|---|---|---|---|---|---|---|---|---|---|---|---|---|---|---|---|---|---|---|
| C | . | . | . | T | . | . | . | . | . | . | . | . | . | . | . | . | . | . | . | . | . | . | . | . | . |
| . | . | . | . | . | . | . | . | . | T | . | . | . | . | . | . | T | . | . | . | . | . | . | . | . | . |
| . | . | . | . | . | . | . | . | . | . | . | . | . | . | . | . | . | . | . | . | . | . | . | . | . | . |
| . | . | . | . | . | . | . | . | . | . | . | . | . | . | . | . | . | . | . | . | . | . | . | . | . | . |
| . | . | . | . | . | . | . | . | . | T | . | . | . | . | . | . | . | . | . | . | . | . | . | . | . | . |
| . | . | . | . | . | . | . | . | . | . | . | . | . | . | . | . | . | . | . | . | . | . | . | . | . | . |
| . | . | . | . | . | . | . | . | . | T | . | . | . | T | . | . | . | . | . | . | . | . | . | . | . | . |
| . | . | . | . | . | . | . | . | . | . | . | . | . | . | . | . | . | . | . | . | . | . | . | . | . | . |
| . | . | . | . | . | . | . | . | . | T | . | . | . | . | . | . | . | . | . | . | . | . | . | . | . | . |
| . | . | . | . | . | . | . | . | . | T | . | . | . | . | . | . | . | . | . | . | . | . | . | . | . | . |
| . | . | . | . | . | . | . | . | . | . | . | . | . | . | . | . | . | . | . | . | . | . | . | . | . | . |
| . | . | . | . | . | . | G | . | . | . | . | . | . | . | . | . | . | . | . | . | . | . | . | . | . | . |
| . | . | . | . | . | . | G | . | . | . | . | . | . | . | . | . | . | . | . | . | . | . | . | . | . | . |
| . | . | . | . | T | . | . | A | . | . | . | . | . | . | . | . | . | . | . | . | . | . | . | . | . | . |
| . | . | . | . | T | . | . | . | . | . | . | . | . | . | . | . | . | . | . | . | . | . | . | . | . | . |
| . | . | . | . | T | . | . | . | . | . | . | . | . | . | . | . | . | . | . | . | . | . | . | . | . | . |
| . | . | . | . | T | . | . | A | . | . | . | . | . | . | . | . | . | . | . | . | . | . | . | . | . | . |
| . | . | . | . | T | . | . | . | . | . | . | . | . | . | . | . | . | . | . | . | . | . | . | . | . | . |
| . | . | . | . | T | . | . | . | . | . | . | . | . | . | . | . | . | . | . | . | . | . | . | . | . | . |
| . | . | . | . | . | . | . | . | . | . | . | . | T | . | . | . | . | . | . | . | . | . | . | . | . | . |
| . | . | . | . | T | . | . | A | . | . | . | . | . | . | . | . | . | . | . | . | . | . | . | . | . | . |
| . | . | . | . | T | . | . | . | . | . | . | . | . | . | . | . | . | . | . | . | . | . | . | . | . | . |
| . | . | . | . | . | . | . | . | . | T | . | . | . | . | . | . | . | . | . | . | . | . | . | . | . | . |
| . | . | . | . | . | . | . | . | . | T | . | . | . | T | . | . | . | . | . | . | . | . | . | . | . | . |
| . | . | . | . | . | . | . | . | . | T | . | . | . | . | . | . | . | . | . | . | . | . | . | . | . | . |
| . | . | . | . | . | . | . | . | . | T | . | . | . | . | . | . | . | . | . | . | . | . | . | . | . | . |
| . | . | . | . | . | . | . | . | . | T | . | . | . | . | . | . | . | . | . | . | . | . | . | . | . | . |
| . | . | . | . | . | . | . | . | . | T | . | . | . | T | . | . | . | . | . | . | . | . | . | . | . | . |
| . | . | . | . | . | . | . | . | . | T | . | . | . | . | . | . | . | . | . | . | . | . | . | . | . | . |
| . | . | . | . | T | . | . | . | . | . | . | . | . | . | . | . | . | . | . | . | . | . | . | . | . | . |
| . | . | . | . | . | . | . | . | . | T | . | . | . | T | . | . | . | . | . | . | . | . | . | . | . | . |
| . | . | . | . | . | . | . | . | . | T | . | . | . | T | . | . | . | . | . | . | . | . | . | . | . | . |
| . | . | . | . | T | . | . | . | . | . | . | . | . | . | . | . | . | . | . | . | . | . | . | . | . | . |
| . | . | . | . | . | . | . | . | . | T | . | . | . | . | . | . | . | . | . | . | . | . | . | . | . | . |
| . | . | . | . | T | . | . | . | . | . | . | . | . | . | . | . | . | . | . | . | . | . | . | . | . | . |
| . | . | . | . | T | . | . | . | . | . | . | . | . | . | . | . | . | . | . | . | . | . | . | . | . | . |
| . | . | . | . | T | . | . | . | . | . | . | . | . | . | . | . | . | . | . | . | . | . | . | . | . | . |
| . | . | . | . | T | . | . | . | . | . | . | . | . | . | . | . | . | . | . | . | . | . | . | . | . | . |
| . | . | . | . | T | . | . | . | . | . | . | . | . | . | . | . | . | . | . | . | . | . | . | . | . | . |
| . | . | . | . | T | . | . | . | . | . | . | . | . | . | . | . | . | . | . | . | . | . | . | . | . | . |
| . | . | . | . | . | . | . | . | . | . | . | . | . | . | . | . | . | . | . | . | . | . | . | . | . | . |
| . | . | . | . | . | . | . | . | . | . | . | . | . | . | . | . | . | . | . | . | . | . | . | . | . | . |
| . | . | . | . | . | . | . | . | . | T | . | . | . | . | . | . | . | . | . | . | . | . | . | . | . | . |
| . | . | . | . | . | . | . | . | . | T | . | . | . | . | . | . | . | . | . | . | . | . | . | . | . | . |
| . | . | . | . | . | . | . | . | . | T | . | . | . | . | . | . | . | . | . | . | . | . | . | . | . | . |
| . | . | . | . | . | . | . | . | . | T | . | . | . | . | . | . | . | . | . | . | . | . | . | . | . | . |
| . | . | . | . | . | . | . | . | . | T | . | . | . | . | . | . | . | . | . | . | . | . | . | . | . | . |
| . | . | . | . | . | . | . | . | . | T | . | . | . | . | . | . | . | . | . | . | . | . | . | . | . | . |
| . | . | . | . | . | . | . | . | . | T | . | . | . | . | . | . | . | . | . | . | . | . | . | . | . | . |
| . | . | . | . | . | . | . | . | . | T | . | . | . | . | . | . | . | . | . | . | . | . | . | . | . | . |
| . | . | . | . | . | . | . | . | . | T | . | . | . | . | . | . | . | . | . | . | . | . | . | . | . | . |
| . | . | . | . | . | . | . | . | . | T | . | . | . | . | . | . | . | . | . | . | . | . | . | . | . | . |
| . | . | . | . | . | . | . | . | . | T | . | . | . | . | . | . | . | . | . | . | . | . | . | . | . | . |
| . | . | . | . | . | . | . | . | . | T | . | . | . | . | . | . | . | . | . | . | . | . | . | . | . | . |
| . | . | . | . | . | . | . | . | . | T | . | . | . | . | . | . | . | . | . | . | . | . | . | . | . | . |
| . | . | . | . | . | . | . | . | . | T | . | . | . | . | . | . | . | . | . | . | . | . | . | . | . | . |
| . | . | . | . | T | . | . | . | . | T | . | . | . | . | . | . | . | . | . | . | . | . | . | . | . | . |
| . | . | . | . | . | . | . | . | . | . | . | . | . | . | . | . | . | . | . | . | . | . | . | . | . | . |
| . | . | . | . | . | . | . | . | . | T | . | . | . | . | . | . | . | . | . | . | . | . | . | . | . | . |
| . | . | . | . | . | . | . | . | . | T | . | . | . | . | . | . | . | . | . | . | . | . | . | . | . | . |
| . | . | . | . | . | . | . | . | . | T | . | . | . | . | . | A | . | . | . | . | . | . | . | . | . | . |
| . | . | . | . | . | . | . | . | . | . | . | . | . | . | . | . | . | . | . | . | . | . | . | . | . | . |

| | | | | | | | | | | | | | | | | | | | | | | | | | |
|---|---|---|---|---|---|---|---|---|---|---|---|---|---|---|---|---|---|---|---|---|---|---|---|---|---|
| . | . | . | . | . | . | . | . | . | T | . | . | . | . | . | . | . | . | . | . | . | . | . | . | . | . |
| . | . | . | . | . | . | . | . | . | T | . | . | . | . | . | . | . | . | . | . | . | . | . | . | . | . |
| . | . | . | . | . | . | . | . | . | T | . | . | . | . | . | . | . | . | . | . | . | . | . | . | . | . |
| . | . | . | . | . | . | . | . | . | T | . | . | . | . | . | . | . | . | . | . | . | . | . | . | . | . |
| . | . | . | . | . | . | . | . | . | T | . | . | . | . | . | . | . | . | . | . | . | . | . | . | . | . |
| . | . | . | . | T | . | . | . | . | . | . | . | . | . | . | . | . | . | . | . | . | . | . | . | . | . |
| . | . | . | . | . | . | . | . | . | . | . | . | . | . | . | . | . | . | . | . | . | . | . | . | . | . |
| . | . | . | . | . | . | . | . | . | T | . | . | . | . | . | . | . | . | . | . | . | . | . | . | . | . |
| . | . | . | . | T | . | . | . | . | T | . | . | . | . | . | . | . | . | . | . | . | . | . | . | . | . |
| . | . | . | . | . | . | . | . | . | T | . | . | . | . | . | . | . | . | . | . | . | . | . | . | . | . |
| . | . | . | . | . | . | . | . | . | T | . | . | . | . | . | . | . | . | . | . | . | . | . | . | . | . |
| . | . | T | . | . | . | . | . | . | T | . | . | . | . | . | . | . | . | . | . | . | . | . | . | . | . |
| . | . | . | . | . | . | . | . | . | T | . | . | . | . | . | . | . | . | . | . | . | . | . | . | . | . |
| . | . | . | . | . | . | . | . | . | T | . | . | . | . | . | A | . | . | . | . | . | . | . | . | . | . |
| . | . | . | . | . | . | . | . | . | T | . | . | . | . | . | A | . | . | . | . | . | . | . | . | . | . |
| . | . | . | . | . | . | . | . | . | . | . | . | . | . | . | . | . | . | . | . | . | . | . | . | . | . |
| . | . | . | . | . | . | . | . | . | T | . | . | . | . | . | . | . | . | . | . | . | . | . | . | . | . |
| . | . | T | . | . | . | . | . | . | T | . | . | . | . | . | . | . | . | . | . | . | . | . | . | . | . |
| . | . | T | . | . | . | . | . | . | T | . | . | . | . | . | . | . | . | . | . | . | . | . | . | . | . |
| . | . | . | . | . | . | . | . | . | T | . | . | . | . | . | . | . | . | . | . | . | . | . | . | . | . |
| . | . | T | . | . | . | . | . | . | T | . | . | . | . | . | . | . | . | . | . | . | . | . | . | . | . |
| . | . | . | . | T | . | . | . | . | T | . | . | . | . | . | . | . | . | . | . | . | . | . | . | . | . |
| . | . | . | . | . | . | . | . | . | T | . | . | . | . | . | . | . | . | . | . | . | . | . | . | . | . |
| . | . | . | . | . | . | . | . | . | . | . | . | . | . | . | . | . | . | . | . | . | . | . | . | . | . |
| . | . | . | . | . | . | . | . | . | . | . | . | . | . | . | . | . | . | . | . | . | . | . | . | . | . |
| . | . | . | . | . | . | . | . | . | . | . | . | . | . | . | . | . | . | . | . | . | . | . | . | . | . |
| . | . | . | . | . | . | . | . | . | T | . | . | . | T | . | . | . | . | . | . | . | . | C | . | . | . |
| . | . | . | . | . | . | . | . | . | T | . | . | . | . | . | . | . | . | . | . | . | . | . | . | . | . |
| . | . | . | . | . | . | . | . | . | T | . | . | . | . | . | . | . | . | . | . | . | . | . | . | . | . |
| . | . | . | . | . | . | . | . | . | . | . | . | . | . | . | . | . | . | . | . | . | . | . | . | . | . |
| . | . | . | . | . | . | . | . | . | . | . | . | . | . | . | . | . | . | . | . | . | . | . | . | . | . |
| . | . | . | . | . | . | . | . | . | . | . | . | . | . | . | . | . | . | . | . | . | . | . | . | . | . |
| . | . | . | . | . | . | . | . | . | . | . | . | . | . | . | . | . | . | . | . | . | . | . | . | . | . |
| . | . | . | . | . | . | . | . | . | . | . | . | . | . | . | . | . | . | . | . | . | . | . | . | . | . |
| . | . | . | . | . | . | . | . | . | T | . | . | . | T | . | . | . | . | . | . | . | . | C | . | . | . |
| . | . | . | . | . | . | . | . | . | . | . | . | . | . | . | . | . | . | . | . | . | . | . | . | . | . |
| . | . | . | . | . | . | . | . | . | . | . | . | . | . | . | . | . | . | . | . | . | . | . | . | . | . |
| . | . | . | . | T | . | . | . | . | . | . | . | . | . | . | . | . | T | . | . | . | . | . | . | . | . |
| . | . | . | . | . | . | . | . | . | . | . | . | . | . | . | . | . | . | . | . | . | . | . | . | . | . |
| . | . | . | . | . | . | . | . | . | T | . | . | . | . | . | . | . | . | . | . | . | . | . | . | . | . |
| . | . | . | . | . | . | . | . | . | T | . | . | . | . | . | . | . | . | . | . | . | . | . | . | . | . |
| . | . | . | . | . | . | . | . | . | T | . | . | . | . | . | . | . | . | . | . | . | . | . | . | . | . |
| . | . | . | . | . | . | . | . | . | T | . | . | . | . | . | . | . | . | . | . | . | . | . | . | . | . |
| . | . | . | . | . | . | . | . | . | T | . | . | . | . | . | . | . | . | . | . | . | . | . | . | . | . |
| . | . | . | . | . | . | . | . | . | T | . | . | . | . | . | . | . | . | . | . | . | . | . | . | . | . |
| . | . | . | . | . | . | . | . | . | T | . | . | . | . | . | . | . | . | . | . | . | . | . | . | . | . |
| . | . | . | . | . | . | . | . | . | T | . | . | . | . | . | . | . | . | . | . | . | . | . | . | . | . |
| . | . | . | . | . | . | . | . | . | T | . | . | . | . | . | . | . | . | . | . | . | . | . | . | . | . |
| . | . | . | . | . | . | . | . | . | T | . | . | . | . | . | . | . | . | . | . | . | . | . | . | . | . |
| . | . | . | . | . | . | . | . | . | T | . | . | . | . | . | . | . | . | . | . | . | . | . | . | . | . |
| . | . | . | . | . | . | . | . | . | T | . | . | . | . | . | . | . | . | . | . | . | . | . | . | . | . |
| . | . | . | . | . | . | . | . | . | T | . | . | . | . | . | . | . | . | . | . | . | . | . | . | . | . |
| . | . | . | . | . | . | . | . | . | T | . | . | . | . | . | . | . | . | . | . | . | . | . | . | . | . |
| . | . | . | . | . | . | . | . | . | T | . | . | . | . | . | . | . | . | . | . | . | . | . | . | . | . |
| . | . | . | . | . | . | . | . | . | T | . | . | . | . | . | . | . | . | . | . | . | . | . | . | . | . |
| . | . | . | . | . | . | . | . | . | T | . | . | . | . | . | . | . | . | . | . | . | . | . | . | . | . |
| . | . | . | . | T | . | . | . | . | . | . | . | . | . | . | . | . | . | . | . | . | . | . | . | . | . |
| . | . | . | . | . | . | . | . | . | T | . | . | . | . | . | . | . | . | . | . | . | . | . | . | . | . |
| . | . | . | . | . | . | . | . | . | T | . | . | . | . | . | . | . | . | . | . | . | . | . | . | . | . |
| . | . | . | . | . | . | . | . | . | T | . | . | . | . | . | . | . | . | . | . | . | . | . | . | . | . |
| . | . | . | . | . | . | . | . | . | T | . | . | . | . | . | . | . | . | . | . | . | . | . | . | . | . |
| . | . | . | . | . | . | . | . | . | T | . | . | . | . | . | . | . | . | . | . | . | . | . | . | . | . |

```
. . . . . . . . . T . . . . . . . . . . . . . . . .
. . . . . . . . . T . . . . . . . . . . . . . . . .
. . . . . . . . . T . . . . . . . . . . . . . . . .
. . . . . . . . . T . . . . . . . . . . . . . . . .
. . . . . . . . . T . . . . . . . . . . . . . . . .
. . . . . . . . . T . . . . . . . . . . . . . . . .
. . . . . . . . . T . . . . . . . . . . . . . . . .
. . . . . . . . . T . . . . . . . . . . . . . . . .
. . . . . . . . . . . . . . . . . . . . . . . . . .
. . . . . . . . . T . . . . . . . . . . . . . . . .
. . . . . . . . . T . . . T . . . . . . . . . . . .
. . . . . . . . . T . . . T . . . . . . . . . . . .
. . . . . . . . . T . . . T . . . . . . . . . . . .
. . . . . . . . . T . . . . . . . . . . . T . . . .
. . . . . . . . . T . . . . . . . . . . . T . . . .
. . . . . . . . . T . . . . . . . . . . . T . . . .
. . . . . . . . . T . . . . . . . . . . . T . . . .
. . . . . . . . . T . . . . . . . . . . . T . . . .
. . . . . . . . . T . . . . . . . . . . . T . . . .
. . . . . . . . . T . . . . . . . . . . . T . . . .
. . . . . . . . . T . . . . . . . . . . . T . . . .
. . . . T . . . . . . . . . . . . . . . . . . . . .
. . . . T . . . . . . . . . . . . . . . . . . . . .
. . . . . . . . . T . . . . . . . . . . . . . . . .
. . . . . . . . . T . . . . . . . . . . . . . . . .
. . . . T . . . . . . . . . . . . . . . . . . . . .
. . . . T . . . . . . . . . . . . . . . . . . . . .
. . . . T . . . . . . . . . . . . . . . . . . . . .
. . . . T . . . . . . . . . . . . . . . . . . . . .
. . . . T . . . . . . . . . . . . . . . . . . . . .
. . . . T . . . . T . . . . . . . . . . . . . . . .
. . . . T . . . . . . . . . . . . . . . . . . . . .
. . . . T . . . . . . . . . . . . . . . . . . . . .
. . . . T . . . . . . . . . . . . . . . . . . . . .
. . . . T . . . . . . . . . . . . . . . . . . . . .
. . . . T . . . . . . . . . . . . . . . . . . . . .
. T . . T . . . . . . . . . . . . . . . . . . . . .
. . . . . . . . . . . . . . . . . . . . . . . . . .
. . . . . . . . . . . . . . . . . . . . . . . . . .
. . . . . . . . . T . . . . . . . . . . . . . . . .
. . . . . . . . . T . . . . . . . . . . . . . . . .
. . . . . . . . . T . . . . . . . . . . . . . . . .
. . . . . . . . . T . . . . . . . . . . . . . . . .
. . . . . . . . . T . . . . . . . . . . . . . . . .
. . . . . . . . . T . . . . . . . . . . . . . . . .
. . . . . . . . . T . . . . . . . . . . . . . . . .
. . . . . . . . . T . . . . . . . . . . . . . . . .
. . . . . . . . . T . . . . . . . . . . . . . . . .
. . . . . . . . . T . . . . . . . . . . . . . . . .
. . . . . . . . . T . . . . . . . . . . . . . . . .
. . . . . . . . . T . . . . . . . . . . . . . . . .
. . . . . . . . . T . . . . . . . . . . . . . . . .
. . . . . . . . . T . . . . . . . . . . . . . . . .
. . . . . . . . . T . . . . . . . . . . . . . . . .
. . . . . . . . . T . . . . . . . . . . . . . . . .
. . . . . . . . . T . . . . . . . . . . . . . . . .
. . . . . . . . . T . . . . . . . . . . . . . . . .
. . . . . . . . . T . . . . . . . . . . . . . . . .
. . . . . . . . . T . . . . . . . . . . . . . . . .
. . . . . . . . . T . . . . . . . . . . . . . . . .
. . . . . . . . . T . . . T . . . . . . . . . . . .
. . . . . . . . . T . . . . . . . . . . . . . . . .
```

| | | | | | | | | | | | | | | | | | | | | | | | | | |
|---|---|---|---|---|---|---|---|---|---|---|---|---|---|---|---|---|---|---|---|---|---|---|---|---|---|
| . | . | . | . | . | . | . | . | . | . | . | . | . | . | . | . | . | . | . | . | . | . | . | . | . | . |
| . | . | . | . | . | . | . | . | . | T | . | . | . | . | . | . | . | . | . | . | . | . | . | . | . | . |
| . | . | . | . | . | . | . | . | . | T | . | . | . | . | . | . | . | . | . | . | . | . | . | . | . | . |
| . | . | . | . | . | . | . | . | . | T | . | . | . | . | . | . | . | . | . | . | . | . | . | . | . | . |
| . | . | . | . | . | . | . | . | . | . | . | . | . | . | . | . | . | . | . | . | . | . | . | . | . | . |
| . | . | . | . | T | . | . | . | . | . | . | . | . | . | . | . | . | . | . | . | . | . | . | . | . | . |
| . | . | . | . | Y | . | . | . | . | Y | . | . | . | . | . | . | . | . | . | . | . | . | . | . | . | . |
| . | . | . | . | Y | . | . | . | . | Y | . | . | . | . | . | . | . | . | . | . | . | . | . | . | . | . |
| . | . | . | . | T | . | . | . | . | . | . | . | . | . | R | . | . | . | . | . | . | . | . | . | . | . |
| . | . | . | . | T | . | . | . | . | . | . | . | . | . | . | . | . | . | . | . | . | . | . | . | . | . |
| . | . | . | . | T | . | . | . | . | . | . | . | . | . | R | . | . | . | . | . | . | . | . | . | . | . |
| . | . | . | . | T | . | . | . | . | . | . | . | . | . | R | . | . | . | . | . | . | . | . | . | . | . |
| . | . | . | . | T | . | . | . | . | . | . | . | . | . | R | . | . | . | . | . | . | . | . | . | . | . |
| . | . | . | . | T | . | . | . | . | . | . | . | . | . | R | . | . | . | . | . | . | . | . | . | . | . |
| . | . | . | . | T | . | . | . | . | . | . | . | . | . | R | . | . | . | . | . | . | . | . | . | . | . |
| . | . | . | . | T | . | . | . | . | . | . | . | . | . | . | . | . | . | . | . | . | . | . | . | . | . |
| . | . | . | . | T | . | . | . | . | . | . | . | . | . | ? | . | . | . | . | . | . | . | . | . | . | . |
| . | . | . | . | Y | . | . | . | . | ? | . | . | . | . | . | . | . | . | . | . | . | . | . | . | . | . |
| . | . | . | . | T | . | . | . | . | . | . | . | . | . | ? | . | . | . | . | . | . | . | . | . | . | . |
| . | . | . | . | T | . | . | . | . | . | . | . | . | . | . | . | . | . | . | . | . | . | . | . | . | . |
| . | . | . | . | T | . | . | . | . | . | . | . | . | . | R | . | . | . | . | . | . | . | . | . | . | . |
| . | . | . | . | T | . | . | . | . | . | . | . | . | . | R | . | . | . | . | . | . | . | . | . | . | . |
| . | . | . | . | T | . | . | . | . | . | . | . | . | . | R | . | . | . | . | . | . | . | . | . | . | . |
| . | . | . | . | T | . | . | . | . | . | . | . | . | . | . | . | . | . | . | . | . | . | . | . | . | . |
| . | . | . | . | T | . | . | . | . | . | . | . | . | . | R | . | . | . | . | . | . | . | . | . | . | . |
| . | . | . | . | T | . | . | . | . | . | . | . | . | . | . | . | . | . | . | . | . | . | . | . | . | . |
| . | . | . | . | T | . | . | . | . | . | . | . | . | . | R | . | . | . | . | . | . | . | . | . | . | . |
| . | . | . | . | T | . | . | . | . | . | . | . | . | . | R | . | . | . | . | . | . | . | . | . | . | . |
| . | . | . | . | T | . | . | . | . | . | . | . | . | . | . | . | . | . | . | . | . | . | . | . | . | . |
| . | . | . | . | T | . | . | . | . | . | . | . | . | . | R | . | . | . | . | . | . | . | . | . | . | . |
| . | . | . | . | T | . | . | . | . | . | . | . | . | . | R | . | . | . | . | . | . | . | . | . | . | . |
| . | . | . | . | . | . | . | . | . | T | . | . | . | . | . | . | . | . | . | . | T | . | . | . | . | . |
| . | . | . | . | . | . | . | . | . | T | . | . | . | . | . | . | . | . | . | . | T | . | . | . | . | . |
| . | . | . | . | . | . | . | . | . | T | . | . | . | . | . | . | . | . | . | . | . | . | . | . | . | . |
| . | . | . | . | . | . | . | . | . | T | . | . | . | T | . | . | . | . | . | . | . | . | . | . | . | . |
| . | . | . | . | . | . | . | . | . | . | . | . | . | . | . | . | . | . | . | . | . | . | . | . | . | . |
| . | . | . | . | . | . | . | . | . | T | . | . | . | T | . | . | . | . | . | . | . | . | . | . | . | . |
| . | . | . | . | . | . | . | . | . | T | . | . | . | . | . | . | . | . | . | . | . | . | . | . | . | . |
| . | . | . | . | . | . | . | . | . | T | . | . | . | . | . | . | . | . | A | . | . | . | . | . | . | . |
| . | . | . | . | . | . | . | . | . | T | . | . | . | . | . | . | . | . | . | . | . | . | . | . | . | . |
| . | . | . | . | . | . | . | . | . | T | . | . | . | T | . | . | . | . | . | . | . | . | . | . | . | . |
| . | . | . | . | . | . | . | . | . | T | . | . | . | . | . | . | . | . | . | . | . | . | . | . | . | . |
| . | . | . | . | . | . | . | . | . | T | . | . | . | . | . | . | . | . | . | . | . | . | . | . | . | . |
| . | . | . | . | . | . | . | . | . | T | . | . | . | T | . | . | . | . | . | . | . | . | . | . | . | . |
| . | . | . | . | . | . | . | . | . | T | . | . | . | T | . | . | . | . | . | . | . | . | . | . | . | . |
| . | . | . | . | . | . | . | . | . | T | . | . | . | . | . | . | . | . | . | . | . | . | . | . | . | . |
| . | . | . | . | . | . | . | . | . | T | . | . | . | T | . | . | . | . | . | . | . | . | . | . | . | . |
| . | . | . | . | . | . | . | . | . | T | . | . | . | . | . | . | . | . | . | . | . | . | . | . | . | . |
| . | . | . | . | . | G | . | . | . | T | . | . | . | . | . | . | . | . | A | . | . | . | . | . | . | . |
| . | . | . | . | . | . | . | . | . | T | . | . | . | T | . | . | . | . | . | . | . | . | . | . | . | . |
| . | . | . | . | . | . | . | . | . | T | . | G | . | . | . | . | . | . | . | . | . | . | . | . | . | . |
| . | . | . | . | . | . | . | . | . | T | . | G | . | . | . | . | . | . | . | . | . | . | . | . | . | . |
| . | . | . | . | . | . | . | . | . | T | . | . | . | . | . | . | . | . | . | . | . | . | . | . | . | . |
| . | . | . | . | . | . | . | . | . | T | . | . | . | . | . | . | . | . | . | . | . | . | . | . | . | . |
| . | . | . | . | . | . | . | . | . | T | . | . | . | . | . | . | . | . | . | . | . | . | . | . | . | . |
| . | . | . | . | . | . | . | . | . | T | . | . | . | . | . | . | . | . | . | . | . | . | . | . | . | . |
| . | . | . | . | . | . | . | . | . | T | . | . | . | T | . | . | . | . | . | . | . | . | . | . | . | . |
| . | . | . | . | . | . | . | . | . | T | . | . | . | . | . | . | . | . | . | . | . | . | . | . | . | . |
| . | . | . | . | . | . | . | . | . | T | . | . | . | . | . | . | . | . | . | . | . | . | . | . | . | . |
| . | . | . | . | . | . | . | . | . | T | . | . | . | T | . | . | . | . | . | . | . | . | . | . | . | . |

| 134 | 135 | 136 | 137 | 138 | 139 | 140 | 141 | 142 | 143 | 144 | 145 | 146 | 147 | 148 | 149 | 150 | 151 | 152 | 153 | 154 | 155 | 156 | 157 | 158 | 159 |
|---|---|---|---|---|---|---|---|---|---|---|---|---|---|---|---|---|---|---|---|---|---|---|---|---|---|
| ORF1ab | ORF1ab | ORF1ab | ORF1ab | ORF1ab | ORF1ab | ORF1ab | ORF1ab | ORF1ab | ORF1ab | ORF1ab | ORF1ab | ORF1ab | ORF1ab | ORF1ab | ORF1ab | ORF1ab | ORF1ab | ORF1ab | ORF1ab | ORF1ab | ORF1ab | ORF1ab | ORF1ab | ORF1ab | ORF1ab |
| 17135 | 17440 | 17722 | 17747 | 17858 | 17959 | 18052 | 18129 | 18380 | 18395 | 18457 | 18486 | 18511 | 18877 | 19086 | 19185 | 19344 | 19417 | 19524 | 19679 | 19684 | 19816 | 19872 | 19983 | 20006 | 20063 |
| C | C | G | C | A | A | A | C | G | C | C | C | C | C | G | C | T | G | C | G | G | G | G | C | G | T |
| . | . | . | . | . | . | . | . | . | . | . | . | . | . | . | . | . | . | . | . | . | . | . | . | . | . |
| . | . | . | . | . | . | . | . | . | . | . | . | . | . | . | . | . | . | . | . | . | . | . | . | . | . |
| . | . | . | . | . | . | . | . | . | . | . | . | . | . | . | . | . | . | . | . | . | . | . | . | . | . |
| . | . | . | . | . | . | . | . | . | . | . | . | . | . | . | . | . | . | . | . | . | . | . | . | . | . |
| . | . | . | . | . | . | . | . | . | . | . | . | . | . | . | . | . | . | . | . | . | . | . | . | . | . |
| . | . | . | . | . | . | . | . | . | . | . | . | . | . | . | . | . | . | . | . | . | . | . | . | . | . |
| . | . | . | . | . | . | . | . | . | . | . | . | . | . | . | . | . | . | . | . | . | . | . | . | . | . |
| . | . | . | . | . | . | . | . | . | . | . | . | . | . | . | . | . | . | . | . | . | . | . | . | . | . |
| . | . | . | . | . | . | . | . | . | . | . | . | . | . | . | . | . | . | . | . | . | . | . | . | . | . |
| . | . | . | . | . | . | . | . | . | . | . | . | . | . | . | . | . | . | . | . | . | . | . | . | . | . |
| . | . | . | . | . | . | . | . | . | . | . | . | . | . | . | . | . | . | . | . | . | . | . | . | . | . |
| . | . | . | . | . | . | . | . | . | . | . | . | . | . | . | . | . | . | . | . | . | . | . | . | . | . |
| . | . | . | . | . | . | . | . | . | . | . | . | . | . | . | . | . | . | . | . | . | . | . | . | . | . |
| . | . | . | . | . | . | . | . | . | . | . | . | . | . | . | . | . | . | . | . | . | . | . | . | . | . |
| . | . | . | . | . | . | . | . | . | . | . | . | . | . | . | . | . | . | . | . | . | . | . | . | . | . |
| . | . | . | . | . | . | . | . | . | . | . | . | . | . | . | . | . | . | . | . | . | . | . | . | . | . |
| . | . | . | . | . | . | . | . | . | . | . | . | . | . | . | . | . | . | . | . | . | . | . | . | . | . |
| . | . | . | . | . | . | . | . | . | . | . | . | . | . | . | . | . | . | . | . | . | . | . | . | . | . |
| . | . | . | . | . | . | . | . | . | . | . | . | . | . | . | . | . | . | . | . | . | . | . | . | . | . |
| . | . | . | . | . | . | . | . | . | . | . | . | . | . | . | . | . | . | . | . | . | . | . | . | . | . |
| . | . | . | . | . | . | . | . | . | . | . | . | . | . | . | . | . | . | . | . | . | . | . | . | . | . |
| . | . | . | . | . | . | . | . | . | . | . | . | . | . | . | . | . | . | . | . | . | . | . | . | . | . |
| . | . | . | . | . | . | . | . | . | . | . | . | . | . | . | . | . | . | . | . | . | . | . | . | . | . |
| . | . | . | . | . | . | . | . | . | . | . | . | . | . | . | . | . | . | . | . | . | . | . | . | . | . |
| . | . | . | . | . | . | . | . | . | . | . | . | . | . | . | . | . | . | . | . | . | . | . | . | . | . |
| . | . | . | . | . | . | . | . | . | . | . | . | . | . | . | . | . | . | . | . | . | . | . | . | . | . |
| . | . | . | . | . | . | . | . | . | . | . | . | . | . | . | . | . | . | . | . | . | . | . | . | . | . |
| . | . | . | . | . | . | . | . | . | . | . | . | . | . | . | . | . | . | . | . | . | . | . | . | . | . |
| . | . | . | . | . | . | . | . | . | . | . | . | . | . | . | . | . | . | . | . | . | . | . | . | . | . |
| . | . | . | . | . | . | . | . | . | . | . | . | . | . | . | . | . | . | . | . | . | . | . | . | . | . |
| . | . | . | . | . | . | . | . | . | . | . | . | . | . | . | . | . | . | . | . | . | . | . | . | . | . |
| . | . | . | . | . | . | . | . | . | . | . | . | . | . | . | . | . | . | . | . | . | . | . | . | . | . |
| . | . | . | . | . | . | . | . | . | . | . | . | . | . | . | . | . | . | . | . | . | . | . | . | . | . |
| . | . | . | . | . | . | . | . | . | . | . | . | . | . | . | . | . | . | . | . | . | . | . | . | . | . |
| . | . | . | . | . | . | . | . | . | . | . | . | . | . | . | . | . | . | . | . | . | . | . | . | . | . |
| . | . | . | . | . | . | . | . | . | . | . | . | . | . | . | . | . | . | . | . | . | . | . | . | . | . |
| . | . | . | . | . | . | . | . | ? | ? | ? | ? | ? | . | . | . | . | . | . | . | . | . | . | . | . | . |
| . | . | . | . | . | . | . | . | . | . | . | . | . | . | . | . | . | . | . | . | . | . | . | . | . | . |
| . | . | . | . | . | . | . | . | . | . | . | . | . | . | . | . | . | . | . | . | . | . | . | . | . | . |
| . | . | . | . | . | . | . | . | . | . | . | . | . | . | . | . | . | . | . | . | . | . | . | . | . | . |
| . | . | . | . | . | . | . | . | . | . | . | . | . | . | . | . | . | . | . | . | . | . | . | . | . | . |
| . | . | . | . | . | . | . | . | ? | ? | ? | ? | ? | . | . | . | . | . | . | ? | ? | . | . | . | . | . |
| . | . | . | . | . | . | . | . | . | . | . | . | . | . | . | . | . | . | . | . | . | . | . | . | . | . |
| . | . | . | . | . | . | . | . | . | . | . | . | . | . | . | . | . | . | . | . | . | . | . | . | . | . |
| . | . | . | . | . | . | . | . | . | . | . | . | . | . | . | . | . | . | . | . | . | . | . | . | . | . |
| . | . | . | . | . | . | . | . | . | . | . | . | . | . | . | . | . | . | . | . | . | . | . | . | . | . |
| . | . | . | . | . | . | . | . | . | . | . | . | . | . | . | . | . | . | . | . | . | . | . | . | . | . |
| . | . | . | . | . | . | . | . | . | . | . | . | . | . | . | . | . | . | . | . | . | . | . | . | . | . |
| . | . | . | . | . | . | . | . | . | . | . | . | . | . | . | . | . | . | . | . | . | . | . | . | . | . |
| . | . | . | . | . | . | . | . | . | . | . | . | . | . | . | . | . | . | . | . | . | . | . | . | . | . |
| . | . | . | . | . | . | . | . | . | . | . | . | . | . | . | . | . | . | Y | . | . | . | . | . | . | . |
| . | . | . | . | . | . | . | . | . | . | . | . | . | . | . | . | . | . | . | . | . | . | . | . | . | . |
| . | . | . | . | . | . | . | . | . | . | . | . | . | . | . | . | . | . | . | . | . | . | . | . | . | . |
| . | . | . | . | . | . | . | . | . | . | . | . | . | . | . | . | . | . | . | . | . | . | . | . | . | . |

| | | | | | | | | | | | | | | | | | | | | | | | | | |
|---|---|---|---|---|---|---|---|---|---|---|---|---|---|---|---|---|---|---|---|---|---|---|---|---|---|
| . | . | . | . | . | . | . | . | . | . | . | . | . | . | . | . | . | . | . | . | . | . | . | . | . | . |
| . | . | . | T | . | . | . | . | . | . | . | . | . | . | . | . | . | . | . | . | . | . | . | . | . | . |
| . | . | . | . | . | . | . | . | . | . | . | . | . | . | . | . | . | . | . | . | . | . | T | . | . | C |
| . | . | . | . | . | . | . | . | . | . | . | . | . | . | . | . | . | . | . | . | . | . | . | . | . | . |
| . | . | . | . | . | . | . | . | . | . | . | . | . | T | . | . | . | . | . | . | . | . | . | . | . | . |
| . | . | . | . | . | . | . | . | . | . | . | . | . | . | . | . | . | . | . | . | . | . | . | . | . | . |
| . | . | . | . | . | . | . | . | . | . | . | . | . | . | . | . | . | . | . | . | . | . | . | . | . | . |
| . | . | . | . | . | . | . | . | . | . | . | . | . | . | . | . | . | . | . | . | . | . | . | . | . | . |
| . | . | . | . | . | . | . | . | . | . | . | . | . | . | . | . | . | . | . | . | . | . | . | . | . | . |
| . | . | . | . | . | . | . | . | . | . | . | . | . | . | . | . | . | . | . | . | . | . | . | . | . | . |
| . | . | . | . | . | . | . | . | . | . | . | . | . | . | . | . | . | . | . | . | . | . | . | . | . | . |
| . | . | . | . | . | . | . | . | . | . | . | . | . | T | . | . | . | . | . | . | . | . | . | . | . | . |
| . | . | . | . | . | . | . | . | . | . | . | . | . | T | . | . | . | . | . | . | . | . | . | . | . | . |
| . | . | . | . | a | . | . | . | . | . | . | . | . | T | . | . | . | . | . | . | . | . | . | . | . | . |
| . | . | . | . | . | . | . | . | . | . | . | . | . | T | . | . | . | . | . | . | . | . | . | . | . | . |
| . | . | . | . | . | . | . | . | . | . | . | . | . | T | . | . | . | . | . | . | . | . | . | . | . | . |
| . | . | . | . | . | . | . | . | . | . | . | . | . | . | . | . | . | . | . | . | . | . | . | . | . | . |
| . | . | . | . | . | . | . | . | . | T | . | . | . | . | . | . | . | . | . | . | . | . | . | . | . | . |
| . | . | . | . | . | . | . | . | . | . | . | . | . | Y | . | . | . | . | . | . | . | . | . | . | . | . |
| . | . | . | . | . | . | . | . | . | . | . | . | . | . | . | . | . | . | ? | . | . | . | . | . | . | . |
| . | . | . | . | . | . | . | . | . | . | . | . | . | . | . | . | . | . | . | . | . | . | . | . | . | . |
| . | . | . | . | . | . | . | . | . | . | . | ? | . | . | . | . | . | . | . | . | . | . | . | . | . | . |
| . | . | . | . | . | . | . | . | . | . | . | . | . | . | . | . | . | . | . | . | . | . | . | . | . | . |
| . | . | . | . | . | . | . | . | . | . | . | . | . | . | . | . | . | . | T | . | . | . | . | . | . | . |
| . | . | . | . | . | . | . | . | . | . | . | . | . | . | . | . | . | . | . | . | . | . | . | . | . | . |
| . | . | . | . | . | . | . | . | . | . | . | . | . | . | . | . | . | . | . | . | . | . | . | . | . | . |
| . | . | . | . | . | . | . | . | . | . | . | . | . | . | . | . | . | . | . | . | . | . | . | . | . | . |
| . | . | . | . | . | . | . | . | . | . | . | . | . | . | . | . | . | . | ? | . | . | . | . | . | . | . |
| . | . | . | . | . | . | . | . | . | . | . | Y | . | . | . | . | . | . | . | . | . | . | . | . | . | . |
| . | . | . | . | . | . | . | . | . | . | . | . | . | . | . | . | . | . | . | . | . | . | . | . | . | . |
| . | . | . | . | . | . | . | . | . | . | . | . | . | . | . | . | . | . | T | . | . | . | . | . | . | . |
| . | . | . | . | . | . | . | . | . | . | . | . | . | . | . | . | . | . | T | . | . | . | . | . | . | . |
| . | . | . | . | . | . | . | . | . | . | . | . | . | . | . | . | . | . | . | . | . | . | . | . | . | . |
| . | . | . | . | . | . | . | . | . | . | . | . | . | . | . | . | . | . | ? | . | . | . | . | . | . | . |
| . | . | . | . | . | . | . | . | . | . | . | ? | . | . | g | . | . | . | Y | . | . | . | . | . | . | . |
| . | . | . | . | . | . | . | . | . | . | . | . | . | . | . | . | . | . | . | . | . | . | . | . | . | . |
| . | . | . | . | . | . | . | . | . | . | . | . | . | . | . | . | . | . | . | . | . | . | . | . | . | . |
| . | . | . | . | . | . | . | . | . | . | . | . | . | . | . | . | . | . | . | . | . | . | . | . | . | . |
| . | . | . | . | . | . | . | . | . | . | . | . | . | . | . | . | . | . | . | . | . | . | . | . | . | . |
| . | . | . | . | . | . | . | . | . | . | . | . | . | . | g | . | . | . | . | . | . | . | . | . | . | . |
| . | . | . | . | . | . | . | . | . | . | . | . | . | . | . | . | . | . | . | . | . | . | . | . | . | . |
| . | . | . | . | . | . | . | . | . | . | . | . | . | . | . | . | . | . | . | . | . | . | . | . | . | . |
| . | . | . | . | . | . | . | . | . | . | . | . | . | . | . | . | . | . | . | . | . | . | . | . | . | . |
| . | . | . | . | . | . | . | . | . | . | . | . | . | . | . | . | . | . | . | . | . | . | . | . | . | . |
| . | . | . | . | . | . | . | . | . | . | . | . | . | . | . | . | . | . | . | . | . | . | . | . | . | . |
| . | . | . | . | . | . | . | . | . | . | . | . | . | . | . | . | . | . | . | . | . | . | . | . | . | . |
| . | . | . | . | . | . | . | . | . | . | . | . | . | . | . | . | . | . | . | . | . | . | . | . | . | . |
| . | . | . | . | . | . | . | . | . | . | . | . | . | . | . | . | . | . | . | . | . | . | . | . | . | . |
| . | . | . | . | . | . | . | . | . | . | . | . | . | . | . | . | . | . | . | . | . | . | . | . | . | . |
| . | . | . | Y | G | . | . | . | . | . | . | . | . | . | . | . | . | . | . | . | . | . | . | . | . | . |
| . | . | . | . | . | . | . | . | . | . | . | . | . | . | . | . | . | . | . | . | . | . | . | . | . | . |
| . | . | . | . | . | . | . | . | . | . | . | . | . | . | . | . | . | . | Y | . | . | . | . | . | . | . |
| . | . | . | . | . | . | . | . | . | T | . | . | . | . | . | . | ? | ? | ? | . | . | . | . | . | . | . |
| . | . | . | . | . | . | . | . | . | . | . | . | . | . | . | . | t | . | t | . | . | . | . | . | . | . |
| . | . | . | . | . | . | . | . | . | . | . | . | . | . | . | . | . | . | ? | . | . | . | . | . | . | . |
| . | . | . | . | . | . | . | . | . | . | . | Y | . | . | . | . | ? | ? | ? | . | . | . | . | . | . | . |
| . | . | . | . | . | . | . | . | . | . | . | . | . | . | . | . | t | g | ? | . | . | . | . | . | . | . |
| . | . | . | . | . | . | . | . | . | . | . | . | . | . | . | . | t | ? | ? | . | . | . | . | . | . | . |
| . | . | . | . | . | . | . | . | . | . | . | . | . | . | . | . | t | g | c | . | . | . | . | . | . | . |
| . | . | . | . | . | . | . | . | . | . | . | . | . | . | . | . | t | g | c | . | . | . | . | . | . | . |
| . | . | . | . | . | . | . | . | . | . | . | . | . | . | . | . | . | . | T | . | . | . | . | . | . | . |
| . | . | . | . | . | . | . | . | . | . | . | . | . | . | . | . | . | . | T | . | . | . | . | . | . | . |

| | | | | | | | | | | | | | | | | | | | | | | | | | |
|---|---|---|---|---|---|---|---|---|---|---|---|---|---|---|---|---|---|---|---|---|---|---|---|---|---|
| . | . | . | . | . | . | . | . | . | . | . | . | . | . | . | . | t | g | t | . | . | . | . | . | . | . |
| . | . | . | . | . | . | . | . | . | . | . | . | . | . | . | . | . | . | T | . | . | . | . | . | . | . |
| . | . | . | . | . | . | . | . | . | . | . | . | . | . | g | . | . | . | . | . | . | . | . | . | . | . |
| . | . | . | . | . | . | . | . | . | . | . | . | . | . | g | . | . | . | c | . | . | . | . | . | . | . |
| . | . | . | . | . | . | . | . | . | . | . | . | . | . | . | . | ? | g | c | . | . | . | . | . | . | . |
| . | . | . | . | . | . | . | . | . | . | . | . | . | . | . | . | . | g | t | . | . | . | . | . | . | . |
| . | . | . | . | . | . | . | . | . | . | . | . | . | . | . | . | ? | ? | ? | . | . | . | . | . | . | . |
| . | . | . | . | . | . | . | . | . | . | . | . | . | . | . | . | . | . | T | . | . | . | . | . | . | . |
| . | . | . | . | . | . | . | . | . | . | . | . | . | . | . | . | . | . | . | . | T | . | . | . | . | . |
| . | . | . | . | . | . | . | . | . | . | . | . | . | . | . | . | t | g | c | . | . | . | . | . | . | . |
| . | . | . | . | . | . | . | . | . | . | . | . | . | . | . | . | t | g | c | . | . | . | . | . | . | . |
| . | . | . | . | . | . | . | . | . | . | . | . | . | . | . | . | t | g | c | . | . | . | . | . | . | . |
| . | . | . | . | . | . | . | . | . | . | . | . | . | . | . | . | . | . | . | . | . | . | . | . | . | . |
| . | . | . | . | . | . | . | . | . | . | . | Y | . | . | . | . | t | g | c | . | . | . | . | . | . | . |
| . | . | . | . | . | . | . | . | . | . | . | . | . | . | . | . | t | g | t | . | . | . | . | . | . | . |
| . | . | . | . | . | . | . | . | . | . | . | . | . | . | g | . | t | g | c | . | . | . | . | . | . | . |
| . | . | . | . | . | . | . | . | . | . | . | . | . | . | . | . | . | . | T | . | . | . | . | . | . | . |
| . | . | . | . | . | . | . | . | . | . | . | . | . | . | . | . | . | . | T | . | . | . | . | . | . | . |
| . | . | . | . | . | . | . | . | . | . | . | . | . | . | . | . | . | . | . | . | . | . | . | . | . | . |
| . | . | . | . | . | . | . | . | . | . | . | . | . | . | . | . | . | . | T | . | . | . | . | . | . | . |
| . | . | . | . | . | . | . | . | . | . | . | . | . | . | . | . | t | g | c | . | . | . | . | . | . | . |
| . | . | . | . | . | . | . | . | . | . | . | . | . | . | . | . | . | . | . | . | . | . | . | . | . | . |
| . | . | . | . | . | . | . | . | . | . | . | . | . | . | . | . | t | g | c | . | . | . | . | . | . | . |
| . | . | . | . | . | . | . | . | . | . | . | . | . | . | . | . | t | g | c | . | . | . | . | . | . | . |
| . | . | . | . | . | . | . | . | . | . | . | . | . | . | g | . | . | . | . | . | T | . | . | . | . | . |
| . | . | . | . | . | . | . | . | . | . | . | . | . | . | . | . | . | . | . | . | . | . | . | . | . | . |
| . | . | . | . | . | . | . | . | . | . | . | . | . | . | . | . | . | . | . | . | . | . | . | . | . | . |
| . | . | . | . | . | . | . | . | . | . | . | . | . | T | . | . | . | . | . | . | . | . | . | . | . | . |
| . | . | . | . | . | . | . | . | g | c | c | . | . | . | . | . | . | . | . | . | . | . | . | . | . | . |
| . | . | . | . | . | . | . | . | . | . | . | . | . | . | . | . | . | . | . | . | . | . | . | . | . | . |
| . | . | . | . | . | . | . | . | . | . | . | . | . | . | . | . | . | . | . | . | . | . | . | . | . | . |
| . | . | . | . | . | . | . | . | . | . | . | . | . | . | . | . | ? | ? | ? | . | . | . | . | . | . | . |
| . | . | . | . | . | . | . | . | . | . | . | . | . | . | . | . | . | . | T | . | . | . | . | . | . | . |
| . | . | . | . | . | . | . | . | . | . | . | . | . | T | . | . | . | . | . | . | . | . | . | . | . | . |
| . | . | . | . | . | . | . | . | . | . | . | . | . | . | . | . | . | . | . | . | . | . | . | . | . | . |
| . | . | . | . | . | . | . | . | . | . | . | . | . | . | . | . | . | . | . | . | . | . | . | . | . | . |
| . | . | . | . | . | . | . | . | . | . | . | . | . | . | . | . | . | . | . | . | . | . | . | . | . | . |
| . | . | . | . | . | . | . | . | . | . | . | . | . | T | . | . | . | . | . | . | . | . | . | . | . | . |
| . | . | . | . | . | . | . | . | . | . | . | . | . | T | . | . | . | . | . | . | . | . | . | . | . | . |
| . | . | . | . | . | . | . | . | . | . | . | . | . | . | . | . | . | . | . | . | . | . | . | . | . | . |
| . | . | . | . | . | . | . | . | . | . | . | . | . | . | . | . | . | . | . | . | . | . | . | . | . | . |
| . | . | . | . | . | . | . | . | . | . | . | . | . | T | . | . | . | . | . | . | . | . | . | . | . | . |
| . | . | . | . | . | . | . | . | . | . | . | . | . | . | . | . | . | . | . | . | . | . | . | . | . | . |
| . | . | . | . | . | . | . | . | . | . | . | . | . | . | . | . | . | . | . | . | . | . | . | . | . | . |
| . | . | . | . | . | . | . | . | . | . | . | . | . | T | . | . | . | . | . | . | . | . | . | . | . | . |
| . | . | . | . | . | . | . | . | . | . | . | . | . | . | . | . | . | . | . | . | . | . | . | . | . | . |
| . | . | . | . | . | . | . | . | . | . | . | . | . | T | . | . | . | . | . | . | . | . | . | . | . | . |
| . | . | . | . | . | . | . | . | . | . | . | . | . | . | . | . | . | . | . | . | . | . | . | . | . | . |
| . | . | . | . | . | . | . | . | . | . | . | . | . | T | . | . | . | . | . | . | . | . | . | . | . | . |
| . | . | . | . | . | . | . | . | . | . | . | . | . | . | . | . | . | . | . | . | . | . | . | . | . | . |
| . | . | . | . | . | . | . | . | . | . | . | . | . | . | . | . | . | . | . | . | . | . | . | . | . | . |
| . | . | . | . | . | . | . | . | . | . | . | . | . | T | . | . | . | . | . | . | . | . | . | . | . | . |
| . | . | . | . | . | . | . | . | . | . | . | . | . | T | . | . | . | . | . | . | . | . | . | . | . | . |
| . | . | . | . | . | . | . | . | . | . | . | . | . | . | . | . | . | . | . | . | . | . | . | . | . | . |
| . | . | . | . | . | . | . | . | . | . | . | . | . | T | . | . | . | . | . | . | . | . | . | . | . | . |
| . | . | . | . | . | . | . | . | . | . | . | . | . | . | . | . | . | . | . | . | . | . | . | . | . | . |
| . | . | . | . | . | . | . | . | . | . | . | . | . | T | . | . | . | . | . | . | . | . | . | . | . | . |
| . | . | . | . | . | . | . | . | . | . | . | . | . | . | . | . | . | . | . | . | . | . | . | . | . | . |
| . | . | . | . | . | . | . | . | . | . | . | . | . | . | . | . | . | . | . | . | . | . | . | . | . | . |
| . | . | . | . | . | . | . | . | . | . | . | . | . | T | . | . | . | . | . | . | . | . | . | . | . | . |
| . | . | . | . | . | . | . | . | . | . | . | . | . | T | . | . | . | . | . | . | . | . | . | . | . | . |
| . | . | . | . | . | . | . | . | . | . | . | . | . | T | . | . | . | . | . | . | . | . | . | . | . | . |

| | | | | | | | | | | | | | | | | | | | | | | | | | |
|---|---|---|---|---|---|---|---|---|---|---|---|---|---|---|---|---|---|---|---|---|---|---|---|---|---|
| . | . | . | . | . | . | . | . | . | . | . | . | . | T | . | . | . | . | . | . | . | . | . | . | . | . |
| . | . | . | . | . | . | . | . | . | . | . | . | . | . | . | . | . | . | . | . | . | . | . | . | . | . |
| . | . | . | . | . | . | . | . | . | . | . | . | . | T | . | . | . | . | . | . | . | . | . | . | . | . |
| . | . | . | . | . | . | . | . | . | . | . | . | . | T | . | . | . | . | . | . | . | . | . | . | . | . |
| . | . | . | . | . | . | . | . | . | . | . | . | . | T | . | . | . | . | . | . | . | . | . | . | . | . |
| . | . | . | . | . | . | . | . | . | . | . | . | . | T | . | . | . | . | . | . | . | . | . | . | . | . |
| . | . | . | . | . | . | . | . | . | . | . | . | . | . | . | . | . | . | . | . | . | . | . | . | . | . |
| . | . | . | . | . | . | . | . | . | . | . | . | . | T | . | . | . | . | . | . | . | . | . | . | . | . |
| . | . | . | . | . | . | . | . | . | . | . | . | . | . | . | . | . | . | . | . | . | . | . | . | . | . |
| . | . | . | . | . | . | . | . | . | . | . | . | . | T | . | . | . | . | . | . | . | . | . | . | . | . |
| . | . | . | . | . | . | . | . | . | . | . | . | . | . | . | . | . | . | . | . | . | . | . | . | . | . |
| . | . | . | . | . | . | . | . | . | . | . | . | . | . | . | . | . | . | . | . | . | . | . | . | . | . |
| . | . | . | . | . | . | . | . | . | . | . | . | . | . | . | . | . | . | . | . | . | . | . | . | . | C |
| . | . | . | . | . | . | . | . | . | . | . | . | . | T | . | . | . | . | . | . | . | . | . | . | . | . |
| . | . | . | . | . | . | . | . | . | . | . | . | . | . | . | . | . | . | . | . | . | . | . | . | . | . |
| . | . | . | . | . | . | . | . | . | . | . | . | . | T | . | . | . | . | . | . | . | . | . | . | . | . |
| . | . | . | . | . | . | . | . | . | . | . | . | . | . | . | . | . | . | . | . | . | . | . | . | . | . |
| . | . | . | . | . | . | . | . | . | . | . | . | . | T | . | . | . | . | . | . | . | . | . | . | . | . |
| . | . | . | . | . | . | . | . | . | . | . | . | . | T | . | . | . | . | . | . | . | . | . | . | . | . |
| . | . | . | . | . | . | . | . | . | . | . | . | . | T | . | . | . | . | . | . | . | . | . | . | . | . |
| . | . | . | . | . | . | . | . | . | . | . | . | . | . | . | . | . | . | . | . | . | . | . | . | . | . |
| . | . | . | . | . | . | . | . | . | . | . | . | . | . | . | . | . | . | . | . | . | . | . | . | . | . |
| . | . | . | . | . | . | . | . | . | . | . | . | . | . | . | . | . | . | . | . | . | . | . | . | . | . |
| . | . | . | . | . | . | . | . | . | . | . | . | . | T | . | . | . | . | . | . | . | . | . | . | . | . |
| . | . | . | . | . | . | . | . | . | . | . | . | . | T | . | . | . | . | . | . | . | . | . | . | . | . |
| . | . | . | . | . | . | . | . | . | . | . | . | . | T | . | . | . | . | . | . | . | . | . | . | . | . |
| . | . | . | . | . | . | . | . | . | . | . | . | . | T | . | . | . | . | . | . | . | . | . | . | . | . |
| . | . | . | . | . | . | . | . | . | . | . | . | . | T | . | . | . | . | . | . | . | . | . | . | . | . |
| . | . | . | . | . | . | . | . | . | . | . | . | . | . | . | . | . | . | . | . | . | . | . | . | . | . |
| . | . | . | . | . | . | . | . | . | . | . | . | . | T | . | . | . | . | . | . | . | . | . | . | . | . |
| . | . | . | . | . | . | . | . | . | . | . | . | . | T | . | . | . | . | . | . | . | . | . | . | . | . |
| . | . | . | . | . | . | . | . | . | . | . | . | . | . | . | . | . | . | . | . | . | . | . | . | . | . |
| . | . | . | . | . | . | . | . | . | . | . | . | . | T | . | . | . | . | . | . | . | . | . | . | . | . |
| . | . | . | . | . | . | . | . | . | . | . | . | . | T | . | . | . | . | . | . | . | . | . | . | . | . |
| . | . | . | . | . | . | . | . | . | . | . | . | . | T | . | . | . | . | . | . | . | . | . | . | . | . |
| . | . | . | . | . | . | . | . | . | . | . | . | . | T | . | . | . | . | . | . | . | . | . | . | . | . |
| . | . | . | . | . | . | . | . | . | . | . | . | . | T | . | . | . | . | . | . | . | . | . | . | . | . |
| . | . | . | . | . | . | . | . | . | . | . | . | . | . | . | . | . | . | . | . | . | . | . | . | . | . |
| . | . | . | . | . | . | . | . | . | . | . | . | . | T | . | . | . | . | . | . | . | . | . | . | . | . |
| . | . | . | . | . | . | . | . | . | . | . | . | . | T | . | . | . | . | . | . | . | . | . | . | . | . |
| . | . | . | . | . | . | . | . | . | . | . | . | . | T | . | . | . | . | . | . | . | . | . | . | . | . |
| . | . | . | . | . | . | . | . | . | . | . | . | . | T | . | . | . | . | . | . | . | . | . | . | . | . |
| . | . | . | . | . | . | . | . | . | . | . | . | . | . | . | . | . | . | . | . | . | . | . | . | . | C |
| . | . | . | . | . | . | . | . | . | . | . | . | . | T | . | . | . | T | . | . | . | . | T | . | . | . |
| . | . | . | . | . | . | . | . | . | . | . | . | . | T | . | . | . | . | . | . | . | . | . | . | . | . |
| . | . | . | . | . | . | . | . | . | . | . | . | . | T | . | . | . | . | . | . | . | . | . | . | . | . |
| . | . | . | . | . | . | . | . | . | . | . | . | . | T | . | . | . | . | . | . | . | . | . | . | . | . |
| . | . | . | . | . | . | . | . | . | . | . | . | . | T | . | . | . | . | . | . | . | . | . | . | . | . |
| . | . | . | . | . | . | . | . | . | . | . | . | . | T | . | . | . | . | . | . | . | . | . | . | . | . |
| . | . | . | . | . | . | . | . | . | . | . | . | . | T | . | . | . | . | . | . | . | . | . | . | . | . |
| . | . | . | . | . | . | . | . | . | . | . | . | . | T | . | . | . | . | . | . | . | . | . | . | . | . |
| . | . | . | . | . | . | . | . | . | . | . | . | . | T | . | . | . | . | . | . | . | . | . | . | . | . |
| . | . | . | . | . | . | . | . | . | . | . | . | . | T | . | . | . | . | . | . | . | . | . | . | . | . |
| . | . | . | . | . | . | . | . | . | . | . | . | . | T | . | . | . | . | . | . | . | . | . | . | . | . |
| . | . | . | . | . | . | . | . | . | . | . | . | . | . | . | . | . | . | . | . | . | . | . | . | . | . |
| . | . | . | . | . | . | . | . | . | . | . | . | . | . | . | . | . | . | . | . | . | . | . | . | . | . |
| . | . | . | . | . | . | . | . | . | . | . | . | . | T | . | . | . | . | . | . | . | . | . | . | . | . |
| . | . | . | . | . | . | . | . | . | . | . | . | . | . | . | . | . | . | . | . | . | . | . | . | . | C |
| . | . | . | . | . | . | . | . | . | . | . | . | . | T | . | . | . | . | . | . | . | . | . | . | . | . |
| . | . | . | . | . | . | . | . | . | . | . | . | . | . | . | . | . | . | . | . | . | . | . | . | . | . |
| . | . | . | . | . | . | . | . | . | . | . | . | . | . | . | . | . | . | . | . | . | . | . | . | . | . |
| . | . | . | . | . | . | . | . | . | . | . | . | . | . | . | . | . | . | . | . | . | . | . | . | . | . |

| | | | | | | | | | | | | | | | | | | | | | | | | | |
|---|---|---|---|---|---|---|---|---|---|---|---|---|---|---|---|---|---|---|---|---|---|---|---|---|---|
| . | . | . | . | . | . | . | . | . | . | . | . | . | T | . | . | . | T | . | . | . | . | T | . | . | . |
| . | . | . | . | . | . | . | . | . | . | . | . | . | T | . | . | . | . | . | . | . | . | . | . | . | . |
| . | . | . | . | . | . | . | . | . | . | . | . | . | . | . | . | . | . | . | . | . | . | . | . | . | . |
| . | . | . | . | . | . | . | . | . | . | . | . | . | T | . | . | . | . | . | . | . | . | . | . | . | . |
| . | . | . | . | . | . | . | . | . | . | . | . | . | . | . | . | . | . | . | . | . | . | . | . | . | . |
| . | . | . | . | . | . | . | . | . | . | . | . | . | . | . | . | . | . | . | . | . | . | . | . | . | . |
| . | . | . | . | . | . | . | . | . | . | . | . | . | . | . | . | . | . | . | . | . | . | . | . | . | . |
| . | . | . | . | . | . | . | . | . | . | . | . | . | . | . | . | . | . | . | . | . | . | . | . | . | . |
| . | . | . | . | . | . | . | . | . | . | . | . | . | . | . | . | . | . | . | . | . | . | . | . | . | . |
| . | . | . | . | . | . | . | . | . | . | . | . | . | . | . | . | . | . | T | . | . | . | . | . | . | . |
| . | . | . | . | . | . | . | . | . | . | . | . | . | . | . | . | . | . | . | . | . | . | . | . | . | . |
| . | . | . | . | . | . | . | . | . | . | . | . | . | . | . | . | . | . | . | . | . | . | . | . | . | . |
| . | . | . | . | . | . | . | . | . | . | . | . | . | . | . | . | . | . | . | . | . | . | . | . | . | . |
| . | . | . | . | . | . | . | . | . | . | . | . | . | . | . | . | . | . | T | . | . | . | . | . | . | . |
| . | . | . | . | . | . | . | . | . | . | . | . | . | . | . | . | . | . | . | . | . | . | . | . | . | . |
| . | . | . | . | . | . | . | . | . | . | . | . | . | . | . | . | . | . | . | . | . | . | . | . | . | . |
| . | . | . | . | . | . | . | . | . | . | . | . | . | . | . | . | . | . | . | . | . | . | . | . | . | . |
| . | . | . | . | . | . | . | . | . | . | . | . | . | . | . | . | . | . | . | . | . | . | . | . | . | . |
| . | . | . | . | . | . | . | . | . | . | . | . | . | . | . | . | . | . | T | . | . | . | . | . | . | . |
| . | . | . | . | . | . | . | . | . | . | . | . | . | . | . | . | . | . | . | . | . | . | . | . | . | . |
| . | . | . | . | . | . | . | . | . | . | . | . | . | . | . | . | . | . | . | . | . | . | . | . | . | . |
| . | . | . | . | . | . | . | . | . | . | . | T | . | . | . | . | C | . | . | . | . | . | . | . | . | . |
| . | . | . | . | . | . | . | . | . | . | . | . | . | . | . | . | . | . | . | . | . | . | . | . | . | . |
| . | . | . | . | . | . | . | . | . | . | . | . | T | . | . | . | . | . | . | T | . | . | . | . | . | . |
| . | . | . | . | . | . | . | . | . | . | . | . | . | . | . | . | . | . | . | T | . | . | . | . | . | . |
| . | . | . | . | . | . | . | . | . | . | . | . | . | . | . | . | . | . | . | . | . | . | . | . | . | . |
| . | . | . | . | . | . | . | . | . | . | . | . | . | . | . | . | . | . | . | . | . | . | . | . | . | . |
| . | . | . | . | . | . | . | . | . | . | . | . | T | . | . | . | . | . | T | . | . | . | . | . | . | . |
| . | . | . | . | . | . | . | . | . | . | . | . | . | . | . | . | . | . | . | . | . | . | . | . | . | . |
| . | . | . | . | . | G | . | . | . | . | . | . | . | . | . | . | . | . | T | . | . | . | . | . | . | . |
| . | . | . | . | . | . | . | . | . | . | . | . | . | . | . | . | . | . | . | . | . | . | . | . | . | . |
| . | . | . | . | . | . | . | . | . | . | . | . | . | . | . | . | . | . | . | . | . | . | . | . | . | . |
| . | . | . | . | . | . | . | . | . | . | . | . | . | . | . | . | . | . | . | . | . | . | . | . | . | . |
| . | . | . | . | . | . | . | . | . | . | . | . | . | . | . | . | . | . | . | . | . | . | . | . | . | . |
| . | . | . | . | . | . | . | . | . | . | . | . | . | . | . | . | . | . | . | . | . | . | . | . | . | . |
| . | . | . | . | . | . | . | . | . | . | . | . | . | . | . | . | . | . | . | . | . | . | . | . | . | . |
| . | . | . | . | . | . | . | . | . | . | . | T | . | . | . | . | . | . | . | . | . | . | . | . | . | . |
| . | . | . | . | . | . | . | . | . | . | . | . | . | . | . | . | . | . | . | . | . | . | . | . | . | . |
| . | . | . | . | . | . | . | . | . | . | . | . | . | . | . | . | . | . | . | . | . | . | . | . | . | . |
| . | . | . | . | . | . | . | . | . | . | . | . | . | . | . | . | . | . | . | . | . | . | . | . | . | . |
| . | . | . | . | . | . | . | . | . | . | . | . | . | . | . | . | . | . | . | . | . | . | . | . | . | . |
| . | . | . | . | . | . | . | . | . | . | . | . | . | . | . | . | . | . | . | . | . | . | . | . | . | . |
| . | . | . | . | . | . | . | . | . | . | . | . | . | . | . | . | . | . | . | . | . | . | . | . | . | . |
| . | . | . | . | . | . | . | . | . | . | . | . | . | . | . | . | . | . | T | . | . | . | . | . | . | . |
| . | . | . | . | . | . | . | . | . | . | . | . | . | . | . | . | . | . | . | . | . | . | . | . | . | . |
| . | . | . | . | . | . | . | . | . | . | . | . | . | . | . | . | . | . | T | . | . | . | . | . | . | . |
| . | . | . | . | . | . | . | . | . | . | . | . | . | . | . | . | . | . | . | . | . | . | . | . | . | . |
| . | . | . | . | . | . | . | . | . | . | . | . | . | . | . | . | . | . | T | . | . | . | . | . | . | . |
| . | . | . | . | . | . | . | . | . | . | . | . | . | . | . | . | . | . | T | . | . | . | . | . | . | . |
| . | . | . | . | . | . | . | . | . | . | . | . | . | . | . | . | . | . | . | . | . | . | . | . | . | . |
| . | . | . | . | . | . | . | . | . | . | . | . | . | . | . | . | . | . | T | . | . | . | . | . | . | . |
| . | . | . | . | . | . | . | . | . | . | . | . | . | . | . | . | . | . | . | T | . | . | . | . | . | . |
| . | . | . | . | . | . | . | . | . | . | . | . | . | . | . | . | . | . | . | ? | . | . | . | . | . | . |
| . | . | . | . | . | . | . | . | . | . | . | . | . | . | . | . | . | . | . | . | . | . | . | . | . | . |
| . | . | . | . | . | . | . | . | . | . | . | . | . | . | . | . | . | . | . | . | . | . | . | . | . | . |
| . | . | . | . | . | . | . | . | . | . | . | . | . | . | . | . | . | . | . | . | . | . | . | . | . | . |
| . | . | . | . | . | . | . | . | . | . | . | . | . | . | . | . | . | . | T | . | . | . | . | . | . | . |
| . | . | . | . | . | . | . | . | . | . | . | . | . | . | . | . | . | . | . | . | . | . | . | . | . | . |
| . | . | . | . | . | . | . | . | . | . | . | . | . | . | . | . | . | . | . | . | . | . | . | . | . | . |
| . | . | . | . | . | . | . | . | . | . | . | . | . | . | . | . | . | . | . | . | . | . | . | . | . | . |
| . | . | . | . | . | . | . | . | . | . | . | . | . | . | . | . | . | . | T | . | . | . | . | . | . | . |

| | | | | | | | | | | | | | | | | | | | | | | | | | |
|---|---|---|---|---|---|---|---|---|---|---|---|---|---|---|---|---|---|---|---|---|---|---|---|---|---|
| . | . | . | . | . | . | . | . | . | . | . | . | . | . | . | . | . | . | . | . | . | . | . | . | . | . |
| . | . | . | . | . | . | . | . | . | . | . | . | . | . | . | . | . | . | T | . | . | . | . | . | . | . |
| . | . | . | . | . | . | . | . | . | . | . | . | . | . | . | . | . | . | . | . | . | . | . | . | . | . |
| . | . | . | . | . | . | . | . | . | . | . | . | . | . | . | . | . | . | . | . | . | . | . | . | . | . |
| . | . | . | . | . | . | . | . | . | . | . | . | . | . | . | . | . | . | . | . | . | . | . | . | . | . |
| . | . | . | . | . | . | . | . | . | . | . | . | . | . | . | . | . | . | . | . | . | . | . | . | . | . |
| . | . | . | . | . | . | . | . | . | . | . | . | . | . | . | . | . | . | . | . | . | . | . | . | . | . |
| . | . | . | . | . | . | . | . | . | . | . | . | . | T | . | . | . | . | . | . | . | . | . | . | . | . |
| . | . | . | . | . | . | . | . | . | . | . | . | . | T | . | . | . | . | . | . | . | . | . | . | . | . |
| . | . | . | . | . | . | . | . | . | . | . | . | . | . | . | . | . | . | . | . | . | . | . | . | . | . |
| T | . | . | . | . | . | . | . | . | . | . | . | . | . | T | . | . | . | . | . | . | . | . | . | . | . |
| T | . | . | . | . | . | . | . | . | . | . | . | . | . | T | . | . | . | . | . | . | . | . | . | . | . |
| . | . | . | . | . | . | . | . | . | . | . | . | . | . | . | . | . | . | T | . | . | . | . | . | . | . |
| . | . | . | . | . | . | . | . | . | . | . | . | . | . | . | . | . | . | . | . | . | . | . | . | . | . |
| . | . | . | . | . | . | . | . | . | . | . | . | . | T | . | . | . | . | . | . | . | . | . | . | . | . |
| . | . | . | . | . | . | . | . | . | . | . | . | . | . | . | . | . | . | . | . | . | . | . | . | . | . |
| . | . | . | . | . | . | . | . | . | . | . | . | . | . | . | . | . | . | . | . | . | . | . | . | . | . |
| . | . | . | . | . | . | . | . | . | . | . | . | . | T | . | . | . | . | . | . | . | . | . | . | . | . |
| . | . | . | . | . | . | . | . | . | . | . | . | . | . | . | . | . | . | . | . | . | . | . | . | . | . |
| . | . | . | . | . | . | . | . | . | . | . | . | . | T | . | . | . | . | . | . | . | . | . | . | . | . |
| . | . | . | . | . | . | . | . | . | . | . | . | . | T | . | . | . | . | . | . | . | . | . | . | . | . |
| . | . | . | . | . | . | . | . | T | . | . | . | . | T | . | . | . | . | . | . | . | . | . | . | . | . |
| . | . | . | . | . | . | . | . | . | . | . | . | . | . | . | . | . | . | T | . | . | . | . | . | . | . |
| . | . | . | . | . | . | . | . | . | . | . | . | . | . | . | . | . | . | . | . | . | . | . | . | . | . |
| . | . | . | . | . | . | . | . | . | . | . | . | . | . | . | . | . | . | T | . | . | . | . | . | . | . |
| . | . | . | . | . | . | . | . | . | . | . | . | . | . | . | . | . | . | . | . | . | . | . | . | . | . |
| . | . | . | . | . | . | . | . | . | . | . | . | . | . | . | . | . | . | . | . | . | . | . | . | . | . |
| . | . | . | . | . | . | . | . | . | . | . | . | . | . | . | . | . | . | T | . | . | . | . | . | . | . |
| . | . | . | . | . | . | . | . | . | . | . | . | . | T | . | . | . | . | . | . | . | . | . | . | . | . |
| . | . | . | . | . | . | . | . | . | . | . | . | . | . | . | . | . | . | T | . | . | . | . | . | . | . |
| . | . | . | . | . | . | . | . | . | . | . | . | . | T | . | . | . | . | . | . | . | . | . | . | . | . |
| . | . | . | . | . | . | . | . | . | . | . | . | . | . | . | . | . | . | . | . | . | . | . | . | . | . |
| . | . | . | . | . | . | . | . | . | . | . | . | . | . | . | . | . | . | . | . | . | . | . | . | . | . |
| c | c | g | c | a | a | a | c | g | c | c | c | c | c | g | c | t | g | c | g | g | g | g | c | g | t |
| c | c | g | c | a | a | a | c | g | c | c | c | c | c | g | c | t | g | c | g | g | g | g | c | g | t |
| c | c | g | c | a | a | a | c | g | c | c | c | c | c | g | c | t | g | c | g | g | g | g | c | g | t |
| c | c | g | c | a | a | a | c | g | c | c | c | c | c | g | c | t | g | c | g | g | g | g | c | g | t |
| c | c | g | c | a | a | a | c | g | c | c | c | c | c | g | c | t | g | c | g | g | g | g | c | g | t |
| c | c | g | c | a | a | a | c | g | c | c | c | c | c | g | c | t | g | c | g | g | g | g | c | g | t |
| c | c | g | c | a | a | a | c | g | c | c | c | c | c | g | c | t | g | c | g | g | g | g | c | g | t |
| c | c | g | c | a | a | a | c | g | c | c | c | c | c | g | c | t | g | c | g | g | g | g | c | g | t |
| c | c | g | c | a | a | a | c | g | c | c | c | c | c | g | c | t | g | c | g | g | g | g | c | g | t |
| c | c | g | t | g | a | a | c | g | c | c | c | c | c | g | c | t | g | c | g | g | g | g | c | g | t |
| c | c | g | c | a | a | a | c | g | c | c | c | c | c | g | c | t | g | c | g | g | g | g | c | g | t |
| c | t | g | c | a | a | a | c | g | c | c | c | c | c | g | c | t | g | c | g | g | g | g | c | g | t |
| c | t | g | c | a | a | a | c | g | c | c | c | c | c | g | c | t | g | c | g | g | t | g | c | g | t |
| c | c | g | c | a | a | a | c | g | c | c | c | c | c | g | c | t | g | c | g | g | g | g | c | g | t |
| c | c | g | c | a | a | a | c | g | c | c | c | c | c | g | c | t | g | c | g | g | g | g | c | g | t |
| c | c | g | c | a | a | a | c | g | c | c | c | c | c | g | c | t | g | t | g | g | g | g | c | g | t |
| c | c | g | c | a | a | a | c | g | c | c | c | c | c | g | c | t | g | c | g | g | g | g | c | g | t |
| c | c | g | c | a | a | a | c | g | c | c | c | c | c | g | c | t | g | c | g | g | g | g | c | g | t |
| c | c | g | c | a | a | a | c | g | c | c | c | c | c | g | c | t | g | c | g | g | g | g | c | g | t |
| c | c | g | c | a | a | g | c | g | c | c | c | c | c | g | c | t | g | c | g | g | g | g | c | g | t |
| c | c | g | c | a | a | a | c | g | c | c | c | c | c | g | c | t | g | c | g | g | g | g | c | g | t |
| c | c | g | c | a | a | a | c | g | c | c | c | c | c | g | c | t | g | t | g | g | g | g | c | g | t |
| c | c | g | c | a | a | a | c | g | c | c | c | c | c | g | c | t | g | c | g | g | g | g | c | g | t |
| c | c | g | c | a | a | a | c | g | c | c | c | c | c | g | c | t | g | c | g | g | g | g | c | g | t |
| . | . | . | . | . | . | . | . | . | . | . | . | . | . | . | . | ? | ? | ? | . | . | . | . | . | . | . |
| . | . | . | . | . | . | . | . | . | . | . | . | . | . | . | T | . | . | . | . | . | . | . | . | . | . |
| . | . | . | . | . | . | . | . | . | . | . | . | . | . | . | T | ? | ? | ? | . | . | . | . | . | . | . |
| . | . | . | . | . | . | . | . | . | . | . | . | . | . | . | . | ? | ? | ? | . | . | . | . | . | . | . |
| . | . | . | . | . | . | . | . | . | . | . | . | . | . | . | . | ? | . | . | . | . | . | . | . | . | . |

| | | | | | | | | | | | | | | | | | | | | | | | | | |
|---|---|---|---|---|---|---|---|---|---|---|---|---|---|---|---|---|---|---|---|---|---|---|---|---|---|
| . | . | . | . | . | . | . | . | . | . | . | . | . | . | . | . | . | . | ? | . | . | . | . | . | . | . |
| . | . | . | . | . | . | . | . | . | . | . | . | . | . | . | . | . | . | . | . | . | . | . | . | . | . |
| . | . | . | . | . | . | G | . | . | . | . | . | . | . | . | . | . | . | . | . | . | . | . | . | . | . |
| . | . | . | . | . | . | G | . | . | . | . | . | . | . | . | . | . | . | . | . | . | . | . | . | . | . |
| . | . | . | . | . | . | . | . | . | . | . | . | . | . | . | . | . | . | . | . | . | . | . | . | . | . |
| . | . | . | . | . | . | . | . | . | . | . | . | . | . | . | . | . | . | . | . | . | . | . | . | . | . |
| . | . | . | . | . | . | . | . | . | . | . | . | . | . | . | . | . | . | . | . | . | . | . | . | . | . |
| . | T | . | . | . | . | . | . | . | . | . | . | . | . | . | . | . | . | . | . | . | . | . | . | . | . |
| . | . | . | . | . | . | . | . | . | . | . | . | . | . | . | . | . | . | . | . | . | . | . | . | . | . |
| . | . | . | . | . | . | . | . | . | . | . | . | . | . | . | . | . | . | . | . | . | . | . | . | . | . |
| . | T | . | . | . | . | . | . | . | . | . | . | . | . | . | . | . | . | . | . | . | T | . | . | . | . |
| . | T | . | . | . | . | . | . | . | . | . | . | . | . | . | . | . | . | . | . | . | T | . | . | . | . |
| . | T | . | . | . | . | . | . | . | . | . | . | . | . | . | . | . | . | . | . | . | T | . | . | . | . |
| . | . | . | . | . | . | . | . | . | . | . | . | . | . | . | . | . | . | . | . | . | . | . | . | . | . |
| . | . | . | . | . | . | . | . | . | . | . | . | . | . | . | . | . | . | T | . | . | . | . | . | . | . |
| . | . | . | . | . | . | . | . | . | . | . | . | . | . | . | . | . | . | T | . | . | . | . | . | . | . |
| . | . | . | . | . | . | . | . | . | . | . | . | . | . | . | . | . | . | . | . | . | . | . | . | . | . |
| . | . | . | . | . | . | . | . | . | . | . | . | . | . | . | . | . | . | . | . | . | . | . | . | . | . |
| . | . | . | . | . | . | . | . | . | . | . | . | . | . | . | . | . | . | T | . | . | . | . | . | . | . |
| . | . | . | . | . | . | . | . | . | . | . | . | . | . | . | . | . | . | . | . | . | . | . | . | . | . |
| . | . | . | . | . | . | . | . | . | . | . | . | . | . | . | . | . | . | . | . | . | . | . | . | . | . |
| . | . | . | . | . | . | . | . | . | . | . | . | . | . | . | . | . | . | T | . | . | . | . | . | . | . |
| . | . | . | . | . | . | . | . | . | . | . | . | . | . | . | . | . | . | . | . | . | . | . | . | . | . |
| . | . | . | . | . | . | . | . | . | . | . | . | . | . | . | . | . | . | . | . | . | . | . | . | . | . |
| . | . | . | . | . | . | . | . | . | . | . | . | . | . | . | . | . | . | . | . | . | . | . | . | . | . |
| . | . | . | . | . | . | . | . | . | . | . | . | . | . | . | . | . | . | . | . | . | . | . | . | . | . |
| . | . | . | . | . | . | . | . | . | . | . | . | . | . | . | . | . | . | . | . | . | . | . | . | . | . |
| . | . | . | . | . | . | . | . | . | . | . | . | . | . | . | . | . | . | . | . | . | . | . | . | . | . |
| . | . | . | . | . | . | . | . | . | . | . | . | . | . | . | . | . | . | . | . | . | . | . | . | . | . |
| . | . | . | . | . | . | . | . | . | . | . | . | . | . | . | . | . | . | . | . | . | . | . | . | . | . |
| . | . | . | . | . | . | . | . | . | . | . | . | . | . | . | . | . | . | . | . | . | . | . | . | . | . |
| . | . | . | . | . | . | . | . | . | . | . | . | . | . | . | . | . | . | . | . | . | . | . | . | . | . |
| . | . | . | . | . | . | . | . | . | . | . | . | . | . | . | . | . | . | . | . | . | . | . | . | . | . |
| . | . | . | . | . | . | . | . | . | . | . | . | . | . | . | . | . | . | . | . | . | . | . | . | . | . |
| . | . | . | . | . | . | . | . | . | . | . | . | . | . | . | . | . | . | . | . | . | . | . | . | . | . |
| . | . | . | . | . | . | . | . | . | . | . | . | . | . | . | . | . | . | ? | . | . | . | . | . | . | . |
| . | . | . | . | . | . | . | . | . | . | . | . | . | . | . | . | . | . | T | . | . | . | . | . | . | . |
| . | . | . | . | . | . | . | . | . | . | . | T | . | . | . | . | . | . | . | . | . | . | . | . | . | . |
| . | . | . | . | . | . | . | . | . | . | . | . | . | . | . | . | . | . | . | . | . | . | . | . | . | . |
| . | . | . | . | . | . | . | . | . | . | . | . | . | . | . | . | . | . | . | . | . | . | . | . | . | . |
| . | . | . | . | . | . | . | . | . | . | . | . | . | . | . | . | . | . | . | . | . | . | . | . | . | . |
| . | . | . | . | . | . | . | . | . | . | . | . | . | T | . | . | . | . | . | . | . | . | . | . | . | . |
| . | . | . | . | . | . | . | . | . | . | . | . | . | . | . | . | . | . | . | . | . | . | . | . | . | . |
| . | . | . | . | . | . | . | . | . | . | . | . | . | . | . | . | . | . | . | . | . | . | . | . | . | . |
| . | . | . | . | . | . | . | . | . | . | . | . | . | . | . | . | . | . | . | . | . | . | . | . | . | . |
| . | . | . | . | . | . | . | . | . | . | . | . | . | T | . | . | . | . | . | . | . | . | . | . | . | . |
| . | . | . | . | . | . | . | . | . | . | . | . | . | . | . | . | . | . | . | . | . | . | . | . | . | . |
| . | . | . | . | . | . | . | . | . | . | . | . | . | T | . | . | . | . | . | . | . | . | . | . | . | . |
| . | . | . | . | . | . | . | . | . | . | . | . | . | . | . | . | . | . | . | . | . | . | . | . | . | . |
| . | . | . | . | . | . | . | . | . | . | . | . | . | . | . | . | . | . | . | . | . | . | . | . | . | . |
| . | . | . | . | . | . | . | . | . | . | . | . | . | . | . | . | . | . | . | . | . | . | . | . | . | . |
| . | . | . | . | . | . | . | . | . | . | . | . | . | T | . | . | . | . | . | . | . | . | . | . | . | . |
| . | . | . | . | . | . | . | . | . | . | . | . | . | T | . | . | . | . | . | . | . | . | . | . | . | . |
| . | . | . | . | . | . | . | . | . | . | . | . | . | . | . | . | . | . | . | . | . | . | . | . | . | . |
| . | . | . | . | . | . | . | . | . | . | . | . | . | . | . | . | . | . | . | . | . | . | . | . | . | . |
| . | . | . | . | . | . | . | . | . | . | . | . | . | . | . | . | . | . | . | . | . | . | . | . | . | . |
| . | . | . | . | . | . | . | . | . | . | . | . | . | . | . | . | . | . | . | . | . | . | . | . | . | . |
| . | . | . | . | . | . | . | . | . | . | . | . | . | . | . | . | . | . | . | . | . | . | . | . | . | . |
| . | . | . | . | . | . | . | . | . | . | . | . | . | . | . | . | . | . | . | . | . | . | . | . | . | . |
| . | . | . | . | . | . | . | . | . | . | . | . | . | . | . | . | . | . | . | . | . | . | . | . | . | . |
| . | . | . | . | . | . | . | . | . | . | . | . | . | . | . | . | . | . | . | . | . | . | . | . | . | . |

T
T
T
T
T
T
T
T
T
T
T

| | | | | | | | | | | | | | | | | | | | | | | | | | |
|---|---|---|---|---|---|---|---|---|---|---|---|---|---|---|---|---|---|---|---|---|---|---|---|---|---|
| . | . | . | . | . | . | . | . | . | . | . | . | . | . | . | . | . | . | . | . | . | . | . | . | . | . |
| . | . | . | . | . | . | . | . | . | . | . | . | . | . | . | . | . | . | . | . | . | . | . | . | . | . |
| . | . | . | . | . | . | . | T | . | . | . | . | . | . | . | . | . | . | . | . | . | . | . | . | . | . |
| . | . | . | . | . | . | . | T | . | . | . | . | . | . | . | . | . | . | . | . | . | . | . | . | . | . |
| . | . | T | . | . | . | . | . | . | . | . | . | . | . | . | . | . | . | . | . | . | . | . | . | . | . |
| . | . | . | . | . | . | . | . | . | . | . | . | . | . | . | . | . | . | . | . | . | . | . | . | . | . |
| . | . | . | . | . | . | . | . | . | . | . | . | . | . | . | . | . | . | . | . | . | . | . | . | . | . |
| . | . | . | . | . | . | . | . | . | . | . | . | . | . | . | . | . | . | . | . | . | . | . | . | . | . |
| . | . | . | . | . | . | . | . | . | . | . | . | . | . | . | . | . | . | . | . | . | . | . | . | . | . |
| . | . | T | . | . | . | . | . | . | . | . | . | . | . | . | . | . | . | . | . | . | . | . | . | . | . |
| . | . | . | . | . | . | . | . | . | . | . | . | . | . | . | . | . | . | . | . | . | . | . | . | . | . |
| . | . | . | . | . | . | . | . | . | . | . | . | . | . | . | . | . | . | . | . | . | . | . | . | . | . |
| . | . | . | . | . | . | . | . | . | . | . | . | . | . | . | . | . | . | . | . | . | . | . | . | . | . |
| . | . | . | . | . | . | . | . | . | . | . | . | . | . | . | . | . | . | . | . | . | . | . | . | . | . |
| . | . | . | . | . | . | . | . | . | . | . | . | . | . | . | . | . | . | . | . | . | . | . | . | . | . |
| . | . | . | . | . | . | . | . | . | . | . | . | . | . | . | . | . | . | . | . | . | . | . | . | . | . |
| . | . | . | . | . | . | . | . | . | . | . | . | . | . | . | . | . | . | . | . | . | . | . | . | . | . |
| . | . | . | . | . | . | . | . | . | . | . | . | . | . | . | . | . | . | . | . | . | . | . | . | . | . |
| . | . | . | . | . | . | . | . | . | . | . | . | . | . | . | . | . | . | . | . | . | . | . | . | . | . |
| . | . | . | . | . | . | . | . | . | . | . | . | . | . | . | . | . | . | . | . | . | . | . | . | . | . |
| . | . | . | . | . | . | . | . | . | . | . | . | . | . | . | . | . | . | . | . | . | . | . | . | . | . |
| . | . | . | . | . | . | . | . | . | . | . | . | . | . | . | . | . | . | T | . | . | . | . | . | . | . |
| . | . | . | . | . | . | . | . | . | . | . | . | . | . | . | . | . | . | T | . | . | . | . | . | . | . |
| . | . | . | . | . | . | . | . | . | . | . | . | . | . | . | . | . | . | . | . | . | . | . | . | . | . |
| . | . | . | . | . | . | . | . | . | . | . | . | . | . | . | . | . | . | . | . | . | . | . | . | . | . |
| . | . | . | . | . | . | . | . | . | . | . | . | . | . | . | . | . | . | . | . | . | . | . | . | . | . |
| . | . | . | . | . | . | . | . | . | . | . | . | . | . | . | . | . | . | . | . | . | . | . | . | . | . |
| . | . | . | . | . | . | . | . | . | . | . | . | . | . | . | . | . | . | T | . | . | . | . | . | . | . |
| . | . | . | . | . | . | . | . | . | . | . | . | . | . | . | . | . | . | T | . | . | . | . | . | . | . |
| . | . | . | . | . | G | . | . | . | . | . | . | . | . | . | . | . | . | T | . | . | . | . | . | . | . |
| . | . | . | . | . | . | . | . | . | . | . | . | . | . | . | . | . | . | . | . | . | . | . | . | . | . |
| . | . | . | . | . | . | . | . | . | . | . | . | . | . | . | . | . | . | . | . | . | . | . | . | . | . |
| . | . | . | . | . | . | . | . | . | . | . | . | . | . | . | . | . | . | T | . | . | . | . | . | . | . |
| . | . | . | . | . | . | . | . | . | . | . | . | . | . | . | . | . | . | T | . | . | . | . | . | . | . |
| . | . | . | . | . | . | . | . | . | . | . | . | . | . | . | . | . | . | . | . | . | . | . | . | . | . |
| . | . | . | . | . | . | . | . | . | . | . | . | . | . | . | . | . | . | T | . | . | . | . | . | . | . |
| . | . | . | . | . | . | . | . | . | . | . | . | . | . | . | . | . | . | . | . | . | . | . | . | . | . |
| . | . | . | . | . | . | . | . | . | . | . | . | . | . | . | . | . | . | . | . | . | . | . | . | . | . |
| . | . | . | . | . | . | . | . | . | . | . | . | . | T | . | . | . | . | . | . | . | . | . | . | . | . |
| . | . | . | . | . | . | . | . | . | . | . | . | . | T | . | . | . | . | . | . | . | . | . | . | . | . |
| . | . | . | . | . | . | . | . | . | . | . | . | . | T | . | . | . | . | . | . | . | . | . | . | . | . |
| . | . | . | . | . | . | . | . | . | . | . | . | . | T | . | . | . | . | . | . | . | . | . | . | . | . |
| . | . | . | . | . | . | . | . | . | . | . | . | . | T | . | . | . | . | . | . | . | . | . | . | . | . |
| . | . | . | . | . | . | . | . | . | . | . | . | . | T | . | . | . | . | . | . | . | . | . | . | . | . |
| . | . | . | . | . | . | . | . | . | . | . | . | . | T | . | . | . | . | . | . | . | . | . | . | . | . |
| . | . | . | . | . | . | . | . | . | . | . | . | . | T | . | . | . | . | . | . | . | . | . | . | . | . |
| . | . | . | . | . | . | . | . | . | . | . | . | . | T | . | . | . | . | . | . | . | . | . | . | . | . |
| . | . | . | . | . | . | . | . | . | . | . | . | . | T | . | . | . | . | . | . | . | . | . | . | . | . |
| . | . | . | . | . | . | . | . | . | . | . | . | . | . | . | . | . | . | . | . | . | . | . | . | . | . |
| . | . | . | . | . | . | . | . | . | . | . | . | . | T | . | . | . | . | . | . | . | . | . | . | . | . |
| . | . | . | . | . | . | . | . | . | . | . | . | . | T | . | . | . | . | . | . | . | . | . | . | . | . |
| . | . | . | . | . | . | . | . | . | . | . | . | . | T | . | . | . | . | . | . | . | . | . | . | . | . |
| . | . | . | . | . | . | . | . | . | . | . | . | . | T | . | . | . | . | . | . | . | . | . | . | . | . |
| . | . | . | . | . | . | . | . | . | . | . | . | . | T | . | . | . | . | . | . | . | . | . | . | . | . |
| . | . | . | . | . | . | . | . | . | . | . | . | . | T | . | . | . | . | . | . | . | . | . | . | . | . |
| . | . | . | . | . | . | . | . | . | . | . | . | . | T | . | . | . | . | . | . | . | . | . | . | . | . |
| . | . | . | . | . | . | . | . | . | . | . | . | . | . | . | . | . | . | . | . | . | . | . | . | . | . |
| . | . | . | . | . | . | . | . | . | . | . | . | . | . | . | . | . | . | . | . | . | . | . | . | . | . |
| . | . | . | . | . | . | . | . | . | . | . | . | . | . | . | . | . | . | . | . | . | . | . | . | . | . |
| . | . | . | . | . | . | . | . | . | . | . | . | . | . | . | . | . | . | . | . | . | . | . | . | . | . |
| . | . | . | . | . | . | . | . | . | . | . | . | . | . | . | . | . | . | . | . | . | . | . | . | . | . |
| . | . | . | . | . | . | . | ? | . | . | . | . | . | . | . | . | . | . | . | . | . | . | . | . | . | . |

| | | | | | | | | | | | | | | | | | | | | | | | | | |
|---|---|---|---|---|---|---|---|---|---|---|---|---|---|---|---|---|---|---|---|---|---|---|---|---|---|
| . | . | . | . | . | . | . | ? | . | . | . | . | . | T | . | . | . | . | . | . | . | . | . | . | . | . |
| . | . | . | . | . | . | . | . | . | . | . | . | . | T | . | . | . | . | . | . | . | . | . | . | . | . |
| . | . | . | . | . | . | . | . | . | . | . | . | . | T | . | . | . | . | . | . | . | . | . | . | . | . |
| . | . | . | . | . | . | . | . | . | . | . | . | . | T | . | . | . | . | . | . | . | . | . | . | . | . |
| . | . | . | . | . | . | . | . | . | . | . | . | . | . | . | . | . | . | . | T | . | . | . | . | . | . |
| . | . | . | . | . | . | . | . | . | . | . | . | . | . | . | . | . | . | . | . | . | . | . | . | . | . |
| . | . | . | . | . | . | . | . | . | . | . | . | . | . | . | . | . | . | . | . | . | . | . | T | . | . |
| . | . | . | . | . | . | . | . | . | . | . | . | . | . | . | . | . | . | . | . | . | . | . | . | . | . |
| . | . | . | . | . | . | . | . | . | . | . | . | . | . | . | . | . | . | . | . | . | . | . | . | . | . |
| . | . | . | . | . | . | . | . | . | . | . | . | . | . | . | . | . | . | . | . | . | . | . | . | . | . |
| . | . | . | . | . | . | . | . | . | . | . | . | . | . | . | . | t | g | c | . | . | . | . | . | . | . |
| . | . | . | . | . | . | . | . | . | . | . | . | . | . | . | . | . | g | c | . | . | . | . | . | . | t |
| . | . | . | . | . | . | . | . | . | . | . | . | . | . | . | . | . | . | . | . | . | . | . | . | . | . |
| . | . | . | . | . | . | . | . | . | . | . | . | . | . | . | . | . | . | c | . | . | . | . | . | . | . |
| . | . | . | . | . | . | . | . | . | . | . | . | . | . | . | . | . | . | . | . | . | . | . | . | . | . |
| . | . | . | . | . | . | . | . | . | . | . | . | . | . | . | . | t | g | c | . | . | . | . | . | . | . |
| . | . | . | . | . | . | . | . | . | . | . | . | . | . | . | . | t | . | . | . | . | . | . | . | . | . |
| . | . | . | . | . | . | . | . | . | . | . | . | . | . | . | . | . | . | . | . | . | . | . | T | . | . |
| . | . | . | . | . | . | . | . | . | . | . | . | . | . | . | . | t | g | c | . | . | . | . | . | . | . |
| . | . | . | . | . | . | . | . | . | . | . | . | . | . | . | . | t | g | c | . | . | . | . | . | . | . |
| . | . | . | . | . | . | . | . | . | . | . | . | . | . | . | . | t | g | c | . | . | . | . | . | . | . |
| . | . | . | . | . | . | . | . | . | . | . | . | . | . | . | . | t | g | c | . | . | . | . | . | . | . |
| . | . | . | . | . | . | . | . | . | . | . | . | . | . | . | . | t | g | c | . | . | . | . | . | . | . |
| . | . | . | . | . | . | . | . | . | . | . | . | . | . | . | . | t | g | . | . | . | . | . | . | . | . |
| . | . | . | . | . | . | . | . | . | . | . | . | . | . | . | . | . | . | T | . | . | . | . | . | . | . |
| . | . | . | . | . | . | . | . | . | . | . | . | . | . | . | . | C | . | ? | . | . | . | . | . | . | . |
| . | . | . | . | . | . | . | . | . | . | . | . | . | . | . | . | . | ? | ? | . | . | . | . | . | . | . |
| . | . | . | . | . | . | . | . | . | . | . | . | . | . | . | . | t | g | c | . | . | . | . | . | . | . |
| . | . | . | . | . | . | . | . | . | . | . | . | . | . | . | . | . | g | c | . | . | . | . | . | . | . |
| . | . | . | . | . | . | . | . | . | . | . | . | . | . | . | . | t | g | c | . | . | . | . | ? | . | . |
| . | . | . | . | . | . | . | . | . | . | . | . | . | . | . | . | . | . | c | . | . | . | . | . | . | . |
| . | . | . | . | . | . | . | . | . | . | . | . | . | . | . | . | . | . | . | . | . | . | . | . | . | . |
| . | . | . | . | . | . | . | . | . | . | . | . | . | . | . | . | . | . | . | . | . | . | . | . | . | . |
| . | . | . | . | . | . | . | . | . | . | . | . | . | T | . | . | . | . | . | . | . | . | . | . | . | . |
| . | . | . | . | . | . | . | . | . | . | . | . | . | . | . | . | . | . | . | . | . | . | . | . | . | . |
| . | . | . | . | . | . | . | . | . | . | . | . | . | . | . | . | . | . | . | . | . | . | . | . | . | . |
| . | . | . | . | . | . | . | . | . | . | . | . | . | . | . | . | . | . | . | . | . | . | . | . | . | . |
| . | . | . | . | . | . | . | . | . | . | . | . | . | T | T | . | . | . | . | . | . | . | . | . | . | . |
| . | . | . | . | . | . | . | . | . | . | . | . | . | T | . | . | . | . | . | . | . | . | . | . | . | . |
| . | . | . | . | . | . | . | . | . | . | . | . | . | . | . | . | . | . | . | . | . | . | . | . | A | . |
| . | . | . | . | . | . | . | . | . | . | . | . | . | . | . | . | . | . | . | . | . | . | . | . | A | . |
| . | . | . | . | . | . | . | . | . | . | . | . | . | T | . | . | . | . | . | . | . | . | . | . | . | . |
| . | . | . | . | . | . | . | . | . | . | . | . | . | T | . | . | . | . | . | . | . | . | . | . | . | . |
| . | . | . | . | . | . | . | . | . | . | . | . | . | . | . | . | . | . | . | . | . | . | . | . | . | . |
| . | . | . | . | . | . | . | . | . | . | . | . | . | T | . | . | . | . | . | . | . | . | . | . | . | . |
| . | . | . | . | . | . | . | . | . | . | . | . | . | . | . | . | . | . | . | . | . | . | . | . | . | . |
| . | . | . | . | . | . | . | . | . | . | . | . | . | . | . | . | . | . | . | . | . | . | . | . | . | . |
| . | . | . | . | . | . | . | . | . | . | . | . | . | . | . | . | . | . | . | . | . | . | . | . | . | . |
| . | . | . | . | . | . | . | . | . | . | . | . | . | T | . | . | . | . | . | . | . | . | . | . | . | . |
| . | . | . | . | . | . | . | . | . | . | . | . | . | . | . | . | . | . | . | . | . | . | . | . | . | . |
| . | . | . | . | . | . | . | . | . | . | . | . | . | . | . | . | . | . | . | . | . | . | . | . | . | . |
| . | . | . | . | . | . | . | . | . | . | . | . | . | . | . | . | . | . | . | . | . | . | . | . | . | . |
| . | . | . | . | . | . | . | . | . | . | . | . | . | T | . | . | . | . | . | . | . | . | . | . | . | . |
| . | . | . | . | . | . | . | . | . | . | . | . | . | T | . | . | . | . | . | . | . | . | . | . | . | . |
| . | . | . | . | . | . | . | . | . | . | T | . | . | T | . | . | . | . | . | . | . | . | . | . | . | . |
| . | . | . | . | . | . | . | . | . | . | T | . | . | T | . | . | . | . | . | . | . | . | . | . | . | . |
| . | . | . | . | . | . | . | . | . | . | . | . | . | . | . | . | . | . | . | . | . | . | . | . | . | . |
| . | . | . | . | . | . | . | . | . | . | . | . | . | T | . | . | . | . | . | . | . | . | . | . | . | . |
| . | . | . | . | . | . | . | . | . | . | . | . | . | . | . | . | . | . | . | . | . | . | . | . | . | . |
| . | . | . | . | . | . | . | . | . | . | . | . | . | T | . | . | . | . | . | . | . | . | . | . | . | . |

| 160 | 161 | 162 | 163 | 164 | 165 | 166 | 167 | 168 | 169 | 170 | 171 | 172 | 173 | 174 | 175 | 176 | 177 | 178 | 179 | 180 | 181 | 182 | 183 | 184 | 185 |
|---|---|---|---|---|---|---|---|---|---|---|---|---|---|---|---|---|---|---|---|---|---|---|---|---|---|
| ORF1ab | ORF1ab | ORF1ab | ORF1ab | ORF1ab | ORF1ab | ORF1ab | ORF1ab | S | S | S | S | S | S | S | S | S | S | S | S | S | S | S | S | S | S |
| 20087 | 20151 | 20355 | 20773 | 21004 | 21137 | 21550 | 21551 | 21575 | 21627 | 21628 | 21646 | 21724 | 21792 | 21795 | 21890 | 22093 | 22289 | 22343 | 22374 | 22444 | 22468 | 22530 | 22663 | 23120 | 23236 |
| G | T | A | G | G | A | A | A | C | C | T | C | G | A | G | T | G | G | G | A | C | G | C | C | G | G |
| . | . | . | . | . | . | . | . | . | . | . | . | . | . | . | . | . | . | . | . | . | . | . | . | . | . |
| . | . | . | . | . | . | . | . | . | . | . | . | . | . | . | . | . | . | . | . | . | . | . | . | . | . |
| . | . | . | . | . | . | . | . | . | . | . | . | . | . | . | . | . | . | . | . | . | . | . | . | . | . |
| . | . | . | . | . | . | . | . | . | . | . | . | . | . | . | . | . | . | . | . | . | . | . | . | . | . |
| . | . | . | . | . | . | . | . | . | . | . | . | . | . | . | . | . | . | . | . | . | . | . | . | . | . |
| . | . | . | . | . | . | . | . | . | . | . | . | . | . | . | . | . | . | . | . | . | . | . | . | . | . |
| . | . | . | . | . | . | . | . | . | . | . | . | . | . | . | . | . | . | . | . | . | . | . | . | . | . |
| . | . | . | . | . | . | . | . | . | . | . | . | . | . | . | . | . | . | . | . | . | . | . | . | . | . |
| . | . | . | . | . | . | . | . | . | . | . | . | . | . | . | . | . | . | . | . | . | . | . | . | . | . |
| . | . | . | . | . | . | . | . | . | . | . | . | . | . | . | . | . | . | . | . | . | . | . | . | . | . |
| . | . | . | . | . | . | . | . | . | . | . | . | . | . | . | . | . | . | . | . | . | . | . | . | . | . |
| . | . | . | . | . | . | . | . | . | . | . | . | . | . | . | . | . | . | . | . | . | . | . | . | . | . |
| . | . | . | . | . | . | . | . | . | . | . | . | . | . | . | . | . | . | . | . | . | . | . | . | . | . |
| . | . | . | . | . | . | . | . | . | . | . | . | . | . | . | . | . | . | . | . | . | . | . | . | . | . |
| . | . | . | . | . | . | . | . | . | . | . | . | . | . | . | . | . | . | . | . | . | . | . | . | . | . |
| . | . | . | . | . | . | . | . | . | . | . | . | . | . | . | . | . | . | . | . | . | . | . | . | . | . |
| . | . | . | . | . | . | . | . | . | . | . | . | . | . | . | . | . | . | . | . | . | . | . | . | . | . |
| . | . | . | . | . | . | . | . | . | . | . | . | . | . | . | . | . | . | . | . | . | . | . | . | . | . |
| . | . | . | . | . | . | . | . | . | . | . | . | . | . | . | . | . | . | . | . | . | . | . | . | . | . |
| . | . | . | . | . | . | . | . | . | . | . | . | . | . | . | . | . | . | . | . | . | . | . | . | . | . |
| . | . | . | . | . | . | . | . | . | . | . | . | . | . | . | . | . | . | . | . | . | . | . | . | . | . |
| . | . | . | . | . | . | . | . | . | . | . | . | . | . | . | . | . | . | . | . | . | . | . | . | . | . |
| . | . | . | . | . | . | . | . | . | . | . | . | . | . | . | . | . | . | . | . | . | . | . | . | . | . |
| . | . | . | . | . | . | . | . | . | . | . | . | . | . | . | . | . | . | . | . | . | . | . | . | . | . |
| . | . | . | . | . | . | . | . | . | . | . | . | . | . | . | . | . | . | . | . | . | . | . | . | . | . |
| . | . | . | . | . | . | . | . | . | . | . | . | . | . | . | . | . | . | . | . | . | . | . | . | . | . |
| . | . | . | . | . | . | . | . | . | . | . | . | . | . | . | . | . | . | . | . | . | . | . | . | . | . |
| . | . | . | . | . | . | . | . | . | . | . | . | . | . | . | . | . | . | . | . | . | . | . | . | . | . |
| . | . | . | . | . | . | . | . | . | . | . | . | . | . | . | . | . | . | . | . | . | . | . | . | . | . |
| . | . | . | . | . | . | . | . | . | . | . | . | . | . | . | . | . | . | . | . | . | . | . | . | . | . |
| . | . | G | . | . | . | . | . | . | . | . | . | . | . | . | . | . | . | . | . | . | . | . | . | . | . |
| . | . | G | . | . | . | . | . | . | . | . | . | . | . | . | . | . | . | . | . | . | . | . | . | . | . |
| . | . | . | . | . | . | . | . | . | . | . | . | . | . | . | . | . | . | . | . | . | . | . | . | . | . |
| . | . | . | . | . | . | . | . | . | . | . | . | . | . | . | . | . | . | . | G | . | . | . | . | . | . |
| . | . | . | . | . | . | . | . | . | . | . | . | . | . | . | . | . | . | . | G | . | . | . | . | . | . |
| . | . | . | . | . | . | . | . | . | . | . | . | . | . | . | . | . | . | . | . | . | . | . | . | . | . |
| . | . | ? | . | . | . | ? | ? | . | . | . | . | . | . | . | . | . | . | . | . | . | . | . | ? | . | . |
| . | . | . | . | . | . | . | . | . | . | . | . | . | . | . | . | . | . | . | . | . | . | . | . | . | . |
| . | . | . | . | . | . | . | . | . | . | . | . | . | . | . | . | . | . | . | . | . | . | . | . | . | . |
| . | . | . | . | . | . | . | . | . | . | . | . | . | . | . | . | . | . | . | . | . | . | . | . | . | . |
| . | . | . | . | . | . | . | . | . | . | . | . | . | . | . | . | . | . | . | . | . | . | . | . | . | . |
| . | . | ? | . | . | . | ? | ? | ? | ? | ? | ? | . | . | . | . | ? | . | . | . | . | . | . | ? | . | ? |
| . | . | . | . | . | . | . | . | . | . | . | . | . | . | . | . | . | . | . | . | . | . | . | . | . | . |
| . | . | . | . | . | . | . | . | . | . | . | . | . | . | . | . | . | . | . | . | . | . | . | . | . | . |
| . | . | . | . | . | . | . | . | . | . | . | . | . | . | . | . | . | . | . | . | . | . | . | . | . | . |
| . | . | . | . | . | . | . | . | . | . | . | . | . | . | . | . | . | . | . | . | . | . | . | . | . | . |
| . | . | . | . | . | . | . | . | . | . | . | . | . | . | . | . | . | . | . | . | . | . | . | . | . | . |
| . | . | . | . | . | . | . | . | . | . | . | . | . | . | . | . | . | . | . | . | . | . | . | . | . | . |
| . | . | . | . | . | . | . | . | . | . | . | . | . | . | . | . | . | . | . | . | . | . | . | . | . | . |
| . | . | . | . | . | . | . | . | . | . | . | . | . | . | . | . | . | . | . | . | . | . | . | . | . | . |
| . | . | . | . | . | . | . | . | . | . | . | . | . | . | . | . | . | . | . | . | . | . | . | . | . | . |
| . | . | . | . | . | . | . | . | . | . | . | . | . | . | . | . | . | . | . | . | . | . | . | . | . | . |
| . | . | . | . | . | . | . | . | . | . | . | . | . | . | . | . | . | . | . | . | . | . | . | . | . | . |
| . | . | . | . | . | . | . | . | . | . | . | . | . | . | . | . | . | . | . | . | . | . | . | . | . | . |
| . | . | . | . | . | . | . | . | . | . | . | . | . | . | . | . | . | . | . | . | . | . | . | . | . | . |

| | | | | | | | | | | | | | | | | | | | | | | | | | |
|---|---|---|---|---|---|---|---|---|---|---|---|---|---|---|---|---|---|---|---|---|---|---|---|---|---|
| . | . | . | . | . | . | . | . | . | . | . | . | . | . | . | . | . | . | . | . | . | . | . | . | . | . |
| . | . | . | . | . | . | . | . | . | . | . | . | . | . | . | . | . | . | . | . | . | . | . | . | . | . |
| . | . | . | . | . | . | . | . | . | . | . | . | . | . | . | . | . | . | . | . | . | . | . | . | . | . |
| . | . | . | . | . | . | . | . | . | . | . | . | . | . | . | . | . | . | . | . | . | . | . | . | . | . |
| . | . | . | . | . | . | . | . | . | . | . | . | . | . | . | . | . | . | . | . | T | . | . | . | . | . |
| . | . | . | . | . | . | . | . | . | . | . | . | . | . | . | . | . | . | . | . | . | . | . | . | . | . |
| . | . | . | . | . | . | . | . | . | . | . | . | . | . | . | . | . | . | . | . | . | . | . | . | . | . |
| . | . | . | . | . | . | . | . | . | . | . | . | . | . | . | . | . | . | . | . | . | . | . | . | . | . |
| . | . | . | . | . | . | . | . | . | . | . | . | . | . | . | . | . | . | . | . | . | . | . | . | . | . |
| . | . | . | . | . | . | . | . | . | . | . | . | . | . | . | . | g | . | . | . | . | . | . | . | . | . |
| . | . | . | . | . | . | . | . | . | . | . | . | . | . | . | . | g | . | . | . | . | . | . | . | . | . |
| . | . | . | . | . | . | . | . | . | . | . | . | . | . | . | . | . | . | . | . | T | . | . | . | . | . |
| . | . | . | . | . | . | . | . | . | . | . | . | . | . | . | . | . | . | . | . | T | . | . | . | . | . |
| . | . | . | . | . | . | . | . | . | . | . | . | . | . | . | . | . | . | . | . | T | . | . | . | . | . |
| . | . | a | . | . | . | . | . | . | . | . | . | . | . | . | . | . | . | . | . | T | . | . | . | . | . |
| . | . | . | . | . | . | . | . | . | . | . | . | . | . | . | . | . | . | . | . | T | . | . | . | . | . |
| . | . | . | . | . | . | . | . | . | . | . | . | . | . | . | . | . | . | . | . | . | . | . | . | . | . |
| . | . | . | . | . | . | . | . | . | . | . | . | . | . | . | . | . | . | . | . | . | . | . | . | . | . |
| . | . | . | . | . | . | a | a | c | c | t | c | . | . | . | . | . | . | . | a | c | g | . | . | . | . |
| . | . | . | . | . | . | . | . | . | . | . | . | . | ? | . | . | . | . | . | . | . | . | . | . | . | . |
| . | . | . | . | . | . | a | a | . | . | . | . | . | . | . | . | . | . | . | . | . | . | . | . | . | . |
| . | . | . | . | . | . | a | a | c | . | . | . | . | . | . | . | . | . | . | . | . | . | . | . | . | . |
| . | . | . | . | . | . | . | . | . | . | . | . | . | . | . | . | . | . | . | . | . | . | . | . | . | . |
| . | . | . | . | . | . | . | . | . | c | t | c | . | . | . | . | . | . | . | . | . | . | . | . | . | . |
| . | . | . | . | . | . | . | . | . | . | . | . | . | . | . | . | . | . | . | . | . | . | . | . | . | . |
| . | . | . | . | . | . | . | . | . | . | . | . | . | . | . | . | . | . | . | . | . | . | . | . | . | . |
| . | . | . | . | . | . | . | . | . | . | . | . | . | . | . | . | . | . | . | . | . | . | . | . | . | . |
| . | . | . | . | . | . | . | . | . | . | . | . | . | . | . | . | . | . | . | . | . | . | . | . | . | . |
| . | . | . | . | . | . | . | . | . | . | . | . | . | . | . | . | . | . | . | . | . | . | . | . | . | . |
| . | . | . | . | . | . | a | a | c | c | t | c | . | . | . | . | . | . | . | . | . | . | . | . | . | . |
| . | . | . | . | . | . | . | . | . | . | . | . | . | . | . | . | . | . | . | . | . | . | . | . | . | . |
| . | . | . | . | . | . | . | . | . | . | . | . | . | . | . | . | . | . | . | . | . | . | . | . | . | . |
| . | . | . | . | . | . | . | . | . | . | . | . | . | . | . | . | . | . | . | . | . | . | . | . | . | . |
| . | . | . | . | . | . | . | . | . | . | . | . | . | . | . | . | . | . | . | . | . | . | . | . | . | . |
| . | . | . | . | . | . | . | . | . | . | . | . | . | . | . | . | . | . | . | . | . | . | . | . | . | . |
| . | . | . | . | . | . | . | . | . | . | . | . | . | . | . | . | . | . | . | . | . | . | . | . | . | . |
| . | . | . | . | . | . | . | . | . | . | . | . | . | . | . | . | . | . | . | . | . | . | . | . | . | . |
| . | . | . | . | . | . | . | . | . | . | . | . | . | . | . | . | . | . | . | . | . | . | . | . | . | . |
| . | . | . | . | . | . | . | . | . | . | . | . | . | . | . | . | . | . | . | . | . | . | . | . | . | . |
| . | . | . | . | . | . | . | . | . | . | . | . | . | . | . | . | . | . | . | . | . | . | . | . | . | g |
| . | . | . | . | . | . | . | . | . | . | . | . | . | . | . | . | . | . | . | . | . | . | . | . | . | . |
| . | . | . | . | . | . | . | . | . | . | . | . | . | . | . | . | . | . | . | . | . | . | . | . | . | . |
| . | . | . | . | . | . | . | . | . | . | . | . | . | . | . | . | . | . | . | . | . | . | . | . | . | . |
| . | . | . | . | . | . | . | . | . | . | . | . | . | . | . | . | . | . | . | . | . | . | . | . | . | . |
| . | . | . | . | . | . | . | . | . | . | . | . | . | . | . | . | . | . | . | . | . | . | . | . | . | . |
| . | . | . | . | . | . | . | . | . | . | . | . | . | . | . | . | . | . | . | . | c | . | . | . | . | . |
| . | . | . | . | . | . | a | a | . | . | . | . | . | . | . | . | . | . | . | . | . | . | . | . | . | . |
| . | . | . | . | . | . | . | . | T | . | . | . | . | . | . | . | . | . | . | . | . | . | . | . | . | . |
| . | . | . | . | . | . | . | . | . | . | . | . | . | . | . | . | . | . | . | . | . | . | . | . | . | . |
| . | . | . | . | . | . | . | . | . | . | . | . | . | . | . | . | . | . | . | . | . | . | . | . | . | . |
| . | . | . | . | . | . | . | . | . | . | . | . | . | . | . | . | . | . | . | . | . | . | . | . | . | . |
| . | . | . | . | . | . | . | . | . | . | . | . | . | . | . | . | . | . | . | . | . | . | . | . | . | . |
| . | . | . | . | . | . | . | . | . | . | . | . | . | . | . | . | . | . | . | . | . | . | . | . | . | . |
| . | . | . | . | . | G | . | . | . | . | . | . | . | . | . | . | . | . | . | . | . | . | . | . | . | . |
| . | . | . | . | . | . | . | . | . | . | . | . | . | . | . | . | . | . | . | . | . | . | . | . | . | . |
| . | . | . | . | . | . | . | . | . | . | . | . | . | . | . | . | . | . | . | . | . | . | . | . | . | . |
| . | . | . | . | . | . | . | . | . | . | . | . | . | . | . | . | . | . | . | . | . | . | . | . | . | . |
| . | . | . | . | . | . | . | . | . | . | . | . | . | T | . | . | . | . | . | . | . | . | . | . | . | . |
| . | . | . | . | . | . | . | . | . | . | . | . | . | . | . | . | . | . | . | . | . | . | . | . | . | . |
| . | . | . | . | . | G | . | . | . | . | . | . | . | . | . | . | . | . | . | . | . | . | . | . | . | . |
| . | . | . | . | . | . | . | . | . | . | . | . | . | . | . | . | . | . | . | . | . | . | . | . | . | . |
| . | . | . | . | . | . | . | . | . | . | . | . | . | . | . | . | . | . | . | . | . | . | . | . | . | . |

| | | | | | | | | | | | | | | | | | | | | | | | | | |
|---|---|---|---|---|---|---|---|---|---|---|---|---|---|---|---|---|---|---|---|---|---|---|---|---|---|
| . | . | . | . | . | . | . | . | . | . | . | . | . | . | . | . | . | . | . | . | . | . | . | . | . | . |
| . | . | . | . | . | . | . | . | . | . | . | . | . | . | . | . | . | T | . | . | . | . | . | . | . | . |
| . | . | a | . | . | . | . | . | . | . | . | . | . | . | . | . | . | . | . | . | . | . | . | . | . | . |
| . | . | . | . | . | . | . | . | . | . | . | . | . | . | . | . | . | . | . | . | . | . | . | . | . | . |
| . | . | a | . | . | . | . | . | . | . | . | . | . | . | . | . | . | . | . | . | . | . | . | . | . | . |
| . | . | . | . | . | . | . | . | . | . | . | . | . | . | . | . | . | . | . | . | . | . | . | . | . | . |
| . | . | . | . | . | . | . | . | . | . | . | . | . | . | . | . | . | . | . | . | . | . | . | . | . | . |
| . | . | . | . | . | . | . | . | . | . | . | . | . | . | . | . | . | . | . | . | . | . | . | . | . | . |
| . | . | . | . | . | . | . | . | . | . | . | . | . | . | . | . | . | . | . | . | . | . | . | . | . | . |
| . | . | . | . | . | . | . | . | . | . | . | . | . | . | . | . | . | . | . | . | . | . | . | . | . | . |
| . | . | . | . | . | . | . | . | . | . | . | . | . | . | . | . | . | . | . | . | . | . | . | . | . | . |
| . | . | . | . | . | . | . | . | . | . | . | . | . | . | . | . | . | . | . | . | . | . | . | . | . | . |
| . | . | . | . | . | . | . | . | . | . | . | . | . | . | . | . | . | . | . | . | c | . | . | . | . | . |
| . | . | . | . | . | . | . | . | . | . | . | . | . | . | . | . | . | . | . | . | . | . | . | . | . | . |
| . | . | . | . | . | . | . | . | . | . | . | . | . | . | . | . | . | . | . | a | c | g | . | . | . | . |
| . | . | . | . | . | . | . | . | . | . | . | . | . | . | . | . | . | . | . | . | . | . | . | . | . | . |
| . | . | . | . | . | . | . | . | . | . | . | . | . | . | . | . | . | . | . | . | . | . | . | . | . | . |
| . | . | . | g | . | . | . | . | . | . | . | . | . | . | . | . | . | T | . | . | . | . | . | . | . | . |
| . | . | . | . | . | . | . | . | . | . | . | . | . | T | . | . | . | . | . | . | . | . | . | . | . | . |
| . | . | . | . | . | . | . | . | . | . | . | . | . | . | . | . | . | . | . | . | . | . | . | . | . | . |
| . | . | . | . | . | . | . | . | . | . | . | . | . | . | . | . | . | . | . | . | . | . | . | . | . | . |
| . | . | . | . | . | . | . | . | . | . | . | . | . | . | . | . | . | . | . | . | . | . | . | . | . | . |
| . | . | . | . | . | . | . | . | . | . | . | . | . | . | . | . | . | . | . | . | c | g | . | . | . | . |
| . | . | . | . | . | . | . | . | . | . | . | . | . | . | . | . | . | . | . | . | . | . | . | . | . | . |
| . | . | . | g | . | . | . | . | . | . | . | . | . | . | . | . | . | . | . | . | . | . | . | . | . | . |
| . | . | . | . | . | . | . | . | . | . | . | . | . | . | . | . | . | . | . | . | . | . | . | . | . | . |
| . | . | . | . | . | . | . | . | . | . | . | . | . | . | . | . | . | . | . | . | . | . | . | . | . | . |
| . | . | . | . | . | . | . | . | . | . | . | . | . | . | . | . | . | . | . | . | . | . | . | . | . | . |
| . | . | . | . | . | . | . | . | . | . | . | . | . | . | . | . | . | . | . | . | . | . | . | . | . | . |
| . | . | a | . | . | . | . | . | . | . | . | . | . | . | . | . | . | . | . | . | . | . | . | . | . | . |
| . | . | . | . | . | . | . | . | . | . | . | . | . | . | . | . | . | . | . | . | . | . | . | . | . | . |
| . | . | . | . | . | . | . | . | . | . | . | . | . | . | . | . | . | . | . | . | . | . | . | . | . | . |
| . | . | . | . | . | . | . | . | . | . | . | . | . | . | . | . | . | . | . | . | . | . | . | . | . | . |
| . | . | . | . | . | . | . | . | . | . | . | . | . | . | . | . | . | . | . | . | . | . | . | . | . | . |
| . | . | . | . | . | . | . | . | . | . | . | . | . | . | . | . | . | . | . | . | . | T | . | . | . | . |
| . | . | . | . | . | . | . | . | . | . | . | . | . | . | . | . | . | . | . | . | . | . | . | . | . | . |
| . | . | . | . | . | . | . | . | . | . | . | . | . | . | . | . | . | . | . | . | . | . | . | . | . | . |
| . | . | . | . | . | . | . | . | . | . | . | . | . | . | . | . | . | . | . | . | T | . | . | . | . | . |
| . | . | . | . | . | . | . | . | . | . | . | . | . | . | . | . | . | . | . | . | T | . | . | . | . | . |
| . | . | . | . | . | . | . | . | . | . | . | . | . | . | . | . | . | . | . | . | . | . | . | . | . | . |
| . | . | . | . | . | . | . | . | . | . | . | . | . | . | . | . | . | . | . | . | . | . | . | . | . | . |
| . | . | . | . | . | . | . | . | . | . | . | . | . | . | . | . | . | . | . | . | . | . | . | . | . | . |
| . | . | . | . | . | . | . | . | . | . | . | . | . | . | . | . | . | . | . | . | . | . | . | . | . | . |
| . | . | . | . | . | . | . | . | . | . | . | . | . | . | . | . | . | . | . | . | . | . | . | . | . | . |
| . | . | . | . | . | . | . | . | . | . | . | . | T | . | . | . | . | . | . | . | T | . | . | . | . | . |
| . | . | . | . | . | . | . | . | . | . | G | . | . | . | . | . | . | . | . | . | . | . | . | . | . | . |
| . | . | . | . | . | . | . | . | . | . | G | . | . | . | . | . | . | . | . | . | T | . | . | . | . | . |
| . | . | . | . | . | . | . | . | . | . | . | . | . | . | . | . | . | . | . | . | . | . | . | . | . | . |
| . | . | . | . | . | . | . | . | . | . | . | . | . | . | . | . | . | . | . | . | . | . | . | . | . | . |
| . | . | . | . | . | . | . | . | . | . | . | . | . | . | . | . | . | . | . | . | . | . | . | . | . | . |
| . | . | . | . | . | . | . | . | . | . | . | . | . | . | . | . | . | . | . | . | . | . | . | . | . | . |
| . | . | . | . | . | . | . | . | . | . | . | . | . | . | . | . | . | . | . | . | . | . | . | . | . | . |
| . | . | . | . | . | . | . | . | . | . | . | . | . | . | . | . | . | . | . | . | . | . | . | . | . | . |
| . | . | . | . | . | . | . | . | . | . | . | . | . | . | . | . | . | . | . | . | . | . | . | . | . | . |
| . | . | . | . | . | . | . | . | . | . | . | . | . | . | . | . | . | . | . | . | T | . | . | . | . | . |
| . | . | . | . | . | . | . | . | . | . | . | . | . | . | . | . | . | . | . | . | . | . | . | . | . | . |
| . | . | . | . | . | . | . | . | . | . | . | . | . | . | . | . | . | . | . | . | T | . | . | . | . | . |
| . | . | . | . | . | . | . | . | . | . | . | . | . | . | . | C | . | . | . | . | . | . | . | . | . | . |
| . | . | . | . | . | . | . | . | . | . | . | . | . | . | . | . | . | . | . | . | . | . | . | . | . | . |
| . | . | . | . | . | . | . | . | . | . | . | . | . | . | . | . | . | . | . | . | T | . | . | . | . | . |
| . | . | . | . | . | . | . | . | . | . | . | . | . | . | . | . | . | . | . | . | T | . | . | . | . | . |
| . | . | . | . | . | . | . | . | . | . | . | . | . | . | . | . | . | . | . | . | T | . | . | . | . | . |

| | | | | | | | | | | | | | | | | | | | | | | | | | |
|---|---|---|---|---|---|---|---|---|---|---|---|---|---|---|---|---|---|---|---|---|---|---|---|---|---|
| . | . | . | . | . | . | . | . | . | . | . | . | . | . | . | . | . | . | . | . | . | . | . | . | . | . |
| . | . | . | . | . | . | . | . | . | . | . | . | . | . | . | . | . | . | . | . | . | . | . | . | . | . |
| . | . | . | . | . | . | . | . | . | . | . | . | . | . | . | . | . | . | . | . | T | . | . | . | . | . |
| . | . | . | . | . | . | . | . | . | . | . | . | . | . | . | . | . | . | . | . | . | . | . | . | . | . |
| . | . | . | . | . | . | . | . | . | . | . | . | . | . | . | . | . | . | . | . | . | . | . | . | . | . |
| . | . | . | . | . | . | . | . | . | . | . | . | . | . | . | . | . | . | . | . | T | . | . | . | . | . |
| . | . | . | . | . | . | . | . | . | . | . | . | . | . | . | . | . | . | . | . | . | . | . | . | . | . |
| . | . | . | . | . | . | . | . | . | . | . | . | . | . | . | . | . | . | . | . | . | . | . | . | . | . |
| . | . | . | . | . | . | . | . | . | . | . | . | . | . | . | C | . | . | . | . | . | . | . | . | . | . |
| . | . | . | . | . | . | . | . | . | . | . | . | . | . | . | . | . | . | . | . | T | . | . | . | . | . |
| . | . | . | . | . | . | . | . | . | . | . | . | . | . | . | . | . | . | . | . | . | . | . | . | . | . |
| . | . | . | . | . | . | . | . | . | . | . | . | . | . | . | . | . | . | . | . | . | . | . | . | . | . |
| . | . | . | . | . | . | . | . | . | . | . | . | . | . | . | . | . | . | . | . | . | . | . | . | . | . |
| . | . | . | . | . | . | . | . | . | . | . | . | . | . | . | . | . | . | . | . | . | . | . | . | . | . |
| . | . | . | . | . | . | . | . | . | . | . | . | . | . | . | . | . | . | . | . | . | . | . | . | . | . |
| . | . | . | . | . | . | . | . | . | . | . | . | . | . | . | . | . | . | . | . | . | . | . | . | . | . |
| . | . | . | . | . | . | . | . | . | . | . | . | . | . | . | . | . | . | . | . | . | . | . | . | . | . |
| . | . | . | . | . | . | . | . | . | . | . | . | . | . | T | . | . | . | . | . | . | . | . | . | . | . |
| . | . | . | . | . | . | . | . | . | . | . | . | . | . | T | . | . | . | . | . | . | . | . | . | . | . |
| . | . | . | . | . | . | . | . | . | . | . | . | T | . | . | . | . | . | . | . | T | . | . | . | . | . |
| . | . | . | . | . | . | . | . | . | . | . | . | . | . | . | . | . | . | . | . | . | . | . | . | . | . |
| . | . | . | . | . | . | . | . | . | . | . | . | . | . | . | . | . | . | . | . | . | . | . | . | . | . |
| . | . | . | . | . | . | ? | ? | ? | . | . | . | . | . | . | . | . | . | . | . | . | . | . | . | . | . |
| . | . | . | . | . | . | . | . | . | . | . | . | T | . | . | . | . | . | . | . | T | . | . | . | . | . |
| . | C | . | . | . | . | ? | ? | ? | . | . | . | . | . | . | . | . | . | . | . | T | . | . | . | . | . |
| . | . | . | . | . | . | . | . | . | . | . | . | . | . | . | . | . | . | . | . | T | . | . | . | . | . |
| . | . | . | . | . | . | . | . | . | . | . | . | . | . | . | . | . | . | . | . | . | . | . | . | . | . |
| . | . | . | . | . | . | . | . | . | . | . | . | . | . | . | . | . | . | . | . | . | . | . | . | . | . |
| . | . | . | . | . | . | . | . | . | . | . | . | . | . | . | . | . | . | . | . | . | . | . | . | . | . |
| . | . | . | . | . | . | . | . | . | . | . | . | . | . | . | . | . | . | . | . | T | . | . | . | . | . |
| . | . | . | . | . | . | . | . | . | . | . | . | . | . | . | . | T | . | . | . | T | . | . | . | . | . |
| . | . | . | . | . | . | . | . | . | . | . | . | . | . | . | . | T | . | . | . | T | . | . | . | . | . |
| . | . | . | . | . | . | ? | ? | . | . | . | . | . | . | . | . | . | . | . | . | . | . | . | . | . | . |
| . | C | . | . | . | . | . | . | . | . | . | . | . | . | . | . | . | . | . | . | T | . | . | . | . | . |
| . | C | . | . | . | . | . | . | . | . | . | . | . | . | . | . | . | . | . | . | T | . | . | . | . | . |
| . | . | . | . | . | . | . | . | . | . | . | . | . | . | . | . | . | . | . | . | T | . | . | . | . | . |
| . | . | . | . | . | . | . | . | . | . | . | . | T | . | . | . | . | . | . | . | T | . | . | . | . | . |
| . | . | . | . | . | . | . | . | . | . | . | . | . | . | . | . | . | . | . | . | . | . | . | . | . | . |
| . | . | . | . | . | . | . | . | . | . | . | . | . | . | . | . | . | . | . | . | T | . | . | . | . | . |
| . | . | . | . | . | . | . | . | . | . | . | . | . | . | . | . | . | . | . | . | T | . | . | . | . | . |
| . | . | . | . | . | . | . | . | . | . | . | . | . | . | . | . | . | . | . | . | T | . | . | . | . | . |
| . | . | . | . | . | . | . | . | . | . | . | . | . | . | . | . | . | . | . | . | T | . | . | . | . | . |
| . | . | . | . | . | . | . | . | . | . | . | . | . | . | . | . | . | . | . | . | . | . | . | . | . | . |
| . | . | . | . | . | . | . | . | . | . | . | . | . | . | . | . | . | . | . | . | . | . | . | . | . | . |
| . | . | . | . | . | . | . | . | . | . | . | . | . | . | T | . | . | . | . | . | . | . | . | . | . | . |
| . | . | . | . | . | . | . | . | . | . | . | . | . | . | T | . | . | . | . | . | . | . | . | . | . | . |
| . | . | . | . | . | . | . | . | . | . | . | . | . | . | . | . | . | . | . | . | . | . | . | . | . | . |
| . | . | . | . | . | . | . | . | . | . | . | . | . | . | T | . | . | . | . | . | . | . | . | . | . | . |
| . | . | . | . | . | . | . | . | . | . | . | . | . | . | . | . | . | . | . | . | . | . | . | . | . | . |
| . | . | . | . | . | . | . | . | . | . | . | . | . | . | . | . | . | . | . | . | . | . | . | . | . | . |
| . | . | . | . | . | . | . | . | . | . | . | . | . | . | T | . | . | . | . | . | . | . | . | . | . | . |
| . | . | . | . | . | . | . | . | . | . | . | . | . | . | T | . | . | . | . | . | . | . | . | . | . | . |
| . | . | . | . | . | . | . | . | . | . | . | . | . | . | T | . | . | . | . | . | . | . | . | . | . | . |
| . | . | . | . | . | . | . | . | . | . | . | . | . | . | T | . | . | . | . | . | . | . | . | . | . | . |
| . | . | . | . | . | . | . | . | . | . | . | . | . | . | . | . | . | . | . | . | . | . | . | . | . | . |
| . | . | . | . | . | . | . | . | . | . | . | . | . | . | . | . | . | . | . | . | . | . | . | . | . | . |
| . | . | . | . | . | . | . | . | . | . | . | . | . | . | . | . | . | . | . | . | T | . | . | . | . | . |
| . | . | . | . | . | . | . | . | . | . | . | . | . | . | . | . | . | . | . | . | . | . | . | . | . | . |
| . | . | . | . | . | . | . | . | . | . | . | . | . | . | . | . | . | . | . | . | T | . | . | . | . | . |
| . | . | . | . | . | . | . | . | . | . | . | . | . | . | . | . | . | . | . | . | . | . | . | . | . | . |
| . | . | . | . | . | . | . | . | . | . | . | . | . | . | . | . | . | . | . | . | . | . | . | . | . | . |
| . | . | . | . | . | . | . | . | . | . | . | . | . | . | . | . | . | . | . | . | . | . | . | . | . | . |

| | | | | | | | | | | | | | | | | | | | | | | | | | |
|---|---|---|---|---|---|---|---|---|---|---|---|---|---|---|---|---|---|---|---|---|---|---|---|---|---|
| . | . | . | . | . | . | . | . | . | . | . | . | . | . | . | . | . | . | . | . | . | . | . | . | . | . |
| . | . | . | . | . | . | . | . | . | . | . | . | . | . | . | . | . | . | . | . | T | . | . | . | . | . |
| . | . | . | . | . | . | . | . | . | . | . | . | . | . | . | . | . | . | . | . | . | . | . | . | . | . |
| . | . | . | . | . | . | . | . | . | . | . | . | . | . | . | . | . | . | . | . | . | . | . | . | . | . |
| . | . | . | . | . | . | . | . | . | . | . | . | . | T | . | . | . | . | . | . | . | . | . | . | . | . |
| . | . | . | . | . | . | . | . | . | . | . | . | . | . | . | . | . | . | . | . | . | . | . | . | . | . |
| . | . | . | . | . | . | . | . | . | . | . | . | . | . | . | . | . | . | . | . | . | . | . | . | . | . |
| . | . | . | . | . | . | . | . | . | . | . | . | . | . | . | . | . | . | . | . | . | . | . | . | . | . |
| . | . | . | . | . | . | . | . | . | . | . | . | . | . | . | . | . | . | . | . | . | . | . | . | . | . |
| . | . | . | . | . | . | . | . | . | . | . | . | . | . | . | . | . | . | . | . | . | . | . | . | . | . |
| . | . | . | . | . | . | . | . | . | . | . | . | . | . | . | . | . | . | . | . | . | . | . | . | . | . |
| . | . | . | . | . | . | . | . | . | . | . | . | . | . | . | . | . | . | . | . | . | . | . | . | . | . |
| . | . | . | . | . | . | . | . | . | . | . | . | . | . | . | . | . | . | . | . | . | . | . | . | . | . |
| . | . | . | . | . | . | . | . | . | . | . | . | . | . | . | . | . | . | . | . | . | . | . | . | . | . |
| . | . | . | . | . | . | . | . | . | . | . | . | . | T | . | . | . | . | . | . | . | . | . | . | . | . |
| . | . | . | . | . | . | . | . | . | . | . | . | . | . | . | . | . | . | . | . | . | . | . | . | . | . |
| . | . | . | . | . | . | . | . | . | . | . | . | . | . | . | . | . | . | . | . | . | . | . | . | . | . |
| . | . | . | . | . | . | . | . | . | . | . | . | . | . | . | . | . | . | . | . | . | . | . | . | . | . |
| . | . | . | . | . | . | . | . | . | . | . | . | . | . | . | . | . | . | . | . | . | . | . | . | . | . |
| . | . | . | . | . | . | . | . | . | . | . | . | . | T | . | . | . | . | . | . | . | . | . | . | . | . |
| . | . | . | . | . | . | . | . | . | . | . | . | . | . | . | . | . | . | . | . | . | . | . | . | . | . |
| . | . | . | . | . | . | . | . | . | . | . | . | . | . | . | . | . | . | . | . | . | . | . | . | . | . |
| . | . | . | . | . | . | . | . | . | . | . | . | . | . | . | . | . | . | . | . | . | . | . | . | . | . |
| . | . | . | . | . | . | . | . | . | . | . | . | . | . | . | . | . | . | . | . | . | . | . | A | . | . |
| . | . | . | . | . | . | . | . | . | . | . | . | . | . | . | . | . | . | . | . | . | . | . | . | . | . |
| . | . | . | . | . | . | . | . | . | . | . | . | . | . | . | . | . | . | . | . | . | . | . | . | . | . |
| . | . | . | . | . | . | . | . | . | . | . | . | . | . | . | . | . | . | . | . | . | . | . | . | . | . |
| . | . | . | . | . | . | . | . | . | . | . | . | . | . | . | . | . | . | . | . | . | . | . | . | . | . |
| . | . | . | . | . | . | . | . | . | . | . | . | . | . | . | . | . | . | . | . | . | . | . | . | . | . |
| . | . | . | . | . | . | . | . | . | . | . | . | . | . | . | . | . | . | . | . | . | . | . | . | . | . |
| . | . | . | . | . | . | . | . | . | . | . | . | . | . | . | . | . | . | . | . | . | . | . | . | . | . |
| . | . | . | . | . | . | . | . | . | . | . | . | . | . | . | . | . | . | . | . | . | . | . | . | . | . |
| . | . | . | . | . | . | . | . | . | . | . | . | . | . | . | . | . | . | . | . | . | . | . | . | . | . |
| . | . | . | . | . | . | . | . | . | . | . | . | . | . | . | . | . | . | . | . | . | . | . | . | . | . |
| . | . | . | . | . | . | . | . | . | . | . | . | . | . | . | . | . | . | . | . | . | . | . | . | . | . |
| . | . | . | . | . | . | . | . | . | . | . | . | . | . | . | . | . | . | . | . | . | . | . | . | . | . |
| . | . | . | . | . | . | . | . | . | . | . | . | . | . | . | . | . | . | . | . | . | . | . | A | . | . |
| . | . | . | . | . | . | . | . | . | . | . | . | . | . | . | . | . | . | . | . | . | . | . | . | . | . |
| . | . | . | . | . | . | . | . | . | . | . | . | . | . | . | . | . | . | . | . | . | . | . | . | . | . |
| . | . | . | . | . | . | . | . | . | . | . | . | . | T | . | . | . | . | . | . | . | . | . | . | . | . |
| . | . | . | . | . | . | . | . | . | . | . | . | . | . | . | . | . | . | . | . | . | . | . | . | . | . |
| . | . | . | . | . | . | . | . | . | . | . | . | . | . | . | . | . | . | . | . | . | . | . | . | . | . |
| . | . | . | . | . | . | . | . | . | . | . | . | . | . | . | . | . | . | . | . | . | . | . | . | . | . |
| . | . | . | . | . | . | . | . | . | . | . | . | . | . | . | . | . | . | . | . | . | . | . | . | . | . |
| . | . | . | . | . | . | . | . | . | . | . | . | . | . | . | . | . | . | . | . | . | . | . | . | . | . |
| . | . | . | . | . | . | . | . | . | . | . | . | . | T | . | . | . | . | . | . | . | . | . | . | . | . |
| . | . | . | . | . | . | . | . | . | . | . | . | . | . | . | . | . | . | . | . | . | . | . | . | . | . |
| . | . | . | . | . | . | . | . | . | . | . | . | . | T | . | . | . | . | . | . | . | . | . | . | . | . |
| . | . | . | . | . | . | . | . | . | . | . | . | . | . | . | . | . | . | . | . | . | . | . | . | . | . |
| . | . | . | . | . | . | . | . | . | . | . | . | . | . | . | . | . | . | . | . | . | . | . | . | . | . |
| . | . | . | . | . | . | . | . | . | . | . | . | . | . | . | . | . | . | . | . | . | . | . | . | . | . |
| . | . | . | . | . | . | . | . | . | . | . | . | . | . | . | . | . | . | . | . | . | . | . | . | . | . |
| . | . | . | . | . | . | . | . | . | . | . | . | . | . | . | . | . | . | . | . | . | . | . | . | . | . |
| . | . | . | . | . | . | . | . | . | . | . | . | . | . | . | . | . | . | . | . | . | . | . | . | . | . |
| . | . | . | . | . | . | . | . | . | . | . | . | . | . | . | . | . | . | . | . | . | . | . | . | . | . |
| . | . | . | . | . | . | . | . | . | . | . | . | . | . | . | . | . | . | . | . | . | . | . | . | . | . |
| . | . | . | . | . | . | . | . | . | . | . | . | . | . | . | . | . | . | . | . | . | . | . | . | . | . |
| . | . | . | . | . | . | . | . | . | . | . | . | . | . | . | . | . | . | . | . | . | . | . | . | . | . |
| . | . | . | . | . | . | . | . | . | . | . | . | . | . | . | . | . | . | . | . | . | . | . | . | . | . |
| . | . | . | . | . | . | . | . | . | . | . | . | . | . | . | . | . | . | . | . | . | T | . | . | . | . |
| . | . | . | . | . | . | . | . | . | . | . | . | . | . | . | . | . | . | . | . | . | . | . | . | . | . |
| . | . | . | . | . | . | . | . | . | . | . | . | . | . | . | . | . | . | . | . | . | . | . | . | . | . |

| | | | | | | | | | | | | | | | | | | | | | | | | | |
|---|---|---|---|---|---|---|---|---|---|---|---|---|---|---|---|---|---|---|---|---|---|---|---|---|---|
| . | . | . | . | . | . | . | . | . | . | . | . | . | . | . | . | . | . | . | . | . | . | . | . | . | . |
| . | . | . | . | . | . | . | . | . | . | . | . | . | . | . | . | . | . | . | . | . | . | . | . | . | . |
| . | . | . | . | . | . | . | . | . | . | . | . | . | . | . | . | . | . | . | . | . | . | . | . | . | . |
| . | . | . | . | . | . | M | W | . | . | . | . | . | T | . | . | . | . | . | . | . | . | . | . | . | . |
| . | . | . | . | . | . | . | . | . | . | . | . | . | . | . | . | . | . | . | . | . | . | . | . | . | . |
| . | . | . | . | . | . | . | . | . | . | . | . | . | . | . | . | . | . | . | . | . | . | . | . | . | . |
| . | . | . | . | . | . | . | . | . | . | . | . | . | . | . | . | . | . | . | . | . | . | . | . | . | . |
| . | . | . | . | . | . | . | . | . | . | . | . | . | . | . | . | . | . | . | . | . | . | . | . | . | . |
| . | . | . | . | . | . | . | . | . | . | . | . | . | . | T | . | . | . | . | . | T | . | . | . | . | . |
| ? | . | . | . | . | . | ? | ? | . | . | . | . | . | . | . | . | . | . | . | . | . | . | . | . | . | . |
| . | . | . | . | . | . | . | . | . | . | . | . | . | . | . | . | . | . | . | . | . | . | . | . | . | . |
| . | . | . | . | . | . | . | . | . | . | . | . | . | . | . | . | . | . | . | . | . | . | . | . | . | . |
| . | . | . | . | . | . | . | . | . | . | . | . | . | . | . | . | . | . | . | . | . | . | . | . | . | . |
| . | . | . | . | . | . | . | . | . | . | . | . | . | . | . | . | . | . | . | . | . | . | . | . | . | . |
| . | . | . | . | . | . | . | . | . | . | . | . | . | . | . | . | . | . | . | . | . | . | . | . | . | . |
| . | . | . | . | . | . | . | . | . | . | . | . | . | . | . | . | . | . | . | . | . | T | . | . | . | . |
| . | . | . | . | . | . | . | . | . | . | . | . | . | . | . | . | . | . | . | . | . | T | . | . | . | . |
| . | . | . | . | . | . | . | . | . | . | . | . | . | . | . | . | . | . | . | . | T | . | . | . | . | . |
| . | . | . | . | . | . | . | . | T | . | . | . | . | . | . | . | . | . | . | . | . | . | . | . | . | . |
| . | . | . | . | . | . | . | . | . | . | . | . | . | . | . | . | . | . | . | . | T | . | . | . | . | . |
| . | . | . | . | . | . | . | . | . | . | . | . | . | . | . | . | . | . | . | . | T | . | . | . | . | . |
| . | . | . | . | . | . | . | . | . | . | . | . | T | . | . | . | . | . | . | . | T | . | . | . | . | . |
| . | . | . | . | . | . | . | . | . | . | . | . | . | . | . | . | . | . | . | . | . | . | . | . | . | . |
| . | . | . | . | . | . | . | . | . | . | . | . | . | . | . | . | . | . | . | . | . | . | . | . | . | . |
| . | . | . | . | . | . | . | . | . | . | . | . | . | . | . | . | . | . | . | . | . | . | . | . | . | . |
| . | . | . | . | . | . | . | . | . | . | . | . | . | . | . | . | . | . | . | . | . | . | . | . | . | . |
| . | . | . | . | . | . | . | . | . | . | . | . | . | . | . | . | . | . | . | . | . | . | . | . | . | . |
| . | . | . | . | . | . | . | . | . | . | . | . | . | . | . | . | . | . | . | . | . | . | . | . | . | . |
| . | . | . | . | . | . | . | . | . | . | . | . | . | . | . | . | . | . | . | . | . | . | . | . | . | . |
| . | . | . | . | . | . | . | . | . | . | . | . | . | . | . | . | . | . | . | . | . | . | . | . | . | . |
| . | . | . | . | . | . | . | . | . | . | . | . | . | . | . | . | . | . | . | . | T | . | . | . | . | . |
| . | . | . | . | . | . | . | . | . | . | . | . | . | . | . | . | . | . | . | . | . | . | . | . | . | . |
| . | . | . | . | . | . | . | . | . | . | . | . | . | . | . | . | . | . | . | . | . | . | . | . | . | . |
| g | t | a | g | g | a | a | a | c | c | t | c | g | a | g | t | g | g | g | a | c | g | c | c | g | g |
| g | t | a | g | g | a | a | a | c | c | t | c | g | a | g | t | g | g | g | a | c | g | c | c | g | g |
| g | t | a | t | g | a | a | a | c | t | t | c | g | a | g | t | g | g | g | a | c | g | c | c | g | g |
| g | t | a | t | g | a | a | a | c | t | t | c | g | a | g | t | g | g | g | a | c | g | c | c | g | g |
| g | t | a | g | g | a | a | a | c | c | t | c | g | a | g | t | g | g | g | a | c | g | c | c | g | g |
| g | t | a | g | g | a | a | a | c | c | t | c | g | a | g | t | g | g | g | a | c | g | c | c | g | g |
| g | t | a | g | g | a | a | a | c | c | t | c | g | a | g | t | g | g | g | a | c | g | c | c | g | g |
| g | t | a | g | g | a | a | a | c | c | t | c | g | a | g | t | g | g | g | a | c | g | c | c | g | g |
| g | t | a | g | g | a | a | a | c | c | t | c | g | a | g | t | g | g | g | a | c | g | c | c | g | g |
| g | t | a | g | g | a | a | a | c | c | t | c | g | a | g | t | g | g | g | a | c | g | c | c | g | g |
| g | t | a | g | g | a | a | a | c | c | t | c | g | a | g | t | g | g | g | a | c | t | c | c | g | g |
| g | t | a | g | g | a | a | a | c | c | t | c | g | a | g | t | g | g | g | a | c | t | c | c | g | g |
| g | t | a | g | g | a | a | a | c | c | t | c | g | a | g | t | g | g | g | a | c | t | c | c | g | g |
| g | t | a | g | g | a | a | a | c | c | t | c | g | a | g | t | g | g | g | a | c | g | c | c | g | g |
| g | t | a | g | g | a | a | a | c | c | t | c | g | a | g | t | g | g | g | a | c | g | c | c | g | g |
| g | t | a | g | g | a | a | a | c | c | t | c | g | a | g | t | g | g | g | a | c | g | c | c | g | g |
| g | t | a | g | g | a | a | a | c | c | t | c | g | a | g | t | g | g | g | a | c | g | c | c | g | t |
| g | t | a | g | g | a | a | a | c | c | t | c | g | a | g | t | g | g | g | a | c | g | c | c | g | g |
| g | t | a | g | g | a | a | a | c | c | t | c | g | a | g | t | g | g | g | a | c | g | c | c | g | g |
| g | t | a | g | g | a | a | a | c | c | t | c | g | a | g | t | g | g | g | a | c | g | c | c | g | g |
| g | t | a | g | g | a | a | a | c | c | t | c | g | a | g | t | g | g | g | a | c | g | c | c | g | g |
| g | t | a | g | g | a | a | a | c | c | t | c | g | a | g | t | g | g | g | a | c | g | c | c | g | g |
| g | t | a | g | g | g | a | a | c | c | t | c | g | a | g | t | g | g | g | a | c | g | c | c | g | g |
| g | t | a | g | g | a | a | a | c | c | t | c | g | a | g | t | g | g | g | a | c | g | c | c | g | g |
| . | . | ? | . | . | . | . | . | . | . | . | . | . | . | . | . | . | . | . | . | . | . | . | . | . | . |
| . | . | . | . | . | . | . | . | . | . | . | . | . | ? | ? | ? | . | . | . | . | . | . | . | . | . | . |
| . | . | . | . | . | . | . | . | . | . | . | . | . | . | . | . | . | . | . | . | . | . | . | . | . | . |
| . | . | . | . | . | . | . | . | . | . | . | . | . | . | . | . | . | . | . | . | . | . | . | . | . | . |
| . | . | . | . | . | . | . | . | . | . | . | . | . | . | . | . | . | . | . | . | . | . | . | . | . | . |

| | | | | | | | | | | | | | | | | | | | | | | | | | |
|---|---|---|---|---|---|---|---|---|---|---|---|---|---|---|---|---|---|---|---|---|---|---|---|---|---|
| . | . | . | . | . | . | . | . | . | . | . | . | . | . | . | . | . | . | . | . | . | . | . | . | . | . |
| . | . | . | . | . | . | . | T | . | . | . | . | . | . | . | . | . | . | . | . | . | . | . | . | . | . |
| C | . | . | . | . | . | . | . | . | . | . | T | . | . | . | . | . | . | . | . | . | . | . | . | . | . |
| . | . | . | . | . | . | . | . | . | . | . | . | . | . | . | . | . | . | . | . | . | . | . | . | . | . |
| . | . | . | . | . | . | . | . | . | . | . | . | . | . | . | . | . | . | . | . | . | . | . | . | . | . |
| . | . | . | . | . | . | . | . | . | . | . | . | . | . | . | . | . | . | . | . | . | . | . | . | . | . |
| . | . | . | . | . | . | . | . | . | . | . | . | . | . | . | . | . | . | . | . | . | . | . | . | . | . |
| . | . | . | . | . | . | . | . | . | . | . | . | . | . | . | . | . | . | . | . | . | T | . | . | . | . |
| . | . | . | . | . | . | . | . | . | . | . | . | . | . | . | . | . | . | . | . | . | . | . | . | . | . |
| C | . | . | . | . | . | . | . | . | . | . | . | . | . | . | . | . | . | . | . | . | . | . | . | . | . |
| . | . | . | . | . | . | . | . | . | . | . | . | . | . | . | . | . | . | . | . | . | T | . | . | . | . |
| . | . | . | . | . | . | . | . | . | . | . | . | . | . | . | . | . | . | . | . | . | T | . | . | . | . |
| . | . | . | . | . | . | . | . | . | . | . | . | . | . | . | . | . | . | . | . | . | T | . | . | . | . |
| . | . | . | . | . | . | . | . | . | . | . | . | . | . | . | . | . | . | . | . | . | . | . | . | . | . |
| . | . | . | . | . | . | . | . | . | . | . | . | . | . | . | . | . | . | . | . | . | . | . | . | . | . |
| . | . | . | . | . | . | . | . | . | . | . | . | . | . | . | . | . | . | . | . | . | . | . | . | . | . |
| . | . | . | . | . | . | . | . | . | . | . | . | . | . | . | . | . | . | . | . | . | . | . | . | . | . |
| . | . | . | . | . | . | . | . | . | . | . | . | . | . | . | . | . | . | . | . | . | . | . | . | . | . |
| . | . | . | . | . | . | . | . | . | . | . | . | . | . | . | . | . | . | . | . | . | . | . | . | . | . |
| . | . | . | . | . | . | . | . | . | . | . | . | . | . | . | . | . | . | . | . | . | . | . | . | . | . |
| . | . | . | . | . | . | . | . | . | . | . | . | . | . | . | . | . | . | . | . | . | . | . | . | . | . |
| . | . | . | . | . | . | . | . | . | . | . | . | . | . | . | . | . | . | . | . | . | . | . | . | . | . |
| . | . | . | . | . | . | . | . | . | . | . | . | . | . | . | . | . | . | . | . | . | . | . | . | . | . |
| . | . | . | . | . | . | . | . | . | . | . | . | . | . | . | . | . | . | . | . | . | . | . | . | . | . |
| . | . | . | . | . | . | . | . | . | . | . | . | . | . | . | . | . | . | . | . | . | . | . | . | . | . |
| . | . | . | . | . | . | . | . | . | . | . | . | . | . | . | . | . | . | . | . | . | . | . | . | . | . |
| . | . | . | . | . | . | . | . | . | . | . | . | . | . | . | . | . | . | . | . | . | . | . | . | . | . |
| . | . | . | . | . | . | . | . | . | . | . | . | . | . | . | . | . | . | . | . | . | . | . | . | . | . |
| . | . | . | . | . | . | . | . | . | . | . | . | . | . | . | . | . | . | . | . | . | . | . | . | . | . |
| . | . | . | . | . | . | . | . | . | . | . | . | . | . | . | . | . | . | . | . | . | . | . | . | . | . |
| . | . | . | . | . | . | . | . | . | . | . | . | . | . | . | . | . | . | . | . | . | . | . | . | . | . |
| . | . | . | . | . | . | . | . | . | . | . | . | . | . | . | . | . | . | . | . | . | . | . | . | . | . |
| . | . | . | . | . | . | . | . | . | . | . | . | . | . | . | . | . | . | . | . | . | . | . | . | . | . |
| . | . | . | . | . | . | . | . | . | . | . | . | . | . | . | . | . | . | . | . | . | . | . | . | . | . |
| . | . | . | . | . | . | . | . | . | . | . | . | . | . | . | . | . | . | . | . | . | . | . | . | . | . |
| . | ? | . | . | . | . | . | . | . | . | . | . | . | . | . | . | . | . | . | . | . | . | . | . | . | . |
| . | . | . | . | . | . | . | . | . | . | . | . | . | . | . | . | . | . | . | . | . | . | . | . | . | . |
| . | . | . | . | . | . | . | . | . | . | . | . | . | . | . | . | . | . | . | . | . | . | . | . | . | . |
| . | . | . | . | . | . | . | . | . | . | . | . | . | . | . | . | . | . | . | . | . | . | . | . | . | . |
| . | . | . | . | . | . | ? | ? | . | . | . | . | . | . | . | . | . | . | . | . | . | T | . | . | . | . |
| . | . | . | . | . | . | . | . | . | . | . | . | . | . | . | . | . | . | . | . | . | T | . | . | . | . |
| . | . | . | . | . | . | . | . | . | . | . | . | . | . | . | . | . | . | . | . | . | . | . | . | . | . |
| . | . | . | . | . | . | . | . | . | . | . | . | . | . | . | . | . | . | . | . | . | . | . | . | . | . |
| . | . | . | . | . | . | . | . | . | . | . | . | . | . | . | . | . | . | . | . | . | . | . | . | . | . |
| . | . | . | . | . | . | . | . | . | . | . | . | . | . | . | . | . | . | . | . | . | . | . | . | . | . |
| . | . | . | . | . | . | . | . | . | . | . | . | . | . | . | . | . | . | . | . | . | . | . | . | . | . |
| . | . | . | . | . | . | . | . | . | . | . | . | . | . | . | . | . | . | . | . | . | . | . | . | . | . |
| . | . | . | . | . | . | . | . | . | . | . | . | . | . | . | . | . | . | . | . | . | . | . | . | . | . |
| . | . | . | . | . | . | . | . | . | . | . | . | . | . | . | . | . | . | . | . | . | . | . | . | . | . |
| . | . | . | . | . | . | . | . | . | . | . | . | . | . | . | . | . | . | . | . | . | . | . | . | . | . |
| . | . | . | . | . | . | . | . | . | . | . | . | . | . | . | . | . | . | . | . | . | . | . | . | . | . |
| . | . | . | . | . | . | . | . | . | . | . | . | . | . | . | . | . | . | . | . | . | . | . | . | . | . |
| . | . | . | . | . | . | . | . | . | . | . | . | . | . | . | . | . | . | . | . | . | . | . | . | . | . |
| . | . | . | . | . | . | . | . | . | . | . | . | . | . | . | . | . | . | . | . | . | . | . | . | . | . |
| . | . | . | . | . | . | . | . | . | . | . | . | . | . | . | . | . | . | . | . | . | . | . | . | . | . |
| . | . | . | . | . | . | . | . | . | . | . | . | . | . | . | . | . | . | . | . | . | . | . | . | . | . |
| . | . | . | . | . | . | . | . | . | . | . | . | . | . | . | . | . | . | . | . | . | T | . | . | . | . |
| . | . | . | . | . | . | . | . | . | . | . | . | . | . | . | . | . | . | . | . | . | . | . | . | . | . |
| . | . | . | . | . | . | . | . | . | . | . | . | . | . | . | . | . | . | . | . | . | . | . | . | . | . |
| . | . | . | . | . | . | . | . | . | . | . | . | . | . | . | . | . | . | A | . | . | . | . | . | . | . |
| . | . | . | . | . | . | . | . | . | . | . | . | . | . | . | . | . | . | . | . | . | T | . | . | . | . |

| | | | | | | | | | | | | | | | | | | | | | | | | | |
|---|---|---|---|---|---|---|---|---|---|---|---|---|---|---|---|---|---|---|---|---|---|---|---|---|---|
| . | . | . | . | . | . | . | . | . | . | . | . | . | . | . | . | . | . | . | . | . | . | . | . | . | . |
| . | . | . | . | . | . | . | . | . | . | . | . | . | . | . | . | . | . | . | . | . | . | . | . | . | . |
| . | . | . | . | . | . | . | . | . | . | . | . | . | . | . | . | . | . | . | . | . | . | . | . | . | . |
| . | . | . | . | . | . | . | . | . | . | . | . | . | . | . | . | . | . | . | . | . | . | . | . | . | . |
| . | . | . | . | . | . | . | . | . | . | . | . | . | . | . | . | . | . | . | . | . | . | T | . | . | . |
| . | . | . | . | . | . | . | . | . | . | . | . | . | . | . | . | . | . | . | . | . | . | . | . | . | . |
| . | . | . | . | . | . | . | . | . | . | . | . | . | . | . | . | . | . | . | . | . | T | . | . | . | . |
| . | . | . | . | . | . | . | . | . | . | . | . | . | . | . | . | . | . | . | . | . | . | . | . | . | . |
| . | . | . | . | . | . | . | . | . | . | . | . | . | . | . | . | . | . | . | . | . | . | . | . | . | . |
| . | . | . | . | . | . | . | . | . | . | . | . | . | . | . | . | . | . | . | . | . | . | . | . | . | . |
| . | . | . | . | . | . | . | . | . | . | . | . | . | . | . | . | . | . | . | . | . | . | T | . | . | . |
| . | . | . | . | . | . | . | . | . | . | . | . | . | . | . | . | . | . | . | . | . | . | . | . | . | . |
| . | . | . | . | . | . | . | . | . | . | . | . | . | . | . | . | . | . | . | . | . | . | T | . | . | . |
| . | . | . | . | . | . | . | . | . | . | . | . | . | . | . | . | . | . | A | . | . | . | . | . | . | . |
| . | . | . | . | . | . | . | . | . | . | . | . | . | . | . | . | . | . | A | . | . | . | . | . | . | . |
| . | . | . | . | . | . | . | . | . | . | . | . | . | . | . | . | . | . | . | . | . | T | . | . | . | . |
| . | . | . | . | . | . | . | . | . | . | . | . | . | . | . | . | . | . | . | . | . | . | . | . | . | . |
| . | . | . | . | . | . | . | . | . | . | . | . | . | . | . | . | . | . | . | . | . | . | . | . | . | . |
| . | . | . | . | . | . | . | . | . | . | . | . | . | . | . | . | . | . | . | . | . | . | . | . | . | . |
| . | . | . | . | . | . | . | . | . | . | . | . | . | . | . | . | . | . | . | . | . | . | . | . | . | . |
| . | . | . | . | . | . | . | . | . | . | . | . | . | . | . | . | . | . | . | . | . | . | . | . | . | . |
| . | . | . | . | . | . | . | . | . | . | . | . | . | . | . | . | . | . | . | . | . | . | . | . | . | . |
| . | . | . | . | . | . | . | . | . | . | . | . | . | . | . | . | . | . | . | . | . | . | . | . | . | . |
| . | . | . | . | . | . | . | . | . | . | . | . | . | . | . | . | . | . | . | . | . | T | . | . | . | . |
| . | . | . | . | . | . | . | . | . | . | . | . | . | . | . | . | . | . | . | . | . | T | . | . | . | . |
| . | . | . | . | . | . | . | . | . | . | . | . | . | . | . | . | . | . | . | . | . | T | . | . | . | . |
| . | . | . | . | . | . | . | . | . | . | . | T | . | . | . | . | . | . | . | . | . | . | . | . | . | . |
| . | . | . | . | . | . | . | . | . | . | . | . | . | . | . | . | . | . | . | . | . | . | . | . | . | . |
| . | . | . | . | . | . | . | . | . | . | . | . | . | . | . | . | . | . | . | . | . | . | . | . | . | . |
| . | . | . | . | . | . | . | . | . | . | . | . | . | . | . | . | . | . | . | . | . | . | . | . | . | . |
| . | . | . | . | . | . | . | . | . | . | . | . | . | . | . | . | . | . | . | . | . | T | . | . | . | . |
| . | . | . | . | . | . | . | . | . | . | . | . | . | . | . | . | . | . | . | . | . | T | . | . | . | . |
| . | . | . | . | . | . | . | . | . | . | . | . | . | . | . | . | . | . | . | . | . | T | . | . | . | . |
| . | . | . | . | . | . | . | . | . | . | . | . | . | . | . | . | . | . | . | . | . | T | . | . | . | . |
| . | . | . | . | . | . | . | . | . | . | . | T | . | . | . | . | . | . | . | . | . | . | . | . | . | . |
| . | . | . | . | . | . | . | . | . | . | . | . | . | . | . | . | . | . | . | . | . | T | . | . | . | . |
| . | . | . | . | . | . | . | . | . | . | . | . | . | . | . | . | . | . | . | . | . | T | . | . | . | . |
| . | . | . | . | . | . | . | . | . | . | . | . | . | . | . | . | . | . | . | . | . | . | . | . | . | . |
| . | . | . | . | . | . | . | . | . | . | . | . | . | . | . | . | . | . | . | . | . | T | . | . | . | . |
| . | . | . | . | . | . | . | . | . | . | . | . | . | . | . | . | . | . | . | . | . | . | . | . | . | . |
| . | . | . | . | . | . | . | . | . | . | . | . | . | . | . | . | . | . | . | . | . | . | . | . | . | . |
| . | . | . | . | . | . | . | . | . | . | . | . | . | . | . | . | . | . | . | . | . | . | . | . | . | . |
| . | . | . | . | . | . | . | . | . | . | . | . | . | . | . | . | . | . | . | . | . | . | . | . | . | . |
| . | . | . | . | . | . | . | . | . | . | . | . | . | . | . | . | . | . | . | . | . | . | . | . | . | . |
| . | . | . | . | . | . | . | . | . | . | . | . | . | . | . | . | . | . | . | . | . | . | . | . | . | . |
| . | . | . | . | . | . | . | . | . | . | . | . | . | . | . | . | . | . | . | . | . | . | . | . | . | . |
| . | . | . | . | . | . | . | . | . | . | . | . | . | . | . | . | . | . | . | . | . | . | . | . | . | . |
| . | . | . | . | . | . | . | . | . | . | . | . | . | . | . | . | . | . | . | . | . | . | . | . | . | . |
| . | . | . | . | . | . | . | . | . | . | . | . | . | . | . | . | . | . | . | . | . | . | . | . | . | . |
| . | . | . | . | . | . | . | . | . | . | . | . | . | . | . | . | . | . | . | . | . | . | . | . | . | . |
| . | . | . | . | . | . | . | . | . | . | . | . | . | . | . | . | . | . | . | . | . | . | . | . | . | . |
| . | . | . | . | . | . | . | . | . | . | . | . | . | . | . | . | . | . | . | . | . | . | . | . | . | . |
| . | . | . | . | . | . | . | . | . | . | . | . | . | . | . | . | . | . | . | . | . | . | . | . | . | . |
| . | . | . | . | . | . | . | . | . | . | . | . | . | . | . | . | . | . | . | . | . | . | . | . | . | . |
| . | . | . | . | . | . | . | . | . | . | . | . | . | . | . | . | . | . | . | . | . | . | . | . | . | . |
| . | . | . | . | . | . | . | . | . | . | . | . | . | . | . | . | . | . | . | . | . | . | . | . | . | T |
| . | . | . | . | . | . | . | . | . | . | . | . | . | . | . | . | . | . | . | . | . | . | . | . | . | . |
| . | . | . | . | . | . | . | . | . | . | . | . | . | . | . | . | . | . | . | . | . | . | . | . | . | . |
| . | . | . | . | . | . | . | . | . | . | . | . | . | . | . | . | . | . | . | . | . | . | . | . | . | . |
| . | . | . | . | . | . | . | . | . | . | . | . | . | . | . | . | . | . | . | . | . | . | . | . | . | . |
| . | . | . | . | . | . | . | . | . | . | . | . | . | . | . | . | . | . | . | . | . | . | . | . | . | . |

| | | | | | | | | | | | | | | | | | | | | | | | | | |
|---|---|---|---|---|---|---|---|---|---|---|---|---|---|---|---|---|---|---|---|---|---|---|---|---|---|
| . | . | . | . | . | . | . | . | . | . | . | . | . | . | . | . | . | . | . | . | . | . | . | . | . | . |
| . | . | . | . | . | . | . | . | . | . | . | . | . | . | . | . | . | . | . | . | . | . | . | . | . | . |
| . | . | . | . | . | . | . | . | . | . | . | . | . | . | . | . | . | . | . | . | . | . | . | . | . | . |
| . | . | . | . | . | . | . | . | . | . | . | . | . | . | . | . | . | . | . | . | . | . | . | . | . | . |
| . | . | . | . | . | . | C | . | . | . | . | . | . | . | . | . | . | . | . | . | . | . | . | . | T | . |
| . | . | . | . | . | . | C | T | . | . | . | . | . | . | . | . | . | . | . | . | . | . | . | . | T | . |
| . | . | . | . | . | . | . | . | . | . | . | . | . | . | . | . | . | . | . | . | . | . | . | . | T | . |
| . | . | . | . | . | . | C | T | . | . | . | . | . | . | . | . | . | . | . | . | . | . | . | . | T | . |
| . | . | . | . | . | . | . | . | . | . | . | . | . | . | . | . | . | . | . | . | . | . | . | . | T | . |
| . | . | . | . | . | . | C | T | . | . | . | . | . | . | . | . | . | . | . | . | . | . | . | . | T | . |
| . | . | . | . | . | . | C | . | . | . | . | . | . | . | . | . | . | . | . | . | . | . | . | . | . | . |
| . | . | . | . | . | . | . | T | . | . | . | . | . | . | . | . | . | . | . | . | . | . | . | . | . | . |
| . | . | . | . | . | . | C | T | . | . | . | . | . | . | . | . | . | . | . | . | . | . | . | . | . | . |
| . | . | . | . | . | . | . | . | . | . | . | . | . | . | . | . | . | . | . | . | . | . | . | . | . | . |
| . | . | . | . | . | . | . | . | . | . | . | . | . | . | . | . | . | . | . | . | . | . | . | . | . | . |
| . | . | . | . | . | . | . | . | . | . | . | . | . | . | . | . | . | . | . | . | . | . | . | . | . | . |
| . | . | . | . | . | . | . | . | . | . | . | . | . | . | . | . | . | . | . | . | . | . | . | . | . | . |
| . | . | . | . | . | . | . | . | . | . | . | . | . | . | . | . | . | . | . | . | . | . | . | . | . | . |
| . | . | . | . | . | . | C | T | . | . | . | . | . | . | . | . | . | . | . | . | . | . | . | . | . | . |
| . | . | . | . | . | . | . | . | . | . | . | . | . | . | . | . | . | . | . | . | . | . | . | . | . | . |
| . | . | . | . | . | . | . | . | . | . | . | . | . | . | . | . | . | . | . | . | . | . | . | . | . | . |
| . | . | . | . | . | . | . | . | . | . | . | . | . | . | . | . | . | . | . | . | . | . | . | . | . | . |
| . | . | . | . | . | . | . | . | . | . | . | . | . | . | . | . | . | . | . | . | . | . | . | . | . | . |
| . | . | . | . | . | . | . | . | . | . | . | . | . | . | . | . | . | . | . | . | . | . | . | . | . | . |
| . | . | . | . | . | . | . | . | . | . | . | . | . | . | . | . | . | . | . | . | . | . | . | . | . | . |
| . | . | . | . | . | . | . | . | . | . | . | . | . | . | . | . | . | . | . | . | . | . | . | . | . | . |
| . | . | . | . | . | . | . | . | . | . | . | . | . | . | . | . | . | . | . | . | . | . | . | . | . | . |
| . | . | . | . | . | G | . | . | . | . | . | . | . | . | . | . | . | . | . | . | . | . | . | . | . | . |
| . | . | . | . | . | . | . | . | . | . | . | . | . | . | . | . | . | . | . | . | . | . | . | . | . | . |
| . | . | . | . | . | . | . | . | . | . | . | . | . | . | . | . | . | . | . | . | . | . | . | . | . | . |
| . | . | . | . | . | . | . | . | . | . | . | . | . | . | . | . | . | . | . | . | . | . | . | . | . | . |
| . | . | . | . | . | . | M | W | . | . | . | . | . | T | . | . | . | . | . | . | . | . | . | . | . | . |
| . | . | . | . | . | . | M | W | . | . | . | . | . | . | . | . | . | . | . | . | . | . | . | . | . | . |
| . | . | . | . | . | . | M | W | . | . | . | . | . | . | . | . | . | . | . | . | . | . | . | . | . | . |
| . | . | . | . | . | . | . | . | . | . | . | . | . | . | . | . | . | . | . | . | . | . | . | . | . | . |
| . | . | . | . | . | . | . | . | . | . | . | . | . | . | . | . | . | . | . | . | . | . | . | . | . | . |
| . | . | . | . | . | . | M | W | . | . | . | . | . | . | . | . | . | . | . | . | . | . | . | . | . | . |
| . | . | . | . | . | . | . | . | . | . | . | . | . | . | . | . | . | . | . | . | . | . | . | . | . | . |
| . | . | . | . | . | . | . | . | . | . | . | . | . | . | . | . | . | . | . | . | T | . | . | . | . | . |
| . | . | . | . | . | . | . | . | . | . | . | . | . | . | . | . | . | . | . | . | . | . | . | . | . | . |
| . | . | . | . | . | . | . | . | . | . | . | . | . | . | . | . | . | . | . | . | T | . | . | . | . | . |
| . | . | . | . | . | . | . | . | . | . | . | . | . | . | . | . | . | . | . | . | . | . | . | . | . | . |
| . | . | . | . | . | . | . | . | . | . | . | . | . | . | . | . | . | . | . | . | T | . | . | . | . | . |
| . | . | . | . | . | . | . | . | . | . | . | . | . | . | . | . | . | . | . | . | T | . | . | . | . | . |
| . | . | . | . | . | . | . | . | . | . | . | . | . | . | . | . | . | . | . | . | T | . | . | . | . | . |
| . | . | . | . | . | . | . | . | . | . | . | . | . | . | . | . | . | . | . | . | T | . | . | . | . | . |
| . | . | . | . | . | . | . | . | . | . | . | . | . | . | . | . | . | . | . | . | T | . | . | . | . | . |
| . | . | . | . | . | . | . | . | . | . | . | . | . | . | . | . | . | . | . | . | T | . | . | . | . | . |
| . | . | . | . | . | . | . | . | . | . | . | . | . | . | . | . | . | . | . | . | . | . | . | . | . | . |
| . | . | . | . | . | . | . | . | . | . | . | . | . | . | . | . | . | . | . | . | T | . | . | . | . | . |
| . | . | . | . | . | . | . | . | . | . | . | . | . | . | . | . | . | . | . | . | T | . | . | . | . | . |
| . | . | . | . | . | . | . | . | . | . | . | . | . | . | . | . | . | . | . | . | T | . | . | . | . | . |
| . | . | . | . | T | . | . | . | . | . | . | . | T | . | . | . | . | . | . | . | T | . | . | . | . | . |
| . | . | . | . | T | . | . | . | . | . | . | . | T | . | . | . | . | . | . | . | . | . | . | . | . | . |
| . | . | . | . | . | . | . | . | . | . | . | . | . | . | . | . | . | . | . | . | . | . | . | . | . | . |
| . | . | . | . | . | . | . | . | T | . | . | . | . | . | . | . | . | . | . | . | T | . | . | . | . | . |
| . | . | . | . | . | . | . | . | . | . | . | . | . | . | . | . | . | . | . | . | . | . | . | . | . | . |
| . | . | . | . | . | . | . | . | . | . | . | . | . | . | . | . | . | . | . | . | . | . | . | . | . | . |
| . | . | . | . | . | . | . | . | . | . | . | . | . | . | . | . | . | . | . | . | . | . | . | . | . | . |
| . | . | . | . | . | . | . | . | . | . | . | . | . | . | . | . | . | . | . | . | . | . | . | . | . | . |
| . | . | . | . | . | . | . | . | . | . | . | . | . | . | . | . | . | . | . | . | . | . | . | . | . | . |
| . | . | . | . | . | . | . | . | . | . | . | . | . | . | . | . | . | . | . | . | . | . | . | . | . | . |

```
. . . . . . . . . . . . . . . . . . . . . . . . . .
. . . . . . . . . . . . . . . . . . . . . . . . . .
. . . . . . . . . . . . T . . . . . . . T . . . . .
. . . . . . . . . . . . T . . . . . . . T . . . . .
. . . . . . . . . . . . . . . . . . . . . . . . . .
. . . . . . . . . . . . . . . . . . . . . . . . . .
. . . . . . . . . . . . . . . . . . . . . . . . . .
. . . . . . . . . . . . . . . . . . . . . . . . . .
. . . . . . . . . . . . . . . . . . . . . . . . . .
. . . . . . . . . . . . . . . . . . . . . . . . . .
. . . . . . M W . . . . . . . . . . . . . . . . . .
. . . . . . ? ? . . . . . . . . . . . . . . . . . .
. . . . . . . . . . . . . . . . . . . . . . . . . .
. . . . . . . . . . . . . . . . . . . . . . . . . .
. . . . . . . . . . . . . . . . . . . . . . . . . .
. . . . . . . . . . . . . ? . . . . . . . . . . . .
. . . . . . . . . . . . . . . . . . . . . . . . . .
. . . . . . . . . . . . . . . . . . . . . . . . . .
. . . . . . . . . . . . . . . . . . . . . . . . . .
. . . . . . M W . . . . . . . . . . . . . . . . . .
. . . . . . . . . . . . . . . . . . . . . . . . . .
. . . . . . M W . . . . . . . . . . . . . . . . . .
. . . . . . M W . . . . . . . . . . . . . . . . . .
. . . . . . M W . . . . . . . . . . . . . . . . . .
. . . . . . . . . . . . . ? . . . . . . . . . . . .
. . . . . . M W . . . . . . . . . . . . . . . . . .
. . . . . . . . . . . . . . . . . . . . . . . . . .
. . . . . . ? ? . . . . . . . . . . . . . . . . . .
. . . . . . . . . . . . . . . . . . . . . . . . . .
. . . . . . . . . . . . . . . . . . . . . . . . . .
. . . . . . . . . . . . . . . . . . . . . . . . . .
. . . . . . . . . . . . . . . . . . . . . . . . . .
. . . . . . . . . . . . . . . . . . . . . . . . . .
. . . . . . . . . . . . . . . . . . . . . . . . . .
. . . . . . . . . . . . . . . . . . . . . . . . . .
. . . . . . . . . . . . . . . . . . . . . T . . . .
. . . . . . . . . . . . . . . . . . . . . . . . . .
. . . . . . . . . . . . . . . . . . . . T . . . . .
. . . . . . . . . . . . . . . . . . . . T . . . . .
. . . . . . . . . . . . . . . . . . . . . . . . . .
. . . . . . . . . . . . . . . . . . . . . . . . . .
. . . . . . . . . . . . . . . . . . . . T . . . . .
. . . . . . . . . . . . . . . . . . . . T . . . . .
. . . . . . . . . . . . . . . . . . . . T . . . . .
. . . . . . . . . . . . . . . . . . . . T . . . . .
. . . . . . . . . . . . . . . . . . . . . . . . . .
. . . . . . . . . . . . . . . . . . . . . . . . . .
. . . . . . . . . . . . . . . . . . . . . . . . . .
. . . . . . . . . . . . . . . . . . . . T . . . . .
. . . . . . . . . . . . . . . . . . . . . . . . . .
. . . . . . . . . . . . . . . . . . . . . . . . . .
. . . . . . . . . . . . . . . . . . . . . . . . . .
. . . . . . . . . . . . . . . . . . . . T . . . . .
. . . . . . . . . . . . . . . . . . . . T . . . . .
. . . . . . . . . . . . . . . . . . . . T . . . . .
. . . . . . . . . . . . . . . . . . . . T . . . . .
. . . . . . . . . . . . . . . . . . . . T . . . . .
. . . . . . . . . . . . . . . . . . . . . . . . . .
. . . . . . . . . . . . . . . . . . . . . . . . . .
. . . . . . . . . . . . . . . . . . . . . . . . . .
```

| 186 | 187 | 188 | 189 | 190 | 191 | 192 | 193 | 194 | 195 | 196 | 197 | 198 | 199 | 200 | 201 | 202 | 203 | 204 | 205 | 206 | 207 | 208 | 209 | 210 | 211 |
|---|---|---|---|---|---|---|---|---|---|---|---|---|---|---|---|---|---|---|---|---|---|---|---|---|---|
| S | S | S | S | S | S | S | S | S | S | S | S | S | S | S | ORF3a | ORF3a | ORF3a | ORF3a | ORF3a | ORF3a | ORF3a | ORF3a | ORF3a | ORF3a | NC |
| 23277 | 23311 | 23403 | 23593 | 23638 | 23678 | 23815 | 23821 | 23929 | 24811 | 24933 | 25098 | 25290 | 25314 | 25381 | 25445 | 25461 | 25513 | 25528 | 25563 | 25596 | 25613 | 25855 | 25904 | 26144 | 26226 |
| C | G | A | G | C | G | T | G | C | T | G | T | G | G | A | G | T | C | C | G | A | C | G | C | G | A |
| . | . | . | . | . | . | . | . | . | . | . | . | . | . | . | . | . | . | . | . | . | . | . | . | . | . |
| . | . | . | . | . | . | . | . | . | . | . | . | . | . | . | . | . | . | . | . | . | . | . | . | . | . |
| . | . | G | . | . | . | . | . | . | . | . | . | . | . | . | . | . | . | . | . | . | . | . | . | . | . |
| . | . | G | . | . | . | . | . | . | . | . | . | . | . | . | . | . | . | . | . | . | . | . | . | . | . |
| . | . | G | . | . | . | . | . | . | . | . | . | . | . | . | . | . | . | . | . | . | . | . | . | . | . |
| . | . | G | . | . | . | . | . | . | . | . | . | . | . | . | . | . | . | . | . | . | . | . | . | . | . |
| . | . | G | . | . | . | . | . | . | . | . | . | . | . | . | . | . | . | . | . | . | . | . | . | . | . |
| . | . | G | . | . | . | . | . | . | . | . | . | . | . | . | . | . | . | . | . | . | . | . | . | . | . |
| . | . | G | . | . | . | . | . | . | . | . | . | . | . | . | . | . | . | . | . | . | . | . | . | . | . |
| . | . | G | . | . | . | . | . | . | . | . | . | . | . | . | . | . | . | . | . | . | . | . | . | . | . |
| . | . | G | . | . | . | . | . | . | . | . | . | . | . | . | . | . | . | . | . | . | . | . | . | . | . |
| . | . | G | . | . | . | . | . | . | . | . | . | . | . | . | . | . | . | . | . | . | . | . | . | . | . |
| . | . | G | . | . | . | . | . | . | . | . | . | . | . | . | . | . | . | . | . | . | . | . | . | . | . |
| . | . | G | . | . | . | . | . | . | . | . | . | . | . | . | . | . | . | . | . | . | . | . | . | . | . |
| . | . | G | . | . | . | . | . | . | . | . | . | . | . | . | . | . | . | . | . | . | . | . | . | . | . |
| . | . | G | . | . | . | . | . | . | . | . | . | . | . | . | . | . | . | . | . | . | . | . | . | . | . |
| . | . | . | . | . | . | . | . | . | . | . | . | . | . | . | . | . | . | . | . | . | . | . | . | . | . |
| . | . | . | . | . | . | . | . | . | . | . | . | . | . | . | . | . | . | . | . | . | . | . | . | . | . |
| . | . | . | . | . | . | . | . | . | . | . | . | . | . | . | . | . | . | . | . | . | . | . | . | . | . |
| . | . | . | . | . | . | . | . | . | . | . | . | . | . | . | . | . | . | . | . | . | . | . | . | . | . |
| . | . | . | . | . | . | . | . | . | . | . | . | . | . | . | . | . | . | . | . | . | . | . | . | . | . |
| . | . | . | . | . | . | . | . | . | . | . | . | . | . | . | . | . | . | . | . | . | . | . | . | . | . |
| . | . | . | . | . | . | . | . | . | . | . | . | . | . | . | . | . | . | . | . | . | . | . | T | . | . |
| . | . | . | . | . | . | . | . | . | . | . | . | . | . | . | . | . | . | . | . | . | . | . | . | . | . |
| . | . | . | . | . | . | . | . | . | . | . | . | . | . | . | . | . | . | . | . | . | . | . | . | . | . |
| . | . | . | . | . | . | . | . | . | . | . | . | . | . | . | . | . | . | . | . | . | . | . | . | . | G |
| . | . | . | . | . | . | . | . | . | . | . | . | . | . | . | . | . | . | . | . | . | . | . | . | . | . |
| . | . | G | . | . | . | . | . | . | . | . | . | . | . | . | . | . | . | . | . | . | . | . | . | . | . |
| . | . | . | . | . | . | . | . | . | . | . | . | . | . | . | . | . | . | . | . | . | . | . | . | . | G |
| . | . | G | . | . | . | . | . | . | . | . | . | . | . | . | . | . | . | . | . | . | . | . | . | . | . |
| . | . | G | . | . | . | . | . | . | . | . | . | . | . | . | . | . | . | . | . | . | . | . | . | . | . |
| . | . | G | . | . | . | . | . | . | . | . | . | . | . | . | . | . | . | . | . | . | . | . | . | . | . |
| . | . | G | . | . | . | . | . | . | . | . | . | . | . | . | . | . | . | . | . | . | . | . | . | . | . |
| . | . | G | . | . | . | . | . | . | . | . | . | . | . | . | . | . | . | . | . | . | . | . | . | . | . |
| . | . | G | . | . | . | . | . | . | . | . | . | . | . | . | . | . | . | . | . | . | . | . | . | . | . |
| . | . | G | . | . | . | . | . | . | . | . | . | . | . | . | . | . | . | . | . | . | . | ? | . | . | . |
| . | . | G | . | . | . | . | . | . | . | . | . | . | . | . | . | . | . | . | . | . | . | . | . | . | . |
| . | . | G | . | . | . | . | . | . | . | . | . | . | . | . | . | . | . | . | . | . | . | . | . | . | . |
| . | . | . | . | . | . | . | . | T | . | . | . | . | . | . | . | . | . | . | . | . | . | . | . | . | . |
| . | . | . | . | . | . | . | . | T | . | . | . | . | . | . | . | . | . | . | . | . | . | . | . | . | . |
| . | . | G | . | . | . | . | . | . | . | . | . | . | . | . | . | . | . | . | . | . | . | ? | . | . | . |
| . | . | . | . | . | . | . | . | T | . | . | . | . | . | . | . | . | . | . | . | . | . | . | . | . | . |
| . | . | . | . | . | . | . | . | T | . | . | . | . | . | . | . | . | . | . | . | . | . | . | . | . | . |
| . | . | G | . | . | . | . | . | . | . | T | . | . | . | . | . | . | . | . | . | . | . | . | . | . | . |
| . | . | G | . | . | . | . | . | . | . | . | . | . | . | . | . | . | . | . | . | . | . | . | . | . | . |
| . | . | G | . | . | . | . | . | . | . | . | . | . | . | . | . | . | . | . | . | . | . | . | . | . | . |
| . | . | G | . | . | . | . | . | . | . | . | . | . | . | . | . | . | . | . | . | . | . | . | . | . | . |
| . | . | G | . | . | . | . | . | . | . | T | . | . | . | . | . | . | . | . | . | . | . | . | . | . | . |
| . | . | . | . | . | . | . | . | . | . | . | . | . | . | . | . | . | . | . | . | . | . | . | . | . | . |
| . | . | G | . | . | . | . | . | . | . | . | . | . | . | . | . | . | . | . | . | . | . | . | . | . | . |
| . | . | . | . | . | . | . | . | T | . | . | . | . | . | . | . | . | . | . | . | . | . | . | . | . | . |
| . | . | G | . | . | . | . | . | . | . | . | . | . | . | . | . | . | . | . | . | . | . | . | . | . | . |
| . | . | G | . | . | . | . | . | . | . | . | . | . | . | . | . | . | . | . | . | . | . | . | . | . | . |
| . | . | G | . | . | . | . | . | . | . | . | . | . | . | . | . | . | . | . | . | . | . | . | . | . | . |

| | | | | | | | | | | | | | | | | | | | | | | | | | |
|---|---|---|---|---|---|---|---|---|---|---|---|---|---|---|---|---|---|---|---|---|---|---|---|---|---|
| . | . | G | . | . | . | . | . | . | . | . | . | . | . | . | . | . | . | . | T | . | . | . | . | . | . |
| . | . | G | . | . | . | . | . | . | . | . | . | . | . | . | . | . | . | . | T | . | . | . | . | . | . |
| . | . | G | . | T | . | . | . | . | . | . | . | . | . | . | . | . | . | . | . | . | . | . | . | . | . |
| . | . | G | . | . | . | . | . | . | . | . | . | . | . | . | . | . | . | . | . | . | . | . | . | . | . |
| . | . | G | . | . | . | . | . | . | . | . | . | . | . | . | . | . | . | . | T | . | . | . | . | . | . |
| . | . | G | . | . | . | . | . | . | . | . | . | . | . | . | . | . | . | . | . | . | . | . | . | . | . |
| . | . | G | . | . | . | . | . | . | . | . | . | . | . | . | . | . | . | . | . | . | . | . | . | . | . |
| . | . | G | . | . | . | . | . | . | . | g | . | . | . | . | . | . | . | . | . | . | . | . | . | . | . |
| . | . | G | . | . | . | . | . | . | . | . | . | . | . | . | . | . | . | . | . | . | . | . | . | . | . |
| . | . | G | . | . | . | . | . | . | . | . | . | . | . | . | . | . | . | . | T | . | . | . | . | . | . |
| . | . | G | . | . | . | . | . | . | . | . | . | . | . | . | . | . | . | . | T | . | . | . | . | . | . |
| . | . | G | . | . | . | . | . | . | . | . | . | . | . | . | . | . | . | . | T | . | . | . | . | . | . |
| . | . | G | . | . | . | . | . | . | . | . | . | . | . | . | . | . | . | . | T | . | . | . | . | . | . |
| . | . | G | . | . | . | . | . | . | . | . | . | . | . | . | . | . | . | . | T | . | . | . | . | . | . |
| . | . | G | . | . | . | . | . | . | . | . | . | . | . | . | . | . | . | . | T | . | . | . | . | . | . |
| . | . | G | . | . | . | . | . | . | . | . | . | . | . | . | . | . | . | . | T | . | . | . | . | . | . |
| . | . | G | . | . | . | . | . | . | . | . | . | . | . | . | . | . | . | . | . | . | . | . | . | . | . |
| . | . | G | . | . | . | . | . | . | . | . | . | . | . | . | . | . | . | . | . | . | . | . | . | . | . |
| . | . | G | . | . | . | . | . | . | . | . | . | . | . | . | . | . | . | . | . | . | . | . | . | . | . |
| . | . | . | . | . | . | . | . | T | . | . | . | . | . | ? | . | . | . | . | . | . | . | . | . | . | . |
| . | . | . | . | . | . | . | . | c | . | . | . | . | . | . | . | . | . | . | . | . | . | . | . | . | . |
| . | . | . | . | . | . | . | . | T | . | . | . | . | . | . | . | . | . | . | . | . | . | . | . | . | . |
| . | . | . | . | . | . | . | . | Y | . | . | . | . | . | . | . | . | . | . | . | . | . | . | . | . | . |
| . | . | . | . | . | . | . | . | T | . | . | . | . | . | . | g | t | . | . | . | . | . | . | . | . | . |
| . | . | . | . | . | . | . | . | Y | . | . | . | . | . | . | . | . | . | . | . | . | . | . | . | . | . |
| . | . | . | . | . | . | . | . | c | . | . | . | . | . | . | . | . | . | . | . | . | . | . | . | . | . |
| . | . | . | . | . | . | . | . | c | . | . | . | . | . | . | . | . | . | . | . | . | . | . | . | . | . |
| . | . | . | . | . | . | . | . | ? | . | . | . | . | . | . | . | . | . | . | . | . | . | . | . | . | . |
| . | . | . | . | . | . | . | . | t | . | . | . | . | . | . | . | . | . | . | . | . | . | . | . | . | . |
| . | . | . | . | . | . | . | . | T | . | . | . | . | . | . | . | . | . | . | . | . | . | . | . | . | . |
| . | . | . | . | . | . | . | . | T | . | . | . | . | . | . | . | . | . | . | . | . | . | . | . | . | . |
| . | . | . | . | . | . | . | . | T | . | . | . | . | . | . | . | . | . | . | . | . | . | . | . | . | . |
| . | . | . | . | . | . | . | . | T | . | . | . | . | . | . | . | . | . | . | . | . | . | . | . | . | . |
| . | . | . | . | . | . | . | . | T | . | . | . | . | . | . | . | . | . | . | . | . | . | . | . | . | . |
| . | . | . | . | . | . | . | . | T | . | . | . | . | . | . | . | . | . | . | . | . | . | . | . | . | . |
| . | . | . | . | . | . | . | . | T | . | . | . | . | . | . | . | . | . | . | . | . | . | . | . | . | . |
| . | . | . | . | . | . | . | . | T | . | . | . | . | . | . | . | . | . | . | . | . | . | . | . | . | . |
| . | . | a | . | . | . | . | . | T | . | . | . | . | . | . | . | . | . | . | . | . | . | . | . | . | . |
| . | . | . | . | . | . | . | . | T | . | . | . | . | . | . | . | . | . | . | . | . | . | . | . | . | . |
| c | g | . | . | . | . | . | . | T | . | . | . | . | . | . | . | . | . | . | . | . | . | . | . | . | . |
| . | . | . | . | . | . | . | . | T | . | . | . | . | . | . | g | t | c | c | . | . | . | . | . | . | . |
| . | . | . | . | . | . | . | . | . | . | . | . | . | . | . | . | C | . | . | . | . | . | . | . | . | . |
| . | . | . | . | . | . | . | . | . | . | . | . | . | . | . | . | C | . | . | . | . | . | . | . | . | . |
| . | . | . | . | . | . | . | . | . | . | . | . | . | . | . | . | C | . | . | . | . | . | . | . | . | . |
| . | . | . | . | . | . | . | . | . | . | . | . | . | . | . | . | C | . | . | . | . | . | . | . | . | . |
| . | . | . | . | . | . | . | . | . | . | . | . | . | . | . | . | C | . | . | . | . | . | . | . | . | . |
| . | . | . | . | . | . | . | . | . | . | . | . | . | . | . | . | . | . | . | . | . | . | . | . | . | . |
| . | . | . | . | . | . | . | . | . | . | . | . | . | . | . | . | . | . | . | . | . | . | . | . | T | . |
| . | . | . | . | . | . | . | . | . | . | . | . | . | . | . | . | . | . | . | . | . | . | . | . | T | . |
| . | . | . | . | . | . | . | . | c | . | . | . | . | . | . | . | . | . | . | . | . | . | . | . | . | . |
| . | . | . | . | . | . | . | . | . | . | . | . | . | . | . | . | . | . | . | . | . | . | . | . | . | . |
| . | . | . | . | . | . | . | . | T | . | . | . | . | . | . | . | . | . | . | . | . | . | . | . | . | . |
| . | . | G | . | . | . | . | . | . | . | . | . | . | . | . | . | . | . | . | . | . | . | . | . | . | . |
| . | . | . | . | . | . | . | . | T | . | . | . | . | . | . | . | . | . | . | . | . | . | . | . | . | . |
| . | . | . | . | . | . | . | . | T | . | . | . | . | . | . | . | . | . | . | . | . | . | . | . | . | . |
| . | . | . | . | . | . | . | . | T | . | . | . | . | . | . | . | . | . | . | . | . | . | . | . | . | . |
| . | . | . | . | . | . | . | . | T | . | . | . | . | . | . | . | . | . | . | . | . | . | . | . | . | . |
| . | . | . | . | . | . | . | . | T | . | . | . | . | . | . | . | . | . | . | . | . | . | . | . | . | . |
| . | . | . | . | . | . | . | . | T | . | . | . | . | . | . | . | . | . | . | . | . | . | . | . | . | . |
| . | . | . | . | . | . | . | . | T | . | . | . | . | . | . | . | . | . | . | . | . | . | . | . | . | . |
| . | . | . | . | . | . | . | . | T | . | . | . | . | . | . | . | . | . | . | . | . | . | . | . | . | . |
| . | . | . | . | . | . | . | . | T | . | . | . | . | . | . | . | . | . | . | . | . | . | . | . | . | . |

| | | | | | | | | | | | | | | | | | | | | | | | | | |
|---|---|---|---|---|---|---|---|---|---|---|---|---|---|---|---|---|---|---|---|---|---|---|---|---|---|
| . | . | . | . | . | . | . | . | T | . | . | . | . | . | . | . | . | . | . | . | . | . | . | . | . | . |
| . | . | . | . | . | . | . | . | T | . | . | . | . | . | . | . | . | . | . | . | . | . | . | . | . | . |
| . | . | . | . | . | . | . | . | T | . | . | . | . | . | . | . | . | . | . | . | . | . | . | . | . | . |
| . | . | . | . | . | . | . | . | T | . | . | . | . | . | . | . | . | . | . | . | . | . | . | . | . | . |
| . | . | . | . | . | . | . | . | T | . | . | . | . | . | . | . | . | . | . | . | . | . | . | . | . | . |
| . | . | . | . | . | . | . | . | T | . | . | . | . | . | . | . | . | . | . | . | . | . | . | . | . | . |
| . | . | . | . | . | . | . | . | T | . | . | . | . | . | . | . | . | . | . | . | . | . | . | . | . | . |
| . | . | . | . | . | . | . | . | T | . | . | . | . | . | . | . | . | . | . | . | . | . | . | . | . | . |
| . | . | . | . | . | . | . | . | Y | . | . | . | . | . | . | . | . | . | . | . | . | . | . | . | . | . |
| . | . | . | . | . | . | . | . | T | . | . | . | . | . | . | . | . | . | . | . | . | . | . | . | . | . |
| . | . | . | . | . | . | . | . | T | . | . | . | . | . | . | . | . | . | . | . | . | . | . | . | . | . |
| . | . | . | . | . | . | . | . | T | . | . | . | . | . | . | . | . | . | . | . | . | . | . | . | . | . |
| . | . | . | . | . | . | . | . | T | . | . | . | . | . | . | . | . | . | . | . | . | . | . | . | . | . |
| . | . | . | . | . | . | . | . | T | . | . | . | . | . | . | . | . | . | . | . | . | . | . | . | . | . |
| . | . | . | . | . | . | . | . | T | . | . | . | . | . | . | . | . | . | . | . | . | . | . | . | g | . |
| . | . | . | . | . | . | . | . | T | . | . | . | . | . | . | . | . | . | . | . | . | . | . | . | . | . |
| . | . | . | . | . | . | . | . | t | . | . | . | . | . | . | . | . | . | . | . | . | . | . | . | . | . |
| . | . | . | . | . | . | . | . | T | . | . | . | . | . | . | . | . | . | . | . | . | . | . | . | . | . |
| . | . | . | . | . | . | . | . | T | . | . | . | . | . | . | . | . | . | . | . | . | . | . | . | . | . |
| . | . | . | . | . | . | . | . | T | . | . | . | . | . | . | . | . | . | . | . | . | . | . | . | . | . |
| . | . | . | . | . | . | . | . | T | . | . | . | . | . | . | . | . | . | . | . | . | . | . | . | . | . |
| . | . | ? | . | . | . | . | . | . | . | . | . | . | . | . | . | . | . | . | . | . | . | . | . | . | . |
| . | . | G | . | . | . | . | . | . | . | . | . | . | . | . | . | . | . | . | . | . | . | . | . | . | . |
| . | . | G | . | . | . | . | . | . | . | . | . | . | . | . | . | . | . | . | . | . | . | . | . | . | . |
| . | . | G | . | . | . | . | . | . | . | . | . | . | . | . | . | . | . | . | . | . | . | . | . | . | . |
| . | . | G | . | . | . | . | . | . | . | . | . | . | . | . | . | . | . | . | . | . | . | . | . | . | . |
| . | . | R | . | . | . | . | . | . | . | . | . | . | . | . | . | . | . | . | . | . | . | . | . | . | . |
| . | . | G | . | . | . | . | . | . | . | . | . | . | . | . | . | . | . | . | T | . | . | . | . | . | . |
| . | . | G | . | . | . | . | . | . | . | . | . | . | . | . | . | . | . | . | . | . | . | . | . | . | . |
| . | . | G | . | . | . | . | . | . | . | . | . | . | . | . | . | . | . | . | . | . | . | . | . | . | . |
| . | . | G | . | . | . | . | . | . | . | . | . | . | . | . | . | . | . | . | . | . | . | . | . | . | . |
| . | . | R | . | . | . | . | . | . | . | . | . | . | . | . | . | . | . | . | . | . | . | . | . | . | . |
| . | . | . | . | . | . | . | . | T | . | . | . | . | . | . | . | . | . | . | . | . | . | . | . | . | . |
| . | . | G | . | . | . | . | . | . | . | . | . | . | . | . | . | . | . | . | T | . | . | . | . | . | . |
| . | . | . | . | . | . | . | . | . | . | . | . | . | . | . | . | . | . | . | . | . | . | . | . | . | . |
| . | . | . | . | . | . | . | . | T | . | . | . | . | . | . | . | . | . | . | . | . | . | . | . | . | . |
| . | . | . | . | . | . | . | . | T | . | . | . | . | . | . | . | . | . | . | . | . | . | . | . | . | . |
| . | . | G | . | . | T | . | . | . | . | . | . | T | . | . | . | . | . | . | T | . | . | T | . | . | . |
| . | . | G | . | . | . | . | . | . | . | . | . | T | . | . | . | . | . | . | T | . | . | T | . | . | . |
| . | . | G | . | . | . | . | . | . | . | . | . | . | . | . | . | . | . | . | . | . | . | . | . | . | . |
| . | . | G | . | . | . | . | . | . | . | . | . | . | . | . | . | . | . | . | . | . | . | . | . | . | . |
| . | . | G | . | . | . | . | . | . | . | . | . | . | . | . | . | . | . | . | T | . | . | . | . | . | . |
| . | . | G | T | . | . | . | . | . | . | . | . | . | . | . | . | . | . | . | . | . | . | . | . | . | . |
| . | . | G | . | . | . | . | . | . | . | . | . | . | . | . | . | . | . | . | . | . | . | . | . | . | . |
| . | . | G | . | . | . | . | . | . | . | . | . | . | . | . | . | . | . | . | T | . | . | . | . | . | . |
| . | . | . | . | . | . | . | . | T | . | . | . | . | . | . | . | . | . | . | . | . | . | . | . | . | . |
| . | T | . | . | . | . | . | . | T | . | . | . | . | . | . | . | . | . | . | T | . | . | . | T | . | . |
| . | . | G | . | . | . | . | . | . | . | . | . | . | . | . | . | . | . | . | . | . | . | . | . | . | . |
| . | . | G | . | . | . | . | . | . | . | . | . | . | . | . | . | . | . | . | T | . | . | . | . | . | . |
| . | . | . | . | . | . | . | . | T | . | . | . | . | . | . | . | . | . | . | . | . | . | . | . | . | . |
| . | . | . | . | . | . | . | . | T | . | . | . | . | . | . | . | . | . | . | . | . | . | . | . | . | . |
| . | . | G | . | . | . | . | . | . | . | . | . | . | . | . | . | . | . | . | T | . | . | . | . | . | . |
| . | . | G | . | . | . | . | . | . | . | . | . | . | . | . | . | . | . | . | T | . | . | . | . | . | . |
| . | . | G | . | . | . | . | . | . | . | . | . | . | . | . | . | . | . | . | . | . | . | . | . | . | . |
| . | T | G | . | . | . | . | . | . | . | . | . | . | . | . | . | . | . | . | T | . | . | . | . | . | . |
| . | . | G | . | . | . | . | . | . | . | . | . | . | . | . | . | . | . | . | . | . | . | . | . | . | . |
| . | . | G | . | . | . | . | . | . | . | . | . | . | . | . | . | . | . | . | T | . | . | . | . | . | . |
| . | . | G | . | . | . | . | . | . | . | . | . | . | . | . | . | . | . | . | . | . | . | . | . | . | . |
| . | . | G | T | . | . | . | . | . | . | . | . | . | . | . | . | . | . | . | . | . | . | . | . | . | . |
| . | . | G | . | . | . | . | . | . | . | . | . | . | . | . | . | . | . | . | T | . | . | . | . | . | . |
| T | . | G | . | . | . | . | . | . | . | . | . | . | . | . | . | . | . | . | T | . | . | . | . | . | . |
| T | . | G | . | . | . | . | . | . | . | . | . | . | . | . | . | . | . | . | T | . | . | . | . | . | . |

| | | | | | | | | | | | | | | | | | | | | | | | | | |
|---|---|---|---|---|---|---|---|---|---|---|---|---|---|---|---|---|---|---|---|---|---|---|---|---|---|
| . | . | G | . | . | . | . | . | . | . | . | . | . | . | . | . | . | . | . | T | . | . | . | . | . | . |
| . | . | G | . | . | . | . | . | . | . | . | . | . | . | . | . | . | . | . | . | . | . | . | . | . | . |
| . | T | G | . | . | . | . | . | . | . | . | . | . | . | . | . | . | . | . | T | . | . | . | . | . | . |
| . | . | G | . | . | . | . | . | . | . | . | . | . | . | . | . | . | . | . | T | . | . | . | . | . | . |
| . | . | G | . | . | . | . | . | . | . | . | . | . | . | . | . | . | . | . | T | . | . | . | . | . | . |
| . | . | G | . | . | . | . | . | . | . | . | . | . | . | . | . | . | . | . | T | . | . | . | . | . | . |
| . | . | G | . | . | . | . | . | . | . | . | . | . | . | . | . | . | . | . | . | . | . | . | . | . | . |
| . | . | G | . | . | . | . | . | . | . | . | . | . | . | . | . | . | . | . | T | . | . | . | . | . | . |
| . | . | G | . | . | . | . | . | . | . | . | . | . | . | . | . | . | . | . | . | . | . | . | . | . | . |
| T | . | G | . | . | . | . | . | . | . | . | . | . | . | . | . | . | . | . | T | . | . | . | . | . | . |
| . | . | G | . | . | . | . | . | . | . | . | . | . | . | . | . | . | . | . | . | . | . | . | . | . | . |
| . | . | G | . | . | . | . | . | . | . | . | . | . | . | . | . | . | . | . | . | . | . | . | . | . | . |
| . | . | G | . | T | . | . | . | . | . | . | . | . | . | . | . | . | . | . | . | . | . | . | . | . | . |
| . | . | G | . | . | . | . | . | . | . | . | . | . | . | . | . | . | . | . | T | . | . | . | . | . | . |
| . | . | G | . | . | . | . | . | . | . | . | . | . | . | . | . | . | . | . | . | . | . | . | . | . | . |
| . | . | G | . | . | . | . | . | . | . | . | . | . | . | . | . | . | . | . | T | . | . | . | . | . | . |
| . | . | G | . | . | . | . | . | . | . | . | . | . | . | . | . | . | . | . | . | . | . | . | . | . | . |
| . | . | G | . | . | . | . | . | . | . | . | . | . | . | . | . | . | . | . | T | . | . | . | . | . | . |
| . | . | G | . | . | . | . | . | . | . | . | . | . | . | . | . | . | . | . | T | . | . | . | . | . | . |
| . | . | G | . | . | . | . | . | . | . | . | . | . | . | . | . | . | . | . | T | . | . | . | . | . | . |
| . | . | G | . | . | . | . | . | . | . | . | . | . | . | . | . | . | . | . | . | . | . | . | . | . | . |
| . | . | G | . | . | . | . | . | . | . | . | . | . | . | . | . | . | . | . | . | . | . | . | . | . | . |
| . | . | G | . | . | . | . | . | . | . | . | . | . | . | . | . | . | . | . | . | . | . | . | . | . | . |
| . | . | G | . | . | . | . | . | . | . | . | . | . | . | . | . | . | . | . | T | . | . | . | . | . | . |
| . | T | G | . | . | . | . | . | . | . | . | . | . | . | . | . | . | . | . | T | . | . | . | . | . | . |
| T | . | G | . | . | . | . | . | . | . | . | . | . | . | . | . | . | . | . | T | . | . | . | . | . | . |
| . | . | G | . | . | . | . | . | . | . | . | . | . | . | . | . | . | . | . | T | . | . | . | . | . | . |
| . | . | G | . | . | . | . | . | . | . | . | . | . | . | . | . | . | . | . | T | . | . | . | . | . | . |
| . | . | G | . | . | . | . | . | . | . | . | . | . | . | . | . | . | . | . | . | . | . | . | . | . | . |
| . | . | G | . | . | . | . | . | . | . | . | . | . | . | . | . | . | . | . | T | . | . | . | . | . | . |
| . | . | G | . | . | . | . | . | . | . | . | . | . | . | . | . | . | . | . | T | . | . | . | . | . | . |
| . | . | G | . | . | . | . | . | . | . | . | . | . | . | . | . | . | . | . | T | . | . | . | . | . | . |
| . | . | G | . | . | . | . | . | . | . | . | . | . | . | . | . | . | . | . | T | . | . | . | . | . | . |
| . | T | G | . | . | . | . | . | . | . | . | . | . | . | . | . | . | . | . | T | . | . | . | . | . | . |
| . | T | G | . | . | . | . | . | . | . | . | . | . | . | . | . | . | . | . | T | . | . | . | . | . | . |
| . | . | G | . | . | . | . | . | . | . | . | . | . | . | . | . | . | . | . | T | . | . | . | . | . | . |
| . | . | G | . | . | . | . | . | . | . | . | . | . | . | . | . | . | . | . | T | . | . | . | . | . | . |
| . | . | G | . | . | . | . | . | . | . | . | . | . | . | . | . | . | . | . | . | . | . | . | . | . | . |
| . | . | G | . | . | . | . | . | . | . | . | . | . | . | . | . | . | . | . | T | . | . | . | . | . | . |
| . | . | G | . | . | T | . | . | . | . | . | . | . | . | . | . | . | . | . | T | . | . | . | . | . | . |
| . | . | G | . | . | . | . | . | . | . | . | . | . | . | . | . | . | . | . | T | . | . | . | . | . | . |
| . | . | G | . | . | . | . | . | . | . | . | . | . | . | . | . | . | . | . | T | . | . | . | . | . | . |
| . | . | G | . | T | . | . | . | . | . | . | . | . | . | . | . | . | . | . | . | . | . | . | . | . | . |
| . | . | G | . | . | . | . | . | . | . | . | . | . | . | . | . | . | . | . | T | . | . | . | . | . | . |
| . | . | G | . | . | . | . | . | . | . | . | . | . | . | . | . | . | . | . | T | . | . | . | . | . | . |
| . | . | G | . | . | . | . | . | . | . | . | . | . | . | . | . | . | . | . | T | . | . | . | . | . | . |
| . | . | G | . | . | . | . | . | . | . | . | . | . | . | . | . | . | . | . | T | . | . | . | . | . | . |
| . | . | G | . | . | . | . | . | . | . | . | . | . | . | . | . | . | . | . | T | . | . | . | . | . | . |
| . | . | G | . | . | . | . | . | . | . | . | . | . | . | . | . | . | . | . | T | . | . | . | . | . | . |
| . | . | G | . | . | . | . | . | . | . | . | . | . | . | . | . | . | . | . | T | . | . | . | . | . | . |
| . | . | G | . | . | . | . | . | . | . | . | . | . | . | . | . | . | . | . | T | . | . | . | . | . | . |
| . | . | G | . | . | . | . | . | . | . | . | . | . | . | . | . | . | . | . | T | . | . | . | . | . | . |
| . | . | G | . | . | . | . | . | . | . | . | . | . | . | . | . | . | . | . | T | . | . | . | . | . | . |
| . | . | G | . | . | . | . | . | . | . | . | . | . | . | . | . | . | . | . | T | . | . | . | . | . | . |
| . | . | G | . | . | . | . | . | . | . | . | . | . | . | . | . | . | . | . | . | . | . | . | . | . | . |
| . | . | G | . | . | . | . | . | . | . | . | . | . | . | . | . | . | . | . | . | . | . | . | . | . | . |
| . | T | G | . | . | . | . | . | . | . | . | . | . | . | . | . | . | . | . | T | . | . | . | . | . | . |
| . | . | G | . | T | . | . | . | . | . | . | . | . | . | . | . | . | . | . | . | . | . | . | . | . | . |
| . | . | G | . | . | . | . | . | . | . | . | . | . | . | . | . | . | . | . | T | . | . | . | . | . | . |
| . | . | G | . | . | . | . | . | . | . | . | . | . | . | . | . | . | . | . | . | . | . | . | . | . | . |
| . | . | G | . | . | . | . | . | . | . | . | . | . | . | . | . | . | . | . | . | . | . | . | . | . | . |
| . | . | G | . | . | . | . | . | . | . | . | . | . | . | . | . | . | . | . | . | . | . | . | . | . | . |

| | | | | | | | | | | | | | | | | | | | | | | | | | |
|---|---|---|---|---|---|---|---|---|---|---|---|---|---|---|---|---|---|---|---|---|---|---|---|---|---|
| . | . | G | . | . | . | . | . | . | . | . | . | . | . | . | . | . | . | . | T | . | . | . | . | . | . |
| . | . | G | . | . | . | . | . | . | . | . | . | . | . | . | . | . | . | . | T | . | . | . | . | . | . |
| . | . | G | . | . | . | . | . | . | . | . | . | . | . | . | . | . | . | . | . | . | . | . | . | . | . |
| . | . | G | . | . | . | . | . | . | . | . | . | . | . | . | . | . | . | . | T | . | . | . | . | . | . |
| . | . | . | . | . | . | . | . | T | . | . | . | . | . | . | . | . | . | . | . | . | . | . | . | . | . |
| . | . | . | . | . | . | . | . | T | . | . | . | . | . | . | . | . | . | . | . | . | . | . | . | . | . |
| . | . | . | . | . | . | . | . | T | . | . | . | . | . | . | . | . | . | . | . | . | . | . | . | . | . |
| . | . | . | . | . | . | . | . | T | . | . | . | . | . | . | . | . | . | . | . | . | . | . | . | . | . |
| . | . | . | . | . | . | . | . | T | . | . | . | . | . | . | . | . | . | . | . | . | . | . | . | . | . |
| . | . | . | . | . | . | . | . | T | . | . | . | . | . | . | . | . | . | . | . | . | . | . | . | . | . |
| . | . | . | . | . | . | . | . | T | . | . | . | . | . | . | . | . | . | . | . | . | . | . | . | . | . |
| . | . | . | . | . | . | . | . | T | . | . | . | . | . | . | . | . | . | . | . | . | . | . | . | . | . |
| . | . | . | . | . | . | . | . | T | . | . | . | . | . | . | . | . | . | . | . | . | . | . | . | . | . |
| . | . | . | . | . | . | . | . | . | . | . | . | . | . | . | . | . | . | . | . | . | . | . | . | . | . |
| . | . | . | . | . | . | . | . | T | . | . | . | . | . | . | . | . | . | . | . | . | . | . | . | . | . |
| . | . | . | . | . | . | . | . | T | . | . | . | . | . | . | . | . | . | . | . | . | . | . | . | . | . |
| . | . | . | . | . | . | . | . | T | . | . | . | . | . | . | . | . | . | . | . | . | . | . | . | . | . |
| . | . | . | . | . | . | . | . | T | . | . | . | . | . | . | . | . | . | . | . | . | . | . | . | . | . |
| . | . | . | . | . | . | . | . | T | . | . | . | . | . | . | . | . | . | . | . | . | . | . | . | . | . |
| . | . | . | . | . | . | . | . | T | . | . | . | . | . | . | . | . | . | . | . | . | . | . | . | . | . |
| . | . | . | . | . | . | . | . | T | . | . | . | . | . | . | . | . | . | . | . | . | . | . | . | . | . |
| . | . | . | . | . | . | . | . | T | . | . | . | . | . | . | . | . | . | . | . | . | . | . | . | . | . |
| . | . | . | . | . | . | . | . | T | . | . | . | . | . | . | . | . | . | . | . | . | . | . | . | . | . |
| . | . | . | . | . | . | . | . | T | . | . | . | . | . | . | . | . | . | . | . | . | . | . | . | . | . |
| . | . | . | . | . | . | . | . | T | . | . | . | . | . | . | . | . | . | . | . | . | . | . | . | . | . |
| . | . | . | . | . | . | . | . | T | . | . | . | . | . | . | . | . | . | . | . | . | . | . | . | . | . |
| . | . | . | . | . | . | . | . | T | . | . | . | . | . | . | . | . | . | . | . | . | . | . | . | . | . |
| . | . | . | . | . | . | . | . | T | . | . | . | . | . | . | . | . | . | . | . | . | . | . | . | . | . |
| . | . | . | . | . | . | . | . | T | . | . | . | . | . | . | . | . | . | . | . | . | . | . | . | . | . |
| . | . | . | . | . | . | . | . | T | . | . | . | . | . | . | . | . | . | . | . | . | . | . | . | . | . |
| . | . | . | . | . | . | . | . | T | . | . | . | . | . | . | . | . | . | . | . | . | . | . | . | . | . |
| . | . | . | . | . | . | . | . | T | . | . | . | . | . | . | . | . | . | . | . | . | . | . | . | . | . |
| . | . | . | . | . | . | . | . | T | . | . | . | . | . | . | . | . | . | . | . | . | T | . | . | . | . |
| . | . | . | . | . | . | . | . | T | . | . | . | . | . | . | . | . | . | . | . | . | T | . | . | . | . |
| . | . | . | . | . | . | . | . | T | . | . | . | . | . | . | . | . | . | . | . | . | T | . | . | . | . |
| . | . | G | . | . | . | . | . | . | . | . | . | . | . | . | . | . | . | . | . | . | . | . | . | . | . |
| . | . | . | . | . | . | . | . | T | . | . | . | . | . | . | . | . | . | . | . | . | . | . | . | . | . |
| . | . | . | . | . | . | . | . | T | . | . | . | . | . | . | . | . | . | . | . | . | . | . | . | . | . |
| . | . | . | . | . | . | . | . | T | . | . | . | . | . | . | . | . | . | . | . | . | . | . | . | . | . |
| . | . | . | . | . | . | . | . | T | . | . | . | . | . | . | . | . | . | . | . | . | . | . | . | . | . |
| . | . | . | . | . | . | . | . | . | . | . | . | . | . | . | . | . | . | . | . | . | . | . | . | . | . |
| . | . | . | . | . | . | . | . | T | . | . | . | . | . | . | . | . | . | . | . | . | . | . | . | . | . |
| . | . | . | . | . | . | . | . | . | . | . | . | . | . | . | . | . | . | . | . | . | . | . | . | . | . |
| . | . | . | . | . | . | . | . | T | . | . | . | . | . | . | . | . | . | . | . | . | . | . | . | . | . |
| . | . | . | . | . | . | . | . | T | . | . | . | . | . | . | . | . | . | . | . | . | . | . | . | . | . |
| . | . | . | . | . | . | . | . | T | . | . | . | . | . | . | . | . | . | . | . | . | . | . | . | . | . |
| . | . | . | . | . | . | . | . | T | . | . | . | . | . | . | . | . | . | . | . | . | . | . | . | . | . |
| . | . | . | . | . | . | . | . | T | . | . | . | . | . | . | . | . | . | . | . | . | . | . | . | . | . |
| . | . | . | . | . | . | . | . | T | . | . | . | . | . | . | . | . | . | . | . | . | . | . | . | . | . |
| . | . | . | . | . | . | . | . | T | . | . | . | . | . | . | . | . | . | . | . | . | . | . | . | . | . |
| . | . | . | . | . | . | . | . | T | . | . | . | . | . | . | . | . | . | . | . | . | . | . | . | . | . |
| . | . | . | . | . | . | . | . | T | . | . | . | . | . | . | . | . | . | . | . | . | . | . | . | . | . |
| . | . | . | . | . | . | . | . | T | . | . | . | . | . | . | . | . | . | . | . | . | . | . | . | . | . |
| . | . | . | . | . | . | . | . | T | . | . | . | . | . | . | . | . | . | . | . | . | . | . | . | . | . |
| . | . | . | . | . | . | . | . | T | . | . | . | . | . | . | . | . | . | . | . | . | . | . | . | . | . |
| . | . | . | . | . | . | . | . | T | . | . | . | . | . | . | . | . | . | . | . | . | . | . | . | . | . |
| . | . | . | . | . | . | . | . | T | . | . | . | . | . | . | . | . | . | . | . | . | . | . | . | . | . |
| . | . | . | . | . | . | . | . | T | . | . | . | . | . | . | . | . | . | . | . | . | . | . | . | . | . |
| . | . | G | . | . | . | . | . | . | . | . | . | . | . | . | . | . | . | . | . | . | . | . | . | . | . |
| . | . | . | . | . | . | . | . | . | . | . | . | . | . | . | . | . | . | . | . | . | . | . | . | . | . |
| . | . | G | . | . | . | . | . | . | . | . | . | . | . | . | . | . | . | . | . | . | . | . | . | . | . |
| . | . | . | . | . | . | . | . | T | . | . | . | . | . | . | . | . | . | . | . | . | . | . | . | . | . |

| | | | | | | | | | | | | | | | | | | | | | | | | | |
|---|---|---|---|---|---|---|---|---|---|---|---|---|---|---|---|---|---|---|---|---|---|---|---|---|---|
| . | . | . | . | . | . | . | . | T | . | . | . | . | . | . | . | . | . | . | . | . | . | . | . | . | . |
| . | . | . | . | . | . | . | . | T | . | . | . | . | . | . | . | . | . | . | . | . | . | . | . | . | . |
| . | . | . | . | . | . | . | . | T | . | . | . | . | . | . | . | . | . | . | . | . | . | . | . | . | . |
| . | . | . | . | . | . | . | . | T | . | . | . | . | . | . | . | . | . | . | . | . | . | . | . | . | . |
| . | . | . | . | . | . | . | . | T | . | . | . | . | . | . | . | . | . | . | . | . | . | . | . | . | . |
| . | . | . | . | . | . | . | . | T | . | . | . | . | . | . | . | . | . | . | . | . | . | . | . | . | . |
| . | . | G | . | . | . | . | . | . | . | . | . | . | . | . | . | . | . | . | . | . | . | . | . | . | . |
| . | T | G | . | . | . | . | . | . | A | . | . | . | . | . | . | . | . | . | T | . | . | . | . | . | . |
| . | . | G | . | . | . | . | . | . | A | . | . | . | . | . | . | . | . | . | T | . | . | . | . | . | . |
| . | . | . | . | . | . | . | . | T | . | . | . | . | . | . | . | . | . | . | . | . | . | . | . | . | . |
| . | . | G | . | . | . | . | . | . | . | . | . | . | . | . | . | . | . | . | . | . | . | . | . | . | . |
| . | . | G | . | . | . | . | . | . | . | . | . | . | . | . | . | . | . | . | . | . | . | . | . | . | . |
| . | . | . | T | . | . | . | . | T | . | . | . | . | . | . | . | . | . | . | . | . | . | . | . | . | . |
| . | . | G | . | . | . | . | . | . | . | . | . | . | . | . | . | . | . | . | . | . | . | . | . | . | . |
| . | . | G | . | . | . | . | . | . | . | . | . | . | . | . | . | . | . | . | T | . | . | . | . | . | . |
| . | . | . | . | . | . | . | . | . | . | . | . | . | . | . | . | . | T | . | . | . | . | . | . | . | . |
| . | . | . | . | . | . | . | . | . | . | . | . | . | . | . | . | . | T | . | . | . | . | . | . | . | . |
| . | . | G | . | . | . | . | . | . | . | . | . | . | . | . | . | . | . | . | T | . | . | . | . | . | . |
| . | . | G | . | . | . | . | . | . | . | . | . | . | . | . | . | . | . | . | . | . | . | . | . | . | . |
| . | . | G | . | . | . | . | . | . | . | . | . | . | . | . | . | . | . | . | T | . | . | . | . | . | . |
| . | . | G | . | . | . | . | . | . | . | . | . | . | . | . | . | . | . | . | T | . | . | . | . | . | . |
| . | . | G | . | . | . | . | . | . | . | . | . | . | . | . | . | . | . | . | T | . | . | . | . | . | . |
| . | . | . | . | . | . | . | . | T | . | . | . | . | . | . | . | . | . | . | . | . | . | . | . | . | . |
| . | . | G | . | . | . | . | . | . | . | . | . | . | . | . | . | . | . | . | . | . | . | . | . | . | . |
| . | . | . | . | . | . | . | . | T | . | . | . | . | . | . | . | . | . | . | . | . | . | . | . | . | . |
| . | . | G | . | . | . | . | . | . | . | . | . | . | . | . | . | . | . | . | . | . | . | . | . | . | . |
| . | . | G | . | . | . | . | . | . | . | . | . | . | . | . | . | . | . | . | . | . | . | . | . | . | . |
| . | . | . | . | . | . | . | . | T | . | . | . | . | . | . | . | . | . | . | . | . | . | . | . | . | . |
| . | . | G | . | . | . | . | . | . | . | . | . | . | . | . | . | . | . | . | T | . | . | . | . | . | . |
| . | . | . | . | . | . | . | . | T | . | . | . | . | . | . | . | . | . | . | . | . | . | . | . | . | . |
| . | . | G | . | . | . | . | . | . | . | . | . | . | . | . | . | . | . | . | T | . | . | . | . | . | . |
| . | . | G | . | . | . | . | . | . | . | . | . | . | . | . | . | . | . | . | . | . | . | . | . | . | . |
| . | . | . | . | . | . | . | . | T | . | . | . | . | . | . | . | . | . | . | . | . | . | . | . | . | . |
| c | g | g | g | c | g | t | g | c | t | g | t | g | g | a | g | t | c | c | g | a | c | g | c | g | a |
| c | g | g | g | c | g | t | g | c | t | g | t | g | g | a | g | t | c | c | g | a | c | g | c | g | a |
| c | g | g | g | c | g | t | g | c | t | g | t | g | g | a | g | t | c | c | g | a | c | g | c | g | a |
| c | g | g | g | c | g | t | g | c | t | g | t | g | g | a | g | t | c | c | g | a | c | g | c | g | a |
| c | g | g | t | c | g | c | g | c | t | g | t | g | g | a | g | t | c | c | g | a | c | g | c | g | a |
| c | g | g | g | c | g | c | g | c | t | g | t | g | g | a | g | t | c | c | g | a | c | g | c | g | a |
| c | g | g | g | c | g | t | g | c | t | g | t | g | g | a | g | t | c | c | g | a | c | g | c | g | a |
| c | g | g | g | c | g | c | g | c | t | g | t | g | g | a | g | t | c | c | g | a | c | g | c | g | a |
| c | g | g | g | c | g | t | g | c | t | g | t | g | g | a | g | t | c | c | g | a | c | g | c | g | a |
| c | g | a | g | c | g | t | g | c | t | g | t | g | g | a | g | t | c | c | g | a | c | g | c | g | a |
| c | g | a | g | c | g | t | g | c | t | g | t | g | g | a | g | t | c | c | g | a | c | g | c | g | a |
| c | g | a | g | c | g | t | g | c | t | g | t | g | g | a | g | t | c | c | g | a | c | g | c | g | a |
| c | g | a | g | c | g | t | g | c | t | g | t | g | g | a | g | t | c | c | g | a | c | g | c | g | a |
| c | g | g | g | c | g | t | g | c | t | g | t | g | g | a | g | t | c | c | g | a | c | g | c | g | a |
| c | g | a | g | c | g | t | g | t | t | g | t | g | g | a | g | t | c | c | g | a | c | g | c | g | a |
| c | g | a | g | c | g | t | g | t | t | g | t | g | g | a | g | t | c | c | g | a | c | g | c | g | a |
| c | g | g | c | c | g | t | g | c | t | g | t | g | g | a | g | t | c | c | g | a | c | g | c | g | a |
| c | g | g | g | c | g | t | g | c | t | g | t | g | g | a | g | t | c | c | g | a | c | g | c | g | a |
| c | g | g | g | c | g | t | g | c | t | g | t | g | g | a | g | t | c | c | g | a | c | g | c | g | a |
| c | g | a | g | c | g | t | g | c | t | g | t | g | g | a | g | t | c | c | g | g | c | g | c | g | a |
| c | g | g | g | c | g | t | g | c | t | g | t | g | g | a | g | t | c | c | g | a | c | g | c | g | a |
| c | g | a | t | c | g | t | g | t | t | g | t | g | g | a | g | t | c | c | g | a | c | g | c | g | a |
| c | g | g | g | c | g | t | g | c | t | g | t | g | g | a | g | t | c | c | g | a | c | g | c | g | a |
| c | g | g | g | c | g | t | g | c | t | g | t | g | g | a | g | t | c | c | g | a | c | g | c | g | a |
| . | . | G | . | . | . | . | . | . | . | . | . | . | . | . | . | . | . | . | . | . | . | . | . | . | . |
| . | . | G | . | . | . | . | . | . | . | . | . | . | . | . | . | . | . | . | . | . | . | . | . | . | . |
| . | . | G | . | . | . | . | . | . | . | . | . | T | . | . | . | . | . | . | . | . | . | . | . | . | . |
| . | . | . | . | . | . | . | . | T | . | . | . | . | . | . | . | . | . | . | . | . | . | . | . | . | . |
| . | . | G | . | . | . | . | . | . | . | . | . | . | . | . | . | . | . | . | . | . | . | . | . | . | . |

| | | | | | | | | | | | | | | | | | | | | | | | | | |
|---|---|---|---|---|---|---|---|---|---|---|---|---|---|---|---|---|---|---|---|---|---|---|---|---|---|
| . | . | . | . | . | . | . | . | T | . | . | . | . | . | . | . | . | . | . | . | . | . | . | . | . | . |
| . | . | G | . | . | . | . | . | . | . | . | . | . | . | . | . | . | . | . | . | . | . | . | . | . | . |
| . | . | . | . | . | . | . | . | . | . | . | . | . | . | . | . | . | . | . | . | G | . | . | . | . | . |
| . | . | . | . | . | . | . | . | . | . | . | . | . | . | . | . | . | . | . | . | G | . | . | . | . | . |
| . | . | G | . | . | . | . | . | . | . | . | . | . | . | . | . | . | . | . | . | . | . | . | . | . | . |
| . | . | . | . | . | . | . | . | . | . | . | A | . | . | . | . | . | . | . | . | . | . | . | . | . | . |
| . | . | G | . | . | . | . | . | . | . | . | . | . | . | . | . | . | . | . | . | . | . | . | . | . | . |
| . | . | . | . | . | . | . | . | . | . | . | . | . | . | . | . | . | . | . | . | . | . | . | . | . | . |
| . | . | G | . | . | . | . | . | . | . | . | . | . | . | . | . | . | . | . | . | . | . | . | . | . | . |
| . | . | G | . | . | . | . | . | . | . | . | . | . | . | . | . | . | . | . | . | . | . | . | . | . | . |
| . | . | . | . | . | . | . | . | . | . | . | . | . | . | . | . | . | . | . | . | . | . | . | . | . | . |
| . | . | . | . | . | . | . | . | . | . | . | . | . | . | . | . | . | . | . | . | . | . | . | . | . | . |
| . | . | . | . | . | . | . | . | . | . | . | . | . | . | . | . | . | . | . | . | . | . | . | . | . | . |
| . | . | . | . | . | . | . | . | T | . | . | . | . | . | . | . | . | . | . | . | . | . | . | . | . | . |
| . | . | . | . | . | . | . | . | T | . | . | . | . | . | . | . | . | . | . | . | . | . | . | . | . | . |
| . | . | . | . | . | . | . | . | T | . | . | . | . | . | . | . | . | . | . | . | . | . | . | . | . | . |
| . | . | . | . | . | . | . | . | T | . | . | . | . | . | . | . | . | . | . | . | . | . | . | . | . | . |
| . | . | . | . | . | . | . | . | T | . | . | . | . | . | . | . | . | . | . | . | . | . | . | . | . | . |
| . | . | . | . | . | . | . | . | T | . | . | . | . | . | . | . | . | . | . | . | . | . | . | . | . | . |
| . | . | . | . | . | . | . | . | . | . | . | . | . | . | . | . | . | . | . | . | . | . | . | . | T | . |
| . | . | . | . | . | . | . | . | T | . | . | . | . | . | . | . | . | . | . | . | . | . | . | . | . | . |
| . | . | . | . | . | . | . | . | T | . | . | . | . | . | . | . | . | . | . | . | . | . | . | . | . | . |
| . | . | G | . | . | . | . | . | . | . | . | . | . | . | . | . | . | . | . | . | . | . | . | . | . | . |
| . | . | G | . | . | . | . | . | . | . | . | . | . | . | . | . | . | . | . | . | . | . | . | . | . | . |
| . | . | G | . | . | . | . | . | . | . | . | A | . | . | . | . | . | . | . | . | . | . | . | . | . | . |
| . | . | G | . | . | . | . | . | . | . | . | . | . | . | . | . | . | . | . | . | . | . | . | . | . | . |
| . | . | G | . | . | . | . | . | . | . | . | . | . | . | . | . | . | . | . | . | . | . | . | . | . | . |
| . | . | G | . | . | . | . | . | . | . | . | . | . | . | . | . | . | . | . | . | . | . | . | . | . | . |
| . | . | G | . | . | . | . | . | . | . | . | . | . | . | . | . | . | . | . | . | . | . | . | . | . | . |
| . | . | . | . | . | . | . | . | T | . | . | . | . | . | . | . | . | . | . | . | . | . | . | . | . | . |
| . | . | G | . | . | . | . | . | . | . | . | . | . | . | . | . | . | . | . | . | . | . | . | . | . | . |
| . | . | G | . | . | . | . | . | . | . | . | . | . | . | . | . | . | . | . | . | . | . | . | . | . | . |
| . | . | . | . | . | . | . | . | T | . | . | . | . | . | . | . | . | . | . | . | . | . | . | . | . | . |
| . | . | . | . | . | . | . | . | . | . | . | . | . | . | . | . | . | . | . | . | . | . | . | . | . | . |
| . | . | . | . | . | . | . | . | T | . | . | . | . | . | . | . | . | . | . | . | . | . | . | . | . | . |
| . | . | . | . | . | . | . | . | T | . | . | . | . | . | . | . | . | . | . | . | . | . | . | . | . | . |
| . | . | . | . | . | . | . | . | T | . | . | . | . | . | . | . | . | . | . | . | . | . | . | . | . | . |
| . | . | . | . | . | . | . | . | T | . | . | . | . | . | . | . | . | . | . | . | . | . | . | . | . | . |
| . | . | . | . | . | . | . | . | T | . | . | . | . | . | . | . | . | . | . | . | . | . | . | . | . | . |
| . | . | . | . | . | . | . | . | T | . | . | . | . | . | . | . | . | . | . | . | . | . | . | . | . | . |
| . | . | . | . | . | . | . | . | . | . | . | . | . | . | . | . | . | T | . | . | . | . | . | . | . | . |
| . | . | . | . | . | . | . | . | . | . | . | . | . | . | . | . | . | T | . | . | . | . | . | . | . | . |
| . | . | G | . | . | . | . | . | . | . | . | . | . | . | . | . | . | . | . | T | . | . | . | . | . | . |
| . | . | G | . | . | . | . | . | . | . | . | . | . | . | . | . | . | . | . | . | . | . | . | . | . | . |
| . | . | G | . | . | . | . | . | . | . | . | . | . | . | . | . | . | . | . | . | . | . | . | . | . | . |
| . | . | G | . | . | . | . | . | . | . | . | . | . | . | . | . | . | . | . | . | . | . | . | . | . | . |
| . | . | G | . | . | . | . | . | . | . | . | . | . | . | . | . | . | . | . | T | . | . | . | . | . | . |
| . | . | G | . | . | . | . | . | . | . | . | . | . | . | . | . | . | . | . | . | . | . | . | . | . | . |
| . | . | G | . | . | . | . | . | . | . | . | . | . | . | . | . | . | . | . | T | . | . | . | . | . | . |
| . | . | G | . | . | . | . | . | . | . | . | . | . | . | . | . | . | . | . | . | . | . | . | . | . | . |
| . | . | G | . | . | . | . | . | . | . | . | . | . | . | . | . | . | . | . | . | . | . | . | . | . | . |
| . | . | G | . | . | . | . | . | . | . | . | . | . | . | . | . | . | . | . | . | . | . | . | . | . | . |
| . | . | G | . | . | . | . | . | . | . | . | . | . | . | . | . | . | . | . | T | . | . | . | . | . | . |
| . | . | G | . | . | . | . | . | . | . | . | . | . | . | . | . | . | . | . | T | . | . | . | . | . | . |
| . | . | G | . | . | . | . | . | . | . | . | . | . | . | . | . | . | . | . | . | . | . | . | . | . | . |
| . | . | G | . | . | . | . | . | . | . | . | . | . | . | . | . | . | . | . | . | . | . | . | . | . | . |
| . | . | G | . | . | . | . | . | . | . | . | . | . | . | . | . | . | . | . | . | . | . | . | . | . | . |
| . | . | . | . | . | . | . | . | . | . | . | . | . | . | . | . | . | . | . | . | . | . | . | . | . | . |
| . | . | G | . | . | . | . | . | . | . | . | . | . | . | . | . | . | . | . | . | . | . | . | . | . | . |
| . | . | G | . | . | . | . | . | . | . | . | . | . | . | . | . | . | . | . | . | . | . | . | . | . | . |
| . | . | G | . | . | . | . | . | . | . | . | . | . | . | . | . | . | . | . | . | . | . | . | . | . | . |
| . | . | . | . | . | . | . | . | . | . | . | . | . | . | . | . | . | . | . | . | . | . | . | . | . | . |

| | | | | | | | | | | | | | | | | | | | | | | | | | |
|---|---|---|---|---|---|---|---|---|---|---|---|---|---|---|---|---|---|---|---|---|---|---|---|---|---|
| . | . | G | . | . | . | . | . | . | . | . | . | . | . | . | . | . | . | . | . | . | . | . | . | . | . |
| . | . | G | . | . | . | . | . | . | . | . | . | . | . | . | . | . | . | . | . | . | . | . | . | . | . |
| . | . | G | . | . | . | . | . | . | . | . | . | . | . | . | . | . | . | . | . | . | . | . | . | . | . |
| . | . | G | . | . | . | . | A | . | . | . | . | . | . | . | . | . | . | . | . | . | . | . | . | . | . |
| . | . | G | . | . | . | . | . | . | . | . | . | . | . | . | . | . | . | . | . | . | . | . | . | . | . |
| . | . | . | . | . | . | . | . | T | . | . | . | . | . | . | . | . | . | . | . | . | . | . | . | . | . |
| . | . | . | . | . | . | . | . | . | . | . | . | . | . | . | . | . | . | . | . | . | . | . | . | . | . |
| . | . | G | . | . | . | . | . | . | . | . | . | . | . | . | . | . | . | . | . | . | . | . | . | . | . |
| . | . | G | . | . | . | . | . | . | . | . | . | . | . | . | . | . | . | . | . | . | . | . | . | . | . |
| . | . | G | . | . | . | . | . | . | . | . | . | . | . | . | . | . | . | . | . | . | . | . | . | . | . |
| . | . | G | . | . | . | . | . | . | . | . | . | . | . | . | . | . | . | . | . | . | . | . | . | . | . |
| . | . | G | . | . | . | . | . | . | . | . | . | . | . | . | . | . | . | . | . | . | . | . | . | . | . |
| . | . | G | . | . | . | . | . | . | . | . | . | . | . | . | . | . | . | . | . | . | . | . | . | . | . |
| . | . | G | . | . | . | . | . | . | . | . | . | . | . | . | . | . | . | . | . | . | . | . | . | . | . |
| . | . | G | . | . | . | . | . | . | . | . | . | . | . | . | . | . | . | . | . | . | . | . | . | . | . |
| . | . | . | . | . | . | . | . | . | . | . | . | . | . | . | . | . | . | . | . | . | . | . | . | . | . |
| . | . | G | . | . | . | . | . | . | . | . | . | . | . | . | . | . | . | . | . | . | . | . | . | . | . |
| . | . | G | . | . | . | . | . | . | . | . | . | . | . | . | . | . | . | . | . | . | . | . | . | . | . |
| . | . | G | . | . | . | . | . | . | . | . | . | . | . | . | . | . | . | . | . | . | . | . | . | . | . |
| . | . | G | . | . | . | . | A | . | . | . | . | . | . | . | . | . | . | . | . | . | . | . | . | . | . |
| . | . | G | . | . | . | . | . | . | . | . | . | . | . | . | . | . | . | . | . | . | . | . | . | . | . |
| . | . | G | . | . | . | . | . | . | . | . | . | . | . | . | . | . | . | . | . | . | . | . | . | . | . |
| . | . | G | . | . | . | . | . | . | . | . | . | . | . | . | . | . | . | . | . | . | . | . | . | . | . |
| . | . | . | . | . | . | . | . | . | . | . | . | . | . | C | . | . | . | . | . | . | . | . | . | . | . |
| . | . | . | . | . | . | . | . | . | . | . | . | . | . | . | . | . | . | . | . | . | . | . | . | . | . |
| . | . | . | . | . | . | . | . | . | . | . | . | . | . | C | . | . | . | . | . | . | . | . | . | . | . |
| . | . | G | . | . | . | . | . | . | . | . | . | . | . | . | . | . | . | . | . | . | . | . | . | . | . |
| . | . | G | . | . | . | . | . | . | . | . | . | . | . | . | . | . | . | . | . | . | . | . | . | . | . |
| . | . | G | . | . | . | . | . | . | . | . | . | . | . | . | . | . | . | . | . | . | . | . | . | . | . |
| . | . | . | . | . | . | . | . | . | . | . | . | . | . | . | . | . | . | . | . | . | . | . | . | . | . |
| . | . | . | . | . | . | . | . | . | . | . | . | . | . | . | . | . | . | . | . | . | . | . | . | . | . |
| . | . | . | . | . | . | . | . | . | . | . | . | . | . | . | . | . | . | . | . | . | . | . | . | . | . |
| . | . | . | . | . | . | . | . | . | . | . | . | . | . | . | . | . | . | . | . | . | . | . | . | . | . |
| . | . | . | . | . | . | . | . | . | . | . | . | . | . | . | . | . | . | . | . | . | . | . | . | . | . |
| . | . | G | . | . | . | . | . | . | . | . | . | . | . | . | . | . | . | . | . | . | . | . | . | . | . |
| . | . | . | . | . | . | . | . | . | . | . | . | . | . | . | . | . | . | . | . | . | . | . | . | . | . |
| . | . | . | . | . | . | . | . | . | . | . | . | . | . | . | . | . | . | . | . | . | . | . | . | . | . |
| . | . | . | . | . | . | . | . | T | . | . | . | . | . | . | . | . | . | . | . | . | . | . | . | . | . |
| . | . | . | . | . | . | . | . | . | . | . | . | . | . | . | . | . | . | . | . | . | . | . | . | . | . |
| . | . | G | . | . | . | . | . | . | . | . | . | . | . | C | . | . | . | . | . | . | . | . | . | . | . |
| . | . | G | . | . | . | . | . | . | . | . | . | . | . | . | . | . | . | . | T | . | . | . | . | . | . |
| . | . | G | . | . | . | . | . | . | . | . | . | . | . | . | . | . | . | . | T | . | . | . | . | . | . |
| . | . | G | . | . | . | . | . | . | . | . | . | . | . | . | . | . | . | . | T | . | . | . | . | . | . |
| . | . | G | . | . | . | . | . | . | . | . | . | . | . | . | . | . | . | . | T | . | . | . | . | . | . |
| . | . | G | . | . | . | . | . | . | . | . | . | . | . | . | . | . | . | . | T | . | . | . | . | . | . |
| . | . | G | . | . | . | . | . | . | . | . | . | . | . | . | . | . | . | . | T | . | . | . | . | . | . |
| . | . | G | . | . | . | . | . | . | . | . | . | . | . | . | . | . | . | . | T | . | . | . | . | . | . |
| . | . | G | . | . | . | . | . | . | . | . | . | . | . | . | . | . | . | . | T | . | . | . | . | . | . |
| . | . | G | . | . | . | . | . | . | . | . | . | . | . | . | . | . | . | . | T | . | . | . | . | . | . |
| . | . | G | . | . | . | . | . | . | . | . | . | . | . | . | . | . | . | . | . | . | . | . | . | . | . |
| . | . | G | . | . | . | . | . | . | . | . | . | . | . | . | . | . | . | . | . | . | . | . | . | . | . |
| . | . | G | . | . | . | . | . | . | . | . | . | . | . | . | . | . | . | . | . | . | . | . | . | . | . |
| . | . | G | . | . | . | . | . | . | . | . | . | . | . | . | . | . | . | . | . | . | . | . | . | . | . |
| . | . | G | . | . | . | . | . | . | . | . | . | . | . | . | . | . | . | . | . | . | . | . | . | . | . |
| . | . | G | . | . | . | . | . | . | . | . | . | . | . | . | . | . | . | . | . | . | . | . | . | . | . |
| . | . | G | . | . | . | . | . | . | . | . | . | . | . | . | . | . | . | . | . | . | . | . | . | . | . |
| . | . | . | . | . | . | . | . | T | . | . | . | . | . | . | . | . | . | . | . | . | . | . | . | . | . |
| . | . | G | . | . | . | . | . | . | . | . | . | . | . | . | . | . | . | . | . | . | . | . | . | . | . |
| . | . | G | . | . | . | . | . | . | . | . | . | . | . | . | . | . | . | . | . | . | . | . | . | . | . |
| . | . | G | . | . | . | . | . | . | . | . | . | . | . | . | . | . | . | . | . | . | . | . | . | . | . |
| . | . | G | . | . | . | . | . | . | . | . | . | . | . | . | . | . | . | . | . | . | . | . | . | . | . |
| . | . | G | . | . | . | . | . | . | . | . | . | . | . | . | . | . | . | . | . | . | . | . | . | . | . |

| | | | | | | | | | | | | | | | | | | | | | | | | | |
|---|---|---|---|---|---|---|---|---|---|---|---|---|---|---|---|---|---|---|---|---|---|---|---|---|---|
| . | . | G | . | . | . | . | . | . | . | . | . | . | . | . | . | . | . | . | . | . | . | . | . | . | . |
| . | . | G | . | . | . | . | . | . | . | . | . | . | . | . | . | . | . | . | . | . | . | . | . | . | . |
| . | . | G | . | . | . | . | . | . | . | . | . | . | . | . | . | . | . | . | . | . | . | . | . | . | . |
| . | . | G | . | . | . | . | . | . | . | . | . | . | . | . | . | . | . | . | . | . | . | . | . | . | . |
| . | . | G | . | . | . | . | . | . | . | . | . | . | T | . | . | . | . | T | . | . | . | . | . | . | . |
| . | . | G | . | . | . | . | . | . | . | . | . | . | T | . | . | . | . | T | . | . | . | . | . | . | . |
| . | . | G | . | . | . | . | . | . | . | . | . | . | T | . | . | . | . | T | . | . | . | . | . | . | . |
| . | . | G | . | . | . | . | . | . | . | . | . | . | T | . | . | . | . | T | . | . | . | . | . | . | . |
| . | . | G | . | . | . | . | . | . | . | . | . | . | T | . | . | . | . | T | . | . | . | . | . | . | . |
| . | . | G | . | . | . | . | . | . | . | . | . | . | T | . | . | . | . | T | . | . | . | . | . | . | . |
| . | . | G | . | . | . | . | . | . | . | . | . | . | . | . | . | . | . | . | . | . | . | . | . | . | . |
| . | . | G | . | . | . | . | . | . | . | . | . | . | . | . | . | . | . | . | . | . | . | . | . | . | . |
| . | . | G | . | . | . | . | . | . | . | . | . | . | . | . | . | . | . | . | . | . | . | . | . | . | . |
| . | . | G | . | . | . | . | . | . | . | . | . | . | . | . | . | . | . | . | . | . | . | . | . | . | . |
| . | . | G | . | . | . | . | . | . | . | . | . | . | . | . | . | . | . | . | . | . | . | . | . | . | . |
| . | . | G | . | . | . | . | . | . | . | . | . | . | . | . | . | . | . | . | . | . | . | . | . | . | . |
| . | . | G | . | . | . | . | . | . | . | . | . | . | . | . | . | . | . | . | . | . | . | . | . | . | . |
| . | . | G | . | . | . | . | . | . | . | . | . | . | . | . | . | . | . | . | . | . | . | . | . | . | . |
| . | . | G | . | . | . | . | . | . | . | . | . | . | . | . | . | . | . | . | . | . | . | . | . | . | . |
| . | . | G | . | . | . | . | . | . | . | . | . | . | . | . | . | . | . | . | . | . | . | . | . | . | . |
| . | . | G | . | . | . | . | . | . | . | . | . | . | . | C | . | . | . | . | . | . | . | . | . | . | . |
| . | . | . | . | . | . | . | . | T | . | . | . | . | . | . | . | . | . | . | . | . | . | . | . | . | . |
| . | . | . | . | . | . | . | . | T | . | . | . | . | . | . | . | . | . | . | . | . | . | . | . | . | . |
| . | . | G | . | . | . | . | . | . | . | . | . | . | . | . | . | . | . | . | . | . | . | . | . | . | . |
| . | . | G | . | . | . | . | . | . | . | . | . | . | . | . | . | . | . | . | . | . | . | . | . | . | . |
| . | . | . | . | . | . | . | . | T | . | . | . | . | . | . | . | . | . | . | . | . | . | . | . | . | . |
| . | . | . | . | . | . | . | . | T | . | . | . | . | . | . | T | . | . | . | . | . | . | . | . | . | . |
| . | . | . | . | . | . | . | . | T | . | . | . | . | . | . | . | . | . | . | . | . | . | . | . | . | . |
| . | . | . | . | . | . | . | . | T | . | . | . | . | . | . | . | . | . | . | . | . | . | . | . | . | . |
| . | . | . | . | . | . | . | . | T | . | . | . | . | . | . | . | . | . | . | . | . | . | . | . | . | . |
| . | . | . | . | . | . | . | . | T | . | . | . | . | . | . | . | . | . | . | . | . | . | . | . | . | . |
| . | . | . | . | . | . | . | . | T | . | . | . | . | . | . | . | . | . | . | . | . | . | . | . | . | . |
| . | . | . | . | . | . | . | . | T | . | . | . | . | . | . | . | . | . | . | . | . | . | . | . | . | . |
| . | . | . | . | . | . | . | . | T | . | . | . | . | . | . | . | . | . | . | . | . | . | . | . | . | . |
| . | . | . | . | . | . | . | . | T | . | . | . | . | . | . | T | . | . | . | . | . | . | . | . | . | . |
| . | . | . | . | . | . | . | . | T | . | . | . | . | . | . | . | . | . | . | . | . | . | . | . | . | . |
| . | . | . | . | . | . | . | . | T | . | . | . | . | . | . | . | . | . | . | . | . | . | . | . | . | . |
| . | . | . | . | . | . | . | . | . | . | . | . | . | . | . | . | . | . | . | . | . | . | . | . | . | . |
| . | . | G | . | . | . | . | . | . | . | . | . | . | . | . | . | . | . | . | T | . | . | . | . | . | . |
| . | . | G | . | . | . | . | . | . | . | . | . | . | . | . | . | . | . | . | T | . | . | . | . | . | . |
| . | . | G | . | . | . | . | . | . | . | . | . | . | . | . | . | . | . | . | T | . | . | . | . | . | . |
| . | . | G | . | . | . | . | . | . | . | . | . | . | . | . | . | . | . | . | T | . | . | . | . | . | . |
| T | . | G | . | . | . | . | . | . | . | . | . | . | . | . | . | . | . | . | T | . | . | . | . | . | . |
| T | . | G | . | . | . | . | . | . | . | . | . | . | . | . | . | . | . | . | T | . | . | . | . | . | . |
| . | . | G | . | . | . | . | . | . | . | . | . | . | . | . | . | . | . | . | T | . | . | . | . | . | . |
| . | . | G | . | . | . | . | . | . | . | . | . | . | . | . | . | . | . | . | . | . | . | . | . | . | . |
| . | . | G | . | . | . | . | . | . | . | . | . | . | . | . | . | . | . | . | T | . | . | . | . | . | . |
| . | . | G | . | . | . | . | . | . | . | . | . | . | . | . | . | . | . | . | T | . | . | . | . | . | . |
| . | . | G | . | . | . | . | . | . | . | . | . | . | . | . | . | . | . | . | . | . | . | . | . | . | . |
| . | . | G | . | . | . | . | . | . | . | . | . | . | . | . | . | . | . | . | T | . | . | . | . | . | . |
| . | . | G | . | . | . | . | . | . | . | . | . | . | . | . | . | . | . | . | T | . | . | . | . | . | . |
| . | . | G | . | . | . | . | . | . | . | . | . | . | . | . | . | . | . | . | T | . | . | . | . | . | . |
| . | . | G | . | . | . | . | . | . | . | . | . | . | . | . | . | . | . | . | T | . | . | . | . | . | . |
| . | . | G | . | . | . | . | . | . | . | . | . | . | . | . | . | . | . | . | T | . | . | . | . | . | . |
| . | . | G | . | . | . | . | . | . | . | . | . | . | . | . | . | . | . | . | T | . | . | . | . | . | . |
| T | . | G | . | . | . | . | . | . | . | . | . | . | . | . | . | . | . | . | T | . | . | . | . | . | . |
| . | . | G | . | . | . | . | . | . | . | . | . | . | . | . | . | . | . | . | . | . | . | . | . | . | . |
| . | . | G | . | . | . | . | . | . | . | . | . | . | . | . | . | . | . | . | . | . | . | . | . | . | . |
| . | . | G | . | . | . | . | . | . | . | . | . | . | . | . | . | . | . | . | . | . | . | . | . | . | . |
| . | . | G | . | . | . | . | . | . | . | . | . | . | . | . | . | . | . | . | . | . | . | . | . | . | . |
| . | . | G | . | . | . | . | . | . | . | . | . | . | . | . | . | . | . | . | . | . | . | . | . | . | . |
| . | . | G | . | . | . | . | . | . | . | . | . | . | . | . | . | . | . | . | . | . | . | . | . | . | . |

| | | | | | | | | | | | | | | | | | | | | | | | | | |
|---|---|---|---|---|---|---|---|---|---|---|---|---|---|---|---|---|---|---|---|---|---|---|---|---|---|
| . | . | . | . | . | . | . | . | . | . | . | . | . | . | . | . | . | . | . | T | . | . | . | . | . | . |
| . | . | G | . | . | . | . | . | . | . | . | . | . | . | . | . | . | . | . | T | . | . | . | . | . | . |
| . | . | G | . | . | . | . | . | . | . | . | . | . | . | . | . | . | . | . | T | . | . | . | . | . | . |
| . | . | G | . | . | . | . | . | . | . | . | . | . | . | . | . | . | . | . | T | . | . | . | . | . | . |
| . | . | . | . | . | . | . | . | . | . | . | . | . | . | . | . | . | . | . | . | . | . | . | . | . | . |
| . | . | . | . | . | . | . | . | T | . | . | . | . | . | M | . | . | . | . | . | . | . | . | . | . | . |
| . | . | R | . | . | . | . | . | . | . | . | . | . | . | . | . | . | . | . | . | . | . | . | . | . | . |
| . | . | R | . | . | . | . | . | . | . | . | . | . | . | . | . | . | . | . | . | . | . | . | . | . | . |
| . | . | . | . | . | . | . | . | T | . | . | . | . | . | . | . | . | . | . | . | . | . | . | . | . | . |
| . | . | . | . | . | . | . | . | Y | . | . | . | . | . | . | . | . | . | . | . | . | . | . | . | . | . |
| . | . | . | . | . | . | . | . | T | . | . | . | . | . | . | . | . | . | . | . | . | . | . | . | . | . |
| . | . | . | . | . | . | . | . | T | . | . | . | . | . | . | . | . | . | . | . | . | . | . | . | . | . |
| . | . | . | . | . | . | . | . | T | . | . | . | . | . | . | . | . | . | . | . | . | . | . | . | . | . |
| . | . | . | . | . | . | . | . | T | . | . | . | . | . | . | . | . | . | . | . | . | . | . | . | . | . |
| . | . | . | . | . | . | . | . | T | . | . | . | . | . | . | . | . | . | . | . | . | . | . | . | . | . |
| . | . | . | . | . | . | . | . | T | . | . | . | . | . | . | . | . | . | . | . | . | . | . | . | . | . |
| . | . | . | . | . | . | . | . | T | . | . | . | . | . | . | . | . | . | . | . | . | . | . | . | . | . |
| . | . | R | . | . | . | . | . | Y | . | . | . | . | . | ? | . | . | . | . | . | . | . | . | . | . | . |
| . | . | . | . | . | . | . | . | T | . | . | . | . | . | . | . | . | . | . | . | . | . | . | . | . | . |
| . | . | . | . | . | . | . | . | T | . | . | . | . | . | . | . | . | . | . | . | . | . | . | . | . | . |
| . | . | . | . | . | . | . | . | T | . | . | . | . | . | . | . | . | . | . | . | . | . | . | . | . | . |
| . | . | . | . | . | . | . | . | T | . | . | . | . | . | . | . | . | . | . | . | . | . | . | . | . | . |
| . | . | . | . | . | . | . | . | T | . | . | . | . | . | . | . | . | . | . | . | . | . | . | . | . | . |
| . | . | . | . | . | . | . | . | T | . | . | . | . | . | . | . | . | . | . | . | . | . | . | . | . | . |
| . | . | . | . | . | . | . | . | T | . | . | . | . | . | . | . | . | . | . | . | . | . | . | . | . | . |
| . | . | . | . | . | . | . | . | T | . | . | . | . | . | . | . | . | . | . | . | . | . | . | . | . | . |
| . | . | . | . | . | . | . | . | T | . | . | . | . | . | . | . | . | . | . | . | . | . | . | . | . | . |
| . | . | . | . | . | . | . | . | T | . | . | . | . | . | . | . | . | . | . | . | . | . | . | . | . | . |
| . | . | . | . | . | . | . | . | T | . | . | . | . | . | . | . | . | . | . | . | . | . | . | . | . | . |
| . | . | . | . | . | . | . | . | T | . | . | . | . | . | . | . | . | . | . | . | . | . | . | . | . | . |
| . | . | . | . | . | . | . | . | T | . | . | . | . | . | . | . | . | . | . | . | . | . | . | . | . | . |
| . | . | G | . | . | . | . | . | . | . | . | . | . | . | . | . | . | . | . | . | . | . | . | . | . | . |
| . | . | G | . | . | . | . | . | . | . | . | . | . | . | . | . | . | . | . | . | . | . | . | . | . | . |
| . | . | G | . | . | . | . | . | . | . | . | . | . | . | . | . | . | . | . | T | . | . | . | . | . | . |
| . | . | G | . | . | . | . | . | . | . | . | . | . | . | . | . | . | . | . | . | . | . | . | . | . | . |
| . | . | . | . | . | . | . | . | . | . | . | . | . | . | . | . | . | . | . | . | . | . | . | . | . | . |
| . | . | G | . | . | . | . | . | . | . | . | . | . | . | . | . | . | . | . | . | . | . | . | . | . | . |
| T | . | G | . | . | . | . | . | . | . | . | . | . | . | . | . | . | . | . | T | . | . | . | . | . | . |
| . | T | G | . | . | . | . | . | . | . | . | . | . | . | . | . | . | . | . | T | . | . | . | . | . | . |
| . | . | G | . | . | . | . | . | . | . | . | . | . | . | . | . | . | . | . | . | . | . | . | . | . | . |
| . | . | G | . | . | . | . | . | . | . | . | . | . | . | . | . | . | . | . | . | . | . | . | . | . | . |
| . | . | G | . | . | . | . | . | . | . | . | . | . | . | . | . | . | . | . | T | . | . | . | . | . | . |
| . | . | G | . | . | . | . | . | . | . | . | . | . | . | . | . | . | . | . | T | . | . | . | . | . | . |
| . | . | G | . | . | . | . | . | . | . | . | . | . | . | . | . | . | . | . | . | . | . | . | . | . | . |
| . | . | G | . | . | . | . | . | . | . | . | . | . | . | . | . | . | . | . | . | . | . | . | . | . | . |
| . | . | G | . | . | . | . | . | . | . | . | . | . | . | . | . | . | . | . | T | . | . | . | . | . | . |
| . | . | G | . | . | . | . | . | . | . | . | . | . | . | . | . | . | . | . | . | . | . | . | . | . | . |
| . | . | G | . | . | . | . | . | . | . | . | . | . | . | . | . | . | . | . | T | . | . | . | . | . | . |
| . | T | G | . | . | . | . | . | . | . | . | . | . | . | . | . | . | . | . | T | . | . | . | T | . | . |
| . | . | G | . | . | . | . | . | . | . | . | . | . | . | . | . | . | . | . | . | . | . | . | . | . | . |
| . | . | G | . | . | . | . | . | . | . | . | . | . | . | . | . | . | . | . | . | . | . | . | . | . | . |
| . | . | G | . | . | . | . | . | . | . | . | . | . | . | . | . | . | . | . | . | . | . | . | . | . | . |
| . | . | G | . | . | . | . | . | . | . | . | . | . | . | . | . | . | . | . | T | . | . | . | . | . | . |
| . | . | G | . | . | . | . | . | . | . | . | . | . | . | . | . | . | . | . | T | . | . | . | . | . | . |
| . | . | G | . | . | . | . | . | . | . | . | . | . | . | . | . | . | . | . | T | . | . | . | . | . | . |
| . | . | G | . | . | . | . | . | . | . | . | . | . | . | . | . | . | . | . | T | . | . | . | . | . | . |
| . | . | G | . | . | . | . | . | . | . | . | . | . | . | . | . | . | . | . | . | . | . | . | . | . | . |
| . | . | G | . | . | . | . | . | . | . | . | . | . | . | . | . | . | . | . | T | . | . | . | . | . | . |
| . | . | G | . | . | . | . | . | . | . | . | . | . | . | . | . | . | . | . | . | . | . | . | . | . | . |
| . | . | G | . | . | . | . | . | . | . | . | . | . | . | . | . | . | . | . | T | . | . | . | . | . | . |

| 212 | 213 | 214 | 215 | 216 | 217 | 218 | 219 | 220 | 221 | 222 | 223 | 224 | 225 | 226 | 227 | 228 | 229 | 230 | 231 | 232 | 233 | 234 | 235 | 236 | 237 |
|---|---|---|---|---|---|---|---|---|---|---|---|---|---|---|---|---|---|---|---|---|---|---|---|---|---|
| E | E | E | E | E | M | M | M | M | M | M | ORF6 | ORF6 | ORF6 | ORF6 | ORF6 | ORF7a | ORF7b | NC | ORF8 | ORF8 | ORF8 | ORF8 | ORF8 | ORF8 | ORF8 |
| 26330 | 26338 | 26375 | 26376 | 26467 | 26530 | 26681 | 26730 | 26735 | 27110 | 27191 | 27213 | 27379 | 27382 | 27383 | 27384 | 27613 | 27874 | 27889 | 28001 | 28077 | 28083 | 28144 | 28221 | 28253 | 28254 |
| T | G | G | C | G | A | C | G | C | C | A | C | A | G | A | T | G | C | C | G | G | G | T | G | C | A |
| . | . | . | . | . | . | . | . | . | . | . | . | . | . | . | . | . | . | . | . | . | . | . | . | . | . |
| . | . | . | . | . | . | . | . | . | . | . | . | . | . | . | . | . | . | . | . | . | . | C | . | . | . |
| . | . | . | . | . | . | . | . | . | . | . | . | . | . | . | . | . | . | . | . | . | . | . | . | . | . |
| . | . | . | . | . | . | . | . | . | . | . | . | . | . | . | . | . | . | . | . | . | . | . | . | . | . |
| . | . | . | . | . | . | . | . | . | . | . | . | . | . | . | . | . | . | . | . | . | . | . | . | . | . |
| . | . | . | . | . | . | . | . | . | . | . | . | . | . | . | . | . | . | . | . | . | . | . | . | . | . |
| . | . | . | . | . | . | . | . | . | . | . | . | . | . | . | . | . | . | . | . | . | . | . | . | . | . |
| . | . | . | . | . | . | . | . | . | . | . | . | . | . | . | . | . | . | . | . | . | . | . | . | . | . |
| . | . | . | . | . | . | . | . | . | . | . | . | . | . | . | . | . | . | . | . | . | . | . | . | . | . |
| . | . | . | . | . | . | . | . | . | . | . | . | . | . | . | . | . | . | . | . | . | . | . | . | . | . |
| . | . | . | . | . | . | . | . | . | . | . | . | . | . | . | . | . | . | . | . | . | . | . | . | . | . |
| . | . | . | . | . | . | . | . | . | . | . | . | . | . | . | . | . | . | . | . | . | . | . | . | . | . |
| . | . | . | . | . | . | . | . | . | . | . | . | . | . | . | . | . | . | . | . | . | . | . | . | . | . |
| . | . | . | . | . | . | . | . | . | . | . | . | . | . | . | . | . | . | . | . | . | . | . | . | . | . |
| . | . | . | . | . | . | . | . | . | . | . | . | . | . | . | . | . | . | . | . | . | . | . | . | . | . |
| . | . | . | . | . | . | . | . | . | . | . | . | . | . | . | . | . | . | . | . | . | . | . | . | . | . |
| . | . | . | . | . | . | . | . | . | . | . | . | . | . | . | . | T | . | . | . | . | . | . | . | . | . |
| . | . | . | . | . | . | . | . | . | . | . | . | . | . | . | . | T | . | . | . | . | . | . | . | . | . |
| . | . | . | . | . | . | . | . | . | . | . | . | . | . | . | . | T | . | . | . | . | . | . | . | . | . |
| . | . | . | . | . | . | . | . | . | . | . | . | . | . | . | . | . | . | . | . | . | . | . | . | . | . |
| . | . | . | . | . | . | . | . | . | . | . | . | . | . | . | . | . | . | . | . | . | . | . | . | . | . |
| . | . | . | . | . | . | . | . | . | . | . | . | . | . | . | . | T | . | . | . | . | . | . | . | . | . |
| . | . | . | . | . | . | . | . | . | . | . | T | . | . | . | . | . | . | . | . | . | . | . | . | . | . |
| . | . | . | . | . | . | . | . | . | . | . | . | . | . | . | . | . | . | . | . | . | . | . | . | . | . |
| . | . | . | . | . | . | . | . | . | . | . | . | . | . | . | . | . | . | . | . | . | . | . | . | A | C |
| . | . | . | . | . | . | . | . | . | . | . | . | . | . | . | . | . | . | . | . | . | . | . | . | . | . |
| . | . | . | . | . | . | . | . | . | . | . | . | . | . | . | . | . | . | . | . | . | . | . | . | . | . |
| . | . | . | . | . | . | . | . | . | . | . | . | . | . | . | . | T | . | . | . | . | . | . | . | . | . |
| . | . | . | . | . | . | . | . | . | . | . | . | . | . | . | . | . | . | . | . | . | . | . | . | . | . |
| . | . | . | . | . | . | . | . | . | . | . | . | . | . | . | . | . | . | . | . | . | . | . | . | . | . |
| . | . | . | . | . | G | . | . | . | . | . | . | . | . | . | . | . | . | . | . | . | . | . | . | . | . |
| . | . | . | . | . | G | . | . | . | . | . | . | . | . | . | . | . | . | . | . | . | . | . | . | . | . |
| . | . | . | . | . | . | . | . | . | . | . | . | . | . | . | . | . | . | . | . | . | . | . | . | . | . |
| . | . | . | . | . | . | . | . | . | . | . | . | . | . | . | . | . | . | . | . | . | . | . | . | . | . |
| . | . | . | . | . | . | . | . | . | . | . | . | . | . | . | . | . | . | . | . | . | . | . | . | . | . |
| . | . | . | . | T | . | . | . | . | . | . | . | . | . | . | . | . | . | . | . | . | . | . | . | . | . |
| . | . | . | . | ? | . | . | . | . | . | . | . | . | . | . | . | . | ? | . | . | . | . | . | ? | ? | ? |
| . | . | . | . | T | . | . | . | . | . | . | . | . | . | . | . | . | . | . | . | . | . | . | . | . | . |
| . | . | . | . | T | . | . | . | . | . | . | . | . | . | . | . | . | . | . | . | . | . | . | . | . | . |
| . | . | . | . | T | . | . | . | . | . | . | . | . | . | . | . | . | ? | . | . | . | . | . | . | . | . |
| . | . | . | . | ? | . | . | . | . | . | . | . | . | . | . | . | . | . | . | . | . | . | . | . | . | . |
| . | . | . | . | ? | . | . | . | . | . | . | . | . | . | . | . | . | ? | ? | ? | ? | ? | ? | ? | ? | ? |
| . | . | . | . | T | . | . | . | . | . | . | . | . | . | . | . | . | . | . | . | . | . | . | . | . | . |
| . | . | . | . | T | . | . | . | . | . | . | . | . | . | . | . | . | . | . | . | . | . | . | . | . | . |
| . | . | . | . | . | . | . | . | . | . | . | . | . | . | . | . | . | . | . | . | . | . | . | . | . | . |
| . | . | . | . | . | . | . | . | . | . | . | . | . | . | . | . | . | . | . | . | . | . | . | . | . | . |
| . | . | . | . | . | . | . | . | . | . | . | . | . | . | . | . | . | . | . | . | . | . | . | . | . | . |
| . | . | . | . | . | . | . | . | . | . | . | . | . | . | . | . | . | . | . | . | . | . | . | . | . | . |
| . | . | . | . | . | . | . | . | . | . | . | . | . | . | . | . | . | . | . | . | . | . | . | . | . | . |
| . | . | . | . | . | . | . | . | . | . | . | . | . | . | . | . | . | . | . | . | . | . | . | . | . | . |
| . | . | . | . | . | . | . | . | . | . | . | . | . | . | . | . | . | . | . | . | . | . | . | . | . | . |
| . | . | . | . | . | . | . | . | . | . | . | . | . | . | . | . | . | . | . | . | . | . | . | . | . | . |
| . | . | . | . | . | . | . | . | . | . | . | . | . | . | . | . | . | . | . | . | . | . | . | . | . | . |
| . | . | . | . | . | . | . | . | . | . | . | . | . | . | . | . | . | . | . | . | . | . | . | . | . | . |
| . | . | . | . | . | . | . | . | . | . | . | . | . | . | . | . | . | . | . | . | . | . | . | . | . | . |

| | | | | | | | | | | | | | | | | | | | | | | | | | |
|---|---|---|---|---|---|---|---|---|---|---|---|---|---|---|---|---|---|---|---|---|---|---|---|---|---|
| . | . | . | . | . | . | . | . | . | . | . | . | . | . | . | . | . | . | . | . | . | . | . | T | . | . |
| . | . | . | . | . | . | . | . | . | . | . | . | . | . | . | . | . | . | . | . | . | . | . | T | . | . |
| . | . | . | . | . | . | . | . | . | . | . | . | . | . | . | . | . | . | . | . | . | . | . | . | . | . |
| . | . | . | . | . | . | . | . | . | . | . | . | . | . | . | . | . | . | . | . | . | . | . | . | . | . |
| . | . | . | . | . | . | . | . | T | . | . | . | . | . | . | . | . | . | . | . | . | . | . | . | . | . |
| . | . | . | . | . | . | . | . | . | . | . | . | . | . | . | . | . | . | . | . | . | . | . | . | . | . |
| . | . | . | . | . | . | . | . | . | . | . | . | . | . | . | . | g | . | . | . | . | . | . | . | . | . |
| . | . | . | . | . | . | . | . | . | . | . | . | . | . | . | . | g | . | . | . | . | . | . | . | . | . |
| . | . | . | . | . | . | . | . | . | . | . | . | . | . | . | . | g | . | . | . | . | . | . | . | . | . |
| . | . | . | . | . | . | . | . | . | . | . | . | . | . | . | . | . | . | . | . | . | . | . | . | . | . |
| . | . | . | . | . | . | . | . | . | . | . | . | . | . | . | . | . | . | . | . | . | . | . | . | . | . |
| . | . | . | . | . | . | . | . | T | . | . | . | . | . | . | . | . | . | . | . | . | . | . | . | . | . |
| . | . | . | . | . | . | . | . | T | . | . | . | . | . | . | . | . | . | . | . | . | . | . | . | . | . |
| . | . | . | . | . | . | . | . | T | . | . | . | . | . | . | . | . | . | . | . | . | . | . | . | . | . |
| . | . | . | . | . | . | . | . | T | . | . | . | . | . | . | . | . | . | . | . | . | . | . | . | . | . |
| . | . | . | . | . | . | . | . | T | . | . | . | . | . | . | . | . | . | . | . | . | . | . | . | . | . |
| . | . | . | . | . | . | . | . | . | . | . | . | . | . | . | . | . | . | . | . | . | . | . | . | . | . |
| . | . | . | . | . | . | . | . | . | . | . | . | . | . | . | . | . | . | . | . | . | . | . | . | . | . |
| . | . | . | . | . | . | . | . | . | . | . | . | . | . | . | . | g | . | . | . | . | . | . | . | . | . |
| . | . | . | . | . | . | . | . | . | . | . | . | . | . | . | . | . | . | . | . | . | . | . | . | . | . |
| . | . | . | . | . | . | . | . | . | . | . | . | . | . | . | . | g | . | . | . | . | . | . | . | . | . |
| . | . | . | . | . | . | . | . | . | . | . | . | . | . | . | . | ? | . | . | . | . | . | . | . | . | . |
| . | . | . | . | . | . | . | . | . | . | . | . | . | . | . | . | . | . | . | . | . | . | . | . | . | . |
| . | . | . | . | . | . | . | . | . | . | . | . | . | . | . | . | ? | . | . | . | . | . | . | . | . | . |
| . | . | . | . | . | . | . | . | . | . | . | . | . | . | . | . | g | . | . | . | . | . | . | . | . | . |
| . | . | . | . | . | . | . | . | . | . | . | . | . | . | . | . | g | . | . | . | . | . | . | . | . | . |
| . | . | . | . | . | . | . | . | . | . | . | . | . | . | . | . | g | . | . | . | . | . | . | . | . | . |
| . | . | . | . | . | . | . | . | . | . | . | . | . | . | . | . | g | . | . | . | . | . | . | . | . | . |
| . | . | . | . | . | . | . | . | . | . | . | . | . | . | . | . | g | . | . | . | . | . | . | . | . | . |
| . | . | . | . | . | . | . | . | . | . | . | . | . | . | . | . | ? | . | . | . | . | . | . | . | . | . |
| . | . | . | . | . | . | . | . | . | . | . | . | . | . | . | . | g | . | . | . | . | . | . | . | . | . |
| . | . | . | . | . | . | . | . | . | . | . | . | . | . | . | . | . | . | . | . | . | . | . | . | . | . |
| . | . | . | . | . | . | . | . | . | . | . | . | . | . | . | . | g | . | . | . | . | . | . | . | . | . |
| . | . | . | . | . | . | . | . | . | . | . | . | . | . | . | . | . | . | . | . | . | . | . | . | . | . |
| . | . | . | . | . | . | . | . | . | . | . | . | . | . | . | . | . | . | . | . | . | . | . | . | . | . |
| . | . | . | . | . | . | . | . | . | . | . | . | . | . | . | . | . | . | . | . | . | . | . | . | . | . |
| . | . | . | . | . | . | . | . | . | . | . | . | . | . | . | . | . | . | . | . | . | . | . | . | . | . |
| . | . | . | . | . | . | . | . | . | . | . | . | . | . | . | . | . | . | . | . | . | . | . | . | . | . |
| . | . | . | . | . | . | . | . | . | . | . | . | . | . | . | . | g | . | . | . | . | . | . | . | . | . |
| . | . | . | . | . | . | . | . | . | . | . | . | . | . | . | . | . | . | . | . | . | . | . | . | . | . |
| . | . | . | . | . | . | . | . | . | . | . | . | . | . | . | . | g | . | . | . | . | . | . | . | . | . |
| . | . | . | . | . | . | . | . | . | . | G | . | . | . | . | . | . | . | . | . | . | . | . | . | . | . |
| . | . | . | . | . | . | . | . | . | . | G | . | . | . | . | . | g | . | . | . | . | . | . | . | . | . |
| . | . | . | . | . | . | . | . | . | . | R | . | . | . | . | . | . | . | . | . | . | . | . | . | . | . |
| . | . | . | . | . | . | . | . | . | . | . | . | . | . | . | . | g | . | . | . | . | . | . | . | . | . |
| . | . | . | . | . | . | . | . | . | . | . | . | . | . | . | . | . | . | . | . | . | . | . | . | . | . |
| . | . | . | . | . | . | . | . | . | . | . | . | . | . | . | . | ? | . | . | . | . | . | . | . | . | . |
| . | . | . | . | . | . | . | . | . | . | . | . | . | . | . | . | g | . | . | . | . | . | . | . | . | . |
| . | . | . | . | . | . | . | . | . | . | . | . | . | . | . | . | . | . | . | . | . | . | . | . | . | . |
| . | . | . | . | . | . | . | . | . | . | . | . | . | . | . | . | ? | . | . | . | . | . | C | . | . | . |
| . | . | . | . | . | . | . | . | . | . | . | . | . | . | . | . | g | . | . | . | . | . | C | . | . | . |
| . | . | . | . | . | . | . | . | . | . | . | . | . | . | . | . | . | . | . | . | . | . | . | . | . | . |
| . | . | . | . | . | . | . | . | . | . | . | . | . | . | . | . | . | . | . | . | . | . | . | . | . | . |
| . | . | . | . | . | . | . | . | . | . | . | . | . | . | . | . | . | . | . | . | . | . | . | . | . | . |
| . | . | . | . | . | . | . | . | . | . | . | . | . | . | . | . | . | . | . | . | . | . | . | . | . | . |
| . | . | . | . | . | . | . | . | . | . | . | . | . | . | . | . | . | . | . | . | . | . | . | . | . | . |
| . | . | . | . | . | . | . | . | . | . | . | . | . | . | . | . | g | . | . | . | . | . | . | . | . | . |
| . | . | . | . | . | . | . | . | . | . | . | . | . | . | . | . | . | T | . | . | . | . | . | . | . | . |
| . | . | . | . | . | . | . | . | . | . | . | . | . | . | . | . | . | . | . | . | . | . | . | . | . | . |
| . | . | . | . | . | . | . | . | . | . | . | . | . | . | . | . | g | . | . | . | . | . | . | . | . | . |
| . | . | . | . | . | . | . | . | . | . | . | . | . | . | . | . | . | . | . | . | . | . | . | . | . | . |
| . | . | . | . | . | . | . | . | . | . | . | . | . | . | . | . | . | . | . | . | . | . | . | . | . | . |

| | | | | | | | | | | | | | | | | | | | | | | | | | |
|---|---|---|---|---|---|---|---|---|---|---|---|---|---|---|---|---|---|---|---|---|---|---|---|---|---|
| . | . | . | . | . | . | . | . | . | . | . | . | . | . | . | . | . | . | . | . | . | . | . | . | . | . |
| . | . | . | . | . | . | . | . | . | . | . | . | . | . | . | . | . | . | . | . | . | . | . | . | . | . |
| . | . | . | . | . | . | . | . | . | . | . | . | . | . | . | . | g | . | . | . | . | . | . | . | . | . |
| . | . | . | . | . | . | . | . | . | . | . | . | . | . | . | . | . | . | . | . | . | . | . | . | . | . |
| . | . | . | . | . | . | . | . | . | . | . | . | . | . | . | . | . | . | . | . | . | . | . | . | . | . |
| . | . | . | . | . | . | . | . | . | . | . | . | . | . | . | . | . | . | . | . | . | . | . | . | . | . |
| . | . | . | . | . | . | . | . | . | . | . | . | . | . | . | . | . | . | . | . | . | . | . | . | . | . |
| . | . | . | . | . | . | . | . | . | . | . | . | . | . | . | . | . | . | . | . | . | . | . | . | . | . |
| . | . | . | . | . | . | . | . | . | . | . | . | . | K | . | . | ? | . | . | . | S | . | Y | . | . | . |
| . | . | . | . | . | . | . | . | . | . | . | . | . | . | . | . | . | . | . | . | . | . | . | . | . | . |
| . | . | . | . | . | . | . | . | . | . | . | . | . | . | . | . | ? | . | . | . | . | . | . | . | . | . |
| . | . | . | . | ? | . | . | . | . | . | . | . | . | . | . | . | . | . | . | . | . | . | . | . | . | . |
| . | . | . | . | . | . | . | . | . | . | . | . | . | . | . | . | . | . | . | . | . | . | . | . | . | . |
| . | . | . | . | . | . | . | . | . | . | . | . | . | . | . | . | . | . | . | . | . | . | . | . | . | . |
| . | . | . | . | . | . | . | . | . | . | . | . | . | . | . | . | . | . | . | . | . | . | . | . | . | . |
| . | . | . | . | . | . | . | . | . | . | . | . | . | . | . | . | . | . | . | . | . | . | . | . | . | . |
| . | . | . | . | . | . | . | . | . | . | . | . | . | . | . | . | . | . | . | . | . | . | . | . | . | . |
| . | . | . | . | . | . | . | . | . | . | . | . | . | . | . | . | . | . | . | . | . | . | . | . | . | . |
| . | . | . | . | . | . | . | . | . | . | . | . | . | . | . | . | . | . | . | . | . | . | . | . | . | . |
| . | . | . | . | . | . | . | . | . | . | . | . | . | . | . | . | . | . | . | . | . | . | . | . | c | a |
| . | . | . | . | . | . | . | . | . | . | . | . | . | . | . | . | . | . | . | . | . | . | . | . | . | . |
| . | . | . | . | g | . | . | . | . | . | . | . | . | . | . | . | ? | . | . | . | . | . | . | . | . | . |
| . | . | . | . | . | . | . | . | . | . | . | . | . | . | . | . | . | . | . | . | . | . | . | . | c | a |
| . | . | . | . | . | . | . | . | . | . | . | . | . | . | . | . | . | . | . | . | . | . | . | . | . | . |
| . | . | . | . | . | . | . | . | . | . | . | . | . | . | . | . | . | . | . | . | . | . | . | . | T | . |
| . | . | . | . | . | . | . | . | . | . | . | . | . | . | . | . | . | . | . | . | . | . | . | . | . | . |
| . | . | . | . | . | . | . | . | . | . | . | . | . | . | . | . | . | . | . | . | . | . | . | . | . | . |
| . | . | . | . | . | . | . | . | T | . | . | . | . | . | . | . | . | . | . | . | . | . | . | . | . | . |
| . | . | . | . | . | . | . | . | . | . | . | . | . | . | . | . | . | . | . | . | . | . | . | . | . | . |
| . | . | . | . | . | . | . | . | . | . | . | . | . | . | . | . | . | . | . | . | . | . | . | . | . | . |
| . | . | . | . | . | . | . | . | . | . | . | . | . | . | . | . | . | . | . | . | . | . | . | . | . | . |
| . | . | . | . | . | . | . | . | . | . | . | . | . | . | . | . | g | . | . | . | . | . | . | . | . | . |
| . | . | . | . | . | . | . | . | . | . | . | . | . | . | . | . | . | . | . | . | . | . | . | . | . | . |
| . | . | . | . | . | . | . | . | T | . | . | . | . | . | . | . | . | . | . | . | . | . | . | . | . | . |
| . | . | . | . | . | . | . | . | . | . | . | . | . | . | . | . | . | . | . | . | . | . | C | . | . | . |
| . | . | . | . | . | . | . | . | . | . | . | . | . | . | . | . | . | . | . | . | . | . | . | . | . | . |
| . | . | . | . | . | . | . | . | . | . | . | . | . | . | . | . | . | . | . | . | . | . | . | . | . | . |
| . | . | . | . | . | . | . | . | T | . | . | . | . | . | . | . | . | . | . | . | . | . | . | . | . | . |
| . | . | . | . | . | . | . | . | T | . | . | . | . | . | . | . | . | . | . | . | . | . | . | . | . | . |
| . | . | . | . | . | . | . | . | . | . | . | . | . | . | . | . | . | . | . | . | . | . | . | . | . | . |
| . | . | . | . | . | . | . | . | . | . | . | . | . | . | . | . | . | . | . | . | . | . | . | . | . | . |
| . | . | . | . | . | . | . | . | T | . | . | . | . | . | . | . | . | . | . | . | . | . | . | . | . | . |
| . | . | . | . | . | . | . | . | . | . | . | . | . | . | . | . | . | . | . | . | . | . | . | . | . | . |
| . | . | . | . | . | . | . | . | . | . | . | . | . | . | . | . | . | . | . | . | . | . | . | . | . | . |
| . | . | . | . | . | . | . | . | T | . | . | . | . | . | . | . | . | . | . | . | . | . | . | . | . | . |
| . | . | . | . | . | . | . | . | . | . | . | . | . | . | . | . | . | . | . | . | . | . | . | . | . | . |
| . | . | . | . | . | . | . | . | . | . | . | . | . | . | . | . | . | . | . | . | . | . | . | . | . | . |
| . | . | . | . | . | . | . | . | . | . | . | . | . | . | . | . | . | . | . | . | . | . | . | . | . | . |
| . | . | . | . | . | . | . | . | . | . | . | . | . | . | . | . | . | . | . | . | . | . | . | . | . | . |
| . | . | . | . | . | . | . | . | . | . | . | . | . | . | . | . | . | . | . | . | . | . | . | . | . | . |
| . | . | . | . | . | . | . | . | . | . | . | . | . | . | . | . | . | . | . | . | . | . | . | . | . | . |
| . | . | . | . | . | . | . | . | T | . | . | . | . | . | . | . | . | . | . | . | . | . | . | . | . | . |
| . | . | . | . | . | . | . | . | T | . | . | . | . | . | . | . | . | . | . | . | . | . | . | . | . | . |
| . | . | . | . | . | . | . | . | . | . | . | . | . | . | . | . | . | . | . | . | . | . | . | . | . | . |
| . | . | . | . | . | . | . | . | T | . | . | . | . | . | . | . | . | . | . | . | . | . | . | . | . | . |
| . | . | . | . | . | . | . | . | . | . | . | . | . | . | . | . | . | . | . | . | . | . | . | . | . | . |
| . | . | . | . | . | . | . | . | T | . | . | . | . | . | . | . | . | . | . | . | . | . | . | . | . | . |
| . | . | . | . | . | . | . | . | . | . | . | . | . | . | . | . | . | . | . | . | . | . | . | . | . | . |
| . | . | . | . | . | . | . | . | . | . | . | . | . | . | . | . | . | . | . | . | . | . | . | . | . | . |
| . | . | . | . | . | . | . | . | T | . | . | . | . | . | . | . | . | . | . | . | . | . | . | . | . | . |
| . | . | . | . | . | . | . | . | T | . | . | . | . | . | . | . | . | . | . | . | . | . | . | . | . | . |
| . | . | . | . | . | . | . | . | T | . | . | . | . | . | . | . | . | . | . | . | . | . | . | . | . | . |

| | | | | | | | | | | | | | | | | | | | | | | | | | |
|---|---|---|---|---|---|---|---|---|---|---|---|---|---|---|---|---|---|---|---|---|---|---|---|---|---|
| . | . | . | . | . | . | . | . | T | . | . | . | . | . | . | . | . | . | . | . | . | . | . | . | . | . |
| . | . | . | . | . | . | . | . | . | . | . | . | . | . | . | . | . | . | . | . | . | . | . | . | . | . |
| . | . | . | . | . | . | . | . | T | . | . | . | . | . | . | . | . | . | . | . | . | . | . | . | . | . |
| . | . | . | . | . | . | . | . | T | . | . | . | . | . | . | . | . | . | . | . | . | . | . | . | . | . |
| . | . | . | . | . | . | . | . | T | . | . | . | . | . | . | . | . | . | . | . | . | . | . | . | . | . |
| . | . | . | . | . | . | . | . | T | . | . | . | . | . | . | . | . | . | . | . | . | . | . | . | . | . |
| . | . | . | . | . | . | . | . | . | . | . | . | . | . | . | . | . | . | . | . | . | . | . | . | . | . |
| . | . | . | . | . | . | . | . | T | . | . | . | . | . | . | . | . | . | . | . | . | . | . | . | . | . |
| . | . | . | . | . | . | . | . | . | . | . | . | . | . | . | . | . | . | . | . | . | . | . | . | . | . |
| . | . | . | . | . | . | . | . | T | . | . | . | . | . | . | . | . | . | . | . | . | . | . | . | . | C |
| . | . | . | . | . | . | . | . | . | . | . | . | . | . | . | . | . | . | . | . | . | . | . | . | . | . |
| . | . | . | . | . | . | . | . | . | . | . | . | . | . | . | . | . | . | . | . | . | . | . | . | . | . |
| . | . | . | . | . | . | . | . | . | . | . | . | . | . | . | . | . | . | . | . | . | . | . | . | . | . |
| . | . | . | . | . | . | . | . | T | . | . | . | . | . | . | . | . | . | . | . | . | . | . | . | . | . |
| . | . | . | . | . | . | . | . | . | . | . | . | . | . | . | . | . | . | . | . | . | . | . | . | . | . |
| . | . | . | . | . | . | . | . | T | . | . | . | . | . | . | . | . | . | . | . | . | . | . | . | . | . |
| . | . | . | . | . | . | . | . | . | . | . | . | . | . | . | . | . | . | . | . | . | . | . | . | . | . |
| . | . | . | . | . | . | . | . | T | . | . | . | . | . | . | . | . | . | . | . | . | . | . | . | . | . |
| . | . | . | . | . | . | . | . | T | . | . | . | . | . | . | . | . | . | . | . | . | . | . | . | . | . |
| . | . | . | . | . | . | . | . | T | . | . | . | . | . | . | . | . | . | . | . | . | . | . | . | . | . |
| . | . | . | . | . | . | . | . | . | . | . | . | . | . | . | . | . | . | . | . | . | . | . | . | . | . |
| . | . | . | . | . | . | . | . | . | . | . | . | . | . | . | . | . | . | . | . | . | . | . | . | . | . |
| . | . | . | . | . | . | . | . | . | . | . | . | . | . | . | . | . | . | . | . | . | . | . | . | . | . |
| . | . | . | . | . | . | . | . | T | . | . | . | . | . | . | . | . | . | . | . | . | T | . | . | . | . |
| . | . | . | . | . | . | . | . | T | . | . | . | . | . | . | . | . | . | . | . | . | . | . | . | . | . |
| . | . | . | . | . | . | . | . | T | . | . | . | . | . | . | . | . | . | . | . | . | . | . | . | . | . |
| . | . | . | . | . | . | . | . | T | . | . | . | . | . | . | . | . | . | . | . | . | . | . | . | . | . |
| . | . | . | . | . | . | . | . | T | . | . | . | . | . | . | . | . | . | . | . | . | . | . | . | . | . |
| . | . | . | . | . | . | . | . | . | . | . | . | . | . | . | . | . | . | . | . | . | . | . | . | . | . |
| . | . | . | . | . | . | . | . | T | . | . | . | . | . | . | . | . | . | . | . | . | . | . | . | . | . |
| . | . | . | . | . | . | . | . | T | . | . | . | . | . | . | . | . | . | . | . | . | . | . | . | . | . |
| . | . | . | . | . | . | . | . | T | . | . | . | . | . | . | . | . | . | . | . | . | . | . | . | . | . |
| . | . | . | . | . | . | . | . | T | . | . | . | . | . | . | . | . | . | . | . | . | . | . | . | . | . |
| . | . | . | . | . | . | . | . | T | . | . | . | . | . | . | . | . | . | . | . | . | . | . | . | . | . |
| . | . | . | . | . | . | . | . | T | . | . | . | . | . | . | . | . | . | . | . | . | . | . | . | . | . |
| . | . | . | . | . | . | . | . | T | . | . | . | . | . | . | . | . | . | . | . | . | . | . | . | . | . |
| . | . | . | . | . | . | . | . | T | . | . | . | . | . | . | . | . | . | . | . | . | . | . | . | . | . |
| . | . | . | . | . | . | . | . | . | . | . | . | . | . | . | . | . | . | . | . | . | . | . | . | . | . |
| . | . | . | . | . | . | . | . | T | . | . | . | . | . | . | . | . | . | . | . | T | . | . | . | . | . |
| . | . | . | . | . | . | . | . | T | . | . | . | . | . | . | . | . | . | . | . | . | . | . | . | . | . |
| . | . | . | . | . | . | . | . | T | . | . | . | . | . | . | . | . | . | . | . | . | . | . | . | . | . |
| . | . | . | . | . | . | . | . | T | . | . | . | . | . | . | . | . | . | . | . | . | . | . | . | . | . |
| . | . | . | . | . | . | . | . | . | . | . | . | . | . | . | . | . | . | . | . | . | . | . | . | . | . |
| . | . | . | . | . | . | . | . | T | . | . | . | . | . | . | . | . | . | . | . | . | . | . | . | . | . |
| . | . | . | . | . | . | . | . | T | . | . | . | . | . | . | . | . | . | . | . | . | . | . | . | . | . |
| . | . | . | . | . | . | . | . | T | . | . | . | . | . | . | . | . | . | . | . | . | . | . | . | . | . |
| . | . | . | . | . | . | . | . | T | . | . | . | . | . | . | . | . | . | . | . | . | . | . | . | . | . |
| . | . | . | . | . | . | . | . | T | . | . | . | . | . | . | . | . | . | . | . | . | . | . | . | . | . |
| . | . | . | . | . | . | . | . | T | . | . | . | . | . | . | . | . | . | . | . | . | . | . | . | . | . |
| . | . | . | . | . | . | . | . | T | . | . | . | . | . | . | . | . | . | . | . | . | . | . | . | . | . |
| . | . | . | . | . | . | . | . | T | . | . | . | . | . | . | . | . | . | . | . | . | . | . | . | . | . |
| . | . | . | . | . | . | . | . | T | . | . | . | . | . | . | . | . | . | . | . | . | . | . | . | . | . |
| . | . | . | . | . | . | . | . | T | . | . | . | . | . | . | . | . | . | . | . | . | . | . | . | . | . |
| . | . | . | . | . | . | . | . | T | . | . | . | . | . | . | . | . | . | . | . | . | . | . | . | . | . |
| . | . | . | . | . | . | . | . | . | . | . | . | . | . | . | . | . | . | . | . | . | . | . | . | . | . |
| . | . | . | . | . | . | . | . | . | . | . | . | . | . | . | . | . | . | . | . | . | . | . | . | . | . |
| . | . | . | . | . | . | . | . | T | . | . | . | . | . | . | . | . | . | . | . | . | . | . | . | . | . |
| . | . | . | . | . | . | . | . | . | . | . | . | . | . | . | . | . | . | . | . | . | . | . | . | . | . |
| . | . | . | . | . | . | . | . | T | . | . | . | . | . | . | . | . | . | . | . | . | . | . | . | . | . |
| . | . | . | . | . | . | . | . | . | . | . | . | . | . | . | . | . | . | . | . | . | . | . | . | . | . |
| . | . | . | . | . | . | . | . | . | . | . | . | . | . | . | . | . | . | . | . | . | . | . | . | . | . |
| . | . | . | . | . | . | . | . | . | . | . | . | . | . | . | . | . | . | . | . | . | . | . | . | . | . |

```
. . . . . . . . T . . . . . . . . . . . . . . . . .
. . . . . . . . T . . . . . . . . . . . . . . . . .
. . . . . . . . . . . . . . . . . . . . . . . . . .
. . . . . . . . T . . . . . . . . . . . . . . . . .
. . . . . . . . . . . . . . . . . . . . . . . . . .
. . . . . . . . . . . . . . . . . . . . . . . . . .
. . . . . . . . . . . . . . . . . . . . . . . . . .
. . . . . . . . . . . . . . . . . . . . . . . . . .
. . . . . . . . . . . . . . . . . . . . . . . . . .
. . . . . . . . . . . . . . . . . . . . . . . . . .
. . . . . . . . . . . . . . . . . . . . . . . . . .
. . . . . . . . . . . . . . . . . . . . . . . . . .
. . . . . . . . . . . . . . . . . . . . . . . . . .
. . . . . . . . . . . . . . . . . . . . . . . . . .
. . . . . . . . . . . . . . . . . . . . . . . . . .
. . . . . . . . . . . . . . . . . . . . . . . . . .
. . . . . . . . . . . . . . . . . . . . . . . . . .
. . . . . . . . . . . . . . . . . . . . . . . . . .
. . . . . . . . . . . . . . . . . . . . . . . . . .
. . . . . . . . . . . . . . . . . . . . . . . . . .
. . . . . . . . . . . . . . . . . . . . . . . . . .
. . . . . . . . . . . . . . . . . . . . . . . . . .
. . . . . . . . . . . . . . . . . . . . . . . . . .
. . . . . . . . . . . . . . . . . . . . . . . . . .
. . . . . . . . . . . . . . . . . . . . . . . . . .
. . . . . . . . . . . . . . . . . . . . . . . . . .
. . . . . . . . . . . . . . . . . . . . . . . . . .
. . . . . . . . . . . . . . . . . . . . . . . . . .
. . . . . . . . . . . . . . . . . . . . . . . . . .
. . . . . . . . . . . . . . . . . . . . . . . . . .
. . . . . . . . . . . . . . . . . . . . . . . . . .
. . . . . . . . . . . . . . . . . . . . . . . . . .
. . . . . . . . . . . . . . . . . . . . . . . . . .
. . . . . . . . . . . . . . . . . . . . . . . . . .
. . . . . . . . . . . . . . . . . . . . . . . . . .
. . . . . . . . . T . . . . . C . . . . . . . . . .
. . . . . . . . . . . . . . . . . . . . . . . . . .
. . . . . . . . . . . . . . . . . . . . . . . . . .
. . . . . . . . . . . . . . . . . . . . . . . . T .
. . . . . . . . . . . . . . . . . . . . . . . . . .
. . . . . . . . . . . . . . . . . . . . T . . . . .
. . . . . . . . . . . . . . . . . . . . . . . . . .
. . . . . . . . . . . . . . . . . . . . . . . . . .
. . . . . . . . . . . . . . . . . . . . . . . . . .
. . . . . . . . . . . . . . . . . . . . . . . . . .
. . . . . . . . . . . . . . . . . . . . . . . . . .
. . . . . . . . . . . . . . . . . . . . . . . . . .
. . . . . . . . . . . T . . . . . . . . . . . . . .
. . . . . . . . . . . . . . . . . . . . . . . . . .
. . . . . . . . . . . . . . . . . . . . . . . . . .
. . . . . . . . . . . . . . . . . . . . . . . . . .
. . . . . . . . . . . . . . . . . . . . . . . . . .
. . . . . . . . . . . . . . . . . . . . . . . . . .
. . . . . . . . . . . . . . . . . . . . . . . . . .
. . . . . . . . . . . . . . . . . . . . . . . . . .
. . . . . . . . . . . . . . . . . . . . T . . . . .
. . . . . . . . . . . . . . . . . . . . . . . . . .
. . . . . . . . . . . . . . . . . . . . . . . . . .
. . . . . . . . . . . . . . . . . . . . . . . . . .
. . . . . . . . . . . . . . . . . . . . . . C . . .
. . . . . . . . . . . . . . . . . . . . . . . . . .
. . . . . . . . . . . . . . . . . . . . . . . . . .
```

| | | | | | | | | | | | | | | | | | | | | | | | | | |
|---|---|---|---|---|---|---|---|---|---|---|---|---|---|---|---|---|---|---|---|---|---|---|---|---|---|
| . | . | . | . | . | . | . | . | . | . | . | . | . | . | . | . | . | . | . | . | . | . | . | . | . | . |
| . | . | . | . | . | . | . | . | . | . | . | . | . | . | . | . | . | . | . | . | . | . | . | . | . | . |
| . | . | . | . | . | . | . | . | . | . | . | . | . | . | . | . | . | . | . | . | . | . | . | . | . | . |
| . | . | . | . | . | . | . | . | . | . | . | . | . | . | . | . | . | . | . | . | . | . | . | . | . | . |
| . | . | . | . | . | . | . | . | . | . | . | . | . | . | . | . | . | . | . | . | . | . | . | . | . | . |
| . | . | . | . | . | . | . | . | . | . | . | . | . | . | . | . | . | . | . | . | . | . | . | . | . | . |
| . | . | . | . | . | . | . | . | . | . | . | . | . | . | . | . | . | . | . | . | . | . | . | . | . | . |
| . | . | . | . | . | . | . | . | T | . | . | . | . | . | . | . | . | . | . | . | . | . | . | . | . | . |
| . | . | . | . | . | . | . | . | T | . | . | . | . | . | . | . | . | . | . | . | . | . | . | . | . | . |
| . | . | . | . | . | . | . | . | . | . | . | . | . | . | . | . | . | . | . | . | . | . | . | . | . | . |
| . | . | . | . | . | . | . | . | . | . | . | . | . | . | . | . | . | . | . | . | . | . | . | . | . | . |
| . | . | . | . | . | . | . | . | . | . | . | . | . | . | . | . | . | . | . | . | . | . | . | . | . | . |
| . | . | . | . | . | . | . | . | . | . | . | . | . | . | . | . | . | . | . | . | . | . | . | . | . | . |
| . | . | . | . | . | . | . | . | . | . | . | . | . | . | . | . | . | . | . | . | . | . | . | . | . | . |
| . | . | . | . | . | . | . | . | T | . | . | . | . | . | . | . | . | . | . | . | . | . | . | . | . | . |
| . | . | . | . | . | . | . | . | . | . | . | . | . | . | . | . | . | . | . | . | . | . | C | . | . | . |
| . | . | . | . | . | . | . | . | . | . | . | . | . | . | . | . | . | . | . | . | . | . | C | . | . | . |
| . | . | . | . | . | . | . | . | T | . | . | . | . | . | . | . | . | . | . | . | . | . | . | . | . | . |
| . | . | . | . | . | . | . | . | . | . | . | . | . | . | . | . | . | . | . | . | . | . | . | . | . | . |
| . | . | . | . | . | . | . | . | T | . | . | . | . | . | . | . | . | . | . | . | . | . | . | . | . | . |
| . | . | . | . | . | . | . | . | T | . | . | . | . | . | . | . | . | . | . | . | . | . | . | . | . | . |
| . | . | . | . | . | . | . | . | T | . | . | . | . | . | . | . | . | . | . | . | . | . | . | . | . | . |
| . | . | . | . | . | . | . | . | . | . | . | . | . | . | . | . | . | . | . | . | . | . | . | . | . | . |
| . | . | . | . | . | . | . | . | . | . | . | . | . | . | . | . | . | . | . | . | . | . | . | . | . | . |
| . | . | . | . | . | . | . | . | . | . | . | . | . | . | . | . | . | . | . | . | . | . | . | . | . | . |
| . | . | . | . | . | . | . | . | . | . | . | . | . | . | . | . | . | . | . | . | . | . | . | . | . | . |
| . | . | . | . | . | . | . | . | . | . | . | . | . | . | . | . | . | . | . | . | . | . | . | . | . | . |
| . | . | . | . | . | . | . | . | . | . | . | . | . | . | . | . | . | . | . | . | . | . | . | . | . | . |
| . | . | . | . | . | . | . | . | T | . | . | . | . | . | . | . | . | . | . | . | . | . | . | . | . | . |
| . | . | . | . | . | . | . | . | . | . | . | . | . | . | . | . | . | . | . | . | . | . | . | . | . | . |
| . | . | . | . | . | . | . | . | T | . | . | . | . | . | . | . | . | . | . | . | . | . | . | . | . | . |
| . | . | . | . | . | . | . | . | . | . | . | . | . | . | . | . | . | . | . | . | . | . | . | . | . | . |
| . | . | . | . | . | . | . | . | . | . | . | . | . | . | . | . | . | . | . | . | . | . | . | . | . | . |
| t | g | g | c | g | a | c | g | c | c | a | c | a | g | a | t | g | c | c | g | g | g | t | g | c | a |
| t | g | g | c | g | a | c | g | c | c | a | c | a | g | a | t | g | c | c | g | g | g | t | g | c | a |
| t | g | g | c | g | a | c | g | c | c | a | c | a | g | a | t | g | c | c | g | g | g | t | g | c | a |
| t | g | g | c | g | a | c | g | c | c | a | c | a | g | a | t | g | c | c | g | g | g | t | g | c | a |
| t | g | g | c | g | a | c | g | c | c | a | c | a | g | a | t | g | c | c | g | g | g | t | g | c | a |
| t | g | g | c | g | a | c | g | c | c | a | c | a | g | a | t | g | c | c | g | g | g | t | g | c | a |
| t | g | g | c | g | a | c | g | c | c | a | c | a | g | a | t | g | c | c | g | g | g | t | g | c | a |
| t | g | g | c | g | a | c | g | c | c | a | c | a | g | a | t | g | c | c | g | g | g | t | g | c | a |
| t | g | g | c | g | a | c | g | c | c | a | c | a | g | a | t | g | c | t | g | g | g | t | g | c | a |
| t | g | g | c | g | a | c | g | c | c | a | c | a | g | a | t | g | c | c | g | g | g | c | g | c | a |
| t | g | g | c | g | a | c | g | c | c | a | c | a | g | a | t | g | c | c | g | g | g | c | g | c | a |
| t | g | g | c | g | a | c | g | c | c | a | c | a | g | a | t | g | c | c | g | g | g | c | g | c | a |
| t | g | g | c | g | a | c | g | c | c | a | c | a | g | a | t | g | c | c | g | g | g | c | g | c | a |
| t | g | g | c | g | a | c | g | c | c | a | c | a | g | a | t | g | c | c | g | g | g | t | g | c | a |
| t | g | g | c | g | a | c | g | c | c | a | c | a | g | a | t | g | c | c | g | g | g | t | g | c | a |
| t | g | g | c | g | a | c | g | c | c | a | c | a | g | a | t | g | c | c | g | g | g | t | g | c | a |
| t | g | g | c | g | a | c | g | c | c | a | c | a | g | a | t | g | c | c | g | g | g | t | g | c | a |
| t | g | g | c | g | a | c | g | c | c | a | c | a | g | a | t | g | c | c | g | g | g | t | g | c | a |
| t | g | g | c | g | a | c | g | c | c | a | c | a | g | a | t | g | c | c | g | g | g | t | g | c | a |
| t | g | g | c | g | a | c | g | c | c | a | c | a | g | a | t | g | c | c | g | g | g | t | g | c | a |
| t | g | g | c | g | a | c | g | c | c | a | c | a | g | a | t | g | c | c | g | g | g | t | g | c | a |
| t | g | g | c | g | a | c | g | c | c | a | c | a | g | a | t | g | c | c | g | g | g | t | g | c | a |
| t | g | g | c | g | a | c | g | c | c | a | c | a | g | a | t | g | c | c | g | g | g | t | g | c | a |
| t | g | g | c | g | a | c | g | c | c | a | c | a | g | a | t | g | c | c | g | g | g | t | g | c | a |
| . | . | . | . | . | . | . | . | . | . | . | . | . | . | . | . | . | . | . | . | . | . | . | . | . | . |
| . | . | . | . | . | . | . | . | . | . | . | . | . | . | . | . | . | . | . | . | . | . | . | . | . | . |
| . | . | . | . | . | . | . | . | . | . | . | . | . | . | . | . | . | . | . | . | . | . | . | . | . | . |
| . | . | . | . | . | . | . | . | . | . | . | . | . | . | . | . | . | . | . | . | . | . | . | . | . | . |
| . | . | . | . | . | . | . | . | . | . | . | . | . | . | . | . | . | . | . | . | . | . | . | . | . | . |

| | | | | | | | | | | | | | | | | | | | | | | | | | |
|---|---|---|---|---|---|---|---|---|---|---|---|---|---|---|---|---|---|---|---|---|---|---|---|---|---|
| . | . | . | . | . | . | . | . | . | . | . | . | . | . | . | . | . | . | . | . | . | . | . | . | . | . |
| . | . | . | . | . | . | . | . | . | . | . | . | . | . | . | . | . | . | T | . | . | . | . | . | . | . |
| . | . | . | . | . | . | . | . | . | . | . | . | . | . | . | . | . | . | . | . | . | . | . | . | . | . |
| . | . | . | . | . | . | . | . | . | . | . | . | . | . | . | . | . | . | . | . | . | . | . | . | . | . |
| . | . | . | . | . | G | . | . | . | . | . | . | . | . | . | . | . | . | . | . | . | . | . | . | . | . |
| . | . | . | . | . | . | . | . | . | . | . | . | . | . | . | . | . | . | . | . | . | . | . | . | . | . |
| . | ? | . | . | . | . | . | . | . | . | . | . | . | . | . | . | . | . | . | . | . | . | . | . | . | . |
| . | T | . | . | . | . | . | . | . | . | . | . | . | . | . | . | . | . | . | . | . | . | C | . | . | . |
| . | . | . | . | . | . | . | . | . | . | . | . | . | . | . | . | . | . | . | . | . | . | . | . | . | . |
| . | . | . | . | . | . | . | . | . | . | . | . | . | . | . | . | . | . | . | . | . | . | . | . | . | . |
| . | . | . | . | . | . | . | . | . | . | . | . | . | . | . | . | . | . | . | . | . | . | C | . | . | . |
| . | T | . | . | . | . | . | . | . | . | . | . | . | . | . | . | . | . | . | . | . | . | C | . | . | . |
| G | ? | A | A | . | . | . | . | . | . | . | . | . | . | . | . | . | . | . | . | . | . | C | . | . | . |
| . | . | . | . | . | . | . | . | . | . | . | . | . | . | . | . | . | . | . | . | . | . | . | . | . | . |
| . | . | . | . | . | . | . | . | . | . | . | . | . | . | . | . | . | . | . | . | . | . | . | . | . | . |
| . | . | . | . | . | . | . | . | . | . | . | . | . | . | . | . | . | . | . | . | . | . | . | . | . | . |
| . | . | . | . | . | . | . | . | . | . | . | . | . | . | . | . | . | . | . | . | . | . | . | . | . | . |
| . | . | . | . | . | . | . | . | . | . | . | . | . | . | . | . | . | . | . | . | . | . | . | . | T | . |
| . | . | . | . | . | . | . | . | . | . | . | . | . | . | . | . | . | . | . | . | . | . | . | . | . | . |
| . | . | . | . | . | . | . | . | . | . | . | . | . | . | . | . | . | . | . | . | . | . | . | . | . | . |
| . | . | . | . | . | . | . | . | . | . | . | . | . | . | . | . | . | . | . | . | . | . | . | . | . | . |
| . | . | . | . | . | . | . | . | . | . | . | . | . | . | . | . | . | . | . | . | . | . | . | . | . | . |
| . | . | . | . | . | . | . | . | . | . | . | . | . | . | . | . | . | . | . | . | . | . | . | . | . | . |
| . | . | . | . | . | . | . | . | . | . | . | . | . | . | . | . | . | . | . | . | . | . | . | . | . | . |
| . | . | . | . | . | . | . | . | . | . | . | . | . | . | . | . | . | . | . | . | . | . | . | . | . | . |
| . | . | A | A | . | . | . | . | . | . | . | . | . | . | . | . | . | . | . | . | . | . | . | . | . | . |
| . | . | . | . | . | . | . | . | . | . | . | . | . | . | . | . | . | . | . | . | . | . | . | . | . | . |
| . | . | . | . | . | . | . | . | . | . | . | . | . | . | . | . | . | . | . | . | . | . | . | . | . | . |
| . | . | . | . | . | . | . | . | . | . | . | . | . | . | . | . | . | . | . | . | . | . | . | . | . | . |
| . | . | . | . | . | . | . | . | . | . | . | . | . | . | . | . | . | . | . | . | . | . | . | . | . | . |
| G | ? | . | . | . | . | . | . | . | . | . | . | . | . | . | . | . | . | . | . | . | . | . | . | . | . |
| . | . | . | . | . | . | . | . | . | . | . | . | . | . | . | . | . | T | . | . | . | . | . | . | . | . |
| . | . | . | . | . | . | . | . | . | . | . | . | . | . | . | . | . | . | . | . | . | . | . | . | . | . |
| . | . | . | . | . | . | . | . | . | . | . | . | . | . | . | . | . | . | . | . | . | . | . | . | . | . |
| . | . | . | . | . | . | . | . | . | . | . | . | . | . | . | . | . | . | . | . | . | . | . | . | . | . |
| . | . | . | . | . | . | . | . | . | . | . | . | . | . | . | . | . | . | . | . | . | . | . | . | . | . |
| . | . | . | . | . | . | . | . | . | . | . | . | . | . | . | . | . | . | . | . | . | . | . | . | . | . |
| . | . | . | . | . | . | . | . | . | . | . | . | . | . | . | . | . | . | . | . | . | . | . | . | . | . |
| . | . | . | . | . | . | . | . | . | . | . | . | . | . | . | . | . | . | . | . | . | . | . | . | . | . |
| . | . | . | . | . | . | . | . | . | . | . | . | . | . | . | . | . | . | . | . | . | . | . | . | . | . |
| . | . | . | . | . | . | . | . | . | . | . | . | . | . | . | . | . | . | . | . | . | . | C | . | . | . |
| . | . | . | . | . | . | . | . | . | . | . | . | . | . | . | . | . | . | . | . | . | . | C | . | . | . |
| . | . | . | . | . | . | . | . | T | . | . | . | . | . | . | . | . | . | . | . | . | . | . | . | . | . |
| . | . | . | . | . | . | . | . | . | . | . | . | . | . | . | . | . | . | . | . | . | . | . | . | . | . |
| . | . | . | . | . | . | . | . | . | . | . | . | . | . | . | . | . | . | . | . | . | . | . | . | . | . |
| . | . | . | . | . | . | . | . | . | . | . | . | . | . | . | . | . | . | . | . | . | . | . | . | . | . |
| . | . | . | . | . | . | . | . | T | . | . | . | . | . | . | . | . | . | . | . | . | . | . | . | . | . |
| . | . | . | . | . | . | . | . | . | . | . | . | . | . | . | . | . | . | . | . | . | . | . | . | . | . |
| . | . | . | . | . | . | . | . | T | . | . | . | . | . | . | . | . | . | . | . | . | . | . | . | . | . |
| . | . | . | . | . | . | . | . | . | . | . | . | . | . | . | . | . | . | . | . | . | . | . | . | . | . |
| . | . | . | . | . | . | . | . | . | . | . | . | . | . | . | . | . | . | . | . | . | . | . | . | . | . |
| . | . | . | . | . | . | . | . | . | . | . | . | . | . | . | . | . | . | . | . | . | . | . | . | . | . |
| . | . | . | . | . | . | . | . | T | . | . | . | . | . | . | . | . | . | . | . | . | . | . | . | . | . |
| . | . | . | . | . | . | . | . | T | . | . | . | . | . | . | . | . | . | . | . | . | . | . | . | . | . |
| . | . | . | . | . | . | . | . | . | . | . | . | . | . | . | . | . | . | . | . | . | . | . | T | . | . |
| . | . | . | . | . | . | . | . | . | . | . | . | . | . | . | . | . | . | . | . | . | . | . | . | . | . |
| . | . | . | . | . | . | . | . | . | . | . | . | . | . | . | . | . | . | . | . | . | . | . | . | . | . |
| . | . | . | . | . | . | . | . | . | . | . | . | . | . | . | . | . | . | . | . | . | . | C | . | . | . |
| . | . | . | . | . | . | . | . | . | . | . | . | . | . | . | . | . | . | . | . | . | . | . | T | . | . |
| . | . | . | . | . | . | . | . | . | . | . | . | . | . | . | . | . | . | . | . | . | . | . | . | . | . |
| . | . | . | . | . | . | . | . | . | . | . | . | . | . | . | . | . | . | . | . | . | . | . | . | . | . |
| . | . | . | . | . | . | . | . | . | . | . | . | . | . | . | . | . | . | . | . | . | . | C | . | . | . |

| | | | | | | | | | | | | | | | | | | | | | | | | | |
|---|---|---|---|---|---|---|---|---|---|---|---|---|---|---|---|---|---|---|---|---|---|---|---|---|---|
| . | . | . | . | . | . | . | . | . | . | . | . | . | . | . | . | . | . | . | . | . | . | . | . | . | . |
| . | . | . | . | . | . | . | . | . | . | . | . | . | . | . | . | . | . | . | . | . | . | . | . | . | . |
| . | . | . | . | . | . | . | . | . | . | . | . | . | . | . | . | . | . | . | . | . | . | . | . | . | . |
| . | . | . | . | . | . | . | T | . | . | . | . | . | . | . | . | . | . | . | . | . | . | . | . | . | . |
| . | . | . | . | . | . | . | . | . | . | . | . | . | . | . | . | . | . | . | . | . | . | . | . | . | . |
| . | . | . | . | . | . | . | . | . | . | . | . | . | . | . | . | . | . | . | . | . | . | . | . | . | . |
| . | . | . | . | . | . | . | . | . | . | . | . | . | . | . | . | . | . | . | . | . | . | C | . | . | . |
| . | . | . | . | . | . | . | . | . | . | . | . | . | . | . | . | . | . | . | . | . | . | . | . | . | . |
| . | . | . | . | . | . | . | . | . | . | . | . | . | . | . | . | . | . | . | . | . | . | . | . | . | . |
| . | . | . | . | . | . | . | . | . | . | . | . | . | . | . | . | . | . | . | . | . | . | . | T | . | . |
| . | . | . | . | . | . | . | T | . | . | . | . | . | . | . | . | . | . | . | . | . | . | . | . | . | . |
| . | . | . | . | . | . | . | . | . | . | . | . | . | . | . | . | . | . | . | . | . | . | . | . | . | . |
| . | . | . | . | . | . | . | T | . | . | . | . | . | . | . | . | . | . | . | . | . | . | . | . | . | . |
| . | . | . | . | . | . | . | . | . | . | . | . | . | . | . | . | . | . | . | . | . | . | . | . | . | . |
| . | . | . | . | . | . | . | . | . | . | . | . | . | . | . | . | . | . | . | . | . | . | . | . | . | . |
| . | . | . | . | . | . | T | . | . | . | . | . | . | . | . | . | . | . | . | . | . | . | C | . | . | . |
| . | . | . | . | . | . | . | . | . | . | . | . | . | . | . | . | . | . | . | . | . | . | . | . | . | . |
| . | . | . | . | . | . | . | . | . | . | . | . | . | . | . | . | . | . | . | . | . | . | . | . | . | . |
| . | . | . | . | . | . | . | . | . | . | . | . | . | . | . | . | . | . | . | . | . | . | . | . | . | . |
| . | . | . | . | . | . | . | T | . | . | . | . | . | . | . | . | . | . | . | . | . | . | . | . | . | . |
| . | . | . | . | . | . | . | . | . | . | . | . | . | . | . | . | . | . | . | . | . | . | . | . | . | . |
| . | . | . | . | . | . | . | . | . | . | . | . | . | . | . | . | . | . | . | . | . | . | . | . | . | . |
| . | . | . | . | . | . | . | . | . | . | . | . | . | . | . | . | . | . | . | . | . | . | . | . | . | . |
| . | . | . | . | . | . | . | . | . | . | . | . | . | . | . | . | . | . | . | . | . | . | C | . | . | . |
| . | . | . | . | . | . | . | . | . | . | . | . | . | . | . | . | ? | . | . | . | . | . | C | . | . | . |
| . | . | . | . | . | . | . | . | . | . | . | . | . | . | . | . | . | . | . | . | ? | . | C | . | . | . |
| . | . | . | . | . | . | . | . | . | . | . | . | . | . | . | . | . | . | . | T | . | . | . | . | . | . |
| . | . | . | . | . | . | . | . | . | . | . | . | . | . | . | . | . | . | . | . | . | . | . | . | . | . |
| . | . | . | . | . | . | . | . | . | . | . | . | . | . | . | . | . | . | . | ? | . | . | . | . | . | . |
| . | . | . | . | . | . | . | . | . | . | . | . | . | . | . | . | . | . | . | . | . | . | C | . | . | . |
| . | . | . | . | . | . | . | . | . | . | . | . | . | . | . | . | . | . | . | . | . | . | C | . | . | . |
| . | . | . | . | . | . | . | . | . | . | . | . | . | . | . | . | . | . | . | . | . | . | C | . | . | . |
| . | . | . | . | . | . | . | . | . | . | . | . | . | . | . | . | . | . | . | . | . | . | C | . | . | . |
| . | . | . | . | . | . | . | . | . | . | . | . | . | . | . | . | . | . | . | . | . | . | C | . | . | . |
| . | . | . | . | . | . | . | . | . | . | . | . | . | . | . | . | . | . | . | T | . | . | . | . | . | . |
| . | . | . | . | . | . | . | . | . | . | . | . | . | . | . | . | . | . | . | . | . | . | C | . | . | . |
| . | . | . | . | . | . | . | . | . | . | . | . | . | . | . | . | . | . | . | . | . | . | C | . | . | . |
| . | . | . | . | . | . | . | . | . | . | . | . | . | . | . | . | . | . | . | . | . | . | . | . | . | . |
| . | . | . | . | . | . | . | . | . | . | . | . | . | . | . | . | . | . | . | . | . | . | C | . | . | . |
| . | . | . | . | . | . | . | . | . | . | . | . | . | . | . | . | . | . | . | . | . | . | . | . | . | . |
| . | . | . | . | . | . | . | . | T | . | . | . | . | . | . | . | . | . | . | . | . | . | . | . | . | . |
| . | . | . | . | . | . | . | . | T | . | . | . | . | . | . | . | . | . | . | . | . | . | . | . | . | . |
| . | . | . | . | . | . | . | . | T | . | . | . | . | . | . | . | . | . | . | . | . | . | . | . | . | . |
| . | . | . | . | . | . | . | . | T | . | . | . | . | . | . | . | . | . | . | . | . | . | . | . | . | . |
| . | . | . | . | . | . | . | . | T | . | . | . | . | . | . | . | . | . | . | . | . | . | . | . | . | . |
| . | . | . | . | . | . | . | . | T | . | . | . | . | . | . | . | . | . | . | . | . | . | . | . | . | . |
| . | . | . | . | . | . | . | . | T | . | . | . | . | . | . | . | ? | . | . | . | . | . | . | . | T | . |
| . | . | . | . | . | . | . | . | T | . | . | . | . | . | . | . | . | . | . | . | . | . | . | . | . | . |
| . | . | . | . | . | . | . | . | T | . | . | . | . | . | . | . | . | . | . | . | . | . | . | . | . | . |
| . | . | . | . | . | . | . | . | . | . | . | . | . | . | . | . | . | . | . | . | . | . | . | . | . | . |
| . | . | . | . | . | . | . | . | . | . | . | . | . | . | . | . | . | . | . | . | . | . | . | . | . | . |
| . | . | . | . | . | . | . | . | . | . | . | . | . | . | . | . | . | . | . | . | . | . | . | . | . | . |
| . | . | . | . | . | . | . | . | . | . | . | . | . | . | . | . | . | . | . | . | . | . | . | . | . | . |
| . | . | . | . | . | . | . | . | . | . | . | . | . | . | . | . | . | . | . | . | . | . | . | . | . | . |
| . | . | . | . | . | . | . | . | . | . | . | . | . | . | . | . | . | . | . | . | . | . | . | . | . | . |
| . | . | . | . | . | . | . | . | . | . | . | . | . | . | . | . | . | . | . | . | . | . | . | . | . | . |
| . | . | . | . | . | . | . | . | . | . | . | . | . | . | . | . | . | . | . | . | . | . | . | . | . | . |
| . | . | . | . | . | . | . | . | . | . | . | . | . | . | . | . | . | . | . | . | . | . | . | . | . | . |
| . | . | . | . | . | . | . | . | . | . | . | . | . | . | . | . | . | . | . | . | . | . | . | . | . | . |
| . | . | . | . | . | . | . | . | . | T | . | . | . | . | . | C | . | . | . | . | . | . | . | . | . | . |
| . | . | . | . | . | . | . | . | . | T | . | . | . | . | . | C | . | . | . | . | . | . | . | . | . | . |
| . | . | . | . | . | . | . | . | . | . | . | . | . | . | . | . | . | . | . | . | . | . | . | . | . | . |

| | | | | | | | | | | | | | | | | | | | | | | | | | |
|---|---|---|---|---|---|---|---|---|---|---|---|---|---|---|---|---|---|---|---|---|---|---|---|---|---|
| . | . | . | . | . | . | . | . | . | T | . | . | . | . | . | C | . | . | . | . | . | . | . | . | . | . |
| . | . | . | . | . | . | . | . | . | T | . | . | . | . | . | C | . | . | . | . | . | . | . | . | . | . |
| . | . | . | . | . | . | . | . | . | . | . | . | . | . | . | . | . | . | . | . | . | . | . | . | . | . |
| . | . | . | . | . | . | . | . | . | . | . | . | . | . | . | . | . | . | . | . | . | . | . | . | . | . |
| . | . | . | . | . | . | . | . | . | . | . | . | . | . | . | . | . | . | . | . | . | . | . | . | . | . |
| . | . | . | . | . | . | . | . | . | . | . | . | . | . | . | . | . | . | . | . | . | . | . | . | . | . |
| . | . | . | . | . | . | . | . | . | . | . | . | . | . | . | . | . | . | . | . | . | . | . | . | . | . |
| . | . | . | . | . | . | . | . | . | . | . | . | . | . | . | . | . | . | . | . | . | . | . | . | . | . |
| . | . | . | . | . | . | . | . | . | . | . | . | . | . | . | . | . | . | . | . | . | . | . | . | . | . |
| . | . | . | . | . | . | . | . | . | . | . | . | . | . | . | . | . | . | . | . | . | . | . | . | . | . |
| . | . | . | . | . | . | . | . | . | . | . | . | . | . | . | . | . | . | . | . | . | . | . | . | . | . |
| . | . | . | . | . | . | . | . | . | . | . | . | . | . | . | . | . | . | . | . | . | . | . | . | . | . |
| . | . | . | . | . | . | . | . | . | . | . | . | . | . | . | . | . | . | . | . | . | . | . | . | . | . |
| . | . | . | . | . | . | . | . | . | . | . | . | . | . | . | . | . | . | . | . | . | . | . | . | . | . |
| . | . | . | . | . | . | . | . | . | . | . | . | . | . | . | . | . | . | . | . | . | . | . | . | . | . |
| . | . | . | . | . | . | . | . | . | . | . | . | . | . | . | . | . | . | . | . | . | . | . | . | . | . |
| . | . | . | . | . | . | . | . | . | . | . | . | G | C | T | C | . | . | . | . | . | . | . | . | . | . |
| . | . | . | . | . | . | . | . | . | . | . | . | G | C | T | C | . | . | . | . | . | . | . | . | . | . |
| . | . | . | . | . | . | . | . | . | . | . | . | . | . | . | . | . | . | . | . | . | . | . | . | . | . |
| . | . | . | . | . | . | . | . | . | . | . | . | . | . | . | . | . | . | . | . | . | . | . | . | . | . |
| . | . | . | . | . | . | . | . | . | . | . | . | . | . | . | . | . | . | . | . | . | . | . | . | . | . |
| . | . | . | . | . | . | . | . | . | . | . | . | . | . | . | . | . | . | . | . | . | . | . | . | . | . |
| . | . | . | . | . | . | . | . | . | . | . | . | . | . | . | . | . | . | . | . | . | . | . | . | . | . |
| . | . | . | . | . | . | . | . | . | . | . | . | . | . | . | . | . | . | . | . | . | . | . | . | . | . |
| . | . | . | . | . | . | . | . | . | . | . | . | . | . | . | . | . | . | . | . | . | . | . | . | . | . |
| . | . | . | . | . | . | . | . | . | . | . | . | . | . | . | . | . | . | . | . | . | . | . | . | . | . |
| . | . | . | . | . | . | . | . | . | . | . | . | . | . | . | . | . | . | . | . | . | . | . | . | . | . |
| . | . | . | . | . | . | . | . | . | . | . | . | . | . | . | . | . | . | . | . | . | . | . | . | . | . |
| . | . | . | . | . | . | . | . | . | . | . | . | . | . | . | . | . | . | . | . | . | . | . | . | . | . |
| . | . | . | . | . | . | . | . | . | . | . | . | . | . | . | . | . | . | . | . | . | . | . | . | . | . |
| . | . | . | . | . | . | . | . | . | . | . | . | . | . | . | . | . | . | . | . | . | . | . | . | . | . |
| . | . | . | . | . | . | . | . | . | . | . | T | . | . | . | . | . | . | . | . | . | . | . | . | . | . |
| . | . | . | . | . | . | . | . | . | . | . | . | . | . | . | . | . | . | . | . | . | . | . | . | . | . |
| . | . | . | . | . | . | . | . | . | . | . | . | . | . | . | . | . | . | . | . | . | . | . | . | . | . |
| . | . | . | . | . | . | . | . | . | . | . | . | . | . | . | . | . | . | . | . | . | . | . | . | . | . |
| . | . | . | . | . | . | . | . | . | . | . | . | . | . | . | . | . | . | . | . | . | . | . | . | . | . |
| . | . | . | . | . | . | . | . | . | . | . | . | . | . | . | . | . | . | . | . | . | . | . | . | . | . |
| . | . | . | . | . | . | . | . | . | . | . | . | . | . | . | . | . | . | . | . | . | . | . | . | . | . |
| . | . | . | . | . | . | . | . | T | . | . | . | . | . | . | . | . | . | . | . | . | . | . | . | . | . |
| . | . | . | . | . | . | . | . | T | . | . | . | . | . | . | . | . | . | . | . | . | . | . | . | . | . |
| . | . | . | . | . | . | . | . | T | . | . | . | . | . | . | . | . | . | . | . | . | . | . | . | . | . |
| . | . | . | . | . | . | . | . | T | . | . | . | . | . | . | . | . | . | . | . | . | . | . | . | . | . |
| . | . | . | . | . | . | . | . | T | . | . | . | . | . | . | . | . | . | . | . | . | . | . | . | . | . |
| . | . | . | . | . | . | . | . | . | . | . | . | . | . | . | . | . | . | . | . | . | . | . | . | . | . |
| . | . | . | . | . | . | . | . | T | . | . | . | . | . | . | . | . | . | . | . | . | . | . | . | . | . |
| . | . | . | . | . | . | . | . | T | . | . | . | . | . | . | . | . | . | . | . | . | . | . | . | . | . |
| . | . | . | . | . | . | . | . | T | . | . | . | . | . | . | . | . | . | . | . | . | . | . | . | . | . |
| . | . | . | . | . | . | . | . | T | . | . | . | . | . | . | . | . | . | . | . | . | . | . | . | . | . |
| . | . | . | . | . | . | . | . | . | . | . | . | . | . | . | . | . | . | . | . | . | . | . | . | . | . |
| . | . | . | . | . | . | . | . | T | . | . | . | . | . | . | . | . | . | . | . | . | . | . | . | . | . |
| . | . | . | . | . | . | . | . | T | . | . | . | . | . | . | . | . | . | . | . | . | . | . | . | . | . |
| . | . | . | . | . | . | . | . | T | . | . | . | . | . | . | . | . | . | . | . | . | . | . | . | . | . |
| . | . | . | . | . | . | . | . | T | . | . | . | . | . | . | . | . | . | . | . | . | . | . | . | . | . |
| . | . | . | . | . | . | . | . | . | . | . | . | . | . | . | . | . | . | . | . | . | . | . | . | . | . |
| . | . | . | . | . | . | . | . | T | . | . | . | . | . | . | . | . | . | . | . | . | . | . | . | . | . |
| . | . | . | . | . | . | . | . | T | . | . | . | . | . | . | . | . | . | . | . | . | . | . | . | . | . |
| . | . | . | . | . | . | . | . | . | . | . | . | . | . | . | . | . | . | . | . | . | . | . | . | . | . |
| . | . | . | . | . | . | . | . | . | . | . | . | . | . | . | . | . | . | . | . | . | . | . | . | . | . |
| . | . | . | . | . | . | . | . | . | . | . | . | . | . | . | . | . | . | . | . | . | . | . | . | . | . |
| . | . | . | . | . | . | . | . | . | . | . | . | . | . | . | . | . | . | . | . | . | . | . | . | . | . |
| . | . | . | . | . | . | . | . | . | . | . | . | . | . | . | . | . | . | . | . | . | . | . | . | . | . |
| . | . | . | . | . | . | . | . | . | . | . | . | . | . | . | . | . | . | . | . | . | . | . | . | . | . |

| | | | | | | | | | | | | | | | | | | | | | | | | | |
|---|---|---|---|---|---|---|---|---|---|---|---|---|---|---|---|---|---|---|---|---|---|---|---|---|---|
| . | . | . | . | . | . | . | . | T | . | . | . | . | . | . | . | . | . | . | . | . | . | . | . | . | . |
| . | . | . | . | . | . | T | . | . | . | . | . | . | . | . | . | . | . | . | . | . | . | . | . | . | . |
| . | . | . | . | . | . | . | . | T | . | . | . | . | . | . | . | . | . | . | . | . | T | . | . | . | . |
| . | . | . | . | . | . | . | . | T | . | . | . | . | . | . | . | . | . | . | . | . | T | . | . | . | . |
| . | . | . | . | . | . | . | . | . | . | . | . | . | . | . | . | . | . | . | . | . | . | . | . | . | . |
| . | . | . | . | . | . | . | . | . | . | . | . | . | . | . | . | . | . | . | . | S | . | . | . | . | . |
| . | . | . | . | . | . | . | . | . | . | . | . | . | . | . | . | . | . | . | . | . | . | . | . | . | . |
| . | . | . | . | . | . | . | . | . | . | . | . | . | . | . | . | . | . | . | . | . | . | . | . | . | . |
| . | . | . | . | . | . | . | . | . | . | . | . | . | . | . | . | . | . | . | . | . | . | . | . | . | . |
| . | . | . | . | . | . | . | . | . | . | . | . | . | . | . | . | . | . | . | . | . | . | . | . | . | . |
| . | . | . | . | . | . | . | . | . | . | . | . | . | . | . | . | ? | . | . | . | . | . | . | . | . | . |
| . | . | . | . | . | . | . | . | . | . | . | . | . | . | . | . | . | . | . | . | . | . | . | . | . | . |
| . | . | . | . | . | . | . | . | . | . | . | . | . | . | . | . | . | . | . | . | . | . | . | . | . | . |
| . | . | . | . | . | . | . | . | . | . | . | . | . | . | . | . | . | . | . | . | . | . | . | . | . | . |
| . | . | . | . | . | . | . | . | . | . | . | . | . | . | . | . | . | . | . | . | . | . | . | . | . | . |
| . | . | . | . | . | . | . | . | . | . | . | . | . | . | . | . | . | . | . | . | . | . | . | . | . | . |
| . | . | . | . | . | . | . | . | . | . | . | . | . | . | . | . | . | . | . | . | . | . | . | . | . | . |
| . | . | . | . | . | . | . | . | . | . | . | . | . | . | . | . | . | . | . | . | . | . | . | . | . | . |
| . | . | . | . | . | . | . | . | . | . | . | . | . | . | . | . | . | . | . | . | . | . | . | . | . | . |
| . | . | . | . | . | . | . | . | . | . | . | . | . | . | . | . | . | . | . | . | . | . | . | . | . | . |
| . | . | . | . | . | . | . | . | . | . | . | . | . | . | . | . | g | . | . | . | . | . | . | . | . | . |
| . | . | . | . | . | . | . | . | . | . | . | . | . | . | . | . | g | . | . | . | . | . | . | . | . | . |
| . | . | . | . | . | . | . | . | . | . | . | . | . | . | . | . | . | . | . | . | . | . | . | . | . | . |
| . | . | . | . | . | . | . | . | . | . | . | . | . | . | . | . | . | . | . | . | . | . | . | . | . | . |
| . | . | . | . | . | . | . | . | . | . | . | . | . | . | . | . | . | . | . | . | . | . | . | . | . | . |
| . | . | . | . | . | . | . | . | . | . | . | . | . | . | . | . | . | . | . | . | . | . | . | . | . | . |
| . | . | . | . | . | . | . | . | . | . | . | . | . | . | . | . | . | . | . | . | . | . | . | . | . | . |
| . | . | . | . | . | . | . | . | . | . | . | . | . | . | . | . | g | . | . | . | . | . | . | . | . | . |
| . | . | . | . | . | . | . | . | . | . | . | . | . | . | . | . | . | . | . | . | . | . | . | . | . | . |
| . | . | . | . | . | . | . | . | . | . | . | . | . | . | . | . | . | . | . | . | . | . | . | . | . | . |
| . | . | . | . | . | . | . | . | . | . | . | . | . | . | . | . | g | . | . | . | . | . | . | . | . | . |
| . | . | . | . | . | . | . | . | . | . | . | . | . | . | . | . | . | . | . | . | . | . | . | . | . | . |
| . | . | . | . | . | . | . | . | . | . | . | . | . | . | . | . | . | . | . | . | . | . | . | . | . | . |
| . | . | . | . | . | . | . | . | T | . | . | . | . | . | . | . | . | . | . | . | . | . | . | . | . | . |
| . | . | . | . | . | . | . | . | . | . | . | . | . | . | . | . | . | . | . | . | . | . | . | . | . | . |
| . | . | . | . | . | . | . | . | . | . | . | . | . | . | . | . | . | . | . | . | . | . | C | . | . | . |
| . | . | . | . | . | . | . | . | . | . | . | . | . | . | . | . | . | . | . | . | . | . | . | . | . | . |
| . | . | . | . | . | . | . | . | T | . | . | . | . | . | . | . | . | . | . | . | . | . | . | . | . | . |
| . | . | . | . | . | . | . | . | T | . | . | . | . | . | . | . | . | . | . | . | . | . | . | . | . | . |
| . | . | . | . | . | . | . | . | . | . | . | . | . | . | . | . | . | . | . | . | . | . | . | . | . | . |
| . | . | . | . | . | . | . | . | . | . | . | . | . | . | . | . | . | . | . | . | . | . | . | . | . | . |
| . | . | . | . | . | . | . | . | T | . | . | . | . | . | . | . | . | . | . | . | . | . | . | . | . | . |
| . | . | . | . | . | . | . | . | T | . | . | . | . | . | . | . | . | . | . | . | . | . | . | . | . | . |
| . | . | . | . | . | . | . | . | . | . | . | . | . | . | . | . | . | . | . | . | . | . | . | . | . | . |
| . | . | . | . | . | . | . | . | T | . | . | . | . | . | . | . | . | . | . | . | . | . | . | . | . | . |
| . | . | . | . | . | . | . | . | . | . | . | . | . | . | . | . | . | . | . | . | . | . | . | . | . | . |
| . | . | . | . | . | . | . | . | . | . | . | . | . | . | . | . | . | . | . | . | . | . | . | . | . | . |
| . | . | . | . | . | . | . | . | . | . | . | . | . | . | . | . | . | . | . | . | . | . | . | . | . | . |
| . | . | . | . | . | . | . | . | T | . | . | . | . | . | . | . | . | . | . | . | . | . | . | . | . | . |
| . | . | . | . | . | . | . | . | . | . | . | . | . | . | . | . | . | . | . | . | . | . | . | . | . | . |
| . | . | . | . | . | . | . | . | . | . | . | . | . | . | . | . | . | . | . | . | . | . | . | . | . | . |
| . | . | . | . | . | . | . | . | . | . | . | T | . | . | . | . | . | . | . | . | . | . | . | . | . | . |
| . | . | . | . | . | . | . | . | T | . | . | . | . | . | . | . | . | . | . | . | . | . | . | . | . | . |
| . | . | . | . | . | . | . | . | T | . | . | . | . | . | . | . | . | . | . | . | . | . | . | . | . | . |
| . | . | . | . | . | . | . | . | T | . | . | . | . | . | . | . | . | . | . | . | . | . | . | . | . | . |
| . | . | . | . | . | . | . | . | T | . | . | . | . | . | . | . | . | . | . | . | . | . | . | . | . | . |
| . | . | . | . | . | . | . | . | . | . | . | . | . | . | . | . | . | . | . | . | . | . | . | . | . | . |
| . | . | . | . | . | . | . | . | T | . | . | . | . | . | . | . | . | . | . | . | . | . | . | . | . | . |
| . | . | . | . | . | . | . | . | . | . | . | . | . | . | . | . | . | . | . | . | . | . | . | . | . | . |
| . | . | . | . | . | . | . | . | . | . | . | . | . | . | . | . | . | . | . | . | . | . | . | . | . | . |

| 238 | 239 | 240 | 241 | 242 | 243 | 244 | 245 | 246 | 247 | 248 | 249 | 250 | 251 | 252 | 253 | 254 | 255 | 256 | 257 | 258 | 259 | 260 | 261 | 262 | 263 |
|---|---|---|---|---|---|---|---|---|---|---|---|---|---|---|---|---|---|---|---|---|---|---|---|---|---|
| N | N | N | N | N | N | N | N | N | N | N | N | N | N | N | N | N | N | N | N | NC | NC | NC | 3'UTR | 3'UTR | 3'UTR |
| 28289 | 28311 | 28312 | 28326 | 28371 | 28396 | 28688 | 28795 | 28854 | 28878 | 28881 | 28882 | 28883 | 28948 | 29039 | 29188 | 29197 | 29236 | 29451 | 29474 | 29543 | 29555 | 29557 | 29722 | 29734 | 29742 |
| C | C | C | G | G | G | T | A | C | G | G | G | G | C | A | A | C | C | C | G | G | C | G | C | G | G |
| . | . | . | . | . | . | . | . | . | . | . | . | . | . | . | . | . | . | . | . | . | . | . | . | . | . |
| . | . | . | . | . | . | . | . | . | . | . | . | . | . | . | . | . | . | . | . | . | . | . | . | . | . |
| . | . | . | . | . | . | . | . | . | . | . | . | . | . | . | . | . | . | . | . | . | . | . | . | . | . |
| . | . | . | . | . | . | . | . | . | . | . | . | . | . | . | . | . | . | . | . | . | . | . | . | . | . |
| . | . | . | . | . | . | . | . | . | . | . | . | . | . | . | . | . | . | . | . | . | . | . | . | . | . |
| . | . | . | . | . | . | . | . | . | . | . | . | . | . | . | . | . | . | . | . | . | . | . | . | . | . |
| . | . | . | . | . | . | . | . | . | . | . | . | . | . | . | . | . | . | . | . | . | . | . | . | . | . |
| . | . | . | . | . | . | . | . | . | . | . | . | . | . | . | . | . | . | . | . | . | . | . | . | . | . |
| . | . | . | . | . | . | . | . | . | . | . | . | . | . | . | . | . | . | . | . | . | . | . | . | . | . |
| . | . | . | . | . | . | . | . | . | . | . | . | . | . | . | . | . | . | . | . | . | . | . | . | . | . |
| . | . | . | . | . | . | . | . | . | . | . | . | . | . | . | . | . | . | . | . | . | . | . | . | . | . |
| . | . | . | . | . | . | . | . | . | . | . | . | . | . | . | . | . | . | . | . | . | . | . | . | . | . |
| . | . | . | . | . | . | . | . | . | . | . | . | . | . | . | . | . | . | . | . | . | . | . | . | . | . |
| . | . | . | . | . | . | . | . | . | . | . | . | . | . | . | . | . | . | . | . | . | . | . | . | . | . |
| . | . | . | . | . | . | . | . | . | . | A | A | C | . | . | . | . | . | . | . | . | . | . | . | . | . |
| . | . | . | . | . | . | . | . | . | . | A | A | C | . | . | . | . | . | . | . | . | . | . | . | . | . |
| . | . | . | . | . | . | C | . | . | . | . | . | . | . | . | . | . | . | . | . | . | . | . | . | . | T |
| . | . | . | . | . | . | C | . | . | . | . | . | . | . | . | . | . | . | . | . | . | . | . | . | . | T |
| . | . | . | . | . | . | C | . | . | . | . | . | . | . | . | . | . | . | . | . | . | . | . | . | . | T |
| . | . | . | . | . | . | C | . | . | . | . | . | . | . | . | . | . | . | . | . | . | . | . | . | . | T |
| . | . | . | . | . | . | C | . | . | . | . | . | . | . | . | . | . | . | . | . | . | . | . | . | . | T |
| . | . | . | . | . | . | C | . | . | . | . | . | . | . | . | . | . | . | . | . | . | . | . | . | . | T |
| . | . | . | . | . | . | C | . | . | . | . | . | . | . | . | . | . | . | . | . | . | . | . | . | . | T |
| . | . | . | . | . | . | C | . | . | . | . | . | . | . | . | . | . | . | . | . | . | . | . | . | . | T |
| . | . | . | . | . | . | C | . | . | . | . | . | . | . | . | . | . | . | . | . | . | . | . | . | . | T |
| . | . | . | . | . | . | C | . | . | . | . | . | . | . | . | . | . | . | . | . | . | . | . | . | . | T |
| . | . | . | . | . | . | C | . | . | . | . | . | . | . | . | . | . | . | . | . | . | . | . | . | . | T |
| . | . | . | . | . | . | C | . | . | . | . | . | . | . | . | . | . | . | . | . | . | . | . | . | . | T |
| . | . | . | . | . | . | . | . | . | . | . | . | . | . | . | . | . | . | . | . | . | . | . | . | . | . |
| . | . | . | . | . | . | C | . | . | . | . | . | . | . | . | . | . | . | . | . | . | . | . | . | . | T |
| . | . | . | . | . | . | . | . | . | . | . | . | . | . | . | . | . | . | . | . | . | . | . | . | . | . |
| . | . | . | . | . | . | . | . | . | . | . | . | . | . | . | . | . | . | . | . | . | . | . | . | . | . |
| . | . | . | . | . | . | . | . | . | . | A | A | C | . | . | . | . | . | . | . | . | . | . | . | . | . |
| . | . | . | . | . | A | . | . | . | . | . | . | . | . | . | . | . | . | T | . | . | . | . | . | . | . |
| . | . | . | . | . | A | . | . | . | . | . | . | . | . | . | . | . | . | T | . | . | . | . | . | . | . |
| . | . | . | . | . | . | . | . | . | . | A | A | C | . | . | . | . | . | . | . | . | . | . | . | . | . |
| . | . | . | . | . | . | . | ? | ? | ? | ? | ? | ? | ? | . | . | . | . | . | . | . | . | . | . | . | . |
| . | . | . | . | . | . | . | . | . | . | A | A | C | . | . | . | . | . | . | . | . | . | . | . | . | . |
| . | . | . | . | . | . | . | . | . | . | A | A | C | . | . | . | . | . | . | . | . | . | . | . | . | . |
| . | T | . | . | . | . | . | . | . | . | . | . | . | . | . | . | . | . | . | . | . | . | . | . | . | . |
| . | T | . | . | . | . | . | . | . | . | . | . | . | . | . | . | . | . | . | . | . | . | . | . | . | . |
| . | . | . | . | . | . | . | ? | ? | ? | ? | ? | ? | ? | . | . | . | . | . | . | . | . | . | - | - | - |
| . | T | . | . | . | . | . | . | . | . | . | . | . | . | . | . | . | . | . | . | . | . | . | . | . | . |
| . | T | . | . | . | . | . | . | . | . | . | . | . | . | . | . | . | . | . | . | . | . | . | . | . | . |
| . | . | . | . | . | . | . | . | . | . | A | A | C | . | . | . | . | . | . | . | . | . | . | . | . | . |
| . | . | . | . | . | . | . | . | . | . | A | A | C | . | . | . | . | . | . | . | . | . | . | . | . | . |
| . | . | . | . | . | . | . | . | . | . | . | . | . | . | . | . | . | . | . | . | . | . | . | . | . | . |
| . | . | . | . | . | . | . | . | . | . | . | . | . | . | . | . | . | . | . | . | . | . | . | . | . | . |
| . | . | . | . | . | . | . | . | . | . | A | A | C | . | . | . | . | . | . | . | . | . | . | . | . | . |
| . | . | . | . | . | . | . | . | . | . | . | . | . | . | . | . | . | . | . | . | . | . | . | . | . | . |
| . | . | . | . | . | . | . | . | . | . | . | . | . | . | . | . | . | . | . | . | . | . | . | . | . | . |
| . | T | . | . | . | . | . | . | . | . | . | . | . | . | . | . | . | . | . | . | . | . | . | . | . | . |
| . | . | . | . | . | . | . | . | . | . | A | A | C | . | . | . | . | . | . | . | . | . | . | . | . | . |
| . | . | . | . | . | . | . | . | . | . | . | . | . | . | . | . | . | . | . | . | . | . | . | . | . | . |
| . | . | . | . | . | . | . | . | . | . | . | . | . | . | . | . | . | . | . | . | . | . | . | . | . | . |

| | | | | | | | | | | | | | | | | | | | | | | | | | |
|---|---|---|---|---|---|---|---|---|---|---|---|---|---|---|---|---|---|---|---|---|---|---|---|---|---|
| . | . | . | . | T | . | . | . | . | . | . | . | . | . | . | . | . | . | . | . | . | . | . | . | . | . |
| . | . | . | . | T | . | . | . | . | . | . | . | . | . | . | . | . | . | . | . | . | . | . | . | . | . |
| . | . | . | . | . | . | . | . | . | . | . | . | . | . | . | . | . | . | . | . | . | . | . | . | . | . |
| . | . | . | . | . | . | . | . | . | . | . | . | . | . | . | . | . | . | . | . | . | . | . | . | . | . |
| . | . | . | . | . | . | . | . | T | . | . | . | . | . | . | . | . | . | . | . | . | . | . | . | . | . |
| . | . | . | . | . | . | . | . | . | . | . | . | . | . | . | . | . | . | . | . | . | . | . | . | . | . |
| . | . | . | . | . | . | . | . | . | . | . | . | . | . | . | . | . | . | . | . | . | . | . | . | . | . |
| . | . | . | . | . | . | . | . | . | . | . | . | . | . | . | . | . | . | . | . | . | . | . | . | . | . |
| . | . | . | . | . | . | . | . | ? | . | ? | ? | ? | . | . | . | . | . | . | . | . | . | . | . | . | . |
| . | . | . | . | . | . | . | . | . | . | . | . | . | . | . | . | . | . | . | . | . | c | g | . | . | . |
| . | . | . | . | . | . | . | . | . | . | . | . | . | . | . | . | . | . | . | . | . | . | . | . | . | . |
| . | . | . | . | . | . | . | . | T | . | . | . | . | . | . | . | . | . | . | . | . | . | . | . | . | . |
| . | . | . | . | . | . | . | . | T | . | . | . | . | . | . | . | . | . | . | . | . | . | . | . | . | . |
| . | . | . | . | . | . | . | . | T | . | . | . | . | . | . | . | . | . | . | . | . | . | . | . | . | . |
| . | . | . | . | . | . | . | . | T | . | . | . | . | . | . | . | . | . | . | . | . | . | . | . | . | . |
| . | . | . | . | . | . | . | . | T | . | . | . | . | . | . | . | . | . | . | . | . | . | . | . | . | . |
| . | . | . | . | . | . | . | . | . | . | A | A | C | . | . | . | . | . | . | . | . | . | . | . | . | . |
| . | . | . | . | . | . | . | . | . | . | A | A | C | . | . | . | . | . | . | . | . | . | . | . | g | g |
| . | . | . | . | . | . | . | . | . | . | A | A | C | . | . | . | . | . | . | . | . | . | . | c | g | g |
| . | T | . | . | . | . | . | . | . | . | . | . | . | . | . | . | . | . | . | . | . | . | . | . | . | . |
| . | Y | . | . | . | . | . | . | . | . | . | . | . | . | . | . | . | . | . | . | . | . | . | . | . | . |
| . | Y | . | . | . | . | . | . | . | . | . | . | . | . | . | . | . | . | . | . | . | . | . | . | . | . |
| . | T | . | . | . | . | . | . | . | . | . | . | . | . | . | . | . | . | . | . | . | . | . | . | . | . |
| . | T | . | . | . | . | . | . | . | . | . | . | . | . | . | . | . | . | . | . | . | . | . | . | . | . |
| . | T | . | . | . | . | . | . | . | . | . | . | . | . | . | . | . | . | . | . | . | . | . | . | . | . |
| . | T | . | . | . | . | . | . | . | . | . | . | . | . | . | . | . | . | . | . | . | . | . | . | . | . |
| . | T | . | . | . | . | . | . | . | . | . | . | . | . | . | . | . | . | . | . | . | . | . | . | . | . |
| . | T | . | . | . | . | . | . | . | . | . | . | . | . | . | . | . | . | . | . | . | . | . | . | . | . |
| . | T | . | . | . | . | . | . | . | . | . | . | . | . | . | . | . | . | . | . | . | . | . | . | . | . |
| . | T | . | . | . | . | . | . | . | . | . | . | . | . | . | . | . | . | . | . | . | . | . | . | . | . |
| . | T | . | . | . | . | . | . | . | . | . | . | . | . | . | . | . | . | . | . | . | . | . | . | . | . |
| . | T | . | . | . | . | . | . | . | . | . | . | . | . | . | . | . | . | . | . | . | . | . | . | . | . |
| . | T | . | . | . | . | . | . | . | . | . | . | . | . | . | . | . | . | . | . | . | . | . | . | . | g |
| . | T | . | . | . | . | . | . | . | . | . | . | . | . | . | . | . | . | . | . | . | . | . | . | . | . |
| . | T | . | . | . | . | . | . | . | . | . | . | . | . | . | . | . | . | . | . | . | . | . | . | . | . |
| . | T | . | . | . | . | . | . | . | . | . | . | . | . | . | . | . | . | . | . | . | . | . | . | . | . |
| . | T | . | . | . | . | . | . | . | . | . | . | . | . | . | . | . | . | . | . | . | . | . | . | . | . |
| . | T | . | . | . | . | . | . | . | . | . | . | . | . | . | . | . | . | . | . | . | . | . | . | . | . |
| . | T | . | . | . | . | . | . | . | . | . | . | . | . | . | . | . | . | . | . | . | . | . | . | . | . |
| . | T | . | . | . | . | . | . | . | . | . | . | . | . | . | . | . | . | . | . | . | . | . | . | . | . |
| . | T | . | . | . | . | . | . | . | . | . | . | . | . | . | . | . | . | . | . | . | . | . | . | . | . |
| . | . | . | . | . | . | C | . | . | . | . | . | . | . | . | . | . | . | . | . | . | . | . | . | . | T |
| . | . | . | . | . | . | Y | . | . | . | . | . | . | . | . | . | . | . | . | . | . | . | . | . | . | T |
| . | . | . | . | . | . | ? | . | . | . | . | . | . | . | . | . | . | . | . | . | . | . | . | . | . | T |
| . | . | . | . | . | . | C | . | . | . | . | . | . | . | . | . | . | . | . | . | . | . | . | . | . | T |
| . | . | . | . | . | . | Y | . | . | . | . | . | . | . | . | . | . | . | . | . | . | . | . | . | . | T |
| . | . | . | . | . | . | Y | . | . | . | . | . | . | . | . | . | . | . | . | . | . | . | . | . | . | T |
| . | . | . | . | . | . | . | . | . | . | . | . | . | . | . | . | . | . | . | . | . | . | . | . | . | . |
| . | . | . | . | . | . | . | . | . | . | . | . | . | . | . | . | . | . | . | . | . | . | . | - | - | - |
| . | . | . | . | . | . | . | . | . | . | . | . | . | . | . | . | . | . | . | . | . | . | . | . | . | . |
| . | . | . | . | . | . | . | . | . | . | . | . | . | . | . | . | . | . | . | . | . | . | . | . | . | . |
| . | T | . | . | . | . | . | . | . | . | . | . | . | . | . | . | . | . | . | . | . | . | . | . | . | . |
| . | . | . | . | . | . | . | . | . | . | A | A | C | . | . | . | . | . | . | . | . | . | . | . | g | g |
| . | ? | . | . | . | . | . | . | . | . | . | . | . | . | . | . | . | . | . | . | . | T | . | . | . | . |
| . | T | . | . | . | . | . | . | . | . | . | . | . | . | . | . | . | . | . | . | . | . | . | . | . | . |
| . | T | . | . | . | . | . | . | . | . | . | . | . | . | . | . | . | . | . | . | . | . | . | . | . | . |
| . | T | . | . | . | . | . | . | . | . | . | . | . | . | . | . | . | . | . | . | . | . | . | . | . | . |
| . | T | . | . | . | . | . | . | . | . | . | . | . | . | . | . | . | . | . | . | . | . | . | . | . | . |
| . | T | . | . | . | . | . | . | . | . | . | . | . | . | . | . | . | . | . | . | . | . | . | . | . | . |
| . | Y | . | . | . | . | . | . | . | . | . | . | . | . | . | . | . | . | . | . | . | T | . | . | . | . |
| . | T | . | . | . | . | . | . | . | . | . | . | . | . | . | . | . | . | . | . | . | . | . | . | . | . |
| . | T | . | . | . | . | . | . | . | . | . | . | . | . | . | . | . | . | . | . | . | . | . | . | . | . |

| | | | | | | | | | | | | | | | | | | | | | | | | | |
|---|---|---|---|---|---|---|---|---|---|---|---|---|---|---|---|---|---|---|---|---|---|---|---|---|---|
| . | T | . | . | . | . | . | . | . | . | . | . | . | . | . | . | . | . | . | . | . | . | . | . | . | . |
| . | T | . | . | . | . | . | . | . | . | . | . | . | . | . | . | . | . | . | . | . | . | . | . | . | . |
| . | T | . | . | . | . | . | . | . | . | . | . | . | . | . | . | . | . | . | . | . | . | . | . | . | . |
| . | Y | . | . | . | . | . | . | . | . | . | . | . | . | . | . | . | . | . | . | . | . | . | . | . | . |
| . | T | . | . | . | . | . | . | . | . | . | . | . | . | . | . | . | . | . | . | . | . | . | . | . | . |
| . | T | . | . | . | . | . | . | . | . | . | . | . | . | . | . | . | . | . | . | . | . | . | c | g | g |
| . | T | . | . | . | . | . | . | . | . | . | . | . | . | . | . | . | . | . | . | . | . | . | . | . | . |
| . | T | . | . | . | . | . | . | . | . | . | . | . | . | . | . | . | . | . | . | . | . | . | . | . | . |
| . | Y | . | . | . | . | . | . | . | . | g | . | . | . | . | . | . | . | . | . | . | . | . | . | . | . |
| . | T | . | . | . | . | . | . | . | . | . | . | . | . | . | . | . | . | . | . | . | . | . | . | . | . |
| . | Y | . | . | . | . | . | . | . | . | . | . | . | . | . | . | . | . | . | . | . | . | . | . | . | . |
| . | Y | . | . | . | . | . | . | . | . | . | . | . | . | . | . | . | . | . | . | . | . | . | . | . | . |
| . | T | . | . | . | . | . | . | . | . | . | . | . | . | . | . | . | . | . | . | . | . | . | . | . | . |
| . | T | . | . | . | . | . | . | . | . | . | . | . | . | . | . | . | . | . | . | . | . | . | . | . | . |
| . | Y | . | . | . | . | . | . | . | . | . | . | . | . | . | . | . | . | . | . | . | . | . | . | . | . |
| . | T | . | . | . | . | . | . | . | . | . | . | . | . | . | . | . | . | . | . | . | . | . | . | . | . |
| . | T | . | . | . | . | . | . | . | . | . | . | . | . | . | a | c | c | . | . | . | . | . | . | . | . |
| . | T | . | . | . | . | . | . | . | . | . | . | . | . | . | . | . | . | . | . | . | . | . | . | . | . |
| . | T | . | . | . | . | . | . | . | . | . | . | . | . | . | . | . | . | . | . | . | . | . | . | . | . |
| c | T | . | . | . | . | . | . | . | . | . | . | . | . | . | . | . | . | . | . | . | . | . | . | . | . |
| . | T | . | . | . | . | . | . | . | . | . | . | . | . | . | . | . | . | . | . | . | . | . | . | . | . |
| . | . | . | . | . | . | . | a | c | g | g | g | g | c | . | . | . | . | . | . | . | . | . | . | . | . |
| c | c | . | . | . | . | . | . | . | . | . | . | . | . | . | . | . | . | . | . | . | . | . | . | . | . |
| . | . | . | . | . | . | . | . | . | . | . | . | . | . | . | . | . | . | . | . | . | . | . | . | . | . |
| . | . | . | . | . | . | . | . | . | . | . | . | . | . | . | . | . | . | . | . | . | . | . | . | . | . |
| . | . | . | . | . | . | . | . | . | . | . | . | . | . | . | . | . | . | . | . | . | . | . | c | g | g |
| . | Y | . | . | . | . | . | . | . | . | . | . | . | . | . | . | . | . | . | . | . | . | . | . | . | . |
| . | . | . | . | . | . | . | . | . | . | . | . | . | . | . | . | . | . | . | . | . | . | . | . | . | . |
| . | . | . | . | . | . | . | . | . | . | . | . | . | . | . | . | . | . | . | . | . | . | . | . | . | . |
| . | . | . | . | . | . | . | . | . | . | . | . | . | . | . | . | . | . | . | . | . | . | . | . | . | . |
| . | . | . | . | . | . | . | . | . | . | . | . | . | . | . | . | . | . | . | . | . | . | . | . | . | . |
| . | Y | . | . | . | . | . | a | c | g | g | g | g | . | . | . | . | . | . | . | . | . | . | . | . | . |
| . | T | . | . | . | . | . | . | . | . | . | . | . | . | . | . | . | . | . | . | . | . | . | . | . | . |
| . | . | . | . | . | . | . | . | . | . | . | . | . | . | . | . | . | . | . | . | . | . | . | . | . | . |
| . | . | . | . | . | . | . | . | . | A | . | . | . | . | . | . | . | . | . | . | . | . | . | . | . | A |
| . | T | . | . | . | . | . | . | . | . | . | . | . | . | . | . | . | . | . | . | . | . | . | . | . | . |
| . | T | . | . | . | . | . | . | . | . | . | . | . | . | . | . | . | . | . | . | . | . | . | . | . | . |
| . | . | . | . | . | . | . | . | T | . | . | . | . | . | . | . | . | . | . | . | . | . | . | . | . | . |
| . | . | . | . | . | . | . | . | T | . | . | . | . | . | . | . | . | . | . | . | . | . | . | . | . | . |
| . | . | . | . | . | . | . | . | . | . | . | . | . | . | . | . | . | . | . | . | . | . | . | . | . | . |
| . | . | . | . | . | . | . | . | . | . | . | . | . | . | . | . | . | . | . | . | . | . | . | . | . | . |
| . | . | . | . | . | . | . | . | . | . | . | . | . | . | . | . | . | . | . | . | . | . | . | . | . | . |
| . | . | . | . | . | . | . | . | . | . | . | . | . | . | . | . | . | . | . | . | . | . | . | . | . | . |
| . | . | . | . | . | . | . | . | . | . | . | . | . | . | . | . | . | . | . | . | . | . | . | . | . | . |
| . | . | . | . | . | . | . | . | T | . | . | . | . | . | . | . | . | . | . | . | . | . | . | . | . | . |
| . | T | . | . | . | . | . | . | . | . | . | . | . | . | . | . | . | . | . | . | . | . | . | . | . | . |
| . | T | . | . | . | . | . | . | . | . | . | . | . | . | . | . | . | . | . | . | . | . | . | . | . | . |
| . | . | . | . | . | . | . | . | . | . | . | . | . | . | . | . | . | . | . | . | . | . | . | . | . | . |
| . | . | . | . | . | . | . | . | . | . | . | . | . | . | . | . | . | . | . | . | . | . | . | . | . | . |
| . | T | . | . | . | . | . | . | . | . | . | . | . | . | . | . | . | . | . | . | . | . | . | . | . | . |
| . | T | . | . | . | . | . | . | . | . | . | . | . | . | . | . | . | . | . | . | . | . | . | . | . | . |
| . | . | . | . | . | . | . | . | . | . | . | . | . | . | . | . | . | . | . | . | . | . | . | . | . | . |
| . | . | . | . | . | . | . | . | . | . | . | . | . | . | . | . | . | . | . | . | . | . | . | . | . | . |
| . | . | . | . | . | . | . | . | . | . | . | . | . | . | . | . | . | . | . | . | . | . | . | . | . | . |
| . | . | . | . | . | . | . | . | T | . | . | . | . | . | . | . | . | . | . | . | . | . | . | . | . | . |
| . | . | . | . | . | . | . | . | . | . | . | . | . | . | . | . | . | . | . | . | . | . | . | . | . | . |
| . | . | . | . | . | . | . | . | T | . | . | . | . | . | . | . | . | . | . | . | . | . | . | . | . | . |
| . | . | . | . | . | . | . | . | . | . | . | . | . | . | . | . | . | . | . | . | . | . | . | . | . | . |
| . | . | . | . | . | . | . | . | . | . | . | . | . | . | . | . | . | . | . | . | . | . | . | . | . | . |
| . | . | . | . | . | . | . | . | T | . | . | . | . | . | . | . | . | . | . | . | . | . | . | . | . | . |
| . | . | . | . | . | . | . | . | T | . | . | . | . | . | . | . | . | . | . | . | . | . | . | . | . | . |
| . | . | . | . | . | . | . | . | T | . | . | . | . | . | . | . | . | . | . | . | . | . | . | . | . | . |

```
. . . . . . . . . . . . . . . . . . . . . . . . . .
. . . . . . . . . . . . . . . . . . . . . . . . . .
. . . . . . . . T . . . . . . . . . . . . . . . . .
. . . . . . . . . . . . . . . . . . . . . . . . . .
. . . . . . . . . . . . . . . . . . . . . . . . . .
. . . . . . . . T . . . . . . . . . . . . . . . . .
. . . . . . . . . . . . . . . . . . . . . . . . . .
. . . . . . . . . . . . . . T . . . . . . . . . . .
. . . . . . . . . . . . . . . . . . . . . . . . . .
. . . . . . . . T . . . . . . . . . . . . . . . . .
. . . . . . . . . . . . . . . . . . . . . . . . . .
. . . . . . . . . . . . . . . . . . . . . . . . . .
. . . . . . . . . . . . . . . . . . . . . . . . . .
. . . . . . . . . . . . . . T . . . . . . . . . . .
. . . . . . . . . . . . . . . . . . . . . . . . . .
. T . . . . . . . . . . . . . . . . . . . . . . . .
. . . . . . . . . . . . . . . . . . . . . . . . . .
. . . . . . . . . . . . . . . . . . . . . . . . . .
. . . . . . . . . . . . . . . . . . . . . . . . . .
. . . . . . . . T . . . . . . . . . . . . . . . . .
. . . . . . . . . . . . . . . . . . . . . . . . . .
. . . . . . . . . . . . . . . . . . . . . . . . . .
. . . . . . . . . . . . . . . . . . . . . . . . . .
. . . . . . . . T . . . . . . . . . . . . . . . . .
. . . . . . . . T . . . . . . . . . . . . . . . . .
. . . . . . . . T . . . . . . . . . . . . . . . . .
. . . . . . . . . . . . . . . . . . . . . . . . . .
. . . . . . . . . . . . . . . . . . . . . . . . . .
. . . . . . . . . . . . . . . . . . . . . . . . . .
. . . . . . . . T . . . . . . . . . . . . . . . . .
. . . . . . . . T . . . . . . . . . . . . . . . . .
. . . . . . . . . . . . . . . . . . . . . . . . . .
. . . . . . . . . . . . . . . . . . . . . . . . . ?
. . . . . . . . T . . . . . . . . . . . . . . . . .
. . . . . . . . T . . . . . . . . . . . . . . . . .
. . . . . . . . T . . . . . . . . . . . . . . . . .
. . . . . . . . T . . . . . . . . . . . . . . . . .
. . . . . . . . . . . . . . . . . . . . . . . . . .
. . . . . . . . T . . . . . . . . . . . . . . . . .
. . . . . . . . T . . . . . . . . . . . . . . . . .
. . . . . . . . T . . . . . . . . . . . . . . . . .
. . . . . . . . T . . . . . . . . . . . . . . . . .
. . . . . . . . . . . . . . . . . . . . . . . . . .
. . . . . . . . . . . . . . T . . . . . . . . . . .
. . . . . . . . . . . . . . . . . . . . . . . . . .
. . . . . . . . . . . . . . . . . . . . . . . . . .
. . . . . . . . . . . . . . . . . . . ? . . . . . .
. . . . . . . . . . . . . . . . . . . ? . . . . . .
. . . . . . . . . . . . . . . . . . . . . . . . . .
. . . . . . . . . . . . . . . . . . . . . . . . . .
. . . . . . . . . . . . . . . . . . . . . . . . . .
. . . . . . . . . . . . . . . . . . . . . . . . . .
. . . . . . . . . . . . . . . . . . . . . . . . . .
. . . . . . . . . . . . . . . . . . . . . . . . . .
. . . . . . . . . . . . . . . . . . . . . . . . . .
. . . . . . . . . . . . . . . . . . . . . . . . . .
. . . . . . . . T . . . . . . . . . . . . . . . . .
. . . . . . . . . . . . . . . . . . . . . . . . . .
. . . . . . . . T . . . . . . . . . . . . . . . . .
. . . . . . . . . . . . . . . . . . . . . . . . . .
. . . . . . . . . . . . . . . . . . . . . . . . . .
. . . . . . . . . . . . . . . . . . . . . . . . . .
```

| | | | | | | | | | | | | | | | | | | | | | | | | | |
|---|---|---|---|---|---|---|---|---|---|---|---|---|---|---|---|---|---|---|---|---|---|---|---|---|---|
| . | . | . | . | . | . | . | . | . | . | . | . | . | T | . | . | . | . | . | . | . | . | . | . | . | . |
| . | . | . | . | . | . | . | . | T | . | . | . | . | . | . | . | . | . | . | . | . | . | . | . | . | . |
| . | . | . | . | . | . | . | . | . | . | . | . | . | . | . | . | . | . | . | . | . | . | . | . | . | . |
| . | . | . | . | . | . | . | . | . | . | . | . | . | . | . | . | . | . | . | . | . | . | . | . | . | . |
| . | T | . | . | . | . | . | . | . | . | . | . | . | . | . | . | . | . | . | . | . | . | . | T | . | . |
| . | T | . | . | . | . | . | . | . | . | . | . | . | . | . | . | . | . | . | . | . | . | . | . | . | . |
| . | T | . | . | . | . | . | . | . | . | . | . | . | . | . | . | . | . | . | . | . | . | . | . | . | . |
| . | T | . | . | . | . | . | . | . | . | . | . | . | . | . | . | . | . | . | . | . | . | . | . | . | . |
| . | T | . | . | . | . | . | . | . | . | . | . | . | . | . | . | . | . | . | . | . | . | . | . | . | . |
| . | T | . | . | . | . | . | . | . | . | . | . | . | . | . | . | . | . | . | . | . | . | . | . | . | . |
| . | T | . | . | . | . | . | . | . | . | . | . | . | . | . | . | . | . | . | . | . | . | . | . | . | . |
| . | T | . | . | . | . | . | . | . | . | . | . | . | . | . | . | . | . | . | . | . | . | . | . | . | . |
| . | T | . | . | . | . | . | . | . | . | . | . | . | . | . | . | . | . | . | . | . | . | . | . | . | . |
| . | T | . | . | . | . | . | . | . | . | . | . | . | . | . | . | . | . | . | . | . | . | . | . | . | . |
| . | T | . | . | . | . | . | . | . | . | . | . | . | . | . | . | . | . | . | . | . | . | . | T | . | . |
| . | T | . | . | . | . | . | . | . | . | . | . | . | . | . | . | . | . | . | . | . | . | . | . | . | . |
| . | T | . | . | . | . | . | . | . | . | . | . | . | . | . | . | . | . | . | . | . | . | . | . | . | . |
| . | T | . | . | . | . | . | . | . | . | . | . | . | . | . | . | . | . | . | . | . | . | . | . | . | . |
| . | T | . | . | . | . | . | . | . | . | . | . | . | . | . | . | . | . | . | . | . | . | . | . | . | . |
| . | T | . | . | . | . | . | . | . | . | . | . | . | . | . | . | . | . | . | . | . | . | . | . | . | . |
| . | T | . | . | . | . | . | . | . | . | . | . | . | . | . | . | . | . | . | . | . | . | . | . | . | . |
| . | T | . | . | . | . | . | . | . | . | . | . | . | . | . | . | . | . | . | . | . | . | . | . | . | . |
| . | T | . | . | . | . | . | . | . | . | . | . | . | . | . | . | . | . | . | . | . | . | . | . | . | . |
| . | T | . | . | . | . | . | . | . | . | . | . | . | . | . | . | . | . | . | . | . | . | . | . | . | . |
| . | T | . | . | . | . | . | . | . | . | . | . | . | . | . | . | . | . | . | . | . | . | . | . | . | . |
| . | T | . | . | . | . | . | . | . | . | . | . | . | . | . | . | . | . | . | . | . | . | . | . | . | . |
| . | T | . | . | . | . | . | . | . | . | . | . | . | . | . | . | . | . | . | . | . | . | . | . | . | . |
| . | T | . | . | . | . | . | . | . | . | . | . | . | . | . | . | . | . | . | . | . | . | . | . | . | . |
| . | T | . | . | . | . | . | . | . | . | . | . | . | . | . | . | . | . | . | . | . | . | . | . | . | . |
| . | T | . | . | . | . | . | . | . | . | . | . | . | . | . | . | . | . | . | . | . | . | . | . | . | . |
| . | T | . | . | . | . | . | . | . | . | . | . | . | . | . | . | . | . | . | . | . | . | . | . | . | . |
| . | T | . | . | . | . | . | . | . | . | . | . | . | . | . | . | . | . | . | . | . | . | . | . | . | . |
| . | T | . | . | . | . | . | . | . | . | . | . | . | . | . | . | . | . | . | . | . | . | . | . | . | . |
| . | T | . | . | . | . | . | . | . | . | . | . | . | . | . | . | . | . | . | . | . | . | . | . | . | . |
| . | . | . | . | . | . | . | . | . | . | A | A | C | . | . | . | . | . | . | . | . | . | . | . | . | . |
| . | T | . | . | . | . | . | . | . | . | . | . | . | . | . | . | . | . | . | . | . | . | . | . | . | . |
| . | T | . | . | . | . | . | . | . | . | . | . | . | . | . | . | . | . | . | . | . | . | . | . | . | . |
| . | T | . | . | . | . | . | . | . | . | . | . | . | . | . | . | . | . | . | . | . | . | . | . | . | . |
| . | T | . | . | . | . | . | . | . | . | . | . | . | . | . | . | . | . | . | . | . | . | . | T | . | . |
| . | T | . | . | . | . | . | . | . | . | . | . | . | . | . | . | . | . | . | . | . | . | . | . | . | . |
| . | T | . | . | . | . | . | . | . | . | . | . | . | . | . | . | . | . | . | . | . | . | . | . | . | . |
| . | T | . | . | . | . | . | . | . | . | . | . | . | . | . | . | . | . | . | . | . | . | . | . | . | . |
| . | T | . | . | . | . | . | . | . | . | . | . | . | . | . | . | . | . | . | . | . | . | . | . | . | . |
| . | T | . | . | . | . | . | . | . | . | . | . | . | . | . | . | . | . | . | . | . | . | . | . | . | . |
| . | T | . | . | . | . | . | . | . | . | . | . | . | . | . | . | . | . | . | . | . | . | . | . | . | . |
| . | T | . | . | . | . | . | . | . | . | . | . | . | . | . | . | . | . | . | . | . | . | . | . | . | . |
| . | T | . | . | . | . | . | . | . | . | . | . | . | . | . | . | . | . | . | . | . | . | . | . | . | . |
| . | T | . | . | . | . | . | . | . | . | . | . | . | . | . | . | . | . | . | . | . | . | . | . | . | . |
| . | T | . | . | . | . | . | . | . | . | . | . | . | . | . | . | . | . | . | . | . | . | . | . | . | . |
| . | T | . | . | . | . | . | . | . | . | . | . | . | . | . | . | . | . | . | . | . | . | . | . | . | . |
| . | T | . | . | . | . | . | . | . | . | . | . | . | . | . | . | . | . | . | . | . | . | . | . | . | . |
| . | T | . | . | . | . | . | . | . | . | . | . | . | . | . | . | . | . | . | . | . | . | . | . | . | . |
| . | T | . | . | . | . | . | . | . | . | . | . | . | . | . | . | . | . | . | . | . | . | . | . | . | . |
| . | T | . | . | . | . | . | . | . | . | . | . | . | . | . | . | . | . | . | . | . | . | . | . | . | . |
| . | T | . | . | . | . | . | . | . | . | . | . | . | . | . | . | . | . | . | . | . | . | . | . | . | . |
| . | T | . | . | . | . | . | . | . | . | . | . | . | . | . | . | . | . | . | . | . | . | . | . | . | . |
| . | T | . | . | . | . | . | . | . | . | . | . | . | . | . | . | . | . | . | . | . | . | . | . | . | . |
| . | . | . | . | . | . | . | . | . | . | . | . | . | . | . | . | . | . | . | . | . | . | . | . | . | . |
| . | . | . | . | . | . | . | . | . | A | . | . | . | . | . | . | . | . | . | . | . | . | . | . | . | A |
| . | . | . | . | . | . | . | . | . | . | . | . | . | . | . | . | . | . | . | . | . | . | . | . | . | . |
| . | T | . | . | . | . | . | . | . | . | . | . | . | . | . | . | . | . | . | . | . | . | . | . | . | . |

| | | | | | | | | | | | | | | | | | | | | | | | | | |
|---|---|---|---|---|---|---|---|---|---|---|---|---|---|---|---|---|---|---|---|---|---|---|---|---|---|
| . | T | . | . | . | . | . | . | . | . | . | . | . | . | . | . | . | . | . | . | . | . | . | . | . | . |
| . | T | . | . | . | . | . | . | . | . | . | . | . | . | . | . | . | . | . | . | . | . | . | . | . | . |
| . | T | . | . | . | . | . | . | . | . | . | . | . | . | . | . | . | . | . | . | . | . | . | . | . | . |
| . | T | . | . | . | . | . | . | . | . | . | . | . | . | . | . | . | . | . | . | . | . | . | . | . | . |
| . | T | . | . | . | . | . | . | . | . | . | . | . | . | . | . | . | . | . | . | . | . | . | . | . | . |
| . | T | . | . | . | . | . | . | . | . | . | . | . | . | . | . | . | . | . | . | . | . | . | . | . | . |
| . | . | . | . | . | . | . | . | . | . | . | . | . | . | . | . | . | . | . | . | . | . | . | . | . | . |
| . | . | . | . | . | . | . | . | . | . | . | . | . | . | . | . | . | . | . | . | . | . | . | . | . | . |
| . | . | . | . | . | . | . | . | T | . | . | . | . | . | . | . | . | . | . | . | . | . | . | . | . | . |
| . | T | . | . | . | . | . | . | . | . | . | . | . | . | . | . | . | . | . | . | . | . | . | . | . | . |
| . | . | . | . | . | . | . | . | . | . | A | A | C | . | . | . | . | . | . | . | . | . | . | . | . | . |
| . | . | . | . | . | . | . | . | . | . | A | A | C | . | . | . | . | . | . | . | . | . | . | . | . | . |
| . | T | . | . | . | . | . | . | . | . | . | . | . | . | . | . | . | . | . | . | . | . | . | . | . | . |
| . | . | . | . | . | . | . | . | . | . | . | . | . | . | . | . | . | . | . | . | . | . | . | . | . | . |
| . | . | . | . | . | . | . | . | . | . | . | . | . | . | . | . | . | . | . | . | . | . | . | . | . | . |
| . | . | . | . | . | . | . | . | . | A | . | . | . | . | . | . | . | . | . | . | . | . | . | . | . | A |
| . | . | . | . | . | . | . | . | . | A | . | . | . | . | . | . | . | . | . | . | . | . | . | . | . | A |
| . | . | . | . | . | . | . | . | T | . | . | . | . | . | . | . | . | . | . | . | . | . | . | . | T | . |
| . | . | . | . | . | . | . | . | . | . | . | . | . | . | . | . | . | . | . | . | . | . | . | . | . | . |
| . | . | . | . | . | . | . | . | T | . | . | . | . | . | . | . | . | . | . | . | . | . | . | . | T | . |
| . | . | . | . | . | . | . | . | T | . | . | . | . | . | . | . | . | . | . | . | . | . | . | . | T | . |
| . | . | . | . | . | . | . | . | T | . | . | . | . | . | . | . | . | . | . | . | . | . | . | . | . | . |
| . | T | . | . | . | . | . | . | . | . | . | . | . | . | . | . | . | . | . | . | . | . | . | . | . | . |
| . | . | . | . | . | . | . | . | . | . | . | . | . | . | . | . | . | . | . | . | . | . | . | . | . | . |
| . | T | . | . | . | . | . | . | . | . | . | . | . | . | . | . | . | . | . | . | . | . | . | . | . | . |
| . | . | . | . | . | . | . | . | . | . | . | . | . | . | . | . | . | . | . | . | . | . | . | . | . | . |
| . | . | . | . | . | . | . | . | . | . | . | . | . | . | . | . | . | . | . | . | . | . | . | . | . | . |
| . | T | . | . | . | . | . | . | . | . | . | . | . | . | . | . | . | . | . | . | . | . | . | . | . | . |
| . | . | . | . | . | . | . | . | . | . | . | . | . | . | . | . | . | . | . | . | . | . | . | . | . | . |
| . | T | . | . | . | . | . | . | . | . | . | . | . | . | . | . | . | . | . | . | . | . | . | . | . | . |
| . | . | . | . | . | . | . | . | T | . | . | . | . | . | . | . | . | . | . | . | . | . | . | . | . | . |
| . | . | . | . | . | . | . | . | . | . | . | . | . | . | . | . | . | . | . | . | . | . | . | . | . | . |
| A | T | . | . | . | . | . | . | . | . | . | . | . | . | . | . | . | . | . | . | . | . | . | . | . | . |
| c | c | c | g | g | g | t | a | c | g | a | a | c | c | a | a | c | c | c | g | g | c | g | c | g | g |
| c | c | c | g | g | g | t | a | c | g | a | a | c | c | a | a | c | c | c | g | g | c | g | c | g | g |
| c | c | c | g | g | g | t | a | c | g | g | g | g | c | a | a | c | c | c | g | g | c | g | c | g | g |
| c | c | c | g | g | g | t | a | c | g | g | g | g | c | a | a | c | c | c | g | g | c | g | c | g | g |
| c | c | c | g | g | g | t | a | c | g | a | a | c | c | a | a | c | c | c | g | g | c | g | c | g | g |
| c | c | c | g | g | g | t | a | c | g | a | a | c | c | a | a | c | c | c | g | g | c | g | c | g | g |
| c | c | c | g | g | g | t | a | c | g | a | a | c | c | a | a | c | c | c | g | g | c | g | c | g | g |
| c | c | c | g | g | g | t | a | c | g | a | a | c | c | a | a | c | c | c | g | g | c | g | c | g | g |
| c | c | c | g | g | g | t | a | c | g | a | a | c | c | a | a | c | c | c | g | g | c | g | c | g | g |
| c | c | c | g | g | g | t | a | c | g | g | g | g | c | a | a | c | c | c | g | g | c | g | c | g | g |
| c | c | c | g | g | g | t | a | c | a | g | g | g | c | a | a | c | c | c | g | g | c | g | c | g | a |
| c | c | c | g | g | g | t | a | c | a | g | g | g | c | a | a | c | c | c | g | g | c | g | c | g | a |
| c | c | c | g | g | g | t | a | c | a | g | g | g | c | a | a | c | c | c | g | g | c | g | c | g | a |
| c | c | c | g | g | g | t | a | c | g | a | a | c | c | a | a | c | c | c | g | g | c | g | c | g | g |
| c | t | c | g | g | g | t | a | c | g | g | g | g | c | a | a | c | c | c | g | g | c | g | c | g | g |
| c | t | c | g | g | g | t | a | c | g | g | g | g | c | a | a | c | c | c | g | g | c | g | c | g | g |
| c | c | c | g | g | g | t | a | c | g | a | a | c | c | a | a | c | c | c | g | g | c | g | c | g | g |
| c | c | c | g | g | g | t | a | c | g | a | a | c | c | a | a | c | c | c | g | g | c | g | c | g | g |
| c | c | c | g | g | g | t | a | c | g | a | a | c | c | a | a | c | c | c | g | g | c | g | c | g | g |
| c | c | c | g | g | g | c | g | c | g | g | g | g | c | a | a | c | c | c | g | g | c | g | c | g | t |
| c | c | c | g | g | g | t | a | c | g | g | g | g | c | a | a | c | c | c | g | g | c | g | c | g | g |
| c | t | c | g | g | g | t | a | c | g | g | g | g | c | a | a | c | c | c | g | g | c | g | c | g | g |
| c | c | c | g | g | g | t | a | c | g | a | a | c | c | a | a | c | c | c | g | g | c | g | c | g | g |
| c | c | c | g | g | g | t | a | c | g | g | g | g | c | a | a | c | c | c | g | g | c | g | c | g | g |
| . | . | . | . | . | . | . | . | . | . | . | . | . | . | . | . | . | . | . | . | . | . | . | . | . | . |
| . | . | . | . | . | . | . | . | . | . | . | . | . | . | . | . | . | . | . | . | . | . | . | . | . | . |
| . | . | . | . | . | . | . | . | . | . | . | . | . | . | . | . | . | . | . | . | . | . | . | . | . | . |
| . | T | . | . | . | . | . | . | . | . | . | . | . | . | . | . | . | . | . | . | . | . | . | . | . | . |
| . | . | . | . | . | . | . | . | . | . | . | . | . | . | . | . | . | . | . | . | . | . | . | . | . | . |

| | | | | | | | | | | | | | | | | | | | | | | | | | |
|---|---|---|---|---|---|---|---|---|---|---|---|---|---|---|---|---|---|---|---|---|---|---|---|---|---|
| . | T | . | . | . | . | . | . | . | . | . | . | . | . | . | . | . | . | . | . | . | . | . | . | . | . |
| . | . | . | . | . | . | . | . | . | . | A | A | C | . | . | . | . | . | . | . | . | . | . | . | . | . |
| . | . | . | . | . | . | C | G | . | ? | ? | ? | ? | . | . | . | . | . | . | . | . | . | . | . | . | T |
| . | . | . | . | . | . | C | G | . | . | . | . | . | . | . | . | . | . | . | . | . | . | . | . | . | T |
| . | . | . | . | . | . | . | . | . | . | . | . | . | . | . | . | . | . | . | . | . | . | . | . | . | . |
| . | . | . | . | . | . | C | . | . | . | . | . | . | . | . | . | . | . | . | . | . | . | . | . | . | T |
| . | . | . | . | . | . | . | . | . | . | . | . | . | . | . | . | . | . | . | . | . | . | . | . | . | . |
| . | . | . | . | . | . | . | . | . | A | . | . | . | . | . | . | . | . | . | . | . | . | . | . | . | A |
| . | . | . | . | . | . | . | . | . | . | . | . | . | . | . | . | . | . | . | . | . | . | . | . | . | . |
| . | . | . | . | . | . | . | . | . | . | . | . | . | . | . | . | . | . | . | . | . | . | . | . | C | . |
| . | . | . | . | . | . | . | . | . | A | . | . | . | . | . | . | . | . | . | . | . | . | . | ? | ? | ? |
| . | . | . | . | . | . | . | . | . | A | . | . | . | . | . | . | . | . | . | . | . | . | . | . | . | A |
| . | . | . | . | . | . | . | . | . | A | . | . | . | . | . | . | . | . | . | . | . | . | . | . | . | A |
| . | T | . | . | . | . | . | . | . | . | . | . | . | . | . | . | . | . | . | . | . | . | . | . | . | . |
| . | T | . | . | . | . | . | . | . | . | . | . | . | . | . | . | . | . | . | . | . | . | . | . | . | . |
| . | T | . | . | . | . | . | . | . | . | . | . | . | . | . | . | . | . | . | . | . | . | . | . | . | . |
| . | T | . | . | . | . | . | . | . | . | . | . | . | . | . | . | . | . | . | . | . | . | . | . | . | . |
| . | T | . | . | . | . | . | . | . | . | . | . | . | . | . | . | . | . | . | . | . | . | . | . | . | . |
| . | T | . | . | . | . | . | . | . | . | . | . | . | . | . | . | . | . | . | . | . | . | . | . | . | . |
| . | . | . | . | . | . | . | . | . | . | . | . | . | . | . | . | . | . | . | . | . | . | . | . | . | . |
| . | T | . | . | . | . | . | . | . | . | . | . | . | . | . | . | . | . | . | . | . | . | . | . | . | . |
| . | T | . | . | . | . | . | . | . | . | . | . | . | . | . | . | . | . | . | . | . | . | . | . | . | . |
| . | . | . | . | . | . | . | . | . | . | A | A | C | . | . | . | . | . | . | . | . | . | . | . | . | . |
| . | . | . | . | . | . | . | . | . | . | . | . | . | . | . | . | . | . | . | . | . | . | . | . | . | . |
| . | . | . | . | . | . | . | . | . | . | A | A | C | . | . | . | . | . | . | . | . | . | . | . | . | . |
| . | . | . | . | . | . | . | . | . | . | A | A | C | . | . | . | . | . | . | . | . | . | . | . | . | . |
| . | . | . | . | . | . | . | . | . | . | A | A | C | . | . | . | . | . | . | . | . | . | . | . | . | . |
| . | . | . | . | . | . | . | . | . | . | . | . | . | . | . | . | . | . | . | . | . | . | . | . | . | . |
| . | . | . | . | . | . | C | . | . | . | A | A | C | . | . | . | . | . | . | . | . | . | . | . | . | . |
| . | T | . | . | . | . | . | . | . | . | . | . | . | . | . | . | . | . | . | . | . | . | . | . | . | . |
| . | . | . | . | . | . | . | . | . | . | . | . | . | . | . | . | . | . | . | . | . | . | . | . | . | . |
| . | . | . | . | . | . | . | . | . | . | . | . | . | . | . | . | . | . | . | . | . | . | . | . | . | . |
| . | T | . | . | . | . | . | . | . | . | . | . | . | . | . | . | . | . | . | . | . | . | . | . | . | . |
| . | Y | . | . | . | . | . | . | . | . | . | . | . | . | . | . | . | . | . | . | . | . | . | . | . | . |
| . | T | . | . | . | . | . | . | . | . | . | . | . | . | . | . | . | . | . | . | . | . | . | . | . | . |
| . | T | . | . | . | . | . | . | . | . | . | . | . | . | . | . | . | . | . | . | . | . | . | . | . | ? |
| . | T | . | . | . | . | . | . | . | . | . | . | . | . | . | . | . | . | . | . | . | . | . | . | . | . |
| . | T | . | . | . | . | . | . | . | . | . | . | . | . | . | . | . | . | . | . | . | . | . | . | . | . |
| . | T | . | . | . | . | . | . | . | . | . | . | . | . | . | . | . | . | . | . | . | . | . | . | . | . |
| . | T | . | . | . | . | . | . | . | . | . | . | . | . | . | . | . | . | . | . | . | . | . | . | . | . |
| . | . | . | . | . | . | . | . | . | A | . | . | . | . | . | . | . | . | . | . | . | . | . | . | . | A |
| . | . | . | . | . | . | . | . | . | A | . | . | . | . | . | . | . | . | . | . | . | . | . | . | . | A |
| . | . | . | . | . | . | . | . | . | . | . | . | . | . | . | . | . | . | . | . | . | . | . | . | . | . |
| . | . | . | . | . | . | . | . | . | . | . | . | . | . | . | . | . | . | . | . | . | . | . | . | . | . |
| . | . | . | . | . | . | . | . | . | . | . | . | . | . | . | . | . | . | . | . | . | . | . | . | . | . |
| . | . | . | . | . | . | . | . | . | . | . | . | . | . | . | . | . | . | . | . | . | . | . | . | . | . |
| . | . | . | . | . | . | . | . | . | . | . | . | . | . | . | . | . | . | . | . | . | . | . | . | . | . |
| . | . | . | . | . | . | . | . | . | . | . | . | . | . | . | . | . | . | . | . | . | . | . | . | . | . |
| . | . | . | . | . | . | . | . | . | . | . | . | . | . | . | . | . | . | . | . | . | . | . | . | . | . |
| . | . | . | . | . | . | . | . | . | . | . | . | . | . | . | . | . | . | . | . | . | . | . | . | . | . |
| . | . | . | . | . | . | . | . | . | . | . | . | . | . | . | . | . | . | . | . | . | . | . | . | . | . |
| . | . | . | . | . | . | . | . | . | . | . | . | . | . | . | . | . | . | . | . | . | . | . | . | . | . |
| . | . | . | . | . | . | . | . | . | . | . | . | . | . | . | . | . | . | . | . | . | . | . | . | . | . |
| . | . | . | . | . | . | . | . | . | . | . | . | . | . | . | . | . | . | . | . | . | . | . | . | . | . |
| . | . | . | . | . | . | . | . | . | . | . | . | . | . | . | . | . | . | . | . | . | . | . | ? | ? | ? |
| . | . | . | . | . | . | . | . | . | . | . | . | . | . | . | . | . | . | . | . | . | . | . | . | . | . |
| . | . | . | . | . | . | . | . | . | . | . | . | . | . | . | . | . | . | . | . | . | . | . | . | . | . |
| . | . | . | . | . | . | . | . | . | A | . | . | . | . | . | . | . | . | . | . | . | . | . | . | . | A |
| . | . | . | . | . | . | . | . | . | . | . | . | . | . | . | . | . | . | . | . | . | . | . | . | . | . |
| . | . | . | . | . | . | . | . | . | . | . | . | . | . | . | . | . | . | . | . | . | . | . | . | . | . |
| . | . | . | . | . | . | . | . | . | . | . | . | . | . | . | . | . | . | . | . | . | . | . | . | . | . |
| . | . | . | . | . | . | . | . | . | A | . | . | . | . | . | . | . | . | . | . | . | . | . | . | . | A |

| | | | | | | | | | | | | | | | | | | | | | | | | | |
|---|---|---|---|---|---|---|---|---|---|---|---|---|---|---|---|---|---|---|---|---|---|---|---|---|---|
| . | . | . | . | . | . | . | . | . | . | . | . | . | . | . | . | . | . | . | . | . | . | . | . | . | . |
| . | . | . | . | . | . | . | . | . | . | . | . | . | . | . | . | . | . | . | . | . | . | . | . | . | . |
| . | . | . | . | . | . | . | . | . | . | . | . | . | . | . | . | . | . | . | . | . | . | . | . | . | . |
| . | . | . | . | . | . | . | . | . | . | . | . | . | . | . | . | . | . | . | . | . | . | . | . | . | . |
| . | . | . | . | . | . | . | . | . | . | . | . | . | . | . | . | T | . | . | . | . | . | . | . | . | . |
| . | T | . | . | . | . | . | . | . | . | . | . | . | . | . | . | . | . | . | . | . | . | . | . | . | . |
| . | . | . | . | . | . | . | . | . | A | . | . | . | . | . | . | . | . | . | . | . | . | . | . | . | A |
| . | . | . | . | . | . | . | . | . | . | . | . | . | . | . | . | T | . | . | . | . | . | . | . | . | . |
| . | . | . | . | . | . | . | . | . | . | . | . | . | . | . | . | . | . | . | . | . | . | . | . | . | . |
| . | . | . | . | . | . | . | . | . | . | . | . | . | . | . | . | . | . | . | . | . | . | . | . | . | . |
| . | . | . | . | . | . | . | . | . | . | . | . | . | . | . | . | . | . | . | . | . | . | . | . | . | . |
| . | . | . | . | . | . | . | . | . | . | . | . | . | . | . | . | . | . | . | . | T | . | . | . | . | . |
| . | . | . | . | . | . | . | . | . | . | . | . | . | . | . | . | . | . | . | . | . | . | . | . | . | . |
| . | . | . | . | . | . | . | . | . | . | . | . | . | . | . | . | . | . | . | . | . | . | . | . | . | . |
| . | . | . | . | . | . | . | . | . | . | . | . | . | . | . | . | . | . | . | . | . | . | . | . | . | . |
| . | . | . | . | . | . | . | . | . | A | . | . | . | . | . | . | . | . | . | . | . | . | . | . | . | A |
| . | . | . | . | . | . | . | . | . | . | . | . | . | . | . | . | . | . | . | . | . | . | . | . | . | . |
| . | . | . | . | . | . | . | . | . | . | . | . | . | . | . | . | . | . | . | . | T | . | . | . | . | . |
| . | . | . | . | . | . | . | . | . | . | . | . | . | . | . | . | . | . | . | . | T | . | . | . | . | . |
| . | . | . | . | . | . | . | . | . | . | . | . | . | . | . | . | . | . | . | . | . | . | . | . | . | . |
| . | . | . | . | . | . | . | . | . | . | . | . | . | . | . | . | . | . | . | . | T | . | . | . | . | . |
| . | . | . | . | . | . | . | . | . | . | . | . | . | . | . | . | . | . | . | . | . | . | . | . | . | . |
| . | . | . | . | . | . | . | . | . | . | . | . | . | . | . | . | . | . | . | . | . | . | . | . | . | . |
| . | . | . | T | . | . | . | . | . | A | . | . | . | . | . | . | . | . | ? | . | . | . | . | - | - | - |
| . | . | . | T | . | . | . | . | . | A | . | . | . | . | T | - | - | - | - | - | - | - | - | - | - | - |
| . | . | . | T | . | . | . | . | . | A | . | . | . | . | . | . | . | . | . | . | . | . | . | - | - | - |
| . | . | . | . | . | . | . | . | . | . | . | . | . | . | . | . | . | . | . | . | . | . | . | - | - | - |
| . | . | . | . | . | . | . | . | . | . | . | . | . | . | T | . | . | . | . | . | . | . | . | - | - | - |
| . | . | . | . | . | . | . | . | . | . | . | . | . | . | . | . | . | . | . | . | . | . | . | - | - | - |
| . | . | . | T | . | . | . | . | . | A | . | . | . | . | T | . | . | . | . | - | - | - | - | - | - | - |
| . | . | . | T | . | . | . | . | . | A | . | . | . | . | . | . | . | . | . | . | . | . | . | - | - | - |
| . | . | . | T | . | . | . | . | . | A | . | . | . | . | . | . | . | . | . | . | . | . | . | - | - | - |
| . | . | . | T | . | . | . | . | . | A | . | . | . | . | . | . | . | . | . | . | . | . | . | . | . | A |
| . | . | . | T | . | . | . | . | . | A | . | . | . | . | . | . | . | . | . | . | . | . | . | - | - | - |
| . | . | . | . | . | . | . | . | . | . | . | . | . | . | . | . | . | . | . | . | . | . | . | - | - | - |
| . | . | . | T | . | . | . | . | . | A | . | . | . | . | . | . | . | . | . | . | . | . | . | - | - | - |
| . | . | . | T | . | . | . | . | . | A | . | . | . | . | . | . | . | . | . | . | . | . | . | - | - | - |
| . | T | . | . | . | . | . | . | . | . | . | . | . | . | . | . | . | . | . | . | . | . | . | - | - | - |
| . | . | . | T | . | . | . | . | . | A | . | . | . | . | . | . | . | . | . | . | . | . | . | - | - | - |
| . | . | . | . | . | . | . | . | . | . | A | A | C | . | T | . | . | . | . | . | . | . | . | - | - | - |
| . | . | . | . | . | . | . | . | . | . | . | . | . | . | . | . | . | . | . | . | . | . | . | . | . | . |
| . | . | . | . | . | . | . | . | . | . | . | . | . | . | . | . | . | . | . | . | . | . | . | - | - | - |
| . | . | . | . | . | . | . | . | . | . | . | . | . | . | . | . | . | . | . | . | . | . | . | - | - | - |
| . | . | . | . | . | . | . | . | . | . | . | . | . | . | . | . | . | . | . | . | . | . | . | . | . | . |
| . | . | . | . | . | . | . | . | . | . | . | . | . | . | . | . | . | . | . | . | . | . | . | . | . | . |
| . | . | . | . | . | . | . | . | . | . | . | . | . | . | . | . | . | . | . | . | . | . | . | . | . | . |
| . | . | . | . | . | . | . | . | . | . | . | . | . | . | . | . | . | . | . | . | . | . | . | - | - | - |
| . | . | . | . | . | . | . | . | . | . | . | . | . | . | T | . | . | . | . | . | . | . | . | - | - | - |
| . | . | . | . | . | . | . | . | . | . | . | . | . | . | . | . | . | . | . | . | . | . | . | - | - | - |
| . | . | . | . | . | . | . | . | . | . | . | . | . | . | . | . | . | . | . | . | . | . | . | - | - | - |
| . | . | . | . | . | . | . | . | . | . | . | . | . | . | . | . | . | . | . | . | . | . | . | - | - | - |
| . | . | . | . | . | . | . | . | . | . | A | A | C | . | . | . | . | . | . | . | . | . | . | . | . | . |
| . | . | . | . | . | . | . | . | . | . | A | A | C | . | . | . | . | . | . | . | . | . | . | . | . | . |
| . | . | . | . | . | . | . | . | . | . | A | A | C | . | . | . | . | . | . | . | . | . | . | . | . | . |
| . | . | . | . | . | . | . | . | . | . | A | A | C | . | . | . | . | . | . | . | . | . | . | . | . | . |
| . | . | . | . | . | . | . | . | . | . | A | A | C | . | . | . | . | . | . | . | . | . | . | . | . | . |
| . | T | . | . | . | . | . | . | . | . | . | . | . | . | . | . | . | . | . | . | . | . | . | . | . | . |
| . | . | . | . | . | . | . | . | . | . | A | A | C | . | . | . | . | . | . | . | . | . | . | . | . | . |
| . | . | . | . | . | . | . | . | . | . | A | A | C | . | . | . | . | . | . | . | . | . | . | . | . | . |
| . | . | . | . | . | . | . | . | . | . | A | A | C | . | . | . | . | . | . | . | . | . | . | . | . | . |
| . | . | . | . | . | . | . | . | . | . | A | A | C | . | . | . | . | . | . | . | . | . | . | . | . | . |
| . | . | . | . | . | . | . | . | . | C | A | A | C | . | . | . | . | . | . | . | . | . | . | . | . | . |

| | | | | | | | | | | | | | | | | | | | | | | | | | |
|---|---|---|---|---|---|---|---|---|---|---|---|---|---|---|---|---|---|---|---|---|---|---|---|---|---|
| . | . | . | . | . | . | . | . | . | . | A | A | C | . | . | . | . | . | . | . | . | . | . | . | . | . |
| . | . | . | . | . | . | . | . | . | . | A | A | C | . | . | . | . | . | . | . | . | . | . | . | . | . |
| . | . | . | . | . | . | . | . | . | . | A | A | C | . | . | . | . | . | . | . | . | . | . | . | . | . |
| . | . | . | . | . | . | . | . | . | . | A | A | C | . | . | . | . | . | . | . | . | . | . | . | . | . |
| . | . | . | . | . | . | . | . | . | . | A | A | C | . | . | . | . | . | . | . | . | . | . | . | . | . |
| . | . | . | . | . | . | . | . | . | . | A | A | C | . | . | . | . | . | . | . | . | . | . | . | . | . |
| . | . | . | . | . | . | . | . | . | . | A | A | C | . | . | . | . | . | . | . | . | . | . | . | . | . |
| . | . | . | . | . | . | . | . | . | . | A | A | C | . | . | . | . | . | . | . | . | . | . | . | . | . |
| . | . | . | . | . | . | . | . | . | . | A | A | C | . | . | . | . | . | . | . | . | . | . | . | . | . |
| . | . | . | . | . | . | . | . | . | . | A | A | C | . | . | . | . | . | . | . | . | . | . | . | . | . |
| . | . | . | . | . | . | . | . | . | . | . | . | . | . | . | . | . | . | . | . | . | . | . | . | . | . |
| . | . | . | . | . | . | . | . | . | . | . | . | . | . | . | . | . | . | . | . | . | . | . | . | . | . |
| . | . | . | . | . | . | . | . | . | . | . | . | . | . | . | . | . | . | . | . | . | . | . | . | . | . |
| . | . | . | . | . | . | . | . | . | . | A | A | C | . | . | . | . | . | . | . | . | . | . | . | . | . |
| . | . | . | . | . | . | . | . | . | . | A | A | C | . | . | . | . | . | . | . | . | . | . | . | . | . |
| . | . | . | . | . | . | . | . | . | . | A | A | C | . | . | . | . | . | . | T | . | . | . | . | . | . |
| . | . | . | . | . | . | . | . | . | . | A | A | C | . | . | . | . | . | . | T | . | . | . | . | . | . |
| . | . | . | . | . | . | . | . | . | . | A | A | C | . | . | . | . | . | . | . | . | . | . | . | . | . |
| . | . | . | . | . | . | . | . | . | . | A | A | C | . | . | . | . | . | . | . | . | . | T | . | . | . |
| . | . | . | . | . | . | . | . | . | . | A | A | C | . | . | . | . | . | . | . | . | . | T | . | . | . |
| . | . | . | . | . | . | . | . | . | . | A | A | C | . | . | . | . | . | . | . | . | . | T | . | . | . |
| . | T | . | . | . | . | . | . | . | . | . | . | . | . | . | . | . | . | . | . | . | . | . | . | . | . |
| . | T | . | . | . | . | . | . | . | . | . | . | . | . | . | . | . | . | . | . | . | . | . | . | . | . |
| . | . | . | . | . | . | . | . | . | . | . | . | . | . | . | . | . | . | . | . | . | . | . | . | . | . |
| . | . | . | . | . | . | . | . | . | . | . | . | . | . | . | . | . | . | . | . | . | . | . | . | . | . |
| . | T | . | . | . | . | . | . | . | . | . | . | . | . | . | . | . | . | . | . | . | . | . | . | . | . |
| . | T | T | . | . | . | . | . | . | . | . | . | . | . | . | . | . | . | . | . | . | . | . | . | . | . |
| . | T | . | . | . | . | . | . | . | . | . | . | . | . | . | . | . | . | . | . | . | . | . | . | . | . |
| . | T | . | . | . | . | . | . | . | . | . | . | . | . | . | . | . | . | . | . | . | . | . | . | . | . |
| . | T | . | . | . | . | . | . | . | . | . | . | . | . | . | . | . | . | . | . | . | . | . | . | . | . |
| . | T | . | . | . | . | . | . | . | . | . | . | . | . | . | . | . | . | . | . | . | . | . | . | . | . |
| . | T | . | . | . | . | . | . | . | . | . | . | . | . | . | . | . | . | . | . | . | . | . | . | . | . |
| . | T | . | . | . | . | . | . | . | . | . | . | . | . | . | . | . | . | . | . | . | . | . | . | . | . |
| . | T | . | . | . | . | . | . | . | . | . | . | . | . | . | . | . | . | . | . | . | . | . | . | . | . |
| . | T | T | . | . | . | . | . | . | . | . | . | . | . | . | . | . | . | . | . | . | . | . | . | . | . |
| . | T | . | . | . | . | . | . | . | . | . | . | . | . | . | . | . | . | . | . | . | . | . | . | . | . |
| . | T | . | . | . | . | . | . | . | . | . | . | . | . | . | . | . | . | . | . | . | . | . | . | . | . |
| . | . | . | . | . | . | . | . | . | . | . | . | . | . | . | . | . | . | . | . | . | . | . | . | . | . |
| . | . | . | . | . | . | . | . | T | . | . | . | . | . | . | . | . | . | . | . | . | . | . | . | . | . |
| . | . | . | . | . | . | . | . | . | . | . | . | . | . | . | . | . | . | . | . | . | . | . | . | . | . |
| . | . | . | . | . | . | . | . | T | . | . | . | . | . | . | . | . | . | . | . | . | . | . | . | . | . |
| . | . | . | . | . | . | . | . | . | . | . | . | . | . | . | . | . | . | . | . | . | . | . | . | . | . |
| . | . | . | . | . | . | . | . | T | . | . | . | . | . | . | . | . | . | . | . | . | . | . | . | . | . |
| . | . | . | . | . | . | . | . | T | . | . | . | . | . | . | . | . | . | . | . | . | . | . | . | . | . |
| . | . | . | . | . | . | . | . | T | . | . | . | . | . | . | . | . | . | . | . | . | . | . | . | . | . |
| . | . | . | . | . | . | . | . | T | . | . | . | . | . | . | . | . | . | . | . | . | . | . | . | . | . |
| . | . | . | . | . | . | . | . | T | . | . | . | . | . | . | . | . | . | . | . | . | . | . | . | . | . |
| . | . | . | . | . | . | . | . | T | . | . | . | . | . | . | . | . | . | . | . | . | . | . | . | . | . |
| . | . | . | . | . | . | . | . | . | . | . | . | . | . | . | . | . | . | . | . | . | . | . | . | . | . |
| . | . | . | . | . | . | . | . | T | . | . | . | . | . | . | . | . | . | . | . | . | . | . | . | . | . |
| . | . | . | . | . | . | . | . | T | . | . | . | . | . | . | . | . | . | . | . | . | . | . | . | . | . |
| . | . | . | . | . | . | . | . | T | . | . | . | . | . | . | . | . | . | . | . | . | . | . | . | . | . |
| . | . | . | . | . | . | . | . | T | . | . | . | . | . | . | . | . | . | . | . | . | . | . | . | . | . |
| . | . | . | . | . | . | . | . | T | . | . | . | . | . | . | . | . | . | . | . | . | . | . | . | . | . |
| . | . | . | . | . | . | . | . | . | . | . | . | . | . | . | . | . | . | . | . | . | . | . | . | . | . |
| . | . | . | . | . | . | . | . | T | . | . | . | . | . | . | . | . | . | . | . | . | . | . | . | . | . |
| . | . | . | . | . | . | . | . | . | . | . | . | . | . | . | . | . | . | . | . | . | . | . | . | . | . |
| . | . | . | . | . | . | . | . | . | . | . | . | . | . | . | . | . | . | . | . | . | . | . | . | . | . |
| . | . | . | . | . | . | . | . | . | . | . | . | . | . | . | T | . | . | . | . | . | . | . | . | . | . |
| . | . | . | . | . | . | . | . | . | . | . | . | . | . | . | T | . | . | . | . | . | . | . | . | . | . |
| . | . | . | . | . | . | . | . | . | . | . | . | . | . | . | . | . | . | . | . | . | . | . | . | . | . |
| . | . | . | . | . | . | . | . | . | . | . | . | . | . | . | . | . | . | . | . | . | . | . | . | . | . |

| | | | | | | | | | | | | | | | | | | | | | | | | | |
|---|---|---|---|---|---|---|---|---|---|---|---|---|---|---|---|---|---|---|---|---|---|---|---|---|---|
| A | . | . | . | . | . | . | . | . | . | . | . | . | . | . | . | . | . | . | . | . | . | . | . | T | . |
| . | . | . | . | . | . | . | . | T | . | . | . | . | . | . | . | . | . | . | . | . | . | . | . | . | . |
| . | . | . | . | . | . | . | . | T | . | . | . | . | . | . | . | . | . | . | . | . | . | . | . | . | . |
| . | . | . | . | . | . | . | . | T | . | . | . | . | . | . | . | . | . | . | . | . | . | . | . | . | . |
| . | T | . | . | . | . | . | . | . | . | . | . | . | . | . | . | . | . | . | . | . | . | . | . | . | . |
| . | T | . | . | . | . | . | . | . | . | . | . | . | . | . | . | . | . | . | . | . | . | . | . | . | . |
| . | . | . | . | . | . | . | . | . | . | A | A | C | . | . | . | . | . | . | . | . | . | . | . | . | . |
| . | Y | . | . | . | . | . | . | . | . | A | A | C | . | . | . | . | . | . | . | . | . | . | . | . | . |
| . | T | . | . | . | . | . | . | . | . | . | . | . | . | . | . | . | . | . | . | . | . | . | . | . | . |
| . | Y | . | . | . | . | . | . | . | . | A | A | C | . | . | . | . | . | . | . | . | . | . | . | . | . |
| . | T | . | . | . | . | . | . | . | . | . | . | . | . | . | . | . | . | . | . | . | . | . | . | . | . |
| . | T | . | . | . | . | . | . | . | . | . | . | . | . | . | . | . | . | . | . | . | . | . | . | . | . |
| . | T | . | . | . | . | . | . | . | . | . | . | . | . | . | . | . | . | . | . | . | . | . | . | . | . |
| . | T | . | . | . | . | . | . | . | . | . | . | . | . | . | . | . | . | . | . | . | . | . | . | . | . |
| . | T | . | . | . | . | . | . | . | . | R | R | S | . | . | . | . | . | . | . | . | . | . | . | . | . |
| . | T | . | . | . | . | . | . | . | . | . | . | . | . | . | . | . | . | . | . | . | . | . | . | . | . |
| . | T | . | . | . | . | . | . | . | . | . | . | . | . | . | . | . | . | . | . | . | . | . | . | . | . |
| . | Y | . | . | . | . | . | . | . | . | A | A | C | . | . | . | . | . | . | . | . | . | . | . | . | . |
| . | T | . | . | . | . | . | . | . | . | . | . | . | . | . | . | . | . | . | . | . | . | . | . | . | . |
| . | T | . | . | . | . | . | . | . | . | . | . | . | . | . | . | . | . | . | K | . | . | . | . | . | . |
| . | T | . | . | . | . | . | . | . | . | . | . | . | . | . | . | . | . | . | . | . | . | . | . | . | . |
| . | T | . | . | . | . | . | . | . | . | . | . | . | . | . | . | . | . | . | . | . | . | . | . | . | . |
| . | T | . | . | . | . | . | . | . | . | . | . | . | . | . | . | . | . | . | . | . | . | . | . | . | . |
| . | T | . | . | . | . | . | . | . | . | . | . | . | . | . | . | . | . | . | . | . | . | . | . | . | . |
| . | T | . | . | . | . | . | . | . | . | . | . | . | . | . | . | . | . | . | . | . | . | . | . | . | . |
| . | T | . | . | . | . | . | . | . | . | . | . | . | . | . | . | . | . | . | . | . | . | . | . | . | . |
| . | T | . | . | . | . | . | . | . | . | . | . | . | . | . | . | . | . | . | . | . | . | . | . | . | . |
| . | T | . | . | . | . | . | . | . | . | . | . | . | . | . | . | . | . | . | . | . | . | . | . | . | . |
| . | T | . | . | . | . | . | . | . | . | . | . | . | . | . | . | . | . | . | . | . | . | . | . | . | . |
| . | T | . | . | . | . | . | . | . | . | . | . | . | . | . | . | . | . | . | . | . | . | . | . | . | . |
| . | T | . | . | . | . | . | . | c | . | . | . | . | . | . | . | . | . | . | . | . | . | . | . | . | . |
| . | . | . | . | . | . | . | . | . | . | . | . | . | . | . | . | . | . | . | . | . | . | . | . | . | . |
| . | . | . | . | . | . | . | . | . | . | . | . | . | . | . | . | . | . | . | . | . | . | . | . | . | . |
| . | . | . | . | . | . | . | . | . | . | . | . | . | . | . | . | . | . | . | . | . | . | . | . | . | . |
| . | . | . | . | . | . | . | . | . | . | . | . | . | . | . | . | . | . | . | . | . | . | . | . | . | . |
| . | . | . | . | . | . | . | . | . | A | . | . | . | . | . | . | . | . | . | . | . | . | . | . | . | . |
| . | . | . | . | . | . | . | . | . | . | . | . | . | . | . | . | . | . | . | . | . | . | . | . | . | . |
| . | . | . | . | . | . | . | . | T | . | . | . | . | . | . | . | . | . | . | . | . | . | . | . | . | . |
| . | . | . | . | . | . | . | . | T | . | . | . | . | . | . | . | . | . | . | . | . | . | . | . | . | . |
| . | . | . | . | . | . | . | . | . | . | . | . | . | . | . | . | . | . | . | . | . | . | . | . | . | . |
| . | . | . | . | . | . | . | . | . | . | . | . | . | . | . | . | . | . | . | . | . | . | . | . | . | . |
| . | . | . | . | . | . | . | . | T | . | . | . | . | . | . | . | . | . | . | . | . | . | . | . | . | . |
| . | . | . | . | . | . | . | . | T | . | . | . | . | . | . | . | . | . | . | . | . | . | . | . | . | . |
| . | . | . | . | . | . | . | . | . | . | . | . | . | . | . | . | . | . | . | . | . | . | . | . | . | . |
| . | . | . | . | . | . | . | . | T | . | . | . | . | . | . | . | . | . | . | . | . | . | . | . | . | . |
| . | . | . | . | . | . | . | . | . | . | . | . | . | . | . | . | . | . | . | . | . | . | . | . | . | . |
| . | . | . | . | . | . | . | . | T | . | . | . | . | . | . | . | . | . | . | . | . | . | . | . | . | . |
| . | . | . | . | . | . | . | . | . | . | . | . | . | . | . | . | . | . | . | . | . | . | . | . | . | . |
| . | . | . | . | . | . | . | . | T | . | . | . | . | . | . | . | . | . | . | . | . | . | . | . | . | . |
| . | . | . | . | . | . | . | . | T | . | . | . | . | . | . | . | . | . | . | . | . | . | . | . | . | . |
| . | . | . | . | . | . | . | . | . | . | . | . | . | . | . | . | . | . | . | . | . | . | . | . | . | . |
| . | . | . | . | . | . | . | . | . | . | . | . | . | . | . | . | . | . | . | . | . | . | . | . | . | . |
| . | . | . | . | . | . | . | . | T | . | . | . | . | . | . | . | . | . | . | . | . | . | . | . | . | . |
| . | . | . | . | . | . | . | . | T | . | . | . | . | . | . | . | . | . | . | . | . | . | . | . | . | . |
| . | . | . | . | . | . | . | . | T | . | . | . | . | . | . | . | . | T | . | . | . | . | . | . | . | . |
| . | . | . | . | . | . | . | . | T | . | . | . | . | . | . | . | . | T | . | . | . | . | . | . | . | . |
| . | . | . | . | . | . | . | . | T | . | . | . | . | . | . | . | . | . | . | . | . | . | . | . | . | . |
| . | . | . | . | . | . | . | . | . | . | . | . | . | . | . | . | . | . | . | . | . | . | . | . | . | . |
| . | . | . | . | . | . | . | . | . | . | . | . | . | . | . | . | . | . | . | . | . | . | . | . | . | . |
| . | . | . | . | . | . | . | . | T | . | . | . | . | . | . | . | . | . | . | . | . | . | . | . | . | . |

| 264 | 265 | 266 | 267 | 268 | 269 | 270 |
|---|---|---|---|---|---|---|
| 3'UTR | 3'UTR | 3'UTR | 3'UTR | 3'UTR | 3'UTR | 3'UTR |
| 29743 | 29774 | 29827 | 29829 | 29830 | 29870 | 29874 |
| C | C | A | T | G | C | A |
| . | . | . | . | . | . | - |
| . | . | . | . | . | . | - |
| . | . | . | . | . | . | - |
| . | . | . | . | . | . | . |
| . | . | . | . | . | . | - |
| . | . | . | . | . | . | . |
| . | . | . | . | . | - | - |
| . | . | . | . | . | . | . |
| . | . | . | . | . | . | - |
| . | . | . | . | . | . | . |
| . | . | . | . | . | . | - |
| . | . | . | . | . | . | . |
| . | . | . | . | . | . | - |
| . | . | . | . | . | . | . |
| . | . | . | . | . | . | - |
| . | . | . | . | . | . | . |
| . | . | . | . | . | . | - |
| . | . | . | . | . | . | - |
| . | . | . | . | . | . | - |
| . | . | . | . | . | . | . |
| . | . | . | . | . | . | - |
| . | . | . | . | . | . | - |
| . | . | . | . | . | . | . |
| . | . | . | . | . | . | - |
| . | . | . | . | . | - | - |
| . | . | . | . | . | . | - |
| . | . | . | . | . | . | - |
| . | . | . | . | . | . | - |
| . | . | . | . | . | . | - |
| . | . | . | . | . | . | - |
| . | . | . | . | . | . | - |
| . | . | . | . | . | . | . |
| . | . | . | . | . | . | - |
| . | . | . | . | . | . | . |
| . | . | . | . | . | . | . |
| . | . | . | . | . | - | - |
| . | . | . | . | . | - | - |
| . | . | . | . | . | - | - |
| . | . | . | . | . | - | - |
| . | . | . | . | . | - | - |
| . | . | . | . | . | - | - |
| - | - | - | - | - | - | - |
| . | . | . | . | . | - | - |
| . | . | . | . | . | - | - |
| . | . | . | . | . | . | - |
| . | . | . | . | . | . | - |
| . | . | . | . | . | . | - |
| . | . | . | . | . | . | - |
| . | . | . | . | . | . | - |
| . | . | . | . | . | A | . |
| . | . | . | . | . | . | . |
| . | . | . | . | . | A | . |
| . | . | . | . | . | M | . |
| . | . | . | . | . | ? | ? |
| . | . | . | . | . | ? | ? |

| | | | | | | |
|---|---|---|---|---|---|---|
| . | . | . | . | . | ? | ? |
| . | . | . | . | . | ? | ? |
| . | . | . | . | . | ? | ? |
| . | . | . | . | . | ? | ? |
| . | . | . | . | . | ? | ? |
| . | . | . | . | . | ? | ? |
| . | . | . | . | . | - | - |
| . | . | . | . | . | - | - |
| . | . | . | . | . | - | - |
| . | . | . | . | . | - | - |
| . | . | . | . | . | - | - |
| . | . | . | . | . | - | - |
| . | . | . | . | . | - | - |
| . | . | . | . | . | - | - |
| . | . | . | . | . | - | - |
| . | . | . | . | . | c | - |
| . | . | . | . | . | - | - |
| c | . | . | . | . | - | - |
| c | c | a | t | g | - | - |
| . | . | - | - | - | - | - |
| . | . | . | . | . | - | - |
| . | . | . | . | . | - | - |
| . | . | . | . | . | - | - |
| . | . | . | . | . | - | - |
| . | . | . | . | . | - | - |
| . | . | . | . | . | - | - |
| . | . | . | . | . | - | - |
| . | . | . | . | . | - | - |
| . | . | . | . | . | - | - |
| . | . | . | . | . | - | - |
| . | . | . | . | . | - | - |
| . | . | . | . | . | - | - |
| c | . | . | . | . | - | - |
| . | . | . | . | . | - | - |
| . | . | . | . | . | - | - |
| . | . | . | . | . | - | - |
| . | . | . | . | . | - | - |
| . | . | . | . | . | - | - |
| . | . | . | . | . | - | - |
| . | . | . | . | . | c | - |
| . | . | . | . | . | - | - |
| . | . | . | . | . | - | - |
| . | . | . | . | . | - | - |
| . | . | . | . | . | - | - |
| . | . | . | . | . | - | - |
| . | . | . | . | . | - | - |
| . | . | . | . | . | - | - |
| . | . | . | . | . | - | - |
| - | - | - | - | - | - | - |
| . | . | . | . | . | - | - |
| . | . | . | . | . | - | - |
| . | . | . | . | . | - | - |
| c | . | . | . | . | - | - |
| . | . | . | . | . | - | - |
| . | . | . | . | . | c | a |
| . | . | . | . | . | c | - |
| . | . | . | . | . | - | - |
| . | . | . | . | . | - | - |
| . | . | . | . | . | - | - |
| . | . | . | . | . | - | - |
| . | . | . | . | . | - | - |
| . | . | . | . | . | - | - |

| | | | | | | |
|---|---|---|---|---|---|---|
| . | . | . | . | . | - | - |
| . | . | . | . | . | - | - |
| . | . | . | . | . | - | - |
| . | . | . | . | . | - | - |
| . | . | . | . | . | - | - |
| c | c | . | . | . | - | - |
| . | . | . | . | . | - | - |
| . | . | . | . | . | - | - |
| . | . | . | . | . | - | - |
| . | . | . | . | . | - | - |
| Y | . | . | . | . | - | - |
| . | . | . | . | . | - | - |
| . | . | . | . | . | - | - |
| . | . | . | . | . | - | - |
| . | . | . | . | . | - | - |
| . | . | . | . | . | c | - |
| . | . | . | . | . | c | a |
| . | . | . | . | . | - | - |
| . | . | . | . | . | c | a |
| . | . | . | . | . | - | - |
| . | . | . | . | . | - | - |
| . | . | . | . | . | - | - |
| . | . | . | . | . | - | - |
| . | . | . | . | . | c | - |
| . | . | . | . | . | c | a |
| c | c | . | . | . | - | - |
| . | . | . | . | . | - | - |
| . | . | . | . | . | - | - |
| . | . | . | . | . | - | - |
| . | . | . | . | . | - | - |
| . | . | . | . | . | c | - |
| . | . | . | . | . | - | - |
| . | . | . | . | . | ? | ? |
| . | . | . | . | . | ? | ? |
| . | . | . | . | . | ? | ? |
| . | . | . | . | . | ? | ? |
| . | . | . | . | . | ? | ? |
| . | . | . | . | . | ? | ? |
| . | . | . | . | . | ? | ? |
| . | . | . | . | . | ? | ? |
| . | . | . | . | . | ? | ? |
| . | . | . | . | . | . | ? |
| . | . | . | . | . | ? | ? |
| . | . | . | . | . | ? | ? |
| . | . | . | . | . | . | . |
| . | . | . | . | . | ? | ? |
| . | . | . | . | . | ? | ? |
| . | ? | ? | ? | ? | A | . |
| . | . | . | . | . | A | . |
| . | . | . | . | . | A | . |
| . | . | . | . | . | A | . |
| . | . | . | . | . | ? | ? |
| . | . | . | . | . | . | . |
| . | . | . | . | . | ? | ? |
| . | . | . | . | . | ? | ? |
| . | . | . | . | . | ? | ? |
| . | . | . | . | . | ? | ? |
| . | . | . | . | . | ? | ? |
| . | . | . | . | . | ? | ? |
| . | . | . | . | . | ? | ? |
| . | . | . | . | . | ? | ? |
| . | . | . | . | . | ? | ? |

| | | | | | | |
|---|---|---|---|---|---|---|
| . | . | . | . | . | ? | ? |
| . | . | . | . | . | ? | ? |
| . | . | . | . | . | ? | ? |
| . | . | . | . | . | ? | ? |
| . | . | . | . | . | ? | ? |
| . | . | . | . | . | ? | ? |
| . | . | . | . | . | ? | ? |
| . | . | . | . | . | ? | ? |
| . | . | . | . | . | ? | ? |
| . | . | . | . | . | ? | ? |
| . | . | . | . | . | ? | ? |
| . | . | . | . | . | ? | ? |
| . | . | . | . | . | ? | ? |
| . | . | . | . | . | ? | ? |
| . | . | . | . | . | ? | ? |
| . | . | . | . | . | ? | ? |
| . | . | . | . | . | ? | ? |
| . | . | . | . | . | . | . |
| . | . | . | . | . | ? | ? |
| . | . | . | . | . | ? | ? |
| . | . | . | . | . | ? | ? |
| . | . | . | . | . | ? | ? |
| . | . | . | . | . | ? | ? |
| . | . | . | . | . | ? | ? |
| . | . | . | . | . | ? | ? |
| . | . | . | . | . | ? | ? |
| . | . | . | . | . | ? | ? |
| . | . | . | . | . | ? | ? |
| . | . | . | . | . | ? | ? |
| . | . | . | . | . | ? | ? |
| . | . | . | . | . | ? | ? |
| . | . | . | . | . | ? | ? |
| ? | . | . | . | . | ? | ? |
| . | . | . | . | . | ? | ? |
| . | . | . | . | . | ? | ? |
| . | . | . | . | . | ? | ? |
| . | . | . | . | . | ? | ? |
| . | . | . | . | . | ? | ? |
| . | . | . | . | . | ? | ? |
| . | . | . | . | . | ? | ? |
| . | . | . | . | . | ? | ? |
| . | . | . | . | . | ? | ? |
| . | . | . | . | . | ? | ? |
| . | . | . | . | . | ? | ? |
| . | . | . | . | . | ? | ? |
| . | . | . | . | . | ? | ? |
| . | . | ? | ? | ? | ? | ? |
| . | . | ? | ? | ? | ? | ? |
| . | . | ? | ? | ? | ? | ? |
| . | . | ? | ? | ? | ? | ? |
| . | . | . | . | . | ? | ? |
| . | . | . | . | . | ? | ? |
| . | . | . | . | . | ? | ? |
| . | . | . | . | . | ? | ? |
| . | . | . | . | . | ? | ? |
| . | . | . | . | . | ? | ? |
| . | . | . | . | . | ? | ? |
| . | . | . | . | . | ? | ? |
| . | . | . | . | . | ? | ? |
| . | . | . | . | . | ? | ? |
| . | . | . | . | . | ? | ? |
| . | . | . | . | . | ? | ? |

| | | | | | | |
|---|---|---|---|---|---|---|
| . | . | . | . | . | ? | ? |
| . | . | . | . | . | ? | ? |
| . | . | . | . | . | ? | ? |
| . | . | . | . | . | ? | ? |
| . | . | . | . | . | - | - |
| . | . | . | . | . | . | . |
| . | . | . | . | . | . | . |
| . | . | . | . | . | . | - |
| . | . | . | . | . | . | . |
| . | . | . | . | . | . | . |
| . | . | . | . | . | . | . |
| . | . | . | . | . | . | . |
| . | . | . | . | . | . | . |
| . | . | . | . | . | - | - |
| . | . | . | . | . | . | . |
| . | . | . | . | . | . | . |
| . | . | . | . | . | . | . |
| . | . | . | . | . | . | . |
| . | . | . | . | . | . | . |
| . | . | . | . | . | . | . |
| . | T | . | . | . | . | . |
| . | . | . | . | . | . | . |
| . | . | . | . | . | - | - |
| . | . | . | . | . | . | . |
| . | . | . | . | . | . | . |
| . | . | . | . | . | . | . |
| . | . | . | . | . | . | . |
| . | T | . | . | . | . | . |
| . | . | . | . | . | - | - |
| . | . | . | . | . | . | . |
| . | . | . | . | . | . | . |
| . | . | . | . | . | . | . |
| . | . | . | . | . | . | . |
| . | . | . | . | . | . | . |
| . | . | . | . | . | . | . |
| . | . | . | . | . | . | . |
| . | . | . | . | . | - | - |
| . | . | . | . | . | - | - |
| . | . | . | . | . | - | - |
| . | . | . | . | . | - | - |
| . | . | . | . | . | - | - |
| . | . | . | . | . | - | - |
| . | . | . | . | . | - | - |
| . | . | . | . | . | - | - |
| . | . | . | . | . | - | - |
| . | . | . | . | . | - | - |
| . | . | . | . | . | - | - |
| . | . | . | . | . | - | - |
| . | . | . | . | . | - | - |
| . | . | . | . | . | - | - |
| . | . | . | . | . | - | - |
| . | . | . | . | . | - | - |
| . | . | . | . | . | - | - |
| . | . | . | . | . | - | - |
| . | . | . | . | . | - | - |
| . | . | . | . | . | - | - |
| . | . | . | . | . | - | - |
| . | . | . | . | . | - | - |
| . | . | . | . | . | - | - |
| . | . | . | . | . | - | - |
| . | . | . | . | . | - | - |
| . | . | . | . | . | - | - |

| | | | | | | |
|---|---|---|---|---|---|---|
| . | . | . | . | . | - | - |
| . | . | . | . | . | - | - |
| . | . | . | . | . | - | - |
| . | . | . | . | . | - | - |
| . | . | . | . | . | - | - |
| . | . | . | . | . | - | - |
| . | . | . | . | . | ? | ? |
| . | . | . | . | . | ? | ? |
| . | . | . | . | . | ? | ? |
| . | . | . | . | . | . | ? |
| . | . | . | . | . | ? | ? |
| . | . | . | . | . | ? | ? |
| . | . | . | . | . | ? | ? |
| . | . | . | . | . | ? | ? |
| T | . | . | . | . | ? | ? |
| . | . | . | . | . | ? | ? |
| . | . | . | . | . | . | ? |
| . | . | . | . | . | ? | ? |
| . | . | . | . | . | ? | ? |
| . | . | . | . | . | ? | ? |
| . | . | . | . | . | ? | ? |
| . | . | . | . | . | ? | ? |
| . | . | . | . | . | ? | ? |
| . | . | . | . | . | ? | ? |
| . | . | . | . | . | ? | ? |
| . | . | . | . | . | ? | ? |
| . | . | . | . | . | ? | ? |
| . | . | . | . | . | ? | ? |
| . | . | . | . | . | ? | ? |
| . | . | . | . | . | ? | ? |
| . | . | . | . | . | ? | ? |
| . | . | . | . | . | ? | ? |
| . | . | . | . | . | ? | ? |
| c | c | t | t | t | - | - |
| c | c | t | t | t | - | - |
| c | c | t | t | t | - | - |
| c | c | t | t | t | - | - |
| c | c | a | t | t | - | - |
| c | c | a | t | t | - | - |
| c | c | t | t | t | - | - |
| c | c | a | t | t | - | - |
| c | c | t | t | t | - | - |
| c | c | a | t | t | - | - |
| c | c | t | t | t | - | - |
| c | c | t | t | t | - | - |
| c | c | t | t | t | - | - |
| c | c | t | t | c | - | - |
| c | c | t | t | t | - | - |
| c | c | t | a | t | - | - |
| c | c | t | a | t | - | - |
| c | c | a | t | t | - | - |
| c | c | t | t | t | - | - |
| c | c | t | t | t | - | - |
| c | c | t | t | t | - | - |
| c | c | t | t | t | - | - |
| c | c | a | t | - | - | - |
| c | c | a | t | - | - | - |
| . | . | . | . | . | - | - |
| . | . | . | . | ? | - | - |
| . | - | - | - | - | - | - |
| . | . | . | . | . | - | - |
| . | . | . | . | . | - | - |

| | | | | | | |
|---|---|---|---|---|---|---|
| . | . | . | . | . | - | - |
| . | . | . | . | T | ? | ? |
| . | . | . | . | T | ? | ? |
| . | . | T | . | C | ? | ? |
| . | . | . | . | T | ? | ? |
| . | . | T | . | T | ? | ? |
| . | . | T | . | T | ? | ? |
| . | . | T | . | T | ? | ? |
| . | . | T | . | T | ? | ? |
| . | . | T | . | T | ? | ? |
| ? | ? | ? | ? | ? | ? | ? |
| . | . | . | . | T | ? | ? |
| . | . | T | . | T | ? | ? |
| . | . | T | . | T | ? | ? |
| . | . | . | . | T | ? | ? |
| . | . | . | . | . | ? | ? |
| . | . | . | . | T | ? | ? |
| . | . | T | . | ? | ? | ? |
| . | . | T | . | T | ? | ? |
| . | . | T | . | T | ? | ? |
| . | . | T | . | T | ? | ? |
| . | . | . | . | T | ? | ? |
| . | . | T | . | T | ? | ? |
| . | . | . | . | T | ? | ? |
| . | . | . | . | . | ? | ? |
| . | . | T | ? | ? | ? | ? |
| . | . | T | . | T | ? | ? |
| . | . | . | . | . | ? | ? |
| . | . | T | . | T | ? | ? |
| . | . | . | . | . | ? | ? |
| . | . | . | . | C | ? | ? |
| . | . | T | . | ? | ? | ? |
| . | . | . | . | . | . | . |
| . | . | . | . | . | ? | ? |
| . | . | . | . | . | . | ? |
| . | . | ? | ? | ? | - | - |
| . | . | . | . | . | ? | ? |
| . | . | . | . | . | ? | ? |
| . | . | . | . | . | ? | ? |
| . | . | . | . | . | ? | ? |
| . | . | . | . | . | ? | ? |
| . | . | . | . | . | ? | ? |
| . | . | . | . | . | ? | ? |
| . | . | . | . | . | ? | ? |
| . | . | . | . | . | ? | ? |
| . | . | . | . | . | ? | ? |
| . | . | . | . | . | ? | ? |
| . | . | . | . | . | ? | ? |
| . | . | . | . | . | ? | ? |
| . | . | . | . | . | ? | ? |
| . | . | . | . | . | ? | ? |
| . | . | . | . | . | ? | ? |
| . | . | . | . | . | . | ? |
| . | . | . | . | . | . | . |
| ? | ? | ? | ? | ? | . | . |
| . | . | . | . | . | . | ? |
| . | . | . | . | . | ? | ? |
| . | . | . | . | . | . | ? |
| . | . | . | . | . | . | . |
| . | . | . | . | . | . | ? |
| . | . | . | . | . | . | ? |
| . | . | . | . | . | ? | ? |

| | | | | | | |
|---|---|---|---|---|---|---|
| . | . | . | . | . | . | . |
| . | . | . | . | . | . | ? |
| . | . | . | . | . | . | ? |
| . | . | . | . | . | . | ? |
| . | . | . | . | . | . | ? |
| . | . | . | . | . | . | ? |
| . | . | . | . | . | . | . |
| . | . | . | . | . | . | ? |
| . | . | . | . | . | . | ? |
| . | . | . | . | . | . | ? |
| . | . | . | . | . | ? | ? |
| . | . | . | . | . | . | ? |
| . | . | . | . | . | . | ? |
| . | . | . | . | . | . | ? |
| . | . | . | . | . | . | ? |
| . | . | . | . | . | . | ? |
| . | . | . | . | . | . | ? |
| . | . | . | . | . | ? | ? |
| . | . | . | . | . | . | ? |
| . | . | . | . | . | . | ? |
| . | . | . | . | . | . | ? |
| . | . | . | . | . | ? | ? |
| . | . | . | . | . | ? | ? |
| - | - | - | - | - | - | - |
| - | - | - | - | - | - | - |
| - | - | - | - | - | - | - |
| - | - | - | - | - | - | - |
| - | - | - | - | - | - | - |
| - | - | - | - | - | - | - |
| - | - | - | - | - | - | - |
| - | - | - | - | - | - | - |
| - | - | - | - | - | - | - |
| . | . | . | . | . | - | - |
| - | - | - | - | - | - | - |
| - | - | - | - | - | - | - |
| - | - | - | - | - | - | - |
| - | - | - | - | - | - | - |
| - | - | - | - | - | - | - |
| - | - | - | - | - | - | - |
| - | - | - | - | - | - | - |
| . | . | . | . | . | - | - |
| - | - | - | - | - | - | - |
| - | - | - | - | - | - | - |
| . | . | . | . | . | - | - |
| . | . | - | - | - | - | - |
| . | . | - | - | - | - | - |
| - | - | - | - | - | - | - |
| - | - | - | - | - | - | - |
| - | - | - | - | - | - | - |
| - | - | - | - | - | - | - |
| - | - | - | - | - | - | - |
| . | . | . | . | . | - | - |
| . | . | . | . | . | - | - |
| . | . | . | . | . | - | - |
| . | . | . | . | . | - | - |
| . | . | . | . | . | - | - |
| . | . | . | . | . | - | - |
| . | . | . | . | . | - | - |
| . | . | . | . | . | - | - |
| . | . | . | . | . | - | - |
| . | . | . | . | . | - | - |
| . | . | . | . | . | - | - |

| | | | | | | |
|---|---|---|---|---|---|---|
| . | . | . | . | . | - | - |
| . | . | . | . | . | - | - |
| . | . | . | . | . | - | - |
| . | . | . | . | . | - | - |
| . | . | . | . | . | - | - |
| . | . | . | . | . | - | - |
| . | . | . | . | . | - | - |
| . | . | . | . | . | - | - |
| . | . | . | . | . | - | - |
| . | . | . | . | . | - | - |
| . | . | . | . | . | - | - |
| . | . | . | . | . | - | - |
| . | . | . | . | . | - | - |
| . | . | . | . | . | - | - |
| . | . | . | . | . | - | - |
| . | . | . | . | . | - | - |
| . | . | . | . | . | - | - |
| . | . | . | . | . | - | - |
| . | . | . | . | . | - | - |
| . | . | . | . | . | - | - |
| . | . | . | . | . | - | - |
| . | . | . | . | . | - | - |
| . | . | . | . | . | - | - |
| . | . | . | . | . | - | - |
| . | . | . | . | . | - | - |
| . | . | . | . | . | - | - |
| . | . | . | . | . | - | - |
| . | . | . | . | . | - | - |
| . | . | . | . | . | - | - |
| . | . | . | . | . | - | - |
| . | . | . | . | . | - | - |
| . | . | . | . | . | - | - |
| . | . | . | . | . | - | - |
| . | . | . | . | . | - | - |
| . | . | . | . | . | - | - |
| . | . | . | . | . | - | - |
| . | . | . | . | . | - | - |
| . | . | . | . | . | - | - |
| . | . | . | . | . | ? | ? |
| . | . | . | . | . | ? | ? |
| . | . | . | . | . | ? | ? |
| . | . | . | . | . | ? | ? |
| . | . | . | . | . | ? | ? |
| . | . | . | . | . | ? | ? |
| . | . | . | . | . | ? | ? |
| . | . | . | . | . | ? | ? |
| . | . | . | . | . | ? | ? |
| . | . | . | . | . | ? | ? |
| . | . | . | . | . | ? | ? |
| . | . | . | . | . | ? | ? |
| . | . | . | . | . | ? | ? |
| . | . | . | . | . | ? | ? |
| . | . | . | . | . | ? | ? |
| . | . | . | . | . | ? | ? |
| . | . | . | . | . | ? | ? |
| . | . | . | . | . | ? | ? |
| . | . | . | . | . | ? | ? |
| . | . | . | . | . | ? | ? |
| . | . | . | . | . | ? | ? |
| . | . | . | . | . | ? | ? |
| . | . | . | . | . | ? | ? |
| . | . | . | . | . | ? | ? |

| | | | | | | |
|---|---|---|---|---|---|---|
| T | . | . | . | . | ? | ? |
| . | . | . | . | . | ? | ? |
| . | . | . | . | . | ? | ? |
| . | . | . | . | . | ? | ? |
| . | T | . | . | . | - | - |
| . | . | . | . | . | ? | ? |
| . | . | . | . | . | - | - |
| . | . | . | . | . | ? | ? |
| . | . | . | . | . | ? | a |
| . | . | . | . | . | c | - |
| . | . | . | . | . | c | . |
| . | . | . | . | . | ? | a |
| . | . | . | . | . | c | . |
| . | . | . | . | . | c | a |
| . | . | . | . | . | ? | a |
| . | . | . | . | . | ? | . |
| . | . | . | . | . | ? | . |
| . | . | . | . | . | . | . |
| . | . | . | . | . | ? | . |
| . | . | . | . | . | ? | . |
| . | . | . | . | . | ? | a |
| . | . | . | . | . | ? | . |
| . | . | . | . | . | ? | . |
| . | . | . | . | . | ? | . |
| . | . | . | . | . | ? | ? |
| . | . | . | . | . | ? | . |
| . | . | . | . | . | ? | ? |
| . | . | . | . | . | ? | a |
| . | . | . | . | . | ? | . |
| . | . | . | . | . | ? | . |
| . | . | . | . | . | ? | . |
| . | . | . | . | . | ? | ? |
| . | . | . | . | . | ? | ? |
| . | . | . | . | . | ? | ? |
| . | . | . | . | . | . | . |
| . | . | . | . | . | . | . |
| . | . | . | . | . | ? | ? |
| . | . | . | . | . | ? | ? |
| . | . | . | . | . | ? | ? |
| . | . | . | . | . | . | G |
| . | . | . | . | . | . | G |
| . | . | . | . | . | ? | ? |
| . | . | . | . | . | ? | ? |
| . | . | . | . | . | ? | ? |
| . | . | . | . | . | ? | ? |
| . | . | . | . | . | ? | ? |
| . | . | . | . | . | ? | ? |
| . | . | . | . | . | ? | ? |
| . | . | . | . | . | ? | ? |
| . | . | . | . | . | ? | ? |
| . | . | . | . | . | ? | ? |
| . | . | . | . | . | ? | ? |
| . | . | . | . | . | ? | ? |
| . | . | . | . | . | ? | ? |
| . | . | . | . | . | ? | ? |
| . | . | . | . | . | ? | ? |
| . | . | . | . | . | ? | ? |
| . | . | . | . | . | ? | ? |
| . | . | . | . | . | ? | ? |
| . | . | . | . | . | ? | ? |

**Supplementary file 4: Details of lineage analysis of studied genomes**

| | GISAID | | | Genome Detective | | | NCBI BLAST | PANGOLIN | | | | | | | COVIDEX | | COVIDEX | | COVIDEX | | |
|---|---|---|---|---|---|---|---|---|---|---|---|---|---|---|---|---|---|---|---|---|---|
| S No | Genome ID | Accession ID | State | Genome Length | NT Identity (%) | AA Identity (%) | Per. ident | Lineage | Bootstrap | SH-aLRT | Most common countries | Number of taxa | Date range | Days since last sampling | NEXTSTRAIN Nomenclature prediccion | NEXTSTRAIN probability | Rambault prediccion | Rambault probability | rotype Nomenclature | FMDV probability | Haplogroup |
| 1 | NC 045512.2 | NC 045512.2 | Wuhan, China | 29903 | 100 | 100 | 0 | B | 99 | 100 | UK, China, USA | 1016 | December-24, April-23 | 22 | A7 | 0.371 | B | 1 | A | 0.234 | A1 |
| 4 | 420543 | EPI_ISL_420543 | Maharashtra | 29869 | 99.9799 | 99.9721 | 99.98 | B.1 | 88 | 95 | UK, USA, Australia | 7440 | January-24, May-21 | 48 | A2a | 0.98 | B.1 | 0.952 | A | 0.232 | A1.1.1 |
| 5 | 420544 | EPI_ISL_420544 | Maharashtra | 29903 | 99.9833 | 99.9721 | 99.98 | B.1 | 97 | 100 | UK, USA, Australia | 6733 | January-24, April-23 | 25 | A2a | 1 | B.1 | 0.988 | A | 0.233 | A1.1 |
| 6 | 420545 | EPI_ISL_420545 | Maharashtra | 29862 | 99.9799 | 99.9721 | 99.98 | B.1 | 95 | 95 | UK, USA, Australia | 7440 | January-24, May-21 | 48 | A2a | 0.98 | B.1 | 0.953 | A | 0.232 | A1.1.1 |
| 7 | 420546 | EPI_ISL_420546 | Maharashtra | 29902 | 99.9833 | 99.9721 | 99.98 | B.1 | 96 | 95 | UK, USA, Australia | 7440 | January-24, May-21 | 48 | A2a | 1 | B.1 | 0.988 | A | 0.233 | A1.1 |
| 8 | 420547 | EPI_ISL_420547 | Maharashtra | 29851 | 99.9833 | 99.9721 | 99.98 | B.1 | 98 | 100 | UK, USA, Australia | 7440 | January-24, May-21 | 48 | A2a | 0.999 | B.1 | 0.992 | A | 0.233 | A1.1 |
| 9 | 420548 | EPI_ISL_420548 | Maharashtra | 29876 | 99.9833 | 99.9721 | 99.98 | B.1 | 97 | 100 | UK, USA, Australia | 7440 | January-24, May-21 | 48 | A2a | 1 | B.1 | 0.988 | A | 0.233 | A1.1 |
| 10 | 420549 | EPI_ISL_420549 | Maharashtra | 29858 | 99.9833 | 99.9721 | 99.98 | B.1 | 97 | 96 | UK, USA, Australia | 7440 | January-24, May-21 | 48 | A2a | 1 | B.1 | 0.99 | A | 0.233 | A1.1 |
| 11 | 420550 | EPI_ISL_420550 | Maharashtra | 29886 | 99.9833 | 99.9721 | 99.98 | B.1 | 96 | 95 | UK, USA, Australia | 7440 | January-24, May-21 | 48 | A2a | 1 | B.1 | 0.988 | A | 0.233 | A1.1 |
| 12 | 420551 | EPI_ISL_420551 | Maharashtra | 29856 | 99.9833 | 99.9721 | 99.98 | B.1 | 95 | 95 | UK, USA, Australia | 7440 | January-24, May-21 | 48 | A2a | 1 | B.1 | 0.99 | A | 0.233 | A1.1 |
| 13 | 420552 | EPI_ISL_420552 | Maharashtra | 29892 | 99.9833 | 99.9721 | 99.98 | B.1 | 97 | 94 | UK, USA, Australia | 7440 | January-24, May-21 | 48 | A2a | 1 | B.1 | 0.988 | A | 0.233 | A1.1 |
| 14 | 420553 | EPI_ISL_420553 | Maharashtra | 29871 | 99.9799 | 99.9721 | 99.98 | B.1 | 96 | 95 | UK, USA, Australia | 7440 | January-24, May-21 | 48 | A2a | 1 | B.1 | 0.989 | A | 0.233 | A1.1 |
| 15 | 420554 | EPI_ISL_420554 | Maharashtra | 29882 | 99.9799 | 99.9721 | 99.98 | B.1 | 95 | 96 | UK, USA, Australia | 7440 | January-24, May-21 | 48 | A2a | 1 | B.1 | 0.988 | A | 0.233 | A1.1 |
| 16 | 420555 | EPI_ISL_420555 | Maharashtra | 29860 | 99.9732 | 99.9651 | 99.97 | B.1.1 | 96 | 100 | UK, Australia, USA | 6287 | February-15, May-20 | 49 | A2a | 1 | B.1.1 | 1 | A | 0.232 | A1 |
| 17 | 420556 | EPI_ISL_420556 | Maharashtra | 29897 | 99.9732 | 99.9651 | 99.97 | B.1.1 | 96 | 96 | UK, Australia, USA | 6287 | February-15, May-20 | 49 | A2a | 1 | B.1.1 | 1 | A | 0.232 | A1 |
| 32 | 424364 | EPI_ISL_424364 | Maharashtra | 29857 | 99.9766 | 99.9791 | 99.98 | B.1 | 95 | 96 | UK, USA, Australia | 7440 | January-24, May-21 | 48 | A2a | 1 | B.1 | 1 | A | 0.233 | A1.2 |
| 33 | 424365 | EPI_ISL_424365 | Maharashtra | 29862 | 99.9766 | 99.9791 | 99.98 | B.1 | 96 | 94 | UK, USA, Australia | 7440 | January-24, May-21 | 48 | A2a | 1 | B.1 | 0.999 | A | 0.233 | A1.2 |
| 34 | 426179 | EPI_ISL_426179 | Maharashtra | 29860 | 99.9732 | 99.9651 | 99.97 | B.1.1 | 96 | 100 | UK, Australia, USA | 6287 | February-15, May-20 | 49 | A2a | 1 | B.1.1 | 1 | A | 0.232 | A1 |
| 37 | 428479 | EPI_ISL_428479 | Karnataka | 29839 | 99.9591 | 99.9224 | 99.95 | B.1.1 | 94 | 94 | UK, Australia, USA | 6287 | February-15, May-20 | 49 | A2a | 0.988 | B.1.1 | 0.761 | A | 0.236 | A1 |
| 42 | 428484 | EPI_ISL_428484 | Karnataka | 29836 | 99.9727 | 99.9223 | 99.97 | B | 100 | 100 | UK, USA, China | 1714 | December-24, May-18 | 51 | A3 | 0.253 | B.6 | 0.674 | A | 0.236 | A1 |
| 46 | 430464 | EPI_ISL_430464 | West Bengal | 29858 | 99.9732 | 99.9581 | 99.97 | B.1.1 | 95 | 95 | UK, Australia, USA | 6287 | February-15, May-20 | 49 | A2a | 1 | B.1.1 | 0.976 | A | 0.232 | A1.3 |
| 47 | 430465 | EPI_ISL_430465 | West Bengal | 29858 | 99.9732 | 99.9651 | 99.97 | B.1.1 | 96 | 96 | UK, Australia, USA | 6287 | February-15, May-20 | 49 | A2a | 1 | B.1.1 | 1 | A | 0.232 | A1.4 |
| 49 | 430467 | EPI_ISL_430467 | West Bengal | 29856 | 99.9766 | 99.9651 | 99.98 | B.1 | 97 | 100 | UK, USA, Australia | 7440 | January-24, May-21 | 48 | A2a | 1 | B.1 | 0.968 | A | 0.232 | A1 |
| 50 | 430468 | EPI_ISL_430468 | West Bengal | 29855 | 99.9732 | 99.9581 | 99.97 | B.1.1 | 96 | 95 | UK, Australia, USA | 6287 | February-15, May-20 | 49 | A2a | 1 | B.1.1 | 0.981 | A | 0.232 | A1.3 |
| 52 | 431102 | EPI_ISL_431102 | Telangana | 29903 | 99.9866 | 99.986 | 99.99 | B.1 | 99 | 100 | UK, USA, Australia | 6733 | January-24, April-23 | 25 | A2a | 1 | B.1 | 1 | A | 0.233 | A1 |
| 53 | 431103 | EPI_ISL_431103 | Telangana | 29903 | 99.9665 | 99.9233 | 99.97 | B.6 | 100 | 100 | Australia, Singapore, USA | 94 | March-04, April-15 | 33 | A3 | 0.291 | B.6 | 0.997 | A | 0.233 | A1 |
| 54 | 431117 | EPI_ISL_431117 | Telangana | 29903 | 99.9732 | 99.9651 | 99.97 | B.1 | 100 | 100 | UK, USA, Australia | 6733 | January-24, April-23 | 25 | A2a | 1 | B.1.1 | 1 | A | 0.232 | A1 |
| 55 | 435049 | EPI_ISL_435049 | Gujarat | 29903 | 99.9866 | 99.9791 | 99.94 | B.1 | 99 | 100 | UK, USA, Australia | 6733 | January-24, April-23 | 25 | A2a | 1 | B.1 | 1 | A | 0.233 | A1 |
| 60 | 435054 | EPI_ISL_435054 | Gujarat | 29903 | 99.9732 | 99.9581 | 99.97 | B.1 | 99 | 100 | UK, USA, Australia | 6733 | January-24, April-23 | 25 | A2a | 1 | B.1 | 1 | A | 0.233 | A1 |
| 61 | 435055 | EPI_ISL_435055 | Gujarat | 29903 | 99.9631 | 99.9581 | 99.96 | B.1.36 | 97 | 100 | Turkey, Finland, UK | 32 | March-10, April-01 | 47 | A2a | 1 | B.1.36 | 0.901 | A | 0.232 | A1.6.1.1.1.1 |
| 62 | 435056 | EPI_ISL_435056 | Gujarat | 29903 | 99.9295 | 99.8883 | 99.84 | B.1 | 96 | 100 | UK, USA, Australia | 6733 | January-24, April-23 | 25 | A2a | 1 | B.1 | 0.937 | A | 0.233 | A1.5 |
| 66 | 435063 | EPI_ISL_435063 | Delhi | 29866 | 99.9464 | 99.8814 | 99.95 | B.1 | 100 | 100 | UK, USA, Australia | 7440 | January-24, May-21 | 48 | A2a | 1 | B.1 | 0.973 | A | 0.231 | A1.6 |
| 67 | 435064 | EPI_ISL_435064 | Delhi | 29868 | 99.9665 | 99.9163 | 99.97 | B.1 | 100 | 100 | UK, USA, Australia | 7440 | January-24, May-21 | 48 | A2a | 0.997 | B.1 | 0.994 | A | 0.233 | A1.6 |
| 68 | 435065 | EPI_ISL_435065 | Delhi | 29867 | 99.8962 | 99.7837 | 99.9 | B.1.36 | 98 | 100 | Saudi_Arabia, UK, Turkey | 159 | March-10, May-07 | 62 | A2a | 0.975 | B.1.36 | 0.784 | A | 0.232 | A1.6.1.1 |
| 69 | 435066 | EPI_ISL_435066 | Delhi | 29866 | 99.933 | 99.9163 | 99.93 | B.1.36 | 99 | 100 | Saudi_Arabia, UK, Turkey | 159 | March-10, May-07 | 62 | A2a | 1 | B.1.36 | 0.908 | A | 0.233 | A1.6.1.1 |
| 71 | 435068 | EPI_ISL_435068 | Delhi | 29866 | 99.9531 | 99.9302 | 99.95 | B.1.36 | 98 | 100 | Saudi_Arabia, UK, Turkey | 159 | March-10, May-07 | 62 | A2a | 1 | B.1.36 | 0.922 | A | 0.231 | A1.6.1.1 |
| 72 | 435069 | EPI_ISL_435069 | Delhi | 29871 | 99.9397 | 99.8884 | 99.94 | B.1.36 | 98 | 100 | Saudi_Arabia, UK, Turkey | 159 | March-10, May-07 | 62 | A2a | 0.997 | B.1.36 | 0.909 | A | 0.232 | A1.6.1.1 |
| 73 | 435070 | EPI_ISL_435070 | Delhi | 29866 | 99.8493 | 99.6651 | 99.85 | B.1.1 | 99 | 95 | UK, Australia, USA | 6287 | February-15, May-20 | 49 | A2a | 0.992 | B.1.1 | 0.936 | A | 0.23 | A1 |
| 74 | 435071 | EPI_ISL_435071 | Delhi | 29866 | 99.9565 | 99.9302 | 99.96 | B.1.1 | 95 | 93 | UK, Australia, USA | 6287 | February-15, May-20 | 49 | A2a | 1 | B.1.1 | 0.983 | A | 0.232 | A1 |
| 76 | 435074 | EPI_ISL_435074 | West Bengal | 29811 | 99.9698 | 99.9302 | 99.97 | B.6 | 100 | 100 | India, Singapore, Australia | 221 | March-04, April-05 | -636 | A6 | 0.274 | B.6 | 0.974 | A | 0.234 | A1 |
| 77 | 435078 | EPI_ISL_435078 | Tamil Nadu | 29866 | 99.8995 | 99.7488 | 99.9 | B | 100 | 100 | UK, USA, China | 1714 | December-24, May-18 | 51 | A7 | 0.265 | B.6 | 0.825 | A | 0.232 | A1 |
| 78 | 435080 | EPI_ISL_435080 | Tamil Nadu | 29866 | 99.8717 | 99.6772 | 99.88 | B.6 | 100 | 95 | India, Singapore, Australia | 221 | March-04, April-05 | -636 | A3 | 0.262 | B.6 | 0.878 | A | 0.232 | A1 |
| 79 | 435081 | EPI_ISL_435081 | West Bengal | 29867 | 99.9464 | 99.8395 | 99.95 | B.6 | 100 | 96 | India, Singapore, Australia | 221 | March-04, April-05 | -636 | A2 | 0.303 | B.6 | 0.877 | A | 0.231 | A1 |
| 81 | 435083 | EPI_ISL_435083 | Tamil Nadu | 29866 | 99.9328 | 99.8318 | 99.92 | B.6 | 98 | 96 | India, Singapore, Australia | 221 | March-04, April-05 | -636 | A3 | 0.258 | B.6 | 0.873 | A | 0.234 | A1 |
| 82 | 435084 | EPI_ISL_435084 | Tamil Nadu | 29866 | 99.9094 | 99.8393 | 99.94 | B | 100 | 100 | UK, USA, China | 1714 | December-24, May-18 | 51 | A3 | 0.273 | B.6 | 0.933 | A | 0.231 | A1 |
| 83 | 435085 | EPI_ISL_435085 | Maharashtra | 29866 | 99.9196 | 99.7767 | 99.92 | B | 100 | 100 | UK, USA, China | 1714 | December-24, May-18 | 51 | A3 | 0.325 | B.6 | 0.915 | A | 0.233 | A1 |
| 84 | 435086 | EPI_ISL_435086 | Maharashtra | 29867 | 99.9163 | 99.8256 | 99.92 | B.6 | 99 | 100 | India, Singapore, Australia | 221 | March-04, April-05 | -636 | A3 | 0.318 | B.6 | 0.92 | A | 0.231 | A1 |
| 85 | 435087 | EPI_ISL_435087 | Tamil Nadu | 29866 | 99.902 | 99.7672 | 99.89 | B.6 | 100 | 96 | India, Singapore, Australia | 221 | March-04, April-05 | -636 | A3 | 0.3 | B.6 | 0.909 | A | 0.231 | A1 |
| 86 | 435088 | EPI_ISL_435088 | Odisha | 29866 | 99.9055 | 99.7263 | 99.91 | B.6 | 100 | 97 | India, Singapore, Australia | 221 | March-04, April-05 | -636 | A3 | 0.243 | B.6 | 0.953 | A | 0.24 | A1 |
| 87 | 435090 | EPI_ISL_435090 | Jammu | 29867 | 99.9397 | 99.8395 | 99.87 | B.6 | 100 | 100 | India, Singapore, Australia | 221 | March-04, April-05 | -636 | A7 | 0.255 | B.6 | 0.985 | A | 0.23 | A1 |
| 88 | 435091 | EPI_ISL_435091 | Tamil Nadu | 29867 | 99.9531 | 99.8744 | 99.95 | B.6 | 100 | 100 | India, Singapore, Australia | 221 | March-04, April-05 | -636 | A3 | 0.293 | B.6 | 0.998 | A | 0.232 | A1 |
| 89 | 435092 | EPI_ISL_435092 | Tamil Nadu | 29866 | 99.9297 | 99.8395 | 99.93 | B.6 | 99 | 100 | India, Singapore, Australia | 221 | March-04, April-05 | -636 | A3 | 0.322 | B.6 | 0.988 | A | 0.23 | A1 |
| 90 | 435093 | EPI_ISL_435093 | Tamil Nadu | 29866 | 99.9431 | 99.8674 | 99.94 | B.6 | 100 | 100 | India, Singapore, Australia | 221 | March-04, April-05 | -636 | A3 | 0.266 | B.6 | 0.963 | A | 0.231 | A1 |
| 91 | 435094 | EPI_ISL_435094 | Tamil Nadu | 29866 | 99.9632 | 99.9163 | 99.96 | B.6 | 100 | 100 | India, Singapore, Australia | 221 | March-04, April-05 | -636 | A3 | 0.278 | B.6 | 0.998 | A | 0.234 | A1 |
| 92 | 435095 | EPI_ISL_435095 | Tamil Nadu | 29866 | 99.9598 | 99.9093 | 99.96 | B.6 | 99 | 97 | India, Singapore, Australia | 221 | March-04, April-05 | -636 | A3 | 0.292 | B.6 | 0.999 | A | 0.233 | A1 |
| 93 | 435096 | EPI_ISL_435096 | Tamil Nadu | 29866 | 99.9699 | 99.9302 | 99.97 | B.6 | 100 | 100 | India, Singapore, Australia | 221 | March-04, April-05 | -636 | A3 | 0.299 | B.6 | 0.999 | A | 0.235 | A1 |
| 94 | 435097 | EPI_ISL_435097 | West Bengal | 29866 | 99.9699 | 99.9233 | 99.97 | B.6 | 100 | 100 | India, Singapore, Australia | 221 | March-04, April-05 | -636 | A3 | 0.295 | B.6 | 1 | A | 0.232 | A1 |
| 95 | 435098 | EPI_ISL_435098 | Delhi | 29866 | 99.9196 | 99.8186 | 99.92 | B.6 | 100 | 100 | India, Singapore, Australia | 221 | March-04, April-05 | -636 | A3 | 0.285 | B.6 | 0.953 | A | 0.234 | A1 |
| 96 | 435099 | EPI_ISL_435099 | Uttar Pradesh | 29873 | 99.9565 | 99.8814 | 99.96 | B.6 | 100 | 100 | India, Singapore, Australia | 221 | March-04, April-05 | -636 | A3 | 0.322 | B.6 | 1 | A | 0.232 | A1 |
| 97 | 435100 | EPI_ISL_435100 | Uttar Pradesh | 29866 | 99.9531 | 99.9093 | 99.95 | B.6 | 99 | 97 | India, Singapore, Australia | 221 | March-04, April-05 | -636 | A3 | 0.311 | B.6 | 0.99 | A | 0.233 | A1 |
| 98 | 435101 | EPI_ISL_435101 | Ladakh | 29866 | 99.9565 | 99.9023 | 99.96 | B.4 | 100 | 100 | Australia, UK, Turkey | 258 | January-18, April-14 | 85 | A3 | 0.986 | B.4 | 0.972 | A | 0.233 | A1.23 |
| 99 | 435102 | EPI_ISL_435102 | Ladakh | 29866 | 99.9262 | 99.8254 | 99.93 | B.4 | 100 | 100 | Australia, UK, Turkey | 258 | January-18, April-14 | 85 | A3 | 0.935 | B.4 | 0.914 | A | 0.233 | A1.23 |
| 100 | 435103 | EPI_ISL_435103 | Ladakh | 29866 | 99.9464 | 99.8605 | 99.95 | B.4 | 100 | 100 | Australia, UK, Turkey | 258 | January-18, April-14 | 85 | A3 | 0.92 | B.4 | 0.918 | A | 0.233 | A1.23 |
| 108 | 435112 | EPI_ISL_435112 | Bihar | 29866 | 99.9297 | 99.8674 | 99.93 | B.6 | 100 | 100 | India, Singapore, Australia | 221 | March-04, April-05 | -636 | A3 | 0.273 | B.6 | 0.955 | A | 0.234 | A1 |
| 109 | 436413 | EPI_ISL_436413 | Uttar Pradesh | 29866 | 99.9358 | 99.8597 | 99.95 | B.1.1 | 97 | 100 | UK, Australia, USA | 6287 | February-15, May-20 | 49 | A2a | 0.999 | B.1.1 | 0.976 | A | 0.23 | A1 |
| 110 | 436414 | EPI_ISL_436414 | West Bengal | 29866 | 99.9297 | 99.8535 | 99.93 | B.6 | 100 | 100 | India, Singapore, Australia | 221 | March-04, April-05 | -636 | A1a | 0.296 | B.6 | 0.516 | A | 0.234 | A1.7 |
| 112 | 436417 | EPI_ISL_436417 | Bihar | 29870 | 99.9561 | 99.9018 | 99.96 | B.6 | 100 | 100 | India, Singapore, Australia | 221 | March-04, April-05 | -636 | A3 | 0.3 | B.6 | 0.964 | A | 0.234 | A1 |
| 113 | 436418 | EPI_ISL_436418 | Tamil Nadu | 29866 | 99.9161 | 99.7834 | 99.93 | B.6 | 100 | 100 | India, Singapore, Australia | 221 | March-04, April-05 | -636 | A3 | 0.274 | B.6 | 0.973 | A | 0.233 | A1 |
| 115 | 436420 | EPI_ISL_436420 | Rajasthan | 29866 | 99.9498 | 99.8744 | 99.95 | B.6 | 100 | 100 | India, Singapore, Australia | 221 | March-04, April-05 | -636 | A3 | 0.289 | B.6 | 0.99 | A | 0.233 | A1 |
| 116 | 436421 | EPI_ISL_436421 | Assam | 29866 | 99.9296 | 99.8116 | 99.93 | B.6 | 97 | 95 | India, Singapore, Australia | 221 | March-04, April-05 | -636 | A1a | 0.294 | B.6 | 0.86 | A | 0.234 | A1.7 |
| 117 | 436422 | EPI_ISL_436422 | Assam | 29866 | 99.9196 | 99.7907 | 99.92 | B.6 | 100 | 100 | India, Singapore, Australia | 221 | March-04, April-05 | -636 | A3 | 0.257 | B.6 | 0.997 | A | 0.234 | A1 |
| 118 | 436424 | EPI_ISL_436424 | Delhi | 29866 | 99.9029 | 99.7767 | 99.9 | B.6 | 100 | 100 | India, Singapore, Australia | 221 | March-04, April-05 | -636 | A3 | 0.243 | B.6 | 0.902 | A | 0.233 | A1 |
| 120 | 436426 | EPI_ISL_436426 | Delhi | 29866 | 99.9196 | 99.8535 | 99.92 | B.6 | 100 | 100 | India, Singapore, Australia | 221 | March-04, April-05 | -636 | A3 | 0.253 | B.6 | 0.959 | A | 0.234 | A1.8 |
| 121 | 436428 | EPI_ISL_436428 | Delhi | 29866 | 99.9363 | 99.8744 | 99.94 | B.6 | 99 | 100 | India, Singapore, Australia | 221 | March-04, April-05 | -636 | A3 | 0.266 | B.6 | 0.955 | A | 0.234 | A1 |
| 122 | 436429 | EPI_ISL_436429 | Delhi | 29866 | 99.9129 | 99.8116 | 99.91 | B.6 | 98 | 93 | India, Singapore, Australia | 221 | March-04, April-05 | -636 | A3 | 0.253 | B.6 | 0.894 | A | 0.234 | A1 |
| 123 | 436430 | EPI_ISL_436430 | Delhi | 29866 | 99.953 | 99.8743 | 99.96 | B.6 | 99 | 100 | India, Singapore, Australia | 221 | March-04, April-05 | -636 | A3 | 0.276 | B.6 | 0.979 | A | 0.234 | A1 |
| 124 | 436431 | EPI_ISL_436431 | Delhi | 29866 | 99.9364 | 99.8395 | 99.94 | B.6 | 100 | 100 | India, Singapore, Australia | 221 | March-04, April-05 | -636 | A3 | 0.268 | B.6 | 0.993 | A | 0.233 | A1 |
| 126 | 436433 | EPI_ISL_436433 | Delhi | 29866 | 99.9296 | 99.8465 | 99.93 | B.6 | 100 | 100 | India, Singapore, Australia | 221 | March-04, April-05 | -636 | A3 | 0.277 | B.6 | 0.96 | A | 0.233 | A1 |
| 128 | 436435 | EPI_ISL_436435 | Delhi | 29866 | 99.8996 | 99.7558 | 99.9 | B.6 | 100 | 97 | India, Singapore, Australia | 221 | March-04, April-05 | -636 | A3 | 0.292 | B.6 | 0.91 | A | 0.232 | A1 |
| 129 | 436436 | EPI_ISL_436436 | Delhi | 29866 | 99.8851 | 99.7052 | 99.89 | B.6 | 97 | 92 | India, Singapore, Australia | 221 | March-04, April-05 | -636 | A3 | 0.21 | B.6 | 0.786 | A | 0.235 | A1 |
| 130 | 436437 | EPI_ISL_436437 | Delhi | 29866 | 99.8822 | 99.7343 | 99.89 | B.6 | 98 | 96 | India, Singapore, Australia | 221 | March-04, April-05 | -636 | A1a | 0.285 | B.6 | 0.838 | A | 0.236 | A1 |
| 133 | 436445 | EPI_ISL_436445 | Delhi | 29866 | 99.9062 | 99.7558 | 99.91 | B.6 | 100 | 100 | India, Singapore, Australia | 221 | March-04, April-05 | -636 | A7 | 0.28 | B.6 | 0.887 | A | 0.234 | A1 |
| 134 | 436447 | EPI_ISL_436447 | Karnataka | 29873 | 99.9397 | 99.8814 | 99.94 | B.6 | 100 | 100 | India, Singapore, Australia | 221 | March-04, April-05 | -636 | A3 | 0.329 | B.6 | 0.991 | A | 0.233 | A1 |
| 135 | 436448 | EPI_ISL_436448 | Delhi | 29874 | 99.9531 | 99.8744 | 99.95 | B.6 | 100 | 100 | India, Singapore, Australia | 221 | March-04, April-05 | -636 | A1a | 0.3 | B.6 | 0.957 | A | 0.234 | A1 |
| 136 | 436449 | EPI_ISL_436449 | Bihar | 29868 | 99.933 | 99.8395 | 99.93 | B.6 | 100 | 100 | India, Singapore, Australia | 221 | March-04, April-05 | -636 | A7 | 0.257 | B.6 | 0.957 | A | 0.233 | A1.8 |
| 138 | 436451 | EPI_ISL_436451 | Delhi | 29866 | 99.9397 | 99.8535 | 99.94 | B.6 | 100 | 100 | India, Singapore, Australia | 221 | March-04, April-05 | -636 | A3 | 0.272 | B.6 | 0.995 | A | 0.233 | A1 |
| 140 | 436453 | EPI_ISL_436453 | Madhya Pradesh | 29867 | 99.9189 | 99.8386 | 99.92 | B.1 | 99 | 100 | UK, USA, Australia | 7440 | January-24, May-21 | 48 | A2a | 0.985 | B.1 | 0.885 | A | 0.238 | A1 |
| 142 | 436455 | EPI_ISL_436455 | Delhi | 29870 | 99.9598 | 99.9233 | 99.96 | B.1 | 99 | 100 | UK, USA, Australia | 7440 | January-24, May-21 | 48 | A2a | 1 | B.1 | 0.96 | A | 0.234 | A1 |
| 147 | 436460 | EPI_ISL_436460 | Madhya Pradesh | 29866 | 99.933 | 99.8674 | 99.93 | B.1 | 100 | 100 | UK, USA, Australia | 7440 | January-24, May-21 | 48 | A2a | 0.964 | B.1 | 0.896 | A | 0.233 | A1 |
| 148 | 436461 | EPI_ISL_436461 | Madhya Pradesh | 29866 | 99.9598 | 99.9233 | 99.96 | B.1 | 100 | 100 | UK, USA, Australia | 7440 | January-24, May-21 | 48 | A2a | 0.998 | B.1 | 0.978 | A | 0.234 | A1 |
| 149 | 436462 | EPI_ISL_436462 | Madhya Pradesh | 29870 | 99.9531 | 99.9302 | 99.95 | B.1 | 100 | 100 | UK, USA, Australia | 7440 | January-24, May-21 | 48 | A2a | 1 | B.1 | 0.982 | A | 0.233 | A1 |
| 150 | 436463 | EPI_ISL_436463 | Madhya Pradesh | 29866 | 99.8817 | 99.6702 | 99.87 | B.1 | 93 | 92 | UK, USA, Australia | 7440 | January-24, May-21 | 48 | A2a | 0.733 | B.1 | 0.702 | A | 0.231 | A1 |
| 154 | 437442 | EPI_ISL_437442 | Gujarat | 29903 | 99.936 | 99.8597 | 99.94 | B.6 | 100 | 96 | Australia, Singapore, USA | 94 | March-04, April-15 | 33 | A1a | 0.293 | B.6 | 0.94 | A | 0.229 | A1.24 |
| 155 | 437444 | EPI_ISL_437444 | Gujarat | 29903 | 99.9732 | 99.9512 | 99.97 | B.6 | 99 | 96 | Australia, Singapore, USA | 94 | March-04, April-15 | 33 | A1a | 0.255 | B.6 | 0.978 | A | 0.233 | A1.24 |
| 158 | 437447 | EPI_ISL_437447 | Gujarat | 29903 | 99.9798 | 99.958 | 99.92 | B.1 | 95 | 100 | UK, USA, Australia | 6733 | January-24, April-23 | 25 | A2a | 0.999 | B.1 | 0.986 | A | 0.233 | A1.9 |
| 159 | 437448 | EPI_ISL_437448 | Gujarat | 29903 | 99.9832 | 99.986 | 99.98 | B.1 | 99 | 100 | UK, USA, Australia | 6733 | January-24, April-23 | 25 | A2a | 1 | B.1 | 0.848 | A | 0.233 | A1.9 |
| 160 | 437449 | EPI_ISL_437449 | Gujarat | 29903 | 99.9698 | 99.9651 | 99.97 | B.1.36 | 76 | 100 | Turkey, Finland, UK | 32 | March-10, April-01 | 47 | A2a | 1 | B.1.36 | 0.79 | A | 0.233 | A1.6.1.5 |
| 166 | 437539 | EPI_ISL_437539 | West Bengal | 29903 | 99.9799 | 99.9791 | 99.98 | B.1 | 98 | 100 | UK, USA, Australia | 6733 | January-24, April-23 | 25 | A2a | 1 | B.1 | 0.98 | A | 0.234 | A1 |
| 167 | 437626 | EPI_ISL_437626 | Telangana | 29903 | 99.9699 | 99.9721 | 99.97 | B.1 | 100 | 100 | UK, USA, Australia | 6733 | January-24, April-23 | 25 | A2a | 1 | B.1 | 1 | A | 0.233 | A1.6 |
| 168 | 438138 | EPI_ISL_438138 | Telangana | 29903 | 99.9732 | 99.9512 | 99.97 | B.6 | 100 | 100 | Australia, Singapore, USA | 94 | March-04, April-15 | 33 | A3 | 0.309 | B.6 | 1 | A | 0.232 | A1 |
| 169 | 438139 | EPI_ISL_438139 | Telangana | 29903 | 99.9799 | 99.9512 | 99.98 | B.6 | 100 | 100 | India, Singapore, Australia | 221 | March-04, April-05 | -636 | A3 | 0.315 | B.6 | 1 | A | 0.233 | A1 |

| | | | | | | | | | | | | | | | | | | | | | |
|---|---|---|---|---|---|---|---|---|---|---|---|---|---|---|---|---|---|---|---|---|---|
| 170 | 444456 | EPI_ISL_444456 | Gujarat | 29903 | 99.9732 | 99.9651 | 99.97 | B.1.36 | 80 | 100 | Turkey, Finland, UK | 32 | March-10, April-01 | 47 | A2a | 1 | B.1.36 | 0.513 | A | 0.233 | A1.6.1.2 |
| 171 | 444457 | EPI_ISL_444457 | Gujarat | 29903 | 99.9732 | 99.9651 | 99.97 | B.1.36 | 76 | 100 | Turkey, Finland, UK | 32 | March-10, April-01 | 47 | A2a | 1 | B.1.36 | 0.527 | A | 0.233 | A1.6.1.2 |
| 174 | 444460 | EPI_ISL_444460 | Gujarat | 29903 | 99.9833 | 99.986 | 99.98 | B.1 | 98 | 100 | UK, USA, Australia | 6733 | January-24, April-23 | 25 | A2a | 0.999 | B.1 | 0.844 | A | 0.233 | A1.9 |
| 175 | 444461 | EPI_ISL_444461 | Gujarat | 29903 | 99.9665 | 99.9651 | 99.97 | B.1.36 | 97 | 100 | Turkey, Finland, UK | 32 | March-10, April-01 | 47 | A2a | 1 | B.1.36 | 0.904 | A | 0.232 | A1.6.1.1.1 |
| 179 | 444465 | EPI_ISL_444465 | Gujarat | 29903 | 99.9598 | 99.9581 | 99.96 | B.1.36 | 99 | 100 | Turkey, Finland, UK | 32 | March-10, April-01 | 47 | A2a | 1 | B.1.36 | 0.866 | A | 0.232 | A1.6.1.1.1.2 |
| 180 | 444466 | EPI_ISL_444466 | Gujarat | 29903 | 99.9631 | 99.9581 | 99.96 | B.1.36 | 96 | 100 | Turkey, Finland, UK | 32 | March-10, April-01 | 47 | A2a | 1 | B.1.36 | 0.868 | A | 0.232 | A1.6.1.1.1.2 |
| 183 | 444469 | EPI_ISL_444469 | Gujarat | 29903 | 99.9564 | 99.9372 | 99.96 | B.1.36 | 99 | 100 | Turkey, Finland, UK | 32 | March-10, April-01 | 47 | A2a | 1 | B.1.36 | 0.868 | A | 0.231 | A1.6.1.1.2 |
| 184 | 444470 | EPI_ISL_444470 | Gujarat | 29903 | 99.9665 | 99.9651 | 99.97 | B.1.36 | 69 | 100 | Turkey, Finland, UK | 32 | March-10, April-01 | 47 | A2a | 1 | B.1.36 | 0.708 | A | 0.233 | A1.6.1.5 |
| 185 | 444471 | EPI_ISL_444471 | Gujarat | 29903 | 99.9765 | 99.9791 | 99.98 | B.1.36 | 82 | 100 | Turkey, Finland, UK | 32 | March-10, April-01 | 47 | A2a | 1 | B.1.36 | 0.514 | A | 0.233 | A1.6.1 |
| 186 | 444472 | EPI_ISL_444472 | Gujarat | 29903 | 99.9665 | 99.9651 | 99.97 | B.1.36 | 97 | 100 | Turkey, Finland, UK | 32 | March-10, April-01 | 47 | A2a | 1 | B.1.36 | 0.903 | A | 0.232 | A1.6.1.1.1 |
| 193 | 444479 | EPI_ISL_444479 | Gujarat | 29903 | 99.9799 | 99.9791 | 99.98 | B.1 | 95 | 100 | UK, USA, Australia | 6733 | January-24, April-23 | 25 | A2a | 1 | B.1 | 0.992 | A | 0.233 | A1.10 |
| 195 | 444481 | EPI_ISL_444481 | Gujarat | 29903 | 99.9866 | 99.986 | 99.99 | B.1 | 100 | 100 | UK, USA, Australia | 6733 | January-24, April-23 | 25 | A2a | 1 | B.1 | 1 | A | 0.233 | A1 |
| 196 | 444482 | EPI_ISL_444482 | Gujarat | 29903 | 99.9732 | 99.9651 | 99.97 | B.1.36 | 78 | 100 | Turkey, Finland, UK | 32 | March-10, April-01 | 47 | A2a | 0.996 | B.1.36 | 0.505 | A | 0.232 | A1.6.1 |
| 197 | 444483 | EPI_ISL_444483 | Gujarat | 29903 | 99.9833 | 99.9721 | 99.98 | B.1 | 99 | 100 | UK, USA, Australia | 6733 | January-24, April-23 | 25 | A2a | 0.999 | B.1 | 0.998 | A | 0.233 | A1.11 |
| 198 | 444484 | EPI_ISL_444484 | Gujarat | 29903 | 99.9698 | 99.972 | 99.97 | B.1.36 | 67 | 77 | Turkey, Finland, UK | 32 | March-10, April-01 | 47 | A2a | 1 | B.1 | 0.478 | A | 0.233 | A1.6.1.3 |
| 199 | 444485 | EPI_ISL_444485 | Gujarat | 29903 | 99.9698 | 99.958 | 99.97 | B.1.36 | 67 | 77 | Turkey, Finland, UK | 32 | March-10, April-01 | 47 | A2a | 1 | B.1 | 0.524 | A | 0.233 | A1.6.1.3 |
| 202 | 447031 | EPI_ISL_447031 | Gujarat | 29903 | 99.9866 | 99.986 | 99.99 | B.1 | 99 | 100 | UK, USA, Australia | 6733 | January-24, April-23 | 25 | A2a | 1 | B.1 | 1 | A | 0.233 | A1 |
| 203 | 447032 | EPI_ISL_447032 | Gujarat | 29903 | 99.9798 | 99.9511 | 99.98 | B.1 | 97 | 100 | UK, USA, Australia | 6733 | January-24, April-23 | 25 | A2a | 1 | B.1 | 0.964 | A | 0.232 | A1 |
| 204 | 447033 | EPI_ISL_447033 | Gujarat | 29903 | 99.9598 | 99.9512 | 99.96 | B.1.36 | 96 | 100 | Turkey, Finland, UK | 32 | March-10, April-01 | 47 | A2a | 1 | B.1.36 | 0.807 | A | 0.23 | A1.6.1.1.1.3 |
| 205 | 447034 | EPI_ISL_447034 | Gujarat | 29903 | 99.9529 | 99.93 | 99.97 | B.1.36 | 96 | 100 | Turkey, Finland, UK | 32 | March-10, April-01 | 53 | A2a | 1 | B.1.36 | 0.672 | A | 0.232 | A1.6.1.1.2 |
| 206 | 447035 | EPI_ISL_447035 | Gujarat | 29903 | 99.9631 | 99.9581 | 99.96 | B.1.36 | 99 | 100 | Turkey, Finland, UK | 32 | March-10, April-01 | 47 | A2a | 1 | B.1.36 | 0.857 | A | 0.232 | A1.6.1.1.1.2 |
| 209 | 447038 | EPI_ISL_447038 | Gujarat | 29903 | 99.9799 | 99.9651 | 99.98 | B.1 | 98 | 100 | UK, USA, Australia | 6733 | January-24, April-23 | 25 | A2a | 0.998 | B.1 | 0.991 | A | 0.233 | A1 |
| 210 | 447039 | EPI_ISL_447039 | Gujarat | 29903 | 99.9631 | 99.9651 | 99.96 | B.1.36 | 99 | 100 | Turkey, Finland, UK | 32 | March-10, April-01 | 47 | A2a | 1 | B.1.36 | 0.877 | A | 0.232 | A1.6.1.1.1 |
| 211 | 447040 | EPI_ISL_447040 | Gujarat | 29903 | 99.9564 | 99.9512 | 99.96 | B.1.36 | 96 | 100 | Turkey, Finland, UK | 32 | March-10, April-01 | 47 | A2a | 0.986 | B.1.36 | 0.753 | A | 0.232 | A1.6.1.1.1 |
| 212 | 447041 | EPI_ISL_447041 | Gujarat | 29903 | 99.9698 | 99.9721 | 99.97 | B.1.36 | 69 | 100 | Turkey, Finland, UK | 32 | March-10, April-01 | 47 | A2a | 0.996 | B.1 | 0.669 | A | 0.232 | A1.6.1.1 |
| 215 | 447044 | EPI_ISL_447044 | Gujarat | 29903 | 99.9598 | 99.9512 | 99.96 | B.1.36 | 95 | 100 | Turkey, Finland, UK | 32 | March-10, April-01 | 47 | A2a | 1 | B.1.36 | 0.87 | A | 0.232 | A1.6.1.1.2 |
| 218 | 447047 | EPI_ISL_447047 | Gujarat | 29903 | 99.9866 | 99.986 | 99.99 | B.1 | 99 | 100 | UK, USA, Australia | 6733 | January-24, April-23 | 25 | A2a | 1 | B.1 | 1 | A | 0.233 | A1 |
| 221 | 447050 | EPI_ISL_447050 | Gujarat | 29903 | 99.9631 | 99.9581 | 99.96 | B.1.36 | 98 | 100 | Turkey, Finland, UK | 32 | March-10, April-01 | 47 | A2a | 1 | B.1.36 | 0.908 | A | 0.232 | A1.6.1.1.1.1 |
| 222 | 447051 | EPI_ISL_447051 | Gujarat | 29903 | 99.9631 | 99.9581 | 99.96 | B.1.36 | 96 | 100 | Turkey, Finland, UK | 32 | March-10, April-01 | 47 | A2a | 1 | B.1.36 | 0.908 | A | 0.232 | A1.6.1.1.1.1 |
| 223 | 447052 | EPI_ISL_447052 | Gujarat | 29903 | 99.9799 | 99.9791 | 99.98 | B.1 | 95 | 100 | UK, USA, Australia | 6733 | January-24, April-23 | 25 | A2a | 1 | B.1 | 0.992 | A | 0.233 | A1.10 |
| 226 | 447535 | EPI_ISL_447535 | Gujarat | 29903 | 99.9665 | 99.9651 | 99.97 | B.1.36 | 68 | 76 | Turkey, Finland, UK | 32 | March-10, April-01 | 47 | A2a | 0.999 | B.1.36 | 0.493 | A | 0.233 | A1.6.1.3 |
| 227 | 447536 | EPI_ISL_447536 | Gujarat | 29903 | 99.9731 | 99.9581 | 99.83 | B.1.36 | 68 | 76 | Turkey, Finland, UK | 32 | March-10, April-01 | 47 | A2 | 0.494 | B.1.36 | 0.321 | A | 0.234 | A1.6.1.5 |
| 229 | 447538 | EPI_ISL_447538 | Gujarat | 29903 | 99.9731 | 99.9721 | 99.97 | B.1.36 | 65 | 79 | Turkey, Finland, UK | 32 | March-10, April-01 | 47 | A2 | 0.493 | B.1.36 | 0.356 | A | 0.234 | A1.6.1.5 |
| 231 | 447540 | EPI_ISL_447540 | Gujarat | 29903 | 99.9698 | 99.9721 | 99.97 | B.1.36 | 75 | 100 | Turkey, Finland, UK | 32 | March-10, April-01 | 47 | A2a | 1 | B.1 | 0.5 | A | 0.233 | A1.6.1.3 |
| 232 | 447541 | EPI_ISL_447541 | Gujarat | 29903 | 99.9665 | 99.9581 | 99.97 | B.1.36 | 64 | 78 | Turkey, Finland, UK | 32 | March-10, April-01 | 47 | A2a | 0.999 | B.1 | 0.505 | A | 0.233 | A1.6.1.3 |
| 233 | 447542 | EPI_ISL_447542 | Gujarat | 29903 | 99.9665 | 99.9581 | 99.97 | B.1.36 | 68 | 78 | Turkey, Finland, UK | 32 | March-10, April-01 | 47 | A2a | 0.999 | B.1 | 0.501 | A | 0.233 | A1.6.1.3 |
| 234 | 447543 | EPI_ISL_447543 | Gujarat | 29903 | 99.9698 | 99.9721 | 99.97 | B.1.36 | 71 | 100 | Turkey, Finland, UK | 32 | March-10, April-01 | 47 | A2a | 1 | B.1 | 0.496 | A | 0.233 | A1.6.1.3 |
| 235 | 447544 | EPI_ISL_447544 | Gujarat | 29903 | 99.9832 | 99.9721 | 99.98 | B.1 | 99 | 100 | UK, USA, Australia | 6733 | January-24, April-23 | 25 | A2a | 1 | B.1 | 1 | A | 0.233 | A1.11 |
| 236 | 447545 | EPI_ISL_447545 | Gujarat | 29903 | 99.9832 | 99.9721 | 99.98 | B.1 | 98 | 100 | UK, USA, Australia | 6733 | January-24, April-23 | 25 | A2a | 1 | B.1 | 1 | A | 0.233 | A1.11 |
| 237 | 447546 | EPI_ISL_447546 | Gujarat | 29903 | 99.9631 | 99.9512 | 99.96 | B.1.36 | 97 | 100 | Turkey, Finland, UK | 32 | March-10, April-01 | 47 | A2a | 1 | B.1.36 | 0.874 | A | 0.232 | A1.6.1.1.2 |
| 238 | 447547 | EPI_ISL_447547 | Gujarat | 29903 | 99.9799 | 99.9791 | 99.98 | B.1 | 98 | 100 | UK, USA, Australia | 6733 | January-24, April-23 | 25 | A2a | 1 | B.1 | 0.992 | A | 0.233 | A1.10 |
| 239 | 447548 | EPI_ISL_447548 | Gujarat | 29903 | 99.9598 | 99.9512 | 99.96 | B.1.36 | 96 | 100 | Turkey, Finland, UK | 32 | March-10, April-01 | 47 | A2a | 1 | B.1.36 | 0.892 | A | 0.232 | A1.6.1.1.1.1 |
| 240 | 447549 | EPI_ISL_447549 | Gujarat | 29903 | 99.9799 | 99.986 | 99.98 | B.1 | 98 | 100 | UK, USA, Australia | 6733 | January-24, April-23 | 25 | A2a | 0.998 | B.1 | 0.828 | A | 0.232 | A1.9.1 |
| 241 | 447550 | EPI_ISL_447550 | Gujarat | 29903 | 99.9799 | 99.986 | 99.98 | B.1 | 96 | 100 | UK, USA, Australia | 6733 | January-24, April-23 | 25 | A2a | 1 | B.1 | 0.845 | A | 0.234 | A1.9 |
| 242 | 447551 | EPI_ISL_447551 | Gujarat | 29903 | 99.9799 | 99.986 | 99.98 | B.1 | 98 | 100 | UK, USA, Australia | 6733 | January-24, April-23 | 25 | A2a | 0.998 | B.1 | 0.828 | A | 0.232 | A1.9.1 |
| 245 | 447554 | EPI_ISL_447554 | Gujarat | 29903 | 99.9866 | 99.986 | 99.99 | B.1 | 99 | 100 | UK, USA, Australia | 6733 | January-24, April-23 | 25 | A2a | 1 | B.1 | 1 | A | 0.233 | A1 |
| 246 | 447555 | EPI_ISL_447555 | Gujarat | 29903 | 99.9765 | 99.9791 | 99.98 | B.1.36 | 78 | 100 | Turkey, Finland, UK | 32 | March-10, April-01 | 47 | A2a | 1 | B.1.36 | 0.515 | A | 0.233 | A1.6.1 |
| 247 | 447556 | EPI_ISL_447556 | Telangana | 29851 | 99.9766 | 99.9442 | 99.98 | B.6 | 100 | 100 | India, Singapore, Australia | 221 | March-04, April-05 | -636 | A3 | 0.318 | B.6 | 0.986 | A | 0.233 | A1 |
| 248 | 447557 | EPI_ISL_447557 | Telangana | 29865 | 99.9699 | 99.9512 | 99.97 | B.6 | 100 | 100 | India, Singapore, Australia | 221 | March-04, April-05 | -636 | A3 | 0.29 | B.6 | 0.999 | A | 0.233 | A1 |
| 249 | 447558 | EPI_ISL_447558 | Telangana | 29865 | 99.9833 | 99.9651 | 99.98 | B.6 | 89 | 97 | India, Singapore, Australia | 221 | March-04, April-05 | -636 | A7 | 0.381 | B.6 | 0.959 | A | 0.233 | A1 |
| 250 | 447559 | EPI_ISL_447559 | Telangana | 29853 | 99.9766 | 99.9233 | 99.98 | B.6 | 100 | 96 | India, Singapore, Australia | 221 | March-04, April-05 | -636 | A3 | 0.305 | B.6 | 0.97 | A | 0.233 | A1 |
| 251 | 447560 | EPI_ISL_447560 | Telangana | 29864 | 99.9833 | 99.9512 | 99.98 | B.6 | 100 | 100 | India, Singapore, Australia | 221 | March-04, April-05 | -636 | A3 | 0.299 | B.6 | 1 | A | 0.233 | A1 |
| 252 | 447561 | EPI_ISL_447561 | Telangana | 29864 | 99.9732 | 99.9233 | 99.97 | B.6 | 100 | 100 | India, Singapore, Australia | 221 | March-04, April-05 | -636 | A3 | 0.299 | B.6 | 0.995 | A | 0.232 | A1 |
| 253 | 447562 | EPI_ISL_447562 | Telangana | 29858 | 99.9833 | 99.9512 | 99.98 | B.6 | 100 | 100 | India, Singapore, Australia | 221 | March-04, April-05 | -636 | A3 | 0.294 | B.6 | 1 | A | 0.233 | A1 |
| 255 | 447564 | EPI_ISL_447564 | Telangana | 29890 | 99.9833 | 99.9512 | 99.98 | B.6 | 100 | 100 | India, Singapore, Australia | 221 | March-04, April-05 | -636 | A3 | 0.322 | B.6 | 1 | A | 0.233 | A1 |
| 256 | 447565 | EPI_ISL_447565 | Telangana | 29834 | 99.9899 | 99.9791 | 99.99 | B | 100 | 100 | UK, USA, China | 1714 | December-24, May-18 | 51 | A7 | 0.39 | B.6 | 0.697 | A | 0.232 | A1 |
| 257 | 447566 | EPI_ISL_447566 | Telangana | 29876 | 99.9766 | 99.9442 | 99.98 | B.6 | 99 | 100 | India, Singapore, Australia | 221 | March-04, April-05 | -636 | A3 | 0.321 | B.6 | 0.986 | A | 0.233 | A1 |
| 258 | 447567 | EPI_ISL_447567 | Telangana | 29865 | 99.9665 | 99.9372 | 99.97 | B.6 | 100 | 97 | India, Singapore, Australia | 221 | March-04, April-05 | -636 | A3 | 0.291 | B.6 | 0.999 | A | 0.233 | A1 |
| 265 | 447574 | EPI_ISL_447574 | Telangana | 29834 | 99.9832 | 99.9512 | 99.98 | B.6 | 100 | 100 | India, Singapore, Australia | 221 | March-04, April-05 | -636 | A3 | 0.284 | B.6 | 1 | A | 0.233 | A1 |
| 266 | 447575 | EPI_ISL_447575 | Telangana | 29892 | 99.9699 | 99.9233 | 99.97 | B.6 | 99 | 96 | India, Singapore, Australia | 221 | March-04, April-05 | -636 | A3 | 0.318 | B.6 | 0.988 | A | 0.233 | A1 |
| 268 | 447577 | EPI_ISL_447577 | Telangana | 29865 | 99.9799 | 99.9372 | 99.98 | B.6 | 100 | 100 | India, Singapore, Australia | 221 | March-04, April-05 | -636 | A3 | 0.284 | B.6 | 0.996 | A | 0.233 | A1 |
| 273 | 447582 | EPI_ISL_447582 | Telangana | 29865 | 99.9833 | 99.9512 | 99.98 | B.6 | 100 | 100 | India, Singapore, Australia | 221 | March-04, April-05 | -636 | A3 | 0.303 | B.6 | 1 | A | 0.233 | A1 |
| 275 | 447584 | EPI_ISL_447584 | Tamil Nadu | 29866 | 99.9766 | 99.9302 | 99.98 | B.6 | 99 | 96 | India, Singapore, Australia | 221 | March-04, April-05 | -636 | A3 | 0.303 | B.6 | 0.998 | A | 0.233 | A1.12 |
| 276 | 447585 | EPI_ISL_447585 | Tamil Nadu | 29868 | 99.9766 | 99.9302 | 99.98 | B.6 | 99 | 97 | India, Singapore, Australia | 221 | March-04, April-05 | -636 | A3 | 0.306 | B.6 | 0.998 | A | 0.233 | A1.12 |
| 277 | 447586 | EPI_ISL_447586 | Tamil Nadu | 29865 | 99.9699 | 99.9302 | 99.97 | B.6 | 99 | 97 | India, Singapore, Australia | 221 | March-04, April-05 | -636 | A3 | 0.286 | B.6 | 0.998 | A | 0.233 | A1.12 |
| 279 | 447847 | EPI_ISL_447847 | Telangana | 29822 | 99.9732 | 99.9372 | 99.97 | B.6 | 100 | 97 | India, Singapore, Australia | 221 | March-04, April-05 | -636 | A3 | 0.271 | B.6 | 0.991 | A | 0.234 | A1 |
| 282 | 447850 | EPI_ISL_447850 | Telangana | 29822 | 99.9765 | 99.9442 | 99.98 | B.6 | 100 | 100 | India, Singapore, Australia | 221 | March-04, April-05 | -636 | A3 | 0.285 | B.6 | 0.986 | A | 0.233 | A1 |
| 286 | 447854 | EPI_ISL_447854 | Telangana | 29825 | 99.9799 | 99.9372 | 99.98 | B.6 | 100 | 96 | India, Singapore, Australia | 221 | March-04, April-05 | -636 | A3 | 0.276 | B.6 | 1 | A | 0.233 | A1 |
| 287 | 447855 | EPI_ISL_447855 | Telangana | 29822 | 99.9732 | 99.9233 | 99.97 | B.6 | 100 | 100 | India, Singapore, Australia | 221 | March-04, April-05 | -636 | A7 | 0.293 | B.6 | 0.997 | A | 0.232 | A1 |
| 288 | 447856 | EPI_ISL_447856 | Telangana | 29822 | 99.9799 | 99.9442 | 99.98 | B.6 | 100 | 100 | India, Singapore, Australia | 221 | March-04, April-05 | -636 | A3 | 0.27 | B.6 | 0.999 | A | 0.233 | A1 |
| 289 | 447857 | EPI_ISL_447857 | Telangana | 29822 | 99.9732 | 99.9233 | 99.97 | B.6 | 100 | 100 | India, Singapore, Australia | 221 | March-04, April-05 | -636 | A7 | 0.272 | B.6 | 0.999 | A | 0.232 | A1 |
| 290 | 447858 | EPI_ISL_447858 | Telangana | 29822 | 99.9732 | 99.9302 | 99.97 | B.6 | 99 | 96 | India, Singapore, Australia | 221 | March-04, April-05 | -636 | A3 | 0.285 | B.6 | 0.995 | A | 0.233 | A1.25 |
| 292 | 447860 | EPI_ISL_447860 | Telangana | 29834 | 99.9799 | 99.9512 | 99.98 | B.6 | 100 | 100 | India, Singapore, Australia | 221 | March-04, April-05 | -636 | A3 | 0.282 | B.6 | 1 | A | 0.232 | A1 |
| 293 | 447861 | EPI_ISL_447861 | Telangana | 29834 | 99.9832 | 99.9512 | 99.98 | B.6 | 100 | 100 | India, Singapore, Australia | 221 | March-04, April-05 | -636 | A3 | 0.284 | B.6 | 1 | A | 0.233 | A1 |
| 294 | 447862 | EPI_ISL_447862 | Telangana | 29840 | 99.9732 | 99.9233 | 99.97 | B.6 | 100 | 100 | India, Singapore, Australia | 221 | March-04, April-05 | -636 | A7 | 0.276 | B.6 | 0.999 | A | 0.233 | A1 |
| 295 | 447863 | EPI_ISL_447863 | Telangana | 29822 | 99.9732 | 99.9233 | 99.97 | B.6 | 99 | 97 | India, Singapore, Australia | 221 | March-04, April-05 | -636 | A3 | 0.296 | B.6 | 0.983 | A | 0.233 | A1 |
| 297 | 447865 | EPI_ISL_447865 | Telangana | 29834 | 99.9765 | 99.9233 | 99.98 | B.6 | 100 | 96 | India, Singapore, Australia | 221 | March-04, April-05 | -636 | A3 | 0.283 | B.6 | 0.985 | A | 0.234 | A1 |
| 299 | 450321 | EPI_ISL_450321 | Maharashtra | 29821 | 99.9765 | 99.9233 | 99.98 | B.6 | 100 | 97 | India, Singapore, Australia | 221 | March-04, April-05 | -636 | A3 | 0.272 | B.6 | 0.978 | A | 0.233 | A1 |
| 300 | 450322 | EPI_ISL_450322 | Maharashtra | 29821 | 99.9799 | 99.9512 | 99.98 | B.6 | 100 | 100 | India, Singapore, Australia | 221 | March-04, April-05 | -636 | A3 | 0.269 | B.6 | 1 | A | 0.232 | A1 |
| 301 | 450323 | EPI_ISL_450323 | Maharashtra | 29829 | 99.9866 | 99.986 | 99.99 | B.1 | 100 | 100 | UK, USA, Australia | 7440 | January-24, May-21 | 48 | A2a | 1 | B.1 | 1 | A | 0.233 | A1 |
| 304 | 450326 | EPI_ISL_450326 | Telangana | 29830 | 99.9732 | 99.9233 | 99.91 | B.6 | 100 | 100 | India, Singapore, Australia | 221 | March-04, April-05 | -636 | A7 | 0.27 | B.6 | 0.997 | A | 0.233 | A1 |
| 305 | 450327 | EPI_ISL_450327 | Telangana | 29859 | 99.9833 | 99.9512 | 99.98 | B.6 | 100 | 100 | India, Singapore, Australia | 221 | March-04, April-05 | -636 | A3 | 0.289 | B.6 | 1 | A | 0.233 | A1 |
| 306 | 450328 | EPI_ISL_450328 | Telangana | 29834 | 99.9799 | 99.9512 | 99.98 | B.6 | 100 | 100 | India, Singapore, Australia | 221 | March-04, April-05 | -636 | A3 | 0.282 | B.6 | 1 | A | 0.232 | A1 |
| 307 | 450329 | EPI_ISL_450329 | Telangana | 29834 | 99.9832 | 99.9512 | 99.98 | B.6 | 100 | 100 | India, Singapore, Australia | 221 | March-04, April-05 | -636 | A3 | 0.284 | B.6 | 1 | A | 0.233 | A1 |
| 308 | 450330 | EPI_ISL_450330 | Telangana | 29830 | 99.9732 | 99.9372 | 99.97 | B.6 | 100 | 100 | India, Singapore, Australia | 221 | March-04, April-05 | -636 | A3 | 0.273 | B.6 | 0.99 | A | 0.232 | A1 |
| 309 | 450331 | EPI_ISL_450331 | Telangana | 29836 | 99.9832 | 99.9512 | 99.98 | B.6 | 100 | 100 | India, Singapore, Australia | 221 | March-04, April-05 | -636 | A3 | 0.277 | B.6 | 1 | A | 0.233 | A1 |
| 310 | 450332 | EPI_ISL_450332 | Telangana | 29852 | 99.9799 | 99.9372 | 99.98 | B.6 | 99 | 97 | India, Singapore, Australia | 221 | March-04, April-05 | -636 | A3 | 0.296 | B.6 | 1 | A | 0.233 | A1 |
| 315 | 450785 | EPI_ISL_450785 | Gujarat | 29903 | 99.9531 | 99.9023 | 99.95 | B.1 | 100 | 100 | UK, USA, Australia | 6733 | January-24, April-23 | 31 | A2a | 0.989 | B.1.1 | 0.959 | A | 0.232 | A1.14.1 |
| 316 | 450786 | EPI_ISL_450786 | Gujarat | 29903 | 99.9497 | 99.8953 | 99.95 | B.1 | 100 | 100 | UK, USA, Australia | 6733 | January-24, April-23 | 31 | A2a | 0.988 | B.1.1 | 0.957 | A | 0.232 | A1.14.1 |
| 318 | 450788 | EPI_ISL_450788 | Gujarat | 29903 | 99.9765 | 99.9581 | 99.98 | B.1 | 96 | 100 | UK, USA, Australia | 6733 | January-24, April-23 | 31 | A2a | 0.999 | B.1 | 0.995 | A | 0.233 | A1.5 |
| 320 | 450790 | EPI_ISL_450790 | Gujarat | 29903 | 99.9732 | 99.9441 | 99.89 | A | 99 | 100 | China, South_Korea, USA | 241 | January-05, April-06 | 48 | B4 | 0.965 | A.p7 | 0.715 | A | 0.233 | A1.26 |
| 321 | 450791 | EPI_ISL_450791 | Gujarat | 29903 | 99.9732 | 99.9301 | 99.97 | A | 100 | 100 | China, South_Korea, USA | 241 | January-05, April-06 | 48 | B4 | 0.992 | A.p7 | 0.876 | A | 0.231 | A1.26 |
| 322 | 451149 | EPI_ISL_451149 | Gujarat | 29903 | 99.9564 | 99.9512 | 99.96 | B.1.36 | 97 | 100 | Turkey, Finland, UK | 32 | March-10, April-01 | 53 | A2a | 0.968 | B.1.36 | 0.871 | A | 0.233 | A1.6.1.1.1.4 |
| 324 | 451151 | EPI_ISL_451151 | Gujarat | 29903 | 99.9598 | 99.9651 | 99.96 | B.1.36 | 99 | 100 | Turkey, Finland, UK | 32 | March-10, April-01 | 53 | A2a | 0.971 | B.1.36 | 0.889 | A | 0.233 | A1.6.1.1.1.4 |
| 326 | 451153 | EPI_ISL_451153 | Gujarat | 29903 | 99.9598 | 99.9512 | 99.96 | B.1.36 | 95 | 100 | Turkey, Finland, UK | 32 | March-10, April-01 | 53 | A2a | 0.999 | B.1.36 | 0.805 | A | 0.232 | A1.6.1.1.1.3 |
| 327 | 451154 | EPI_ISL_451154 | Gujarat | 29903 | 99.9698 | 99.9302 | 99.92 | B.6 | 100 | 100 | Australia, Singapore, USA | 94 | March-04, April-15 | 39 | A7 | 0.276 | B.6 | 0.983 | A | 0.232 | A1.27 |
| 329 | 451156 | EPI_ISL_451156 | Gujarat | 29903 | 99.9698 | 99.9302 | 99.92 | B.6 | 100 | 100 | Australia, Singapore, USA | 94 | March-04, April-15 | 39 | A7 | 0.275 | B.6 | 0.983 | A | 0.232 | A1.27 |
| 331 | 451158 | EPI_ISL_451158 | Gujarat | 29903 | 99.9832 | 99.986 | 99.98 | B.1 | 98 | 100 | UK, USA, Australia | 6733 | January-24, April-23 | 31 | A2a | 1 | B.1 | 0.988 | A | 0.233 | A1.13 |
| 332 | 451159 | EPI_ISL_451159 | Gujarat | 29903 | 99.9698 | 99.9372 | 99.97 | B.6 | 100 | 100 | Australia, Singapore, USA | 94 | March-04, April-15 | 39 | A7 | 0.281 | B.6 | 0.984 | A | 0.232 | A1.27 |
| 333 | 451160 | EPI_ISL_451160 | Gujarat | 29903 | 99.9765 | 99.9791 | 99.98 | B.1.36 | 73 | 100 | Turkey, Finland, UK | 32 | March-10, April-01 | 53 | A2a | 1 | B.1.36 | 0.515 | A | 0.233 | A1.6.1 |
| 334 | 451161 | EPI_ISL_451161 | Gujarat | 29903 | 99.9698 | 99.9232 | 99.88 | B.6 | 100 | 100 | Australia, Singapore, USA | 94 | March-04, April-15 | 39 | A7 | 0.27 | B.6 | 0.978 | A | 0.23 | A1.27 |
| 338 | 452192 | EPI_ISL_452192 | Maharashtra | 29805 | 99.9631 | 99.9512 | 99.97 | B.1.1 | 95 | 95 | UK, Australia, USA | 6287 | February-15, May-20 | 49 | A2a | 0.998 | B.1.1 | 0.996 | A | 0.232 | A1.14 |
| 344 | 452199 | EPI_ISL_452199 | Maharashtra | 29799 | 99.9564 | 99.9302 | 99.94 | B.1.1 | 97 | 95 | UK, Australia, USA | 6287 | February-15, May-20 | 49 | A2a | 0.994 | B.1.1 | 0.981 | A | 0.232 | A1.14 |
| 349 | 452204 | EPI_ISL_452204 | Maharashtra | 29799 | 99.9732 | 99.9651 | 99.96 | A | 100 | 100 | China, South_Korea, USA | 223 | January-05, April-23 | 76 | B4 | 0.892 | A.p7 | 0.56 | A | 0.232 | A1.28 |

| | | | | | | | | | | | | | | | | | | | | | |
|---|---|---|---|---|---|---|---|---|---|---|---|---|---|---|---|---|---|---|---|---|---|
| 350 | 452205 | EPI_ISL_452205 | Maharashtra | 29805 | 99.9698 | 99.9651 | 99.98 | A | 100 | 100 | China, South_Korea, USA | 223 | January-05, April-23 | 76 | B4 | 0.905 | A.p7 | 0.571 | A | 0.232 | A1.29 |
| 351 | 452207 | EPI_ISL_452207 | Maharashtra | 29805 | 99.9631 | 99.9512 | 99.97 | B.1.1 | 95 | 94 | UK, Australia, USA | 6287 | February-15, May-20 | 49 | A2a | 0.998 | B.1.1 | 0.996 | A | 0.232 | A1.14 |
| 352 | 452208 | EPI_ISL_452208 | Maharashtra | 29805 | 99.9765 | 99.9512 | 99.98 | B.6 | 100 | 100 | India, Singapore, Australia | 221 | March-04, April-05 | -636 | A6 | 0.284 | B.6 | 1 | A | 0.234 | A1 |
| 353 | 452209 | EPI_ISL_452209 | Maharashtra | 29805 | 99.9664 | 99.9372 | 99.98 | B.6 | 100 | 100 | India, Singapore, Australia | 221 | March-04, April-05 | -636 | A6 | 0.29 | B.6 | 1 | A | 0.233 | A1 |
| 355 | 452211 | EPI_ISL_452211 | Maharashtra | 29805 | 99.9631 | 99.9512 | 99.97 | B.1.1 | 95 | 93 | UK, Australia, USA | 6287 | February-15, May-20 | 49 | A2a | 0.998 | B.1.1 | 0.996 | A | 0.232 | A1.14 |
| 357 | 452213 | EPI_ISL_452213 | Maharashtra | 29805 | 99.9664 | 99.9442 | 99.97 | B.4 | 96 | 100 | Australia, UK, Turkey | 258 | January-18, April-14 | 85 | A3 | 0.89 | B.4 | 0.878 | A | 0.234 | A1.15 |
| 358 | 452214 | EPI_ISL_452214 | Maharashtra | 29805 | 99.9698 | 99.9791 | 99.98 | B.1 | 100 | 100 | UK, USA, Australia | 7440 | January-24, May-21 | 48 | A2a | 1 | B.1 | 0.977 | A | 0.234 | A1.16 |
| 362 | 452790 | EPI_ISL_452790 | Madhya Pradesh | 29808 | 99.7769 | 99.6633 | 99.96 | B.1 | 95 | 96 | UK, USA, Australia | 7440 | January-24, May-21 | 48 | A2a | 0.918 | B.1 | 0.754 | A | 0.235 | A1 |
| 365 | 452793 | EPI_ISL_452793 | Madhya Pradesh | 29833 | 99.8683 | 99.7545 | 99.93 | B.6 | 100 | 100 | India, Singapore, Australia | 221 | March-04, April-05 | -636 | A3 | 0.333 | B.6 | 0.807 | A | 0.232 | A1.27 |
| 369 | 454525 | EPI_ISL_454525 | Maharashtra | 29903 | 99.916 | 99.8181 | 99.73 | B.4 | 89 | 100 | Australia, UK, Turkey | 258 | January-18, April-14 | 51 | A3 | 0.839 | B.4 | 0.639 | A | 0.233 | A1.15 |
| 370 | 454526 | EPI_ISL_454526 | Maharashtra | 29903 | 99.943 | 99.9093 | 99.95 | B.4 | 88 | 100 | Australia, UK, Turkey | 258 | January-18, April-14 | 51 | A3 | 0.884 | B.4 | 0.878 | A | 0.234 | A1.15 |
| 373 | 454529 | EPI_ISL_454529 | Maharashtra | 29903 | 99.9731 | 99.9791 | 99.91 | B.1 | 100 | 100 | UK, USA, Australia | 7440 | January-24, May-21 | 14 | A2a | 0.999 | B.1 | 0.968 | A | 0.234 | A1.16 |
| 374 | 454531 | EPI_ISL_454531 | Maharashtra | 29903 | 99.9698 | 99.9651 | 99.92 | A | 100 | 100 | China, South_Korea, USA | 223 | January-05, April-23 | 42 | B4 | 0.903 | A.p7 | 0.569 | A | 0.232 | A1.28 |
| 375 | 454532 | EPI_ISL_454532 | Maharashtra | 29903 | 99.9597 | 99.9581 | 99.89 | B.1 | 97 | 95 | UK, USA, Australia | 7440 | January-24, May-21 | 14 | A2a | 0.998 | B.1 | 0.963 | A | 0.234 | A1 |
| 376 | 454533 | EPI_ISL_454533 | Maharashtra | 29903 | 99.9362 | 99.9511 | 99.9 | B.1.5 | 81 | 100 | UK, Spain, Australia | 710 | February-26, May-10 | 25 | A2a | 1 | B.1.5 | 0.522 | A | 0.233 | A1 |
| 377 | 454534 | EPI_ISL_454534 | Maharashtra | 29903 | 99.9293 | 99.9581 | 99.98 | A | 100 | 100 | China, South_Korea, USA | 223 | January-05, April-23 | 42 | B4 | 0.45 | A.p7 | 0.395 | A | 0.233 | A1.29 |
| 382 | 454543 | EPI_ISL_454543 | Maharashtra | 29903 | 99.9496 | 99.8883 | 99.87 | B.6 | 100 | 100 | India, Singapore, Australia | 221 | March-04, April-05 | -670 | A6 | 0.331 | B.6 | 0.967 | A | 0.234 | A1 |
| 384 | 454546 | EPI_ISL_454546 | Maharashtra | 29903 | 99.963 | 99.8951 | 99.9 | B.6 | 99 | 97 | India, Singapore, Australia | 221 | March-04, April-05 | -670 | A6 | 0.326 | B.6 | 0.955 | A | 0.236 | A1 |
| 385 | 454547 | EPI_ISL_454547 | Maharashtra | 29903 | 99.9664 | 99.9233 | 99.97 | B.6 | 100 | 100 | India, Singapore, Australia | 221 | March-04, April-05 | -670 | A6 | 0.298 | B.6 | 0.998 | A | 0.233 | A1 |
| 389 | 454557 | EPI_ISL_454557 | Maharashtra | 29903 | 99.9396 | 99.9233 | 99.95 | B.1.1 | 96 | 94 | UK, Australia, USA | 6287 | February-15, May-20 | 15 | A2a | 0.998 | B.1.1 | 0.981 | A | 0.232 | A1.14 |
| 393 | 454565 | EPI_ISL_454565 | Maharashtra | 29903 | 99.943 | 99.9233 | 99.95 | B.1.1 | 96 | 96 | UK, Australia, USA | 6287 | February-15, May-20 | 15 | A2a | 1 | B.1.1 | 0.975 | A | 0.233 | A1 |
| 398 | 454570 | EPI_ISL_454570 | Maharashtra | 29903 | 99.9698 | 99.9651 | 99.97 | B.1 | 100 | 100 | UK, USA, Australia | 7440 | January-24, May-21 | 14 | A2a | 0.999 | B.1 | 0.962 | A | 0.235 | A1.16 |
| 401 | 454832 | EPI_ISL_454832 | Rajasthan | 29877 | 99.9833 | 99.9512 | 99.98 | B.6 | 100 | 100 | India, Singapore, Australia | 221 | March-04, April-05 | -636 | A3 | 0.302 | B.6 | 1 | A | 0.233 | A1 |
| 402 | 454833 | EPI_ISL_454833 | Rajasthan | 29865 | 99.9698 | 99.9442 | 99.97 | B.6 | 100 | 100 | India, Singapore, Australia | 221 | March-04, April-05 | -636 | A3 | 0.327 | B.6 | 0.972 | A | 0.233 | A1 |
| 404 | 454862 | EPI_ISL_454862 | Haryana | 29903 | 99.9698 | 99.9302 | 99.97 | B.6 | 100 | 100 | India, Singapore, Australia | 221 | March-04, April-05 | -670 | A6 | 0.244 | B.6 | 1 | A | 0.233 | A1 |
| 406 | 454867 | EPI_ISL_454867 | Haryana | 29903 | 99.9765 | 99.9442 | 99.98 | B.6 | 100 | 100 | India, Singapore, Australia | 221 | March-04, April-05 | -670 | A6 | 0.245 | B.6 | 1 | A | 0.234 | A1 |
| 407 | 455015 | EPI_ISL_455015 | Gujarat | 29903 | 99.9731 | 99.9087 | 99.8 | A | 100 | 100 | China, South_Korea, USA | 223 | January-05, April-23 | 42 | B4 | 0.906 | A.p7 | 0.545 | A | 0.232 | A1.26 |
| 408 | 455016 | EPI_ISL_455016 | Gujarat | 29903 | 99.9732 | 99.9442 | 99.97 | A | 100 | 100 | China, South_Korea, USA | 223 | January-05, April-23 | 42 | B4 | 0.994 | A.p7 | 0.827 | A | 0.233 | A1.26 |
| 411 | 455019 | EPI_ISL_455019 | Gujarat | 29903 | 99.9832 | 99.986 | 99.98 | B.1 | 96 | 96 | UK, USA, Australia | 7440 | January-24, May-21 | 14 | A2a | 1 | B.1 | 0.989 | A | 0.233 | A1.13 |
| 413 | 455021 | EPI_ISL_455021 | Gujarat | 29903 | 99.9732 | 99.9791 | 99.97 | B.1.36 | 77 | 100 | Saudi_Arabia, UK, Turkey | 159 | March-10, May-07 | 28 | A2a | 1 | B.1.36 | 0.518 | A | 0.233 | A1.6.1.4 |
| 414 | 455022 | EPI_ISL_455022 | Gujarat | 29903 | 99.9832 | 99.986 | 99.98 | B.1 | 96 | 96 | UK, USA, Australia | 7440 | January-24, May-21 | 14 | A2a | 1 | B.1 | 0.989 | A | 0.233 | A1.13 |
| 416 | 455024 | EPI_ISL_455024 | Gujarat | 29903 | 99.9832 | 99.986 | 99.98 | B.1 | 96 | 96 | UK, USA, Australia | 7440 | January-24, May-21 | 14 | A2a | 1 | B.1 | 0.989 | A | 0.233 | A1.13 |
| 417 | 455025 | EPI_ISL_455025 | Gujarat | 29903 | 99.9799 | 99.9721 | 99.98 | B.1 | 98 | 96 | UK, USA, Australia | 7440 | January-24, May-21 | 14 | A2a | 1 | B.1 | 0.985 | A | 0.233 | A1.13 |
| 419 | 455027 | EPI_ISL_455027 | Gujarat | 29903 | 99.9732 | 99.9791 | 99.97 | B.1.36 | 76 | 100 | Saudi_Arabia, UK, Turkey | 159 | March-10, May-07 | 28 | A2a | 1 | B.1.36 | 0.551 | A | 0.233 | A1.6.1.4 |
| 422 | 455643 | EPI_ISL_455643 | West Bengal | 29903 | 99.9866 | 99.986 | 99.99 | B.1 | 99 | 100 | UK, USA, Australia | 7440 | January-24, May-21 | 12 | A2a | 1 | B.1 | 1 | A | 0.233 | A1 |
| 423 | 455644 | EPI_ISL_455644 | West Bengal | 29903 | 99.9698 | 99.9512 | 99.97 | B.1 | 97 | 97 | UK, USA, Australia | 7440 | January-24, May-21 | 12 | A2a | 0.999 | B.1 | 0.969 | A | 0.233 | A1 |
| 432 | 455653 | EPI_ISL_455653 | West Bengal | 29903 | 99.9732 | 99.9791 | 99.97 | B.1 | 96 | 95 | UK, USA, Australia | 7440 | January-24, May-21 | 12 | A2a | 1 | B.1 | 0.947 | A | 0.233 | A1.30 |
| 434 | 455656 | EPI_ISL_455656 | West Bengal | 29903 | 99.9665 | 99.9233 | 99.97 | B.6 | 100 | 100 | India, Singapore, Australia | 221 | March-04, April-05 | -672 | A3 | 0.282 | B.6 | 0.997 | A | 0.234 | A1 |
| 437 | 455660 | EPI_ISL_455660 | West Bengal | 29903 | 99.9699 | 99.9372 | 99.97 | B.1 | 99 | 100 | UK, USA, Australia | 7440 | January-24, May-21 | 12 | A2a | 1 | B.1 | 0.876 | A | 0.233 | A1.17 |
| 440 | 455667 | EPI_ISL_455667 | West Bengal | 29903 | 99.9732 | 99.986 | 99.97 | B.1 | 94 | 95 | UK, USA, Australia | 7440 | January-24, May-21 | 12 | A2a | 0.992 | B.1 | 0.944 | A | 0.233 | A1.20 |
| 446 | 455673 | EPI_ISL_455673 | West Bengal | 29903 | 99.9732 | 99.9721 | 99.93 | B.1 | 95 | 95 | UK, USA, Australia | 7440 | January-24, May-21 | 12 | A2a | 0.992 | B.1 | 0.945 | A | 0.233 | A1.20 |
| 447 | 455674 | EPI_ISL_455674 | West Bengal | 29903 | 99.9732 | 99.986 | 99.97 | B.1 | 95 | 95 | UK, USA, Australia | 7440 | January-24, May-21 | 12 | A2a | 0.992 | B.1 | 0.945 | A | 0.233 | A1.20 |
| 448 | 455675 | EPI_ISL_455675 | West Bengal | 29903 | 99.9732 | 99.9791 | 99.97 | B.1 | 96 | 95 | UK, USA, Australia | 7440 | January-24, May-21 | 12 | A2a | 1 | B.1 | 0.95 | A | 0.233 | A1.30 |
| 449 | 455676 | EPI_ISL_455676 | West Bengal | 29903 | 99.9732 | 99.986 | 99.97 | B.1 | 95 | 95 | UK, USA, Australia | 7440 | January-24, May-21 | 12 | A2a | 0.992 | B.1 | 0.945 | A | 0.233 | A1.20 |
| 450 | 455678 | EPI_ISL_455678 | West Bengal | 29903 | 99.9699 | 99.9372 | 99.97 | B.1 | 95 | 100 | UK, USA, Australia | 7440 | January-24, May-21 | 12 | A2a | 1 | B.1 | 0.876 | A | 0.233 | A1.17 |
| 451 | 455679 | EPI_ISL_455679 | West Bengal | 29903 | 99.9765 | 99.9581 | 99.98 | B.1 | 99 | 100 | UK, USA, Australia | 7440 | January-24, May-21 | 12 | A2a | 0.989 | B.1 | 0.973 | A | 0.233 | A1 |
| 462 | 455764 | EPI_ISL_455764 | Odisha | 29690 | 99.9697 | 99.9512 | 99.97 | A | 100 | 100 | China, South_Korea, USA | 223 | January-05, April-23 | 76 | B4 | 0.597 | A.p7 | 0.533 | A | 0.233 | A1.31 |
| 464 | 455766 | EPI_ISL_455766 | Odisha | 29689 | 99.9697 | 99.9512 | 99.97 | A | 100 | 100 | China, South_Korea, USA | 223 | January-05, April-23 | 76 | B4 | 0.595 | A.p7 | 0.533 | A | 0.233 | A1.31 |
| 465 | 455767 | EPI_ISL_455767 | Odisha | 29693 | 99.9697 | 99.9512 | 99.97 | A | 100 | 100 | China, South_Korea, USA | 223 | January-05, April-23 | 76 | B4 | 0.595 | A.p7 | 0.521 | A | 0.233 | A1.31 |
| 469 | 455775 | EPI_ISL_455775 | Odisha | 29841 | 99.9765 | 99.9791 | 99.98 | B.1.36 | 69 | 100 | Saudi_Arabia, UK, Turkey | 159 | March-10, May-07 | 62 | A2a | 1 | B.1 | 0.502 | A | 0.233 | A1.6.1 |
| 471 | 455777 | EPI_ISL_455777 | Odisha | 29688 | 99.9764 | 99.9791 | 99.98 | B.1.36 | 77 | 100 | Saudi_Arabia, UK, Turkey | 159 | March-10, May-07 | 62 | A2a | 1 | B.1 | 0.548 | A | 0.233 | A1.6.1 |
| 472 | 455778 | EPI_ISL_455778 | Odisha | 29862 | 99.9765 | 99.9791 | 99.98 | B.1.36 | 82 | 100 | Saudi_Arabia, UK, Turkey | 159 | March-10, May-07 | 62 | A2a | 1 | B.1.36 | 0.536 | A | 0.233 | A1.6.1 |
| 473 | 455779 | EPI_ISL_455779 | Odisha | 29813 | 99.9765 | 99.9791 | 99.98 | B.1.36 | 78 | 100 | Saudi_Arabia, UK, Turkey | 159 | March-10, May-07 | 62 | A2a | 1 | B.1.36 | 0.524 | A | 0.234 | A1.6.1 |
| 474 | 455780 | EPI_ISL_455780 | Odisha | 29824 | 99.9765 | 99.9791 | 99.98 | B.1.36 | 81 | 100 | Saudi_Arabia, UK, Turkey | 159 | March-10, May-07 | 62 | A2a | 1 | B.1 | 0.514 | A | 0.234 | A1.6.1 |
| 475 | 455782 | EPI_ISL_455782 | Odisha | 29685 | 99.9729 | 99.979 | 99.97 | B.1.36 | 79 | 100 | Saudi_Arabia, UK, Turkey | 159 | March-10, May-07 | 62 | A2a | 1 | B.1 | 0.609 | A | 0.233 | A1.6.1 |
| 477 | 455784 | EPI_ISL_455784 | Odisha | 29685 | 99.973 | 99.9651 | 99.9 | B.1.36 | 75 | 100 | Saudi_Arabia, UK, Turkey | 159 | March-10, May-07 | 62 | A2a | 0.999 | B.1 | 0.544 | A | 0.236 | A1.6.1 |
| 478 | 455786 | EPI_ISL_455786 | Odisha | 29685 | 99.9831 | 99.986 | 99.98 | B.1 | 96 | 100 | UK, USA, Australia | 7440 | January-24, May-21 | 48 | A2a | 1 | B.1 | 0.989 | A | 0.233 | A1 |
| 479 | 455787 | EPI_ISL_455787 | Odisha | 29690 | 99.9865 | 99.986 | 99.99 | B.1 | 100 | 100 | UK, USA, Australia | 7440 | January-24, May-21 | 48 | A2a | 1 | B.1 | 1 | A | 0.233 | A1 |
| 480 | 458030 | EPI_ISL_458030 | Tamil Nadu | 29824 | 99.9698 | 99.9581 | 99.97 | B.1.1 | 95 | 95 | UK, Australia, USA | 6287 | February-15, May-20 | 49 | A2a | 1 | B.1.1 | 0.987 | A | 0.232 | A1.4 |
| 481 | 458031 | EPI_ISL_458031 | Tamil Nadu | 29824 | 99.9698 | 99.9512 | 99.97 | B.1.1 | 95 | 96 | UK, Australia, USA | 6287 | February-15, May-20 | 49 | A2a | 1 | B.1.1 | 0.999 | A | 0.232 | A1.4.1 |
| 482 | 458032 | EPI_ISL_458032 | Tamil Nadu | 29824 | 99.9732 | 99.9651 | 99.97 | B.1.1 | 96 | 94 | UK, Australia, USA | 6287 | February-15, May-20 | 49 | A2a | 1 | B.1.1 | 1 | A | 0.232 | A1.4 |
| 483 | 458033 | EPI_ISL_458033 | Tamil Nadu | 29824 | 99.9698 | 99.9512 | 99.97 | B.1.1 | 95 | 96 | UK, Australia, USA | 6287 | February-15, May-20 | 49 | A2a | 1 | B.1.1 | 0.999 | A | 0.232 | A1.4.1 |
| 484 | 458034 | EPI_ISL_458034 | Tamil Nadu | 29824 | 99.9698 | 99.9512 | 99.97 | B.1.1 | 95 | 96 | UK, Australia, USA | 6287 | February-15, May-20 | 49 | A2a | 1 | B.1.1 | 0.999 | A | 0.232 | A1.4.1 |
| 487 | 458037 | EPI_ISL_458037 | Tamil Nadu | 29824 | 99.9665 | 99.9651 | 99.97 | B.1.1 | 99 | 95 | UK, Australia, USA | 6287 | February-15, May-20 | 49 | A2a | 1 | B.1.1 | 0.999 | A | 0.232 | A1.4.2 |
| 488 | 458038 | EPI_ISL_458038 | Tamil Nadu | 29824 | 99.9631 | 99.9512 | 99.96 | B.1.1 | 98 | 94 | UK, Australia, USA | 6287 | February-15, May-20 | 49 | A2a | 0.996 | B.1.1 | 0.994 | A | 0.232 | A1.4.3 |
| 489 | 458039 | EPI_ISL_458039 | Tamil Nadu | 29824 | 99.9598 | 99.9512 | 99.96 | B.1.1 | 98 | 94 | UK, Australia, USA | 6287 | February-15, May-20 | 49 | A2a | 0.995 | B.1.1 | 0.91 | A | 0.232 | A1.4.3 |
| 490 | 458040 | EPI_ISL_458040 | Tamil Nadu | 29824 | 99.9631 | 99.9442 | 99.96 | B.1.1 | 97 | 94 | UK, Australia, USA | 6287 | February-15, May-20 | 49 | A2a | 1 | B.1.1 | 0.989 | A | 0.232 | A1.4.1 |
| 491 | 458041 | EPI_ISL_458041 | Tamil Nadu | 29824 | 99.9598 | 99.9372 | 99.96 | B.1.1 | 96 | 94 | UK, Australia, USA | 6287 | February-15, May-20 | 49 | A2a | 0.996 | B.1.1 | 0.989 | A | 0.231 | A1.4.3 |
| 494 | 458044 | EPI_ISL_458044 | Tamil Nadu | 29824 | 99.9665 | 99.9651 | 99.97 | B.1.1 | 99 | 95 | UK, Australia, USA | 6287 | February-15, May-20 | 49 | A2a | 1 | B.1.1 | 0.999 | A | 0.232 | A1.4.2 |
| 496 | 458046 | EPI_ISL_458046 | Telangana | 29824 | 99.9497 | 99.9233 | 99.95 | B.1.1.8 | 87 | 90 | UK, Australia, Gambia | 65 | March-17, April-28 | 71 | A2a | 0.987 | B.1.1.8 | 0.866 | A | 0.231 | A1.21 |
| 498 | 458048 | EPI_ISL_458048 | Telangana | 29824 | 99.9497 | 99.9233 | 99.95 | B.1.1.8 | 87 | 90 | UK, Australia, Gambia | 65 | March-17, April-28 | 71 | A2a | 0.987 | B.1.1.8 | 0.866 | A | 0.231 | A1.21 |
| 503 | 458053 | EPI_ISL_458053 | Telangana | 29824 | 99.9732 | 99.9791 | 99.97 | B.1 | 100 | 100 | UK, USA, Australia | 7440 | January-24, May-21 | 48 | A2a | 0.999 | B.1 | 0.981 | A | 0.232 | A1.16 |
| 509 | 458059 | EPI_ISL_458059 | Telangana | 29824 | 99.9497 | 99.9302 | 99.95 | B.1.1 | 95 | 100 | UK, Australia, USA | 6287 | February-15, May-20 | 49 | A2a | 0.997 | B.1.1 | 0.989 | A | 0.23 | A1.14.2 |
| 510 | 458060 | EPI_ISL_458060 | Telangana | 29824 | 99.9531 | 99.9233 | 99.95 | B.1.1 | 98 | 94 | UK, Australia, USA | 6287 | February-15, May-20 | 49 | A2a | 0.998 | B.1.1 | 0.991 | A | 0.231 | A1.14.2 |
| 511 | 458061 | EPI_ISL_458061 | Telangana | 29824 | 99.9531 | 99.9372 | 99.95 | B.1.1 | 99 | 94 | UK, Australia, USA | 6287 | February-15, May-20 | 49 | A2a | 0.98 | B.1.1 | 0.993 | A | 0.231 | A1.14.2 |
| 512 | 458062 | EPI_ISL_458062 | Telangana | 29824 | 99.9665 | 99.9163 | 99.97 | B.6 | 100 | 100 | India, Singapore, Australia | 221 | March-04, April-05 | -636 | A7 | 0.253 | B.6 | 0.979 | A | 0.233 | A1 |
| 513 | 458063 | EPI_ISL_458063 | Telangana | 29824 | 99.9698 | 99.9233 | 99.97 | B.6 | 100 | 100 | India, Singapore, Australia | 221 | March-04, April-05 | -636 | A7 | 0.28 | B.6 | 0.998 | A | 0.233 | A1.18 |
| 514 | 458064 | EPI_ISL_458064 | Telangana | 29824 | 99.9732 | 99.9581 | 99.97 | B.1 | 96 | 95 | UK, USA, Australia | 7440 | January-24, May-21 | 48 | A2a | 0.999 | B.1 | 0.925 | A | 0.233 | A1.22 |
| 515 | 458065 | EPI_ISL_458065 | Telangana | 29824 | 99.9732 | 99.9581 | 99.97 | B.1 | 96 | 95 | UK, USA, Australia | 7440 | January-24, May-21 | 48 | A2a | 0.999 | B.1 | 0.925 | A | 0.233 | A1.22 |
| 516 | 458066 | EPI_ISL_458066 | Telangana | 29824 | 99.9832 | 99.9512 | 99.98 | B.6 | 100 | 100 | India, Singapore, Australia | 221 | March-04, April-05 | -636 | A3 | 0.275 | B.6 | 1 | A | 0.233 | A1 |
| 519 | 458069 | EPI_ISL_458069 | Telangana | 29824 | 99.9732 | 99.9233 | 99.97 | B.6 | 100 | 100 | India, Singapore, Australia | 221 | March-04, April-05 | -636 | A7 | 0.259 | B.6 | 0.999 | A | 0.233 | A1 |
| 521 | 458071 | EPI_ISL_458071 | Telangana | 29824 | 99.9799 | 99.9442 | 99.98 | B.6 | 100 | 97 | India, Singapore, Australia | 221 | March-04, April-05 | -636 | A2a | 0.404 | B.6 | 0.804 | A | 0.233 | A1 |
| 522 | 458072 | EPI_ISL_458072 | Telangana | 29839 | 99.9665 | 99.9233 | 99.97 | B.6 | 99 | 97 | India, Singapore, Australia | 221 | March-04, April-05 | -636 | A3 | 0.307 | B.6 | 0.986 | A | 0.232 | A1.25 |
| 523 | 458073 | EPI_ISL_458073 | Telangana | 29839 | 99.9665 | 99.9163 | 99.97 | B.6 | 100 | 100 | India, Singapore, Australia | 221 | March-04, April-05 | -636 | A3 | 0.272 | B.6 | 0.99 | A | 0.232 | A1 |
| 524 | 458074 | EPI_ISL_458074 | Telangana | 29839 | 99.9631 | 99.9163 | 99.96 | B.6 | 100 | 100 | India, Singapore, Australia | 221 | March-04, April-05 | -636 | A7 | 0.283 | B.6 | 0.989 | A | 0.232 | A1.18 |
| 526 | 458076 | EPI_ISL_458076 | Telangana | 29839 | 99.9698 | 99.9163 | 99.97 | B.6 | 100 | 100 | India, Singapore, Australia | 221 | March-04, April-05 | -636 | A3 | 0.271 | B.6 | 0.997 | A | 0.233 | A1 |
| 528 | 458080 | EPI_ISL_458080 | Telangana | 29824 | 100 | 100 | 100 | B | 99 | 100 | UK, USA, China | 1714 | December-24, May-18 | 51 | A7 | 0.388 | B | 1 | A | 0.234 | A1 |
| 530 | 458087 | EPI_ISL_458087 | Gujarat | 29903 | 99.9765 | 99.9791 | 99.98 | B.1.36 | 73 | 100 | Turkey, Finland, UK | 32 | March-10, April-01 | 53 | A2a | 1 | B.1.36 | 0.515 | A | 0.233 | A1.6.1 |
| 531 | 458088 | EPI_ISL_458088 | Gujarat | 29903 | 99.9665 | 99.9651 | 99.97 | B.1.36 | 97 | 100 | Saudi_Arabia, UK, Turkey | 159 | March-10, May-07 | 28 | A2a | 1 | B.1.36 | 0.903 | A | 0.232 | A1.6.1.1.1 |
| 532 | 458089 | EPI_ISL_458089 | Gujarat | 29903 | 99.9732 | 99.9721 | 99.97 | B.1.36 | 77 | 100 | Saudi_Arabia, UK, Turkey | 159 | March-10, May-07 | 28 | A2a | 1 | B.1 | 0.499 | A | 0.233 | A1.6.1 |
| 537 | 458094 | EPI_ISL_458094 | Gujarat | 29903 | 99.9665 | 99.9651 | 99.97 | B.1.36 | 97 | 100 | Saudi_Arabia, UK, Turkey | 159 | March-10, May-07 | 28 | A2a | 1 | B.1.36 | 0.903 | A | 0.232 | A1.6.1.1.1 |
| 538 | 458095 | EPI_ISL_458095 | Gujarat | 29903 | 99.9698 | 99.9651 | 99.97 | B.1.36 | 98 | 100 | Saudi_Arabia, UK, Turkey | 159 | March-10, May-07 | 28 | A2a | 1 | B.1.36 | 0.883 | A | 0.232 | A1.6.1.1 |
| 540 | 458097 | EPI_ISL_458097 | Gujarat | 29903 | 99.9631 | 99.9581 | 99.96 | B.1.36 | 99 | 100 | Saudi_Arabia, UK, Turkey | 159 | March-10, May-07 | 28 | A2a | 1 | B.1.36 | 0.895 | A | 0.231 | A1.6.1.1.1 |
| 541 | 458098 | EPI_ISL_458098 | Gujarat | 29903 | 99.9665 | 99.9651 | 99.97 | B.1.36 | 97 | 100 | Saudi_Arabia, UK, Turkey | 159 | March-10, May-07 | 28 | A2a | 1 | B.1.36 | 0.903 | A | 0.232 | A1.6.1.1.1 |
| 547 | 458104 | EPI_ISL_458104 | Gujarat | 29903 | 99.9799 | 99.9721 | 99.98 | B.1 | 97 | 96 | UK, USA, Australia | 7440 | January-24, May-21 | 14 | A2a | 1 | B.1 | 1 | A | 0.232 | A1.11 |
| 548 | 458105 | EPI_ISL_458105 | Gujarat | 29903 | 99.9832 | 99.9721 | 99.98 | B.1 | 96 | 97 | UK, USA, Australia | 7440 | January-24, May-21 | 14 | A2a | 1 | B.1 | 1 | A | 0.233 | A1.11 |
| 549 | 458106 | EPI_ISL_458106 | Gujarat | 29903 | 99.9799 | 99.9721 | 99.98 | B.1 | 95 | 96 | UK, USA, Australia | 7440 | January-24, May-21 | 14 | A2a | 1 | B.1 | 0.947 | A | 0.233 | A1.11 |
| 550 | 458107 | EPI_ISL_458107 | Gujarat | 29903 | 99.9765 | 99.9721 | 99.98 | B.1 | 95 | 94 | UK, USA, Australia | 7440 | January-24, May-21 | 14 | A2a | 0.999 | B.1 | 0.928 | A | 0.233 | A1.11 |
| 551 | 458108 | EPI_ISL_458108 | Gujarat | 29903 | 99.9799 | 99.986 | 99.98 | B.1 | 100 | 100 | UK, USA, Australia | 7440 | January-24, May-21 | 14 | A2a | 1 | B.1 | 0.998 | A | 0.233 | A1.16 |
| 552 | 458109 | EPI_ISL_458109 | Gujarat | 29903 | 99.9832 | 99.9581 | 99.79 | B.1 | 95 | 100 | UK, USA, Australia | 7440 | January-24, May-21 | 14 | A2a | 0.995 | B.1 | 0.869 | A | 0.233 | A1.14 |
| 555 | 458112 | EPI_ISL_458112 | Gujarat | 29903 | 99.9564 | 99.9512 | 99.96 | B.1.36 | 100 | 100 | Saudi_Arabia, UK, Turkey | 159 | March-10, May-07 | 28 | A2a | 1 | B.1.36 | 0.807 | A | 0.231 | A1.6.1.1.1.3 |
| 557 | 458298 | EPI_ISL_458298 | Telangana | 29824 | 99.9765 | 99.9233 | 99.98 | B | 100 | 100 | UK, USA, China | 1714 | December-24, May-18 | 51 | A3 | 0.283 | B.6 | 0.766 | A | 0.234 | A1 |
| 559 | 459913 | EPI_ISL_459913 | Delhi | 29862 | 99.896 | 99.7558 | 99.9 | B.1.1 | 96 | 94 | UK, Australia, USA | 6287 | February-15, May-20 | 49 | A2a | 0.753 | B.1.1 | 0.817 | A | 0.234 | A1 |
| 561 | 459915 | EPI_ISL_459915 | Delhi | 29903 | 99.9498 | 99.8884 | 99.95 | B.6 | 100 | 100 | India, Singapore, Australia | 221 | March-04, April-05 | -665 | A3 | 0.289 | B.6 | 0.999 | A | 0.233 | A1 |

| | | | | | | | | | | | | | | | | | | | | | |
|---|---|---|---|---|---|---|---|---|---|---|---|---|---|---|---|---|---|---|---|---|---|
| 562 | 459916 | EPI_ISL_459916 | Delhi | 29870 | 99.9431 | 99.8744 | 99.94 | B.1.1 | 95 | 95 | UK, Australia, USA | 6287 | February-15, May-20 | 49 | A2 | 0.271 | B.1.1 | 0.551 | A | 0.231 | A1 |
| 563 | 459917 | EPI_ISL_459917 | Delhi | 29903 | 99.9363 | 99.8603 | 99.94 | B.6 | 100 | 100 | India, Singapore, Australia | 221 | March-04, April-05 | -665 | A7 | 0.271 | B.6 | 0.934 | A | 0.232 | A1 |
| 564 | 459918 | EPI_ISL_459918 | Delhi | 29903 | 99.9331 | 99.8535 | 99.93 | B.6 | 100 | 100 | India, Singapore, Australia | 221 | March-04, April-05 | -665 | A3 | 0.263 | B.6 | 0.944 | A | 0.234 | A1 |
| 565 | 459919 | EPI_ISL_459919 | Delhi | 29903 | 99.9599 | 99.8953 | 99.96 | B.6 | 100 | 100 | India, Singapore, Australia | 221 | March-04, April-05 | -665 | A3 | 0.294 | B.6 | 0.998 | A | 0.233 | A1 |
| 566 | 459920 | EPI_ISL_459920 | Delhi | 29903 | 99.9465 | 99.8674 | 99.95 | B.6 | 100 | 100 | India, Singapore, Australia | 221 | March-04, April-05 | -665 | A3 | 0.29 | B.6 | 0.996 | A | 0.232 | A1 |
| 567 | 459921 | EPI_ISL_459921 | Delhi | 29903 | 99.9465 | 99.8884 | 99.95 | B.6 | 100 | 100 | India, Singapore, Australia | 221 | March-04, April-05 | -665 | A3 | 0.288 | B.6 | 0.724 | A | 0.232 | A1 |
| 568 | 459922 | EPI_ISL_459922 | Delhi | 29903 | 99.9364 | 99.8605 | 99.94 | B.6 | 100 | 100 | India, Singapore, Australia | 221 | March-04, April-05 | -665 | A3 | 0.28 | B.6 | 0.982 | A | 0.233 | A1 |
| 569 | 459923 | EPI_ISL_459923 | Delhi | 29903 | 99.9264 | 99.8395 | 99.93 | B.6 | 100 | 100 | India, Singapore, Australia | 221 | March-04, April-05 | -665 | A3 | 0.272 | B.6 | 0.969 | A | 0.232 | A1 |
| 570 | 459924 | EPI_ISL_459924 | Delhi | 29903 | 99.9331 | 99.8605 | 99.93 | B.1.1 | 96 | 94 | UK, Australia, USA | 6287 | February-15, May-20 | 20 | A2a | 0.691 | B.1.1 | 0.738 | A | 0.231 | A1 |
| 571 | 459926 | EPI_ISL_459926 | Delhi | 29903 | 99.8829 | 99.7488 | 99.88 | B.6 | 99 | 100 | India, Singapore, Australia | 221 | March-04, April-05 | -665 | A3 | 0.3 | B.6 | 0.921 | A | 0.23 | A1 |
| 572 | 459927 | EPI_ISL_459927 | Delhi | 29903 | 99.913 | 99.8186 | 99.91 | B.6 | 100 | 100 | India, Singapore, Australia | 221 | March-04, April-05 | -665 | A7 | 0.314 | B.6 | 0.952 | A | 0.232 | A1 |
| 573 | 459932 | EPI_ISL_459932 | Delhi | 29903 | 99.8661 | 99.686 | 99.87 | B.6 | 100 | 100 | India, Singapore, Australia | 221 | March-04, April-05 | -665 | A3 | 0.269 | B.6 | 0.92 | A | 0.233 | A1 |
| 574 | 459933 | EPI_ISL_459933 | Delhi | 29903 | 99.9164 | 99.8326 | 99.92 | B.6 | 100 | 100 | India, Singapore, Australia | 221 | March-04, April-05 | -665 | A3 | 0.265 | B.6 | 0.91 | A | 0.233 | A1 |
| 575 | 459934 | EPI_ISL_459934 | Delhi | 29903 | 99.9164 | 99.8186 | 99.92 | B.6 | 100 | 100 | India, Singapore, Australia | 221 | March-04, April-05 | -665 | A3 | 0.28 | B.6 | 0.912 | A | 0.232 | A1 |
| 576 | 459935 | EPI_ISL_459935 | Delhi | 29903 | 99.903 | 99.8326 | 99.9 | B.6 | 100 | 100 | India, Singapore, Australia | 221 | March-04, April-05 | -665 | A3 | 0.311 | B.6 | 0.949 | A | 0.231 | A1 |
| 577 | 459937 | EPI_ISL_459937 | Delhi | 29903 | 99.9261 | 99.8182 | 99.93 | B.6 | 100 | 100 | India, Singapore, Australia | 221 | March-04, April-05 | -665 | A3 | 0.253 | B.6 | 0.969 | A | 0.236 | A1 |
| 578 | 459938 | EPI_ISL_459938 | Delhi | 29903 | 99.9123 | 99.8246 | 99.95 | B.6 | 99 | 100 | India, Singapore, Australia | 221 | March-04, April-05 | -665 | A3 | 0.271 | B.6 | 0.886 | A | 0.234 | A1 |
| 579 | 459939 | EPI_ISL_459939 | Delhi | 29903 | 99.9192 | 99.825 | 99.95 | B.6 | 100 | 100 | India, Singapore, Australia | 221 | March-04, April-05 | -665 | A3 | 0.252 | B.6 | 0.876 | A | 0.232 | A1 |
| 580 | 459940 | EPI_ISL_459940 | Delhi | 29903 | 99.8963 | 99.7907 | 99.9 | B.6 | 100 | 100 | India, Singapore, Australia | 221 | March-04, April-05 | -665 | A3 | 0.24 | B.6 | 0.89 | A | 0.234 | A1 |
| 581 | 459941 | EPI_ISL_459941 | Delhi | 29903 | 99.9097 | 99.8186 | 99.91 | B.6 | 100 | 100 | India, Singapore, Australia | 221 | March-04, April-05 | -665 | A3 | 0.284 | B.6 | 0.978 | A | 0.233 | A1 |
| 582 | 459942 | EPI_ISL_459942 | Delhi | 29903 | 99.9331 | 99.8814 | 99.93 | B.6 | 100 | 100 | India, Singapore, Australia | 221 | March-04, April-05 | -665 | A3 | 0.29 | B.6 | 0.991 | A | 0.233 | A1 |
| 583 | 459943 | EPI_ISL_459943 | Delhi | 29903 | 99.9164 | 99.8605 | 99.92 | B.6 | 100 | 100 | India, Singapore, Australia | 221 | March-04, April-05 | -665 | A3 | 0.274 | B.6 | 0.948 | A | 0.232 | A1 |
| 587 | 461481 | EPI_ISL_461481 | Gujarat | 29903 | 99.9799 | 99.986 | 99.98 | B.1 | 100 | 100 | UK, USA, Australia | 7440 | January-24, May-21 | 19 | A2a | 1 | B.1 | 0.998 | A | 0.233 | A1.16 |
| 589 | 461483 | EPI_ISL_461483 | Gujarat | 29903 | 99.9799 | 99.986 | 99.98 | B.1 | 100 | 100 | UK, USA, Australia | 7440 | January-24, May-21 | 19 | A2a | 1 | B.1 | 0.999 | A | 0.233 | A1.16 |
| 596 | 461490 | EPI_ISL_461490 | Gujarat | 29903 | 99.9799 | 99.986 | 99.98 | B.1 | 100 | 100 | UK, USA, Australia | 7440 | January-24, May-21 | 19 | A2a | 1 | B.1 | 0.937 | A | 0.232 | A1.19 |
| 599 | 461493 | EPI_ISL_461493 | Gujarat | 29903 | 99.9732 | 99.9721 | 99.97 | B.1 | 100 | 100 | UK, USA, Australia | 7440 | January-24, May-21 | 19 | A2a | 1 | B.1 | 0.803 | A | 0.233 | A1.16 |
| 602 | 461496 | EPI_ISL_461496 | Gujarat | 29903 | 99.9765 | 99.9721 | 99.98 | B.1 | 100 | 100 | UK, USA, Australia | 7440 | January-24, May-21 | 19 | A2a | 1 | B.1 | 0.803 | A | 0.233 | A1.16 |
| 606 | 461500 | EPI_ISL_461500 | Gujarat | 29903 | 99.9631 | 99.9581 | 99.96 | B.1.36 | 96 | 100 | Saudi_Arabia, UK, Turkey | 159 | March-10, May-07 | 33 | A2a | 1 | B.1.36 | 0.907 | A | 0.23 | A1.6.1.1.1 |
| 609 | 461503 | EPI_ISL_461503 | Gujarat | 29903 | 99.9765 | 99.9721 | 99.98 | B.1 | 100 | 100 | UK, USA, Australia | 7440 | January-24, May-21 | 19 | A2a | 1 | B.1 | 0.617 | A | 0.232 | A1.19 |
| 611 | 461505 | EPI_ISL_461505 | Gujarat | 29903 | 99.9832 | 99.9721 | 99.98 | B.1 | 97 | 100 | UK, USA, Australia | 7440 | January-24, May-21 | 19 | A2a | 1 | B.1 | 0.998 | A | 0.234 | A1 |
